# Single-cell proteomics reveals decreased abundance of proteostasis and meiosis proteins in advanced maternal age oocytes

**DOI:** 10.1101/2024.05.23.595547

**Authors:** S Galatidou, A Petelski, A Pujol, K Lattes, L B Latorraca, T Fair, M Popovic, R Vassena, N Slavov, M Barragan

## Abstract

Advanced maternal age is associated with a decline in oocyte quality, which often leads to reproductive failure in humans. However, the mechanisms behind this age-related decline remain unclear. To gain insights into this phenomenon, we applied plexDIA, a multiplexed, single-cell mass spectrometry method, to analyze the proteome of oocytes from both young women and women of advanced maternal age. Our findings primarily revealed distinct proteomic profiles between immature fully grown germinal vesicle and mature metaphase II oocytes. Importantly, we further show that a woman’s age is associated with changes in her oocyte proteome. Specifically, when compared to oocytes obtained from young women, advanced maternal age oocytes exhibited lower levels of the proteasome and TRiC complex, as well as other key regulators of proteostasis and meiosis. This suggests that aging adversely affects the proteostasis and meiosis networks in human oocytes. The proteins identified in this study hold potential as targets for improving oocyte quality and may guide future studies into the molecular processes underlying oocyte aging.

## INTRODUCTION

Over the past decades, increasing numbers of women have delayed childbearing due to financial, educational, and social factors (Schmidt *et al*., 2011). However, female fertility decreases with age, with a more profound decline after the age of 35 years, commonly referred to as advanced maternal age (AMA) (Menken *et al*., 1986; Baird *et al*., 2005). Consequently, a growing number of women face age-related subfertility and are turning to medically assisted reproduction to enhance their chances of conceiving. However, the success of assisted reproductive technologies remains limited in patients of AMA, as i*n vitro* fertilizatio*n* (IVF) treatment is unable to fully offset the natural decline in fertility associated with female aging (Leridon, 2004). In women of AMA, reproductive failure is primarily attributed to the pivotal role of oocytes. As women age, both the number and quality of oocytes decline, ultimately reducing the chances of successful conception.

Poor oocyte quality is closely linked to meiotic aneuploidies, which typically arise from chromosome missegregation errors during the first meiotic division (Hassold and Hold, 2001; Herbert *et al*., 2015) and from cytoplasmic alterations (Igarashi *et al*., 2015; Reader *et al*., 2017). However, the specific mechanisms underlying the diminished quality of oocytes in women of AMA remain poorly understood. Unraveling the complexities of this process will be vital for developing strategies to address age-related subfertility.

Within the ovaries, oocytes are arrested at the prophase stage of meiosis I until a preovulatory surge of luteinizing hormone (LH) induces germinal vesicle breakdown (GVBD) and initiates the resumption of meiosis. RNA transcription becomes silent once oocyte growth is completed, and this transcriptional inactivity is maintained during GVBD, meiotic maturation, fertilization and the initial cleavage divisions until embryonic genome activation (Gosden and Lee, 2010; Vassena *et al*., 2011; Cornet-Bartolomé *et al*., 2021). Despite this transcriptional silence, the translation of maternal mRNAs continues during oocyte maturation and early embryo development (Gosden and Lee, 2010; Susor *et al*., 2015). Consequently, these processes are mainly regulated by post-transcriptional and (post-) translational mechanisms. Any disruption to these regulatory mechanisms could result in an imbalanced proteome and potentially impact the quality of oocytes and embryos.

Loss of proteostasis, which refers to the disruption of proteome homeostasis, has been associated with the aging process in various cell types (Klaips *et al*., 2017; Hipp *et al*., 2019). In the context of oocyte biology, loss of proteostasis may contribute to the decline in oocyte quality with age (Café *et al*., 2021; Sala *et al*., 2022). This hypothesis is supported by studies on mammalian oocytes, which have shown differential expression of genes related to protein metabolism (Duncan *et al*., 2017) and dysregulation of proteasome activity (Mihalas *et al*., 2018). However, existing proteomic research remains limited and has focused on oocytes from young women, leaving a significant gap concerning the age-related effects on the oocyte proteome (Virant-Klun *et al*., 2016; Guo *et al*., 2022).

Here, we used plexDIA, a multiplexed mass spectrometry method for single-cell proteomic quantification (Derks *et al*., 2022), to evaluate the proteomic profile of single oocytes from both young and AMA women. Our findings shed light on the relationship between protein abundance and aging in human oocytes. Specifically, we reveal a significant reduction in the levels of proteins important for the proteostasis and meiosis networks. Among the proteins with decreased abundance, the proteasome complex stands out and may account for the observed poor quality in oocytes from women of AMA. By elucidating these changes in protein abundance associated with oocyte aging, our research contributes to a better understanding of the mechanisms that underlie age-related subfertility. We also highlight the importance of proteostasis in maintaining oocyte quality, which may pave the way towards the development of targeted interventions and treatments to improve oocyte quality in reproductive aged women.

## MATERIALS AND METHODS

### Ethical approval

Approval to conduct this study was obtained from the Ethics Committee for Research with Medicinal Products, Eugin, Barcelona. All women participating in this study provided their written informed consent prior to inclusion.

### Study population

#### Proteomic analysis

Fifty-two women, who underwent controlled ovarian stimulation from May 2021 to May 2022 at two participating centers, were included in the study. The Young group comprised of women enrolled in the centers’ oocyte donation programme (n= 27). They had a mean age of 24.5 years (SD= 3.9, range 18-30) and a mean ovarian reserve, measured by an antral follicle count (AFC), of 23 (SD= 9.1, range 8-45). The AMA group included patients (n= 25) with a mean age of 39 years (SD= 1.7, range 37-43) and a mean AFC of 10 (SD= 4.0, range 2-16). Some of the women contributed with more than one oocyte (Table S1).

#### Functional analysis

For experiments focused on rescue *in vitro* maturation (rIVM) and proteasome complex inhibition, we included nine young oocyte donors (≤ 35 years) who underwent controlled ovarian stimulation from April 2023 to May 2023. They had a mean age of 28.8 years (SD= 3.9, range 22-34) and the mean ovarian reserve, measured by AFC was 21.8 (SD= 4.8, range 13-27). Some of the women contributed with more than one oocyte (Table S2).

### Ovarian stimulation and oocyte retrieval

All participants had normal ovarian morphology at transvaginal ultrasound and a progressive increase in follicular size in response to ovarian stimulation. All received daily injections of 150 to 300 IU highly purified urinary hMG (Menopur®; Ferring, Spain) or follitropin alpha (Gonal®; MerckSerono, Spain) (Blazquez *et al*., 2014). From day 6 of stimulation, pituitary suppression was achieved by administering a GnRH antagonist (0.25 mg of cetrorelix acetate, Cetrotide®; Merck Serono; or 0.25 mg ganirelix, Orgalutran®; Merck Sharp & Dohme, Madrid, Spain) (Olivennes *et al*., 1995). Oocyte maturation and ovulation was triggered with 0.3 mg of GnRH agonist (Decapeptyl®, Ipsen Pharma S.A., Spain) and 250 μg hCG (Ovitrelle®, Merck, Germany) for Young and AMA women, respectively. Ovulation was triggered when at least three follicles >18 mm, and at least five follicles >16 mm in diameter developed on both ovaries. The oocyte pick-up was carried out 36 hours after the trigger by ultrasound-guided transvaginal follicular aspiration. The retrieved oocytes were denuded by enzymatic (80 IU/ml hyaluronidase in G-MOPS medium, Vitrolife, Sweden) and mechanical treatment. Nuclear maturity was determined by assessing the presence of a polar body. Oocytes were classified as metaphase II (MII) oocytes when one polar body was present, or germinal vesicle (GV) oocytes when the germinal vesicle was intact. MII oocytes were vitrified on the day of oocyte pick-up and were warmed on the day of proteome isolation (Cryotop®, Kitazato®, BioPharma Co., Ltd; Japan).

### Oocytes

#### Proteomic analysis

A total of 68 oocytes were included in the study. These included 36 GVs (n= 18 in the YOUNG and n= 18 in the AMA group) and 32 MII oocytes (n= 18 in the Young and n= 14 in the AMA group) (Table S1). After warming, MII oocytes were incubated in G-2^TM^ PLUS medium (Vitrolife, Sweden) for 3 hours at 37°C and 6% CO_2_ prior to further processing, to allow for the metaphase plate to re-assemble. One MII oocyte from the YOUNG group degenerated after the warming process and was discarded (Table S1).

#### Functional analysis

A total of 19 oocytes from young women were included in the study. These included 10 GVs (n= 5 in the control group and n= 5 in the MG-132 treated group) and 9 meiotic metaphase I (MI) stage oocytes that have undergone GVBD *in vivo* but have not extruded the first polar body (of these, n= 5 in the control group and n= 4 in the MG-132 treated group) (Table S2).

### Single-cell proteomics

Individual oocytes were incubated in Acidic Tyrode’s solution (Sigma-Aldrich, USA) for 5 seconds under a stereoscope to ensure the complete removal of cumulus cells, subsequently washed thoroughly in nuclease-free water and placed in individual tubes with 1 μL nuclease-free water, snap-frozen in liquid nitrogen and stored at −80°C until further processing.

The proteomic analysis was performed by plexDIA as described previously (Derks *et al*., 2022). Briefly, single cells were lysed using the mPOP lysis method, which involves a freeze heat cycle. Protein digestion into peptides proceeded with the addition of trypsin and triethylammonium bicarbonate (TEAB, at pH = 8) to each lysed cell at a final concentration of 10 ng/μl and 100 μM, respectively. The samples were digested for three hours at 37℃ in a thermal cycler. The peptides were then labeled with non-isobaric mass tags called mTRAQ for two hours at room temperature. The mTRAQ labels were resuspended in isopropanol at the manufacturer concentration and buffered with TEAB for a final concentration of 200mM. The labeling reaction was quenched using 0.5% hydroxylamine for one hour at room temperature. Finally, clusters of three single cells were pooled into plexDIA sets for subsequent mass spectrometry analysis (Petelski *et al*., 2021). Each plexDIA set contained either 3 oocytes or 2 oocytes along with a negative water control that had experienced all of the sample preparation steps.

The single oocyte sets were injected at 1-μl volumes using a Dionex UltiMate 3000 UHPLC with online nLC with a 15 cm x 75 um IonOpticks Aurora Series UHPLC column. Xcalibur was used to control the instrument. Upon being separated by liquid chromatography, peptide samples were subjected to electrospray ionization and sprayed into a Q Exactive instrument. In these experiments, Buffer A was 0.1% formic acid diluted in LC-MS-grade water, while Buffer B was 80% acetonitrile and 0.1% formic acid also diluted in LC-MS-grade water. The gradient (total run-time of 95 minutes) was designed as follows: 4% Buffer B (minutes 0 - 11.5), 4-8% Buffer B (minutes 11.5 - 12), 8 - 32% Buffer B (minutes 12 - 75), 32 - 95% Buffer B (minutes 75 - 77), 95% Buffer B (minutes 77 - 80), 95 - 4% Buffer B (minutes 80 - 80.1), 4% Buffer B (minutes 80.1 - 95). During the gradient, the flow remained at 200 nl/min. The duty cycle for each run consisted of one MS1 followed by 25 DIA MS2 windows of variable m/z length (specifically: 18 windows of 20 Th, 2 windows of 40 Th, 3 windows of 80 Th, and 2 windows of 160 Th). The entire span of analysis ranged from 378 to 1370 m/z. Each MS1 scan was conducted at 70,000 resolving power, 3 x 10^6^ AGC maximum, and 300-ms injection time. Each MS2 scan was conducted at 35,000 resolving power, 3 x 10^6^ AGC maximum, and 110-ms injection time. NCE was set to 27% with a default charge state of 2.

MS raw files were searched using the DIA-NN software (version 1.8.1 beta 16) using the human spectral library used in Derks *et al*. 2022. The following parameters were used: scan window = 5, mass accuracy = 10 p.p.m., and MS1 accuracy = 5 p.p.m. Library generation was set to “IDs, RT and IM Profiling” and Quantification Strategy was set to “Peak Height”. Additionally, the following commands were entered into the DIA-NN command line GUI: (1) {-fixed-mod mTRAQ, 140.0949630177, nK} 2) {-channels mTRAQ,0,nK,0:0; mTRAQ, 4, nK, 4.0070994:4.0070994; mTRAQ, 8, nK, 8.0141988132:8.0141988132} 3) −peak-translation 4) {-original-mods} 5) {-report-lib-info} 6) {-ms1-isotope-quant}. After the data was searched using DIA-NN, MS1 level quantitation was used to normalize the precursors, which were then collapsed into protein-level data. The final data contains log2 transformed protein abundances that are relative to the global mean.

After quality control, 13 samples were excluded from further analysis, as they either had low proteome coverage (< 65 %) or did not exhibit good agreement among peptides mapping to the same proteins. The final and excluded oocytes are summarized in Table S1.

### Statistical analysis of proteomic data

The analysis was restricted to proteins identified in at least 80% of the samples within each group. Differentially abundant proteins were detected between the experimental groups using the non-parametric Mann Whitney U Test with Benjamini Hochberg correction applied with fold change set at |FC| > 1.5, and an adjusted p-value (p.adj) of ≤ 0.05. Correlation analysis was performed using the Spearman test, with a strong significant correlation determined by the thresholds |R| ≥ 0.5 and p.adj ≤ 0.2, while correlations with moderate significance were determined by the thresholds 0.3≤ |R| < 0.5 and p-value ≤ 0.05. For protein complexes composed of several subunits, correlation with age was assessed using the mean correlation coefficient (R_mean_) values of their subunits. Statistical validation of the mean correlation coefficient was performed by randomizing the data 10^4^ times and comparing the R_mean_ of our data with the permutation distribution of R_mean_.

### Protein set enrichment analysis

Biological processes and pathways were identified after comparing the levels of total proteins belonging to each gene ontology term between the two groups, by applying with the Mann Whitney U Test. The ontology terms were acquired by the Biological_Process_2021, Cellular_Compartment_2021 and, KEGG_2021_Human libraries (Xie *et al*., 2021). Those with less than five annotated proteins were excluded. Biological processes and pathways were considered to differ significantly between the two groups when they had at least 50% of the proteins of the term identified and a p.adj ≤0.05. Data visualization was performed using R packages, ggplot2 for the volcano plots, scatter plots, histogram and Dot plots and ComplexHeatmap for heatmaps.

### Rescue *in vitro* maturation (rIVM) and proteasome complex inhibition

Human immature denuded oocytes, GV and MI oocytes, were cultured in drops of 20 μL G-2™PLUS (Vitrolife, Sweden) covered with Ovoil (Vitrolife, Sweden) at 37°C and 6% CO_2_ to reach nuclear maturation (*in vitro* matured MII, IVM-MII). Rescue IVM was performed in the absence (0.1% DMSO; control) or presence of 10 μM MG-132, a cell-permeable, potent, and reversible proteasome inhibitor (Sigma Aldrich; MerckSerono, Spain). After 6 hours of culture, the oocytes were transferred to fresh G-2™PLUS medium to continue the rIVM process for up to 48 hours. The oocytes were closely monitored for their maturation status, based on the extrusion of the first polar body.

### Immunofluorescence staining

Following the designated culture period for rIVM, oocytes were rinsed in prewarmed phosphate buffered saline (PBS), fixed in 4% paraformaldehyde (PFA)/PBS for 15 minutes at room temperature (RT), washed in PBST (PBS 0.1% Tween-20) for 10 minutes and stored in PBST at 4 °C until processing. For immunostaining, oocytes were permeabilized with 0.5% Triton X-100 in PBS for 20 minutes, then blocked in a solution of 5% normal goat serum, 2% bovine serum albumin (BSA) and 0.1% Tween-20 in PBS for 3h at RT, and incubated overnight at 4°C with rabbit monoclonal anti-proteasome 20S alpha and beta antibody (1:300; ab22673; Abcam; Cambridge, Germany) or with mouse monoclonal anti-α-tubulin antibody (1:1000; T6199; Sigma Aldrich; Merck KGaA, Darmstadt, Germany) in blocking solution. The negative controls were incubated overnight in a blocking solution. Once washed in PBST, samples were incubated for 1 hour at RT with secondary antibody diluted 1:500 (Alexa Fluor 568 goat anti-rabbit IgG; Invitrogen, Carlsbad, CA, USA or Alexa Fluor 488 goat anti-mouse IgG (H + L); Thermo Fisher Scientific, Waltham, MA, USA). After washing, the DNA was stained with 2 μg/mL Hoechst 33342 (Thermo Fisher Scientific, Waltham, MA, USA). Samples were rinsed in PBST and immediately imaged using μ-slide 15-well glass bottom dishes (81507; ibidi; Grafelfing, Germany).

### Image acquisition and analysis

Stained oocytes were imaged using the Nikon Eclipse T*i*2 stand attached to an Andor Dragonfly 505 high speed confocal microscope equipped with the multimode optical fiber illumination system Boreallis^TM^ Integrated Laser Engine containing the 405, 445, 488, 514, 561, 594 and 637 nm solid-state lasers. Samples were imaged through a 60× water objective. Laser power and photomultiplier settings for each staining were kept constants for all samples. Imaging data were analyzed using the open-source image processing software ImageJ (Rasband, W.S., ImageJ, U. S. National Institutes of Health, Bethesda, Maryland, USA, https://imagej.nih.gov/ij/, 1997-2018) and Imaris software (Oxford instruments).

## RESULTS

### The proteomic landscape of human oocytes changes during meiotic maturation

We primarily focused on characterizing the proteomic profiles of human immature (GV) and mature (MII) oocytes to identify changes upon the resumption of meiosis. To begin, we compare the profiles of GV versus MII oocytes only from young women as a reference since these oocytes are expected to have good quality.

Specifically, in Young oocytes, we identified a total of 1,368 proteins in both maturation stages from which 26 proteins were less abundant (including YBOX2, PS14, RS16) and 28 more abundant in MII oocytes (such as AURKA, WEE2, BUB1B) compared to GV oocytes (Figure 1 a, c; Table S3). The proteins that showed lower abundance in MII oocytes primarily participate in translation process with many of them being ribosomal subunits (Table 1). In contrast, the proteins that displayed high abundance in MII oocytes were mainly associated with the regulation of the cell cycle and microtubule organization (Table 1).

**Figure 1:**
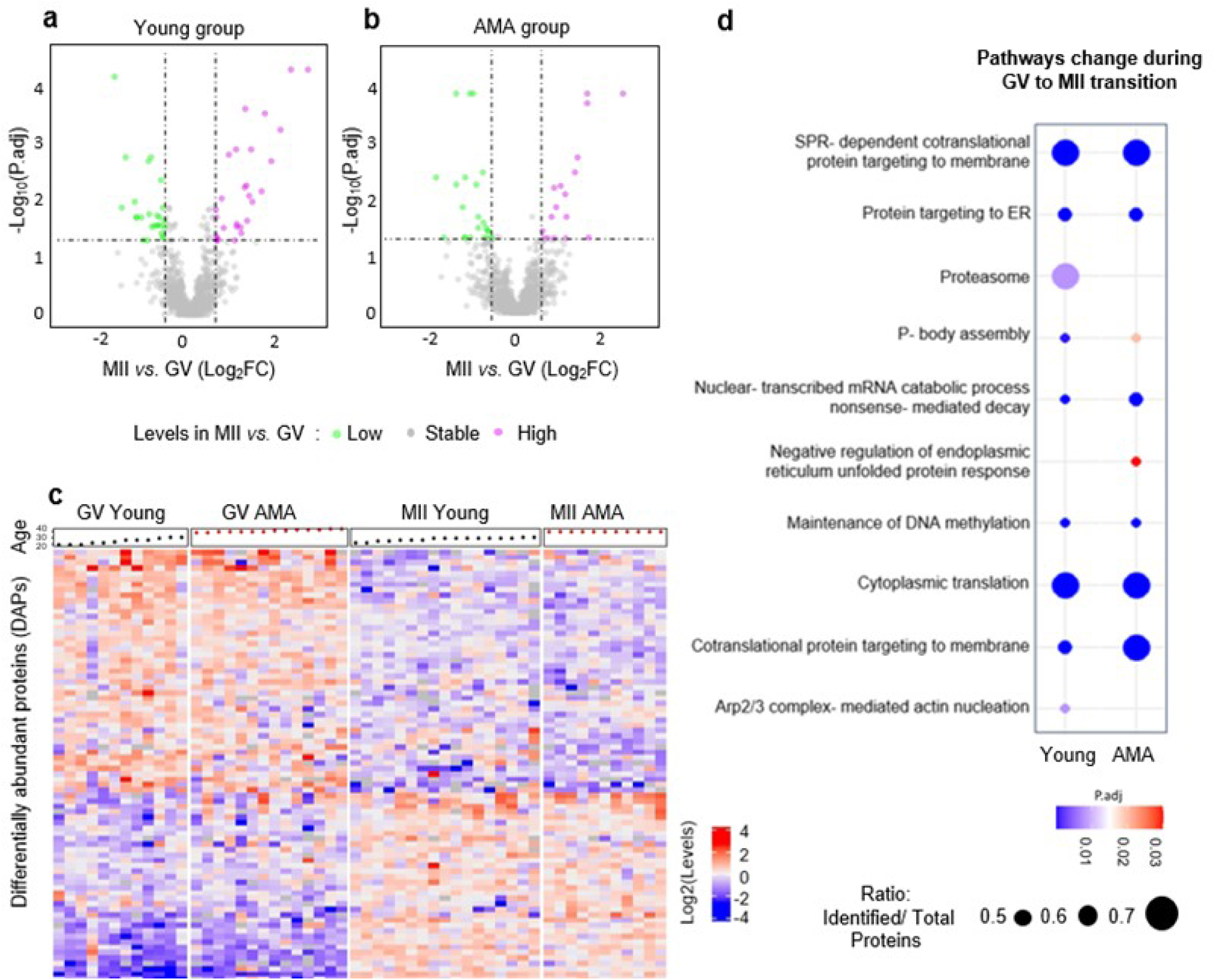
Proteomic changes during the final steps of oocyte maturation. Differentially abundant proteins (DAPs) identified during meiotic maturation of human oocytes. Volcano plots showed DAPs identified by plexDIA between MII and GV oocytes from the a) Young and b) AMA groups (P.adj ≤ 0.05, Wilcoxon test, |fold change| > 1.5). c) Heatmap showing the log2 levels of DAPs for all oocyte groups. d) Ontology terms representing the pathways that statistically differ between GV and MII oocytes in Young and AMA groups (Wilcoxon test, p.adj ≤ 0.05). Dot size reflects the ratio of identified to total pathway proteins that change during GV to MII transition and dot color the p.adj.

**Table 1.** Oocyte maturation-related proteome.

We further evaluated the biological processes, cellular compartments and pathways that undergo overall changes from GV to MII transition by applying protein set enrichment analysis. We found that several pathways, including cytoplasmic translation, maintenance of DNA methylation, Arp2/3 complex mediated actin nucleation and proteasome complex differed significantly between GV and MII oocytes (Figure 1 d; Table S4).

### The proteomic landscape of meiotic maturation is maintained in advanced maternal age oocytes

We performed a similar analysis for oocytes in the AMA group to evaluate whether the overall protein composition and pathways are maintained with AMA. In total 1,451 proteins were identified both in GV and MII AMA oocytes, 23 proteins with lower abundance (including YBOX2, ZAR1, RS18) and 17 proteins with higher abundance in MII when compared to GV (including AURKA, WEE2, BUB1B) (Figure 1 b, c; Table S5). As with the Young group, the differentially abundant proteins were mainly ribosomal subunits (low in MII), cell cycle regulators and microtubule related proteins (high in MII) (Table 1). Furthermore, the majority of pathways undergoing overall changes during GV to MII transition, in AMA oocytes, exhibited similarities to the Young group. Interestingly, the negative regulation of endoplasmic reticulum unfolded protein response, the Arp2/3 complex mediated actin nucleation and the proteasome complex, showed different pattern (Figure 1 d; Table S6).

### AMA disturbs the proteome of immature GV oocytes

Next, we analyzed the relationship between age and the oocyte proteome at each maturation stage. To identify putative protein targets that may explain chromosome segregation errors or early cytoplasmic alterations in oocytes of women of AMA, we focused on GV oocytes. These oocytes have not completed the first meiotic division and retain all pairs of homologous chromosomes. Our analysis revealed a strong correlation (|R|≥ 0.5) with age, showing a negative association for the levels of 12 proteins and a positive association for 8 proteins (Table 2). Additionally, the levels of 134 proteins exhibited a moderate correlation (0.3 ≤|R| <0.5) (Table S7).

**Table 2.** Age-correlated proteins in GV and MII oocytes.

|  | Uniprot ID | Protein Symbol (Name) | R | p.<br>adjusted |
| --- | --- | --- | --- | --- |
| GV<br>OOCYTES | P62195 | PRS8 (26S proteasome regulatory subunit 8) | -0.71 | 0.030 |
|  | P62258 | 1433E (14-3-3 protein epsilon) | -0.70 | 0.031 |
|  | P11142 | HSP7C (Heat shock cognate 71 kDa protein) | -0.65 | 0.117 |
|  | Q9UNS2 | CSN3 (COP9 signalosome complex subunit 3) | -0.64 | 0.117 |
|  | P09936 | UCHL1 (Ubiquitin carboxyl-terminal hydrolase isozyme L1) | -0.63 | 0.117 |
|  | P17980 | PRS6A (26S proteasome regulatory subunit 6A) | -0.62 | 0.117 |
|  | P31948-2 | STIP1 (Stress-induced-phosphoprotein 1) | -0.62 | 0.117 |
|  | Q96FJ2 | DYL2 (Dynein light chain 2, cytoplasmic) | -0.62 | 0.109 |
|  | Q9UIC8 | LCMT1 (Leucine carboxyl methyltransferase 1) | -0.61 | 0.109 |
|  | P47756-2 | CAPZB (F-actin-capping protein subunit beta) | -0.61 | 0.119 |
|  | Q02790 | FKBP4 (Peptidyl-prolyl cis-trans isomerase) | -0.58 | 0.148 |
|  | Q99832 | TCPH (T-complex protein 1 subunit eta) | -0.58 | 0.159 |
|  | P35232 | PHB1 (Prohibitin 1) | 0.57 | 0.180 |
|  | Q9H3N1 | TMX1 (Thioredoxin-related transmembrane protein 1) | 0.58 | 0.196 |
|  | Q16891 | MIC60 (MICOS complex subunit) | 0.58 | 0.148 |
|  | Q9UBS4 | DJB11 (DnaJ homolog subfamily B member 11) | 0.59 | 0.148 |
|  | O75964 | ATP5L (ATP synthase subunit g, mitochondrial) | 0.60 | 0.119 |
|  | O00592 | PODXL (Podocalyxin) | 0.64 | 0.107 |
|  | P18084 | ITB5 (Integrin beta-5) | 0.65 | 0.139 |
|  | Q5VV41 | ARHGG (Rho guanine nucleotide exchange factor 16) | 0.81 | 0.004 |
| MII<br>OOCYTES | P01857 | IGHG1 (Immunoglobulin heavy constant gamma 1) | -0.69 | 0.047 |
|  | Q6UB35 | C1TM (Monofunctional C1-tetrahydrofolate synthase, mitochondrial) | -0.69 | 0.071 |
|  | Q14894 | CRYM (Ketimine reductase mu-crystallin) | -0.68 | 0.047 |
|  | O60547 | GMDS (GDP-mannose 4,6 dehydratase) | -0.61 | 0.145 |
|  | Q9NQI0 | DDX4 (Probable ATP-dependent RNA helicase) | -0.60 | 0.160 |
|  | P80723 | BASP1 (Brain acid soluble protein 1) | 0.67 | 0.070 |
|  | Q9UJS0 | S2513 (Electrogenic aspartate/glutamate antiporter SLC25A13, mitochondrial) | 0.61 | 0.160 |

Among the proteins which abundance declines with age, we identified meiosis key factors, such as 1433E, proteasome subunits, CDK1, regulators of proteostasis (UCHL1, HSP7C, CSN3) and (co)-chaperones (e.g TRiC complex subunits). Conversely, the proteins that showed an increase in abundance with age were primarily associated with mitochondrial functions (e.g., ATP5L, MIC60) (Table 2, Table S7).

We then focused on the proteasome and TRiC complex since these complexes have an important role in meiosis and proteostasis networks. Upon analyzing the proteasome complex comprehensively, 35 subunits were quantified in GV oocytes. Among them, five subunits (PRS8, PRS6A, PRS10, PSA6, PSMF1) showed a negative correlation with age (R from −0.71 to −0.40) (Figure 2 a-e; Table S8). The negative correlation of the five subunits results in a non-random negative pattern with age within the entire complex (R_mean_= −0.14) (Figure 2 f; Table S9).

**Figure 2:**
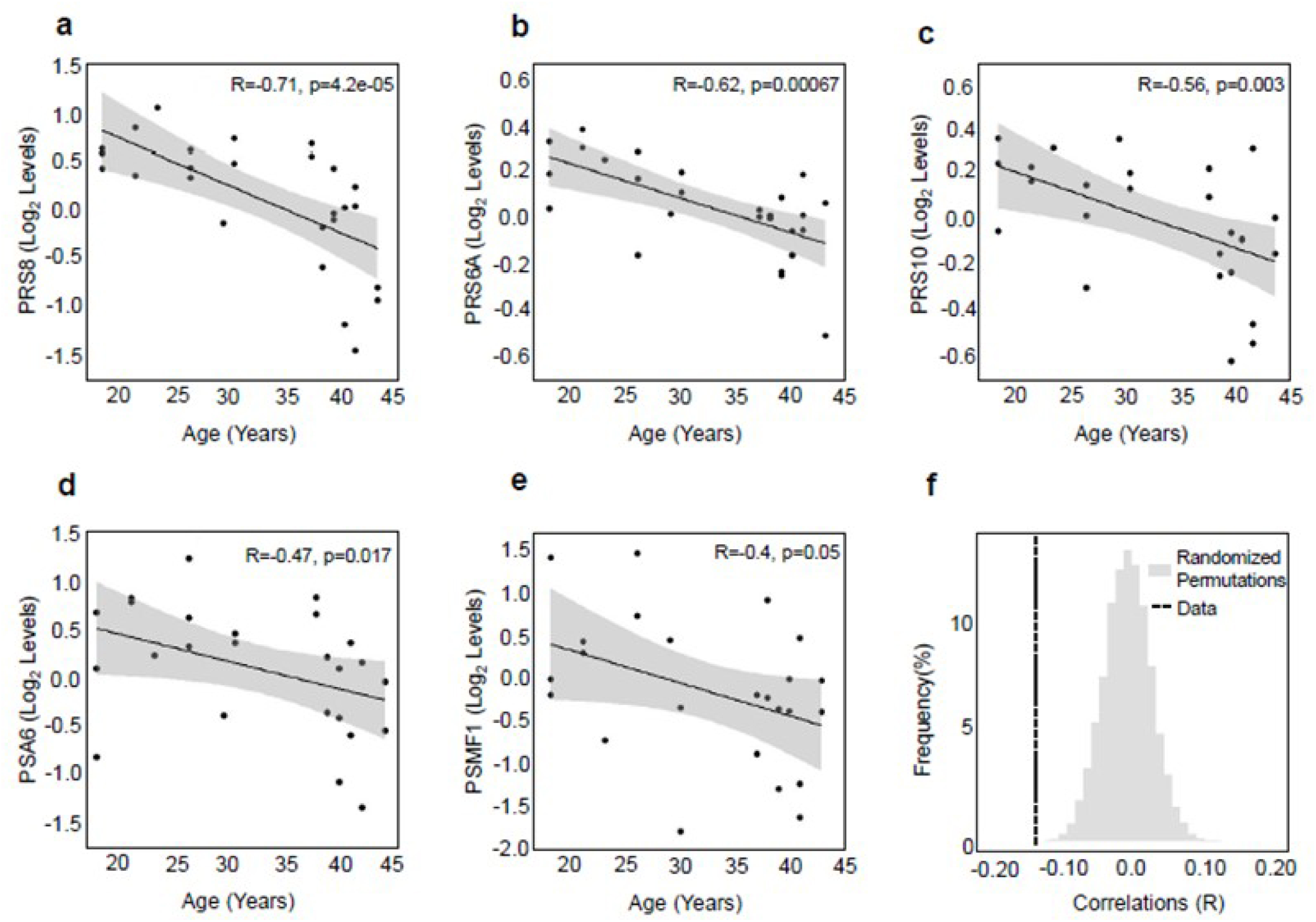
Age effect on the proteasome complex levels. a-e) Scatter plots showing the levels of the proteasome complex subunits which were found to be strongly correlated (p.adj ≤ 0.02, p ≤0.05, |R| ≥ 0.5 with age or to have moderate correlation (p ≤0.05, |R| ≥ 0.3) in GV oocytes. f) Distribution of proteasome subunits mean correlation coefficient (R_mean_) of GV oocytes with age in 10^4^ times randomised data; the proteasome subunits mean correlation coefficient in the original data is represented by the dashed line.

Focusing on the TRiC complex, eight subunits of the complex were quantified in the data (Figure 3 a; Table S10). Of these, four were negatively correlated with age, TCPH, TCPA, TCPQ, TCPE (R= −0.57 to −0.40) (Figure 3 b; Table S10), leading again to a non-random negative pattern in the entire complex with age (R_mean_ = −0.36) (Figure 3 c; Table S9). Finally, pathways such as WNT signaling, regulation of telomerase RNA localization and Arp 2/3 complex mediated actin nucleation pathways appeared to be negatively influenced by age in GV oocytes (Table S11).

**Figure 3:**
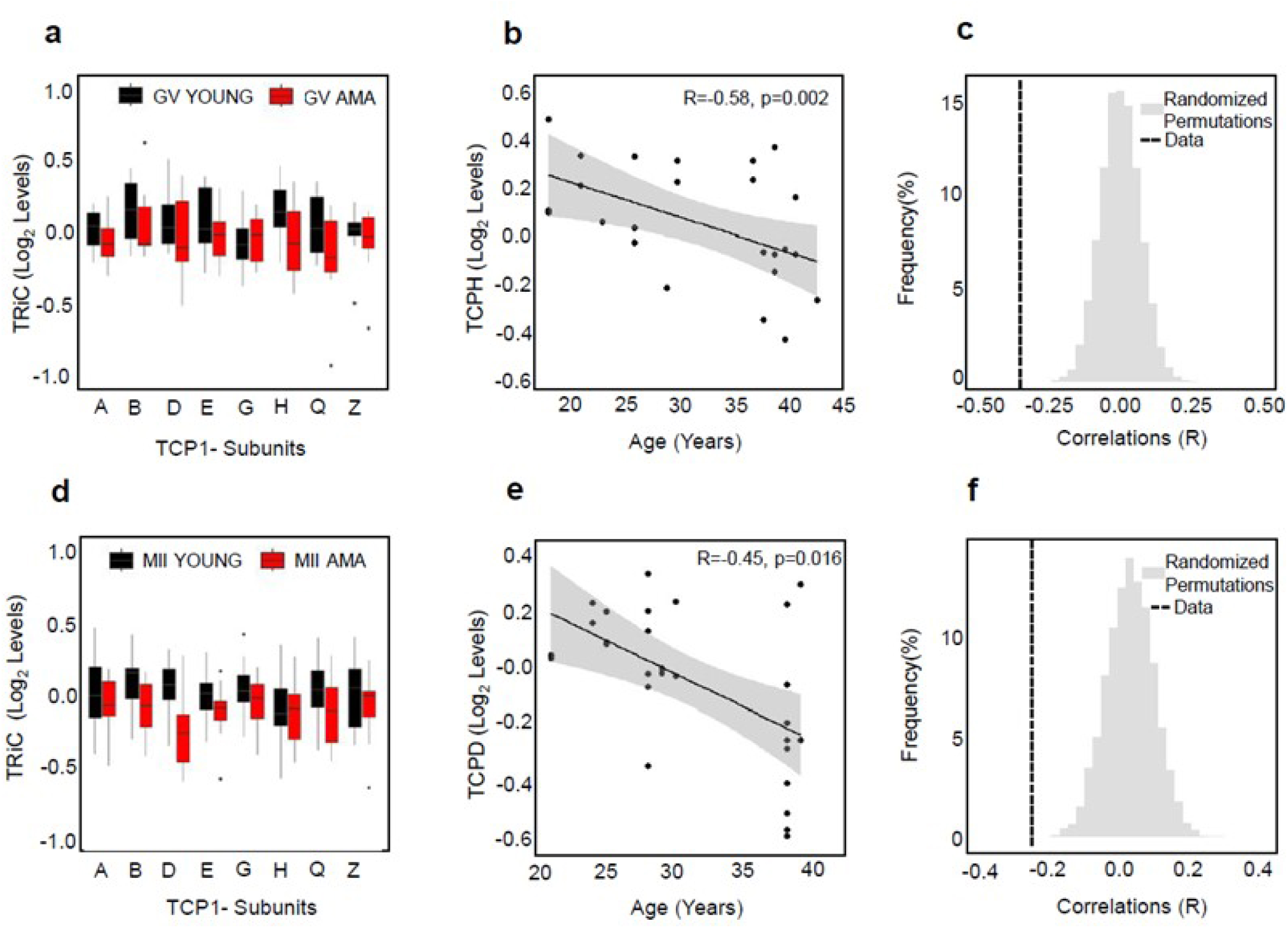
Age effect on the TRiC Complex levels. a) Box plots showing the levels of the TRiC complex subunits in GV oocytes from Young and AMA group. b) Scatter plot showing the levels of the TRiC complex subunit TCPH which was found to be strongly correlated with age in GV oocytes (p.adj ≤ 0.02, p ≤0.05, |R| ≥ 0.5). c) Distribution of TRiC subunits mean correlation coefficient (R_mean_) of GV oocytes with age in 10^4^ times randomised data; the TRiC subunits mean correlation coefficient in the original data is represented by the dashed line. d) Box plots showing the levels of the TRiC complex subunits in MII oocytes from Young and AMA group. b) Scatter plot showing the levels of the TRiC complex subunit TCPD which shown a moderate correlation with age in MII oocytes (p ≤0.05, |R| ≥ 0.3). c) Distribution of TRiC subunits mean correlation coefficient (R_mean_) of MII oocytes with age in 10^4^ times randomised data; the TRiC subunits mean correlation coefficient in the original data is represented by the dashed line.

### Age-related changes in the proteome of mature MII oocytes

Next, we focused on MII oocytes. We aimed to explore whether differences in protein levels in MII oocytes from women with AMA arise after the completion of the first meiotic division or as a consequence of alterations that occurred during the GV stage. For instance, reduced protein abundance in GV oocytes may lead to aberrant abundance of downstream proteins in MII oocytes. Our analysis revealed that the abundance of seven proteins in MII oocytes showed strong age-related correlations. Among them, five proteins were negatively and two positively correlated with age (Table 2). We identified a further 99 proteins that exhibited moderate correlations with age (Table S12).

Moreover, the TriC complex’s abundance exhibited a non-random negative pattern in MII oocytes with maternal age (R_mean_= −0.26) (Figure 3, d-f; Tables S13 and S14). Interestingly, we found that the levels of two isoforms of tubulin beta, TBB5 and TBB8B, which are known targets of the TRiC complex, increase with age (R= 0.47 and 0.40, respectively) (Table S12). Moreover, our analysis reaffirmed that AMA is associated with alterations of the telomerase RNA localization process, as in GV (Table S15).

### Proteasome activity is crucial during the final steps of meiotic maturation

Our findings revealed a decline in the levels of the proteasome complex from the GV to MII stage of oocyte maturation, and we also observed a negative correlation between its abundance and age. Considering the crucial role of the proteasome in meiosis, these results prompted us to undertake further investigations to study its role during oocyte maturation. Understanding the dynamics of the proteasome complex during oocyte development is of significant interest, as the proteasome regulates meiosis progression and plays a pivotal role in protein degradation and maintaining cellular proteostasis.

We primarily stained the proteasome of immature GV oocytes from young women (age ≤ 35 years old) with an antibody against the 20S (alpha and beta) subunits to identify their cellular distribution. We observed that the complex was mainly localized in the nucleus (germinal vesicle) of oocytes, which suggests a possible role in chromatin reorganization and spindle formation before GVBD (Figure 4 a). To further examine this hypothesis, we cultured GV oocytes from young women in rIVM medium in the presence or absence of the proteasome inhibitor MG-132 for 6 hours. After the initial culture, oocytes were washed and placed in fresh rIVM media for an additional 42 hours. Following the extended culture period, the maturation rate was assessed. The oocytes that successfully reached the MII stage (IVM-MII) were then further analyzed to evaluate the alignment of their chromosomes in the metaphase II plate. The results showed that 3 out of 5 GV and 4 out of 4 MI oocytes showed successful maturation (IVM-MII) in the control group, presenting with aligned chromosomes in the metaphase II plate (Figure 4 b to d). Two control GV oocytes were degenerated during the rIVM culture. In contrast, the GV oocytes that were cultured in the presence of the proteasome inhibitor appeared either arrested at the MI stage (2 out of 5) or reached the MII stage (3 out of 5). However, all three mature presented with misaligned chromosomes (Figure 4, b to d).

**Figure 4:**
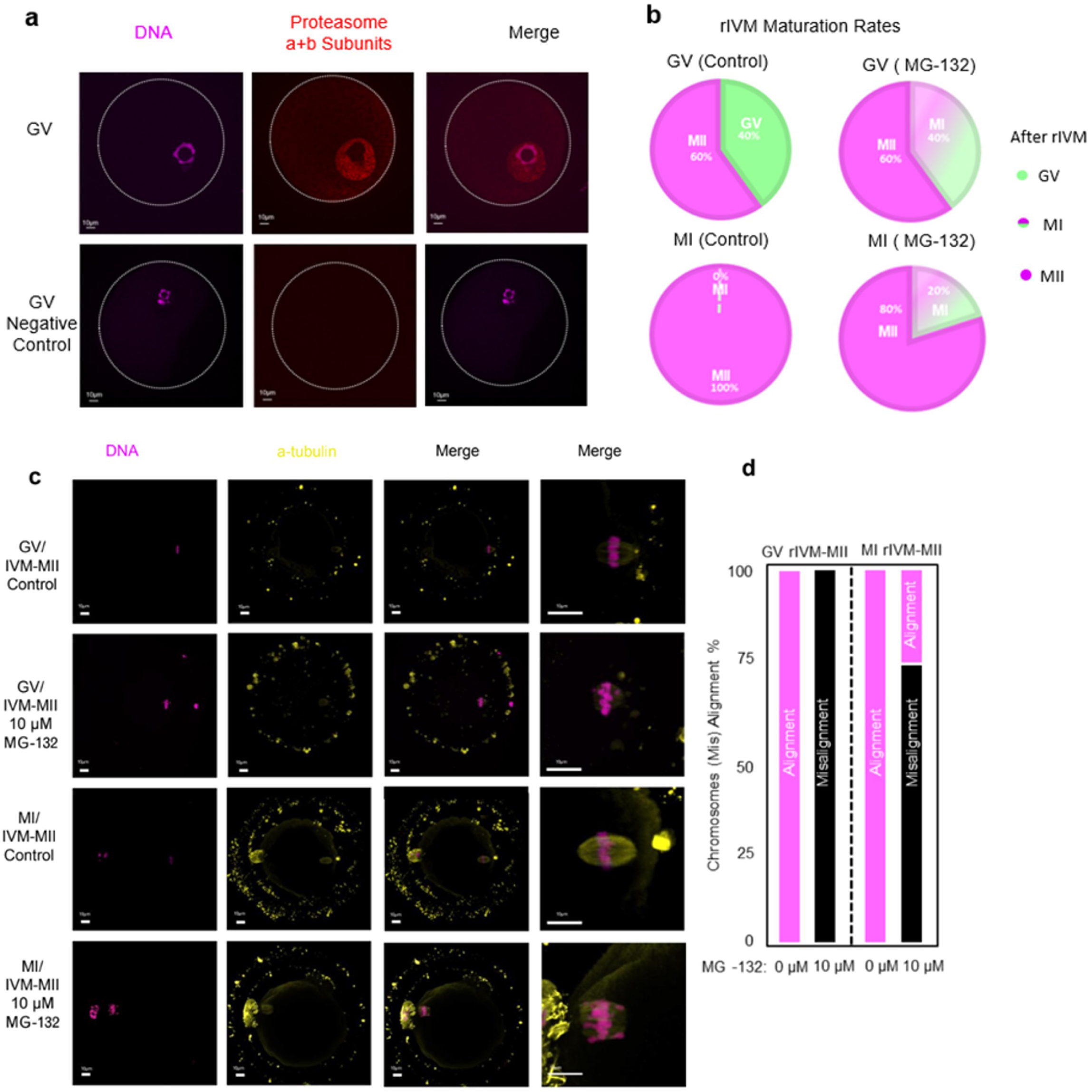
Functional analysis of proteasome complex during the last step of oocyte maturation. (a) Immunofluorescence staining of the proteasome and chromosomes in human GV oocytes. The proteasome complex was stained with an antibody against 20S alpha and beta subunits (red) and DNA were counterstained with Hoechst 33342 (blue). Left panel, DNA; middle panel, proteasome; right panel, merged image. Scale = 10 µm. (b) rIVM rates (%) for GV and MI oocytes cultured in presence or absence of 10 µM of the proteasome inhibitor MG-132, for up to 48 h. (c) Immunofluorescence staining of the spindle with an antibody against TUBA (yellow) and chromosomes (DNA, Hoechst 33342) (pink) of human oocytes after rIVM in presence or absence of 10 µM of MG132. Scale = 10 µm. (d) Percentages of rIVM-MII oocytes with correct alignment (pink bar) or misalignment (black bar) of their chromosomes. correct alignment (pink bar) or misalignment (black bar) of their chromosomes.

To assess if proteasome activity is required for oocyte maturation after GVBD and closer to the time of chromosome segregation, we performed rIVM on MI oocytes. These MI oocytes have already undergone GVBD *in vivo* but had not extruded the first polar body, indicating that they had not completed meiosis I. All MI oocytes cultured in the absence of the proteasome inhibitor reached the IVM-MII stage and displayed correct alignment of their chromosomes in the metaphase plate. However, in the presence of MG-132, four out of five MI oocytes reached the IVM-MII stage and three out of four (75%) exhibited misalignment of their chromosomes, while one IVM-MII oocytes presented aligned chromosomes (Figure 4, b to d).

## DISCUSSION

Women of AMA face subfertility, which is largely attributed to the decreased quality of their oocytes. To gain a deeper understanding into the proteomic basis of the effect of maternal age on oocyte quality, we applied the single-cell plexDIA method to human oocytes.

Using this method, we have characterized the biological processes occurring in oocytes during meiotic maturation and examined the relationship between maternal age and oocyte proteomic content. In total, 2,105 proteins were quantified; building on the findings of two previous single-cell proteomic analyses of human oocytes (Virant-Klun *et al*., 2016; Guo *et al*., 2022).

Our study revealed that human oocytes undergo specific adjustments in the abundance of various proteins during the final stages of meiotic maturation (GV to MII transition). These include, among others, ribosomal subunits, translation factors, cytoskeleton and cell cycle proteins which are likely required for the successful acquisition of developmental competence. These results are in accordance with data from mouse studies, which show that many of the transcripts produced during oocyte growth are stored in translationally inactive ribonucleoprotein particles and translated into proteins at the appropriate time during oocyte maturation (Gosden and Lee, 2010; Susor *et al*., 2015; Luong *et al*., 2020).

In addition, we observed changes in several biological processes including translation, Arp2/3 mediated actin nucleation, maintenance of DNA methylation as well as the proteasome complex between GV and MII oocytes. These findings are also consistent with reports in mice, providing further support for the relevance of these events during oocyte maturation across species (Huo *et al*., 2004; Sun *et al*., 2011; Susor *et al*., 2015; Maenohara *et al*., 2017).

This data provides a comprehensive description of the proteomic landscape of oocyte maturation. Moreover, we saw that this landscape does not undergo major changes with age, since most of the observed protein and pathway alterations during the GV to MII transition exhibit similar patterns in both Young and AMA oocytes. However, we saw a few changes, which could potentially contribute to the loss of developmental competence. In addition, we observed correlations between the abundance of various proteins and maternal age, particularly at the GV stage. Noteworthy is the decline in abundance of several proteasomal subunits with age. The proteasome plays an essential role in oocytes, as it is involved in regulating cell cycle progression (Homer *et al*., 2009) and participates in protein quality control through the ubiquitin-proteasome (UPS) system and maintenance of proteostasis (Café *et al*., 2021).

The proteasome is instrumental in regulating cell cycle progression by targeting key cell cycle factors, including cyclin B and securin. Proteasome-dependent inactivation of the maturation-promoting factor (MPF, composed by cyclin B and CDK1) is required for the successful exit from meiosis I and the transition to meiosis II (Jones, 2004; Li *et al*., 2019). In our study, we noted a drop in the levels of the proteasome complex during the GV to MII transition, with this decline being evident only in young oocytes. Furthermore, we observed a negative correlation between the levels of proteasome subunits and maternal age. Additionally, there was a decline in the levels of CDK1, a critical component of MPF, and its indirect regulator 1433E with age.

Based on these findings, we hypothesize that the age-dependent alteration in the levels of the proteasome, 1433E and CDK1 may lead to dysregulation of MPF activity, potentially resulting in the disruption of meiosis and aneuploidy. Elevated MPF activity has also been observed in aged mouse oocytes, suggested to impact the accurate segregation of chromosomes during oocyte maturation (Koncicka *et al*., 2018). Our hypothesis is further supported by studies on mouse and rat oocytes, which have demonstrated that inhibiting the proteasome results in metaphase I arrest or abnormal meiotic progression, leading to a higher proportion of aneuploid oocytes (Josefsberg *et al*., 2000; Mailhes *et al*. 2002). Similarly, as seen in our study, inhibiting the proteasome in human oocytes led to some oocytes arresting at MI stage, while others reached the MII stage but exhibited misalignment of the chromosomes in the metaphase plate. These observations provide compelling evidence that highlights the importance of proteasome activity during oocyte maturation.

In addition to its role in regulating meiosis and MPF activity, the proteasome complex plays a crucial role in maintaining proteostasis through the UPS system, which is responsible for selectively degrading and eliminating damaged or misfolded proteins (Kelmer *et al*., 2020; Sala *et al*., 2022). Dysfunction of the proteasome and UPS has been associated with aging in various cell types, including oocytes (Keller *et al*., 2000; Mihalas *et al*., 2018; Kelmer *et al*., 2020). We found that proteasomal subunits and UPS-related protein levels, including UCHL1 and CSN3, decline with advancing maternal age. We also identified changes in the abundance of multiple (co)-chaperones, including HSP7C, DJB11, TCPH that participate in protein folding and maintenance of a functional proteome (Chen *et al*., 2017; Fernández-Fernández and Valpuesta, 2018; Grantham, 2020).

The abundance of the TRiC complex, a chaperonin which assists in the folding of about 10% of the proteome, including actin and tubulin isoforms (Sternlicht *et al*., 1993) and participates in proteostatic control of telomerase (Freund *et al*., 2014), was also negatively associated with age in oocytes. Interestingly, the levels of tubulin isoforms TBB5 and TBB8B exhibited a positive correlation in MII with age, which could be potentially attributed to the decreased abundance of the TRiC complex in GV and MII oocytes from AMA women. Whether the activity of TRiC complex is affected in AMA oocytes, resulting in the accumulation of TBB5 and TBB8B, potentially misfolded, is matter of further study.

Taken together, the identified protein alterations suggest a progressive failure of proteostasis in oocytes from women of AMA. The failure of protesostasis could account for the poor quality of oocytes, either independently or by interfering with crucial cellular processes like meiosis, as previously suggested (Sala *et al*., 2022).

Building on our findings, mitochondrial proteins are also influenced by the proteasome-UPS system. Mitochondrial proteins are synthesized in the cytoplasm as unfolded polypeptides and rely on chaperones for proper folding before being imported into the mitochondria (Quiles and Gustafsson, 2020).

This post-translational import mechanism is tightly regulated by the proteasome-UPS system, which ensures the degradation of non-functional polypeptides and damaged proteins that have already been imported into the mitochondria (Krämer *et al*., 2021). In addition, accumulation of mitochondrial precursors in the cytosol has been associated with mitochondria-mediated cell death (mPOS) (Wang and Chen, 2015; Coyne and Chen, 2017). Interestingly, we observed that some mitochondrial proteins displayed increased levels with age in GV oocytes. This accumulation of mitochondrial proteins may be attributed to insufficient post-translational control by the proteasome and chaperones, which are found in lower abundance in GV oocytes from AMA women.

In our analysis of MII oocytes, we identified five proteins with reduced levels and two proteins with increased levels that correlate with age. Notably, among them, IGHG1 and BASP1 have been previously reported to play roles in oocytes. BASP1, is a peptide which probably is involved in fertilization-induced oocyte activation (Zakharova and & Zakharov, 2017) while the immunoglobulins (IGHG1) have been suggested to counteract the increased ROS levels and assist the oocyte to survive in adverse environments (Wang *et al*., 2022). Additionally, DDX4, the human ortholog of VASA, is a well-known germ cell marker that has also been suggested to be involved in the regulation of translation (Castrillon *et al*., 2000; Sundaram *et al*., 2023). The roles of IGHG1, BASP1, and DDX4 in oocyte function and translation regulation suggest that they may be critical factors in maintaining oocyte quality.

Our findings deliver important insights into the proteomic landscape of oocyte maturation and the impact of maternal age on the global oocyte proteome, yet it is essential to acknowledge some limitations. First, to ensure statistical robustness, we chose a stringent criterion of including only proteins present in at least 80% of the samples. While this approach enhanced the reliability of our analysis, it is possible that some proteins in lower abundance were not captured in our study. Secondly, MII oocytes underwent vitrification and warming prior to being included in the study due to clinical protocols. Although these procedures are commonly used in IVF treatment, they may introduce some uncontrolled variability in the proteomic profiles of the oocytes. Additionally, the rIVM maturation of GV and MI oocytes was conducted without the presence of cumulus cells, which may have impacted the maturation process. Nevertheless, the identification of age-correlated proteins and their potential roles in oocyte function provides valuable insights for future research in reproductive medicine. Ultimately, our results shed light on the negative impact of aging on oocyte developmental competence, pointing at the proteostasis and meiosis networks. The altered proteins identified in aged oocytes hold promise as potential targets for interventions aimed at improving oocyte quality and reproductive outcomes in women of AMA. By delving into the complexities of oocyte maturation and aging our study opens new opportunities in the field of reproductive medicine, pave the way for future research into improved treatment options and outcomes for women facing age-related fertility challenges.

## AVAILABILITY OF DATA AND MATERIALS

The data underlying this study will be available from the corresponding author upon reasonable request.

## ACKNOWLEDGMENTS

We would like to thank Laura Sabater from Clinica Eugin for their help in sample handling and all the laboratory staff from the clinics for their support.

## AUTHOR’S ROLE

SG contributed to design the study, collected and processed the samples, analyzed and interpreted the data, and have drafted the manuscript; AP prepared the single oocytes for MS analysis and contributed to the analysis of data; APujol and KL contributed to samples collection; LL contributed to analysis of the immunofluorescence images; TF and MP contributed to the interpretation of the data and substantially revised the manuscript; RV designed the study, contributed in the interpretation of the data, and substantively revised the manuscript; NS contributed in the analysis and interpretation of the data and substantively revised the manuscript; MBM designed the study, analyzed and interpreted the data, and substantively revised the manuscript.

