## Supplementary Tables 1 to 15 for "Single-cell proteomics reveals decreased abundance of proteostasis and meiosis proteins in advanced maternal age oocytes"

**Table S1. Population and Samples for single-cell Proteomic analysis. AFC: antral follicular count; BMI: body mass index; ND: Not determined**

| Woman | Age (Years) | AFC (n) | BMI (kg/m2) | Group | Oocyte # | Maturation Stage | Quality Control |
|---|---|---|---|---|---|---|---|
| Woman 1 | 26 | 13 | 18.07 | Young | oocyte 1 | GV | Pass |
| Woman 2 | 18 | 24 | 21.22 | Young | oocyte 2 | GV | Pass |
| Woman 3 | 30 | 32 | 25.16 | Young | oocyte 3 | GV | Pass |
| Woman 4 | 26 | 36 | 21.52 | Young | oocyte 4 | GV | Pass |
| Woman 5 | 23 | 16 | 22.66 | Young | oocyte 5 | GV | Excluded |
| Woman 6 | 20 | 35 | 22.38 | Young | oocyte 6 | GV | Excluded |
| Woman 7 | 19 | 35 | 25.76 | Young | oocyte 7 | GV | Excluded |
| Woman 8 | 24 | 27 | 24.34 | Young | oocyte 8 | GV | Excluded |
| Woman 9 | 23 | 40 | 24.23 | Young | oocyte 9 | GV | Pass |
| Woman 10 | 21 | 30 | 28.31 | Young | oocyte 10 | GV | Pass |
| Woman 11 | 29 | 13 | 23.98 | Young | oocyte 11 | GV | Pass |
| Woman 12 | 21 | 29 | 23.14 | Young | oocyte 12 | GV | Pass |
| Woman 13 | 29 | 30 | 23.42 | Young | oocyte 13 | GV | Excluded |
| Woman 14 | 30 | 19 | 21.24 | Young | oocyte 14 | GV | Pass |
| Woman 15 | 26 | 24 | 20.53 | Young | oocyte 15 | GV | Pass |
| Woman 16 | 18 | 31 | 19.43 | Young | oocyte 16 | GV | Pass |
| Woman 17 | 18 | 11 | 19.92 | Young | oocyte 17 | GV | Pass |
| Woman 18 | 24 | 12 | 22.57 | Young | oocyte 18 | GV | Excluded |
| Woman 19 | 39 | 7 | 25.6 | AMA | oocyte 19 | GV | Pass |
| Woman 20 | 37 | 7 | 20.7 | AMA | oocyte 20 | GV | Pass |
| Woman 21 | 37 | 14 | 17.42 | AMA | oocyte 21 | GV | Excluded |
| Woman 22 | 37 | 9 | 23.1 | AMA | oocyte 22 | GV | Pass |
| Woman 23 | 41 | 6 | ND | AMA | oocyte 23 | GV | Pass |
| Woman 24 | 40 | 2 | ND | AMA | oocyte 24 | GV | Excluded |
| Woman 25 | 37 | 15 | 22.3 | AMA | oocyte 25 | GV | Excluded |
| Woman 26 | 41 | 8 | 28.38 | AMA | oocyte 26 | GV | Pass |
| Woman 27 | 38 | 5 | 18.34 | AMA | oocyte 27 | GV | Pass |
| Woman 28 | 43 | 8 | 19.3 | AMA | oocyte 28 | GV | Pass |
| Woman 29 | 39 | 15 | 20.7 | AMA | oocyte 29 | GV | Pass |
| Woman 30 | 38 | 5 | 25.3 | AMA | oocyte 30 | GV | Pass |
| Woman 31 | 37 | 14 | 17.92 | AMA | oocyte 31 | GV | Excluded |
| Woman 32 | 39 | 9 | 17.97 | AMA | oocyte 32 | GV | Pass |
| Woman 33 | 41 | 13 | 23.23 | AMA | oocyte 33 | GV | Pass |
| Woman 34 | 43 | 5 | 20.76 | AMA | oocyte 34 | GV | Pass |
| Woman 35 | 40 | 10 | 26.4 | AMA | oocyte 35 | GV | Pass |
| Woman 36 | 40 | 13 | 27.22 | AMA | oocyte 36 | GV | Pass |
| Woman 37 | 29 | 20 | 19.02 | Young | oocyte 37 | IVO-MII | Pass |
| | | | | | oocyte 38 | IVO-MII | Pass |
| Woman 38 | 28 | 22 | 18.44 | Young | oocyte 39 | IVO-MII | Pass |
| | | | | | oocyte 40 | IVO-MII | Pass |
| Woman 39 | 30 | 15 | 21.97 | Young | oocyte 41 | IVO-MII | Pass |
| | | | | | oocyte 42 | IVO-MII | Pass |
| Woman 40 | 28 | 10 | 28.64 | Young | oocyte 43 | IVO-MII | Pass |
| | | | | | oocyte 44 | IVO-MII | Pass |
| Woman 41 | 21 | 30 | 17.18 | Young | oocyte 45 | IVO-MII | Pass |
| | | | | | oocyte 46 | IVO-MII | Pass |
| Woman 42 | 25 | 30 | 29.82 | Young | oocyte 47 | IVO-MII | Pass |
| | | | | | oocyte 48 | IVO-MII | Discarded |
| Woman 43 | 25 | 12 | 24.44 | Young | oocyte 49 | IVO-MII | Pass |
| | | | | | oocyte 50 | IVO-MII | Pass |
| Woman 44 | 28 | 23 | 21.41 | Young | oocyte 51 | IVO-MII | Pass |
| | | | | | oocyte 52 | IVO-MII | Pass |
| Woman 45 | 24 | 8 | 20.69 | Young | oocyte 53 | IVO-MII | Pass |
| | | | | | oocyte 54 | IVO-MII | Pass |
| Woman 46 | 38 | 16 | 16.9 | AMA | oocyte 55 | IVO-MII | Pass |
| | | | | | oocyte 56 | IVO-MII | Excluded |
| Woman 47 | 39 | 13 | 37.11 | AMA | oocyte 57 | IVO-MII | Pass |
| | | | | | oocyte 58 | IVO-MII | Excluded |
| Woman 48 | 38 | 12 | 22.06 | AMA | oocyte 59 | IVO-MII | Pass |
| | | | | | oocyte 60 | IVO-MII | Pass |
| Woman 49 | 39 | 7 | 19.49 | AMA | oocyte 61 | IVO-MII | Excluded |
| | | | | | oocyte 62 | IVO-MII | Pass |
| Woman 50 | 38 | 8 | 20.42 | AMA | oocyte 63 | IVO-MII | Pass |

| | | | | | | | |
|---|---|---|---|---|---|---|---|
| Woman 50 | 38 | 8 | 20.42 | AMA | oocyte 64 | IVO-MII | Pass |
| Woman 51 | 38 | 14 | 19.07 | AMA | oocyte 65 | IVO-MII | Pass |
| | | | | | oocyte 66 | IVO-MII | Pass |
| Woman 52 | 38 | 16 | 22.31 | AMA | oocyte 67 | IVO-MII | Pass |
| | | | | | oocyte 68 | IVO-MII | Pass |

| | | | |
|---|---|---|---|
| mean Young | 24.5 | 23 | 22.57 |
| SD Young | 3.9 | 9.1 | 3.2 |
| mean AMA | 39 | 10 | 22.26 |
| SD AMA | 1.7 | 4 | 4.6 |
| t-test (p-value) | 2.5E-09 | 4.1E-06 | 0.49 |

**Table S2. Population and Samples for functional analysis. AFC: antral follicular count; BMI: body mass index**

| Woman | Age (Years) | AFC (n) | BMI (kg/m2) | Oocyte # | Maturation Stage | [MG-132] (µM) |
|---|---|---|---|---|---|---|
| Woman 1 | 22 | 24 | 17.65 | oocyte 1 | MI | 10 |
| Woman 2 | 34 | 27 | 26.03 | oocyte 2 | GV | 10 |
| | 34 | 27 | 26.03 | oocyte 3 | GV | 10 |
| Woman 3 | 27 | 24 | 18.36 | oocyte 4 | MI | 0 |
| | 27 | 24 | 18.36 | oocyte 5 | MI | 10 |
| Woman 4 | 34 | 27 | 26.03 | oocyte 6 | GV | 0 |
| Woman 5 | 26 | 18 | 25.53 | oocyte 7 | MI | 0 |
| | 26 | 18 | 25.53 | oocyte 8 | MI | 10 |
| Woman 6 | 25 | 27 | 28.52 | oocyte 9 | GV | 0 |
| | 25 | 27 | 28.52 | oocyte 10 | MI | 0 |
| | 25 | 27 | 28.52 | oocyte 11 | MI | 10 |
| | 25 | 27 | 28.52 | oocyte 12 | GV | 0 |
| | 25 | 27 | 28.52 | oocyte 13 | GV | 10 |
| | 25 | 27 | 28.52 | oocyte 14 | GV | 10 |
| Woman 7 | 32 | 13 | 23.92 | oocyte 15 | MI | 0 |
| | 32 | 13 | 23.92 | oocyte 16 | GV | 10 |
| | 32 | 13 | 23.92 | oocyte 17 | MI | 10 |
| Woman 8 | 30 | 18 | 24.92 | oocyte 18 | GV | 0 |
| Woman 9 | 29 | 18 | 22.75 | oocyte 19 | GV | 0 |

**Table S3. Comparative abundance levels of identified proteins during the final step of oocyte maturation (Young group): MII *vs.* GV oocytes. Significant differences: fold change (FC) >| 1.5|; p.adj ≤ 0.05**

| | Uniprot ID | Protein name | Log2 (FC) | P_value | P.adjusted |
|---|---|---|---|---|---|
| Significantly abundant proteins in MII | Q9H568 | ACTL8 | 2.729 | 0.000 | 0.000 |
| | P26358-2 | DNMT1 | 2.335 | 0.000 | 0.000 |
| | Q9Y6A5 | TACC3 | 2.083 | 0.000 | 0.001 |
| | P12004 | PCNA | 1.877 | 0.000 | 0.002 |
| | P0C1S8 | WEE2 | 1.728 | 0.000 | 0.000 |
| | O94988 | FA13A | 1.649 | 0.000 | 0.007 |
| | Q02750 | MP2K1 | 1.433 | 0.000 | 0.011 |
| | Q96MN2 | NALP4 | 1.405 | 0.000 | 0.001 |
| | P19823 | ITIH2 | 1.355 | 0.000 | 0.008 |
| | Q9NWY4 | HPF1 | 1.321 | 0.001 | 0.023 |
| | O14965 | AURKA | 1.296 | 0.000 | 0.005 |
| | A6NKN8 | PC4L1 | 1.266 | 0.000 | 0.000 |
| | O60825-2 | F262 | 1.254 | 0.000 | 0.006 |
| | P27105 | STOM | 1.183 | 0.002 | 0.038 |
| | P19827 | ITIH1 | 1.160 | 0.001 | 0.029 |
| | Q8WTV0-2 | SCRB1 | 1.073 | 0.001 | 0.029 |
| | O00560 | SDCB1 | 1.064 | 0.001 | 0.025 |
| | O60566 | BUB1B | 1.050 | 0.000 | 0.001 |
| | P10909 | CLUS | 1.023 | 0.003 | 0.050 |
| | Q96EA4 | SPDLY | 0.888 | 0.000 | 0.002 |
| | Q8WVJ2 | NUDC2 | 0.763 | 0.001 | 0.030 |
| | Q6UW68 | TM205 | 0.714 | 0.000 | 0.009 |
| | Q13277 | STX3 | 0.622 | 0.003 | 0.048 |
| | P28066 | PSA5 | 0.614 | 0.003 | 0.050 |
| | P58546 | MTPN | 0.601 | 0.002 | 0.042 |
| | O75347 | TBCA | 0.595 | 0.003 | 0.050 |
| | O14737 | PDCD5 | 0.593 | 0.001 | 0.020 |
| | O95433 | AHSA1 | 0.588 | 0.000 | 0.015 |
| | Q06033 | ITIH3 | 1.525 | 0.003 | 0.054 |
| | P07942 | LAMB1 | 1.070 | 0.029 | 0.215 |
| | Q09028 | RBBP4 | 1.053 | 0.033 | 0.231 |
| | O43324 | MCA3 | 1.033 | 0.077 | 0.341 |
| | A6NNZ2 | TBB8B | 1.015 | 0.015 | 0.141 |
| | Q08431 | MFGM | 0.988 | 0.006 | 0.077 |
| | Q92506 | DHB8 | 0.967 | 0.006 | 0.081 |
| | P08572 | CO4A2 | 0.964 | 0.093 | 0.377 |
| | P43007 | SATT | 0.913 | 0.073 | 0.330 |
| | Q99726 | ZNT3 | 0.856 | 0.003 | 0.054 |
| | P18084 | ITB5 | 0.834 | 0.013 | 0.128 |
| | Q9BUN8 | DERL1 | 0.816 | 0.020 | 0.171 |
| | P17302 | CXA1 | 0.804 | 0.227 | 0.582 |
| | Q658Y4 | F91A1 | 0.775 | 0.053 | 0.288 |
| | Q96PU4 | UHRF2 | 0.764 | 0.018 | 0.161 |
| | Q15043 | S39AE | 0.763 | 0.042 | 0.261 |
| | Q9UHF5 | IL17B | 0.757 | 0.053 | 0.288 |

| | | | | |
|---|---|---|---|---|
| Q86WR0 | CCD25 | 0.746 | 0.023 | 0.187 |
| P41222 | PTGDS | 0.735 | 0.107 | 0.400 |
| O60716 | CTND1 | 0.716 | 0.007 | 0.085 |
| Q9H9J2 | RM44 | 0.705 | 0.043 | 0.261 |
| P02748 | CO9 | 0.694 | 0.064 | 0.314 |
| P21754 | ZP3 | 0.685 | 0.018 | 0.161 |
| Q8TDM6 | DLG5 | 0.684 | 0.051 | 0.288 |
| Q8NFF5 | FAD1 | 0.680 | 0.054 | 0.289 |
| P47897 | SYQ | 0.677 | 0.023 | 0.187 |
| Q9Y5J7 | TIM9 | 0.670 | 0.082 | 0.344 |
| Q9BSH5 | HDHD3 | 0.662 | 0.016 | 0.151 |
| Q96EK5 | KBP | 0.657 | 0.007 | 0.086 |
| O00273 | DFFA | 0.651 | 0.141 | 0.469 |
| Q16595 | FRDA | 0.649 | 0.128 | 0.446 |
| Q9Y285 | SYFA | 0.648 | 0.027 | 0.203 |
| P05556 | ITB1 | 0.636 | 0.027 | 0.202 |
| Q9Y4E8 | UBP15 | 0.634 | 0.047 | 0.275 |
| P23229 | ITA6 | 0.623 | 0.009 | 0.101 |
| P04004 | VTNC | 0.607 | 0.034 | 0.231 |
| P60852 | ZP1 | 0.599 | 0.012 | 0.123 |
| Q05996 | ZP2 | 0.592 | 0.030 | 0.215 |
| P56279 | TCL1A | 0.591 | 0.005 | 0.073 |
| O60831 | PRAF2 | 0.588 | 0.272 | 0.635 |
| Q6UXH1 | CREL2 | 0.586 | 0.064 | 0.314 |
| Q7Z7N9 | T179B | 0.585 | 0.286 | 0.636 |
| P36405 | ARL3 | 0.572 | 0.001 | 0.023 |
| Q99614 | TTC1 | 0.572 | 0.064 | 0.314 |
| P62826 | RAN | 0.572 | 0.006 | 0.077 |
| P55327 | TPD52 | 0.569 | 0.093 | 0.377 |
| P06748 | NPM | 0.564 | 0.080 | 0.341 |
| P02652 | APOA2 | 0.564 | 0.073 | 0.330 |
| P63167 | DYL1 | 0.562 | 0.002 | 0.038 |
| Q96JY0 | MAEL | 0.562 | 0.004 | 0.066 |
| Q12836 | ZP4 | 0.559 | 0.227 | 0.582 |
| P37837 | TALDO | 0.551 | 0.011 | 0.111 |
| Q9Y263 | PLAP | 0.547 | 0.000 | 0.013 |
| Q86TP1 | PRUN1 | 0.543 | 0.222 | 0.582 |
| P60520 | GBRL2 | 0.541 | 0.001 | 0.020 |
| Q9NR31 | SAR1A | 0.538 | 0.038 | 0.247 |
| Q15843 | NEDD8 | 0.538 | 0.024 | 0.190 |
| Q13137-4 | CACO2 | 0.537 | 0.042 | 0.261 |
| P12235 | ADT1 | 0.535 | 0.074 | 0.332 |
| Q15005 | SPCS2 | 0.527 | 0.139 | 0.467 |
| P02649 | APOE | 0.526 | 0.140 | 0.467 |
| Q01433 | AMPD2 | 0.511 | 0.225 | 0.582 |
| Q13491-4 | GPM6B | 0.507 | 0.616 | 0.869 |
| Q9BY44 | EIF2A | 0.504 | 0.012 | 0.123 |
| Q9NRX4 | PHP14 | 0.500 | 0.027 | 0.202 |
| P78562 | PHEX | 0.500 | 0.283 | 0.635 |
| P46108 | CRK | 0.499 | 0.043 | 0.261 |
| Q9H300 | PARL | 0.499 | 0.006 | 0.077 |

| | | | | |
|---|---|---|---|---|
| Q86UY8 | NT5D3 | 0.497 | 0.066 | 0.314 |
| Q5XKP0 | MIC13 | 0.497 | 0.305 | 0.643 |
| P09972 | ALDOC | 0.494 | 0.097 | 0.378 |
| P28838 | AMPL | 0.491 | 0.000 | 0.020 |
| Q14894 | CRYM | 0.491 | 0.006 | 0.077 |
| Q9BT09 | CNPY3 | 0.491 | 0.043 | 0.261 |
| Q9Y2Z9 | COQ6 | 0.489 | 0.386 | 0.717 |
| Q5VV41 | ARHGG | 0.487 | 0.305 | 0.643 |
| Q5SW96 | ARH | 0.487 | 0.073 | 0.330 |
| Q9P2B2 | FPRP | 0.487 | 0.059 | 0.302 |
| Q99611 | SPS2 | 0.483 | 0.059 | 0.302 |
| Q8NFU3 | TSTD1 | 0.475 | 0.066 | 0.314 |
| P22307 | SCP2 | 0.473 | 0.140 | 0.467 |
| Q719I0 | AHSA2 | 0.473 | 0.146 | 0.484 |
| Q9UKK9 | NUDT5 | 0.471 | 0.073 | 0.330 |
| P07108-4 | ACBP | 0.470 | 0.499 | 0.790 |
| P01876 | IGHA1 | 0.468 | 0.166 | 0.509 |
| O14562 | UBFD1 | 0.466 | 0.021 | 0.176 |
| Q9BTV4 | TMM43 | 0.466 | 0.201 | 0.572 |
| Q02978 | M2OM | 0.466 | 0.129 | 0.448 |
| O00214-2 | LEG8 | 0.465 | 0.166 | 0.509 |
| Q9Y294 | ASF1A | 0.457 | 0.053 | 0.288 |
| P04792 | HSPB1 | 0.453 | 0.325 | 0.655 |
| P07738 | PMGE | 0.452 | 0.011 | 0.111 |
| O75521 | ECI2 | 0.445 | 0.002 | 0.042 |
| P63172 | DYLT1 | 0.444 | 0.053 | 0.288 |
| Q96GG9 | DCNL1 | 0.442 | 0.481 | 0.781 |
| Q9UMX5 | NENF | 0.440 | 0.140 | 0.467 |
| P61081 | UBC12 | 0.439 | 0.167 | 0.511 |
| P61604 | CH10 | 0.432 | 0.000 | 0.015 |
| P06396-2 | GELS | 0.431 | 0.003 | 0.048 |
| Q9NWV4 | CZIB | 0.430 | 0.152 | 0.489 |
| O43491-4 | E41L2 | 0.428 | 0.212 | 0.578 |
| O00217 | NDUS8 | 0.427 | 0.030 | 0.215 |
| P09382 | LEG1 | 0.426 | 0.030 | 0.215 |
| O75477 | ERLN1 | 0.424 | 0.150 | 0.489 |
| Q3ZCM7 | TBB8 | 0.422 | 0.014 | 0.137 |
| P20339 | RAB5A | 0.421 | 0.443 | 0.747 |
| O75380 | NDUS6 | 0.419 | 0.285 | 0.635 |
| P23919 | KTHY | 0.418 | 0.134 | 0.460 |
| P29218 | IMPA1 | 0.417 | 0.006 | 0.077 |
| Q9C0E8 | LNP | 0.414 | 0.100 | 0.382 |
| Q8NE86 | MCU | 0.412 | 0.478 | 0.778 |
| P04066 | FUCO | 0.411 | 0.227 | 0.582 |
| P10768 | ESTD | 0.411 | 0.080 | 0.341 |
| Q9UBS4 | DJB11 | 0.410 | 0.000 | 0.015 |
| Q8N283 | ANR35 | 0.409 | 0.323 | 0.655 |
| P06396 | GELS | 0.406 | 0.002 | 0.033 |
| Q9BRG1 | VPS25 | 0.401 | 0.384 | 0.715 |
| O95988 | TCL1B | 0.400 | 0.283 | 0.635 |
| Q9Y508 | RN114 | 0.398 | 0.222 | 0.582 |

| | | | | |
|---|---|---|---|---|
| P02768 | ALBU | 0.397 | 0.879 | 0.978 |
| P51148 | RAB5C | 0.397 | 0.088 | 0.365 |
| Q9NTJ5 | SAC1 | 0.396 | 0.034 | 0.231 |
| Q8WVM8 | SCFD1 | 0.394 | 0.053 | 0.288 |
| P46976 | GLYG | 0.394 | 0.006 | 0.077 |
| Q9BV19 | CA050 | 0.393 | 0.059 | 0.302 |
| P30040 | ERP29 | 0.391 | 0.001 | 0.025 |
| A0MZ66 | SHOT1 | 0.391 | 0.018 | 0.161 |
| P07686 | HEXB | 0.390 | 0.556 | 0.835 |
| Q7Z4H3 | HDDC2 | 0.389 | 0.419 | 0.735 |
| Q8N668 | COMD1 | 0.386 | 0.066 | 0.314 |
| P09619 | PGFRB | 0.384 | 0.034 | 0.231 |
| Q16851 | UGPA | 0.384 | 0.003 | 0.050 |
| O00170 | AIP | 0.383 | 0.325 | 0.655 |
| Q16795 | NDUA9 | 0.383 | 0.195 | 0.557 |
| P09493-10 | TPM1 | 0.383 | 0.027 | 0.202 |
| A8MVS5 | HIDE1 | 0.382 | 0.100 | 0.382 |
| Q6DD88 | ATLA3 | 0.382 | 0.042 | 0.261 |
| P36957 | ODO2 | 0.380 | 0.002 | 0.033 |
| P47224 | MSS4 | 0.380 | 0.277 | 0.635 |
| P18206 | VINC | 0.378 | 0.080 | 0.341 |
| P46926 | GNPI1 | 0.377 | 0.024 | 0.190 |
| P22059 | OSBP1 | 0.375 | 0.012 | 0.123 |
| P05026 | AT1B1 | 0.372 | 0.264 | 0.627 |
| P78540 | ARGI2 | 0.371 | 0.002 | 0.042 |
| O95881 | TXD12 | 0.371 | 0.053 | 0.288 |
| Q9BUT1 | BDH2 | 0.371 | 0.128 | 0.446 |
| Q05639 | EF1A2 | 0.370 | 0.904 | 0.985 |
| O60825 | F262 | 0.370 | 0.073 | 0.330 |
| Q92905 | CSN5 | 0.368 | 0.038 | 0.247 |
| Q8WXX5 | DNJC9 | 0.366 | 0.018 | 0.161 |
| P09211 | GSTP1 | 0.365 | 0.211 | 0.576 |
| Q9HD67 | MYO10 | 0.365 | 0.231 | 0.591 |
| P21912 | SDHB | 0.364 | 0.027 | 0.202 |
| O75746 | S2512 | 0.364 | 0.080 | 0.341 |
| Q9BS26 | ERP44 | 0.360 | 0.195 | 0.557 |
| O43684 | BUB3 | 0.358 | 0.183 | 0.538 |
| P61006 | RAB8A | 0.353 | 0.586 | 0.857 |
| P13611 | CSPG2 | 0.353 | 0.329 | 0.663 |
| Q07021 | C1QBP | 0.350 | 0.059 | 0.302 |
| P35754 | GLRX1 | 0.349 | 0.048 | 0.275 |
| Q15814 | TBCC | 0.348 | 0.195 | 0.557 |
| P22695 | QCR2 | 0.345 | 0.053 | 0.288 |
| P11021 | BIP | 0.345 | 0.006 | 0.077 |
| Q9BWD1 | THIC | 0.344 | 0.009 | 0.101 |
| O15371 | EIF3D | 0.344 | 0.399 | 0.719 |
| Q13492 | PICAL | 0.342 | 0.077 | 0.341 |
| P01031 | CO5 | 0.341 | 0.211 | 0.576 |
| Q15084 | PDIA6 | 0.340 | 0.001 | 0.029 |
| P51149 | RAB7A | 0.339 | 0.009 | 0.101 |
| A6NDX5 | ZN840 | 0.339 | 0.011 | 0.111 |

| | | | | |
|---|---|---|---|---|
| Q9NP58 | ABCB6 | 0.335 | 0.225 | 0.582 |
| Q8N2F6 | ARM10 | 0.333 | 0.347 | 0.674 |
| P10645 | CMGA | 0.333 | 0.499 | 0.790 |
| P57105 | SYJ2B | 0.332 | 0.128 | 0.446 |
| P35613 | BASI | 0.330 | 0.303 | 0.641 |
| Q9UMY4 | SNX12 | 0.329 | 0.009 | 0.101 |
| P23229-7 | ITA6 | 0.327 | 0.422 | 0.737 |
| P30038 | AL4A1 | 0.326 | 0.048 | 0.275 |
| Q96T88 | UHRF1 | 0.324 | 0.030 | 0.215 |
| Q5JRA6 | TGO1 | 0.322 | 0.080 | 0.341 |
| O43639 | NCK2 | 0.322 | 0.195 | 0.557 |
| Q5TZA2 | CROCC | 0.321 | 0.152 | 0.489 |
| Q13368 | MPP3 | 0.321 | 0.294 | 0.641 |
| Q6RW13 | ATRAP | 0.320 | 0.128 | 0.446 |
| Q14008 | CKAP5 | 0.319 | 0.027 | 0.202 |
| Q9Y679 | AUP1 | 0.319 | 0.152 | 0.489 |
| P99999 | CYC | 0.316 | 0.018 | 0.161 |
| A4D1E9 | GTPBA | 0.314 | 0.283 | 0.635 |
| Q9ULD0 | OGDHL | 0.313 | 0.200 | 0.570 |
| Q9H3P7 | GCP60 | 0.312 | 0.180 | 0.531 |
| P53999 | TCP4 | 0.312 | 0.223 | 0.582 |
| P05023 | AT1A1 | 0.311 | 0.166 | 0.509 |
| Q9H773 | DCTP1 | 0.310 | 0.303 | 0.641 |
| Q15382 | RHEB | 0.308 | 0.371 | 0.696 |
| O43795 | MYO1B | 0.306 | 0.227 | 0.582 |
| P30101 | PDIA3 | 0.305 | 0.001 | 0.020 |
| Q9Y2Z4 | SYYM | 0.302 | 0.016 | 0.151 |
| Q9H3K6 | BOLA2 | 0.302 | 0.394 | 0.717 |
| P60891 | PRPS1 | 0.301 | 0.174 | 0.526 |
| Q14108 | SCRB2 | 0.300 | 0.211 | 0.576 |
| Q9NZL9 | MAT2B | 0.300 | 0.397 | 0.717 |
| Q96AQ6 | PBIP1 | 0.299 | 0.283 | 0.635 |
| Q9NVT9 | ARMC1 | 0.299 | 0.303 | 0.641 |
| Q8TB36 | GDAP1 | 0.298 | 0.008 | 0.093 |
| O43504 | LTOR5 | 0.296 | 0.716 | 0.928 |
| P26885 | FKBP2 | 0.295 | 0.004 | 0.066 |
| P35232 | PHB1 | 0.295 | 0.048 | 0.275 |
| Q92896 | GSLG1 | 0.295 | 1.000 | 1.000 |
| Q00169 | PIPNA | 0.295 | 0.066 | 0.314 |
| Q8N5M4 | TTC9C | 0.293 | 0.283 | 0.635 |
| P43304 | GPDM | 0.293 | 0.059 | 0.302 |
| P33240 | CSTF2 | 0.292 | 0.342 | 0.674 |
| Q9Y2S7 | PDIP2 | 0.291 | 0.263 | 0.627 |
| Q9NX76 | CKLF6 | 0.291 | 0.428 | 0.744 |
| P41250 | GARS | 0.289 | 0.053 | 0.288 |
| Q9BPX5 | ARP5L | 0.289 | 0.540 | 0.824 |
| Q9H6H4 | REEP4 | 0.288 | 0.371 | 0.696 |
| Q9HD34 | LYRM4 | 0.286 | 0.318 | 0.655 |
| Q9Y2H1 | ST38L | 0.285 | 0.073 | 0.330 |
| P56134 | ATPK | 0.283 | 0.211 | 0.576 |
| Q9NRW7 | VPS45 | 0.283 | 0.456 | 0.763 |

| | | | | |
|---|---|---|---|---|
| Q8WW22 | DNJA4 | 0.282 | 0.174 | 0.526 |
| O14579 | COPE | 0.282 | 0.347 | 0.674 |
| Q9BX68 | HINT2 | 0.281 | 0.073 | 0.330 |
| Q8IW45 | NNRD | 0.277 | 0.205 | 0.576 |
| P36507 | MP2K2 | 0.277 | 0.303 | 0.641 |
| Q7Z2Z2 | EFL1 | 0.276 | 0.195 | 0.557 |
| P00441 | SODC | 0.276 | 0.152 | 0.489 |
| Q99714 | HCD2 | 0.275 | 0.088 | 0.365 |
| Q9UM54-6 | MYO6 | 0.273 | 0.241 | 0.604 |
| P15170-2 | ERF3A | 0.270 | 0.616 | 0.869 |
| O60547 | GMDS | 0.269 | 0.097 | 0.378 |
| Q96PC5 | MIA2 | 0.269 | 0.167 | 0.511 |
| Q9Y450 | HBS1L | 0.268 | 0.403 | 0.724 |
| Q9HCN8 | SDF2L | 0.268 | 0.088 | 0.365 |
| Q15833 | STXB2 | 0.265 | 0.059 | 0.302 |
| Q9BPW8 | NIPS1 | 0.261 | 0.283 | 0.635 |
| P26641 | EF1G | 0.260 | 0.043 | 0.261 |
| Q86WA6 | BPHL | 0.260 | 0.205 | 0.576 |
| O00186 | STXB3 | 0.259 | 0.120 | 0.432 |
| P49321 | NASP | 0.259 | 0.152 | 0.489 |
| Q9NPE2 | NGRN | 0.258 | 0.225 | 0.582 |
| P04114 | APOB | 0.258 | 0.948 | 0.998 |
| Q96DZ1 | ERLEC | 0.257 | 0.073 | 0.330 |
| P04179 | SODM | 0.253 | 0.347 | 0.674 |
| P49419-2 | AL7A1 | 0.253 | 0.140 | 0.467 |
| P61966 | AP1S1 | 0.244 | 0.195 | 0.557 |
| P52565 | GDIR1 | 0.242 | 0.097 | 0.378 |
| O95831 | AIFM1 | 0.242 | 0.001 | 0.027 |
| Q96AB6 | NTAN1 | 0.242 | 0.283 | 0.635 |
| P01009 | A1AT | 0.242 | 0.444 | 0.747 |
| Q12907 | LMAN2 | 0.241 | 0.097 | 0.378 |
| Q9GZP4 | PITH1 | 0.241 | 0.325 | 0.655 |
| Q13438 | OS9 | 0.238 | 0.370 | 0.696 |
| P37268 | FDFT | 0.238 | 0.394 | 0.717 |
| Q8NBS9 | TXND5 | 0.236 | 0.066 | 0.314 |
| P61201 | CSN2 | 0.235 | 0.140 | 0.467 |
| Q5SRE7 | PHYD1 | 0.235 | 0.285 | 0.635 |
| P51570 | GALK1 | 0.233 | 0.303 | 0.641 |
| P53597 | SUCA | 0.232 | 0.030 | 0.215 |
| Q9BV79 | MECR | 0.232 | 0.444 | 0.747 |
| Q9BUR5 | MIC26 | 0.231 | 0.180 | 0.531 |
| P11766 | ADHX | 0.230 | 0.394 | 0.717 |
| Q9BU23 | LMF2 | 0.229 | 0.237 | 0.603 |
| O75489 | NDUS3 | 0.229 | 0.471 | 0.772 |
| Q8IYB8 | SUV3 | 0.225 | 0.195 | 0.557 |
| O60443 | GSDME | 0.225 | 0.370 | 0.696 |
| Q9Y697 | NFS1 | 0.225 | 0.347 | 0.674 |
| Q9NP79 | VTA1 | 0.225 | 0.867 | 0.978 |
| P35749 | MYH11 | 0.225 | 0.394 | 0.717 |
| Q9UII2 | ATIF1 | 0.224 | 0.117 | 0.424 |
| Q9BSD7 | NTPCR | 0.224 | 0.711 | 0.923 |

| | | | | |
|---|---|---|---|---|
| P53985 | MOT1 | 0.224 | 0.616 | 0.869 |
| Q8N1G4 | LRC47 | 0.224 | 0.283 | 0.635 |
| Q969V3 | NCLN | 0.224 | 0.679 | 0.907 |
| P53621 | COPA | 0.223 | 0.128 | 0.446 |
| Q96C36 | P5CR2 | 0.223 | 0.611 | 0.869 |
| P07237 | PDIA1 | 0.222 | 0.034 | 0.231 |
| Q9UJS0 | S2513 | 0.221 | 0.631 | 0.885 |
| A0A1B0GUC4 | MYCOS | 0.221 | 0.370 | 0.696 |
| P49789 | FHIT | 0.221 | 0.166 | 0.509 |
| P17174 | AATC | 0.221 | 0.073 | 0.330 |
| P20674 | COX5A | 0.221 | 0.499 | 0.790 |
| Q8WUD1 | RAB2B | 0.220 | 0.886 | 0.985 |
| Q14696 | MESD | 0.220 | 0.117 | 0.424 |
| Q4G0N4 | NAKD2 | 0.219 | 0.263 | 0.627 |
| Q7Z4G1 | COMD6 | 0.216 | 0.499 | 0.790 |
| P20936 | RASA1 | 0.216 | 0.721 | 0.934 |
| P30086 | PEBP1 | 0.216 | 0.128 | 0.446 |
| O00429 | DNM1L | 0.215 | 0.166 | 0.509 |
| P25685 | DNJB1 | 0.215 | 0.107 | 0.400 |
| P24752 | THIL | 0.212 | 0.030 | 0.215 |
| P14868 | SYDC | 0.211 | 0.347 | 0.674 |
| Q14643-2 | ITPR1 | 0.211 | 0.245 | 0.604 |
| P36915 | GNL1 | 0.210 | 0.568 | 0.848 |
| P04080 | CYTB | 0.209 | 0.227 | 0.582 |
| P04181 | OAT | 0.209 | 0.107 | 0.400 |
| O60826 | CCD22 | 0.209 | 0.802 | 0.955 |
| Q9BRA2 | TXD17 | 0.208 | 0.097 | 0.378 |
| O94788 | AL1A2 | 0.206 | 0.616 | 0.869 |
| Q9Y5X1 | SNX9 | 0.204 | 0.611 | 0.869 |
| Q92734 | TFG | 0.204 | 0.303 | 0.641 |
| Q6PL24 | TMED8 | 0.203 | 0.236 | 0.602 |
| Q02218 | ODO1 | 0.203 | 0.038 | 0.247 |
| Q9UBV2 | SE1L1 | 0.203 | 0.097 | 0.378 |
| P63000 | RAC1 | 0.203 | 0.325 | 0.655 |
| Q13217 | DNJC3 | 0.202 | 0.053 | 0.288 |
| P31947 | 1433S | 0.201 | 0.324 | 0.655 |
| Q12846 | STX4 | 0.198 | 0.507 | 0.798 |
| O60524 | NEMF | 0.197 | 0.781 | 0.955 |
| P00167 | CYB5 | 0.197 | 0.478 | 0.778 |
| P00387 | NB5R3 | 0.196 | 0.117 | 0.424 |
| P56192 | SYMC | 0.195 | 0.227 | 0.582 |
| Q15691 | MARE1 | 0.195 | 0.066 | 0.314 |
| P36955 | PEDF | 0.194 | 0.820 | 0.965 |
| Q9Y333 | LSM2 | 0.194 | 0.263 | 0.627 |
| Q969H8 | MYDGF | 0.194 | 0.303 | 0.641 |
| P12236 | ADT3 | 0.193 | 0.744 | 0.939 |
| P68036 | UB2L3 | 0.193 | 0.444 | 0.747 |
| O15347 | HMGB3 | 0.192 | 0.245 | 0.604 |
| P48506 | GSH1 | 0.191 | 0.499 | 0.790 |
| Q8TDQ7 | GNPI2 | 0.191 | 0.128 | 0.446 |
| P54098 | DPOG1 | 0.190 | 0.285 | 0.635 |

| | | | | |
|---|---|---|---|---|
| Q8IXB1 | DJC10 | 0.190 | 0.544 | 0.828 |
| Q9H9A6 | LRC40 | 0.189 | 0.140 | 0.467 |
| O95168 | NDUB4 | 0.189 | 0.667 | 0.907 |
| Q9H4A5 | GLP3L | 0.188 | 0.180 | 0.531 |
| O75964 | ATP5L | 0.187 | 0.419 | 0.735 |
| Q9NQC3 | RTN4 | 0.186 | 0.152 | 0.489 |
| Q9H1K1 | ISCU | 0.186 | 0.283 | 0.635 |
| Q9UI12 | VATH | 0.186 | 0.879 | 0.978 |
| P34932 | HSP74 | 0.184 | 0.059 | 0.302 |
| Q9GZT3 | SLIRP | 0.184 | 0.711 | 0.923 |
| P09496-2 | CLCA | 0.183 | 0.107 | 0.400 |
| Q9BWS9 | CHID1 | 0.183 | 0.423 | 0.737 |
| O15258 | RER1 | 0.181 | 0.403 | 0.724 |
| P00492 | HPRT | 0.181 | 0.227 | 0.582 |
| Q00610 | CLH1 | 0.181 | 0.059 | 0.302 |
| Q9UMX0 | UBQL1 | 0.180 | 0.581 | 0.857 |
| P02766 | TTHY | 0.180 | 0.698 | 0.923 |
| Q8IWB7 | WDFY1 | 0.179 | 0.227 | 0.582 |
| Q96GD0 | PLPP | 0.179 | 0.732 | 0.939 |
| O00764 | PDXK | 0.179 | 0.370 | 0.696 |
| P20073 | ANXA7 | 0.179 | 0.370 | 0.696 |
| P30405 | PPIF | 0.179 | 0.948 | 0.998 |
| P21796 | VDAC1 | 0.179 | 0.107 | 0.400 |
| P32119 | PRDX2 | 0.178 | 0.245 | 0.604 |
| Q9GZT8 | NIF3L | 0.177 | 0.611 | 0.869 |
| P61020 | RAB5B | 0.177 | 0.444 | 0.747 |
| Q3SXM5 | HSDL1 | 0.177 | 0.397 | 0.717 |
| P22033 | MUTA | 0.176 | 0.088 | 0.365 |
| Q5T124 | UBX11 | 0.175 | 0.443 | 0.747 |
| P49773 | HINT1 | 0.175 | 0.394 | 0.717 |
| Q15369 | ELOC | 0.174 | 0.324 | 0.655 |
| P26640 | SYVC | 0.173 | 0.909 | 0.985 |
| O14818 | PSA7 | 0.173 | 0.303 | 0.641 |
| Q9H9P8 | L2HDH | 0.172 | 0.611 | 0.869 |
| Q16531 | DDB1 | 0.172 | 0.038 | 0.247 |
| Q6GMV3 | PTRD1 | 0.172 | 0.211 | 0.576 |
| Q9UNZ2 | NSF1C | 0.172 | 0.048 | 0.275 |
| P15121 | ALDR | 0.170 | 0.347 | 0.674 |
| Q99624 | S38A3 | 0.170 | 0.302 | 0.641 |
| Q15435 | PP1R7 | 0.169 | 0.811 | 0.955 |
| P14927 | QCR7 | 0.166 | 0.394 | 0.717 |
| P42126 | ECI1 | 0.166 | 0.283 | 0.635 |
| O00483 | NDUA4 | 0.163 | 0.399 | 0.719 |
| Q9NPJ3 | ACO13 | 0.162 | 0.394 | 0.717 |
| P61960 | UFM1 | 0.162 | 0.605 | 0.869 |
| Q7Z6Z7 | HUWE1 | 0.161 | 0.837 | 0.966 |
| P40925 | MDHC | 0.159 | 0.263 | 0.627 |
| Q92538 | GBF1 | 0.158 | 0.799 | 0.955 |
| P00568 | KAD1 | 0.158 | 0.195 | 0.557 |
| Q9UQ13 | SHOC2 | 0.157 | 0.471 | 0.772 |
| P43686 | PRS6B | 0.157 | 0.303 | 0.641 |

| | | | | |
|---|---|---|---|---|
| P37802 | TAGL2 | 0.156 | 0.211 | 0.576 |
| P05141 | ADT2 | 0.156 | 0.394 | 0.717 |
| Q99426 | TBCB | 0.156 | 0.303 | 0.641 |
| Q93034 | CUL5 | 0.156 | 0.117 | 0.424 |
| Q9P032 | NDUF4 | 0.155 | 0.531 | 0.814 |
| P30049 | ATPD | 0.155 | 0.419 | 0.735 |
| P11047 | LAMC1 | 0.154 | 0.227 | 0.582 |
| Q9Y4L1 | HYOU1 | 0.154 | 0.066 | 0.314 |
| O95674 | CDS2 | 0.153 | 0.983 | 1.000 |
| P60981 | DEST | 0.152 | 0.325 | 0.655 |
| P31689 | DNJA1 | 0.151 | 0.211 | 0.576 |
| P09960 | LKHA4 | 0.150 | 0.527 | 0.810 |
| P09669 | COX6C | 0.148 | 0.478 | 0.778 |
| Q9Y696 | CLIC4 | 0.147 | 0.444 | 0.747 |
| O75396 | SC22B | 0.147 | 0.419 | 0.735 |
| Q9NRG9 | AAAS | 0.147 | 0.811 | 0.955 |
| O00534 | VMA5A | 0.145 | 0.527 | 0.810 |
| P08754 | GNAI3 | 0.144 | 1.000 | 1.000 |
| P78406 | RAE1L | 0.143 | 0.586 | 0.857 |
| P23381 | SYWC | 0.143 | 0.211 | 0.576 |
| Q9UI09 | NDUAC | 0.142 | 0.581 | 0.857 |
| Q9NX63 | MIC19 | 0.141 | 0.527 | 0.810 |
| P55011 | S12A2 | 0.141 | 0.397 | 0.717 |
| Q8NBX0 | SCPDL | 0.141 | 0.499 | 0.790 |
| Q9NZ45 | CISD1 | 0.141 | 0.679 | 0.907 |
| P28074 | PSB5 | 0.140 | 0.283 | 0.635 |
| Q9Y5M8 | SRPRB | 0.140 | 0.527 | 0.810 |
| P18859 | ATP5J | 0.139 | 0.347 | 0.674 |
| O95757 | HS74L | 0.139 | 0.263 | 0.627 |
| O95573 | ACSL3 | 0.139 | 0.486 | 0.786 |
| P21291 | CSRP1 | 0.138 | 0.471 | 0.772 |
| P07099 | HYEP | 0.138 | 0.444 | 0.747 |
| P60174 | TPIS | 0.137 | 0.283 | 0.635 |
| Q5T5Y3 | CAMP1 | 0.137 | 0.560 | 0.840 |
| P27797 | CALR | 0.135 | 0.166 | 0.509 |
| O95571 | ETHE1 | 0.135 | 0.527 | 0.810 |
| P00505 | AATM | 0.134 | 0.166 | 0.509 |
| Q13630 | FCL | 0.133 | 0.647 | 0.886 |
| Q96AB3 | ISOC2 | 0.133 | 0.430 | 0.746 |
| A6NDG6 | PGP | 0.133 | 1.000 | 1.000 |
| P04844-2 | RPN2 | 0.133 | 0.811 | 0.955 |
| Q9UL25 | RAB21 | 0.132 | 0.647 | 0.886 |
| P35580 | MYH10 | 0.131 | 0.245 | 0.604 |
| Q8N5K1 | CISD2 | 0.131 | 0.790 | 0.955 |
| Q08722 | CD47 | 0.131 | 0.631 | 0.885 |
| P43034 | LIS1 | 0.130 | 0.325 | 0.655 |
| P35222 | CTNB1 | 0.130 | 0.471 | 0.772 |
| Q5JYT7 | K1755 | 0.130 | 0.616 | 0.869 |
| P60033 | CD81 | 0.130 | 0.732 | 0.939 |
| P10809 | CH60 | 0.130 | 0.073 | 0.330 |
| P09936 | UCHL1 | 0.130 | 0.777 | 0.952 |

| | | | | |
|---|---|---|---|---|
| P28070 | PSB4 | 0.130 | 0.648 | 0.886 |
| O75531 | BAF | 0.129 | 0.913 | 0.985 |
| Q99816 | TS101 | 0.129 | 0.263 | 0.627 |
| P22830 | HEMH | 0.128 | 1.000 | 1.000 |
| Q8N2K0 | ABD12 | 0.128 | 0.913 | 0.985 |
| P49368 | TCPG | 0.128 | 0.097 | 0.378 |
| Q9NYU2 | UGGG1 | 0.126 | 0.283 | 0.635 |
| P08559 | ODPA | 0.126 | 0.073 | 0.330 |
| P00367 | DHE3 | 0.126 | 0.711 | 0.923 |
| Q9HDC9 | APMAP | 0.125 | 0.444 | 0.747 |
| P35606 | COPB2 | 0.125 | 0.043 | 0.261 |
| P07954 | FUMH | 0.124 | 0.263 | 0.627 |
| Q96HY6 | DDRGK | 0.123 | 0.838 | 0.966 |
| O60664 | PLIN3 | 0.122 | 0.845 | 0.966 |
| Q99627 | CSN8 | 0.122 | 0.303 | 0.641 |
| O43776 | SYNC | 0.121 | 0.370 | 0.696 |
| P13861 | KAP2 | 0.121 | 0.419 | 0.735 |
| O14949 | QCR8 | 0.121 | 1.000 | 1.000 |
| P13987 | CD59 | 0.120 | 0.777 | 0.952 |
| Q9BTV5 | FSD1 | 0.120 | 0.959 | 1.000 |
| P11233 | RALA | 0.120 | 0.499 | 0.790 |
| P82909 | KGD4 | 0.119 | 0.616 | 0.869 |
| Q8N4H5 | TOM5 | 0.118 | 0.755 | 0.949 |
| O15116 | LSM1 | 0.117 | 0.647 | 0.886 |
| P23588 | IF4B | 0.117 | 0.732 | 0.939 |
| Q12931 | TRAP1 | 0.116 | 0.166 | 0.509 |
| Q8N0X4 | CLYBL | 0.115 | 0.913 | 0.985 |
| Q9ULR5 | PAI2B | 0.114 | 0.732 | 0.939 |
| Q15631 | TSN | 0.114 | 0.370 | 0.696 |
| P49189 | AL9A1 | 0.114 | 0.166 | 0.509 |
| Q9NTX5 | ECHD1 | 0.113 | 0.586 | 0.857 |
| P54709 | AT1B3 | 0.112 | 0.904 | 0.985 |
| P00390 | GSHR | 0.112 | 0.394 | 0.717 |
| Q8N183 | NDUF2 | 0.112 | 0.647 | 0.886 |
| P13667 | PDIA4 | 0.112 | 0.152 | 0.489 |
| P39656 | OST48 | 0.112 | 0.744 | 0.939 |
| P09622 | DLDH | 0.111 | 0.245 | 0.604 |
| P42765 | THIM | 0.110 | 0.227 | 0.582 |
| Q6ZMN8 | CCNI2 | 0.110 | 0.811 | 0.955 |
| Q15785 | TOM34 | 0.110 | 0.679 | 0.907 |
| P01024 | CO3 | 0.109 | 0.913 | 0.985 |
| O75083 | WDR1 | 0.109 | 0.419 | 0.735 |
| Q6ZMR3 | LDH6A | 0.107 | 0.616 | 0.869 |
| O95822 | DCMC | 0.106 | 0.444 | 0.747 |
| Q9UJZ1 | STML2 | 0.106 | 0.347 | 0.674 |
| Q9H061 | T126A | 0.106 | 0.913 | 0.985 |
| Q96FJ2 | DYL2 | 0.106 | 0.556 | 0.835 |
| P35270 | SPRE | 0.106 | 0.744 | 0.939 |
| P06576 | ATPB | 0.105 | 0.180 | 0.531 |
| P31939 | PUR9 | 0.104 | 0.499 | 0.790 |
| P16435 | NCPR | 0.103 | 0.527 | 0.810 |

| | | | | |
|---|---|---|---|---|
| P33176 | KINH | 0.103 | 0.777 | 0.952 |
| P31040 | SDHA | 0.103 | 0.527 | 0.810 |
| Q15907 | RB11B | 0.102 | 0.943 | 0.998 |
| Q9BZE9 | ASPC1 | 0.102 | 0.744 | 0.939 |
| P12268 | IMDH2 | 0.102 | 0.263 | 0.627 |
| P27348 | 1433T | 0.101 | 0.616 | 0.869 |
| Q86SX6 | GLRX5 | 0.101 | 0.777 | 0.952 |
| Q7L5N1 | CSN6 | 0.101 | 0.744 | 0.939 |
| O94919 | ENDD1 | 0.101 | 0.732 | 0.939 |
| P61221 | ABCE1 | 0.101 | 0.802 | 0.955 |
| P09497-2 | CLCB | 0.101 | 0.647 | 0.886 |
| Q01469 | FABP5 | 0.098 | 0.982 | 1.000 |
| Q9H2U2 | IPYR2 | 0.097 | 0.537 | 0.821 |
| P01178 | NEU1 | 0.097 | 0.983 | 1.000 |
| P31930 | QCR1 | 0.096 | 0.983 | 1.000 |
| O00244 | ATOX1 | 0.096 | 0.845 | 0.966 |
| Q9NR45 | SIAS | 0.096 | 1.000 | 1.000 |
| P10515 | ODP2 | 0.095 | 0.325 | 0.655 |
| P36776 | LONM | 0.095 | 0.302 | 0.641 |
| Q9Y281 | COF2 | 0.095 | 0.811 | 0.955 |
| Q96DV4 | RM38 | 0.094 | 0.677 | 0.907 |
| Q9BV73 | CP250 | 0.094 | 0.777 | 0.952 |
| P35080 | PROF2 | 0.094 | 0.845 | 0.966 |
| O00203 | AP3B1 | 0.092 | 0.732 | 0.939 |
| Q13310 | PABP4 | 0.089 | 0.879 | 0.978 |
| Q15257-2 | PTPA | 0.088 | 0.263 | 0.627 |
| Q8N6M9 | ZFN2A | 0.088 | 0.802 | 0.955 |
| O60220 | TIM8A | 0.088 | 0.325 | 0.655 |
| P62942 | FKB1A | 0.088 | 0.195 | 0.557 |
| Q9NVI7-2 | ATD3A | 0.086 | 0.283 | 0.635 |
| P53992 | SC24C | 0.086 | 0.744 | 0.939 |
| P16885 | PLCG2 | 0.085 | 0.913 | 0.985 |
| P55209 | NP1L1 | 0.085 | 0.679 | 0.907 |
| P23284 | PPIB | 0.083 | 0.419 | 0.735 |
| Q14651 | PLSI | 0.080 | 0.811 | 0.955 |
| P35244 | RFA3 | 0.078 | 0.679 | 0.907 |
| P14618 | KPYM | 0.076 | 0.647 | 0.886 |
| Q86W24 | NAL14 | 0.075 | 0.777 | 0.952 |
| P55884 | EIF3B | 0.075 | 0.595 | 0.869 |
| A0A1W2PR48 | TLE7 | 0.074 | 0.777 | 0.952 |
| P48047 | ATPO | 0.074 | 0.711 | 0.923 |
| Q92581-2 | SL9A6 | 0.074 | 0.322 | 0.655 |
| Q9NZZ3 | CHMP5 | 0.074 | 0.631 | 0.885 |
| O95202 | LETM1 | 0.074 | 0.568 | 0.848 |
| Q16881 | TRXR1 | 0.073 | 0.913 | 0.985 |
| O95292 | VAPB | 0.072 | 0.811 | 0.955 |
| Q9UNF0 | PACN2 | 0.071 | 1.000 | 1.000 |
| Q9Y5T4 | DJC15 | 0.071 | 1.000 | 1.000 |
| P00403 | COX2 | 0.071 | 0.811 | 0.955 |
| Q8NB78 | KDM1B | 0.070 | 0.556 | 0.835 |
| P29401 | TKT | 0.070 | 1.000 | 1.000 |

| | | | | |
|---|---|---|---|---|
| P68366 | TBA4A | 0.068 | 0.586 | 0.857 |
| P54819 | KAD2 | 0.068 | 1.000 | 1.000 |
| O43707 | ACTN4 | 0.067 | 0.227 | 0.582 |
| P34897 | GLYM | 0.066 | 0.527 | 0.810 |
| Q6NVY1 | HIBCH | 0.066 | 0.419 | 0.735 |
| Q16718 | NDUA5 | 0.066 | 0.537 | 0.821 |
| O94826 | TOM70 | 0.066 | 0.948 | 0.998 |
| P61019 | RAB2A | 0.065 | 0.777 | 0.952 |
| P11310 | ACADM | 0.065 | 0.948 | 0.998 |
| P38117 | ETFB | 0.064 | 0.647 | 0.886 |
| O75947 | ATP5H | 0.063 | 0.471 | 0.772 |
| Q9UL26 | RB22A | 0.063 | 0.811 | 0.955 |
| Q96TC7 | RMD3 | 0.062 | 0.283 | 0.635 |
| Q9Y371 | SHLB1 | 0.062 | 0.913 | 0.985 |
| Q13907 | IDI1 | 0.062 | 0.711 | 0.923 |
| Q96A49 | SYAP1 | 0.061 | 0.744 | 0.939 |
| Q16698 | DECR | 0.060 | 0.948 | 0.998 |
| P30044-2 | PRDX5 | 0.060 | 0.647 | 0.886 |
| Q13867 | BLMH | 0.060 | 0.711 | 0.923 |
| P62330 | ARF6 | 0.059 | 0.983 | 1.000 |
| Q00535 | CDK5 | 0.059 | 0.647 | 0.886 |
| P09110 | THIK | 0.059 | 0.845 | 0.966 |
| P31948-2 | STIP1 | 0.059 | 0.370 | 0.696 |
| Q8IVS2 | FABD | 0.058 | 0.809 | 0.955 |
| P15927-3 | RFA2 | 0.058 | 1.000 | 1.000 |
| Q13561 | DCTN2 | 0.057 | 0.879 | 0.978 |
| Q99798 | ACON | 0.057 | 0.647 | 0.886 |
| Q06830 | PRDX1 | 0.057 | 0.444 | 0.747 |
| P43897 | EFTS | 0.056 | 0.471 | 0.772 |
| O60763 | USO1 | 0.056 | 0.913 | 0.985 |
| P30533 | AMRP | 0.056 | 0.227 | 0.582 |
| Q6PIU2 | NCEH1 | 0.055 | 0.744 | 0.939 |
| Q9NXJ5 | PGPI | 0.054 | 0.897 | 0.985 |
| P42766 | RL35 | 0.054 | 0.919 | 0.989 |
| Q86YB8 | ERO1B | 0.054 | 0.616 | 0.869 |
| P00558 | PGK1 | 0.054 | 0.616 | 0.869 |
| P40926 | MDHM | 0.053 | 0.471 | 0.772 |
| P0DOX2 | IGA2 | 0.053 | 0.727 | 0.939 |
| L0R6Q1 | S35U4 | 0.053 | 0.744 | 0.939 |
| Q8WX94-2 | NALP7 | 0.051 | 0.983 | 1.000 |
| Q12904 | AIMP1 | 0.050 | 0.777 | 0.952 |
| Q9BXU1 | STK31 | 0.049 | 0.845 | 0.966 |
| P14314 | GLU2B | 0.049 | 0.845 | 0.966 |
| Q9H9B4 | SFXN1 | 0.049 | 0.913 | 0.985 |
| Q14240 | IF4A2 | 0.048 | 0.913 | 0.985 |
| P13804 | ETFA | 0.048 | 0.647 | 0.886 |
| P51159 | RB27A | 0.047 | 0.647 | 0.886 |
| Q8IYB1 | M21D2 | 0.046 | 0.879 | 0.978 |
| Q14764 | MVP | 0.046 | 0.616 | 0.869 |
| Q9BT78 | CSN4 | 0.046 | 0.711 | 0.923 |
| P14625 | ENPL | 0.045 | 0.444 | 0.747 |

| | | | | |
|---|---|---|---|---|
| P55036 | PSMD4 | 0.043 | 0.767 | 0.952 |
| P35914 | HMGCL | 0.043 | 0.744 | 0.939 |
| Q8NAT1 | PMGT2 | 0.043 | 0.802 | 0.955 |
| P48735 | IDHP | 0.042 | 0.948 | 0.998 |
| Q96BN8 | OTUL | 0.041 | 0.744 | 0.939 |
| Q96IU4 | ABHEB | 0.041 | 0.913 | 0.985 |
| P54652 | HSP72 | 0.038 | 0.983 | 1.000 |
| P04843 | RPN1 | 0.038 | 0.879 | 0.978 |
| P36543 | VATE1 | 0.037 | 0.777 | 0.952 |
| Q15042 | RB3GP | 0.036 | 0.940 | 0.998 |
| Q9BQP7 | MGME1 | 0.034 | 0.919 | 0.989 |
| Q99829 | CPNE1 | 0.034 | 0.983 | 1.000 |
| O43447 | PPIH | 0.034 | 0.664 | 0.906 |
| Q9BQ95 | ECSIT | 0.034 | 0.616 | 0.869 |
| Q96AG4 | LRC59 | 0.033 | 0.845 | 0.966 |
| Q9BRX8 | PXL2A | 0.032 | 0.811 | 0.955 |
| Q9Y230 | RUVB2 | 0.032 | 0.811 | 0.955 |
| P28288 | ABCD3 | 0.032 | 0.982 | 1.000 |
| Q9Y305 | ACOT9 | 0.032 | 0.913 | 0.985 |
| P48444 | COPD | 0.030 | 0.845 | 0.966 |
| P48637 | GSHB | 0.030 | 0.879 | 0.978 |
| P62937 | PPIA | 0.029 | 0.948 | 0.998 |
| Q6TGC4 | PADI6 | 0.029 | 0.647 | 0.886 |
| P21281 | VATB2 | 0.028 | 0.744 | 0.939 |
| Q6FI81 | CPIN1 | 0.028 | 0.982 | 1.000 |
| Q96CN7 | ISOC1 | 0.028 | 0.471 | 0.772 |
| Q99747 | SNAG | 0.028 | 0.744 | 0.939 |
| P49589-3 | SYCC | 0.027 | 0.711 | 0.923 |
| O76024 | WFS1 | 0.027 | 0.873 | 0.978 |
| P04075 | ALDOA | 0.027 | 0.948 | 0.998 |
| Q14257 | RCN2 | 0.027 | 0.556 | 0.835 |
| P06730 | IF4E | 0.026 | 0.948 | 0.998 |
| Q93008 | USP9X | 0.026 | 0.913 | 0.985 |
| P55072 | TERA | 0.026 | 0.811 | 0.955 |
| O95373 | IPO7 | 0.026 | 0.945 | 0.998 |
| Q14974 | IMB1 | 0.025 | 0.913 | 0.985 |
| Q99536 | VAT1 | 0.025 | 0.647 | 0.886 |
| P35080-2 | PROF2 | 0.025 | 0.811 | 0.955 |
| Q96AG3 | S2546 | 0.024 | 0.913 | 0.985 |
| Q9NWU5 | RM22 | 0.024 | 0.777 | 0.952 |
| Q562R1 | ACTBL | 0.024 | 0.983 | 1.000 |
| O00154 | BACH | 0.023 | 0.983 | 1.000 |
| Q9UKG1 | DP13A | 0.023 | 0.838 | 0.966 |
| P49411 | EFTU | 0.023 | 0.948 | 0.998 |
| Q01518 | CAP1 | 0.023 | 0.913 | 0.985 |
| P36404 | ARL2 | 0.023 | 0.711 | 0.923 |
| P45974 | UBP5 | 0.022 | 0.948 | 0.998 |
| P23526 | SAHH | 0.021 | 0.845 | 0.966 |
| P51553 | IDH3G | 0.021 | 0.845 | 0.966 |
| Q8TD30 | ALAT2 | 0.021 | 0.711 | 0.923 |
| Q9HC38 | GLOD4 | 0.020 | 0.777 | 0.952 |

| | | | | |
|---|---|---|---|---|
| P30566 | PUR8 | 0.019 | 0.983 | 1.000 |
| P10253 | LYAG | 0.018 | 0.711 | 0.923 |
| Q13177 | PAK2 | 0.018 | 0.527 | 0.810 |
| P50995 | ANX11 | 0.017 | 0.679 | 0.907 |
| Q08257 | QOR | 0.016 | 0.983 | 1.000 |
| P08195 | 4F2 | 0.016 | 0.811 | 0.955 |
| P40227 | TCPZ | 0.016 | 0.711 | 0.923 |
| Q15046 | SYK | 0.015 | 0.860 | 0.978 |
| Q99623 | PHB2 | 0.015 | 0.983 | 1.000 |
| P38646 | GRP75 | 0.015 | 1.000 | 1.000 |
| P49755 | TMEDA | 0.014 | 0.845 | 0.966 |
| P17152 | TMM11 | 0.014 | 0.983 | 1.000 |
| Q07960 | RHG01 | 0.014 | 0.948 | 0.998 |
| Q16891 | MIC60 | 0.014 | 0.527 | 0.810 |
| P30041 | PRDX6 | 0.013 | 0.845 | 0.966 |
| Q04760 | LGUL | 0.013 | 0.616 | 0.869 |
| P43487 | RANG | 0.013 | 0.811 | 0.955 |
| Q9NP81 | SYSM | 0.013 | 0.711 | 0.923 |
| Q9UHG3 | PCYOX | 0.012 | 0.983 | 1.000 |
| O75608 | LYPA1 | 0.012 | 0.777 | 0.952 |
| P24534 | EF1B | 0.011 | 0.616 | 0.869 |
| P60660 | MYL6 | 0.011 | 0.983 | 1.000 |
| Q9UGP8 | SEC63 | 0.009 | 0.802 | 0.955 |
| O95716 | RAB3D | 0.009 | 0.983 | 1.000 |
| P05089 | ARGI1 | 0.009 | 0.983 | 1.000 |
| Q9Y4W6 | AFG32 | 0.009 | 0.679 | 0.907 |
| P62310 | LSM3 | 0.008 | 0.811 | 0.955 |
| P49748 | ACADV | 0.006 | 0.777 | 0.952 |
| Q99497 | PARK7 | 0.006 | 0.845 | 0.966 |
| P20618 | PSB1 | 0.006 | 0.983 | 1.000 |
| Q9NXA8 | SIR5 | 0.005 | 0.711 | 0.923 |
| O60313-2 | OPA1 | 0.005 | 0.444 | 0.747 |
| Q16204 | CCDC6 | 0.005 | 0.913 | 0.985 |
| Q9Y2Q3 | GSTK1 | 0.004 | 0.347 | 0.674 |
| P51659 | DHB4 | 0.004 | 0.983 | 1.000 |
| P59045 | NAL11 | 0.003 | 0.811 | 0.955 |
| Q9HD33 | RM47 | 0.003 | 0.827 | 0.966 |
| P24539 | AT5F1 | 0.003 | 0.616 | 0.869 |
| Q7Z2K6 | ERMP1 | 0.002 | 0.982 | 1.000 |
| Q9UHK6 | AMACR | 0.002 | 0.913 | 0.985 |
| O00232 | PSD12 | 0.001 | 0.909 | 0.985 |
| Q96FW1 | OTUB1 | 0.000 | 0.679 | 0.907 |
| P36551 | HEM6 | 0.000 | 0.616 | 0.869 |
| P35998 | PRS7 | 0.000 | 0.744 | 0.939 |
| P50213 | IDH3A | -0.001 | 0.879 | 0.978 |
| Q96KP4 | CNDP2 | -0.001 | 0.811 | 0.955 |
| Q9UMS0 | NFU1 | -0.001 | 0.586 | 0.857 |
| Q14697 | GANAB | -0.001 | 0.948 | 0.998 |
| P59047 | NALP5 | -0.001 | 0.983 | 1.000 |
| O60701 | UGDH | -0.002 | 0.948 | 0.998 |
| P08238 | HS90B | -0.002 | 0.711 | 0.923 |

| | | | | |
|---|---|---|---|---|
| P30047 | GFRP | -0.002 | 0.983 | 1.000 |
| P41732 | TSN7 | -0.003 | 0.945 | 0.998 |
| P50990 | TCPQ | -0.003 | 1.000 | 1.000 |
| P11177 | ODPB | -0.004 | 0.471 | 0.772 |
| Q6P996 | PDXD1 | -0.005 | 0.879 | 0.978 |
| Q96J94 | PIWL1 | -0.005 | 0.666 | 0.907 |
| Q9BXS5 | AP1M1 | -0.005 | 0.983 | 1.000 |
| P12277 | KCRB | -0.005 | 0.983 | 1.000 |
| P52209 | 6PGD | -0.006 | 0.913 | 0.985 |
| P62861 | RS30 | -0.006 | 0.948 | 0.998 |
| P08237 | PFKAM | -0.007 | 0.980 | 1.000 |
| P30084 | ECHM | -0.007 | 0.777 | 0.952 |
| P25705 | ATPA | -0.007 | 1.000 | 1.000 |
| P56385 | ATP5I | -0.008 | 0.711 | 0.923 |
| O14578-4 | CTRO | -0.008 | 0.963 | 1.000 |
| P50991 | TCPD | -0.010 | 0.777 | 0.952 |
| P33316 | DUT | -0.012 | 0.499 | 0.790 |
| O43196 | MSH5 | -0.012 | 0.948 | 0.998 |
| P20020 | AT2B1 | -0.012 | 0.913 | 0.985 |
| Q13098 | CSN1 | -0.012 | 0.777 | 0.952 |
| O43488 | ARK72 | -0.012 | 1.000 | 1.000 |
| P15586 | GNS | -0.013 | 0.983 | 1.000 |
| P22061-2 | PIMT | -0.014 | 0.948 | 0.998 |
| P54578 | UBP14 | -0.014 | 0.811 | 0.955 |
| Q14166 | TTL12 | -0.014 | 0.586 | 0.857 |
| P11279 | LAMP1 | -0.015 | 0.586 | 0.857 |
| Q7Z4W1 | DCXR | -0.015 | 0.879 | 0.978 |
| Q9HCU5 | PREB | -0.015 | 0.679 | 0.907 |
| Q96CW1 | AP2M1 | -0.015 | 0.807 | 0.955 |
| P10644 | KAP0 | -0.015 | 0.983 | 1.000 |
| Q9H808 | TLE6 | -0.016 | 0.647 | 0.886 |
| P48556 | PSMD8 | -0.017 | 0.811 | 0.955 |
| Q13526 | PIN1 | -0.017 | 0.777 | 0.952 |
| Q02252 | MMSA | -0.018 | 0.983 | 1.000 |
| P15374 | UCHL3 | -0.019 | 0.845 | 0.966 |
| Q99436 | PSB7 | -0.019 | 0.873 | 0.978 |
| Q9NZ01 | TECR | -0.019 | 0.945 | 0.998 |
| P55084 | ECHB | -0.019 | 0.777 | 0.952 |
| Q9Y6N5 | SQOR | -0.020 | 0.879 | 0.978 |
| P45880 | VDAC2 | -0.020 | 0.983 | 1.000 |
| P61224 | RAP1B | -0.021 | 0.811 | 0.955 |
| P17987 | TCPA | -0.022 | 0.811 | 0.955 |
| P54727 | RD23B | -0.023 | 0.711 | 0.923 |
| P14618-2 | KPYM | -0.025 | 0.983 | 1.000 |
| P11413 | G6PD | -0.025 | 0.983 | 1.000 |
| P62857 | RS28 | -0.025 | 0.556 | 0.835 |
| P61077 | UB2D3 | -0.025 | 1.000 | 1.000 |
| O14561 | ACPM | -0.028 | 1.000 | 1.000 |
| P50053-2 | KHK | -0.028 | 0.879 | 0.978 |
| P11234 | RALB | -0.028 | 0.647 | 0.886 |
| P04406 | G3P | -0.028 | 0.527 | 0.810 |

| | | | | |
|---|---|---|---|---|
| P27824 | CALX | -0.029 | 0.845 | 0.966 |
| P36542 | ATPG | -0.029 | 0.983 | 1.000 |
| P19623 | SPEE | -0.029 | 1.000 | 1.000 |
| Q5T2N8 | ATD3C | -0.029 | 0.777 | 0.952 |
| P07737 | PROF1 | -0.029 | 0.913 | 0.985 |
| Q9UJ70-2 | NAGK | -0.030 | 0.909 | 0.985 |
| Q9BX40 | LS14B | -0.030 | 0.303 | 0.641 |
| Q9Y3B8 | ORN | -0.030 | 0.948 | 0.998 |
| Q9BVK6 | TMED9 | -0.030 | 0.845 | 0.966 |
| Q99598 | TSNAX | -0.031 | 0.556 | 0.835 |
| Q9H853 | TBA4B | -0.031 | 0.599 | 0.869 |
| O95671 | ASML | -0.032 | 0.556 | 0.835 |
| P32969 | RL9 | -0.032 | 0.913 | 0.985 |
| P61981 | 1433G | -0.032 | 0.679 | 0.907 |
| O60610 | DIAP1 | -0.033 | 0.879 | 0.978 |
| P10606 | COX5B | -0.033 | 0.777 | 0.952 |
| P21399 | ACOHC | -0.034 | 0.913 | 0.985 |
| Q06203 | PUR1 | -0.034 | 0.879 | 0.978 |
| Q92820 | GGH | -0.034 | 0.913 | 0.985 |
| Q9UN37 | VPS4A | -0.034 | 0.616 | 0.869 |
| Q9HA64 | KT3K | -0.035 | 0.679 | 0.907 |
| Q9H173 | SIL1 | -0.036 | 1.000 | 1.000 |
| O75874 | IDHC | -0.036 | 0.879 | 0.978 |
| Q9Y5L4 | TIM13 | -0.037 | 0.811 | 0.955 |
| Q9HAB8 | PPCS | -0.038 | 0.811 | 0.955 |
| P40939 | ECHA | -0.038 | 1.000 | 1.000 |
| Q13011 | ECH1 | -0.039 | 0.711 | 0.923 |
| Q9BRQ6 | MIC25 | -0.039 | 0.616 | 0.869 |
| P53007 | TXTP | -0.039 | 0.802 | 0.955 |
| Q6IAA8 | LTOR1 | -0.040 | 0.370 | 0.696 |
| P22314 | UBA1 | -0.041 | 0.744 | 0.939 |
| Q9GZT6 | CC90B | -0.041 | 0.631 | 0.885 |
| O96008 | TOM40 | -0.042 | 0.983 | 1.000 |
| P46782 | RS5 | -0.044 | 0.979 | 1.000 |
| P50395 | GDIB | -0.044 | 0.711 | 0.923 |
| P52788 | SPSY | -0.044 | 0.616 | 0.869 |
| O43752 | STX6 | -0.045 | 0.913 | 0.985 |
| P42704 | LPPRC | -0.045 | 0.679 | 0.907 |
| Q96EY1 | DNJA3 | -0.045 | 1.000 | 1.000 |
| Q9NSE4 | SYIM | -0.045 | 1.000 | 1.000 |
| O43396 | TXNL1 | -0.045 | 0.647 | 0.886 |
| O75832 | PSD10 | -0.047 | 0.444 | 0.747 |
| O43813 | LANC1 | -0.047 | 0.811 | 0.955 |
| O60218 | AK1BA | -0.047 | 0.853 | 0.974 |
| P27144 | KAD4 | -0.047 | 0.471 | 0.772 |
| P62191 | PRS4 | -0.048 | 0.711 | 0.923 |
| Q00325 | MPCP | -0.048 | 0.913 | 0.985 |
| Q9BTW9 | TBCD | -0.048 | 0.527 | 0.810 |
| O15212 | PFD6 | -0.050 | 0.948 | 0.998 |
| P27635 | RL10 | -0.050 | 0.683 | 0.910 |
| Q16401 | PSMD5 | -0.052 | 0.802 | 0.955 |

| | | | | |
|---|---|---|---|---|
| Q9H444 | CHM4B | -0.053 | 0.744 | 0.939 |
| Q13428-4 | TCOF | -0.053 | 0.845 | 0.966 |
| P51114 | FXR1 | -0.054 | 0.647 | 0.886 |
| P11172 | UMPS | -0.054 | 0.711 | 0.923 |
| Q9UBS3 | DNJB9 | -0.055 | 1.000 | 1.000 |
| Q53GQ0 | DHB12 | -0.055 | 0.837 | 0.966 |
| P38606 | VATA | -0.056 | 0.211 | 0.576 |
| Q9Y277 | VDAC3 | -0.057 | 0.499 | 0.790 |
| P54920 | SNAA | -0.057 | 0.679 | 0.907 |
| Q9BPU6 | DPYL5 | -0.059 | 0.586 | 0.857 |
| O75306 | NDUS2 | -0.059 | 0.827 | 0.966 |
| O14745 | NHRF1 | -0.060 | 0.471 | 0.772 |
| P48643 | TCPE | -0.060 | 0.471 | 0.772 |
| Q15165 | PON2 | -0.061 | 0.845 | 0.966 |
| Q9HAV7 | GRPE1 | -0.062 | 0.811 | 0.955 |
| P30048 | PRDX3 | -0.063 | 0.811 | 0.955 |
| Q13442 | HAP28 | -0.064 | 0.527 | 0.810 |
| Q04446 | GLGB | -0.064 | 0.370 | 0.696 |
| P68402 | PA1B2 | -0.065 | 1.000 | 1.000 |
| P25787 | PSA2 | -0.065 | 0.777 | 0.952 |
| Q9Y2Z0 | SGT1 | -0.066 | 0.370 | 0.696 |
| O95336 | 6PGL | -0.066 | 0.777 | 0.952 |
| P49588 | SYAC | -0.067 | 0.527 | 0.810 |
| P62328 | TYB4 | -0.068 | 0.948 | 0.998 |
| P78371 | TCPB | -0.068 | 0.419 | 0.735 |
| P41208 | CETN2 | -0.069 | 0.302 | 0.641 |
| Q8WUM4 | PDC6I | -0.071 | 0.744 | 0.939 |
| Q9UM54-2 | MYO6 | -0.071 | 0.283 | 0.635 |
| Q86W25 | NAL13 | -0.074 | 0.913 | 0.985 |
| O43681 | GET3 | -0.075 | 0.499 | 0.790 |
| Q9Y6G9 | DC1L1 | -0.075 | 0.744 | 0.939 |
| P55789 | ALR | -0.076 | 0.838 | 0.966 |
| P11498 | PYC | -0.076 | 0.879 | 0.978 |
| Q9Y6Y8 | S23IP | -0.077 | 0.963 | 1.000 |
| Q9NUJ1 | ABHDA | -0.078 | 0.586 | 0.857 |
| Q9GZZ9 | UBA5 | -0.078 | 0.732 | 0.939 |
| P20810 | ICAL | -0.079 | 0.742 | 0.939 |
| Q9P0B6 | CC167 | -0.080 | 0.945 | 0.998 |
| Q8NCW5 | NNRE | -0.080 | 0.471 | 0.772 |
| Q9P2R7 | SUCB1 | -0.081 | 0.948 | 0.998 |
| Q9H9Q2 | CSN7B | -0.081 | 0.599 | 0.869 |
| P31946 | 1433B | -0.082 | 0.556 | 0.835 |
| P04899 | GNAI2 | -0.082 | 0.517 | 0.810 |
| Q9BVG9 | PTSS2 | -0.083 | 0.873 | 0.978 |
| Q9Y3C8 | UFC1 | -0.085 | 0.811 | 0.955 |
| P13073 | COX41 | -0.085 | 0.419 | 0.735 |
| Q9HB71 | CYBP | -0.086 | 0.471 | 0.772 |
| Q9Y3F4 | STRAP | -0.088 | 0.614 | 0.869 |
| Q6P1N9 | TATD1 | -0.089 | 0.263 | 0.627 |
| P43307 | SSRA | -0.089 | 0.394 | 0.717 |
| P13693 | TCTP | -0.089 | 0.811 | 0.955 |

| | | | | |
|---|---|---|---|---|
| P56211 | ARP19 | -0.089 | 0.614 | 0.869 |
| O60749 | SNX2 | -0.090 | 0.777 | 0.952 |
| Q9UHV9 | PFD2 | -0.091 | 0.879 | 0.978 |
| P46379 | BAG6 | -0.094 | 0.983 | 1.000 |
| P06744 | G6PI | -0.094 | 0.325 | 0.655 |
| P39019 | RS19 | -0.094 | 0.471 | 0.772 |
| O00592 | PODXL | -0.094 | 0.913 | 0.985 |
| P61018 | RAB4B | -0.095 | 0.879 | 0.978 |
| P29692 | EF1D | -0.095 | 0.527 | 0.810 |
| Q9Y2V2 | CHSP1 | -0.097 | 0.647 | 0.886 |
| P10599 | THIO | -0.097 | 0.913 | 0.985 |
| O43491 | E41L2 | -0.098 | 0.616 | 0.869 |
| Q15286 | RAB35 | -0.098 | 0.474 | 0.776 |
| P33121 | ACSL1 | -0.098 | 0.507 | 0.798 |
| O14964 | HGS | -0.098 | 0.610 | 0.869 |
| P13798 | ACPH | -0.099 | 0.263 | 0.627 |
| Q8WUY1 | THEM6 | -0.099 | 0.980 | 1.000 |
| P17858 | PFKAL | -0.099 | 0.263 | 0.627 |
| P28072 | PSB6 | -0.100 | 0.845 | 0.966 |
| Q15398-3 | DLGP5 | -0.104 | 0.166 | 0.509 |
| Q99460 | PSMD1 | -0.105 | 0.777 | 0.952 |
| Q8WWV3 | RT4I1 | -0.105 | 0.586 | 0.857 |
| O60879 | DIAP2 | -0.106 | 0.948 | 0.998 |
| P51572 | BAP31 | -0.106 | 0.347 | 0.674 |
| Q8IZP0 | ABI1 | -0.108 | 0.959 | 1.000 |
| Q587J8 | KHDC3 | -0.109 | 0.347 | 0.674 |
| O75190-2 | DNJB6 | -0.109 | 0.860 | 0.978 |
| P78344 | IF4G2 | -0.109 | 0.610 | 0.869 |
| Q5VWJ9 | SNX30 | -0.109 | 0.444 | 0.747 |
| P30519 | HMOX2 | -0.109 | 0.845 | 0.966 |
| P60953 | CDC42 | -0.109 | 0.241 | 0.604 |
| O00571 | DDX3X | -0.110 | 0.444 | 0.747 |
| P30085 | KCY | -0.110 | 0.527 | 0.810 |
| P22392-2 | NDKB | -0.110 | 0.683 | 0.910 |
| Q16555 | DPYL2 | -0.111 | 0.444 | 0.747 |
| Q9NQT8 | KI13B | -0.113 | 0.303 | 0.641 |
| P63208 | SKP1 | -0.113 | 0.370 | 0.696 |
| P14406 | CX7A2 | -0.113 | 0.527 | 0.810 |
| Q9NR28 | DBLOH | -0.114 | 0.913 | 0.985 |
| P46459 | NSF | -0.114 | 0.325 | 0.655 |
| Q9NYQ8 | FAT2 | -0.114 | 0.753 | 0.948 |
| P23434 | GCSH | -0.115 | 0.397 | 0.717 |
| P11586 | C1TC | -0.115 | 0.263 | 0.627 |
| Q7RTR0 | NLRP9 | -0.117 | 0.499 | 0.790 |
| Q9NR46 | SHLB2 | -0.117 | 0.556 | 0.835 |
| O43242 | PSMD3 | -0.117 | 0.211 | 0.576 |
| Q9H0U4 | RAB1B | -0.118 | 0.537 | 0.821 |
| Q9Y224 | RTRAF | -0.118 | 0.586 | 0.857 |
| Q96B97 | SH3K1 | -0.118 | 0.444 | 0.747 |
| Q86VP6 | CAND1 | -0.119 | 0.394 | 0.717 |
| P53618 | COPB | -0.120 | 0.647 | 0.886 |

| | | | | |
|---|---|---|---|---|
| Q9UK22 | FBX2 | -0.120 | 0.471 | 0.772 |
| Q93050 | VPP1 | -0.121 | 0.867 | 0.978 |
| Q9UHN1 | DPOG2 | -0.122 | 0.512 | 0.805 |
| P80303 | NUCB2 | -0.122 | 0.982 | 1.000 |
| Q13510 | ASAH1 | -0.124 | 0.873 | 0.978 |
| P15311 | EZRI | -0.125 | 0.325 | 0.655 |
| P55145 | MANF | -0.125 | 0.586 | 0.857 |
| P06753-2 | TPM3 | -0.126 | 0.347 | 0.674 |
| P02751 | FINC | -0.126 | 0.347 | 0.674 |
| Q9Y4I1 | MYO5A | -0.128 | 0.792 | 0.955 |
| Q04837 | SSBP | -0.128 | 0.616 | 0.869 |
| Q16836 | HCDH | -0.128 | 0.811 | 0.955 |
| P25398 | RS12 | -0.130 | 0.303 | 0.641 |
| Q9NQI0 | DDX4 | -0.130 | 0.347 | 0.674 |
| A0AVT1 | UBA6 | -0.131 | 0.325 | 0.655 |
| Q14204 | DYHC1 | -0.134 | 0.347 | 0.674 |
| Q96CT7 | CC124 | -0.134 | 0.370 | 0.696 |
| Q15008 | PSMD6 | -0.135 | 0.556 | 0.835 |
| Q7KZF4 | SND1 | -0.135 | 0.647 | 0.886 |
| A6NGQ2 | OOEP | -0.136 | 0.419 | 0.735 |
| Q9UHR4 | BI2L1 | -0.136 | 0.347 | 0.674 |
| P26196 | DDX6 | -0.138 | 0.325 | 0.655 |
| P49593 | PPM1F | -0.138 | 0.444 | 0.747 |
| Q9NX02 | NALP2 | -0.138 | 0.370 | 0.696 |
| Q16543 | CDC37 | -0.138 | 0.263 | 0.627 |
| P59998 | ARPC4 | -0.139 | 0.711 | 0.923 |
| Q13616 | CUL1 | -0.139 | 0.811 | 0.955 |
| Q2TAA2 | IAH1 | -0.140 | 0.711 | 0.923 |
| Q13615 | MTMR3 | -0.140 | 0.128 | 0.446 |
| Q8N131 | PORIM | -0.141 | 0.767 | 0.952 |
| Q4J6C6 | PPCEL | -0.143 | 0.531 | 0.814 |
| O43405 | COCH | -0.143 | 0.471 | 0.772 |
| P51531 | SMCA2 | -0.143 | 0.664 | 0.906 |
| P62312 | LSM6 | -0.145 | 0.586 | 0.857 |
| P51809 | VAMP7 | -0.145 | 0.755 | 0.949 |
| Q9UKY7 | CDV3 | -0.145 | 0.679 | 0.907 |
| Q5JRX3 | PREP | -0.146 | 0.263 | 0.627 |
| Q9Y639 | NPTN | -0.146 | 0.879 | 0.978 |
| P27361 | MK03 | -0.147 | 0.277 | 0.635 |
| Q9UBQ7 | GRHPR | -0.148 | 0.303 | 0.641 |
| P52292 | IMA1 | -0.148 | 0.499 | 0.790 |
| Q15436 | SC23A | -0.149 | 0.679 | 0.907 |
| P08574 | CY1 | -0.150 | 0.478 | 0.778 |
| P27694 | RFA1 | -0.150 | 0.556 | 0.835 |
| P61970 | NTF2 | -0.152 | 0.180 | 0.531 |
| Q8NC56 | LEMD2 | -0.154 | 0.610 | 0.869 |
| P61923 | COPZ1 | -0.155 | 0.711 | 0.923 |
| Q9UK45 | LSM7 | -0.155 | 0.370 | 0.696 |
| Q15185 | TEBP | -0.157 | 0.873 | 0.978 |
| P61313 | RL15 | -0.157 | 0.419 | 0.735 |
| Q07020 | RL18 | -0.159 | 0.195 | 0.557 |

| | | | | |
|---|---|---|---|---|
| Q9P035 | HACD3 | -0.160 | 0.572 | 0.853 |
| O43852 | CALU | -0.161 | 0.845 | 0.966 |
| Q9NP72 | RAB18 | -0.162 | 0.347 | 0.674 |
| Q96GK7 | FAH2A | -0.163 | 0.303 | 0.641 |
| Q9GZZ1 | NAA50 | -0.163 | 0.478 | 0.778 |
| Q5TAX3 | TUT4 | -0.164 | 0.811 | 0.955 |
| P46779 | RL28 | -0.164 | 0.631 | 0.885 |
| O43399 | TPD54 | -0.165 | 0.616 | 0.869 |
| P31150 | GDIA | -0.165 | 0.227 | 0.582 |
| P52594 | AGFG1 | -0.167 | 0.384 | 0.715 |
| P84550 | SKOR1 | -0.168 | 0.548 | 0.833 |
| Q709F0 | ACD11 | -0.168 | 0.444 | 0.747 |
| Q9H330 | TM245 | -0.168 | 1.000 | 1.000 |
| Q04917 | 1433F | -0.168 | 0.211 | 0.576 |
| Q15102 | PA1B3 | -0.169 | 0.744 | 0.939 |
| Q9ULC6 | PADI1 | -0.169 | 0.370 | 0.696 |
| Q6PKG0 | LARP1 | -0.169 | 0.223 | 0.582 |
| Q10567 | AP1B1 | -0.170 | 0.347 | 0.674 |
| P05362 | ICAM1 | -0.173 | 0.450 | 0.754 |
| P20929-2 | NEBU | -0.174 | 0.647 | 0.886 |
| P54136 | SYRC | -0.174 | 0.245 | 0.604 |
| Q96G97 | BSCL2 | -0.175 | 0.631 | 0.885 |
| Q8TBX8 | PI42C | -0.175 | 0.499 | 0.790 |
| Q9UK41 | VPS28 | -0.175 | 0.324 | 0.655 |
| P21926 | CD9 | -0.177 | 0.419 | 0.735 |
| Q8IVD9 | NUDC3 | -0.177 | 0.303 | 0.641 |
| O00487 | PSDE | -0.178 | 0.152 | 0.489 |
| Q16854 | DGUOK | -0.178 | 0.486 | 0.786 |
| P01834 | IGKC | -0.178 | 0.450 | 0.754 |
| P30626 | SORCN | -0.178 | 0.845 | 0.966 |
| Q9H832 | UBE2Z | -0.178 | 0.347 | 0.674 |
| P55060 | XPO2 | -0.180 | 0.117 | 0.424 |
| Q8IZ83 | A16A1 | -0.180 | 0.776 | 0.952 |
| P28482 | MK01 | -0.180 | 0.370 | 0.696 |
| P61247 | RS3A | -0.181 | 0.303 | 0.641 |
| P34949 | MPI | -0.181 | 0.283 | 0.635 |
| P11142 | HSP7C | -0.182 | 0.080 | 0.341 |
| P26373 | RL13 | -0.182 | 0.300 | 0.641 |
| O15382 | BCAT2 | -0.182 | 0.066 | 0.314 |
| P28331 | NDUS1 | -0.184 | 0.586 | 0.857 |
| Q00688 | FKBP3 | -0.184 | 0.211 | 0.576 |
| E9PAV3 | NACAM | -0.185 | 0.419 | 0.735 |
| Q96A35 | RM24 | -0.186 | 0.647 | 0.886 |
| P25786 | PSA1 | -0.187 | 0.419 | 0.735 |
| P01871 | IGHM | -0.188 | 0.849 | 0.970 |
| P62851 | RS25 | -0.188 | 0.444 | 0.747 |
| Q9NP97 | DLRB1 | -0.188 | 0.325 | 0.655 |
| P17980 | PRS6A | -0.190 | 0.003 | 0.048 |
| O14578-3 | CTRO | -0.190 | 0.324 | 0.655 |
| P25788 | PSA3 | -0.191 | 0.507 | 0.798 |
| P36578 | RL4 | -0.191 | 0.097 | 0.378 |

| | | | | |
|---|---|---|---|---|
| O00231 | PSD11 | -0.192 | 0.027 | 0.202 |
| Q12933 | TRAF2 | -0.193 | 0.384 | 0.715 |
| Q5BJF2 | SGMR2 | -0.196 | 0.666 | 0.907 |
| O43169 | CYB5B | -0.197 | 0.419 | 0.735 |
| Q9Y266 | NUDC | -0.199 | 0.152 | 0.489 |
| Q9Y678 | COPG1 | -0.199 | 0.394 | 0.717 |
| O75390 | CISY | -0.200 | 0.059 | 0.302 |
| P07814 | SYEP | -0.200 | 0.507 | 0.798 |
| Q8N0W3 | FCSK | -0.200 | 0.303 | 0.641 |
| P07900 | HS90A | -0.200 | 0.166 | 0.509 |
| P62273 | RS29 | -0.201 | 0.277 | 0.635 |
| O60383 | GDF9 | -0.201 | 0.370 | 0.696 |
| P27708 | PYR1 | -0.201 | 0.419 | 0.735 |
| Q9UFN0 | NPS3A | -0.201 | 0.499 | 0.790 |
| Q04637 | IF4G1 | -0.201 | 0.394 | 0.717 |
| Q7L0Y3 | TM10C | -0.201 | 0.152 | 0.489 |
| O95299 | NDUAA | -0.202 | 0.878 | 0.978 |
| O43837 | IDH3B | -0.202 | 0.080 | 0.341 |
| P63104 | 1433Z | -0.202 | 0.073 | 0.330 |
| O14804 | TAAR5 | -0.203 | 0.285 | 0.635 |
| Q96T17-2 | MA7D2 | -0.204 | 0.586 | 0.857 |
| O96000 | NDUBA | -0.204 | 0.427 | 0.743 |
| Q15370 | ELOB | -0.205 | 0.527 | 0.810 |
| P49720 | PSB3 | -0.205 | 0.423 | 0.737 |
| O00330 | ODPX | -0.205 | 0.283 | 0.635 |
| Q14558 | KPRA | -0.208 | 0.548 | 0.833 |
| P18124 | RL7 | -0.208 | 0.223 | 0.582 |
| Q12849 | GRSF1 | -0.210 | 0.568 | 0.848 |
| P09417 | DHPR | -0.210 | 0.325 | 0.655 |
| O43615 | TIM44 | -0.212 | 0.283 | 0.635 |
| O60884 | DNJA2 | -0.213 | 0.211 | 0.576 |
| Q9NUQ9 | CYRIB | -0.213 | 0.647 | 0.886 |
| Q9NQR4 | NIT2 | -0.213 | 0.845 | 0.966 |
| Q9NY33 | DPP3 | -0.214 | 0.152 | 0.489 |
| Q99733 | NP1L4 | -0.214 | 0.048 | 0.275 |
| P06753 | TPM3 | -0.214 | 0.325 | 0.655 |
| Q6P1J6 | PLB1 | -0.215 | 0.679 | 0.907 |
| O00425 | IF2B3 | -0.215 | 0.140 | 0.467 |
| P50336 | PPOX | -0.216 | 0.423 | 0.737 |
| P55786 | PSA | -0.216 | 0.245 | 0.604 |
| P05387 | RLA2 | -0.217 | 0.303 | 0.641 |
| P35221 | CTNA1 | -0.219 | 0.047 | 0.275 |
| Q5T160 | SYRM | -0.220 | 0.283 | 0.635 |
| Q7RTY7 | OVCH1 | -0.220 | 0.486 | 0.786 |
| P35268 | RL22 | -0.221 | 0.683 | 0.910 |
| Q9BSQ5 | CCM2 | -0.221 | 0.195 | 0.557 |
| Q13618 | CUL3 | -0.222 | 0.117 | 0.424 |
| P14550 | AK1A1 | -0.223 | 0.303 | 0.641 |
| Q8IW52 | SLIK4 | -0.223 | 0.347 | 0.674 |
| Q9NRY5 | F1142 | -0.224 | 0.527 | 0.810 |
| P07203 | GPX1 | -0.224 | 0.180 | 0.531 |

| | | | | |
|---|---|---|---|---|
| Q13263 | TIF1B | -0.225 | 0.487 | 0.786 |
| Q9NUQ7 | UFSP2 | -0.226 | 0.879 | 0.978 |
| P19404 | NDUV2 | -0.226 | 0.370 | 0.696 |
| P53041 | PPP5 | -0.227 | 0.370 | 0.696 |
| P62820 | RAB1A | -0.227 | 0.080 | 0.341 |
| O43678 | NDUA2 | -0.227 | 0.170 | 0.520 |
| Q9UIC8 | LCMT1 | -0.227 | 0.245 | 0.604 |
| O75165 | DJC13 | -0.228 | 0.140 | 0.467 |
| Q99832 | TCPH | -0.228 | 0.012 | 0.123 |
| Q4VXU2 | PAP1L | -0.228 | 0.038 | 0.247 |
| Q9UG56 | PISD | -0.229 | 0.443 | 0.747 |
| Q9C0C9 | UBE2O | -0.229 | 0.152 | 0.489 |
| Q92499 | DDX1 | -0.229 | 0.066 | 0.314 |
| P62917 | RL8 | -0.230 | 0.043 | 0.261 |
| P49591 | SYSC | -0.230 | 0.018 | 0.161 |
| Q9Y4Z0 | LSM4 | -0.231 | 0.110 | 0.407 |
| O15145 | ARPC3 | -0.234 | 0.146 | 0.484 |
| Q02878 | RL6 | -0.234 | 0.399 | 0.719 |
| Q9H336 | CRLD1 | -0.235 | 0.277 | 0.635 |
| Q9NT62 | ATG3 | -0.235 | 0.043 | 0.261 |
| Q9H8M7 | MINY3 | -0.237 | 0.544 | 0.828 |
| Q8NC96 | NECP1 | -0.238 | 0.211 | 0.576 |
| Q5T4I8 | CF052 | -0.238 | 0.829 | 0.966 |
| P62333 | PRS10 | -0.238 | 0.038 | 0.247 |
| P22570 | ADRO | -0.239 | 0.097 | 0.378 |
| Q9NSD9 | SYFB | -0.239 | 0.059 | 0.302 |
| O95081 | AGFG2 | -0.242 | 0.423 | 0.737 |
| Q02790 | FKBP4 | -0.242 | 0.066 | 0.314 |
| Q5RI15 | COX20 | -0.242 | 0.919 | 0.989 |
| Q15717 | ELAV1 | -0.245 | 0.371 | 0.696 |
| P63241 | IF5A1 | -0.246 | 0.711 | 0.923 |
| P49441 | INPP | -0.246 | 0.283 | 0.635 |
| O43768 | ENSA | -0.248 | 0.837 | 0.966 |
| Q92530 | PSMF1 | -0.248 | 0.330 | 0.663 |
| Q92542 | NICA | -0.250 | 0.373 | 0.697 |
| P52907 | CAZA1 | -0.250 | 0.487 | 0.786 |
| P01857 | IGHG1 | -0.252 | 0.325 | 0.655 |
| O15143 | ARC1B | -0.254 | 0.303 | 0.641 |
| A6NMX2 | I4E1B | -0.257 | 0.152 | 0.489 |
| Q00765 | REEP5 | -0.260 | 0.174 | 0.526 |
| P51665 | PSMD7 | -0.260 | 0.097 | 0.378 |
| P16444 | DPEP1 | -0.260 | 0.245 | 0.604 |
| P62805 | H4 | -0.261 | 0.913 | 0.985 |
| Q9NRW1 | RAB6B | -0.261 | 0.128 | 0.446 |
| Q6UXV4 | MIC27 | -0.262 | 0.180 | 0.532 |
| Q96C10 | DHX58 | -0.264 | 0.302 | 0.641 |
| Q9UNS2 | CSN3 | -0.265 | 0.205 | 0.576 |
| P11055 | MYH3 | -0.265 | 0.507 | 0.798 |
| Q5JSH3 | WDR44 | -0.267 | 0.347 | 0.674 |
| Q13505 | MTX1 | -0.268 | 0.325 | 0.655 |
| Q3ZCQ8 | TIM50 | -0.270 | 0.428 | 0.744 |

| | | | | |
|---|---|---|---|---|
| O75208 | COQ9 | -0.270 | 0.225 | 0.582 |
| P21266 | GSTM3 | -0.271 | 0.097 | 0.378 |
| Q9UNM6 | PSD13 | -0.271 | 0.256 | 0.627 |
| P08865 | RSSA | -0.272 | 0.159 | 0.508 |
| O00442 | RTCA | -0.275 | 0.245 | 0.604 |
| Q6XQN6 | PNCB | -0.275 | 0.501 | 0.793 |
| P61086 | UBE2K | -0.277 | 0.698 | 0.923 |
| Q99417 | MYCBP | -0.279 | 0.034 | 0.231 |
| Q68EM7 | RHG17 | -0.279 | 0.245 | 0.604 |
| Q9P0L0 | VAPA | -0.279 | 0.128 | 0.446 |
| Q9Y6C9 | MTCH2 | -0.281 | 0.845 | 0.966 |
| Q8N8S7 | ENAH | -0.281 | 0.499 | 0.790 |
| Q96RP9 | EFGM | -0.285 | 0.140 | 0.467 |
| O95394 | AGM1 | -0.285 | 0.370 | 0.696 |
| O15126 | SCAM1 | -0.288 | 0.753 | 0.948 |
| Q99962 | SH3G2 | -0.288 | 0.324 | 0.655 |
| P61088 | UBE2N | -0.291 | 0.294 | 0.641 |
| P62258 | 1433E | -0.291 | 0.007 | 0.085 |
| P26440 | IVD | -0.295 | 0.180 | 0.531 |
| P14174 | MIF | -0.296 | 0.423 | 0.737 |
| Q709C8 | VP13C | -0.296 | 0.585 | 0.857 |
| P35241 | RADI | -0.297 | 0.080 | 0.341 |
| Q9UHD1 | CHRD1 | -0.300 | 0.066 | 0.314 |
| Q15365 | PCBP1 | -0.300 | 0.166 | 0.509 |
| A6ND01 | JUNO | -0.300 | 0.444 | 0.747 |
| P18031 | PTN1 | -0.301 | 0.174 | 0.526 |
| Q9Y5X3 | SNX5 | -0.302 | 0.397 | 0.717 |
| O75027 | ABCB7 | -0.302 | 0.347 | 0.674 |
| Q13153 | PAK1 | -0.303 | 0.256 | 0.627 |
| P49821 | NDUV1 | -0.303 | 0.089 | 0.366 |
| P46777 | RL5 | -0.305 | 0.217 | 0.582 |
| P08758 | ANXA5 | -0.311 | 0.347 | 0.674 |
| P61158 | ARP3 | -0.311 | 0.011 | 0.111 |
| Q14165 | MLEC | -0.313 | 0.074 | 0.332 |
| Q03426 | KIME | -0.314 | 0.117 | 0.424 |
| Q9Y265 | RUVB1 | -0.315 | 0.024 | 0.190 |
| P02647 | APOA1 | -0.316 | 0.245 | 0.604 |
| O75369 | FLNB | -0.317 | 0.021 | 0.176 |
| O60869 | EDF1 | -0.317 | 0.021 | 0.176 |
| P30153 | 2AAA | -0.317 | 0.024 | 0.190 |
| P09493-5 | TPM1 | -0.318 | 0.223 | 0.582 |
| P62424 | RL7A | -0.318 | 0.048 | 0.275 |
| P53396 | ACLY | -0.318 | 0.303 | 0.641 |
| P11216 | PYGB | -0.320 | 0.080 | 0.341 |
| P14324 | FPPS | -0.331 | 0.285 | 0.636 |
| O95197 | RTN3 | -0.331 | 0.117 | 0.424 |
| Q7Z3Z3 | PIWL3 | -0.331 | 0.397 | 0.717 |
| P39210 | MPV17 | -0.336 | 0.274 | 0.635 |
| P21695 | GPDA | -0.336 | 0.180 | 0.531 |
| Q9UNE7 | CHIP | -0.337 | 0.008 | 0.093 |
| O14980 | XPO1 | -0.339 | 0.263 | 0.627 |

| | | | | |
|---|---|---|---|---|
| Q96CN9 | GCC1 | -0.340 | 0.873 | 0.978 |
| P80723 | BASP1 | -0.340 | 0.223 | 0.582 |
| O15511 | ARPC5 | -0.342 | 0.325 | 0.655 |
| P16278 | BGAL | -0.343 | 0.073 | 0.330 |
| P09543 | CN37 | -0.344 | 0.260 | 0.627 |
| Q9Y2R5 | RT17 | -0.345 | 0.367 | 0.696 |
| Q92882 | OSTF1 | -0.347 | 0.134 | 0.460 |
| P30520 | PURA2 | -0.348 | 0.330 | 0.663 |
| Q92598 | HS105 | -0.348 | 0.066 | 0.314 |
| P50914 | RL14 | -0.355 | 0.080 | 0.341 |
| Q86UP2 | KTN1 | -0.356 | 0.038 | 0.247 |
| P63244 | RACK1 | -0.356 | 0.066 | 0.314 |
| P25789 | PSA4 | -0.357 | 0.097 | 0.378 |
| Q6HA08 | ASTL | -0.357 | 0.027 | 0.202 |
| P84098 | RL19 | -0.357 | 0.107 | 0.400 |
| P46776 | RL27A | -0.359 | 0.110 | 0.407 |
| Q6X9E4 | FBW12 | -0.360 | 0.444 | 0.747 |
| P35579 | MYH9 | -0.360 | 0.001 | 0.025 |
| P12081 | HARS1 | -0.361 | 0.053 | 0.288 |
| Q9Y512 | SAM50 | -0.361 | 0.048 | 0.275 |
| P27816 | MAP4 | -0.363 | 0.012 | 0.123 |
| P06493 | CDK1 | -0.365 | 0.110 | 0.407 |
| P23528 | COF1 | -0.365 | 0.180 | 0.531 |
| P49327 | FAS | -0.367 | 0.080 | 0.341 |
| P30050 | RL12 | -0.367 | 0.093 | 0.377 |
| Q14847 | LASP1 | -0.369 | 0.272 | 0.635 |
| P84085 | ARF5 | -0.369 | 0.283 | 0.635 |
| Q8TF65 | GIPC2 | -0.369 | 0.241 | 0.604 |
| Q15366 | PCBP2 | -0.369 | 0.211 | 0.576 |
| O14908 | GIPC1 | -0.372 | 0.166 | 0.509 |
| P62280 | RS11 | -0.378 | 0.082 | 0.344 |
| Q9NVA2 | SEP11 | -0.378 | 0.097 | 0.378 |
| Q9Y3I0 | RTCB | -0.378 | 0.059 | 0.302 |
| Q9Y303 | NAGA | -0.379 | 0.093 | 0.377 |
| P31153 | METK2 | -0.380 | 0.236 | 0.602 |
| P31942 | HNRH3 | -0.382 | 0.128 | 0.446 |
| P62495 | ERF1 | -0.383 | 0.212 | 0.578 |
| Q9H0E2 | TOLIP | -0.387 | 0.180 | 0.531 |
| O75822 | EIF3J | -0.387 | 0.080 | 0.341 |
| Q8NBJ4 | GOLM1 | -0.388 | 0.121 | 0.434 |
| P22234 | PUR6 | -0.392 | 0.211 | 0.576 |
| O00515 | LAD1 | -0.392 | 0.001 | 0.027 |
| P23396 | RS3 | -0.393 | 0.016 | 0.151 |
| Q66K74 | MAP1S | -0.396 | 0.009 | 0.101 |
| P18669 | PGAM1 | -0.399 | 0.205 | 0.576 |
| P61353 | RL27 | -0.399 | 0.034 | 0.231 |
| Q14914 | PTGR1 | -0.402 | 0.166 | 0.509 |
| P04264 | K2C1 | -0.403 | 0.397 | 0.717 |
| P60842 | IF4A1 | -0.403 | 0.016 | 0.151 |
| Q96D96 | HVCN1 | -0.406 | 0.180 | 0.531 |
| Q10471 | GALT2 | -0.407 | 0.241 | 0.604 |

| | | | | |
|---|---|---|---|---|
| Q13200 | PSMD2 | -0.407 | 0.001 | 0.029 |
| P49207 | RL34 | -0.409 | 0.118 | 0.426 |
| Q15121 | PEA15 | -0.409 | 0.217 | 0.582 |
| P47755 | CAZA2 | -0.414 | 0.080 | 0.341 |
| P05386 | RLA1 | -0.417 | 0.038 | 0.247 |
| Q13409 | DC1I2 | -0.419 | 0.107 | 0.400 |
| P62913 | RL11 | -0.419 | 0.013 | 0.128 |
| Q10713 | MPPA | -0.422 | 0.467 | 0.772 |
| C9JE40 | PATL2 | -0.424 | 0.008 | 0.093 |
| Q92667 | AKAP1 | -0.426 | 0.003 | 0.050 |
| O95972 | BMP15 | -0.436 | 0.073 | 0.330 |
| Q9UN86 | G3BP2 | -0.438 | 0.100 | 0.382 |
| Q12864 | CAD17 | -0.439 | 0.222 | 0.582 |
| P12956 | XRCC6 | -0.442 | 0.189 | 0.555 |
| Q16864 | VATF | -0.443 | 0.560 | 0.840 |
| O75251 | NDUS7 | -0.444 | 0.133 | 0.460 |
| Q15181 | IPYR | -0.444 | 0.011 | 0.111 |
| P62701 | RS4X | -0.445 | 0.059 | 0.302 |
| P49721 | PSB2 | -0.448 | 0.241 | 0.604 |
| O15144 | ARPC2 | -0.449 | 0.008 | 0.093 |
| P24666 | PPAC | -0.449 | 0.099 | 0.382 |
| Q14738 | 2A5D | -0.450 | 0.432 | 0.747 |
| Q9HC35 | EMAL4 | -0.450 | 0.023 | 0.187 |
| Q9UHD8 | SEPT9 | -0.451 | 0.008 | 0.095 |
| P36871 | PGM1 | -0.451 | 0.018 | 0.161 |
| P13489 | RINI | -0.452 | 0.263 | 0.627 |
| P08833 | IBP1 | -0.458 | 0.324 | 0.655 |
| Q9UKA9 | PTBP2 | -0.460 | 0.080 | 0.341 |
| P26639 | SYTC | -0.461 | 0.100 | 0.382 |
| P46778 | RL21 | -0.462 | 0.017 | 0.160 |
| P08133 | ANXA6 | -0.462 | 0.003 | 0.050 |
| Q96IX5 | ATPMK | -0.464 | 0.217 | 0.582 |
| P61160 | ARP2 | -0.465 | 0.003 | 0.050 |
| P39023 | RL3 | -0.465 | 0.300 | 0.641 |
| O95219 | SNX4 | -0.466 | 0.029 | 0.215 |
| Q16181 | SEPT7 | -0.467 | 0.037 | 0.247 |
| P62277 | RS13 | -0.469 | 0.005 | 0.073 |
| Q86VR2 | RETR3 | -0.470 | 0.397 | 0.717 |
| Q96AX1 | VP33A | -0.471 | 0.046 | 0.275 |
| Q9NQW7 | XPP1 | -0.472 | 0.080 | 0.341 |
| Q9NQC3-2 | RTN4 | -0.473 | 0.373 | 0.697 |
| Q9NX14 | NDUBB | -0.474 | 0.294 | 0.641 |
| O75534 | CSDE1 | -0.478 | 0.134 | 0.460 |
| Q92947 | GCDH | -0.479 | 0.097 | 0.378 |
| P15880 | RS2 | -0.483 | 0.018 | 0.161 |
| O75323 | NIPS2 | -0.485 | 0.212 | 0.578 |
| P05388 | RLA0 | -0.491 | 0.110 | 0.407 |
| Q5T6F2 | UBAP2 | -0.494 | 0.080 | 0.341 |
| Q9H2J4 | PDCL3 | -0.494 | 0.053 | 0.288 |
| P55010 | IF5 | -0.495 | 0.100 | 0.382 |
| P13639 | EF2 | -0.495 | 0.000 | 0.013 |

| | | | | |
|---|---|---|---|---|
| Q9NRA8 | 4ET | -0.498 | 0.000 | 0.015 |
| P06733 | ENOA | -0.500 | 0.001 | 0.027 |
| Q9P2E9 | RRBP1 | -0.501 | 0.160 | 0.508 |
| P47985 | UCRI | -0.503 | 0.679 | 0.907 |
| Q9UMQ6 | CAN11 | -0.503 | 0.222 | 0.582 |
| P62269 | RS18 | -0.503 | 0.002 | 0.033 |
| Q96KA5 | CLP1L | -0.504 | 0.225 | 0.582 |
| P62829 | RL23 | -0.506 | 0.134 | 0.460 |
| Q8N983 | RM43 | -0.507 | 0.285 | 0.635 |
| O60493 | SNX3 | -0.507 | 0.030 | 0.215 |
| Q96FV2 | SCRN2 | -0.507 | 0.047 | 0.275 |
| P55809 | SCOT1 | -0.509 | 0.066 | 0.316 |
| Q86X76 | NIT1 | -0.513 | 0.161 | 0.509 |
| P83731 | RL24 | -0.517 | 0.121 | 0.434 |
| P47914 | RL29 | -0.520 | 0.018 | 0.161 |
| Q15773 | MLF2 | -0.524 | 0.110 | 0.407 |
| Q9NS69 | TOM22 | -0.530 | 0.091 | 0.372 |
| Q9P2J5 | SYLC | -0.530 | 0.024 | 0.190 |
| P47756-2 | CAPZB | -0.536 | 0.000 | 0.013 |
| P62195 | PRS8 | -0.539 | 0.001 | 0.025 |
| Q99766 | ATP5S | -0.542 | 0.272 | 0.635 |
| Q99674 | CGRE1 | -0.545 | 0.034 | 0.231 |
| Q8WW33 | GTSF1 | -0.546 | 0.007 | 0.085 |
| P62888 | RL30 | -0.549 | 0.085 | 0.355 |
| Q6IMN6 | CAPR2 | -0.566 | 0.016 | 0.151 |
| Q86SE8 | NPM2 | -0.573 | 0.053 | 0.288 |
| P02792 | FRIL | -0.573 | 0.097 | 0.378 |
| Q3LXA3 | TKFC | -0.576 | 0.034 | 0.231 |
| Q96QK1 | VPS35 | -0.577 | 0.001 | 0.020 |
| O75436 | VP26A | -0.578 | 0.003 | 0.050 |
| P46940 | IQGA1 | -0.579 | 0.030 | 0.215 |
| Q92832 | NELL1 | -0.582 | 0.267 | 0.632 |
| P07093-2 | GDN | -0.585 | 0.159 | 0.508 |
| P62081 | RS7 | -0.595 | 0.006 | 0.077 |
| P61421 | VA0D1 | -0.598 | 0.058 | 0.302 |
| Q14677 | EPN4 | -0.608 | 0.053 | 0.288 |
| P59190 | RAB15 | -0.609 | 0.005 | 0.074 |
| Q8NA58 | PNDC1 | -0.613 | 0.019 | 0.162 |
| O60927 | PP1RB | -0.613 | 0.820 | 0.965 |
| Q15836 | VAMP3 | -0.615 | 0.043 | 0.261 |
| P18621 | RL17 | -0.617 | 0.007 | 0.086 |
| P62750 | RL23A | -0.618 | 0.004 | 0.066 |
| P40429 | RL13A | -0.631 | 0.004 | 0.066 |
| Q9H4M9 | EHD1 | -0.650 | 0.324 | 0.655 |
| P40616 | ARL1 | -0.652 | 0.267 | 0.632 |
| Q8IYS2 | K2013 | -0.656 | 0.051 | 0.288 |
| Q8IXM3 | RM41 | -0.657 | 0.008 | 0.095 |
| P61163 | ACTZ | -0.681 | 0.048 | 0.275 |
| P13645 | K1C10 | -0.683 | 0.195 | 0.557 |
| Q9UI30 | TR112 | -0.689 | 0.083 | 0.349 |
| Q8N9I0 | SYT2 | -0.690 | 0.068 | 0.326 |

| | | | | | |
|---|---|---|---|---|---|
| | Q99805 | TM9S2 | -0.694 | 0.007 | 0.085 |
| | P62753 | RS6 | -0.717 | 0.053 | 0.288 |
| | P62847 | RS24 | -0.722 | 0.007 | 0.086 |
| | Q8TAE8 | G45IP | -0.742 | 0.024 | 0.190 |
| | O14618 | CCS | -0.758 | 0.038 | 0.247 |
| | P62266 | RS23 | -0.758 | 0.004 | 0.066 |
| | P22626 | ROA2 | -0.798 | 0.020 | 0.171 |
| | Q12765 | SCRN1 | -0.806 | 0.064 | 0.314 |
| | Q9Y490 | TLN1 | -0.827 | 0.042 | 0.261 |
| | O43617 | TPPC3 | -0.847 | 0.154 | 0.494 |
| | Q6P1M0 | S27A4 | -0.851 | 0.025 | 0.196 |
| | Q14019 | COTL1 | -0.885 | 0.006 | 0.077 |
| | P02794 | FRIH | -0.901 | 0.027 | 0.202 |
| | P35527 | K1C9 | -0.927 | 0.222 | 0.582 |
| | P61586 | RHOA | -0.951 | 0.047 | 0.275 |
| | O75600 | KBL | -0.987 | 0.082 | 0.344 |
| | Q14103 | HNRPD | -1.062 | 0.013 | 0.128 |
| | O60925 | PFD1 | -1.148 | 0.010 | 0.105 |
| | O60888 | CUTA | -1.155 | 0.026 | 0.202 |
| | Q9UJC3 | HOOK1 | -1.256 | 0.048 | 0.275 |
| | P08670 | VIME | -1.626 | 0.059 | 0.303 |
| | P09455 | RET1 | -1.693 | 0.006 | 0.082 |
| Significantly abundant proteins in GV | P60900 | PSA6 | -0.603 | 0.001 | 0.029 |
| | P62249 | RS16 | -0.606 | 0.001 | 0.023 |
| | P42025 | ACTY | -0.626 | 0.003 | 0.048 |
| | P21333 | FLNA | -0.644 | 0.000 | 0.013 |
| | P60866 | RS20 | -0.646 | 0.002 | 0.042 |
| | P62241 | RS8 | -0.686 | 0.002 | 0.038 |
| | Q9UBQ0 | VPS29 | -0.691 | 0.001 | 0.027 |
| | Q9H910 | JUPI2 | -0.694 | 0.000 | 0.004 |
| | P62906 | RL10A | -0.725 | 0.001 | 0.020 |
| | P46781 | RS9 | -0.764 | 0.001 | 0.027 |
| | Q9Y2W6 | TDRKH | -0.770 | 0.001 | 0.027 |
| | Q14203 | DCTN1 | -0.788 | 0.000 | 0.019 |
| | P08708 | RS17 | -0.849 | 0.001 | 0.028 |
| | Q96M93 | ADAD1 | -0.899 | 0.001 | 0.029 |
| | Q8IZA3 | H18 | -0.915 | 0.000 | 0.002 |
| | O94760 | DDAH1 | -0.961 | 0.000 | 0.017 |
| | Q9Y2T7 | YBOX2 | -0.977 | 0.000 | 0.002 |
| | P51812 | KS6A3 | -1.015 | 0.003 | 0.050 |
| | P07437 | TBB5 | -1.107 | 0.003 | 0.050 |
| | P43121 | MUC18 | -1.143 | 0.000 | 0.020 |
| | Q96CW5 | GCP3 | -1.262 | 0.001 | 0.020 |
| | Q13185 | CBX3 | -1.264 | 0.001 | 0.020 |
| | P07339 | CATD | -1.296 | 0.000 | 0.011 |
| | Q16658 | FSCN1 | -1.508 | 0.000 | 0.002 |
| | P62263 | RS14 | -1.609 | 0.000 | 0.013 |
| | P07858 | CATB | -1.772 | 0.000 | 0.000 |

**Table S4. Identified changes in GO (Biological Processes, BP; Cellular Compartments, CC) and KEGG pathways. Young group: MII *vs.* GV oocytes.**

| | Terms | Overlap | Ratio | P-value | P.adj | Log2 (FC) | Proteins |
|---|---|---|---|---|---|---|---|
| BP | nuclear-transcribed mRNA catabolic process, nonsense-mediated decay (GO:0000184) | 62/113 | 0.5 | 1E-44 | 2E-41 | -0.389 | RL4; RL5; RL30; RL3; RL34; RLA1; RLA0; RL10A; RL8; RL9; RL6; RL7; RS4X; RS14; RL7A; RS17; RS16; RS19; 2AAA; RS18; RL35; RLA2; RS11; RS13; RS12; RS9; RL21; RS7; RS8; RL23; RS5; RL22; RS6; RL13A; RSSA; RS3A; RL24; RL27; ERF1; RL29; RL28; RL10; RL12; RL11; ERF3A; RL14; RS3; RL13; RL15; RS2; RL18; RL17; RL19; RL23A; RS25; RS28; RS29; RL27A; RS20; RS24; IF4G1; RS23 |
| | cytoplasmic translation (GO:0002181) | 61/93 | 0.7 | 1E-43 | 1E-40 | -0.389 | IF4A2; RL4; IF4A1; RL5; RL30; RL3; RL34; RLA1; RLA0; RL10A; RL8; RL9; RL6; RL7; RS4X; RL7A; RS14; RS17; RS16; RS19; RS18; RL35; RS11; RS13; RS12; RS9; RL21; RS7; RS8; RL23; RS5; RL22; RS6; RL13A; RS3A; RSSA; RL24; ERF1; RL27; RL29; RL28; RL10; RL12; RL11; ERF3A; RL14; RS3; RL13; RL15; RS2; RL18; RL17; RL19; RL23A; RS25; RS28; RS29; RL27A; RS20; RS24; RS23 |
| | SRP-dependent cotranslational protein targeting to membrane (GO:0006614) | 59/90 | 0.7 | 1E-43 | 1E-40 | -0.399 | RL4; RL5; RL30; RL3; RL34; RLA1; RLA0; RL10A; RL8; RL9; RL6; RL7; RS4X; RL7A; RS14; RS17; RS16; RS19; RS18; RL35; RLA2; RS11; RS13; RS12; SEC63; RS9; RL21; RS7; RS8; RL23; RS5; RL22; RS6; RL13A; RS3A; RSSA; RL24; RL27; RL29; RL28; RL10; RL12; RL11; RL14; RS3; RL13; RL15; RS2; RL18; RL17; RL19; RL23A; RS25; RS28; RS29; RL27A; RS20; RS24; RS23 |
| | cotranslational protein targeting to membrane (GO:0006613) | 61/94 | 0.6 | 2E-43 | 2E-40 | -0.388 | RL4; RL5; RL30; RL3; RL34; RLA1; RLA0; RL10A; RL8; RL9; RL6; RL7; RS4X; RL7A; RS14; RS17; RS16; RS19; RS18; RL35; RLA2; RS11; SIL1; RS13; RS12; SEC63; RS9; RL21; RS7; RS8; RL23; RS5; RL22; RS6; RL13A; RS3A; RSSA; RL24; RL27; RL29; RL28; RL10; RL12; RL11; RL14; RS3; RL13; SSRA; RL15; RS2; RL18; RL17; RL19; RL23A; RS25; RS28; RS29; RL27A; RS20; RS24; RS23 |
| | protein targeting to ER (GO:0045047) | 64/103 | 0.6 | 4E-39 | 2E-36 | -0.358 | RL4; RL5; RL30; RL3; RL34; RLA1; ZFN2A; RLA0; RL10A; RL8; RL9; RL6; RL7; RS4X; RS14; RL7A; RS17; RS16; RS19; RS18; RL35; RLA2; RS11; GET3; RS13; RS12; SEC63; RS9; RL21; RS7; RS8; RL23; RS5; RL22; RS6; RL13A; RSSA; RS3A; CHM4B; RL24; RL27; RL29; RL28; RL10; RL12; RL11; RL14; RS3; RL13; RL15; RS2; RL18; RL17; RL19; RL23A; RS25; RS28; SPCS2; RS29; RL27A; SRPRB; RS20; RS24; RS23 |
| | translation (GO:0006412) | 108/214 | 0.5 | 2E-31 | 8E-29 | -0.231 | RL4; RL5; RL30; RL3; RL34; RL8; ACPH; RL10A; RL9; SYYM; RL6; RL7; EF1B; RS14; RS17; RS16; RS19; SYNC; RS18; RL35; RS11; RS13; RS12; SYRM; RS9; SYRC; RL21; RS7; RS8; RL23; RS5; RL22; RS6; RSSA; EFTU; IPYR2; IPYR; SYLC; EF1A2; RL24; RL27; RL29; SYCC; RL28; SYTC; SYAC; ERF3A; SYVC; HARS1; EF2; RM24; AIMP1; RS25; RS28; CSN5; SYEP; RS29; RL27A; RS20; SYFA; RS24; SYFB; RS23; IF4G1; RT17; HBS1L; RLA1; RLA0; RM41; RS4X; TR112; RL7A; SYQ; RLA2; EFTS; PABP4; SYMC; RL13A; RM47; RS3A; RM43; SYDC; EF1G; SYSC; SYSM; EF1D; ERF1; MCA3; RL10; SYK; KGD4; RL12; RL11; RRBP1; RL14; RS3; RL13; SYIM; IF2B3; RL15; RS2; RL18; RL17; RL19; RL23A; SYWC; GARS; ABCE1 |
| | maintenance of DNA methylation (GO:0010216) | 03/06 | 0.5 | 2E-08 | 4E-06 | 1.141 | DNMT1; UHRF2; UHRF1 |
| | P-body assembly (GO:0033962) | 06/12 | 0.5 | 8E-06 | 5E-04 | -0.238 | DDX6; DYHC1; 4ET; LSM4; LSM3; PATL2 |
| | regulation of non-motile cilium assembly (GO:1902855) | 03/06 | 0.5 | 4E-05 | 2E-03 | -0.426 | MAP4; SEPT9; SEPT7 |
| | NIK/NF-kappaB signaling (GO:0038061) | 38/74 | 0.5 | 2E-04 | 5E-03 | -0.111 | PSD10; PSD12; PSD11; PSDE; PSD13; CUL1; PSA7; PSMD8; PSB6; PSB7; PSMD6; PSB4; PSMD7; PSB5; PSMD4; PSB2; PSMD5; PSB3; PSMD2; PSMD3; PSB1; PSMD1; PSMF1; SKP1; TRAF2; PSA5; PSA6; PSA3; PRS8; PSA4; PRS10; PSA1; PRS6A; PSA2; PRS6B; PRS4; PRS7; CSN8 |
| | regulation of cellular amine metabolic process (GO:0033238) | 34/51 | 0.7 | 2E-04 | 5E-03 | -0.114 | PSD10; PSD12; PSD11; PSDE; PSD13; PSA7; PSMD8; PSB6; PSB7; PSMD6; PSB4; PSMD7; PSB5; PSMD4; PSB2; PSMD5; PSB3; PSMD2; PSMD3; PSB1; PSMD1; PSMF1; PSA5; PSA6; PSA3; PRS8; PSA4; PRS10; PSA1; PRS6A; PSA2; PRS6B; PRS4; PRS7 |
| | regulation of cellular amino acid metabolic process (GO:0006521) | 34/54 | 0.6 | 2E-04 | 5E-03 | -0.114 | PSD10; PSD12; PSD11; PSDE; PSD13; PSA7; PSMD8; PSB6; PSB7; PSMD6; PSB4; PSMD7; PSB5; PSMD4; PSB2; PSMD5; PSB3; PSMD2; PSMD3; PSB1; PSMD1; PSMF1; PSA5; PSA6; PSA3; PRS8; PSA4; PRS10; PSA1; PRS6A; PSA2; PRS6B; PRS4; PRS7 |
| | pre-replicative complex assembly (GO:0036388) | 34/64 | 0.5 | 2E-04 | 5E-03 | -0.114 | PSD10; PSD12; PSD11; PSDE; PSD13; PSA7; PSMD8; PSB6; PSB7; PSMD6; PSB4; PSMD7; PSB5; PSMD4; PSB2; PSMD5; PSB3; PSMD2; PSMD3; PSB1; PSMD1; PSMF1; PSA5; PSA6; PSA3; PRS8; PSA4; PRS10; PSA1; PRS6A; PSA2; PRS6B; PRS4; PRS7 |
| | regulation of cellular ketone metabolic process (GO:0010565) | 35/64 | 0.5 | 3E-04 | 7E-03 | -0.111 | PSD10; PSD12; PSD11; PSDE; PSD13; PSA7; PSMD8; PSB6; PSB7; PSMD6; PSB4; PSMD7; PSB5; PSMD4; PSB2; PSMD5; PSB3; PSMD2; PSMD3; PSB1; PSMD1; PSMF1; SIR5; PSA5; PSA6; PSA3; PRS8; PSA4; PRS10; PSA1; PRS6A; PSA2; PRS6B; PRS4; PRS7 |
| | Arp2/3 complex-mediated actin nucleation (GO:0034314) | 08/15 | 0.5 | 4E-04 | 8E-03 | -0.244 | ARP3; ARP2; ARPC2; ARPC3; ARC1B; ARP5L; ARPC4; ARPC5 |
| | negative regulation of cellular response to hypoxia (GO:1900038) | 03/06 | 0.5 | 7E-04 | 1E-02 | -0.436 | DDAH1; HYOU1; ENOA |
| | lysine catabolic process (GO:0006554) | 06/12 | 0.5 | 2E-03 | 3E-02 | 0.160 | GCDH; ODO1; ODO2; CRYM; AL7A1; DLDH |
| | lysine metabolic process (GO:0006553) | 06/12 | 0.5 | 2E-03 | 3E-02 | 0.160 | GCDH; ODO1; ODO2; CRYM; AL7A1; DLDH |
| | 2-oxoglutarate metabolic process (GO:0006103) | 09/214 | 0.6 | 3E-03 | 4E-02 | 0.139 | AATC; KGD4; ALAT2; IDHC; AATM; IDHP; ODO1; ODO2; L2HDH |
| | negative regulation of G2/M transition of mitotic cell cycle (GO:0010972) | 38/66 | 0.6 | 3E-03 | 4E-02 | -0.076 | PSD10; PSD12; PSD11; PSDE; PSD13; VPS4A; CUL1; PSA7; AURKA; PSMD8; PSB6; PSB7; PSMD6; PSB4; PSMD7; PSB5; PSMD4; PSB2; PSMD5; PSB3; PSMD2; PSMD3; PSB1; PSMD1; PSMF1; SKP1; PSA5; PSA6; PSA3; PRS8; PSA4; PRS10; PSA1; PRS6A; PSA2; PRS6B; PRS4; PRS7 |
| | positive regulation of RNA polymerase II transcription preinitiation complex assembly (GO:0045899) | 05/07 | 0.7 | 4E-03 | 4E-02 | -0.162 | PRS8; PRS10; PRS6A; PRS6B; PRS7 |
| | negative regulation of cell cycle G2/M phase transition (GO:1902750) | 37/57 | 0.6 | 4E-03 | 4E-02 | -0.077 | PSMD10; PSMD12; PSMD11; PSMD14; PSMD13; CUL1; PSMA7; AURKA; PSMD8; PSMB6; PSMB7; PSMD6; PSMB4; PSMD7; PSMB5; PSMD4; PSMB2; PSMD5; PSMB3; PSMD2; PSMD3; PSMB1; PSMD1; PSMF1; SKP1; PSMA5; PSMA6; PSMA3; PSMC5; PSMA4; PSMC6; PSMA1; PSMC3; PSMA2; PSMC4; PSMC1; PSMC2 |
| | regulation of RNA polymerase II transcription preinitiation complex assembly (GO:0045898) | 05/10 | 0.5 | 4E-03 | 4E-02 | -0.162 | PRS8; PRS10; PRS6A; PRS6B; PRS7 |
| | regulation of extracellular exosome assembly (GO:1903551) | 03/06 | 0.5 | 4E-03 | 4E-02 | 0.374 | SDCB1; PDC6I; TS101 |
| | antigen processing and presentation of exogenous peptide antigen via MHC class I, TAP-dependent (GO:0002479) | 39/73 | 0.5 | 5E-03 | 5E-02 | -0.083 | PSD10; PSD12; PSD11; ITB5; PSDE; PSD13; PSA7; PSMD8; PSB6; PSB7; PSMD6; PSB4; PSMD7; PSB5; PSMD4; PSB2; PSMD5; PSB3; PSMD2; PSMD3; PSB1; PSMD1; PSMF1; PDIA3; PSA5; PSA6; PSA3; PRS8; PSA4; PRS10; PSA1; PRS6A; PSA2; PRS6B; PRS4; PRS7; CALR; SC22B; VAMP3 |
| KEGG | Proteasome (KEGG: hsa03050) | 32/46 | 0.7 | 4E-04 | 8E-03 | -0.119 | PSD12; PSD11; PSDE; PSD13; PSA7; PSB6; PSMD8; PSB7; PSB4; PSMD6; PSB5; PSMD7; PSB2; PSMD4; PSB3; PSMD2; PSB1; PSMD3; PSMD1; PSMF1; PSA5; PSA6; PSA3; PRS8; PSA4; PRS10; PSA1; PRS6A; PSA2; PRS6B; PRS4; PRS7 |
| | ficolin-1-rich granule lumen (GO:1904813) | 63/123 | 0.5 | 3E-06 | 7E-05 | -0.140 | G6PI; DDX3X; HS90B; A1AT; ACPH; PSMD6; PSMD7; ACTY; FRIH; PSMD2; PSMD3; COTL1; OSTF1; GLYG; CATD; PGM1; CATB; ARP2; HS90A; DYLT1; PGAM1; K2C1; NDKB; MIF; PSA5; ACLY; KPYM; PSA2; ALDOC; ALDOA; PPIA; IMB1; VINC; PA1B2; CYTB; PSD12; TERA; ASAH1; PSD11; PSDE; PSD13; GSTP1; MVP; GNS; PSB7; CAND1; PSB1; MK01; LKHA4; TCPQ; HSP7C; XRCC6; GELS; IDHC; HUWE1; HAP28; ARPC5; EF2; PFKAL; PRS6A; BGAL; IMDH2; PRS7; |
| | cytosolic large ribosomal subunit (GO:0022625) | 34/55 | 0.6 | 3E-22 | 4E-20 | -0.349 | RL4; RL5; RL30; RL3; RL10; RL34; RLA1; RL12; RLA0; RL11; RL10A; RL8; RL9; RL6; RL7; RL7A; RL14; RL35; RLA2; RL13; RL15; RL18; RL17; RL19; RL21; RL23; RL22; RL13A; RL23A; RL27A; RL24; RL27; RL29; RL28; |

| | | | | | | | |
|---|---|---|---|---|---|---|---|
| **CC** | large ribosomal subunit (GO:0015934) | 35/59 | 0.6 | 9E-22 | 7E-20 | -0.337 | RL4; RL5; RL30; RL3; RL10; RL34; RLA1; RL12; RLA0; RL11; RL10A; RL8; RL9; RL6; RL7; RL7A; RL14; RL35; RLA2; RL13; RL15; RL18; RL17; RL19; RL21; RL23; RL22; RL13A; RL23A; RM22; RL27A; RL24; RL27; RL29; RL28; |
| | cytosolic small ribosomal subunit (GO:0022627) | 25/40 | 0.6 | 8E-22 | 7E-20 | -0.447 | RS4X; RS14; RS17; RS16; RS19; RS18; RS3; RS2; RS11; RS13; EIF2A; RS12; RS9; RS7; RS8; RS5; RS6; RS3A; RSSA; RS25; RS28; RS29; RS20; RS24; RS23; |
| | small ribosomal subunit (GO:0015935) | 26/45 | 0.6 | 1E-22 | 2E-20 | -0.444 | RS4X; RS14; RS17; RS16; RS19; RS18; RACK1; RS3; RS2; RS11; RS13; EIF2A; RS12; RS9; RS7; RS8; RS5; RS6; RS3A; RSSA; RS25; RS28; RS29; RS20; RS24; RS23; |
| | oxoglutarate dehydrogenase complex (GO:0045252) | 05/05 | 1.0 | 7E-04 | 8E-03 | 0.225 | KGD4; ODO1; ODO2; OGDHL; DLDH; |
| | rough endoplasmic reticulum membrane (GO:0030867) | 04/06 | 0.7 | 2E-02 | 1E-01 | -0.164 | RS28; RS29; RL27; UBA1; |
| | endoplasmic reticulum tubular network membrane (GO:0098826) | 03/05 | 0.6 | 5E-03 | 4E-02 | 0.325 | ATLA3; LNP; RTN4; |

**Table S5. Comparative abundance levels of identified proteins during the final step of oocyte maturation (AMA group). MII *vs.* GV oocytes. Significant differences: fold change (FC) >|1.5|; p.adj ≤ 0.05**

| | Uniprot ID | Protein name | Log2 (FC) | P_value | P.adjusted |
|---|---|---|---|---|---|
| Significantly abundant proteins in MII | Q9H568 | ACTL8 | 2.480 | 0.000 | 0.000 |
| | A6NNZ2 | TBB8B | 1.673 | 0.001 | 0.046 |
| | P80723 | BASP1 | 1.644 | 0.000 | 0.000 |
| | P12004 | PCNA | 1.642 | 0.000 | 0.000 |
| | Q02750 | MP2K1 | 1.424 | 0.000 | 0.002 |
| | Q96MN2 | NALP4 | 1.354 | 0.000 | 0.003 |
| | Q6RW13 | ATRAP | 1.168 | 0.000 | 0.020 |
| | A6NKN8 | PC4L1 | 1.154 | 0.001 | 0.048 |
| | O14965 | AURKA | 1.129 | 0.000 | 0.008 |
| | P62328 | TYB4 | 1.026 | 0.000 | 0.006 |
| | Q96EA4 | SPDLY | 0.914 | 0.000 | 0.013 |
| | O60566 | BUB1B | 0.870 | 0.000 | 0.006 |
| | P60852 | ZP1 | 0.868 | 0.001 | 0.048 |
| | O14737 | PDCD5 | 0.806 | 0.000 | 0.020 |
| | Q9H853 | TBA4B | 0.725 | 0.001 | 0.047 |
| | Q14108 | SCRB2 | 0.634 | 0.001 | 0.048 |
| | Q6GMV3 | PTRD1 | 0.598 | 0.001 | 0.035 |
| | P26358-2 | DNMT1 | 1.223 | 0.005 | 0.090 |
| | P19823 | ITIH2 | 1.151 | 0.011 | 0.136 |
| | P41222 | PTGDS | 1.113 | 0.033 | 0.239 |
| | Q9P2X3 | IMPCT | 1.098 | 0.116 | 0.444 |
| | Q9Y6A5 | TACC3 | 1.098 | 0.003 | 0.061 |
| | P13645 | K1C10 | 1.096 | 0.029 | 0.220 |
| | Q13137-4 | CACO2 | 1.059 | 0.004 | 0.080 |
| | Q9H6H4 | REEP4 | 1.045 | 0.022 | 0.192 |
| | Q9NWY4 | HPF1 | 1.037 | 0.002 | 0.051 |
| | Q12836 | ZP4 | 1.003 | 0.033 | 0.239 |
| | Q96E29 | MTEF3 | 0.986 | 0.040 | 0.264 |
| | P09493-5 | TPM1 | 0.970 | 0.004 | 0.080 |
| | P00568 | KAD1 | 0.941 | 0.005 | 0.092 |
| | Q96PU4 | UHRF2 | 0.927 | 0.018 | 0.181 |
| | Q9UJW0 | DCTN4 | 0.909 | 0.111 | 0.437 |
| | P04004 | VTNC | 0.895 | 0.015 | 0.168 |
| | Q8N183 | NDUF2 | 0.883 | 0.116 | 0.444 |
| | P04792 | HSPB1 | 0.876 | 0.075 | 0.350 |
| | P35908 | K22E | 0.874 | 0.300 | 0.672 |
| | O75477 | ERLN1 | 0.874 | 0.002 | 0.051 |
| | P68036 | UB2L3 | 0.860 | 0.008 | 0.126 |
| | Q06033 | ITIH3 | 0.853 | 0.202 | 0.545 |
| | P56211 | ARP19 | 0.850 | 0.047 | 0.290 |
| | P02748 | CO9 | 0.848 | 0.016 | 0.170 |
| | P61086 | UBE2K | 0.839 | 0.026 | 0.210 |
| | P0C1S8 | WEE2 | 0.835 | 0.013 | 0.152 |
| | O94788 | AL1A2 | 0.826 | 0.009 | 0.127 |
| | P13611 | CSPG2 | 0.825 | 0.134 | 0.459 |
| | P33240 | CSTF2 | 0.822 | 0.004 | 0.080 |
| | P33121 | ACSL1 | 0.808 | 0.018 | 0.181 |
| | Q8WUD1 | RAB2B | 0.807 | 0.069 | 0.345 |
| | Q8WXX5 | DNJC9 | 0.797 | 0.002 | 0.051 |
| | P21754 | ZP3 | 0.792 | 0.011 | 0.136 |
| | P19827 | ITIH1 | 0.782 | 0.107 | 0.424 |

| | | | | |
|---|---|---|---|---|
| Q96AB3 | ISOC2 | 0.779 | 0.021 | 0.191 |
| P10909 | CLUS | 0.778 | 0.018 | 0.181 |
| O14561 | ACPM | 0.776 | 0.009 | 0.127 |
| P62942 | FKB1A | 0.773 | 0.058 | 0.315 |
| Q15185 | TEBP | 0.768 | 0.118 | 0.444 |
| Q07866 | KLC1 | 0.766 | 0.106 | 0.424 |
| Q9UHB6 | LIMA1 | 0.762 | 0.133 | 0.459 |
| P78562 | PHEX | 0.758 | 0.066 | 0.335 |
| Q05996 | ZP2 | 0.745 | 0.025 | 0.208 |
| P28070 | PSB4 | 0.737 | 0.314 | 0.676 |
| Q9NXJ5 | PGPI | 0.730 | 0.114 | 0.444 |
| Q8WVJ2 | NUDC2 | 0.728 | 0.004 | 0.085 |
| O95988 | TCL1B | 0.723 | 0.085 | 0.379 |
| P46976 | GLYG | 0.717 | 0.015 | 0.168 |
| Q92530 | PSMF1 | 0.711 | 0.011 | 0.139 |
| Q9H3P7 | GCP60 | 0.711 | 0.028 | 0.220 |
| P30047 | GFRP | 0.702 | 0.013 | 0.152 |
| P14868 | SYDC | 0.697 | 0.150 | 0.478 |
| P52888 | THOP1 | 0.692 | 0.106 | 0.424 |
| Q16595 | FRDA | 0.683 | 0.044 | 0.273 |
| Q5BJF2 | SGMR2 | 0.671 | 0.314 | 0.676 |
| Q6P1M0 | S27A4 | 0.670 | 1.000 | 1.000 |
| P40925 | MDHC | 0.658 | 0.018 | 0.181 |
| P09497-2 | CLCB | 0.654 | 0.008 | 0.115 |
| P07738 | PMGE | 0.651 | 0.004 | 0.080 |
| Q9NX14 | NDUBB | 0.649 | 0.159 | 0.504 |
| P09936 | UCHL1 | 0.646 | 0.002 | 0.051 |
| Q9Y294 | ASF1A | 0.640 | 0.058 | 0.315 |
| Q06830 | PRDX1 | 0.635 | 0.044 | 0.273 |
| Q9UJS0 | S2513 | 0.632 | 0.018 | 0.181 |
| P55327 | TPD52 | 0.627 | 0.134 | 0.459 |
| Q9BUN8 | DERL1 | 0.621 | 0.235 | 0.590 |
| P22307 | SCP2 | 0.617 | 0.044 | 0.273 |
| Q96AE7-2 | TTC17 | 0.615 | 0.134 | 0.459 |
| Q8NB78 | KDM1B | 0.614 | 0.002 | 0.055 |
| Q9Y5J7 | TIM9 | 0.611 | 0.104 | 0.424 |
| P07942 | LAMB1 | 0.609 | 0.063 | 0.332 |
| Q86UY8 | NT5D3 | 0.607 | 0.095 | 0.403 |
| P04264 | K2C1 | 0.601 | 0.936 | 0.989 |
| P16112 | PGCA | 0.599 | 0.116 | 0.444 |
| Q8N983 | RM43 | 0.599 | 0.209 | 0.558 |
| O94905 | ERLN2 | 0.597 | 0.069 | 0.345 |
| Q16540 | RM23 | 0.581 | 0.345 | 0.706 |
| P47224 | MSS4 | 0.578 | 0.201 | 0.545 |
| P27105 | STOM | 0.576 | 0.029 | 0.220 |
| Q99624 | S38A3 | 0.574 | 0.022 | 0.192 |
| Q658Y4 | F91A1 | 0.573 | 0.063 | 0.332 |
| O60220 | TIM8A | 0.570 | 0.134 | 0.459 |
| P37837 | TALDO | 0.564 | 0.002 | 0.055 |
| Q9UK76 | JUPI1 | 0.560 | 0.217 | 0.567 |
| P56279 | TCL1A | 0.560 | 0.021 | 0.191 |
| P53041 | PPP5 | 0.558 | 0.075 | 0.350 |
| P61981 | 1433G | 0.558 | 0.033 | 0.239 |
| P35749 | MYH11 | 0.554 | 0.122 | 0.450 |
| P49773 | HINT1 | 0.548 | 0.166 | 0.504 |
| P63172 | DYLT1 | 0.547 | 0.051 | 0.292 |

| | | | | |
|---|---|---|---|---|
| Q9H300 | PARL | 0.544 | 0.006 | 0.101 |
| Q9BUT1 | BDH2 | 0.540 | 0.066 | 0.335 |
| P17152 | TMM11 | 0.539 | 0.134 | 0.459 |
| Q92820 | GGH | 0.532 | 0.044 | 0.273 |
| Q96EK5 | KBP | 0.532 | 0.066 | 0.335 |
| O43768 | ENSA | 0.530 | 0.134 | 0.459 |
| Q9NR46 | SHLB2 | 0.529 | 0.149 | 0.478 |
| Q15631 | TSN | 0.527 | 0.149 | 0.478 |
| O00170 | AIP | 0.527 | 0.063 | 0.332 |
| O43684 | BUB3 | 0.526 | 0.413 | 0.761 |
| Q6P1S2 | CC033 | 0.524 | 0.099 | 0.415 |
| P11055 | MYH3 | 0.521 | 0.063 | 0.332 |
| P46977 | STT3A | 0.521 | 0.201 | 0.545 |
| Q14554 | PDIA5 | 0.517 | 0.217 | 0.567 |
| Q14738 | 2A5D | 0.514 | 0.456 | 0.778 |
| O75531 | BAF | 0.513 | 0.025 | 0.208 |
| Q8WTV0-2 | SCRB1 | 0.512 | 0.082 | 0.376 |
| Q96CN9 | GCC1 | 0.509 | 0.347 | 0.706 |
| Q8N2F6 | ARM10 | 0.508 | 0.149 | 0.478 |
| Q9UNS2 | CSN3 | 0.508 | 0.015 | 0.169 |
| P61077 | UB2D3 | 0.506 | 0.202 | 0.545 |
| Q9Y6N5 | SQOR | 0.504 | 0.026 | 0.210 |
| P12277 | KCRB | 0.500 | 0.058 | 0.315 |
| Q15843 | NEDD8 | 0.495 | 0.007 | 0.115 |
| Q9UMX5 | NENF | 0.495 | 0.009 | 0.127 |
| Q9BV19 | CA050 | 0.492 | 0.149 | 0.478 |
| P11047 | LAMC1 | 0.488 | 0.183 | 0.528 |
| Q9Y3B8 | ORN | 0.488 | 0.002 | 0.051 |
| Q86SX6 | GLRX5 | 0.486 | 0.066 | 0.335 |
| P80192 | M3K9 | 0.482 | 0.034 | 0.240 |
| P04114 | APOB | 0.481 | 0.120 | 0.444 |
| P62937 | PPIA | 0.481 | 0.134 | 0.459 |
| P34949 | MPI | 0.473 | 0.004 | 0.074 |
| Q9BRA2 | TXD17 | 0.472 | 0.002 | 0.055 |
| Q9HCC0 | MCCB | 0.472 | 0.193 | 0.545 |
| Q08431 | MFGM | 0.471 | 0.434 | 0.767 |
| P35606 | COPB2 | 0.469 | 0.009 | 0.127 |
| P08238 | HS90B | 0.466 | 0.134 | 0.459 |
| Q8N5K1 | CISD2 | 0.464 | 0.042 | 0.268 |
| Q14677 | EPN4 | 0.463 | 0.013 | 0.152 |
| P12235 | ADT1 | 0.461 | 0.107 | 0.424 |
| Q9H9B4 | SFXN1 | 0.460 | 0.015 | 0.168 |
| P10768 | ESTD | 0.460 | 0.252 | 0.609 |
| Q9UM54-6 | MYO6 | 0.455 | 0.267 | 0.623 |
| Q99497 | PARK7 | 0.446 | 0.004 | 0.080 |
| Q96CN7 | ISOC1 | 0.445 | 0.058 | 0.315 |
| P60033 | CD81 | 0.445 | 0.149 | 0.478 |
| P53999 | TCP4 | 0.440 | 0.066 | 0.335 |
| O75347 | TBCA | 0.439 | 0.021 | 0.191 |
| Q9UPS6 | SET1B | 0.434 | 0.863 | 0.971 |
| Q99627 | CSN8 | 0.434 | 0.085 | 0.379 |
| Q14008 | CKAP5 | 0.433 | 0.009 | 0.127 |
| Q01433 | AMPD2 | 0.431 | 0.347 | 0.706 |
| Q8WW22 | DNJA4 | 0.430 | 0.151 | 0.479 |
| P60174 | TPIS | 0.429 | 0.317 | 0.676 |
| P49189 | AL9A1 | 0.428 | 0.003 | 0.061 |

| | | | | |
|---|---|---|---|---|
| Q9Y262 | EIF3L | 0.426 | 0.512 | 0.809 |
| Q92734 | TFG | 0.425 | 0.252 | 0.609 |
| P09417 | DHPR | 0.425 | 0.029 | 0.220 |
| P57105 | SYJ2B | 0.425 | 0.166 | 0.504 |
| P26641 | EF1G | 0.425 | 0.033 | 0.239 |
| P49441 | INPP | 0.424 | 0.022 | 0.192 |
| O60443 | GSDME | 0.424 | 0.284 | 0.656 |
| Q96JY0 | MAEL | 0.424 | 0.063 | 0.332 |
| Q96KP4 | CNDP2 | 0.423 | 0.011 | 0.136 |
| Q9Y285 | SYFA | 0.423 | 0.648 | 0.872 |
| Q13510 | ASAH1 | 0.422 | 0.344 | 0.706 |
| Q08257 | QOR | 0.422 | 0.183 | 0.528 |
| Q8NE86 | MCU | 0.421 | 0.373 | 0.725 |
| P62826 | RAN | 0.419 | 0.038 | 0.255 |
| P01031 | CO5 | 0.413 | 0.149 | 0.478 |
| P58546 | MTPN | 0.411 | 0.041 | 0.265 |
| Q15836 | VAMP3 | 0.410 | 0.344 | 0.706 |
| Q12907 | LMAN2 | 0.410 | 0.018 | 0.181 |
| Q9NX76 | CKLF6 | 0.409 | 0.282 | 0.652 |
| Q16881 | TRXR1 | 0.408 | 0.011 | 0.136 |
| Q13491-4 | GPM6B | 0.405 | 0.208 | 0.556 |
| P05023 | AT1A1 | 0.405 | 0.066 | 0.335 |
| P00492 | HPRT | 0.404 | 0.011 | 0.136 |
| Q05639 | EF1A2 | 0.403 | 0.075 | 0.350 |
| P04080 | CYTB | 0.399 | 0.150 | 0.478 |
| P56134 | ATPK | 0.399 | 0.095 | 0.403 |
| P61586 | RHOA | 0.398 | 0.494 | 0.796 |
| Q9UMY4 | SNX12 | 0.397 | 0.030 | 0.229 |
| Q96DV4 | RM38 | 0.395 | 0.767 | 0.944 |
| Q9P2B2 | FPRP | 0.390 | 0.373 | 0.725 |
| Q92581-2 | SL9A6 | 0.389 | 0.418 | 0.767 |
| P05556 | ITB1 | 0.388 | 0.033 | 0.239 |
| Q9HD67 | MYO10 | 0.386 | 0.149 | 0.478 |
| Q9H4A5 | GLP3L | 0.385 | 0.058 | 0.315 |
| Q9Y3C8 | UFC1 | 0.384 | 0.047 | 0.290 |
| P31947 | 1433S | 0.383 | 1.000 | 1.000 |
| P07686 | HEXB | 0.383 | 0.317 | 0.676 |
| Q93034 | CUL5 | 0.380 | 0.007 | 0.115 |
| Q13277 | STX3 | 0.380 | 0.222 | 0.567 |
| P29218 | IMPA1 | 0.378 | 0.149 | 0.478 |
| Q04760 | LGUL | 0.377 | 0.202 | 0.545 |
| P27361 | MK03 | 0.377 | 0.235 | 0.590 |
| P09493-10 | TPM1 | 0.377 | 0.011 | 0.136 |
| P52209 | 6PGD | 0.376 | 0.120 | 0.444 |
| P52565 | GDIR1 | 0.375 | 0.029 | 0.220 |
| Q3ZCM7 | TBB8 | 0.373 | 0.095 | 0.403 |
| P63241 | IF5A1 | 0.373 | 0.373 | 0.725 |
| P51149 | RAB7A | 0.372 | 0.021 | 0.191 |
| O95433 | AHSA1 | 0.370 | 0.030 | 0.229 |
| O60716 | CTND1 | 0.370 | 0.202 | 0.545 |
| Q8IWB7 | WDFY1 | 0.369 | 0.120 | 0.444 |
| Q9Y263 | PLAP | 0.368 | 0.085 | 0.379 |
| A0MZ66 | SHOT1 | 0.368 | 0.035 | 0.248 |
| Q96AB6 | NTAN1 | 0.366 | 0.011 | 0.136 |
| P00441 | SODC | 0.366 | 0.291 | 0.657 |
| Q99426 | TBCB | 0.365 | 0.149 | 0.478 |

| | | | | |
|---|---|---|---|---|
| P08572 | CO4A2 | 0.365 | 0.317 | 0.676 |
| Q00169 | PIPNA | 0.365 | 0.066 | 0.335 |
| O43504 | LTOR5 | 0.361 | 0.026 | 0.210 |
| Q2TAA2 | IAH1 | 0.360 | 0.459 | 0.778 |
| P00558 | PGK1 | 0.360 | 0.727 | 0.922 |
| Q9UNF0 | PACN2 | 0.359 | 0.277 | 0.642 |
| P53990 | IST1 | 0.358 | 0.345 | 0.706 |
| P62857 | RS28 | 0.357 | 0.107 | 0.424 |
| O95168 | NDUB4 | 0.356 | 0.602 | 0.860 |
| Q96T88 | UHRF1 | 0.356 | 0.000 | 0.020 |
| Q86W25 | NAL13 | 0.355 | 0.183 | 0.528 |
| P13693 | TCTP | 0.352 | 0.035 | 0.248 |
| P23229 | ITA6 | 0.345 | 0.344 | 0.706 |
| Q6UXH1 | CREL2 | 0.345 | 0.349 | 0.709 |
| Q9BV79 | MECR | 0.345 | 0.317 | 0.676 |
| Q9Y333 | LSM2 | 0.341 | 0.183 | 0.528 |
| Q99714 | HCD2 | 0.341 | 0.107 | 0.424 |
| P09960 | LKHA4 | 0.341 | 0.069 | 0.345 |
| Q4J6C6 | PPCEL | 0.341 | 0.243 | 0.597 |
| Q6X9E4 | FBW12 | 0.339 | 0.093 | 0.403 |
| P23588 | IF4B | 0.338 | 0.134 | 0.459 |
| O60610 | DIAP1 | 0.334 | 0.134 | 0.459 |
| Q9HD34 | LYRM4 | 0.334 | 0.120 | 0.444 |
| P41732 | TSN7 | 0.333 | 0.536 | 0.819 |
| P06753 | TPM3 | 0.333 | 0.134 | 0.459 |
| Q96IU4 | ABHEB | 0.332 | 0.252 | 0.609 |
| P20929-2 | NEBU | 0.332 | 0.222 | 0.567 |
| O00560 | SDCB1 | 0.332 | 0.609 | 0.860 |
| P49789 | FHIT | 0.329 | 0.120 | 0.444 |
| P23434 | GCSH | 0.329 | 0.431 | 0.767 |
| P11413 | G6PD | 0.328 | 0.222 | 0.567 |
| Q9NVT9 | ARMC1 | 0.327 | 0.531 | 0.819 |
| Q9UNZ2 | NSF1C | 0.326 | 0.015 | 0.168 |
| P61923 | COPZ1 | 0.325 | 0.069 | 0.345 |
| Q8NBS9 | TXND5 | 0.325 | 0.015 | 0.168 |
| O75608 | LYPA1 | 0.325 | 0.222 | 0.567 |
| O00232 | PSD12 | 0.325 | 0.192 | 0.545 |
| P53618 | COPB | 0.325 | 0.025 | 0.208 |
| P62312 | LSM6 | 0.322 | 0.106 | 0.424 |
| O60218 | AK1BA | 0.320 | 0.107 | 0.424 |
| O43752 | STX6 | 0.318 | 0.222 | 0.567 |
| P06396 | GELS | 0.315 | 0.029 | 0.220 |
| Q8IYB1 | M21D2 | 0.315 | 0.149 | 0.478 |
| P08237 | PFKAM | 0.314 | 0.169 | 0.510 |
| Q96FW1 | OTUB1 | 0.313 | 0.082 | 0.376 |
| P25685 | DNJB1 | 0.313 | 0.026 | 0.210 |
| Q9H330 | TM245 | 0.312 | 0.344 | 0.706 |
| P08574 | CY1 | 0.312 | 0.183 | 0.528 |
| P21266 | GSTM3 | 0.311 | 0.222 | 0.567 |
| Q8NC96 | NECP1 | 0.311 | 0.344 | 0.706 |
| O00330 | ODPX | 0.310 | 0.183 | 0.528 |
| Q9Y678 | COPG1 | 0.307 | 0.202 | 0.545 |
| Q8N131 | PORIM | 0.306 | 0.459 | 0.778 |
| Q02224 | CENPE | 0.306 | 0.295 | 0.662 |
| P50395 | GDIB | 0.305 | 0.222 | 0.567 |
| P35613 | BASI | 0.303 | 0.267 | 0.623 |

| | | | | |
|---|---|---|---|---|
| P60953 | CDC42 | 0.302 | 0.082 | 0.376 |
| P17301 | ITA2 | 0.300 | 0.063 | 0.332 |
| P59998 | ARPC4 | 0.299 | 0.647 | 0.872 |
| P32119 | PRDX2 | 0.298 | 0.809 | 0.953 |
| Q96FJ2 | DYL2 | 0.297 | 0.501 | 0.796 |
| O75306 | NDUS2 | 0.297 | 0.347 | 0.706 |
| P55789 | ALR | 0.297 | 0.392 | 0.749 |
| P06396-2 | GELS | 0.295 | 0.572 | 0.842 |
| P31948-2 | STIP1 | 0.295 | 0.107 | 0.424 |
| O14949 | QCR8 | 0.294 | 0.259 | 0.623 |
| P61201 | CSN2 | 0.293 | 0.120 | 0.444 |
| P61604 | CH10 | 0.293 | 0.003 | 0.072 |
| P50053-2 | KHK | 0.293 | 0.167 | 0.505 |
| Q9BSH5 | HDHD3 | 0.293 | 0.222 | 0.567 |
| P02649 | APOE | 0.292 | 0.501 | 0.796 |
| Q9Y4I1 | MYO5A | 0.292 | 0.691 | 0.900 |
| P18859 | ATP5J | 0.292 | 0.075 | 0.350 |
| Q9UBQ0 | VPS29 | 0.292 | 0.851 | 0.963 |
| P36957 | ODO2 | 0.291 | 0.058 | 0.315 |
| P14618 | KPYM | 0.291 | 0.344 | 0.706 |
| P55786 | PSA | 0.289 | 0.051 | 0.292 |
| Q96A57 | TM230 | 0.287 | 0.251 | 0.609 |
| Q9NTJ5 | SAC1 | 0.287 | 0.202 | 0.545 |
| P30086 | PEBP1 | 0.287 | 0.183 | 0.528 |
| P40926 | MDHM | 0.285 | 0.011 | 0.136 |
| P06744 | G6PI | 0.284 | 0.727 | 0.922 |
| O95571 | ETHE1 | 0.284 | 0.134 | 0.459 |
| Q12846 | STX4 | 0.283 | 0.434 | 0.767 |
| Q8TDQ7 | GNPI2 | 0.283 | 0.120 | 0.444 |
| P21291 | CSRP1 | 0.282 | 0.649 | 0.872 |
| O43813 | LANC1 | 0.281 | 0.075 | 0.350 |
| Q15785 | TOM34 | 0.281 | 0.609 | 0.860 |
| Q14643-2 | ITPR1 | 0.280 | 0.095 | 0.403 |
| P30049 | ATPD | 0.278 | 0.107 | 0.424 |
| P63010-3 | AP2B1 | 0.277 | 0.173 | 0.521 |
| P21912 | SDHB | 0.277 | 0.033 | 0.239 |
| P37268 | FDFT | 0.276 | 0.459 | 0.778 |
| P04179 | SODM | 0.276 | 0.222 | 0.567 |
| P61019 | RAB2A | 0.275 | 0.222 | 0.567 |
| P23528 | COF1 | 0.274 | 0.267 | 0.623 |
| P46926 | GNPI1 | 0.272 | 0.075 | 0.350 |
| O00154 | BACH | 0.271 | 0.066 | 0.335 |
| P51809 | VAMP7 | 0.268 | 0.392 | 0.749 |
| O00244 | ATOX1 | 0.267 | 0.316 | 0.676 |
| Q15043 | S39AE | 0.267 | 0.955 | 0.999 |
| P09543 | CN37 | 0.266 | 0.369 | 0.725 |
| Q7L5D6 | GET4 | 0.265 | 0.605 | 0.860 |
| P34932 | HSP74 | 0.265 | 0.038 | 0.255 |
| Q9Y2H1 | ST38L | 0.264 | 0.244 | 0.597 |
| Q9Y639 | NPTN | 0.263 | 0.467 | 0.778 |
| Q86SE8 | NPM2 | 0.262 | 0.066 | 0.335 |
| Q6TGC4 | PADI6 | 0.261 | 0.166 | 0.504 |
| Q9NZ45 | CISD1 | 0.260 | 0.033 | 0.239 |
| P82909 | KGD4 | 0.259 | 0.021 | 0.191 |
| Q99816 | TS101 | 0.259 | 0.015 | 0.168 |
| Q9UIC8 | LCMT1 | 0.259 | 0.291 | 0.657 |

| | | | | |
|---|---|---|---|---|
| O95822 | DCMC | 0.259 | 0.075 | 0.350 |
| O60925 | PFD1 | 0.258 | 0.459 | 0.778 |
| O95674 | CDS2 | 0.255 | 0.291 | 0.657 |
| Q8NC56 | LEMD2 | 0.254 | 0.331 | 0.700 |
| P27797 | CALR | 0.252 | 0.015 | 0.168 |
| P11766 | ADHX | 0.252 | 0.434 | 0.767 |
| Q3SXM5 | HSDL1 | 0.252 | 0.244 | 0.597 |
| Q9NQW7 | XPP1 | 0.250 | 0.449 | 0.778 |
| P98194 | AT2C1 | 0.250 | 0.261 | 0.623 |
| Q8N5M4 | TTC9C | 0.249 | 0.820 | 0.957 |
| P24534 | EF1B | 0.249 | 0.727 | 0.922 |
| Q7L5N1 | CSN6 | 0.248 | 0.467 | 0.778 |
| O75874 | IDHC | 0.248 | 0.095 | 0.403 |
| P28066 | PSA5 | 0.247 | 0.051 | 0.292 |
| P54578 | UBP14 | 0.246 | 0.244 | 0.597 |
| P17987 | TCPA | 0.246 | 0.609 | 0.860 |
| P29692 | EF1D | 0.245 | 0.851 | 0.963 |
| P36507 | MP2K2 | 0.244 | 0.424 | 0.767 |
| O14562 | UBFD1 | 0.243 | 0.252 | 0.609 |
| Q13098 | CSN1 | 0.243 | 0.095 | 0.403 |
| P15121 | ALDR | 0.243 | 0.727 | 0.922 |
| Q15084 | PDIA6 | 0.243 | 0.021 | 0.191 |
| P28838 | AMPL | 0.242 | 0.317 | 0.676 |
| P48637 | GSHB | 0.241 | 0.202 | 0.545 |
| Q9Y6Y8 | S23IP | 0.241 | 0.317 | 0.676 |
| Q01469 | FABP5 | 0.241 | 0.738 | 0.931 |
| P04066 | FUCO | 0.240 | 0.536 | 0.819 |
| O75380 | NDUS6 | 0.240 | 0.976 | 1.000 |
| P36405 | ARL3 | 0.239 | 0.228 | 0.579 |
| Q9UL26 | RB22A | 0.239 | 0.344 | 0.706 |
| Q86WR0 | CCD25 | 0.238 | 0.628 | 0.872 |
| P35080 | PROF2 | 0.238 | 0.202 | 0.545 |
| Q9BY44 | EIF2A | 0.237 | 0.166 | 0.504 |
| P11234 | RALB | 0.236 | 0.291 | 0.657 |
| P63000 | RAC1 | 0.236 | 0.317 | 0.676 |
| Q9BSQ5 | CCM2 | 0.235 | 0.424 | 0.767 |
| P22061-2 | PIMT | 0.235 | 0.183 | 0.528 |
| P04844-2 | RPN2 | 0.233 | 0.467 | 0.778 |
| P63104 | 1433Z | 0.233 | 0.403 | 0.752 |
| O75521 | ECI2 | 0.230 | 0.066 | 0.335 |
| P62258 | 1433E | 0.230 | 0.467 | 0.778 |
| O00203 | AP3B1 | 0.229 | 0.752 | 0.943 |
| P17174 | AATC | 0.229 | 0.222 | 0.567 |
| Q06418 | TYRO3 | 0.227 | 0.267 | 0.623 |
| Q96DZ1 | ERLEC | 0.226 | 0.222 | 0.567 |
| P35580 | MYH10 | 0.226 | 0.183 | 0.528 |
| Q8NBX0 | SCPDL | 0.225 | 0.434 | 0.767 |
| P63092 | GNAS2 | 0.223 | 0.628 | 0.872 |
| P61160 | ARP2 | 0.222 | 0.222 | 0.567 |
| Q14914 | PTGR1 | 0.222 | 0.119 | 0.444 |
| Q99832 | TCPH | 0.222 | 0.809 | 0.953 |
| Q8N6M9 | ZFN2A | 0.221 | 0.243 | 0.597 |
| O14618 | CCS | 0.219 | 0.531 | 0.819 |
| Q9Y625 | GPC6 | 0.219 | 0.557 | 0.842 |
| O43639 | NCK2 | 0.219 | 0.459 | 0.778 |
| Q9BSD7 | NTPCR | 0.219 | 0.317 | 0.676 |

| | | | | |
|---|---|---|---|---|
| O75083 | WDR1 | 0.218 | 0.403 | 0.752 |
| P06730 | IF4E | 0.217 | 0.166 | 0.504 |
| Q9GZT8 | NIF3L | 0.217 | 0.752 | 0.943 |
| P08758 | ANXA5 | 0.217 | 0.609 | 0.860 |
| P06493 | CDK1 | 0.215 | 0.608 | 0.860 |
| O95197 | RTN3 | 0.213 | 0.936 | 0.989 |
| Q9Y4E8 | UBP15 | 0.213 | 0.872 | 0.977 |
| Q9P035 | HACD3 | 0.210 | 0.190 | 0.544 |
| P53985 | MOT1 | 0.210 | 0.434 | 0.767 |
| Q9Y679 | AUP1 | 0.208 | 0.361 | 0.725 |
| P59045 | NAL11 | 0.207 | 0.434 | 0.767 |
| Q9NTX5 | ECHD1 | 0.207 | 0.434 | 0.767 |
| O00214-2 | LEG8 | 0.206 | 0.608 | 0.860 |
| P62310 | LSM3 | 0.205 | 0.727 | 0.922 |
| O14579 | COPE | 0.205 | 0.317 | 0.676 |
| Q9NRW7 | VPS45 | 0.204 | 0.776 | 0.944 |
| O95373 | IPO7 | 0.204 | 0.608 | 0.860 |
| Q5SW96 | ARH | 0.202 | 0.494 | 0.796 |
| O95716 | RAB3D | 0.202 | 0.183 | 0.528 |
| Q9NYU2 | UGGG1 | 0.201 | 0.134 | 0.459 |
| Q9UI09 | NDUAC | 0.199 | 0.841 | 0.963 |
| Q9NTK5 | OLA1 | 0.199 | 0.483 | 0.796 |
| P49321 | NASP | 0.199 | 0.051 | 0.292 |
| Q5JSH3 | WDR44 | 0.198 | 0.291 | 0.657 |
| P07237 | PDIA1 | 0.198 | 0.085 | 0.379 |
| P59047 | NALP5 | 0.198 | 0.120 | 0.444 |
| P78406 | RAE1L | 0.197 | 0.809 | 0.953 |
| Q15042 | RB3GP | 0.196 | 1.000 | 1.000 |
| P07900 | HS90A | 0.196 | 0.267 | 0.623 |
| Q15773 | MLF2 | 0.195 | 0.771 | 0.944 |
| P21281 | VATB2 | 0.194 | 0.403 | 0.752 |
| P61018 | RAB4B | 0.194 | 0.403 | 0.752 |
| P43686 | PRS6B | 0.194 | 0.183 | 0.528 |
| Q86WA6 | BPHL | 0.193 | 0.371 | 0.725 |
| P63167 | DYL1 | 0.192 | 0.687 | 0.900 |
| Q7Z2K6 | ERMP1 | 0.191 | 0.373 | 0.725 |
| Q9UI12 | VATH | 0.191 | 0.647 | 0.872 |
| Q15388 | TOM20 | 0.191 | 0.459 | 0.778 |
| P34896-2 | GLYC | 0.191 | 0.656 | 0.875 |
| Q9NP81 | SYSM | 0.190 | 0.038 | 0.255 |
| P18084 | ITB5 | 0.190 | 0.562 | 0.842 |
| Q9NQC3 | RTN4 | 0.190 | 0.202 | 0.545 |
| P31150 | GDIA | 0.189 | 0.291 | 0.657 |
| P24666 | PPAC | 0.187 | 0.392 | 0.749 |
| P61081 | UBC12 | 0.187 | 0.446 | 0.778 |
| Q96GK7 | FAH2A | 0.186 | 0.727 | 0.922 |
| P11142 | HSP7C | 0.184 | 0.851 | 0.963 |
| P18669 | PGAM1 | 0.184 | 0.651 | 0.872 |
| Q9UGP8 | SEC63 | 0.182 | 0.166 | 0.504 |
| Q9BRQ6 | MIC25 | 0.182 | 0.149 | 0.478 |
| Q00266 | METK1 | 0.181 | 0.841 | 0.963 |
| Q9H9J2 | RM44 | 0.181 | 1.000 | 1.000 |
| O14578-4 | CTRO | 0.179 | 0.512 | 0.809 |
| Q9Y2S7 | PDIP2 | 0.179 | 0.572 | 0.842 |
| Q9NPJ3 | ACO13 | 0.178 | 0.851 | 0.963 |
| Q8WX94-2 | NALP7 | 0.178 | 0.095 | 0.403 |

| | | | | |
|---|---|---|---|---|
| Q96D96 | HVCN1 | 0.178 | 0.403 | 0.752 |
| Q68EM7 | RHG17 | 0.178 | 0.536 | 0.819 |
| Q13630 | FCL | 0.177 | 0.166 | 0.504 |
| Q9NX63 | MIC19 | 0.177 | 0.149 | 0.478 |
| Q14696 | MESD | 0.176 | 0.134 | 0.459 |
| Q9BUR5 | MIC26 | 0.175 | 0.434 | 0.767 |
| Q9H9A6 | LRC40 | 0.175 | 0.166 | 0.504 |
| Q96PC5 | MIA2 | 0.174 | 0.809 | 0.953 |
| P35754 | GLRX1 | 0.174 | 0.459 | 0.778 |
| Q08722 | CD47 | 0.173 | 0.501 | 0.796 |
| Q13153 | PAK1 | 0.172 | 0.569 | 0.842 |
| Q96IQ9 | ZN414 | 0.172 | 0.766 | 0.944 |
| P53621 | COPA | 0.172 | 0.107 | 0.424 |
| Q96B97 | SH3K1 | 0.171 | 0.767 | 0.944 |
| Q562R1 | ACTBL | 0.170 | 0.809 | 0.953 |
| P53992 | SC24C | 0.170 | 0.317 | 0.676 |
| O00429 | DNM1L | 0.169 | 0.267 | 0.623 |
| Q99735 | MGST2 | 0.168 | 0.557 | 0.842 |
| Q9NYQ8 | FAT2 | 0.168 | 0.434 | 0.767 |
| Q02978 | M2OM | 0.168 | 0.228 | 0.579 |
| Q9P0M9 | RM27 | 0.168 | 0.654 | 0.874 |
| A6NDX5 | ZN840 | 0.168 | 0.095 | 0.403 |
| Q16204 | CCDC6 | 0.167 | 0.267 | 0.623 |
| O00273 | DFFA | 0.167 | 0.865 | 0.971 |
| Q13442 | HAP28 | 0.166 | 0.649 | 0.872 |
| Q00610 | CLH1 | 0.165 | 0.120 | 0.444 |
| P39656 | OST48 | 0.164 | 0.317 | 0.676 |
| O75947 | ATP5H | 0.164 | 0.134 | 0.459 |
| Q9Y5X1 | SNX9 | 0.164 | 1.000 | 1.000 |
| P30626 | SORCN | 0.162 | 0.344 | 0.706 |
| Q15257-2 | PTPA | 0.161 | 0.267 | 0.623 |
| P21926 | CD9 | 0.160 | 0.898 | 0.984 |
| Q99436 | PSB7 | 0.160 | 0.727 | 0.922 |
| P30038 | AL4A1 | 0.159 | 0.202 | 0.545 |
| Q13409 | DC1I2 | 0.159 | 0.434 | 0.767 |
| P30040 | ERP29 | 0.158 | 0.183 | 0.528 |
| Q9BPW8 | NIPS1 | 0.157 | 0.244 | 0.597 |
| O95573 | ACSL3 | 0.156 | 0.467 | 0.778 |
| P47985 | UCRI | 0.156 | 0.776 | 0.944 |
| Q86YB8 | ERO1B | 0.156 | 0.647 | 0.872 |
| Q9BT09 | CNPY3 | 0.156 | 0.373 | 0.725 |
| P23526 | SAHH | 0.156 | 0.267 | 0.623 |
| Q96AG4 | LRC59 | 0.155 | 0.095 | 0.403 |
| Q13505 | MTX1 | 0.154 | 0.609 | 0.860 |
| Q14651 | PLSI | 0.153 | 0.467 | 0.778 |
| P07737 | PROF1 | 0.153 | 0.809 | 0.953 |
| O60927 | PP1RB | 0.153 | 0.628 | 0.872 |
| P49368 | TCPG | 0.152 | 0.936 | 0.989 |
| O75348 | VATG1 | 0.150 | 0.512 | 0.809 |
| O60493 | SNX3 | 0.150 | 0.501 | 0.796 |
| Q8N1G4 | LRC47 | 0.149 | 0.647 | 0.872 |
| P22059 | OSBP1 | 0.149 | 0.267 | 0.623 |
| Q96C36 | P5CR2 | 0.148 | 0.841 | 0.963 |
| P40227 | TCPZ | 0.148 | 0.767 | 0.944 |
| Q9NX02 | NALP2 | 0.148 | 0.467 | 0.778 |
| P04406 | G3P | 0.147 | 0.727 | 0.922 |

| | | | | |
|---|---|---|---|---|
| Q9NY33 | DPP3 | 0.147 | 0.501 | 0.796 |
| P00505 | AATM | 0.147 | 0.373 | 0.725 |
| Q9NRX4 | PHP14 | 0.147 | 0.501 | 0.796 |
| Q15833 | STXB2 | 0.146 | 0.166 | 0.504 |
| Q92905 | CSN5 | 0.146 | 0.303 | 0.676 |
| Q8NCW5 | NNRE | 0.145 | 0.531 | 0.819 |
| P60520 | GBRL2 | 0.145 | 0.303 | 0.676 |
| P00390 | GSHR | 0.144 | 0.467 | 0.778 |
| P42261 | GRIA1 | 0.144 | 0.955 | 0.999 |
| Q16851 | UGPA | 0.144 | 0.149 | 0.478 |
| P61221 | ABCE1 | 0.143 | 0.494 | 0.796 |
| P30041 | PRDX6 | 0.142 | 0.979 | 1.000 |
| O95757 | HS74L | 0.141 | 0.244 | 0.597 |
| P62273 | RS29 | 0.141 | 0.483 | 0.796 |
| Q13428-4 | TCOF | 0.141 | 0.317 | 0.676 |
| A0FGR8 | ESYT2 | 0.141 | 0.539 | 0.821 |
| Q99653 | CHP1 | 0.140 | 0.976 | 1.000 |
| P16435 | NCPR | 0.140 | 0.609 | 0.860 |
| O43242 | PSMD3 | 0.139 | 0.373 | 0.725 |
| Q9Y2V2 | CHSP1 | 0.139 | 0.910 | 0.989 |
| P39210 | MPV17 | 0.138 | 0.439 | 0.775 |
| Q9H2W6 | RM46 | 0.136 | 0.365 | 0.725 |
| P30520 | PURA2 | 0.136 | 0.824 | 0.960 |
| Q8WWV3 | RT4I1 | 0.135 | 0.893 | 0.982 |
| Q9HC38 | GLOD4 | 0.135 | 1.000 | 1.000 |
| Q96HY6 | DDRGK | 0.134 | 0.738 | 0.931 |
| Q15691 | MARE1 | 0.134 | 0.373 | 0.725 |
| Q9Y2Z4 | SYYM | 0.133 | 0.291 | 0.657 |
| P07954 | FUMH | 0.133 | 0.434 | 0.767 |
| Q8TB36 | GDAP1 | 0.133 | 0.344 | 0.706 |
| Q9NS69 | TOM22 | 0.132 | 0.497 | 0.796 |
| P43304 | GPDM | 0.132 | 0.403 | 0.752 |
| Q6UW68 | TM205 | 0.130 | 0.373 | 0.725 |
| Q92743 | HTRA1 | 0.130 | 1.000 | 1.000 |
| O15258 | RER1 | 0.130 | 0.483 | 0.796 |
| O60869 | EDF1 | 0.129 | 0.649 | 0.872 |
| P11233 | RALA | 0.129 | 0.647 | 0.872 |
| P55809 | SCOT1 | 0.129 | 0.393 | 0.751 |
| Q15382 | RHEB | 0.129 | 0.652 | 0.872 |
| Q9H444 | CHM4B | 0.129 | 0.467 | 0.778 |
| Q13177 | PAK2 | 0.128 | 0.267 | 0.623 |
| P36542 | ATPG | 0.128 | 0.609 | 0.860 |
| Q13526 | PIN1 | 0.128 | 0.569 | 0.842 |
| P51570 | GALK1 | 0.127 | 0.936 | 0.989 |
| O60831 | PRAF2 | 0.126 | 0.434 | 0.767 |
| Q16401 | PSMD5 | 0.125 | 0.317 | 0.676 |
| O43854 | EDIL3 | 0.124 | 0.898 | 0.984 |
| Q96IX5 | ATPMK | 0.123 | 0.865 | 0.971 |
| O00515 | LAD1 | 0.123 | 0.373 | 0.725 |
| P78540 | ARGI2 | 0.122 | 0.149 | 0.478 |
| P00167 | CYB5 | 0.122 | 0.536 | 0.819 |
| P42025 | ACTY | 0.121 | 0.809 | 0.953 |
| L0R6Q1 | S35U4 | 0.120 | 0.609 | 0.860 |
| O43617 | TPPC3 | 0.120 | 0.695 | 0.902 |
| P09619 | PGFRB | 0.120 | 0.767 | 0.944 |
| O43196 | MSH5 | 0.119 | 0.536 | 0.819 |

| | | | | |
|---|---|---|---|---|
| O75027 | ABCB7 | 0.118 | 0.569 | 0.842 |
| P20073 | ANXA7 | 0.118 | 0.851 | 0.963 |
| Q92882 | OSTF1 | 0.116 | 0.572 | 0.842 |
| Q96AG3 | S2546 | 0.115 | 0.494 | 0.796 |
| Q14203 | DCTN1 | 0.115 | 0.809 | 0.953 |
| P78371 | TCPB | 0.114 | 0.467 | 0.778 |
| Q9H3K6 | BOLA2 | 0.114 | 0.649 | 0.872 |
| P61163 | ACTZ | 0.113 | 0.820 | 0.957 |
| P48047 | ATPO | 0.113 | 0.202 | 0.545 |
| P21796 | VDAC1 | 0.113 | 0.501 | 0.796 |
| P22695 | QCR2 | 0.112 | 0.373 | 0.725 |
| O15116 | LSM1 | 0.112 | 0.317 | 0.676 |
| Q15907 | RB11B | 0.112 | 0.449 | 0.778 |
| P47897 | SYQ | 0.111 | 0.647 | 0.872 |
| O95831 | AIFM1 | 0.110 | 0.291 | 0.657 |
| P26885 | FKBP2 | 0.110 | 0.501 | 0.796 |
| Q5T2N8 | ATD3C | 0.110 | 0.291 | 0.657 |
| Q13561 | DCTN2 | 0.109 | 0.291 | 0.657 |
| P09382 | LEG1 | 0.108 | 0.467 | 0.778 |
| P84550 | SKOR1 | 0.108 | 0.776 | 0.944 |
| O94826 | TOM70 | 0.107 | 0.107 | 0.424 |
| Q86X76 | NIT1 | 0.105 | 0.687 | 0.900 |
| Q9H9Q2 | CSN7B | 0.105 | 0.572 | 0.842 |
| P12268 | IMDH2 | 0.105 | 0.727 | 0.922 |
| Q96GG9 | DCNL1 | 0.104 | 0.691 | 0.900 |
| Q9HCU5 | PREB | 0.104 | 0.467 | 0.778 |
| Q10589 | BST2 | 0.104 | 0.412 | 0.761 |
| Q5T124 | UBX11 | 0.104 | 0.600 | 0.860 |
| P36871 | PGM1 | 0.103 | 0.647 | 0.872 |
| Q02252 | MMSA | 0.103 | 0.647 | 0.872 |
| Q9Y230 | RUVB2 | 0.102 | 0.501 | 0.796 |
| P22392-2 | NDKB | 0.102 | 0.833 | 0.963 |
| P08833 | IBP1 | 0.102 | 0.609 | 0.860 |
| P38117 | ETFB | 0.101 | 0.166 | 0.504 |
| Q9UHR4 | BI2L1 | 0.100 | 0.628 | 0.872 |
| P11021 | BIP | 0.099 | 0.434 | 0.767 |
| P15374 | UCHL3 | 0.099 | 0.569 | 0.842 |
| P07099 | HYEP | 0.097 | 0.609 | 0.860 |
| P02652 | APOA2 | 0.097 | 0.767 | 0.944 |
| P42765 | THIM | 0.097 | 0.572 | 0.842 |
| Q9H808 | TLE6 | 0.097 | 0.317 | 0.676 |
| Q9UNM6 | PSD13 | 0.096 | 0.767 | 0.944 |
| Q5JRA6 | TGO1 | 0.096 | 0.647 | 0.872 |
| Q93008 | USP9X | 0.095 | 0.851 | 0.963 |
| Q9UHV9 | PFD2 | 0.094 | 0.649 | 0.872 |
| Q9UNE7 | CHIP | 0.094 | 0.687 | 0.900 |
| Q15369 | ELOC | 0.094 | 0.740 | 0.932 |
| Q6ZMN8 | CCNI2 | 0.092 | 0.809 | 0.953 |
| O43681 | GET3 | 0.092 | 0.820 | 0.957 |
| P61020 | RAB5B | 0.092 | 1.000 | 1.000 |
| Q8IW45 | NNRD | 0.091 | 0.651 | 0.872 |
| P09622 | DLDH | 0.089 | 0.467 | 0.778 |
| P06753-2 | TPM3 | 0.089 | 0.434 | 0.767 |
| P09669 | COX6C | 0.088 | 0.767 | 0.944 |
| Q6P1J6 | PLB1 | 0.088 | 0.317 | 0.676 |
| Q9NPA0 | EMC7 | 0.088 | 0.695 | 0.902 |

| | | | | |
|---|---|---|---|---|
| Q8WW33 | GTSF1 | 0.086 | 0.569 | 0.842 |
| P38646 | GRP75 | 0.085 | 0.501 | 0.796 |
| Q9BQ95 | ECSIT | 0.085 | 0.893 | 0.982 |
| P10809 | CH60 | 0.084 | 0.317 | 0.676 |
| P17812 | PYRG1 | 0.084 | 0.928 | 0.989 |
| Q9BTV4 | TMM43 | 0.083 | 0.733 | 0.926 |
| Q9Y281 | COF2 | 0.083 | 0.649 | 0.872 |
| Q13162 | PRDX4 | 0.082 | 0.910 | 0.989 |
| P98160 | PGBM | 0.082 | 0.820 | 0.957 |
| Q8IW52 | SLIK4 | 0.082 | 0.776 | 0.944 |
| Q9NRW1 | RAB6B | 0.081 | 0.851 | 0.963 |
| P30153 | 2AAA | 0.080 | 0.569 | 0.842 |
| P54709 | AT1B3 | 0.080 | 0.879 | 0.982 |
| P46108 | CRK | 0.080 | 0.733 | 0.926 |
| P10599 | THIO | 0.078 | 0.955 | 0.999 |
| P01876 | IGHA1 | 0.077 | 1.000 | 1.000 |
| Q7RTR0 | NLRP9 | 0.076 | 0.536 | 0.819 |
| P36915 | GNL1 | 0.076 | 0.379 | 0.733 |
| Q9NSD9 | SYFB | 0.076 | 0.851 | 0.963 |
| Q8N0W3 | FCSK | 0.075 | 0.733 | 0.926 |
| Q00535 | CDK5 | 0.074 | 0.572 | 0.842 |
| Q9H9P8 | L2HDH | 0.074 | 0.776 | 0.944 |
| Q8WUM4 | PDC6I | 0.073 | 0.434 | 0.767 |
| P35222 | CTNB1 | 0.071 | 0.647 | 0.872 |
| Q9UN37 | VPS4A | 0.071 | 1.000 | 1.000 |
| P33176 | KINH | 0.071 | 0.609 | 0.860 |
| P43007 | SATT | 0.070 | 0.893 | 0.982 |
| Q9ULC6 | PADI1 | 0.070 | 0.467 | 0.778 |
| P04843 | RPN1 | 0.070 | 1.000 | 1.000 |
| Q99611 | SPS2 | 0.070 | 0.893 | 0.982 |
| P55060 | XPO2 | 0.068 | 0.687 | 0.900 |
| Q9Y5M8 | SRPRB | 0.067 | 0.936 | 0.989 |
| P09211 | GSTP1 | 0.067 | 0.865 | 0.971 |
| Q9UHG3 | PCYOX | 0.067 | 0.647 | 0.872 |
| P52292 | IMA1 | 0.066 | 0.449 | 0.778 |
| Q16698 | DECR | 0.066 | 0.572 | 0.842 |
| Q8NAT1 | PMGT2 | 0.066 | 0.936 | 0.989 |
| Q16531 | DDB1 | 0.065 | 0.403 | 0.752 |
| P62820 | RAB1A | 0.065 | 0.687 | 0.900 |
| P13667 | PDIA4 | 0.064 | 0.183 | 0.528 |
| P09110 | THIK | 0.064 | 0.647 | 0.872 |
| Q13867 | BLMH | 0.063 | 0.649 | 0.872 |
| P15586 | GNS | 0.062 | 0.851 | 0.963 |
| O14745 | NHRF1 | 0.062 | 0.851 | 0.963 |
| Q9HDC9 | APMAP | 0.061 | 0.572 | 0.842 |
| P48735 | IDHP | 0.060 | 0.572 | 0.842 |
| P20810 | ICAL | 0.060 | 0.879 | 0.982 |
| Q8NCN5 | PDPR | 0.059 | 0.862 | 0.971 |
| Q9NZL9 | MAT2B | 0.059 | 0.973 | 1.000 |
| Q5JYT7 | K1755 | 0.058 | 0.727 | 0.922 |
| P23284 | PPIB | 0.058 | 0.434 | 0.767 |
| Q15008 | PSMD6 | 0.057 | 1.000 | 1.000 |
| P48444 | COPD | 0.056 | 0.727 | 0.922 |
| O60825-2 | F262 | 0.056 | 0.941 | 0.993 |
| P61224 | RAP1B | 0.056 | 0.979 | 1.000 |
| Q9C0C9 | UBE2O | 0.055 | 0.936 | 0.989 |

| | | | | |
|---|---|---|---|---|
| Q9UM54-2 | MYO6 | 0.055 | 0.893 | 0.982 |
| P51553 | IDH3G | 0.055 | 0.936 | 0.989 |
| P48506 | GSH1 | 0.054 | 0.361 | 0.725 |
| Q92520 | FAM3C | 0.052 | 0.923 | 0.989 |
| Q9UMS0 | NFU1 | 0.052 | 0.649 | 0.872 |
| Q9Y490 | TLN1 | 0.051 | 0.691 | 0.900 |
| Q6IAA8 | LTOR1 | 0.051 | 0.809 | 0.953 |
| P38606 | VATA | 0.050 | 0.851 | 0.963 |
| P13473 | LAMP2 | 0.050 | 0.872 | 0.977 |
| P20936 | RASA1 | 0.050 | 0.608 | 0.860 |
| Q14166 | TTL12 | 0.050 | 0.979 | 1.000 |
| Q9UKK9 | NUDT5 | 0.050 | 0.955 | 0.999 |
| P35232 | PHB1 | 0.049 | 0.727 | 0.922 |
| P68366 | TBA4A | 0.048 | 1.000 | 1.000 |
| O00764 | PDXK | 0.046 | 0.687 | 0.900 |
| P11279 | LAMP1 | 0.046 | 1.000 | 1.000 |
| Q7Z4H3 | HDDC2 | 0.046 | 0.767 | 0.944 |
| Q15102 | PA1B3 | 0.044 | 0.649 | 0.872 |
| P53597 | SUCA | 0.044 | 0.767 | 0.944 |
| Q13126 | MTAP | 0.043 | 0.949 | 0.999 |
| P13798 | ACPH | 0.043 | 0.955 | 0.999 |
| P26639 | SYTC | 0.043 | 0.936 | 0.989 |
| O75746 | S2512 | 0.043 | 0.727 | 0.922 |
| P50990 | TCPQ | 0.043 | 0.979 | 1.000 |
| P55072 | TERA | 0.042 | 0.344 | 0.706 |
| P45880 | VDAC2 | 0.042 | 0.687 | 0.900 |
| P16885 | PLCG2 | 0.042 | 0.647 | 0.872 |
| Q16854 | DGUOK | 0.042 | 0.727 | 0.922 |
| Q8WVM8 | SCFD1 | 0.041 | 0.851 | 0.963 |
| O95336 | 6PGL | 0.041 | 0.809 | 0.953 |
| Q9Y4L1 | HYOU1 | 0.041 | 0.647 | 0.872 |
| Q9UBQ7 | GRHPR | 0.041 | 0.851 | 0.963 |
| O43396 | TXNL1 | 0.040 | 1.000 | 1.000 |
| P14314 | GLU2B | 0.039 | 0.373 | 0.725 |
| Q6PIU2 | NCEH1 | 0.039 | 0.936 | 0.989 |
| P09496-2 | CLCA | 0.039 | 0.809 | 0.953 |
| Q9ULD0 | OGDHL | 0.038 | 0.820 | 0.957 |
| O60664 | PLIN3 | 0.038 | 0.609 | 0.860 |
| Q9BS26 | ERP44 | 0.038 | 0.851 | 0.963 |
| O14980 | XPO1 | 0.037 | 0.786 | 0.952 |
| Q8N283 | ANR35 | 0.037 | 0.879 | 0.982 |
| Q9Y2Z0 | SGT1 | 0.036 | 0.687 | 0.900 |
| P21695 | GPDA | 0.036 | 0.936 | 0.989 |
| P24539 | AT5F1 | 0.036 | 1.000 | 1.000 |
| P37802 | TAGL2 | 0.035 | 1.000 | 1.000 |
| Q14697 | GANAB | 0.035 | 0.609 | 0.860 |
| O43399 | TPD54 | 0.035 | 0.820 | 0.957 |
| P61966 | AP1S1 | 0.034 | 0.865 | 0.971 |
| P51665 | PSMD7 | 0.033 | 0.572 | 0.842 |
| A0AVT1 | UBA6 | 0.033 | 0.501 | 0.796 |
| Q9Y3F4 | STRAP | 0.033 | 0.936 | 0.989 |
| Q5TZA2 | CROCC | 0.033 | 0.910 | 0.989 |
| Q14764 | MVP | 0.032 | 0.687 | 0.900 |
| Q12864 | CAD17 | 0.032 | 0.974 | 1.000 |
| O43707 | ACTN4 | 0.032 | 0.979 | 1.000 |
| Q9Y277 | VDAC3 | 0.030 | 0.687 | 0.900 |

| | | | | |
|---|---|---|---|---|
| P54727 | RD23B | 0.030 | 0.979 | 1.000 |
| O15127 | SCAM2 | 0.030 | 0.652 | 0.872 |
| Q04837 | SSBP | 0.029 | 0.536 | 0.819 |
| P30044-2 | PRDX5 | 0.028 | 1.000 | 1.000 |
| Q9BWD1 | THIC | 0.028 | 0.809 | 0.953 |
| P49593 | PPM1F | 0.026 | 0.373 | 0.725 |
| Q9UKY7 | CDV3 | 0.026 | 0.979 | 1.000 |
| P48643 | TCPE | 0.026 | 0.572 | 0.842 |
| P18206 | VINC | 0.026 | 0.936 | 0.989 |
| P06576 | ATPB | 0.025 | 0.647 | 0.872 |
| Q3ZCQ8 | TIM50 | 0.024 | 0.752 | 0.943 |
| Q5T4I8 | CF052 | 0.023 | 0.776 | 0.944 |
| P16333 | NCK1 | 0.023 | 0.862 | 0.971 |
| P43307 | SSRA | 0.023 | 0.752 | 0.943 |
| P27816 | MAP4 | 0.023 | 0.809 | 0.953 |
| P11498 | PYC | 0.023 | 0.893 | 0.982 |
| P24752 | THIL | 0.022 | 0.851 | 0.963 |
| Q07960 | RHG01 | 0.021 | 0.727 | 0.922 |
| Q9UBS4 | DJB11 | 0.021 | 0.979 | 1.000 |
| P01834 | IGKC | 0.021 | 0.608 | 0.860 |
| P54652 | HSP72 | 0.021 | 0.809 | 0.953 |
| Q9NZZ3 | CHMP5 | 0.020 | 0.910 | 0.989 |
| Q8N668 | COMD1 | 0.020 | 0.955 | 0.999 |
| P23381 | SYWC | 0.020 | 0.910 | 0.989 |
| Q92542 | NICA | 0.020 | 0.796 | 0.953 |
| P43034 | LIS1 | 0.019 | 0.727 | 0.922 |
| P60891 | PRPS1 | 0.018 | 0.832 | 0.963 |
| P14625 | ENPL | 0.018 | 0.851 | 0.963 |
| Q92930 | RAB8B | 0.018 | 0.949 | 0.999 |
| Q5XKP0 | MIC13 | 0.017 | 0.893 | 0.982 |
| P49591 | SYSC | 0.015 | 0.687 | 0.900 |
| Q16543 | CDC37 | 0.015 | 1.000 | 1.000 |
| O15144 | ARPC2 | 0.014 | 0.936 | 0.989 |
| O60749 | SNX2 | 0.014 | 1.000 | 1.000 |
| P00367 | DHE3 | 0.014 | 0.893 | 0.982 |
| Q13011 | ECH1 | 0.013 | 0.893 | 0.982 |
| O95881 | TXD12 | 0.013 | 0.893 | 0.982 |
| Q7KZF4 | SND1 | 0.012 | 0.796 | 0.953 |
| P10645 | CMGA | 0.012 | 0.936 | 0.989 |
| Q6PL24 | TMED8 | 0.012 | 0.776 | 0.944 |
| P11172 | UMPS | 0.010 | 0.767 | 0.944 |
| Q3YEC7 | RABL6 | 0.009 | 0.809 | 0.953 |
| Q9NT62 | ATG3 | 0.009 | 0.691 | 0.900 |
| Q8TF65 | GIPC2 | 0.008 | 0.361 | 0.725 |
| P55209 | NP1L1 | 0.008 | 0.936 | 0.989 |
| O75396 | SC22B | 0.008 | 0.936 | 0.989 |
| P45974 | UBP5 | 0.007 | 0.936 | 0.989 |
| P62333 | PRS10 | 0.006 | 1.000 | 1.000 |
| Q9Y697 | NFS1 | 0.005 | 0.459 | 0.778 |
| Q14204 | DYHC1 | 0.005 | 0.936 | 0.989 |
| P99999 | CYC | 0.005 | 0.936 | 0.989 |
| Q9UJZ1 | STML2 | 0.004 | 0.767 | 0.944 |
| Q9H336 | CRLD1 | 0.004 | 0.979 | 1.000 |
| P50995 | ANX11 | 0.004 | 0.936 | 0.989 |
| P30101 | PDIA3 | 0.003 | 0.936 | 0.989 |
| P17302 | CXA1 | 0.003 | 0.820 | 0.957 |

| | | | | |
|---|---|---|---|---|
| P28074 | PSB5 | 0.002 | 0.979 | 1.000 |
| O15212 | PFD6 | 0.002 | 0.809 | 0.953 |
| P51148 | RAB5C | 0.002 | 0.936 | 0.989 |
| Q15436 | SC23A | 0.002 | 0.851 | 0.963 |
| P47756-2 | CAPZB | 0.001 | 0.373 | 0.725 |
| Q96EY1 | DNJA3 | 0.000 | 0.809 | 0.953 |
| P35270 | SPRE | -0.002 | 0.851 | 0.963 |
| O15143 | ARC1B | -0.002 | 1.000 | 1.000 |
| P04181 | OAT | -0.003 | 0.893 | 0.982 |
| A8MVS5 | HIDE1 | -0.003 | 0.977 | 1.000 |
| P14927 | QCR7 | -0.003 | 0.893 | 0.982 |
| Q8IYB8 | SUV3 | -0.003 | 0.851 | 0.963 |
| P30048 | PRDX3 | -0.004 | 0.936 | 0.989 |
| Q5T5Y3 | CAMP1 | -0.005 | 0.464 | 0.778 |
| Q9BT78 | CSN4 | -0.005 | 1.000 | 1.000 |
| P30519 | HMOX2 | -0.006 | 0.467 | 0.778 |
| O14818 | PSA7 | -0.007 | 0.809 | 0.953 |
| P52788 | SPSY | -0.007 | 1.000 | 1.000 |
| P14384 | CBPM | -0.007 | 0.809 | 0.953 |
| O75390 | CISY | -0.008 | 0.609 | 0.860 |
| Q99962 | SH3G2 | -0.009 | 0.931 | 0.989 |
| P62191 | PRS4 | -0.009 | 0.851 | 0.963 |
| P06756 | ITAV | -0.009 | 0.563 | 0.842 |
| Q9HAV7 | GRPE1 | -0.009 | 0.809 | 0.953 |
| Q9Y224 | RTRAF | -0.010 | 0.809 | 0.953 |
| P16444 | DPEP1 | -0.012 | 0.572 | 0.842 |
| P13861 | KAP2 | -0.012 | 0.893 | 0.982 |
| Q6P996 | PDXD1 | -0.013 | 0.851 | 0.963 |
| P49419-2 | AL7A1 | -0.014 | 0.893 | 0.982 |
| P34913 | HYES | -0.015 | 1.000 | 1.000 |
| P35080-2 | PROF2 | -0.015 | 1.000 | 1.000 |
| Q9Y6G9 | DC1L1 | -0.016 | 0.936 | 0.989 |
| O15511 | ARPC5 | -0.017 | 0.536 | 0.819 |
| P28482 | MK01 | -0.017 | 0.501 | 0.796 |
| P25787 | PSA2 | -0.017 | 0.910 | 0.989 |
| Q16555 | DPYL2 | -0.018 | 0.893 | 0.982 |
| Q14974 | IMB1 | -0.019 | 0.536 | 0.819 |
| Q9UL25 | RAB21 | -0.019 | 0.851 | 0.963 |
| P62495 | ERF1 | -0.021 | 1.000 | 1.000 |
| Q15046 | SYK | -0.022 | 0.651 | 0.872 |
| P05026 | AT1B1 | -0.022 | 0.820 | 0.957 |
| O95292 | VAPB | -0.022 | 0.609 | 0.860 |
| Q02218 | ODO1 | -0.023 | 0.767 | 0.944 |
| P23919 | KTHY | -0.024 | 0.863 | 0.971 |
| P30566 | PUR8 | -0.024 | 0.893 | 0.982 |
| Q9BZE9 | ASPC1 | -0.024 | 0.851 | 0.963 |
| Q9C0E8 | LNP | -0.024 | 1.000 | 1.000 |
| P05089 | ARGI1 | -0.025 | 0.936 | 0.989 |
| P55145 | MANF | -0.026 | 0.979 | 1.000 |
| Q9NSE4 | SYIM | -0.027 | 0.936 | 0.989 |
| P31930 | QCR1 | -0.027 | 0.936 | 0.989 |
| Q92499 | DDX1 | -0.029 | 0.936 | 0.989 |
| Q15121 | PEA15 | -0.030 | 0.776 | 0.944 |
| P15311 | EZRI | -0.030 | 0.403 | 0.752 |
| P11177 | ODPB | -0.030 | 0.893 | 0.982 |
| Q9BXS5 | AP1M1 | -0.031 | 0.936 | 0.989 |

| | | | | |
|---|---|---|---|---|
| Q9NXA8 | SIR5 | -0.031 | 0.809 | 0.953 |
| P54136 | SYRC | -0.031 | 0.955 | 0.999 |
| Q9Y4W6 | AFG32 | -0.032 | 0.936 | 0.989 |
| P13804 | ETFA | -0.033 | 0.979 | 1.000 |
| Q99417 | MYCBP | -0.034 | 1.000 | 1.000 |
| P61960 | UFM1 | -0.034 | 0.628 | 0.872 |
| Q9GZT3 | SLIRP | -0.035 | 0.851 | 0.963 |
| Q9BX68 | HINT2 | -0.036 | 0.727 | 0.922 |
| Q969H8 | MYDGF | -0.036 | 0.851 | 0.963 |
| P56385 | ATP5I | -0.036 | 0.767 | 0.944 |
| Q16891 | MIC60 | -0.036 | 0.979 | 1.000 |
| Q00688 | FKBP3 | -0.036 | 0.955 | 0.999 |
| Q99798 | ACON | -0.036 | 0.727 | 0.922 |
| P07203 | GPX1 | -0.038 | 0.851 | 0.963 |
| O95671 | ASML | -0.038 | 0.291 | 0.657 |
| Q9HB71 | CYBP | -0.039 | 0.687 | 0.900 |
| Q02790 | FKBP4 | -0.039 | 0.291 | 0.657 |
| Q9H2J4 | PDCL3 | -0.039 | 0.647 | 0.872 |
| O43488 | ARK72 | -0.040 | 0.936 | 0.989 |
| E9PAV3 | NACAM | -0.041 | 0.536 | 0.819 |
| Q9Y4Z0 | LSM4 | -0.042 | 0.494 | 0.796 |
| Q9H061 | T126A | -0.043 | 0.851 | 0.963 |
| Q07021 | C1QBP | -0.043 | 0.979 | 1.000 |
| Q587J8 | KHDC3 | -0.043 | 0.647 | 0.872 |
| Q6FI81 | CPIN1 | -0.043 | 1.000 | 1.000 |
| P60981 | DEST | -0.045 | 0.910 | 0.989 |
| P43897 | EFTS | -0.045 | 0.809 | 0.953 |
| P25705 | ATPA | -0.045 | 0.647 | 0.872 |
| A6NGQ2 | OOEP | -0.046 | 0.893 | 0.982 |
| Q96AQ6 | PBIP1 | -0.047 | 0.979 | 1.000 |
| Q9Y305 | ACOT9 | -0.047 | 0.767 | 0.944 |
| P13987 | CD59 | -0.048 | 0.936 | 0.989 |
| Q9NWU5 | RM22 | -0.048 | 0.609 | 0.860 |
| Q9UFN0 | NPS3A | -0.049 | 0.851 | 0.963 |
| P36776 | LONM | -0.049 | 0.434 | 0.767 |
| Q9GZZ1 | NAA50 | -0.050 | 0.522 | 0.819 |
| Q8IZ83 | A16A1 | -0.050 | 0.976 | 1.000 |
| P04899 | GNAI2 | -0.051 | 1.000 | 1.000 |
| Q9NP79 | VTA1 | -0.053 | 1.000 | 1.000 |
| O00410 | IPO5 | -0.053 | 1.000 | 1.000 |
| O00231 | PSD11 | -0.053 | 0.809 | 0.953 |
| P14406 | CX7A2 | -0.054 | 0.809 | 0.953 |
| P29401 | TKT | -0.055 | 0.979 | 1.000 |
| Q13217 | DNJC3 | -0.056 | 0.893 | 0.982 |
| Q9Y266 | NUDC | -0.056 | 0.531 | 0.819 |
| P49411 | EFTU | -0.056 | 0.727 | 0.922 |
| P17858 | PFKAL | -0.057 | 0.344 | 0.706 |
| P20674 | COX5A | -0.058 | 0.572 | 0.842 |
| Q9BTW9 | TBCD | -0.059 | 0.647 | 0.872 |
| P22314 | UBA1 | -0.059 | 0.609 | 0.860 |
| P34897 | GLYM | -0.060 | 0.809 | 0.953 |
| Q9BX40 | LS14B | -0.060 | 0.647 | 0.872 |
| O75208 | COQ9 | -0.060 | 0.647 | 0.872 |
| O60763 | USO1 | -0.061 | 0.979 | 1.000 |
| O95394 | AGM1 | -0.062 | 0.910 | 0.989 |
| Q03426 | KIME | -0.063 | 0.979 | 1.000 |

| | | | | |
|---|---|---|---|---|
| Q9BYX2 | TBD2A | -0.064 | 0.412 | 0.761 |
| P59190 | RAB15 | -0.066 | 0.910 | 0.989 |
| Q96J94 | PIWL1 | -0.067 | 0.820 | 0.957 |
| Q15814 | TBCC | -0.068 | 0.893 | 0.982 |
| Q9Y6C9 | MTCH2 | -0.069 | 1.000 | 1.000 |
| P62244 | RS15A | -0.069 | 0.896 | 0.984 |
| Q7Z4G1 | COMD6 | -0.069 | 0.809 | 0.953 |
| Q8NFF5 | FAD1 | -0.070 | 1.000 | 1.000 |
| P28331 | NDUS1 | -0.071 | 0.727 | 0.922 |
| Q9UHF5 | IL17B | -0.071 | 0.647 | 0.872 |
| P41250 | GARS | -0.074 | 0.767 | 0.944 |
| Q96QK1 | VPS35 | -0.074 | 0.267 | 0.623 |
| Q9BRX8 | PXL2A | -0.075 | 0.979 | 1.000 |
| Q15365 | PCBP1 | -0.075 | 0.865 | 0.971 |
| P56192 | SYMC | -0.077 | 0.647 | 0.872 |
| P55011 | S12A2 | -0.078 | 0.536 | 0.819 |
| P47914 | RL29 | -0.078 | 0.687 | 0.900 |
| O00534 | VMA5A | -0.079 | 0.767 | 0.944 |
| P50213 | IDH3A | -0.080 | 0.373 | 0.725 |
| P14550 | AK1A1 | -0.080 | 0.501 | 0.796 |
| P61247 | RS3A | -0.081 | 0.373 | 0.725 |
| Q9GZP4 | PITH1 | -0.082 | 0.647 | 0.872 |
| P53602 | MVD1 | -0.082 | 0.918 | 0.989 |
| Q9NQT8 | KI13B | -0.083 | 0.501 | 0.796 |
| O00571 | DDX3X | -0.083 | 0.403 | 0.752 |
| Q9BPU6 | DPYL5 | -0.084 | 0.767 | 0.944 |
| Q9Y2Q3 | GSTK1 | -0.084 | 0.809 | 0.953 |
| Q96C10 | DHX58 | -0.084 | 0.776 | 0.944 |
| O43837 | IDH3B | -0.085 | 0.222 | 0.567 |
| Q9UHK6 | AMACR | -0.085 | 0.647 | 0.872 |
| Q99674 | CGRE1 | -0.085 | 0.776 | 0.944 |
| P35998 | PRS7 | -0.085 | 0.536 | 0.819 |
| P36955 | PEDF | -0.086 | 0.494 | 0.796 |
| P49748 | ACADV | -0.086 | 0.687 | 0.900 |
| Q9BWS9 | CHID1 | -0.087 | 0.687 | 0.900 |
| P05141 | ADT2 | -0.089 | 0.536 | 0.819 |
| P02792 | FRIL | -0.090 | 0.809 | 0.953 |
| Q7L0Y3 | TM10C | -0.090 | 0.786 | 0.952 |
| O75489 | NDUS3 | -0.091 | 0.687 | 0.900 |
| P54920 | SNAA | -0.092 | 0.373 | 0.725 |
| O15347 | HMGB3 | -0.093 | 0.649 | 0.872 |
| Q9Y3I0 | RTCB | -0.093 | 0.244 | 0.597 |
| Q96KA5 | CLP1L | -0.093 | 0.605 | 0.860 |
| P12081 | HARS1 | -0.094 | 0.851 | 0.963 |
| P25398 | RS12 | -0.095 | 0.893 | 0.982 |
| P63208 | SKP1 | -0.095 | 0.434 | 0.767 |
| P55084 | ECHB | -0.096 | 0.434 | 0.767 |
| Q9NR28 | DBLOH | -0.097 | 0.536 | 0.819 |
| O60826 | CCD22 | -0.099 | 0.531 | 0.819 |
| P46776 | RL27A | -0.099 | 0.767 | 0.944 |
| P22570 | ADRO | -0.100 | 0.434 | 0.767 |
| P01009 | A1AT | -0.102 | 0.501 | 0.796 |
| P27144 | KAD4 | -0.102 | 0.936 | 0.989 |
| P40616 | ARL1 | -0.105 | 0.413 | 0.761 |
| Q9UII2 | ATIF1 | -0.105 | 0.434 | 0.767 |
| P61970 | NTF2 | -0.105 | 0.531 | 0.819 |

| | | | | |
|---|---|---|---|---|
| Q16718 | NDUA5 | -0.106 | 0.501 | 0.796 |
| Q9P0L0 | VAPA | -0.108 | 0.572 | 0.842 |
| Q6P1N9 | TATD1 | -0.108 | 1.000 | 1.000 |
| Q8N1F7 | NUP93 | -0.111 | 0.786 | 0.952 |
| P19623 | SPEE | -0.111 | 0.467 | 0.778 |
| P07814 | SYEP | -0.112 | 0.467 | 0.778 |
| P49721 | PSB2 | -0.112 | 0.709 | 0.917 |
| P35244 | RFA3 | -0.112 | 0.494 | 0.796 |
| P28072 | PSB6 | -0.112 | 0.691 | 0.900 |
| P27824 | CALX | -0.113 | 0.536 | 0.819 |
| Q86UP2 | KTN1 | -0.113 | 0.467 | 0.778 |
| Q5ST30 | SYVM | -0.113 | 0.687 | 0.900 |
| P62195 | PRS8 | -0.114 | 0.536 | 0.819 |
| P46940 | IQGA1 | -0.114 | 0.605 | 0.860 |
| P07108-4 | ACBP | -0.114 | 0.649 | 0.872 |
| O60879 | DIAP2 | -0.115 | 0.501 | 0.796 |
| P00387 | NB5R3 | -0.115 | 0.893 | 0.982 |
| Q9P2R7 | SUCB1 | -0.115 | 0.373 | 0.725 |
| Q9ULR5 | PAI2B | -0.117 | 1.000 | 1.000 |
| P49589-3 | SYCC | -0.118 | 0.609 | 0.860 |
| P14618-2 | KPYM | -0.118 | 0.403 | 0.752 |
| Q9NP72 | RAB18 | -0.119 | 0.536 | 0.819 |
| P20618 | PSB1 | -0.120 | 0.149 | 0.478 |
| Q6NVY1 | HIBCH | -0.120 | 0.244 | 0.597 |
| P49755 | TMEDA | -0.121 | 0.647 | 0.872 |
| Q06323 | PSME1 | -0.121 | 0.948 | 0.999 |
| Q9Y371 | SHLB1 | -0.121 | 0.936 | 0.989 |
| P61158 | ARP3 | -0.122 | 0.501 | 0.796 |
| P30533 | AMRP | -0.122 | 0.501 | 0.796 |
| A0A1W2PR48 | TLE7 | -0.122 | 0.392 | 0.749 |
| O43615 | TIM44 | -0.124 | 0.467 | 0.778 |
| P50991 | TCPD | -0.124 | 0.202 | 0.545 |
| Q9Y265 | RUVB1 | -0.124 | 0.501 | 0.796 |
| P08559 | ODPA | -0.124 | 0.202 | 0.545 |
| Q13615 | MTMR3 | -0.125 | 0.317 | 0.676 |
| P54819 | KAD2 | -0.126 | 0.403 | 0.752 |
| Q9P032 | NDUF4 | -0.126 | 0.674 | 0.899 |
| P10606 | COX5B | -0.126 | 0.222 | 0.567 |
| P27708 | PYR1 | -0.127 | 0.609 | 0.860 |
| P17980 | PRS6A | -0.127 | 0.095 | 0.403 |
| Q9NR45 | SIAS | -0.127 | 0.467 | 0.778 |
| P84085 | ARF5 | -0.129 | 0.820 | 0.957 |
| Q12933 | TRAF2 | -0.129 | 0.606 | 0.860 |
| Q96TC7 | RMD3 | -0.130 | 0.727 | 0.922 |
| Q9H773 | DCTP1 | -0.130 | 0.898 | 0.984 |
| P46782 | RS5 | -0.131 | 0.651 | 0.872 |
| Q9UBV2 | SE1L1 | -0.132 | 0.267 | 0.623 |
| Q8TBX8 | PI42C | -0.133 | 0.166 | 0.504 |
| Q5VWJ9 | SNX30 | -0.135 | 0.472 | 0.784 |
| P53396 | ACLY | -0.135 | 0.865 | 0.971 |
| Q86VP6 | CAND1 | -0.135 | 0.361 | 0.725 |
| P06748 | NPM | -0.136 | 0.647 | 0.872 |
| Q9Y5L4 | TIM13 | -0.137 | 0.424 | 0.767 |
| P60660 | MYL6 | -0.137 | 0.403 | 0.752 |
| P22033 | MUTA | -0.138 | 0.767 | 0.944 |
| P09972 | ALDOC | -0.139 | 0.051 | 0.292 |

| | | | | |
|---|---|---|---|---|
| Q7Z4W1 | DCXR | -0.144 | 0.647 | 0.872 |
| P10515 | ODP2 | -0.145 | 0.403 | 0.752 |
| Q8N0X4 | CLYBL | -0.145 | 0.536 | 0.819 |
| P10253 | LYAG | -0.145 | 0.647 | 0.872 |
| P49821 | NDUV1 | -0.146 | 0.609 | 0.860 |
| Q15165 | PON2 | -0.147 | 0.647 | 0.872 |
| O14964 | HGS | -0.147 | 0.446 | 0.778 |
| O43776 | SYNC | -0.147 | 0.936 | 0.989 |
| Q9HCN8 | SDF2L | -0.150 | 0.344 | 0.706 |
| Q86VE9 | SERC5 | -0.150 | 0.808 | 0.953 |
| O60313-2 | OPA1 | -0.151 | 0.095 | 0.403 |
| O75832 | PSD10 | -0.151 | 0.767 | 0.944 |
| P31939 | PUR9 | -0.152 | 0.459 | 0.778 |
| P51572 | BAP31 | -0.152 | 0.344 | 0.706 |
| P31689 | DNJA1 | -0.153 | 0.536 | 0.819 |
| P51159 | RB27A | -0.153 | 0.536 | 0.819 |
| Q9NR31 | SAR1A | -0.155 | 0.536 | 0.819 |
| P84095 | RHOG | -0.155 | 0.602 | 0.860 |
| P15170-2 | ERF3A | -0.155 | 0.608 | 0.860 |
| Q6YN16 | HSDL2 | -0.156 | 0.235 | 0.590 |
| Q99614 | TTC1 | -0.156 | 0.403 | 0.752 |
| O95182 | NDUA7 | -0.156 | 0.605 | 0.860 |
| P31040 | SDHA | -0.158 | 0.373 | 0.725 |
| Q00765 | REEP5 | -0.158 | 0.569 | 0.842 |
| Q13492 | PICAL | -0.160 | 0.478 | 0.792 |
| P02794 | FRIH | -0.163 | 0.809 | 0.953 |
| A6NMX2 | I4E1B | -0.163 | 0.403 | 0.752 |
| P60900 | PSA6 | -0.163 | 0.494 | 0.796 |
| P48556 | PSMD8 | -0.164 | 0.693 | 0.901 |
| Q15435 | PP1R7 | -0.164 | 0.776 | 0.944 |
| O95081 | AGFG2 | -0.164 | 0.566 | 0.842 |
| Q96T17-2 | MA7D2 | -0.165 | 0.609 | 0.860 |
| P08195 | 4F2 | -0.166 | 0.202 | 0.545 |
| Q9NUQ7 | UFSP2 | -0.167 | 0.403 | 0.752 |
| P16671 | CD36 | -0.168 | 0.744 | 0.936 |
| P30405 | PPIF | -0.168 | 0.267 | 0.623 |
| O60506 | HNRPQ | -0.169 | 0.705 | 0.913 |
| O14791 | APOL1 | -0.171 | 0.824 | 0.960 |
| Q7Z2Z2 | EFL1 | -0.172 | 0.228 | 0.579 |
| P41208 | CETN2 | -0.173 | 0.833 | 0.963 |
| Q9NP58 | ABCB6 | -0.174 | 0.018 | 0.182 |
| Q16795 | NDUA9 | -0.175 | 0.647 | 0.872 |
| A0A1B0GUC4 | MYCOS | -0.175 | 0.820 | 0.957 |
| P04075 | ALDOA | -0.176 | 0.008 | 0.115 |
| Q04446 | GLGB | -0.176 | 0.150 | 0.478 |
| P0C869-4 | PA24B | -0.176 | 0.872 | 0.977 |
| O43795 | MYO1B | -0.177 | 0.392 | 0.749 |
| O43169 | CYB5B | -0.179 | 0.733 | 0.926 |
| O75534 | CSDE1 | -0.179 | 0.508 | 0.806 |
| P01024 | CO3 | -0.180 | 0.693 | 0.901 |
| P52907 | CAZA1 | -0.180 | 0.252 | 0.609 |
| Q16836 | HCDH | -0.180 | 0.202 | 0.545 |
| Q9HAB8 | PPCS | -0.183 | 0.691 | 0.900 |
| Q15286 | RAB35 | -0.185 | 0.422 | 0.767 |
| Q9BXU1 | STK31 | -0.185 | 0.202 | 0.545 |
| Q9GZT6 | CC90B | -0.185 | 0.267 | 0.623 |

| | | | | |
|---|---|---|---|---|
| P55884 | EIF3B | -0.186 | 0.600 | 0.860 |
| Q9UK45 | LSM7 | -0.186 | 0.424 | 0.767 |
| Q9UI30 | TR112 | -0.188 | 0.332 | 0.700 |
| P26373 | RL13 | -0.188 | 0.501 | 0.796 |
| Q9Y512 | SAM50 | -0.189 | 0.727 | 0.922 |
| Q9BU23 | LMF2 | -0.189 | 0.605 | 0.860 |
| Q6DD88 | ATLA3 | -0.191 | 0.536 | 0.819 |
| Q10567 | AP1B1 | -0.193 | 0.317 | 0.676 |
| Q9P0B6 | CC167 | -0.195 | 0.501 | 0.796 |
| Q9H2U2 | IPYR2 | -0.195 | 0.082 | 0.376 |
| P30084 | ECHM | -0.197 | 0.267 | 0.623 |
| Q92506 | DHB8 | -0.197 | 0.786 | 0.952 |
| P27348 | 1433T | -0.198 | 0.955 | 0.999 |
| Q15398-3 | DLGP5 | -0.199 | 0.202 | 0.545 |
| O75165 | DJC13 | -0.200 | 0.013 | 0.152 |
| P08754 | GNAI3 | -0.201 | 0.361 | 0.725 |
| Q13438 | OS9 | -0.203 | 0.467 | 0.778 |
| Q9NQR4 | NIT2 | -0.203 | 0.649 | 0.872 |
| O94919 | ENDD1 | -0.204 | 0.609 | 0.860 |
| Q9NVI7-2 | ATD3A | -0.205 | 0.501 | 0.796 |
| Q9Y696 | CLIC4 | -0.206 | 0.392 | 0.749 |
| O43447 | PPIH | -0.207 | 0.424 | 0.767 |
| P46459 | NSF | -0.208 | 0.107 | 0.424 |
| P30085 | KCY | -0.209 | 0.134 | 0.459 |
| Q9NUJ1 | ABHDA | -0.210 | 0.134 | 0.459 |
| Q99623 | PHB2 | -0.214 | 0.134 | 0.459 |
| P05362 | ICAM1 | -0.214 | 0.373 | 0.725 |
| P54098 | DPOG1 | -0.214 | 0.572 | 0.842 |
| Q99766 | ATP5S | -0.215 | 0.566 | 0.842 |
| Q5VV41 | ARHGG | -0.216 | 0.602 | 0.860 |
| P06733 | ENOA | -0.217 | 0.501 | 0.796 |
| Q5SRE7 | PHYD1 | -0.217 | 0.208 | 0.556 |
| Q92538 | GBF1 | -0.218 | 0.413 | 0.761 |
| Q92598 | HS105 | -0.218 | 0.044 | 0.273 |
| P46379 | BAG6 | -0.221 | 0.183 | 0.528 |
| P23634 | AT2B4 | -0.222 | 0.413 | 0.761 |
| Q4VXU2 | PAP1L | -0.222 | 0.183 | 0.528 |
| Q12904 | AIMP1 | -0.222 | 0.727 | 0.922 |
| Q8IVD9 | NUDC3 | -0.222 | 0.344 | 0.706 |
| Q99747 | SNAG | -0.223 | 0.107 | 0.424 |
| Q02878 | RL6 | -0.223 | 0.467 | 0.778 |
| Q8IXB1 | DJC10 | -0.223 | 0.651 | 0.872 |
| O15145 | ARPC3 | -0.223 | 0.244 | 0.597 |
| P15927-3 | RFA2 | -0.223 | 0.316 | 0.676 |
| Q9UQ13 | SHOC2 | -0.224 | 0.291 | 0.657 |
| P08133 | ANXA6 | -0.224 | 0.107 | 0.424 |
| Q96BN8 | OTUL | -0.226 | 0.459 | 0.778 |
| Q709C8 | VP13C | -0.226 | 0.784 | 0.952 |
| P00403 | COX2 | -0.228 | 0.647 | 0.872 |
| Q9BVK6 | TMED9 | -0.229 | 0.344 | 0.706 |
| P43487 | RANG | -0.230 | 0.379 | 0.733 |
| P55010 | IF5 | -0.231 | 0.456 | 0.778 |
| Q9NZ01 | TECR | -0.231 | 0.403 | 0.752 |
| Q96RP9 | EFGM | -0.231 | 0.687 | 0.900 |
| Q9UK22 | FBX2 | -0.232 | 0.222 | 0.567 |
| Q9NRY5 | F1142 | -0.233 | 0.331 | 0.700 |

| | | | | |
|---|---|---|---|---|
| O15371 | EIF3D | -0.235 | 0.691 | 0.900 |
| P36404 | ARL2 | -0.235 | 0.075 | 0.350 |
| O43405 | COCH | -0.236 | 0.149 | 0.478 |
| O00483 | NDUA4 | -0.236 | 0.562 | 0.842 |
| O75822 | EIF3J | -0.237 | 0.459 | 0.778 |
| Q12931 | TRAP1 | -0.237 | 0.044 | 0.273 |
| O43491-4 | E41L2 | -0.238 | 0.833 | 0.963 |
| O00487 | PSDE | -0.239 | 0.018 | 0.182 |
| P55036 | PSMD4 | -0.240 | 0.501 | 0.796 |
| Q9NRA8 | 4ET | -0.240 | 0.025 | 0.208 |
| P22830 | HEMH | -0.241 | 0.467 | 0.778 |
| P42704 | LPPRC | -0.242 | 0.183 | 0.528 |
| P27694 | RFA1 | -0.242 | 0.572 | 0.842 |
| Q06203 | PUR1 | -0.243 | 0.202 | 0.545 |
| Q13616 | CUL1 | -0.245 | 0.317 | 0.676 |
| P35914 | HMGCL | -0.247 | 0.029 | 0.220 |
| Q99460 | PSMD1 | -0.248 | 0.222 | 0.567 |
| P26196 | DDX6 | -0.248 | 0.033 | 0.239 |
| Q96LD8 | SENP8 | -0.249 | 0.464 | 0.778 |
| P13010 | XRCC5 | -0.250 | 0.381 | 0.735 |
| P61225 | RAP2B | -0.251 | 0.270 | 0.630 |
| P33316 | DUT | -0.251 | 0.029 | 0.220 |
| Q9H1K1 | ISCU | -0.252 | 0.373 | 0.725 |
| P05388 | RLA0 | -0.256 | 0.376 | 0.728 |
| Q9BPX5 | ARP5L | -0.258 | 0.845 | 0.963 |
| O95972 | BMP15 | -0.259 | 0.501 | 0.796 |
| P02751 | FINC | -0.260 | 0.044 | 0.273 |
| Q13263 | TIF1B | -0.260 | 0.531 | 0.819 |
| P40939 | ECHA | -0.262 | 0.018 | 0.181 |
| Q9H0U3 | MAGT1 | -0.262 | 0.186 | 0.535 |
| Q4G0N4 | NAKD2 | -0.263 | 0.095 | 0.403 |
| P78344 | IF4G2 | -0.267 | 0.609 | 0.860 |
| Q9H910 | JUPI2 | -0.267 | 0.134 | 0.459 |
| P13073 | COX41 | -0.268 | 0.095 | 0.403 |
| Q6ZMR3 | LDH6A | -0.268 | 0.058 | 0.315 |
| Q00325 | MPCP | -0.268 | 0.149 | 0.478 |
| P21399 | ACOHC | -0.268 | 0.202 | 0.545 |
| P32969 | RL9 | -0.269 | 0.202 | 0.545 |
| P13489 | RINI | -0.269 | 0.373 | 0.725 |
| Q01518 | CAP1 | -0.269 | 0.051 | 0.292 |
| Q92947 | GCDH | -0.270 | 0.776 | 0.944 |
| P25786 | PSA1 | -0.272 | 0.317 | 0.676 |
| Q15181 | IPYR | -0.273 | 0.075 | 0.350 |
| P10644 | KAP0 | -0.273 | 0.095 | 0.403 |
| Q9GZZ9 | UBA5 | -0.274 | 0.449 | 0.778 |
| P62847 | RS24 | -0.275 | 0.202 | 0.545 |
| Q14240 | IF4A2 | -0.279 | 0.244 | 0.597 |
| P39019 | RS19 | -0.280 | 0.038 | 0.255 |
| Q9NPE2 | NGRN | -0.282 | 0.608 | 0.860 |
| Q13200 | PSMD2 | -0.282 | 0.021 | 0.191 |
| P18124 | RL7 | -0.285 | 0.244 | 0.597 |
| Q96A49 | SYAP1 | -0.287 | 0.979 | 1.000 |
| Q9H0U4 | RAB1B | -0.287 | 0.134 | 0.459 |
| A6ND01 | JUNO | -0.288 | 0.344 | 0.706 |
| Q9NWT8 | AKIP | -0.289 | 0.456 | 0.778 |
| P50914 | RL14 | -0.291 | 0.025 | 0.208 |

| | | | | |
|---|---|---|---|---|
| P19404 | NDUV2 | -0.291 | 0.244 | 0.597 |
| O96000 | NDUBA | -0.292 | 0.166 | 0.504 |
| Q12765 | SCRN1 | -0.293 | 0.776 | 0.944 |
| Q14247 | SRC8 | -0.294 | 1.000 | 1.000 |
| P46778 | RL21 | -0.294 | 0.149 | 0.478 |
| P42126 | ECI1 | -0.296 | 0.183 | 0.528 |
| P31942 | HNRH3 | -0.297 | 0.434 | 0.767 |
| P26640 | SYVC | -0.297 | 0.776 | 0.944 |
| Q9H0E2 | TOLIP | -0.297 | 0.467 | 0.778 |
| Q13907 | IDI1 | -0.298 | 0.347 | 0.706 |
| P26440 | IVD | -0.299 | 0.317 | 0.676 |
| Q9BQP7 | MGME1 | -0.301 | 0.477 | 0.791 |
| Q7RTY7 | OVCH1 | -0.301 | 0.252 | 0.609 |
| Q719I0 | AHSA2 | -0.301 | 0.207 | 0.554 |
| Q66K74 | MAP1S | -0.301 | 0.085 | 0.379 |
| P61421 | VA0D1 | -0.302 | 0.072 | 0.350 |
| P62330 | ARF6 | -0.303 | 0.183 | 0.528 |
| Q9Y3D6 | FIS1 | -0.306 | 0.424 | 0.767 |
| Q5JRX3 | PREP | -0.307 | 0.166 | 0.504 |
| P08708 | RS17 | -0.307 | 0.349 | 0.709 |
| Q13813 | SPTN1 | -0.310 | 0.324 | 0.689 |
| P60842 | IF4A1 | -0.311 | 0.038 | 0.255 |
| O15382 | BCAT2 | -0.311 | 0.244 | 0.597 |
| Q04917 | 1433F | -0.314 | 0.120 | 0.444 |
| O75964 | ATP5L | -0.314 | 0.051 | 0.292 |
| Q3LXA3 | TKFC | -0.315 | 0.093 | 0.403 |
| P53007 | TXTP | -0.316 | 0.183 | 0.528 |
| Q15370 | ELOB | -0.316 | 0.563 | 0.842 |
| Q9BRG1 | VPS25 | -0.317 | 0.413 | 0.761 |
| P11216 | PYGB | -0.320 | 0.267 | 0.623 |
| P51659 | DHB4 | -0.320 | 0.029 | 0.220 |
| P60866 | RS20 | -0.322 | 0.134 | 0.459 |
| O96008 | TOM40 | -0.322 | 0.382 | 0.737 |
| O60547 | GMDS | -0.324 | 0.038 | 0.255 |
| O75369 | FLNB | -0.324 | 0.051 | 0.292 |
| P49720 | PSB3 | -0.325 | 0.314 | 0.676 |
| O00186 | STXB3 | -0.325 | 0.232 | 0.585 |
| Q9UHN1 | DPOG2 | -0.328 | 0.424 | 0.767 |
| Q8N9I0 | SYT2 | -0.329 | 0.202 | 0.545 |
| P11586 | C1TC | -0.329 | 0.051 | 0.292 |
| Q14019 | COTL1 | -0.331 | 0.134 | 0.459 |
| Q9NVA2 | 40787 | -0.332 | 0.312 | 0.676 |
| Q99536 | VAT1 | -0.334 | 0.149 | 0.478 |
| Q8WUY1 | THEM6 | -0.335 | 0.609 | 0.860 |
| Q6UXV4 | MIC27 | -0.335 | 0.539 | 0.821 |
| P27635 | RL10 | -0.337 | 0.243 | 0.597 |
| Q9Y5X3 | SNX5 | -0.338 | 0.150 | 0.478 |
| P05387 | RLA2 | -0.338 | 0.095 | 0.403 |
| O95219 | SNX4 | -0.338 | 0.149 | 0.478 |
| P49207 | RL34 | -0.341 | 0.202 | 0.545 |
| O14908 | GIPC1 | -0.343 | 0.232 | 0.585 |
| Q8N4H5 | TOM5 | -0.346 | 0.134 | 0.459 |
| Q9NWS8-2 | RMND1 | -0.348 | 0.456 | 0.778 |
| P35241 | RADI | -0.350 | 0.029 | 0.220 |
| P12236 | ADT3 | -0.350 | 0.267 | 0.623 |
| Q9UJ70-2 | NAGK | -0.352 | 0.207 | 0.554 |

| | | | | |
|---|---|---|---|---|
| Q04637 | IF4G1 | -0.354 | 0.120 | 0.444 |
| Q15717 | ELAV1 | -0.355 | 0.519 | 0.819 |
| Q3MIX3 | ADCK5 | -0.355 | 0.695 | 0.902 |
| Q96CW1 | AP2M1 | -0.358 | 0.300 | 0.672 |
| P36551 | HEM6 | -0.360 | 0.075 | 0.350 |
| Q9Y303 | NAGA | -0.360 | 0.114 | 0.444 |
| P31946 | 1433B | -0.362 | 0.166 | 0.504 |
| Q7Z3Z3 | PIWL3 | -0.366 | 0.192 | 0.545 |
| Q86TP1 | PRUN1 | -0.366 | 0.376 | 0.728 |
| O60383 | GDF9 | -0.368 | 0.095 | 0.403 |
| Q9H173 | SIL1 | -0.369 | 0.085 | 0.379 |
| Q9UGI8 | TES | -0.372 | 0.464 | 0.778 |
| P07093-2 | GDN | -0.372 | 0.222 | 0.567 |
| P47755 | CAZA2 | -0.374 | 0.107 | 0.424 |
| Q86VR2 | RETR3 | -0.374 | 0.699 | 0.907 |
| O60825 | F262 | -0.376 | 0.051 | 0.292 |
| O00217 | NDUS8 | -0.379 | 0.344 | 0.706 |
| Q8N2K0 | ABD12 | -0.379 | 0.095 | 0.403 |
| Q13618 | CUL3 | -0.379 | 0.166 | 0.504 |
| Q9P2B4 | CT2NL | -0.380 | 0.347 | 0.706 |
| Q9HC35 | EMAL4 | -0.382 | 0.134 | 0.459 |
| P02647 | APOA1 | -0.385 | 0.222 | 0.567 |
| P46779 | RL28 | -0.387 | 0.410 | 0.761 |
| Q9Y450 | HBS1L | -0.387 | 0.331 | 0.700 |
| Q14558 | KPRA | -0.389 | 0.373 | 0.725 |
| P80303 | NUCB2 | -0.389 | 0.166 | 0.504 |
| P46777 | RL5 | -0.399 | 0.051 | 0.292 |
| Q9UNK0 | STX8 | -0.401 | 0.413 | 0.761 |
| Q9H4M9 | EHD1 | -0.401 | 0.471 | 0.783 |
| P35579 | MYH9 | -0.405 | 0.001 | 0.046 |
| Q9UK41 | VPS28 | -0.409 | 0.941 | 0.993 |
| P51114 | FXR1 | -0.411 | 0.166 | 0.504 |
| Q709F0 | ACD11 | -0.411 | 0.021 | 0.191 |
| Q99733 | NP1L4 | -0.411 | 0.038 | 0.255 |
| Q99598 | TSNAX | -0.412 | 0.134 | 0.459 |
| Q9UN86 | G3BP2 | -0.414 | 0.167 | 0.505 |
| O43491 | E41L2 | -0.418 | 0.051 | 0.292 |
| Q8NFU3 | TSTD1 | -0.422 | 0.066 | 0.335 |
| P62280 | RS11 | -0.423 | 0.051 | 0.292 |
| Q86W24 | NAL14 | -0.425 | 0.021 | 0.191 |
| Q9H3N1 | TMX1 | -0.427 | 0.331 | 0.700 |
| P62899 | RL31 | -0.432 | 0.343 | 0.706 |
| P23396 | RS3 | -0.436 | 0.008 | 0.115 |
| P62753 | RS6 | -0.437 | 0.536 | 0.819 |
| Q9UMQ6 | CAN11 | -0.443 | 0.292 | 0.657 |
| P62249 | RS16 | -0.448 | 0.025 | 0.208 |
| Q8TD30 | ALAT2 | -0.450 | 0.038 | 0.255 |
| P11310 | ACADM | -0.453 | 0.058 | 0.315 |
| O76024 | WFS1 | -0.454 | 0.051 | 0.292 |
| O75251 | NDUS7 | -0.456 | 0.088 | 0.394 |
| Q16181 | 39326 | -0.458 | 0.361 | 0.725 |
| O95202 | LETM1 | -0.458 | 0.291 | 0.657 |
| P36578 | RL4 | -0.461 | 0.011 | 0.136 |
| Q96A35 | RM24 | -0.461 | 0.051 | 0.292 |
| O14578-3 | CTRO | -0.461 | 0.107 | 0.424 |
| P68402 | PA1B2 | -0.463 | 0.288 | 0.657 |

| | | | | |
|---|---|---|---|---|
| P62424 | RL7A | -0.465 | 0.013 | 0.152 |
| Q9BZF1 | OSBL8 | -0.465 | 0.123 | 0.452 |
| Q99829 | CPNE1 | -0.469 | 0.317 | 0.676 |
| O75323 | NIPS2 | -0.473 | 0.539 | 0.821 |
| P62851 | RS25 | -0.476 | 0.038 | 0.255 |
| Q09028 | RBBP4 | -0.478 | 0.740 | 0.932 |
| Q15813 | TBCE | -0.480 | 0.193 | 0.545 |
| Q9NRG9 | AAAS | -0.482 | 0.166 | 0.504 |
| Q9NWV4 | CZIB | -0.482 | 0.075 | 0.350 |
| P62913 | RL11 | -0.482 | 0.021 | 0.191 |
| P51531 | SMCA2 | -0.483 | 0.277 | 0.642 |
| Q14847 | LASP1 | -0.484 | 0.122 | 0.450 |
| O75436 | VP26A | -0.485 | 0.107 | 0.424 |
| O00425 | IF2B3 | -0.487 | 0.001 | 0.044 |
| P20020 | AT2B1 | -0.489 | 0.134 | 0.459 |
| P07437 | TBB5 | -0.491 | 0.222 | 0.567 |
| P62829 | RL23 | -0.496 | 0.072 | 0.350 |
| Q14061 | COX17 | -0.498 | 0.129 | 0.459 |
| P62861 | RS30 | -0.501 | 0.106 | 0.424 |
| P23229-7 | ITA6 | -0.501 | 0.119 | 0.444 |
| Q92667 | AKAP1 | -0.504 | 0.021 | 0.191 |
| Q9NP97 | DLRB1 | -0.506 | 0.044 | 0.273 |
| Q14257 | RCN2 | -0.507 | 0.344 | 0.706 |
| P25788 | PSA3 | -0.509 | 0.035 | 0.248 |
| Q6HA08 | ASTL | -0.511 | 0.018 | 0.181 |
| O43852 | CALU | -0.515 | 0.202 | 0.545 |
| P13639 | EF2 | -0.518 | 0.015 | 0.168 |
| Q9P2J5 | SYLC | -0.518 | 0.022 | 0.192 |
| P09651 | ROA1 | -0.522 | 0.582 | 0.857 |
| Q12849 | GRSF1 | -0.522 | 0.222 | 0.567 |
| Q15366 | PCBP2 | -0.526 | 0.041 | 0.265 |
| P18065 | IBP2 | -0.530 | 0.557 | 0.842 |
| O14548 | COX7R | -0.530 | 0.235 | 0.590 |
| Q5T160 | SYRM | -0.531 | 0.120 | 0.444 |
| Q8NBJ4 | GOLM1 | -0.533 | 0.006 | 0.098 |
| P28288 | ABCD3 | -0.534 | 0.183 | 0.528 |
| Q16658 | FSCN1 | -0.535 | 0.149 | 0.478 |
| P62906 | RL10A | -0.536 | 0.021 | 0.191 |
| P51648 | AL3A2 | -0.538 | 0.138 | 0.469 |
| O75190-2 | DNJB6 | -0.547 | 0.267 | 0.623 |
| P40429 | RL13A | -0.550 | 0.044 | 0.273 |
| P62266 | RS23 | -0.553 | 0.011 | 0.136 |
| Q9UKA9 | PTBP2 | -0.557 | 0.572 | 0.842 |
| Q5T013 | HYI | -0.559 | 0.410 | 0.761 |
| Q5T6F2 | UBAP2 | -0.560 | 0.149 | 0.478 |
| Q9Y2Z9 | COQ6 | -0.564 | 0.159 | 0.504 |
| P63244 | RACK1 | -0.564 | 0.005 | 0.092 |
| P49588 | SYAC | -0.566 | 0.058 | 0.315 |
| Q53GQ0 | DHB12 | -0.568 | 0.167 | 0.505 |
| P12956 | XRCC6 | -0.569 | 0.167 | 0.505 |
| P61088 | UBE2N | -0.570 | 0.063 | 0.332 |
| P46783 | RS10 | -0.572 | 0.426 | 0.767 |
| P18621 | RL17 | -0.573 | 0.013 | 0.152 |
| P22626 | ROA2 | -0.574 | 0.063 | 0.332 |
| Q9UG56 | PISD | -0.575 | 0.057 | 0.315 |
| P35221 | CTNA1 | -0.575 | 0.148 | 0.478 |

| | | | | |
|---|---|---|---|---|
| P21333 | FLNA | -0.581 | 0.002 | 0.051 |
| A4D1E9 | GTPBA | -0.582 | 0.120 | 0.444 |
| Q13368 | MPP3 | -0.584 | 0.291 | 0.657 |
| O60701 | UGDH | -0.588 | 0.033 | 0.239 |
| P62277 | RS13 | -0.592 | 0.002 | 0.051 |
| P67936 | TPM4 | -0.593 | 0.075 | 0.350 |
| Q96AX1 | VP33A | -0.593 | 0.036 | 0.251 |
| P62241 | RS8 | -0.597 | 0.025 | 0.208 |
| Q8IXM3 | RM41 | -0.597 | 0.058 | 0.315 |
| P20339 | RAB5A | -0.614 | 0.254 | 0.614 |
| Q02818 | NUCB1 | -0.616 | 0.042 | 0.268 |
| Q96FV2 | SCRN2 | -0.618 | 0.277 | 0.643 |
| P30050 | RL12 | -0.619 | 0.120 | 0.444 |
| O94760 | DDAH1 | -0.620 | 0.058 | 0.315 |
| Q07020 | RL18 | -0.620 | 0.002 | 0.055 |
| P84098 | RL19 | -0.621 | 0.029 | 0.220 |
| Q5TAX3 | TUT4 | -0.621 | 0.222 | 0.567 |
| Q14894 | CRYM | -0.623 | 0.202 | 0.545 |
| P61313 | RL15 | -0.627 | 0.003 | 0.061 |
| Q96GD0 | PLPP | -0.630 | 0.134 | 0.459 |
| Q9BVG9 | PTSS2 | -0.634 | 0.075 | 0.350 |
| Q9Y2R5 | RT17 | -0.636 | 0.002 | 0.059 |
| P50336 | PPOX | -0.638 | 0.041 | 0.265 |
| Q9BW91 | NUDT9 | -0.638 | 0.144 | 0.478 |
| Q9UHD8 | SEPT9 | -0.644 | 0.011 | 0.136 |
| P62701 | RS4X | -0.645 | 0.009 | 0.127 |
| Q9H6T3 | RPAP3 | -0.650 | 0.175 | 0.526 |
| Q9UHD1 | CHRD1 | -0.656 | 0.013 | 0.152 |
| Q9H832 | UBE2Z | -0.669 | 0.207 | 0.554 |
| A6NDG6 | PGP | -0.676 | 0.316 | 0.676 |
| Q13310 | PABP4 | -0.676 | 0.006 | 0.101 |
| P05386 | RLA1 | -0.682 | 0.006 | 0.101 |
| P07858 | CATB | -0.682 | 0.025 | 0.208 |
| P35268 | RL22 | -0.683 | 0.212 | 0.566 |
| O00442 | RTCA | -0.684 | 0.074 | 0.350 |
| O43678 | NDUA2 | -0.687 | 0.082 | 0.376 |
| P25789 | PSA4 | -0.689 | 0.008 | 0.115 |
| Q14165 | MLEC | -0.695 | 0.058 | 0.315 |
| P61353 | RL27 | -0.703 | 0.002 | 0.055 |
| Q9Y5R8 | TPPC1 | -0.707 | 0.140 | 0.478 |
| P15880 | RS2 | -0.730 | 0.002 | 0.051 |
| P36543 | VATE1 | -0.732 | 0.002 | 0.055 |
| Q9HD33 | RM47 | -0.737 | 0.003 | 0.061 |
| Q96M93 | ADAD1 | -0.737 | 0.004 | 0.080 |
| P01178 | NEU1 | -0.740 | 0.085 | 0.379 |
| P35527 | K1C9 | -0.742 | 0.459 | 0.778 |
| P62081 | RS7 | -0.744 | 0.018 | 0.181 |
| Q6PKG0 | LARP1 | -0.746 | 0.018 | 0.181 |
| P09455 | RET1 | -0.770 | 0.193 | 0.545 |
| O00592 | PODXL | -0.776 | 0.003 | 0.061 |
| P51116 | FXR2 | -0.782 | 0.310 | 0.676 |
| P16278 | BGAL | -0.784 | 0.038 | 0.255 |
| Q5JVS0 | HABP4 | -0.786 | 0.101 | 0.422 |
| Q10713 | MPPA | -0.788 | 0.020 | 0.191 |
| P01857 | IGHG1 | -0.790 | 0.075 | 0.350 |
| Q969V3 | NCLN | -0.791 | 0.021 | 0.191 |

| | | | | | |
|---|---|---|---|---|---|
| | Q92896 | GSLG1 | -0.811 | 0.159 | 0.504 |
| | P07339 | CATD | -0.817 | 0.003 | 0.061 |
| | Q93050 | VPP1 | -0.820 | 0.074 | 0.350 |
| | Q14195-2 | DPYL3 | -0.844 | 0.101 | 0.422 |
| | Q13724 | MOGS | -0.854 | 0.006 | 0.098 |
| | Q9UBS3 | DNJB9 | -0.859 | 0.071 | 0.350 |
| | P62888 | RL30 | -0.872 | 0.006 | 0.101 |
| | P31153 | METK2 | -0.875 | 0.055 | 0.314 |
| | Q96G97 | BSCL2 | -0.889 | 0.144 | 0.478 |
| | P16219 | ACADS | -0.893 | 0.041 | 0.265 |
| | P18031 | PTN1 | -0.896 | 0.018 | 0.181 |
| | P14174 | MIF | -0.921 | 0.037 | 0.255 |
| | P46781 | RS9 | -0.939 | 0.006 | 0.101 |
| | Q9UKG1 | DP13A | -0.952 | 0.067 | 0.341 |
| | Q9Y5T4 | DJC15 | -0.954 | 0.003 | 0.061 |
| | Q9HAU5 | RENT2 | -0.959 | 0.056 | 0.315 |
| | P19087 | GNAT2 | -0.981 | 0.080 | 0.375 |
| | Q8NA58 | PNDC1 | -0.982 | 0.003 | 0.072 |
| | Q96CT7 | CC124 | -0.982 | 0.120 | 0.444 |
| | Q9UIJ7 | KAD3 | -0.993 | 0.028 | 0.220 |
| | Q8TF74 | WIPF2 | -1.052 | 0.011 | 0.136 |
| | Q13185 | CBX3 | -1.063 | 0.031 | 0.233 |
| | Q14197 | ICT1 | -1.064 | 0.111 | 0.437 |
| | P62805 | H4 | -1.082 | 0.005 | 0.092 |
| | Q15417 | CNN3 | -1.086 | 0.009 | 0.127 |
| | Q6XQN6 | PNCB | -1.105 | 0.082 | 0.376 |
| | Q6WRI0 | IGS10 | -1.152 | 0.025 | 0.209 |
| | P62263 | RS14 | -1.154 | 0.002 | 0.055 |
| | P02768 | ALBU | -1.182 | 0.403 | 0.752 |
| | O60684 | IMA7 | -1.379 | 0.038 | 0.255 |
| | Q6UB35 | C1TM | -1.603 | 0.002 | 0.055 |
| | Q92832 | NELL1 | -1.649 | 0.006 | 0.098 |
| | P08670 | VIME | -1.775 | 0.031 | 0.231 |
| Significantly abundant proteins in GV | Q8IZA3 | H18 | -0.593 | 0.001 | 0.048 |
| | O60884 | DNJA2 | -0.619 | 0.001 | 0.048 |
| | C9JE40 | PATL2 | -0.683 | 0.001 | 0.035 |
| | Q9NQI0 | DDX4 | -0.683 | 0.001 | 0.035 |
| | P62269 | RS18 | -0.709 | 0.001 | 0.048 |
| | P62750 | RL23A | -0.720 | 0.001 | 0.031 |
| | P62917 | RL8 | -0.766 | 0.001 | 0.046 |
| | P61006 | RAB8A | -0.786 | 0.000 | 0.025 |
| | P08865 | RSSA | -0.800 | 0.000 | 0.003 |
| | P49327 | FAS | -0.903 | 0.001 | 0.031 |
| | Q9BV73 | CP250 | -0.907 | 0.000 | 0.020 |
| | Q6IMN6 | CAPR2 | -0.948 | 0.000 | 0.005 |
| | Q9Y2T7 | YBOX2 | -1.025 | 0.000 | 0.000 |
| | P83731 | RL24 | -1.082 | 0.001 | 0.046 |
| | P43121 | MUC18 | -1.103 | 0.000 | 0.000 |
| | Q99805 | TM9S2 | -1.210 | 0.000 | 0.004 |
| | O75600 | KBL | -1.219 | 0.001 | 0.046 |
| | Q10471 | GALT2 | -1.231 | 0.001 | 0.046 |
| | P22234 | PUR6 | -1.276 | 0.000 | 0.013 |
| | Q9Y2W6 | TDRKH | -1.419 | 0.000 | 0.000 |
| | Q08629 | TICN1 | -1.419 | 0.000 | 0.005 |
| | P00338 | LDHA | -1.710 | 0.001 | 0.046 |
| | Q86SH2 | ZAR1 | -1.899 | 0.000 | 0.004 |

**Table S6. Identified changes in GO (Biological Processes, BP; Cellular Compartments, CC) and KEGG pathways. AMA group: MII *vs.* GV oocytes.**

| | Terms | Overlap | Ratio | P-value | P.adj | Log2 (FC) | Proteins |
|---|---|---|---|---|---|---|---|
| BP | translation (GO:0006412) | 111/214 | 0.5 | 2E-38 | 7E-36 | -0.299 | RL4; RL5; RL30; RL31; RL34; RL8; ACPH; RL10A; RL9; SYYM; RL6; RL7; EF1B; RS14; RS17; RS16; RS19; SYNC; RS18; RS11; RS10; RS13; RS12; SYRM; RS9; SYRC; RL21; RS7; RS8; RL23; RS5; RL22; RS6; RSSA; EFTU; IPYR2; IPYR; SYLC; EF1A2; RL24; RL27; RL29; SYCC; RL28; SYTC; SYAC; ERF3A; SYVC; HARS1; RM27; RM23; EF2; RM24; AIMP1; RS25; RS28; CSN5; SYEP; RS29; RL27A; RS20; SYFA; RS24; SYFB; RS23; IF4G1; RT17; HBS1L; RLA1; RLA0; RM41; RS4X; TR112; RL7A; SYQ; RLA2; EFTS; PABP4; SYMC; RL13A; RM47; RS3A; RM43; SYDC; ZAR1; EF1G; SYSC; SYSM; EF1D; ERF1; RL10; SYK; KGD4; RL12; RL11; RS15A; RL14; RS3; RL13; SYIM; IF2B3; RL15; RS2; RL18; RL17; RL19; NDUA7; RL23A; SYWC; GARS; ABCE1 |
| | cytoplasmic translation (GO:0002181) | 62/93 | 0.7 | 7E-43 | 9E-40 | -0.447 | IF4A2; RL4; IF4A1; RL5; RL30; RL31; RL34; RLA1; RLA0; RL10A; RL8; RL9; RL6; RL7; RS4X; RL7A; RS14; RS17; RS16; RS19; RS18; RS11; RS10; RS13; RS12; RS9; RL21; RS7; RS8; RL23; RS5; RL22; RS6; RL13A; RS3A; RSSA; RL24; ERF1; RL27; RL29; RL28; RL10; RL12; RL11; ERF3A; RS15A; RL14; RS3; RL13; RL15; RS2; RL18; RL17; RL19; RL23A; RS25; RS28; RS29; RL27A; RS20; RS24; RS23 |
| | cotranslational protein targeting to membrane (GO:0006613) | 62/94 | 0.7 | 4E-42 | 4E-39 | -0.443 | RL4; RL5; RL30; RL31; RL34; RLA1; RLA0; RL10A; RL8; RL9; RL6; RL7; RS4X; RL7A; RS14; RS17; RS16; RS19; RS18; RLA2; RS11; RS10; SIL1; RS13; RS12; SEC63; RS9; RL21; RS7; RS8; RL23; RS5; RL22; RS6; RL13A; RS3A; RSSA; RL24; RL27; RL29; RL28; RL10; RL12; RL11; RS15A; RL14; RS3; RL13; SSRA; RL15; RS2; RL18; RL17; RL19; RL23A; RS25; RS28; RS29; RL27A; RS20; RS24; RS23 |
| | protein targeting to ER (GO:0045047) | 64/103 | 0.6 | 1E-38 | 6E-36 | -0.416 | RL4; RL5; RL30; RL31; RL34; RLA1; ZFN2A; RLA0; RL10A; RL8; RL9; RL6; RL7; RS4X; RS14; RL7A; RS17; RS16; RS19; RS18; RLA2; RS11; GET3; RS10; RS13; RS12; SEC63; RS9; RL21; RS7; RS8; RL23; RS5; RL22; RS6; RL13A; RSSA; RS3A; CHM4B; RL24; RL27; RL29; RL28; RL10; RL12; RL11; RS15A; RL14; RS3; RL13; RL15; RS2; RL18; RL17; RL19; RL23A; RS25; RS28; RS29; RL27A; SRPRB; RS20; RS24; RS23 |
| | SRP-dependent cotranslational protein targeting to membrane (GO:0006614) | 60/90 | 0.7 | 7E-42 | 5E-39 | -0.452 | RL4; RL5; RL30; RL31; RL34; RLA1; RLA0; RL10A; RL8; RL9; RL6; RL7; RS4X; RL7A; RS14; RS17; RS16; RS19; RS18; RLA2; RS11; RS10; RS13; RS12; SEC63; RS9; RL21; RS7; RS8; RL23; RS5; RL22; RS6; RL13A; RS3A; RSSA; RL24; RL27; RL29; RL28; RL10; RL12; RL11; RS15A; RL14; RS3; RL13; RL15; RS2; RL18; RL17; RL19; RL23A; RS25; RS28; RS29; RL27A; RS20; RS24; RS23 |
| | nuclear-transcribed mRNA catabolic process, nonsense-mediated decay (GO:0000184) | 64/113 | 0.6 | 3E-43 | 7E-40 | -0.448 | RL4; RL5; RL30; RL31; RL34; RLA1; RLA0; RL10A; RL8; RL9; RL6; RL7; RS4X; RS14; RL7A; RS17; RS16; RS19; 2AAA; RS18; RLA2; RS11; RS10; RS13; RS12; RENT2; RS9; RL21; RS7; RS8; RL23; RS5; RL22; RS6; RL13A; RSSA; RS3A; RL24; RL27; ERF1; RL29; RL28; RL10; RL12; RL11; ERF3A; RS15A; RL14; RS3; RL13; RL15; RS2; RL18; RL17; RL19; RL23A; RS25; RS28; RS29; RL27A; RS20; RS24; IF4G1; RS23 |
| | fatty acid beta-oxidation using acyl-CoA dehydrogenase (GO:0033539) | 08/10 | 0.8 | 2E-03 | 4E-02 | -0.280 | GCDH; ACADV; IVD; ETFA; ACADM; ETFB; ACD11; ACADS |
| | protein deneddylation (GO:0000338) | 08/10 | 0.8 | 2E-04 | 9E-03 | 0.247 | CSN4; CSN3; CSN6; CSN5; CSN2; CSN1; CSN7B; CSN8 |
| | negative regulation of endoplasmic reticulum unfolded protein response (GO:1900102) | 07/13 | 0.5 | 2E-03 | 3E-02 | -0.369 | PTN1; BIP; WFS1; DDRGK; RACK1; DNJB9; NCK1 |
| | P-body assembly (GO:0033962) | 06/12 | 0.5 | 7E-04 | 2E-02 | -0.168 | DDX6; DYHC1; 4ET; LSM4; LSM3; PATL2 |
| | maintenance of DNA methylation (GO:0010216) | 03/06 | 0.5 | 7E-08 | 1E-05 | 0.827 | DNMT1; UHRF2; UHRF1 |
| CC | cytosolic large ribosomal subunit (GO:0022625) | 34/55 | 0.6 | 3E-27 | 5E-25 | -0.485 | RL4; RL5; RL30; RL10; RL31; RL34; RLA1; RL12; RLA0; RL11; RL10A; RL8; RL9; RL6; RL7; RL7A; FXR2; RL14; RLA2; RL13; RL15; RL18; RL17; RL19; RL21; RL23; RL22; RL13A; RL23A; RL27A; RL24; RL27; RL29; RL28 |
| | large ribosomal subunit (GO:0015934) | 35/59 | 0.6 | 4E-27 | 5E-25 | -0.473 | RL4; RL5; RL30; RL10; RL31; RL34; RLA1; RL12; RLA0; RL11; RL10A; RL8; RL9; RL6; RL7; RL7A; FXR2; RL14; RLA2; RL13; RL15; RL18; RL17; RL19; RL21; RL23; RL22; RL13A; RL23A; RM22; RL27A; RL24; RL27; RL29; RL28 |
| | cytosolic small ribosomal subunit (GO:0022627) | 27/40 | 0.7 | 1E-16 | 8E-15 | -0.417 | RS4X; RS14; RS17; RS15A; RS16; RS19; RS18; RS3; RS2; RS11; RS10; RS13; EIF2A; RS12; RS9; RS7; RS8; RS5; RS6; RS3A; RSSA; RS25; RS28; RS29; RS20; RS24; RS23 |
| | small ribosomal subunit (GO:0015935) | 28/45 | 0.6 | 4E-18 | 4E-16 | -0.423 | RS4X; RS14; RS17; RS15A; RS16; RS19; RS18; RACK1; RS3; RS2; RS11; RS10; RS13; EIF2A; RS12; RS9; RS7; RS8; RS5; RS6; RS3A; RSSA; RS25; RS28; RS29; RS20; RS24; RS23 |
| | ribosome (GO:0005840) | 32/62 | 0.5 | 3E-27 | 5E-25 | -0.500 | RL30; RL31; RL11; RL10A; RL8; RL9; RL6; RS4X; RL7A; RS17; LARP1; RS19; RS18; RS3; RL15; RS11; RL18; RS10; RS13; NCK1; RL19; RS9; RS7; RL23; RS5; RL13A; RS25; RS28; RS29; RL24; RL27; RS23 |
| | COPI vesicle coat (GO:0030126) | 07/13 | 0.5 | 2E-05 | 6E-04 | 0.263 | COPB2; COPA; COPB; COPG1; COPZ1; COPE; COPD |
| | piP-body (GO:0071547) | 03/05 | 0.6 | 3E-04 | 5E-03 | -0.568 | TDRKH; DDX4; MAEL |

**Table S7. Proteins showing correlation trends with age in GV oocytes. (|0.3| ≤ R< |0.5|; p≤0.05)**

| Uniprot ID | Protein name | R | P_value | P.adjusted |
|---|---|---|---|---|
| P62333 | PRS10 | -0.559 | 0.003 | 0.203 |
| P21266 | GSTM3 | -0.554 | 0.003 | 0.218 |
| O60925 | PFD1 | -0.543 | 0.005 | 0.289 |
| P30047 | GFRP | -0.539 | 0.005 | 0.291 |
| Q96CW5 | GCP3 | -0.539 | 0.005 | 0.291 |
| Q14677 | EPN4 | -0.536 | 0.006 | 0.294 |
| P61158 | ARP3 | -0.521 | 0.006 | 0.305 |
| Q9Y2Z0 | SGT1 | -0.521 | 0.006 | 0.305 |
| P11216 | PYGB | -0.518 | 0.007 | 0.305 |
| Q96QK1 | VPS35 | -0.514 | 0.007 | 0.317 |
| P06493 | CDK1 | -0.513 | 0.009 | 0.343 |
| Q96CN7 | ISOC1 | -0.510 | 0.008 | 0.329 |
| P17987 | TCPA | -0.502 | 0.009 | 0.343 |
| Q15102 | PA1B3 | -0.502 | 0.011 | 0.367 |
| P53041 | PPP5 | -0.500 | 0.009 | 0.343 |
| P51812 | KS6A3 | -0.496 | 0.014 | 0.389 |
| P49189 | AL9A1 | -0.489 | 0.011 | 0.374 |
| Q96IU4 | ABHEB | -0.488 | 0.013 | 0.389 |
| Q8TDQ7 | GNPI2 | -0.484 | 0.012 | 0.389 |
| P24666 | PPAC | -0.482 | 0.017 | 0.398 |
| Q8WW33 | GTSF1 | -0.481 | 0.015 | 0.396 |
| P62820 | RAB1A | -0.478 | 0.014 | 0.389 |
| P47755 | CAZA2 | -0.477 | 0.014 | 0.389 |
| P49593 | PPM1F | -0.471 | 0.015 | 0.396 |
| P60900 | PSA6 | -0.471 | 0.017 | 0.400 |
| P31150 | GDIA | -0.470 | 0.015 | 0.396 |
| P61160 | ARP2 | -0.469 | 0.016 | 0.396 |
| Q9Y266 | NUDC | -0.465 | 0.019 | 0.404 |
| Q9Y4Z0 | LSM4 | -0.462 | 0.020 | 0.404 |
| P50053-2 | KHK | -0.457 | 0.022 | 0.404 |
| P52788 | SPSY | -0.456 | 0.022 | 0.404 |
| Q9UBQ7 | GRHPR | -0.455 | 0.019 | 0.404 |
| P84085 | ARF5 | -0.454 | 0.023 | 0.404 |
| Q7L5D6 | GET4 | -0.452 | 0.040 | 0.442 |
| P00558 | PGK1 | -0.451 | 0.021 | 0.404 |
| P23528 | COF1 | -0.451 | 0.021 | 0.404 |

| | | | | |
|---|---|---|---|---|
| P02792 | FRIL | -0.451 | 0.021 | 0.404 |
| P02794 | FRIH | -0.450 | 0.021 | 0.404 |
| P13693 | TCTP | -0.450 | 0.024 | 0.404 |
| P50395 | GDIB | -0.447 | 0.022 | 0.404 |
| O94760 | DDAH1 | -0.446 | 0.022 | 0.404 |
| P22061-2 | PIMT | -0.443 | 0.023 | 0.404 |
| P35606 | COPB2 | -0.442 | 0.024 | 0.404 |
| A0AVT1 | UBA6 | -0.441 | 0.024 | 0.404 |
| P52209 | 6PGD | -0.441 | 0.024 | 0.404 |
| P49789 | FHIT | -0.438 | 0.025 | 0.416 |
| Q93034 | CUL5 | -0.436 | 0.029 | 0.441 |
| Q9Y3B8 | ORN | -0.433 | 0.027 | 0.436 |
| P61586 | RHOA | -0.432 | 0.031 | 0.442 |
| Q99962 | SH3G2 | -0.429 | 0.029 | 0.441 |
| O14618 | CCS | -0.427 | 0.033 | 0.442 |
| O60927 | PP1RB | -0.425 | 0.038 | 0.442 |
| Q9H910 | JUPI2 | -0.423 | 0.031 | 0.442 |
| P40925 | MDHC | -0.419 | 0.033 | 0.442 |
| P50990 | TCPQ | -0.419 | 0.033 | 0.442 |
| Q9NT62 | ATG3 | -0.418 | 0.037 | 0.442 |
| P15121 | ALDR | -0.418 | 0.033 | 0.442 |
| Q13561 | DCTN2 | -0.418 | 0.034 | 0.442 |
| P11055 | MYH3 | -0.416 | 0.043 | 0.454 |
| Q9Y3U8 | RL36 | -0.414 | 0.044 | 0.454 |
| P62312 | LSM6 | -0.413 | 0.040 | 0.442 |
| P23526 | SAHH | -0.412 | 0.037 | 0.442 |
| P80723 | BASP1 | -0.411 | 0.037 | 0.442 |
| Q15417 | CNN3 | -0.410 | 0.042 | 0.450 |
| Q15257-2 | PTPA | -0.407 | 0.039 | 0.442 |
| Q99497 | PARK7 | -0.407 | 0.039 | 0.442 |
| Q14019 | COTL1 | -0.407 | 0.039 | 0.442 |
| P06744 | G6PI | -0.407 | 0.039 | 0.442 |
| Q587J8 | KHDC3 | -0.406 | 0.040 | 0.442 |
| Q92530 | PSMF1 | -0.405 | 0.050 | 0.467 |
| O75390 | CISY | -0.402 | 0.042 | 0.450 |
| P48643 | TCPE | -0.401 | 0.042 | 0.450 |
| Q9BRA2 | TXD17 | -0.395 | 0.046 | 0.457 |
| P54652 | HSP72 | -0.390 | 0.049 | 0.467 |

| | | | | |
|---|---|---|---|---|
| P24539 | AT5F1 | 0.390 | 0.049 | 0.467 |
| Q8NBX0 | SCPDL | 0.392 | 0.048 | 0.467 |
| O94919 | ENDD1 | 0.394 | 0.047 | 0.460 |
| Q9Y639 | NPTN | 0.395 | 0.046 | 0.457 |
| Q15043 | S39AE | 0.395 | 0.050 | 0.467 |
| P62277 | RS13 | 0.395 | 0.046 | 0.457 |
| P28838 | AMPL | 0.397 | 0.045 | 0.454 |
| P36551 | HEM6 | 0.397 | 0.045 | 0.454 |
| Q9Y2S7 | PDIP2 | 0.398 | 0.044 | 0.454 |
| O15116 | LSM1 | 0.402 | 0.042 | 0.450 |
| P11021 | BIP | 0.402 | 0.042 | 0.450 |
| P08754 | GNAI3 | 0.404 | 0.050 | 0.467 |
| Q9P2E9 | RRBP1 | 0.405 | 0.045 | 0.454 |
| Q9H300 | PARL | 0.405 | 0.040 | 0.442 |
| Q9UBS3 | DNJB9 | 0.406 | 0.049 | 0.467 |
| O00186 | STXB3 | 0.406 | 0.049 | 0.467 |
| Q16851 | UGPA | 0.407 | 0.039 | 0.442 |
| Q9NQC3 | RTN4 | 0.408 | 0.039 | 0.442 |
| Q9Y277 | VDAC3 | 0.408 | 0.039 | 0.442 |
| P04843 | RPN1 | 0.411 | 0.037 | 0.442 |
| Q10471 | GALT2 | 0.411 | 0.037 | 0.442 |
| Q86W24 | NAL14 | 0.412 | 0.036 | 0.442 |
| Q8NFF5 | FAD1 | 0.414 | 0.045 | 0.454 |
| Q99798 | ACON | 0.415 | 0.035 | 0.442 |
| P39656 | OST48 | 0.417 | 0.034 | 0.442 |
| O95292 | VAPB | 0.418 | 0.034 | 0.442 |
| O60313-2 | OPA1 | 0.419 | 0.033 | 0.442 |
| P61006 | RAB8A | 0.419 | 0.033 | 0.442 |
| P05556 | ITB1 | 0.420 | 0.033 | 0.442 |
| P45880 | VDAC2 | 0.420 | 0.032 | 0.442 |
| Q9H173 | SIL1 | 0.426 | 0.030 | 0.442 |
| P18031 | PTN1 | 0.428 | 0.029 | 0.441 |
| Q9Y2Z9 | COQ6 | 0.434 | 0.034 | 0.442 |
| P09619 | PGFRB | 0.437 | 0.026 | 0.417 |
| O00217 | NDUS8 | 0.439 | 0.028 | 0.439 |
| Q5SW96 | ARH | 0.440 | 0.028 | 0.438 |
| O43615 | TIM44 | 0.443 | 0.024 | 0.404 |
| P04181 | OAT | 0.451 | 0.021 | 0.404 |

| | | | | |
|---|---|---|---|---|
| P05141 | ADT2 | 0.452 | 0.021 | 0.404 |
| P51148 | RAB5C | 0.452 | 0.020 | 0.404 |
| P04844-2 | RPN2 | 0.457 | 0.019 | 0.404 |
| Q8NFU3 | TSTD1 | 0.460 | 0.018 | 0.404 |
| P08195 | 4F2 | 0.465 | 0.017 | 0.396 |
| P62917 | RL8 | 0.465 | 0.017 | 0.396 |
| P40939 | ECHA | 0.465 | 0.017 | 0.396 |
| Q00325 | MPCP | 0.468 | 0.016 | 0.396 |
| P07355 | ANXA2 | 0.475 | 0.029 | 0.441 |
| P08559 | ODPA | 0.478 | 0.014 | 0.389 |
| Q9NX76 | CKLF6 | 0.480 | 0.028 | 0.438 |
| P54098 | DPOG1 | 0.484 | 0.014 | 0.393 |
| A4D1E9 | GTPBA | 0.486 | 0.012 | 0.387 |
| O43795 | MYO1B | 0.486 | 0.014 | 0.389 |
| Q9UJZ1 | STML2 | 0.492 | 0.011 | 0.367 |
| O00483 | NDUA4 | 0.493 | 0.017 | 0.396 |
| P43007 | SATT | 0.498 | 0.010 | 0.345 |
| Q07020 | RL18 | 0.501 | 0.009 | 0.343 |
| Q15084 | PDIA6 | 0.533 | 0.005 | 0.289 |
| Q9BTV4 | TMM43 | 0.534 | 0.009 | 0.343 |
| Q9BU23 | LMF2 | 0.560 | 0.007 | 0.305 |
| Q9HD33 | RM47 | 0.579 | 0.004 | 0.239 |

**Table S8. Correlation values for the abundance of proteasome complex subunits with age in GV oocytes. Age-Correlated Trends: |0.3| ≤ R< |0.5|; p≤0.05. Age-Correlated: R≥ |0.5|, p.adj≤0.2.**

| Uniprot ID | Protein name | R | P-value | P.adjusted |
|---|---|---|---|---|
| P62195 | PRS8 | -0.714 | 0.000 | 0.030 |
| P17980 | PRS6A | -0.623 | 0.001 | 0.107 |
| P62333 | PRS10 | -0.559 | 0.003 | 0.203 |
| P60900 | PSA6 | -0.471 | 0.017 | 0.400 |
| Q92530 | PSMF1 | -0.405 | 0.050 | 0.467 |
| P49720 | PSB3 | -0.376 | 0.070 | 0.509 |
| P51665 | PSMD7 | -0.366 | 0.066 | 0.509 |
| P62191 | PRS4 | -0.348 | 0.081 | 0.516 |
| Q13200 | PSMD2 | -0.295 | 0.143 | 0.634 |
| Q9UNM6 | PSD13 | -0.290 | 0.151 | 0.638 |
| P28072 | PSB6 | -0.282 | 0.172 | 0.662 |
| P48556 | PSMD8 | -0.271 | 0.189 | 0.678 |
| P49721 | PSB2 | -0.254 | 0.211 | 0.682 |
| O00231 | PSD11 | -0.232 | 0.254 | 0.711 |
| O43242 | PSMD3 | -0.190 | 0.352 | 0.769 |
| Q99460 | PSMD1 | -0.173 | 0.399 | 0.803 |
| Q06323 | PSME1 | -0.137 | 0.514 | 0.862 |
| O00232 | PSD12 | -0.133 | 0.518 | 0.866 |
| Q16401 | PSMD5 | -0.102 | 0.620 | 0.915 |
| P25789 | PSA4 | -0.085 | 0.679 | 0.933 |
| O00487 | PSDE | -0.068 | 0.746 | 0.950 |
| Q99436 | PSB7 | -0.045 | 0.826 | 0.965 |
| O75832 | PSD10 | -0.003 | 0.987 | 0.999 |
| P28066 | PSA5 | 0.027 | 0.894 | 0.974 |
| P28070 | PSB4 | 0.033 | 0.879 | 0.970 |
| P25787 | PSA2 | 0.044 | 0.836 | 0.965 |
| P25786 | PSA1 | 0.061 | 0.767 | 0.954 |
| Q15008 | PSMD6 | 0.096 | 0.641 | 0.921 |
| P20618 | PSB1 | 0.125 | 0.544 | 0.876 |
| O14818 | PSA7 | 0.154 | 0.452 | 0.830 |
| P43686 | PRS6B | 0.157 | 0.443 | 0.829 |
| P28074 | PSB5 | 0.158 | 0.440 | 0.829 |
| P25788 | PSA3 | 0.203 | 0.330 | 0.769 |
| P35998 | PRS7 | 0.275 | 0.174 | 0.663 |
| P55036 | PSMD4 | 0.282 | 0.162 | 0.662 |

$R_{mean}$ -0.137

**Table S9. Permutation Results for randomized correlations of Rmean**

| | GV | |
|---|---|---|
| Randomization # | Rmean for Proteasome subunits | Rmean for TRiC subunits |
| 1 | 0.050 | -0.078 |
| 2 | 0.066 | 0.102 |
| 3 | 0.008 | -0.042 |
| 4 | 0.029 | 0.044 |
| 5 | 0.006 | 0.094 |
| 6 | 0.031 | -0.061 |
| 7 | 0.013 | 0.154 |
| 8 | -0.037 | 0.035 |
| 9 | -0.001 | -0.036 |
| 10 | 0.006 | -0.001 |
| 11 | 0.020 | 0.201 |
| 12 | 0.036 | 0.154 |
| 13 | -0.005 | -0.182 |
| 14 | -0.001 | 0.040 |
| 15 | -0.001 | 0.068 |
| 16 | 0.051 | 0.070 |
| 17 | 0.043 | 0.083 |
| 18 | 0.033 | -0.022 |
| 19 | -0.015 | -0.043 |
| 20 | -0.016 | -0.083 |
| 21 | -0.049 | 0.060 |
| 22 | -0.028 | 0.084 |
| 23 | -0.050 | 0.083 |
| 24 | -0.014 | 0.052 |
| 25 | 0.000 | -0.010 |
| 26 | 0.020 | 0.085 |
| 27 | -0.007 | -0.099 |
| 28 | 0.058 | -0.029 |
| 29 | 0.034 | 0.057 |
| 30 | 0.073 | 0.087 |
| 31 | -0.026 | -0.061 |
| 32 | 0.004 | -0.027 |
| 33 | 0.027 | -0.044 |
| 34 | -0.037 | 0.005 |
| 35 | 0.043 | -0.013 |
| 36 | -0.021 | 0.003 |
| 37 | -0.018 | -0.028 |
| 38 | -0.033 | -0.044 |
| 39 | 0.070 | 0.007 |
| 40 | 0.042 | 0.031 |
| 41 | -0.029 | 0.043 |
| 42 | -0.039 | 0.024 |
| 43 | -0.053 | 0.097 |
| 44 | 0.010 | 0.001 |

| | | |
|---|---|---|
| 45 | 0.010 | -0.097 |
| 46 | -0.030 | 0.016 |
| 47 | -0.026 | 0.050 |
| 48 | -0.043 | 0.000 |
| 49 | 0.017 | -0.101 |
| 50 | 0.052 | -0.028 |
| 51 | 0.018 | -0.065 |
| 52 | 0.000 | 0.075 |
| 53 | -0.051 | -0.010 |
| 54 | -0.018 | 0.071 |
| 55 | -0.038 | -0.129 |
| 56 | 0.014 | 0.041 |
| 57 | -0.003 | 0.079 |
| 58 | -0.075 | 0.063 |
| 59 | 0.000 | -0.026 |
| 60 | 0.078 | 0.088 |
| 61 | -0.016 | -0.057 |
| 62 | 0.032 | -0.097 |
| 63 | -0.032 | 0.047 |
| 64 | -0.005 | 0.144 |
| 65 | -0.001 | -0.007 |
| 66 | 0.012 | 0.050 |
| 67 | -0.013 | -0.127 |
| 68 | 0.005 | -0.116 |
| 69 | 0.053 | -0.004 |
| 70 | 0.009 | 0.011 |
| 71 | 0.009 | -0.043 |
| 72 | -0.009 | 0.030 |
| 73 | 0.007 | -0.061 |
| 74 | 0.016 | 0.097 |
| 75 | 0.012 | -0.010 |
| 76 | 0.023 | -0.009 |
| 77 | -0.020 | -0.015 |
| 78 | 0.004 | 0.056 |
| 79 | -0.030 | 0.113 |
| 80 | -0.037 | 0.060 |
| 81 | 0.019 | -0.009 |
| 82 | -0.023 | -0.070 |
| 83 | -0.016 | 0.030 |
| 84 | 0.025 | 0.032 |
| 85 | 0.013 | -0.008 |
| 86 | 0.010 | 0.095 |
| 87 | -0.007 | 0.059 |
| 88 | -0.032 | 0.033 |
| 89 | -0.034 | 0.034 |
| 90 | 0.016 | 0.011 |
| 91 | 0.002 | -0.014 |
| 92 | -0.065 | 0.073 |
| 93 | 0.003 | -0.010 |
| 94 | 0.014 | 0.002 |
| 95 | -0.006 | 0.046 |

| | | |
|---|---|---|
| 96 | -0.001 | -0.101 |
| 97 | -0.010 | -0.042 |
| 98 | 0.076 | -0.054 |
| 99 | 0.004 | -0.059 |
| 100 | -0.027 | 0.032 |
| 101 | -0.013 | -0.053 |
| 102 | -0.002 | 0.006 |
| 103 | -0.002 | -0.051 |
| 104 | 0.028 | 0.030 |
| 105 | -0.079 | -0.150 |
| 106 | 0.008 | -0.022 |
| 107 | -0.072 | 0.109 |
| 108 | 0.008 | -0.059 |
| 109 | 0.036 | 0.042 |
| 110 | 0.061 | -0.093 |
| 111 | 0.002 | 0.058 |
| 112 | -0.011 | 0.022 |
| 113 | 0.037 | -0.079 |
| 114 | -0.047 | -0.021 |
| 115 | 0.018 | 0.042 |
| 116 | 0.004 | 0.139 |
| 117 | 0.034 | 0.037 |
| 118 | -0.012 | 0.023 |
| 119 | 0.039 | 0.035 |
| 120 | -0.047 | 0.015 |
| 121 | 0.028 | -0.106 |
| 122 | -0.033 | 0.034 |
| 123 | 0.011 | 0.052 |
| 124 | 0.055 | 0.026 |
| 125 | 0.039 | -0.105 |
| 126 | 0.054 | -0.015 |
| 127 | -0.024 | 0.023 |
| 128 | -0.005 | -0.010 |
| 129 | 0.061 | 0.087 |
| 130 | 0.038 | -0.009 |
| 131 | -0.032 | 0.035 |
| 132 | 0.008 | 0.067 |
| 133 | 0.044 | 0.179 |
| 134 | 0.046 | 0.085 |
| 135 | 0.033 | 0.162 |
| 136 | -0.009 | 0.050 |
| 137 | -0.007 | 0.006 |
| 138 | -0.010 | 0.003 |
| 139 | -0.019 | 0.103 |
| 140 | -0.019 | -0.033 |
| 141 | -0.056 | 0.020 |
| 142 | -0.027 | 0.112 |
| 143 | 0.005 | -0.006 |
| 144 | 0.019 | -0.024 |
| 145 | 0.004 | -0.052 |
| 146 | -0.043 | 0.065 |

| | | |
|---|---|---|
| 147 | 0.070 | 0.133 |
| 148 | 0.023 | -0.041 |
| 149 | 0.018 | -0.012 |
| 150 | -0.041 | -0.007 |
| 151 | 0.040 | -0.097 |
| 152 | -0.047 | 0.004 |
| 153 | 0.002 | 0.102 |
| 154 | 0.019 | -0.042 |
| 155 | 0.028 | -0.018 |
| 156 | -0.013 | -0.074 |
| 157 | -0.074 | -0.022 |
| 158 | -0.013 | 0.023 |
| 159 | 0.073 | 0.137 |
| 160 | 0.061 | 0.030 |
| 161 | -0.014 | -0.098 |
| 162 | 0.024 | 0.076 |
| 163 | -0.014 | -0.048 |
| 164 | 0.013 | -0.070 |
| 165 | -0.035 | 0.029 |
| 166 | -0.049 | 0.086 |
| 167 | 0.043 | -0.102 |
| 168 | -0.004 | 0.076 |
| 169 | -0.029 | -0.026 |
| 170 | -0.032 | -0.034 |
| 171 | 0.004 | -0.012 |
| 172 | -0.011 | 0.128 |
| 173 | -0.012 | -0.032 |
| 174 | 0.028 | -0.120 |
| 175 | -0.027 | -0.154 |
| 176 | 0.006 | 0.071 |
| 177 | 0.026 | 0.007 |
| 178 | -0.032 | -0.003 |
| 179 | -0.043 | -0.061 |
| 180 | -0.045 | 0.038 |
| 181 | -0.106 | -0.021 |
| 182 | 0.009 | -0.065 |
| 183 | -0.048 | -0.031 |
| 184 | 0.070 | -0.021 |
| 185 | -0.033 | -0.012 |
| 186 | 0.014 | 0.049 |
| 187 | 0.001 | -0.036 |
| 188 | -0.021 | -0.044 |
| 189 | 0.005 | -0.074 |
| 190 | 0.030 | -0.062 |
| 191 | 0.038 | -0.050 |
| 192 | -0.071 | -0.089 |
| 193 | 0.050 | 0.095 |
| 194 | -0.001 | -0.054 |
| 195 | 0.014 | 0.083 |
| 196 | 0.015 | -0.062 |
| 197 | -0.002 | -0.109 |

| | | |
|---|---|---|
| 198 | -0.038 | 0.020 |
| 199 | -0.004 | -0.049 |
| 200 | 0.061 | 0.028 |
| 201 | 0.058 | -0.066 |
| 202 | 0.008 | -0.075 |
| 203 | 0.006 | 0.050 |
| 204 | -0.023 | 0.011 |
| 205 | 0.057 | -0.021 |
| 206 | -0.008 | 0.150 |
| 207 | 0.002 | 0.070 |
| 208 | 0.051 | 0.014 |
| 209 | 0.023 | 0.108 |
| 210 | -0.094 | -0.029 |
| 211 | 0.010 | -0.155 |
| 212 | 0.009 | 0.039 |
| 213 | 0.013 | -0.134 |
| 214 | -0.004 | -0.016 |
| 215 | -0.023 | -0.095 |
| 216 | 0.046 | -0.084 |
| 217 | 0.058 | 0.038 |
| 218 | 0.006 | 0.027 |
| 219 | 0.003 | -0.036 |
| 220 | -0.010 | 0.027 |
| 221 | 0.016 | -0.058 |
| 222 | -0.003 | -0.035 |
| 223 | -0.042 | 0.083 |
| 224 | 0.007 | -0.069 |
| 225 | 0.007 | -0.009 |
| 226 | -0.011 | 0.181 |
| 227 | -0.047 | -0.022 |
| 228 | -0.009 | -0.017 |
| 229 | -0.035 | 0.013 |
| 230 | -0.023 | -0.081 |
| 231 | 0.003 | -0.041 |
| 232 | 0.045 | 0.075 |
| 233 | -0.010 | -0.107 |
| 234 | 0.004 | -0.117 |
| 235 | -0.057 | 0.092 |
| 236 | 0.005 | 0.079 |
| 237 | -0.004 | -0.081 |
| 238 | 0.045 | 0.036 |
| 239 | 0.037 | -0.106 |
| 240 | -0.031 | 0.111 |
| 241 | 0.093 | 0.169 |
| 242 | -0.028 | 0.051 |
| 243 | -0.002 | 0.051 |
| 244 | -0.042 | -0.059 |
| 245 | -0.031 | -0.147 |
| 246 | -0.025 | 0.119 |
| 247 | -0.089 | 0.062 |
| 248 | -0.007 | -0.091 |

| | | |
|---|---|---|
| 249 | 0.006 | -0.133 |
| 250 | 0.001 | -0.149 |
| 251 | -0.030 | -0.120 |
| 252 | 0.020 | 0.016 |
| 253 | -0.044 | -0.217 |
| 254 | 0.000 | -0.089 |
| 255 | 0.059 | 0.071 |
| 256 | 0.031 | -0.014 |
| 257 | -0.017 | 0.001 |
| 258 | -0.062 | -0.057 |
| 259 | -0.029 | 0.035 |
| 260 | 0.017 | -0.047 |
| 261 | -0.011 | -0.054 |
| 262 | -0.014 | 0.020 |
| 263 | -0.021 | -0.001 |
| 264 | -0.026 | -0.048 |
| 265 | 0.037 | 0.004 |
| 266 | 0.045 | -0.038 |
| 267 | -0.008 | 0.025 |
| 268 | -0.034 | 0.038 |
| 269 | 0.002 | -0.125 |
| 270 | 0.021 | 0.001 |
| 271 | -0.041 | -0.051 |
| 272 | -0.003 | 0.037 |
| 273 | -0.039 | -0.002 |
| 274 | -0.045 | 0.009 |
| 275 | 0.025 | -0.152 |
| 276 | 0.007 | -0.074 |
| 277 | 0.017 | 0.043 |
| 278 | -0.029 | -0.052 |
| 279 | 0.001 | -0.087 |
| 280 | -0.016 | -0.034 |
| 281 | -0.025 | -0.054 |
| 282 | 0.053 | -0.016 |
| 283 | 0.051 | -0.012 |
| 284 | 0.045 | -0.192 |
| 285 | 0.038 | 0.052 |
| 286 | 0.007 | -0.011 |
| 287 | -0.053 | 0.084 |
| 288 | 0.007 | -0.065 |
| 289 | -0.018 | -0.073 |
| 290 | -0.016 | -0.003 |
| 291 | 0.014 | 0.009 |
| 292 | 0.052 | 0.097 |
| 293 | -0.020 | -0.014 |
| 294 | -0.017 | -0.065 |
| 295 | 0.017 | -0.012 |
| 296 | -0.007 | 0.059 |
| 297 | -0.049 | 0.096 |
| 298 | -0.001 | -0.121 |
| 299 | 0.042 | 0.056 |

| | | |
|---|---|---|
| 300 | 0.019 | 0.088 |
| 301 | -0.044 | 0.026 |
| 302 | -0.033 | 0.061 |
| 303 | 0.024 | -0.145 |
| 304 | 0.031 | -0.089 |
| 305 | -0.006 | 0.032 |
| 306 | 0.013 | -0.028 |
| 307 | -0.086 | 0.096 |
| 308 | -0.042 | -0.113 |
| 309 | 0.024 | -0.008 |
| 310 | -0.014 | 0.067 |
| 311 | -0.050 | 0.014 |
| 312 | 0.002 | 0.055 |
| 313 | -0.022 | 0.003 |
| 314 | -0.032 | -0.102 |
| 315 | -0.017 | -0.011 |
| 316 | 0.067 | 0.056 |
| 317 | -0.054 | -0.091 |
| 318 | 0.008 | -0.093 |
| 319 | 0.044 | -0.048 |
| 320 | -0.018 | -0.142 |
| 321 | -0.033 | -0.131 |
| 322 | 0.001 | 0.058 |
| 323 | 0.028 | -0.029 |
| 324 | 0.003 | 0.043 |
| 325 | -0.003 | 0.015 |
| 326 | -0.008 | -0.059 |
| 327 | -0.001 | -0.011 |
| 328 | 0.025 | -0.023 |
| 329 | -0.040 | -0.075 |
| 330 | -0.022 | 0.017 |
| 331 | 0.042 | -0.017 |
| 332 | 0.030 | -0.053 |
| 333 | 0.042 | 0.036 |
| 334 | -0.004 | -0.056 |
| 335 | 0.002 | -0.032 |
| 336 | 0.006 | 0.048 |
| 337 | -0.021 | 0.119 |
| 338 | 0.015 | -0.052 |
| 339 | 0.024 | 0.068 |
| 340 | -0.018 | -0.002 |
| 341 | -0.033 | -0.026 |
| 342 | -0.052 | 0.032 |
| 343 | -0.036 | 0.020 |
| 344 | 0.015 | 0.002 |
| 345 | -0.027 | 0.104 |
| 346 | -0.005 | 0.096 |
| 347 | 0.006 | -0.036 |
| 348 | -0.026 | -0.067 |
| 349 | -0.044 | 0.026 |
| 350 | 0.032 | 0.048 |

| | | |
|---|---|---|
| 351 | 0.016 | 0.077 |
| 352 | -0.046 | -0.055 |
| 353 | 0.021 | 0.004 |
| 354 | 0.012 | 0.037 |
| 355 | 0.068 | -0.040 |
| 356 | -0.042 | -0.101 |
| 357 | -0.012 | -0.032 |
| 358 | 0.001 | -0.064 |
| 359 | -0.027 | 0.042 |
| 360 | -0.020 | 0.031 |
| 361 | -0.043 | -0.020 |
| 362 | 0.052 | -0.051 |
| 363 | 0.000 | -0.026 |
| 364 | -0.011 | 0.127 |
| 365 | 0.038 | 0.176 |
| 366 | -0.031 | -0.076 |
| 367 | 0.005 | -0.085 |
| 368 | 0.024 | 0.025 |
| 369 | -0.076 | -0.010 |
| 370 | 0.027 | -0.074 |
| 371 | 0.045 | -0.051 |
| 372 | 0.076 | -0.033 |
| 373 | -0.004 | 0.028 |
| 374 | 0.025 | -0.020 |
| 375 | 0.024 | 0.040 |
| 376 | -0.015 | 0.003 |
| 377 | 0.069 | -0.058 |
| 378 | 0.082 | 0.074 |
| 379 | 0.050 | 0.088 |
| 380 | -0.015 | -0.133 |
| 381 | 0.024 | 0.006 |
| 382 | -0.096 | -0.008 |
| 383 | -0.027 | 0.063 |
| 384 | -0.007 | -0.015 |
| 385 | 0.025 | -0.048 |
| 386 | -0.019 | -0.005 |
| 387 | 0.039 | 0.047 |
| 388 | 0.047 | -0.014 |
| 389 | 0.002 | -0.083 |
| 390 | -0.010 | -0.037 |
| 391 | 0.039 | 0.091 |
| 392 | 0.023 | 0.041 |
| 393 | -0.016 | 0.000 |
| 394 | 0.074 | 0.085 |
| 395 | -0.029 | 0.010 |
| 396 | 0.036 | 0.023 |
| 397 | 0.027 | -0.103 |
| 398 | 0.006 | 0.029 |
| 399 | 0.050 | 0.116 |
| 400 | -0.051 | 0.087 |
| 401 | 0.022 | -0.042 |

| | | |
|---|---|---|
| 402 | -0.004 | -0.060 |
| 403 | 0.006 | 0.076 |
| 404 | -0.082 | 0.089 |
| 405 | -0.014 | -0.047 |
| 406 | -0.005 | -0.062 |
| 407 | -0.004 | -0.052 |
| 408 | -0.062 | 0.012 |
| 409 | -0.030 | -0.017 |
| 410 | 0.004 | 0.022 |
| 411 | -0.032 | 0.081 |
| 412 | -0.005 | -0.052 |
| 413 | -0.030 | 0.034 |
| 414 | 0.044 | 0.048 |
| 415 | -0.052 | 0.048 |
| 416 | 0.000 | -0.011 |
| 417 | -0.067 | -0.041 |
| 418 | 0.027 | 0.038 |
| 419 | 0.002 | 0.003 |
| 420 | -0.068 | -0.074 |
| 421 | -0.016 | 0.090 |
| 422 | 0.012 | 0.055 |
| 423 | -0.013 | -0.101 |
| 424 | -0.058 | -0.005 |
| 425 | 0.039 | 0.122 |
| 426 | 0.017 | -0.001 |
| 427 | 0.063 | 0.057 |
| 428 | 0.022 | -0.082 |
| 429 | 0.009 | -0.090 |
| 430 | -0.007 | 0.046 |
| 431 | 0.080 | 0.025 |
| 432 | 0.031 | -0.024 |
| 433 | 0.029 | 0.059 |
| 434 | 0.001 | -0.127 |
| 435 | 0.063 | -0.010 |
| 436 | -0.023 | -0.048 |
| 437 | -0.014 | -0.052 |
| 438 | -0.033 | -0.041 |
| 439 | 0.019 | -0.016 |
| 440 | -0.010 | 0.035 |
| 441 | 0.022 | 0.023 |
| 442 | 0.001 | 0.033 |
| 443 | 0.020 | 0.011 |
| 444 | -0.025 | 0.048 |
| 445 | 0.027 | -0.127 |
| 446 | -0.006 | -0.030 |
| 447 | 0.028 | 0.065 |
| 448 | -0.009 | -0.026 |
| 449 | 0.002 | 0.037 |
| 450 | 0.031 | -0.061 |
| 451 | 0.002 | -0.075 |
| 452 | -0.003 | -0.014 |

| | | |
|---|---|---|
| 453 | -0.003 | -0.053 |
| 454 | 0.016 | -0.026 |
| 455 | -0.037 | 0.035 |
| 456 | -0.054 | -0.006 |
| 457 | -0.010 | -0.067 |
| 458 | 0.031 | -0.077 |
| 459 | -0.008 | -0.066 |
| 460 | -0.034 | 0.120 |
| 461 | -0.004 | -0.015 |
| 462 | 0.023 | 0.115 |
| 463 | 0.006 | 0.115 |
| 464 | -0.044 | -0.089 |
| 465 | -0.026 | -0.111 |
| 466 | -0.029 | -0.014 |
| 467 | 0.048 | 0.038 |
| 468 | 0.068 | 0.092 |
| 469 | 0.017 | 0.028 |
| 470 | 0.015 | 0.105 |
| 471 | 0.016 | 0.022 |
| 472 | 0.015 | -0.030 |
| 473 | -0.015 | 0.009 |
| 474 | -0.008 | 0.008 |
| 475 | 0.000 | -0.104 |
| 476 | -0.038 | 0.111 |
| 477 | 0.000 | 0.001 |
| 478 | -0.039 | -0.018 |
| 479 | 0.001 | 0.084 |
| 480 | -0.085 | -0.016 |
| 481 | 0.028 | -0.124 |
| 482 | -0.048 | -0.016 |
| 483 | -0.006 | -0.006 |
| 484 | -0.022 | -0.077 |
| 485 | -0.013 | -0.073 |
| 486 | 0.029 | -0.003 |
| 487 | 0.055 | 0.129 |
| 488 | -0.040 | 0.105 |
| 489 | 0.009 | 0.095 |
| 490 | 0.045 | 0.030 |
| 491 | -0.052 | -0.023 |
| 492 | -0.041 | 0.031 |
| 493 | 0.040 | 0.145 |
| 494 | -0.037 | 0.009 |
| 495 | -0.016 | 0.054 |
| 496 | 0.056 | -0.027 |
| 497 | -0.019 | 0.042 |
| 498 | 0.015 | -0.007 |
| 499 | -0.011 | -0.045 |
| 500 | -0.001 | -0.056 |
| 501 | 0.028 | 0.021 |
| 502 | -0.012 | -0.013 |
| 503 | -0.043 | -0.024 |

| | | |
|---|---|---|
| 504 | 0.028 | -0.029 |
| 505 | 0.037 | 0.149 |
| 506 | 0.018 | 0.067 |
| 507 | -0.046 | 0.003 |
| 508 | 0.040 | 0.041 |
| 509 | 0.043 | -0.038 |
| 510 | -0.003 | -0.051 |
| 511 | 0.020 | 0.033 |
| 512 | 0.105 | -0.016 |
| 513 | -0.031 | 0.085 |
| 514 | 0.049 | -0.025 |
| 515 | 0.031 | -0.020 |
| 516 | -0.025 | 0.157 |
| 517 | -0.016 | -0.004 |
| 518 | 0.074 | 0.120 |
| 519 | -0.062 | -0.072 |
| 520 | 0.013 | -0.069 |
| 521 | -0.030 | -0.034 |
| 522 | -0.025 | 0.142 |
| 523 | 0.006 | -0.021 |
| 524 | 0.010 | -0.090 |
| 525 | -0.001 | -0.049 |
| 526 | 0.010 | -0.012 |
| 527 | -0.019 | 0.069 |
| 528 | -0.026 | 0.047 |
| 529 | -0.007 | -0.064 |
| 530 | 0.027 | 0.025 |
| 531 | -0.045 | 0.097 |
| 532 | 0.022 | -0.033 |
| 533 | -0.031 | -0.056 |
| 534 | 0.032 | 0.116 |
| 535 | -0.018 | 0.014 |
| 536 | -0.063 | -0.003 |
| 537 | -0.058 | 0.073 |
| 538 | 0.007 | 0.045 |
| 539 | -0.026 | 0.054 |
| 540 | 0.024 | -0.082 |
| 541 | -0.066 | 0.005 |
| 542 | -0.024 | 0.029 |
| 543 | 0.043 | 0.040 |
| 544 | -0.005 | -0.021 |
| 545 | -0.030 | -0.037 |
| 546 | -0.011 | -0.083 |
| 547 | 0.029 | -0.008 |
| 548 | -0.030 | 0.098 |
| 549 | -0.066 | 0.112 |
| 550 | 0.010 | -0.026 |
| 551 | 0.023 | 0.030 |
| 552 | 0.041 | 0.081 |
| 553 | -0.042 | 0.029 |
| 554 | -0.019 | 0.005 |

| | | |
|---|---|---|
| 555 | -0.017 | 0.040 |
| 556 | 0.079 | -0.033 |
| 557 | -0.017 | -0.142 |
| 558 | 0.027 | 0.043 |
| 559 | -0.028 | 0.061 |
| 560 | 0.048 | -0.079 |
| 561 | -0.049 | -0.048 |
| 562 | -0.056 | 0.004 |
| 563 | 0.051 | 0.008 |
| 564 | -0.024 | 0.003 |
| 565 | 0.013 | -0.071 |
| 566 | 0.003 | -0.064 |
| 567 | -0.033 | 0.051 |
| 568 | -0.004 | -0.094 |
| 569 | 0.005 | 0.026 |
| 570 | -0.047 | -0.015 |
| 571 | -0.013 | 0.038 |
| 572 | -0.032 | 0.116 |
| 573 | -0.008 | 0.065 |
| 574 | -0.038 | -0.091 |
| 575 | 0.032 | 0.112 |
| 576 | 0.022 | -0.069 |
| 577 | 0.025 | -0.078 |
| 578 | 0.029 | -0.044 |
| 579 | 0.015 | -0.063 |
| 580 | 0.014 | 0.208 |
| 581 | -0.022 | 0.071 |
| 582 | -0.001 | -0.071 |
| 583 | 0.002 | 0.005 |
| 584 | -0.013 | 0.022 |
| 585 | -0.006 | 0.002 |
| 586 | -0.047 | 0.058 |
| 587 | 0.000 | 0.036 |
| 588 | -0.009 | -0.011 |
| 589 | 0.023 | 0.093 |
| 590 | -0.012 | -0.064 |
| 591 | 0.020 | 0.061 |
| 592 | -0.023 | -0.091 |
| 593 | 0.006 | 0.112 |
| 594 | 0.034 | 0.084 |
| 595 | 0.016 | -0.143 |
| 596 | -0.014 | 0.051 |
| 597 | 0.004 | 0.145 |
| 598 | 0.071 | 0.000 |
| 599 | 0.010 | 0.045 |
| 600 | -0.004 | -0.111 |
| 601 | 0.043 | -0.147 |
| 602 | 0.025 | -0.060 |
| 603 | 0.050 | 0.011 |
| 604 | 0.038 | -0.097 |
| 605 | -0.022 | -0.035 |

| | | |
|---|---|---|
| 606 | 0.000 | -0.099 |
| 607 | 0.005 | -0.035 |
| 608 | 0.039 | -0.008 |
| 609 | 0.028 | -0.080 |
| 610 | -0.018 | 0.100 |
| 611 | -0.039 | -0.015 |
| 612 | 0.001 | 0.064 |
| 613 | -0.050 | 0.038 |
| 614 | 0.003 | 0.103 |
| 615 | 0.015 | -0.033 |
| 616 | 0.015 | 0.085 |
| 617 | -0.059 | -0.075 |
| 618 | -0.012 | -0.145 |
| 619 | -0.024 | -0.082 |
| 620 | 0.013 | -0.025 |
| 621 | 0.030 | 0.088 |
| 622 | 0.046 | 0.013 |
| 623 | 0.015 | -0.025 |
| 624 | -0.022 | -0.077 |
| 625 | 0.081 | -0.062 |
| 626 | 0.017 | -0.138 |
| 627 | 0.012 | -0.048 |
| 628 | 0.033 | 0.013 |
| 629 | 0.008 | 0.145 |
| 630 | 0.033 | 0.017 |
| 631 | 0.060 | 0.039 |
| 632 | 0.040 | -0.009 |
| 633 | 0.043 | -0.067 |
| 634 | -0.023 | -0.029 |
| 635 | -0.019 | -0.040 |
| 636 | 0.029 | -0.064 |
| 637 | 0.026 | 0.086 |
| 638 | 0.017 | 0.000 |
| 639 | 0.024 | -0.044 |
| 640 | -0.023 | 0.035 |
| 641 | -0.003 | -0.126 |
| 642 | -0.083 | 0.002 |
| 643 | -0.042 | -0.015 |
| 644 | 0.038 | -0.042 |
| 645 | -0.011 | -0.028 |
| 646 | -0.019 | 0.069 |
| 647 | -0.013 | -0.025 |
| 648 | -0.015 | 0.028 |
| 649 | -0.056 | -0.003 |
| 650 | -0.020 | -0.005 |
| 651 | -0.014 | -0.051 |
| 652 | 0.019 | 0.103 |
| 653 | -0.018 | -0.061 |
| 654 | 0.020 | -0.001 |
| 655 | 0.011 | -0.112 |
| 656 | -0.049 | -0.012 |

| | | |
|---|---|---|
| 657 | -0.057 | 0.097 |
| 658 | -0.010 | 0.011 |
| 659 | -0.067 | -0.003 |
| 660 | 0.026 | -0.141 |
| 661 | -0.049 | -0.090 |
| 662 | -0.039 | 0.005 |
| 663 | 0.001 | 0.002 |
| 664 | 0.058 | 0.114 |
| 665 | -0.056 | -0.026 |
| 666 | 0.013 | 0.033 |
| 667 | 0.011 | -0.087 |
| 668 | -0.012 | -0.002 |
| 669 | -0.045 | -0.149 |
| 670 | 0.041 | -0.024 |
| 671 | -0.016 | -0.032 |
| 672 | -0.014 | 0.045 |
| 673 | -0.016 | -0.012 |
| 674 | 0.010 | 0.164 |
| 675 | -0.046 | -0.053 |
| 676 | 0.022 | -0.011 |
| 677 | -0.013 | 0.058 |
| 678 | 0.033 | 0.003 |
| 679 | 0.037 | -0.077 |
| 680 | 0.006 | -0.069 |
| 681 | -0.057 | 0.165 |
| 682 | -0.034 | 0.066 |
| 683 | 0.005 | 0.065 |
| 684 | 0.033 | 0.010 |
| 685 | 0.053 | -0.073 |
| 686 | 0.017 | 0.145 |
| 687 | -0.016 | -0.016 |
| 688 | 0.050 | 0.020 |
| 689 | -0.014 | -0.051 |
| 690 | 0.061 | 0.027 |
| 691 | 0.005 | 0.130 |
| 692 | 0.063 | -0.060 |
| 693 | -0.062 | 0.078 |
| 694 | -0.006 | 0.031 |
| 695 | 0.078 | 0.006 |
| 696 | -0.013 | 0.001 |
| 697 | -0.040 | 0.014 |
| 698 | 0.017 | -0.057 |
| 699 | 0.037 | -0.065 |
| 700 | -0.030 | -0.029 |
| 701 | -0.045 | 0.102 |
| 702 | 0.005 | -0.123 |
| 703 | -0.011 | -0.057 |
| 704 | 0.017 | 0.170 |
| 705 | 0.008 | 0.052 |
| 706 | 0.017 | -0.001 |
| 707 | -0.038 | -0.072 |

| | | |
|---|---|---|
| 708 | -0.021 | -0.091 |
| 709 | -0.064 | -0.029 |
| 710 | 0.035 | 0.049 |
| 711 | -0.035 | 0.109 |
| 712 | -0.059 | 0.078 |
| 713 | -0.053 | 0.070 |
| 714 | -0.021 | 0.006 |
| 715 | 0.038 | 0.122 |
| 716 | 0.019 | 0.015 |
| 717 | -0.044 | 0.018 |
| 718 | -0.007 | 0.078 |
| 719 | -0.046 | -0.009 |
| 720 | -0.059 | 0.071 |
| 721 | 0.032 | 0.023 |
| 722 | -0.046 | 0.017 |
| 723 | -0.061 | -0.026 |
| 724 | 0.065 | 0.104 |
| 725 | -0.049 | -0.008 |
| 726 | 0.033 | 0.005 |
| 727 | 0.007 | -0.087 |
| 728 | -0.007 | 0.097 |
| 729 | 0.031 | 0.008 |
| 730 | 0.009 | 0.027 |
| 731 | -0.059 | 0.071 |
| 732 | -0.018 | 0.039 |
| 733 | -0.029 | 0.085 |
| 734 | 0.026 | -0.053 |
| 735 | 0.025 | -0.070 |
| 736 | 0.029 | 0.107 |
| 737 | 0.032 | 0.057 |
| 738 | -0.028 | 0.027 |
| 739 | 0.026 | -0.093 |
| 740 | -0.018 | 0.006 |
| 741 | -0.027 | 0.005 |
| 742 | 0.033 | 0.079 |
| 743 | -0.021 | 0.052 |
| 744 | 0.009 | -0.151 |
| 745 | -0.043 | -0.146 |
| 746 | -0.036 | 0.086 |
| 747 | 0.035 | -0.014 |
| 748 | -0.046 | 0.008 |
| 749 | 0.037 | 0.030 |
| 750 | 0.058 | 0.042 |
| 751 | -0.027 | -0.060 |
| 752 | -0.026 | -0.093 |
| 753 | -0.038 | 0.106 |
| 754 | 0.010 | 0.054 |
| 755 | 0.034 | 0.064 |
| 756 | 0.020 | 0.035 |
| 757 | 0.005 | 0.014 |
| 758 | -0.045 | 0.019 |

| | | |
|---|---|---|
| 759 | 0.078 | -0.055 |
| 760 | -0.018 | 0.069 |
| 761 | -0.014 | 0.159 |
| 762 | -0.017 | 0.140 |
| 763 | -0.017 | -0.036 |
| 764 | -0.023 | -0.005 |
| 765 | 0.028 | 0.080 |
| 766 | 0.037 | -0.035 |
| 767 | 0.046 | -0.066 |
| 768 | 0.031 | -0.001 |
| 769 | -0.051 | 0.097 |
| 770 | -0.023 | -0.057 |
| 771 | -0.044 | -0.017 |
| 772 | 0.008 | -0.037 |
| 773 | 0.015 | -0.050 |
| 774 | -0.023 | -0.042 |
| 775 | 0.021 | -0.094 |
| 776 | 0.017 | -0.108 |
| 777 | 0.031 | 0.082 |
| 778 | 0.017 | -0.120 |
| 779 | -0.064 | -0.107 |
| 780 | 0.026 | -0.031 |
| 781 | 0.036 | 0.092 |
| 782 | 0.006 | 0.015 |
| 783 | 0.002 | -0.083 |
| 784 | 0.002 | 0.053 |
| 785 | 0.043 | 0.174 |
| 786 | 0.058 | -0.075 |
| 787 | -0.036 | -0.023 |
| 788 | 0.039 | -0.008 |
| 789 | 0.011 | 0.045 |
| 790 | -0.013 | -0.067 |
| 791 | 0.066 | -0.060 |
| 792 | -0.037 | -0.005 |
| 793 | -0.066 | -0.034 |
| 794 | -0.061 | 0.009 |
| 795 | 0.073 | -0.029 |
| 796 | -0.001 | -0.020 |
| 797 | -0.037 | 0.000 |
| 798 | 0.033 | -0.075 |
| 799 | -0.040 | -0.114 |
| 800 | 0.054 | 0.028 |
| 801 | 0.067 | 0.098 |
| 802 | -0.031 | 0.043 |
| 803 | 0.041 | -0.068 |
| 804 | -0.010 | -0.012 |
| 805 | -0.012 | -0.045 |
| 806 | 0.038 | 0.075 |
| 807 | 0.001 | -0.015 |
| 808 | 0.006 | 0.050 |
| 809 | 0.055 | -0.047 |

| | | |
|---|---|---|
| 810 | 0.006 | -0.036 |
| 811 | 0.015 | -0.095 |
| 812 | -0.040 | -0.010 |
| 813 | 0.024 | -0.168 |
| 814 | 0.027 | -0.103 |
| 815 | 0.022 | 0.051 |
| 816 | 0.020 | -0.039 |
| 817 | 0.029 | 0.040 |
| 818 | -0.001 | 0.035 |
| 819 | -0.003 | 0.028 |
| 820 | 0.050 | -0.035 |
| 821 | 0.021 | -0.034 |
| 822 | -0.003 | 0.100 |
| 823 | -0.039 | 0.033 |
| 824 | -0.055 | -0.045 |
| 825 | -0.062 | -0.082 |
| 826 | -0.018 | -0.060 |
| 827 | -0.022 | 0.001 |
| 828 | -0.004 | 0.102 |
| 829 | -0.011 | -0.057 |
| 830 | 0.043 | 0.009 |
| 831 | 0.034 | -0.071 |
| 832 | -0.002 | -0.048 |
| 833 | -0.020 | 0.028 |
| 834 | -0.033 | 0.065 |
| 835 | -0.005 | -0.012 |
| 836 | 0.034 | 0.014 |
| 837 | -0.058 | 0.073 |
| 838 | -0.031 | -0.105 |
| 839 | -0.051 | 0.089 |
| 840 | -0.044 | -0.110 |
| 841 | 0.034 | 0.021 |
| 842 | 0.001 | 0.055 |
| 843 | 0.033 | 0.007 |
| 844 | -0.046 | 0.080 |
| 845 | 0.028 | -0.014 |
| 846 | 0.023 | 0.137 |
| 847 | -0.063 | 0.018 |
| 848 | 0.019 | -0.031 |
| 849 | -0.044 | -0.090 |
| 850 | 0.009 | 0.159 |
| 851 | -0.007 | -0.074 |
| 852 | 0.003 | -0.009 |
| 853 | 0.006 | 0.005 |
| 854 | -0.013 | 0.071 |
| 855 | 0.044 | 0.077 |
| 856 | 0.042 | -0.007 |
| 857 | 0.034 | 0.081 |
| 858 | -0.008 | 0.137 |
| 859 | 0.063 | 0.126 |
| 860 | 0.034 | -0.091 |

| | | |
|---|---|---|
| 861 | -0.037 | 0.126 |
| 862 | 0.021 | 0.017 |
| 863 | -0.008 | -0.026 |
| 864 | -0.017 | 0.015 |
| 865 | 0.031 | 0.014 |
| 866 | -0.033 | 0.008 |
| 867 | -0.007 | 0.132 |
| 868 | 0.032 | 0.029 |
| 869 | -0.033 | -0.006 |
| 870 | 0.000 | 0.046 |
| 871 | -0.020 | 0.053 |
| 872 | -0.040 | -0.114 |
| 873 | 0.012 | 0.096 |
| 874 | 0.044 | 0.071 |
| 875 | 0.029 | -0.030 |
| 876 | 0.053 | -0.003 |
| 877 | -0.060 | 0.067 |
| 878 | -0.019 | -0.005 |
| 879 | 0.042 | -0.034 |
| 880 | -0.011 | 0.115 |
| 881 | -0.006 | 0.019 |
| 882 | 0.036 | 0.045 |
| 883 | 0.015 | 0.091 |
| 884 | 0.040 | -0.035 |
| 885 | -0.047 | 0.025 |
| 886 | -0.007 | 0.001 |
| 887 | 0.045 | 0.012 |
| 888 | -0.023 | -0.039 |
| 889 | -0.034 | -0.050 |
| 890 | -0.007 | -0.089 |
| 891 | 0.001 | -0.035 |
| 892 | -0.062 | 0.030 |
| 893 | -0.034 | -0.041 |
| 894 | -0.024 | -0.066 |
| 895 | 0.025 | -0.110 |
| 896 | -0.024 | 0.076 |
| 897 | 0.025 | 0.063 |
| 898 | 0.031 | 0.004 |
| 899 | -0.005 | -0.050 |
| 900 | -0.014 | 0.029 |
| 901 | -0.034 | 0.005 |
| 902 | 0.084 | 0.097 |
| 903 | -0.048 | 0.019 |
| 904 | -0.014 | 0.033 |
| 905 | -0.011 | -0.038 |
| 906 | -0.003 | -0.004 |
| 907 | 0.002 | -0.074 |
| 908 | -0.004 | -0.035 |
| 909 | -0.038 | -0.093 |
| 910 | -0.020 | 0.027 |
| 911 | 0.004 | 0.014 |

| | | |
|---|---|---|
| 912 | -0.019 | -0.035 |
| 913 | -0.010 | 0.025 |
| 914 | 0.033 | 0.066 |
| 915 | 0.054 | -0.018 |
| 916 | 0.010 | -0.030 |
| 917 | -0.011 | 0.087 |
| 918 | 0.017 | 0.046 |
| 919 | -0.022 | 0.106 |
| 920 | 0.010 | 0.085 |
| 921 | -0.030 | -0.115 |
| 922 | -0.021 | -0.047 |
| 923 | 0.000 | 0.007 |
| 924 | 0.059 | 0.028 |
| 925 | 0.039 | -0.001 |
| 926 | 0.035 | -0.029 |
| 927 | -0.011 | -0.193 |
| 928 | 0.025 | 0.055 |
| 929 | -0.018 | -0.130 |
| 930 | -0.109 | -0.095 |
| 931 | 0.022 | -0.027 |
| 932 | 0.086 | -0.008 |
| 933 | -0.058 | -0.060 |
| 934 | -0.005 | 0.044 |
| 935 | 0.023 | -0.055 |
| 936 | -0.023 | -0.046 |
| 937 | 0.007 | 0.005 |
| 938 | 0.014 | -0.072 |
| 939 | 0.001 | 0.068 |
| 940 | 0.019 | -0.060 |
| 941 | 0.014 | -0.056 |
| 942 | 0.011 | -0.033 |
| 943 | 0.045 | 0.023 |
| 944 | -0.011 | -0.050 |
| 945 | -0.052 | 0.008 |
| 946 | 0.073 | 0.035 |
| 947 | -0.045 | -0.044 |
| 948 | -0.044 | 0.106 |
| 949 | -0.053 | -0.039 |
| 950 | 0.003 | 0.040 |
| 951 | -0.046 | 0.065 |
| 952 | 0.029 | 0.043 |
| 953 | 0.022 | 0.000 |
| 954 | -0.031 | -0.014 |
| 955 | -0.003 | -0.094 |
| 956 | 0.011 | -0.006 |
| 957 | -0.027 | -0.196 |
| 958 | 0.032 | -0.035 |
| 959 | 0.010 | -0.141 |
| 960 | 0.017 | 0.100 |
| 961 | -0.055 | 0.201 |
| 962 | -0.010 | 0.051 |

| | | |
|---|---|---|
| 963 | 0.030 | -0.045 |
| 964 | 0.032 | 0.030 |
| 965 | -0.009 | -0.037 |
| 966 | -0.028 | -0.004 |
| 967 | -0.002 | 0.009 |
| 968 | 0.040 | 0.037 |
| 969 | 0.014 | 0.039 |
| 970 | -0.016 | 0.069 |
| 971 | 0.004 | 0.008 |
| 972 | 0.016 | 0.075 |
| 973 | -0.006 | -0.101 |
| 974 | 0.005 | -0.093 |
| 975 | -0.034 | 0.173 |
| 976 | -0.009 | 0.069 |
| 977 | 0.001 | -0.102 |
| 978 | -0.003 | 0.028 |
| 979 | 0.015 | -0.079 |
| 980 | -0.003 | -0.016 |
| 981 | -0.039 | 0.048 |
| 982 | 0.025 | -0.090 |
| 983 | -0.015 | -0.222 |
| 984 | 0.005 | 0.122 |
| 985 | 0.025 | 0.058 |
| 986 | -0.003 | -0.076 |
| 987 | 0.035 | -0.021 |
| 988 | 0.006 | 0.051 |
| 989 | 0.000 | 0.047 |
| 990 | -0.061 | 0.058 |
| 991 | -0.005 | 0.060 |
| 992 | 0.041 | 0.020 |
| 993 | 0.027 | 0.108 |
| 994 | -0.008 | 0.042 |
| 995 | -0.033 | 0.063 |
| 996 | -0.071 | 0.096 |
| 997 | 0.016 | 0.071 |
| 998 | -0.020 | 0.093 |
| 999 | -0.055 | 0.124 |
| 1000 | 0.014 | -0.065 |
| 1001 | 0.011 | -0.020 |
| 1002 | 0.092 | 0.000 |
| 1003 | -0.012 | -0.056 |
| 1004 | -0.030 | 0.074 |
| 1005 | 0.011 | -0.039 |
| 1006 | 0.036 | -0.038 |
| 1007 | 0.005 | 0.032 |
| 1008 | 0.039 | 0.033 |
| 1009 | 0.027 | -0.012 |
| 1010 | 0.057 | 0.050 |
| 1011 | -0.029 | -0.133 |
| 1012 | -0.043 | 0.075 |
| 1013 | -0.016 | 0.024 |

| | | |
|---|---|---|
| 1014 | -0.021 | -0.050 |
| 1015 | 0.005 | 0.031 |
| 1016 | -0.044 | -0.089 |
| 1017 | -0.025 | -0.007 |
| 1018 | 0.019 | 0.117 |
| 1019 | -0.069 | 0.125 |
| 1020 | 0.029 | -0.064 |
| 1021 | -0.015 | 0.019 |
| 1022 | -0.022 | 0.133 |
| 1023 | 0.019 | 0.049 |
| 1024 | 0.004 | 0.034 |
| 1025 | 0.040 | -0.012 |
| 1026 | -0.002 | 0.039 |
| 1027 | -0.007 | -0.058 |
| 1028 | 0.004 | -0.014 |
| 1029 | 0.018 | 0.078 |
| 1030 | -0.038 | -0.052 |
| 1031 | -0.022 | -0.042 |
| 1032 | -0.040 | 0.127 |
| 1033 | 0.001 | -0.014 |
| 1034 | -0.026 | -0.045 |
| 1035 | 0.032 | -0.089 |
| 1036 | -0.044 | -0.098 |
| 1037 | -0.001 | 0.029 |
| 1038 | 0.052 | 0.121 |
| 1039 | 0.014 | -0.017 |
| 1040 | -0.029 | 0.072 |
| 1041 | 0.000 | 0.027 |
| 1042 | 0.051 | 0.052 |
| 1043 | 0.019 | -0.026 |
| 1044 | -0.046 | 0.029 |
| 1045 | -0.020 | 0.029 |
| 1046 | 0.011 | 0.129 |
| 1047 | -0.019 | -0.121 |
| 1048 | -0.043 | 0.118 |
| 1049 | -0.033 | 0.164 |
| 1050 | 0.000 | -0.105 |
| 1051 | -0.027 | -0.178 |
| 1052 | 0.064 | 0.179 |
| 1053 | 0.005 | 0.013 |
| 1054 | 0.016 | -0.026 |
| 1055 | -0.023 | 0.009 |
| 1056 | 0.030 | 0.038 |
| 1057 | -0.018 | 0.114 |
| 1058 | 0.051 | -0.011 |
| 1059 | 0.073 | -0.160 |
| 1060 | 0.037 | -0.033 |
| 1061 | -0.014 | -0.016 |
| 1062 | 0.017 | -0.047 |
| 1063 | 0.042 | 0.116 |
| 1064 | 0.014 | 0.066 |

| | | |
|---|---|---|
| 1065 | 0.019 | -0.044 |
| 1066 | 0.005 | -0.003 |
| 1067 | 0.046 | 0.049 |
| 1068 | -0.035 | 0.005 |
| 1069 | 0.028 | 0.018 |
| 1070 | 0.061 | -0.025 |
| 1071 | 0.012 | -0.014 |
| 1072 | 0.028 | -0.047 |
| 1073 | -0.071 | 0.067 |
| 1074 | -0.014 | -0.084 |
| 1075 | 0.055 | -0.061 |
| 1076 | -0.074 | -0.070 |
| 1077 | 0.009 | -0.043 |
| 1078 | -0.041 | -0.043 |
| 1079 | -0.013 | 0.091 |
| 1080 | 0.006 | -0.012 |
| 1081 | 0.009 | -0.032 |
| 1082 | -0.039 | 0.028 |
| 1083 | -0.002 | -0.006 |
| 1084 | 0.005 | 0.027 |
| 1085 | 0.003 | -0.115 |
| 1086 | -0.042 | 0.006 |
| 1087 | -0.069 | 0.083 |
| 1088 | 0.023 | 0.098 |
| 1089 | -0.013 | -0.022 |
| 1090 | 0.011 | -0.055 |
| 1091 | -0.041 | -0.054 |
| 1092 | 0.039 | 0.013 |
| 1093 | -0.014 | -0.016 |
| 1094 | 0.061 | -0.006 |
| 1095 | -0.013 | 0.068 |
| 1096 | 0.026 | -0.034 |
| 1097 | -0.024 | 0.018 |
| 1098 | -0.001 | 0.072 |
| 1099 | 0.064 | 0.020 |
| 1100 | -0.030 | 0.043 |
| 1101 | -0.019 | 0.107 |
| 1102 | -0.054 | -0.072 |
| 1103 | -0.014 | 0.038 |
| 1104 | 0.003 | 0.014 |
| 1105 | 0.001 | 0.123 |
| 1106 | -0.061 | 0.024 |
| 1107 | 0.038 | 0.091 |
| 1108 | 0.007 | 0.069 |
| 1109 | 0.024 | 0.110 |
| 1110 | -0.006 | -0.166 |
| 1111 | 0.038 | -0.009 |
| 1112 | 0.049 | 0.037 |
| 1113 | 0.025 | -0.062 |
| 1114 | 0.032 | -0.099 |
| 1115 | 0.015 | 0.040 |

| | | |
|---|---|---|
| 1116 | 0.028 | -0.155 |
| 1117 | 0.015 | -0.035 |
| 1118 | -0.012 | -0.032 |
| 1119 | 0.026 | -0.032 |
| 1120 | -0.037 | -0.027 |
| 1121 | -0.014 | -0.034 |
| 1122 | 0.023 | 0.044 |
| 1123 | -0.040 | -0.102 |
| 1124 | -0.004 | -0.075 |
| 1125 | 0.028 | -0.019 |
| 1126 | -0.038 | 0.071 |
| 1127 | -0.017 | -0.046 |
| 1128 | 0.054 | 0.011 |
| 1129 | 0.040 | -0.039 |
| 1130 | -0.047 | -0.024 |
| 1131 | -0.010 | -0.103 |
| 1132 | -0.010 | 0.173 |
| 1133 | -0.045 | 0.072 |
| 1134 | 0.029 | -0.033 |
| 1135 | -0.067 | 0.010 |
| 1136 | -0.022 | 0.005 |
| 1137 | 0.044 | 0.044 |
| 1138 | 0.008 | -0.019 |
| 1139 | 0.011 | 0.065 |
| 1140 | -0.005 | -0.040 |
| 1141 | -0.009 | 0.119 |
| 1142 | -0.020 | 0.033 |
| 1143 | -0.040 | 0.078 |
| 1144 | -0.016 | -0.164 |
| 1145 | 0.015 | -0.205 |
| 1146 | 0.020 | 0.017 |
| 1147 | -0.014 | -0.065 |
| 1148 | 0.085 | 0.019 |
| 1149 | 0.006 | 0.001 |
| 1150 | -0.036 | -0.112 |
| 1151 | -0.058 | -0.100 |
| 1152 | 0.003 | 0.115 |
| 1153 | -0.014 | -0.033 |
| 1154 | -0.039 | 0.044 |
| 1155 | 0.010 | 0.005 |
| 1156 | -0.089 | -0.035 |
| 1157 | -0.036 | 0.021 |
| 1158 | -0.020 | -0.073 |
| 1159 | -0.058 | 0.030 |
| 1160 | -0.033 | -0.039 |
| 1161 | -0.020 | 0.115 |
| 1162 | 0.012 | -0.062 |
| 1163 | 0.035 | -0.083 |
| 1164 | -0.077 | -0.039 |
| 1165 | -0.002 | -0.066 |
| 1166 | 0.082 | 0.012 |

| | | |
|---|---|---|
| 1167 | -0.007 | -0.093 |
| 1168 | -0.016 | -0.015 |
| 1169 | 0.067 | -0.055 |
| 1170 | 0.017 | 0.120 |
| 1171 | -0.059 | 0.102 |
| 1172 | 0.014 | -0.002 |
| 1173 | 0.009 | 0.042 |
| 1174 | -0.022 | 0.054 |
| 1175 | -0.054 | -0.031 |
| 1176 | -0.036 | -0.001 |
| 1177 | -0.026 | -0.004 |
| 1178 | 0.023 | 0.091 |
| 1179 | -0.043 | -0.040 |
| 1180 | 0.034 | -0.044 |
| 1181 | -0.048 | 0.053 |
| 1182 | -0.011 | 0.032 |
| 1183 | -0.017 | -0.089 |
| 1184 | -0.001 | -0.017 |
| 1185 | 0.016 | -0.123 |
| 1186 | 0.000 | 0.067 |
| 1187 | 0.004 | 0.148 |
| 1188 | 0.061 | -0.045 |
| 1189 | -0.014 | -0.032 |
| 1190 | -0.062 | -0.005 |
| 1191 | -0.037 | 0.030 |
| 1192 | -0.007 | 0.082 |
| 1193 | 0.040 | -0.002 |
| 1194 | -0.020 | -0.020 |
| 1195 | 0.012 | -0.069 |
| 1196 | -0.020 | -0.053 |
| 1197 | 0.025 | 0.051 |
| 1198 | 0.007 | -0.044 |
| 1199 | 0.006 | 0.070 |
| 1200 | 0.003 | -0.039 |
| 1201 | -0.008 | 0.014 |
| 1202 | -0.039 | 0.038 |
| 1203 | 0.082 | -0.136 |
| 1204 | 0.086 | 0.187 |
| 1205 | -0.046 | 0.081 |
| 1206 | -0.036 | 0.086 |
| 1207 | -0.012 | 0.065 |
| 1208 | -0.047 | 0.147 |
| 1209 | 0.032 | -0.034 |
| 1210 | 0.019 | 0.104 |
| 1211 | -0.010 | 0.028 |
| 1212 | -0.049 | 0.068 |
| 1213 | 0.046 | 0.049 |
| 1214 | -0.029 | -0.107 |
| 1215 | -0.017 | -0.144 |
| 1216 | -0.047 | -0.118 |
| 1217 | 0.027 | 0.045 |

| | | |
|---|---|---|
| 1218 | 0.019 | 0.010 |
| 1219 | -0.040 | 0.026 |
| 1220 | 0.033 | 0.039 |
| 1221 | 0.015 | -0.058 |
| 1222 | 0.053 | -0.044 |
| 1223 | 0.033 | 0.061 |
| 1224 | 0.028 | -0.043 |
| 1225 | -0.050 | 0.074 |
| 1226 | 0.012 | -0.135 |
| 1227 | -0.043 | 0.074 |
| 1228 | 0.043 | 0.053 |
| 1229 | 0.025 | -0.148 |
| 1230 | -0.017 | -0.083 |
| 1231 | 0.002 | -0.029 |
| 1232 | -0.077 | 0.062 |
| 1233 | -0.057 | 0.027 |
| 1234 | -0.046 | -0.053 |
| 1235 | 0.032 | 0.070 |
| 1236 | 0.009 | -0.071 |
| 1237 | -0.008 | -0.058 |
| 1238 | -0.012 | -0.058 |
| 1239 | 0.062 | 0.038 |
| 1240 | -0.035 | 0.041 |
| 1241 | -0.004 | -0.094 |
| 1242 | 0.016 | 0.088 |
| 1243 | 0.028 | 0.054 |
| 1244 | -0.019 | -0.031 |
| 1245 | 0.003 | -0.071 |
| 1246 | 0.012 | -0.056 |
| 1247 | 0.024 | 0.104 |
| 1248 | -0.036 | -0.024 |
| 1249 | -0.013 | 0.020 |
| 1250 | 0.004 | -0.029 |
| 1251 | -0.004 | -0.021 |
| 1252 | -0.010 | 0.010 |
| 1253 | 0.035 | 0.066 |
| 1254 | 0.001 | 0.065 |
| 1255 | -0.030 | -0.012 |
| 1256 | -0.076 | 0.078 |
| 1257 | 0.021 | -0.013 |
| 1258 | -0.016 | -0.082 |
| 1259 | -0.005 | -0.133 |
| 1260 | -0.026 | -0.070 |
| 1261 | 0.025 | -0.074 |
| 1262 | -0.042 | 0.107 |
| 1263 | 0.019 | 0.081 |
| 1264 | -0.050 | -0.070 |
| 1265 | -0.030 | 0.077 |
| 1266 | 0.048 | 0.007 |
| 1267 | -0.002 | -0.101 |
| 1268 | 0.019 | -0.041 |

| | | |
|---|---|---|
| 1269 | -0.037 | 0.079 |
| 1270 | -0.042 | -0.016 |
| 1271 | -0.012 | -0.039 |
| 1272 | 0.040 | -0.037 |
| 1273 | 0.000 | -0.137 |
| 1274 | 0.037 | 0.079 |
| 1275 | -0.028 | 0.091 |
| 1276 | -0.019 | 0.022 |
| 1277 | -0.046 | -0.009 |
| 1278 | -0.020 | -0.090 |
| 1279 | 0.032 | -0.005 |
| 1280 | -0.024 | -0.068 |
| 1281 | 0.014 | 0.099 |
| 1282 | 0.032 | -0.024 |
| 1283 | -0.053 | 0.152 |
| 1284 | 0.029 | -0.099 |
| 1285 | -0.061 | -0.077 |
| 1286 | 0.029 | -0.014 |
| 1287 | 0.047 | 0.087 |
| 1288 | 0.051 | 0.115 |
| 1289 | 0.006 | -0.087 |
| 1290 | 0.024 | -0.203 |
| 1291 | -0.005 | 0.003 |
| 1292 | -0.030 | 0.062 |
| 1293 | -0.022 | 0.039 |
| 1294 | -0.024 | -0.123 |
| 1295 | -0.009 | -0.042 |
| 1296 | -0.017 | -0.002 |
| 1297 | 0.025 | -0.073 |
| 1298 | -0.023 | 0.164 |
| 1299 | -0.039 | -0.112 |
| 1300 | 0.004 | 0.070 |
| 1301 | 0.010 | 0.039 |
| 1302 | 0.063 | 0.039 |
| 1303 | -0.029 | 0.052 |
| 1304 | -0.037 | -0.121 |
| 1305 | -0.008 | 0.124 |
| 1306 | -0.016 | -0.007 |
| 1307 | 0.038 | -0.013 |
| 1308 | -0.028 | -0.083 |
| 1309 | -0.011 | -0.030 |
| 1310 | 0.011 | -0.027 |
| 1311 | -0.029 | 0.092 |
| 1312 | 0.023 | 0.055 |
| 1313 | -0.032 | 0.072 |
| 1314 | 0.036 | -0.030 |
| 1315 | -0.022 | 0.062 |
| 1316 | 0.018 | 0.063 |
| 1317 | 0.037 | -0.122 |
| 1318 | -0.032 | -0.032 |
| 1319 | -0.012 | -0.020 |

| | | |
|---|---|---|
| 1320 | -0.015 | 0.005 |
| 1321 | -0.007 | -0.046 |
| 1322 | 0.000 | -0.047 |
| 1323 | -0.043 | 0.061 |
| 1324 | 0.005 | -0.060 |
| 1325 | -0.025 | -0.079 |
| 1326 | 0.044 | -0.090 |
| 1327 | 0.001 | 0.015 |
| 1328 | -0.026 | -0.033 |
| 1329 | -0.012 | -0.097 |
| 1330 | 0.000 | 0.033 |
| 1331 | 0.004 | 0.134 |
| 1332 | -0.008 | 0.017 |
| 1333 | -0.014 | 0.038 |
| 1334 | 0.029 | 0.067 |
| 1335 | -0.009 | -0.053 |
| 1336 | -0.019 | 0.142 |
| 1337 | -0.015 | 0.074 |
| 1338 | -0.025 | 0.053 |
| 1339 | -0.002 | -0.046 |
| 1340 | -0.056 | 0.103 |
| 1341 | 0.023 | 0.022 |
| 1342 | -0.012 | 0.009 |
| 1343 | -0.021 | 0.054 |
| 1344 | 0.017 | 0.001 |
| 1345 | 0.007 | -0.046 |
| 1346 | 0.020 | -0.054 |
| 1347 | -0.028 | 0.142 |
| 1348 | 0.028 | -0.004 |
| 1349 | -0.031 | 0.027 |
| 1350 | -0.031 | 0.116 |
| 1351 | 0.026 | 0.181 |
| 1352 | 0.001 | 0.065 |
| 1353 | 0.007 | -0.105 |
| 1354 | -0.086 | 0.024 |
| 1355 | -0.042 | 0.106 |
| 1356 | 0.018 | -0.106 |
| 1357 | -0.025 | -0.007 |
| 1358 | -0.045 | -0.018 |
| 1359 | -0.008 | 0.156 |
| 1360 | -0.009 | 0.080 |
| 1361 | -0.005 | -0.020 |
| 1362 | 0.029 | -0.197 |
| 1363 | -0.017 | 0.029 |
| 1364 | -0.112 | 0.100 |
| 1365 | -0.005 | 0.070 |
| 1366 | -0.036 | -0.091 |
| 1367 | -0.055 | 0.038 |
| 1368 | 0.052 | -0.144 |
| 1369 | 0.030 | -0.024 |
| 1370 | 0.043 | -0.023 |

| | | |
|---|---|---|
| 1371 | 0.025 | 0.001 |
| 1372 | -0.065 | 0.048 |
| 1373 | 0.000 | 0.023 |
| 1374 | 0.007 | 0.050 |
| 1375 | -0.002 | 0.056 |
| 1376 | -0.033 | -0.031 |
| 1377 | 0.002 | 0.038 |
| 1378 | 0.029 | 0.049 |
| 1379 | 0.016 | -0.110 |
| 1380 | -0.011 | 0.047 |
| 1381 | 0.006 | 0.080 |
| 1382 | -0.023 | 0.114 |
| 1383 | -0.063 | 0.012 |
| 1384 | 0.004 | -0.018 |
| 1385 | 0.010 | -0.014 |
| 1386 | -0.007 | 0.140 |
| 1387 | 0.063 | -0.030 |
| 1388 | 0.002 | -0.041 |
| 1389 | -0.027 | 0.105 |
| 1390 | -0.050 | 0.030 |
| 1391 | 0.034 | -0.045 |
| 1392 | -0.027 | -0.083 |
| 1393 | 0.024 | -0.056 |
| 1394 | 0.051 | -0.003 |
| 1395 | -0.007 | -0.011 |
| 1396 | 0.000 | -0.037 |
| 1397 | 0.005 | -0.008 |
| 1398 | 0.022 | -0.005 |
| 1399 | 0.000 | -0.125 |
| 1400 | 0.006 | -0.036 |
| 1401 | 0.017 | 0.085 |
| 1402 | -0.012 | 0.023 |
| 1403 | -0.029 | -0.055 |
| 1404 | -0.027 | 0.037 |
| 1405 | -0.028 | 0.026 |
| 1406 | 0.055 | 0.017 |
| 1407 | -0.006 | 0.015 |
| 1408 | 0.007 | 0.082 |
| 1409 | 0.025 | -0.048 |
| 1410 | 0.014 | 0.089 |
| 1411 | -0.048 | -0.085 |
| 1412 | -0.014 | -0.072 |
| 1413 | -0.023 | -0.050 |
| 1414 | 0.012 | 0.005 |
| 1415 | -0.050 | -0.036 |
| 1416 | 0.056 | 0.154 |
| 1417 | -0.035 | -0.039 |
| 1418 | -0.008 | 0.034 |
| 1419 | 0.033 | 0.047 |
| 1420 | 0.049 | -0.092 |
| 1421 | 0.001 | 0.012 |

| | | |
|---|---|---|
| 1422 | 0.003 | 0.130 |
| 1423 | -0.002 | 0.050 |
| 1424 | 0.024 | 0.132 |
| 1425 | 0.009 | 0.109 |
| 1426 | -0.042 | 0.042 |
| 1427 | -0.007 | -0.171 |
| 1428 | 0.015 | 0.010 |
| 1429 | -0.015 | -0.080 |
| 1430 | -0.021 | 0.038 |
| 1431 | -0.036 | 0.088 |
| 1432 | -0.008 | -0.011 |
| 1433 | 0.001 | 0.010 |
| 1434 | 0.001 | 0.046 |
| 1435 | 0.001 | -0.039 |
| 1436 | 0.010 | -0.008 |
| 1437 | -0.045 | 0.059 |
| 1438 | 0.013 | 0.020 |
| 1439 | 0.020 | 0.122 |
| 1440 | 0.038 | 0.113 |
| 1441 | -0.031 | 0.008 |
| 1442 | 0.023 | -0.058 |
| 1443 | 0.000 | -0.003 |
| 1444 | 0.010 | -0.065 |
| 1445 | -0.045 | 0.089 |
| 1446 | 0.017 | -0.054 |
| 1447 | 0.051 | 0.034 |
| 1448 | 0.018 | 0.027 |
| 1449 | -0.010 | -0.140 |
| 1450 | 0.004 | 0.079 |
| 1451 | 0.007 | 0.013 |
| 1452 | -0.051 | 0.077 |
| 1453 | 0.020 | -0.072 |
| 1454 | -0.011 | -0.128 |
| 1455 | -0.006 | -0.034 |
| 1456 | -0.016 | -0.063 |
| 1457 | -0.032 | 0.000 |
| 1458 | 0.097 | -0.091 |
| 1459 | 0.037 | -0.022 |
| 1460 | 0.005 | 0.012 |
| 1461 | -0.017 | 0.008 |
| 1462 | 0.003 | -0.011 |
| 1463 | 0.019 | 0.068 |
| 1464 | -0.005 | 0.140 |
| 1465 | 0.007 | -0.048 |
| 1466 | 0.017 | 0.068 |
| 1467 | 0.031 | 0.039 |
| 1468 | -0.002 | 0.052 |
| 1469 | 0.006 | -0.116 |
| 1470 | 0.019 | -0.106 |
| 1471 | -0.003 | 0.014 |
| 1472 | 0.016 | 0.103 |

| | | |
|---|---|---|
| 1473 | -0.059 | 0.007 |
| 1474 | -0.012 | 0.110 |
| 1475 | 0.008 | 0.066 |
| 1476 | 0.031 | 0.029 |
| 1477 | -0.002 | 0.031 |
| 1478 | -0.016 | -0.034 |
| 1479 | -0.063 | -0.036 |
| 1480 | 0.036 | 0.066 |
| 1481 | -0.020 | -0.039 |
| 1482 | -0.038 | 0.031 |
| 1483 | 0.019 | 0.143 |
| 1484 | -0.017 | -0.047 |
| 1485 | 0.021 | -0.070 |
| 1486 | -0.019 | -0.072 |
| 1487 | 0.009 | -0.155 |
| 1488 | 0.031 | 0.010 |
| 1489 | -0.006 | -0.029 |
| 1490 | -0.065 | 0.039 |
| 1491 | 0.011 | 0.020 |
| 1492 | 0.015 | 0.124 |
| 1493 | 0.035 | -0.018 |
| 1494 | 0.005 | 0.042 |
| 1495 | 0.001 | 0.118 |
| 1496 | -0.020 | -0.056 |
| 1497 | 0.008 | -0.076 |
| 1498 | -0.046 | -0.028 |
| 1499 | -0.007 | 0.001 |
| 1500 | 0.016 | -0.025 |
| 1501 | -0.065 | -0.058 |
| 1502 | -0.040 | 0.068 |
| 1503 | 0.004 | 0.081 |
| 1504 | -0.065 | 0.031 |
| 1505 | 0.000 | 0.012 |
| 1506 | 0.015 | -0.043 |
| 1507 | -0.032 | 0.187 |
| 1508 | 0.009 | -0.054 |
| 1509 | 0.033 | 0.027 |
| 1510 | 0.047 | 0.045 |
| 1511 | 0.012 | 0.053 |
| 1512 | -0.014 | 0.032 |
| 1513 | -0.009 | -0.020 |
| 1514 | -0.075 | 0.011 |
| 1515 | -0.012 | 0.056 |
| 1516 | -0.006 | 0.058 |
| 1517 | 0.039 | 0.001 |
| 1518 | -0.028 | 0.087 |
| 1519 | 0.027 | 0.127 |
| 1520 | 0.015 | 0.109 |
| 1521 | 0.025 | 0.016 |
| 1522 | -0.043 | 0.039 |
| 1523 | -0.003 | 0.042 |

| | | |
|---|---|---|
| 1524 | 0.017 | -0.074 |
| 1525 | -0.048 | 0.065 |
| 1526 | 0.034 | -0.016 |
| 1527 | -0.007 | -0.031 |
| 1528 | -0.041 | 0.022 |
| 1529 | -0.022 | -0.107 |
| 1530 | 0.040 | 0.055 |
| 1531 | 0.008 | -0.065 |
| 1532 | -0.010 | -0.127 |
| 1533 | 0.081 | -0.086 |
| 1534 | -0.067 | -0.048 |
| 1535 | -0.015 | -0.035 |
| 1536 | 0.004 | 0.149 |
| 1537 | -0.040 | -0.006 |
| 1538 | -0.083 | -0.041 |
| 1539 | -0.003 | 0.126 |
| 1540 | -0.034 | -0.047 |
| 1541 | -0.024 | 0.037 |
| 1542 | -0.012 | -0.026 |
| 1543 | 0.031 | -0.123 |
| 1544 | -0.020 | -0.052 |
| 1545 | 0.040 | 0.004 |
| 1546 | -0.058 | 0.054 |
| 1547 | 0.029 | -0.025 |
| 1548 | -0.042 | 0.018 |
| 1549 | 0.043 | 0.021 |
| 1550 | -0.020 | -0.084 |
| 1551 | -0.054 | 0.117 |
| 1552 | -0.033 | -0.009 |
| 1553 | 0.019 | 0.055 |
| 1554 | -0.015 | 0.051 |
| 1555 | -0.009 | 0.078 |
| 1556 | -0.055 | -0.030 |
| 1557 | 0.043 | 0.119 |
| 1558 | 0.016 | 0.007 |
| 1559 | -0.027 | 0.004 |
| 1560 | -0.026 | 0.034 |
| 1561 | -0.030 | -0.019 |
| 1562 | -0.052 | -0.183 |
| 1563 | -0.065 | 0.035 |
| 1564 | 0.009 | 0.009 |
| 1565 | 0.011 | 0.098 |
| 1566 | 0.025 | -0.051 |
| 1567 | 0.048 | 0.046 |
| 1568 | 0.031 | -0.015 |
| 1569 | -0.025 | -0.073 |
| 1570 | 0.036 | 0.036 |
| 1571 | 0.000 | 0.147 |
| 1572 | 0.015 | 0.021 |
| 1573 | 0.016 | -0.001 |
| 1574 | -0.040 | 0.077 |

| | | |
|---|---|---|
| 1575 | 0.013 | -0.011 |
| 1576 | -0.033 | -0.099 |
| 1577 | 0.026 | -0.036 |
| 1578 | 0.029 | -0.033 |
| 1579 | -0.007 | -0.007 |
| 1580 | -0.053 | -0.052 |
| 1581 | 0.032 | 0.044 |
| 1582 | -0.003 | 0.011 |
| 1583 | -0.066 | 0.121 |
| 1584 | -0.103 | -0.055 |
| 1585 | -0.018 | -0.032 |
| 1586 | 0.010 | -0.144 |
| 1587 | 0.001 | 0.064 |
| 1588 | 0.038 | -0.020 |
| 1589 | 0.003 | 0.016 |
| 1590 | -0.010 | 0.131 |
| 1591 | 0.011 | -0.119 |
| 1592 | -0.058 | 0.013 |
| 1593 | -0.068 | 0.080 |
| 1594 | 0.062 | -0.022 |
| 1595 | -0.008 | 0.009 |
| 1596 | -0.024 | 0.082 |
| 1597 | -0.073 | -0.054 |
| 1598 | 0.020 | -0.038 |
| 1599 | -0.027 | 0.007 |
| 1600 | 0.036 | 0.007 |
| 1601 | 0.021 | -0.071 |
| 1602 | 0.026 | 0.072 |
| 1603 | -0.012 | -0.064 |
| 1604 | -0.030 | 0.017 |
| 1605 | -0.008 | -0.118 |
| 1606 | 0.021 | 0.026 |
| 1607 | 0.083 | 0.047 |
| 1608 | -0.002 | 0.065 |
| 1609 | 0.052 | -0.055 |
| 1610 | 0.075 | 0.063 |
| 1611 | 0.027 | -0.033 |
| 1612 | 0.035 | -0.015 |
| 1613 | -0.074 | -0.121 |
| 1614 | 0.063 | 0.013 |
| 1615 | 0.005 | -0.020 |
| 1616 | 0.050 | 0.075 |
| 1617 | 0.002 | 0.012 |
| 1618 | 0.009 | 0.046 |
| 1619 | -0.005 | 0.000 |
| 1620 | -0.036 | 0.022 |
| 1621 | -0.007 | 0.122 |
| 1622 | 0.023 | 0.118 |
| 1623 | -0.032 | 0.088 |
| 1624 | 0.006 | 0.031 |
| 1625 | 0.044 | -0.019 |

| | | |
|---|---|---|
| 1626 | 0.025 | -0.107 |
| 1627 | -0.012 | -0.092 |
| 1628 | -0.010 | -0.123 |
| 1629 | -0.005 | 0.037 |
| 1630 | -0.023 | -0.083 |
| 1631 | -0.033 | 0.060 |
| 1632 | 0.073 | -0.018 |
| 1633 | -0.005 | -0.053 |
| 1634 | -0.048 | 0.053 |
| 1635 | 0.002 | -0.074 |
| 1636 | 0.007 | -0.085 |
| 1637 | 0.077 | -0.056 |
| 1638 | 0.071 | -0.053 |
| 1639 | 0.001 | 0.030 |
| 1640 | 0.008 | -0.066 |
| 1641 | 0.012 | -0.145 |
| 1642 | 0.001 | -0.114 |
| 1643 | 0.030 | -0.044 |
| 1644 | -0.084 | -0.097 |
| 1645 | -0.004 | -0.008 |
| 1646 | 0.050 | -0.002 |
| 1647 | -0.014 | -0.036 |
| 1648 | -0.065 | -0.116 |
| 1649 | -0.031 | 0.028 |
| 1650 | -0.042 | 0.051 |
| 1651 | 0.033 | -0.095 |
| 1652 | -0.033 | -0.115 |
| 1653 | 0.015 | -0.069 |
| 1654 | 0.029 | 0.059 |
| 1655 | -0.005 | 0.108 |
| 1656 | 0.023 | -0.054 |
| 1657 | 0.033 | 0.003 |
| 1658 | -0.021 | 0.059 |
| 1659 | 0.001 | 0.010 |
| 1660 | -0.038 | -0.058 |
| 1661 | -0.011 | -0.022 |
| 1662 | 0.032 | 0.017 |
| 1663 | 0.030 | -0.061 |
| 1664 | -0.008 | 0.013 |
| 1665 | 0.012 | 0.050 |
| 1666 | 0.018 | 0.056 |
| 1667 | 0.042 | -0.094 |
| 1668 | -0.046 | -0.061 |
| 1669 | 0.017 | 0.033 |
| 1670 | -0.009 | 0.043 |
| 1671 | 0.027 | 0.005 |
| 1672 | -0.005 | 0.029 |
| 1673 | 0.006 | -0.076 |
| 1674 | -0.019 | -0.013 |
| 1675 | 0.010 | -0.036 |
| 1676 | 0.035 | 0.040 |

| | | |
|---|---|---|
| 1677 | -0.010 | 0.100 |
| 1678 | 0.036 | -0.021 |
| 1679 | -0.035 | 0.079 |
| 1680 | 0.029 | -0.109 |
| 1681 | 0.051 | -0.108 |
| 1682 | 0.016 | -0.034 |
| 1683 | 0.018 | -0.016 |
| 1684 | -0.054 | 0.036 |
| 1685 | -0.010 | -0.087 |
| 1686 | 0.049 | 0.079 |
| 1687 | -0.033 | -0.064 |
| 1688 | -0.080 | -0.106 |
| 1689 | -0.007 | 0.133 |
| 1690 | -0.018 | 0.013 |
| 1691 | 0.030 | 0.129 |
| 1692 | 0.021 | -0.078 |
| 1693 | 0.029 | -0.072 |
| 1694 | 0.056 | 0.058 |
| 1695 | -0.014 | 0.055 |
| 1696 | 0.036 | -0.091 |
| 1697 | -0.040 | 0.092 |
| 1698 | 0.031 | 0.054 |
| 1699 | -0.034 | -0.044 |
| 1700 | 0.047 | -0.175 |
| 1701 | 0.025 | 0.104 |
| 1702 | 0.009 | -0.002 |
| 1703 | -0.037 | -0.138 |
| 1704 | -0.049 | -0.037 |
| 1705 | 0.042 | 0.043 |
| 1706 | -0.057 | -0.055 |
| 1707 | -0.004 | -0.045 |
| 1708 | -0.055 | -0.058 |
| 1709 | -0.014 | 0.015 |
| 1710 | -0.024 | 0.022 |
| 1711 | -0.020 | 0.043 |
| 1712 | 0.023 | 0.021 |
| 1713 | 0.023 | 0.103 |
| 1714 | -0.054 | -0.040 |
| 1715 | 0.048 | 0.104 |
| 1716 | 0.015 | -0.157 |
| 1717 | 0.024 | -0.026 |
| 1718 | -0.035 | 0.111 |
| 1719 | -0.057 | 0.081 |
| 1720 | -0.008 | -0.096 |
| 1721 | -0.019 | -0.062 |
| 1722 | -0.037 | 0.144 |
| 1723 | 0.007 | 0.045 |
| 1724 | 0.029 | 0.020 |
| 1725 | 0.009 | 0.034 |
| 1726 | -0.049 | 0.195 |
| 1727 | -0.024 | 0.108 |

| | | |
|---|---|---|
| 1728 | 0.031 | -0.005 |
| 1729 | 0.053 | -0.059 |
| 1730 | -0.074 | -0.089 |
| 1731 | 0.031 | -0.093 |
| 1732 | -0.032 | 0.041 |
| 1733 | -0.004 | 0.180 |
| 1734 | 0.023 | -0.005 |
| 1735 | 0.006 | 0.040 |
| 1736 | 0.030 | 0.057 |
| 1737 | -0.026 | 0.108 |
| 1738 | -0.001 | 0.081 |
| 1739 | -0.053 | -0.001 |
| 1740 | 0.018 | -0.053 |
| 1741 | -0.042 | -0.010 |
| 1742 | -0.025 | -0.036 |
| 1743 | 0.000 | -0.040 |
| 1744 | -0.018 | 0.002 |
| 1745 | -0.050 | 0.047 |
| 1746 | -0.019 | 0.001 |
| 1747 | -0.025 | 0.021 |
| 1748 | 0.001 | -0.042 |
| 1749 | 0.024 | -0.123 |
| 1750 | 0.008 | 0.110 |
| 1751 | 0.001 | 0.011 |
| 1752 | -0.059 | -0.029 |
| 1753 | 0.027 | 0.037 |
| 1754 | 0.003 | 0.088 |
| 1755 | -0.014 | -0.052 |
| 1756 | 0.033 | -0.133 |
| 1757 | -0.013 | 0.005 |
| 1758 | -0.014 | -0.051 |
| 1759 | 0.010 | -0.086 |
| 1760 | -0.022 | 0.115 |
| 1761 | -0.045 | 0.096 |
| 1762 | -0.004 | -0.166 |
| 1763 | 0.001 | -0.078 |
| 1764 | -0.007 | -0.040 |
| 1765 | 0.021 | -0.034 |
| 1766 | -0.058 | -0.077 |
| 1767 | 0.000 | -0.035 |
| 1768 | 0.002 | 0.081 |
| 1769 | 0.056 | -0.141 |
| 1770 | 0.004 | -0.078 |
| 1771 | -0.026 | 0.096 |
| 1772 | 0.002 | -0.022 |
| 1773 | 0.005 | -0.060 |
| 1774 | 0.033 | -0.067 |
| 1775 | -0.015 | -0.051 |
| 1776 | 0.011 | 0.052 |
| 1777 | -0.019 | 0.044 |
| 1778 | 0.067 | 0.046 |

| | | |
|---|---|---|
| 1779 | 0.013 | 0.047 |
| 1780 | 0.041 | 0.121 |
| 1781 | -0.007 | 0.031 |
| 1782 | -0.049 | 0.019 |
| 1783 | 0.002 | -0.047 |
| 1784 | -0.054 | 0.082 |
| 1785 | -0.037 | -0.121 |
| 1786 | -0.009 | 0.042 |
| 1787 | 0.050 | -0.078 |
| 1788 | 0.099 | -0.037 |
| 1789 | -0.006 | -0.101 |
| 1790 | 0.001 | 0.006 |
| 1791 | -0.008 | -0.047 |
| 1792 | 0.027 | 0.063 |
| 1793 | 0.010 | -0.003 |
| 1794 | -0.054 | 0.013 |
| 1795 | -0.057 | 0.009 |
| 1796 | 0.033 | -0.014 |
| 1797 | -0.016 | 0.036 |
| 1798 | -0.040 | -0.092 |
| 1799 | 0.001 | -0.063 |
| 1800 | 0.006 | 0.037 |
| 1801 | 0.013 | 0.003 |
| 1802 | 0.062 | -0.090 |
| 1803 | 0.047 | 0.038 |
| 1804 | 0.004 | 0.088 |
| 1805 | 0.024 | -0.037 |
| 1806 | 0.037 | 0.008 |
| 1807 | 0.007 | 0.058 |
| 1808 | 0.000 | -0.114 |
| 1809 | 0.053 | -0.031 |
| 1810 | -0.041 | 0.088 |
| 1811 | 0.037 | 0.048 |
| 1812 | -0.016 | 0.041 |
| 1813 | -0.015 | -0.010 |
| 1814 | -0.028 | -0.029 |
| 1815 | 0.017 | 0.067 |
| 1816 | -0.063 | -0.053 |
| 1817 | 0.018 | -0.044 |
| 1818 | -0.027 | -0.022 |
| 1819 | 0.009 | -0.129 |
| 1820 | 0.026 | -0.041 |
| 1821 | -0.044 | 0.121 |
| 1822 | 0.029 | 0.031 |
| 1823 | 0.003 | 0.012 |
| 1824 | -0.076 | 0.093 |
| 1825 | 0.064 | -0.137 |
| 1826 | -0.039 | -0.052 |
| 1827 | 0.037 | -0.003 |
| 1828 | -0.042 | -0.045 |
| 1829 | -0.040 | 0.081 |

| | | |
|---|---|---|
| 1830 | -0.033 | 0.013 |
| 1831 | 0.013 | 0.083 |
| 1832 | -0.025 | 0.052 |
| 1833 | -0.035 | -0.047 |
| 1834 | -0.009 | 0.073 |
| 1835 | -0.005 | -0.054 |
| 1836 | 0.053 | -0.070 |
| 1837 | 0.034 | 0.012 |
| 1838 | 0.038 | -0.071 |
| 1839 | 0.044 | -0.100 |
| 1840 | 0.005 | 0.021 |
| 1841 | 0.035 | 0.015 |
| 1842 | 0.024 | -0.028 |
| 1843 | 0.019 | -0.002 |
| 1844 | 0.012 | 0.057 |
| 1845 | -0.001 | -0.061 |
| 1846 | 0.022 | -0.057 |
| 1847 | -0.040 | -0.112 |
| 1848 | 0.028 | -0.015 |
| 1849 | 0.057 | 0.011 |
| 1850 | 0.075 | -0.060 |
| 1851 | 0.053 | 0.030 |
| 1852 | 0.067 | -0.020 |
| 1853 | -0.006 | -0.073 |
| 1854 | -0.061 | -0.094 |
| 1855 | -0.011 | 0.137 |
| 1856 | 0.009 | 0.104 |
| 1857 | 0.022 | -0.013 |
| 1858 | -0.079 | -0.006 |
| 1859 | -0.004 | 0.063 |
| 1860 | 0.040 | 0.158 |
| 1861 | 0.041 | -0.069 |
| 1862 | -0.015 | -0.039 |
| 1863 | 0.005 | -0.028 |
| 1864 | -0.025 | 0.028 |
| 1865 | -0.050 | -0.010 |
| 1866 | -0.069 | 0.017 |
| 1867 | 0.018 | -0.018 |
| 1868 | -0.054 | 0.028 |
| 1869 | 0.049 | 0.042 |
| 1870 | 0.048 | 0.067 |
| 1871 | 0.056 | 0.002 |
| 1872 | -0.021 | 0.111 |
| 1873 | -0.046 | 0.028 |
| 1874 | -0.071 | -0.108 |
| 1875 | 0.012 | 0.132 |
| 1876 | -0.052 | -0.120 |
| 1877 | 0.016 | -0.040 |
| 1878 | -0.032 | -0.041 |
| 1879 | -0.013 | 0.001 |
| 1880 | 0.025 | -0.001 |

| | | |
|---|---|---|
| 1881 | 0.047 | 0.076 |
| 1882 | -0.029 | -0.054 |
| 1883 | 0.049 | -0.202 |
| 1884 | 0.000 | -0.076 |
| 1885 | -0.012 | -0.010 |
| 1886 | 0.003 | -0.097 |
| 1887 | 0.012 | -0.006 |
| 1888 | 0.044 | -0.042 |
| 1889 | 0.026 | 0.097 |
| 1890 | 0.060 | 0.039 |
| 1891 | 0.023 | 0.050 |
| 1892 | 0.070 | 0.006 |
| 1893 | -0.003 | 0.061 |
| 1894 | -0.055 | 0.087 |
| 1895 | -0.027 | -0.023 |
| 1896 | 0.085 | 0.042 |
| 1897 | -0.030 | 0.025 |
| 1898 | 0.033 | 0.001 |
| 1899 | 0.020 | 0.084 |
| 1900 | -0.046 | 0.020 |
| 1901 | 0.083 | -0.030 |
| 1902 | -0.034 | 0.027 |
| 1903 | 0.048 | 0.008 |
| 1904 | -0.051 | 0.011 |
| 1905 | 0.011 | 0.083 |
| 1906 | 0.002 | 0.001 |
| 1907 | 0.062 | -0.081 |
| 1908 | -0.018 | 0.029 |
| 1909 | -0.027 | -0.002 |
| 1910 | -0.045 | -0.006 |
| 1911 | -0.006 | -0.109 |
| 1912 | 0.018 | 0.017 |
| 1913 | 0.024 | -0.050 |
| 1914 | 0.024 | 0.026 |
| 1915 | -0.047 | -0.104 |
| 1916 | 0.016 | 0.048 |
| 1917 | -0.015 | -0.055 |
| 1918 | 0.027 | -0.040 |
| 1919 | -0.038 | -0.034 |
| 1920 | -0.040 | 0.018 |
| 1921 | -0.015 | 0.242 |
| 1922 | 0.019 | 0.025 |
| 1923 | 0.009 | -0.002 |
| 1924 | 0.003 | 0.075 |
| 1925 | -0.020 | -0.100 |
| 1926 | -0.038 | 0.048 |
| 1927 | 0.007 | 0.052 |
| 1928 | -0.034 | 0.018 |
| 1929 | 0.070 | -0.071 |
| 1930 | -0.011 | -0.036 |
| 1931 | 0.008 | 0.038 |

| | | |
|---|---|---|
| 1932 | -0.010 | -0.003 |
| 1933 | 0.037 | 0.010 |
| 1934 | -0.032 | -0.019 |
| 1935 | -0.027 | 0.058 |
| 1936 | -0.039 | -0.019 |
| 1937 | -0.024 | -0.031 |
| 1938 | 0.006 | 0.015 |
| 1939 | 0.034 | 0.005 |
| 1940 | -0.030 | -0.093 |
| 1941 | -0.053 | 0.006 |
| 1942 | -0.060 | -0.028 |
| 1943 | -0.011 | -0.082 |
| 1944 | 0.093 | -0.055 |
| 1945 | 0.065 | -0.036 |
| 1946 | -0.015 | 0.040 |
| 1947 | 0.003 | 0.019 |
| 1948 | -0.041 | -0.012 |
| 1949 | -0.017 | -0.046 |
| 1950 | -0.068 | -0.094 |
| 1951 | -0.049 | -0.059 |
| 1952 | 0.035 | 0.038 |
| 1953 | 0.091 | -0.093 |
| 1954 | -0.049 | 0.085 |
| 1955 | -0.025 | -0.019 |
| 1956 | -0.001 | 0.077 |
| 1957 | 0.001 | -0.027 |
| 1958 | -0.055 | 0.015 |
| 1959 | -0.030 | 0.146 |
| 1960 | -0.046 | -0.063 |
| 1961 | -0.026 | 0.082 |
| 1962 | 0.004 | -0.045 |
| 1963 | -0.009 | 0.178 |
| 1964 | 0.029 | 0.015 |
| 1965 | 0.004 | -0.032 |
| 1966 | 0.016 | 0.021 |
| 1967 | -0.009 | -0.144 |
| 1968 | -0.001 | -0.043 |
| 1969 | -0.009 | 0.062 |
| 1970 | -0.017 | -0.067 |
| 1971 | 0.036 | 0.100 |
| 1972 | 0.057 | -0.053 |
| 1973 | -0.028 | -0.090 |
| 1974 | 0.021 | 0.021 |
| 1975 | 0.002 | 0.058 |
| 1976 | 0.042 | 0.108 |
| 1977 | 0.025 | -0.064 |
| 1978 | -0.027 | -0.035 |
| 1979 | 0.027 | 0.080 |
| 1980 | 0.015 | -0.070 |
| 1981 | 0.028 | 0.005 |
| 1982 | -0.027 | -0.038 |

| | | |
|---|---|---|
| 1983 | -0.020 | -0.063 |
| 1984 | -0.041 | 0.004 |
| 1985 | 0.036 | 0.010 |
| 1986 | -0.007 | -0.056 |
| 1987 | -0.109 | 0.045 |
| 1988 | -0.041 | -0.012 |
| 1989 | 0.000 | -0.071 |
| 1990 | 0.053 | 0.019 |
| 1991 | -0.018 | -0.059 |
| 1992 | -0.019 | 0.017 |
| 1993 | -0.029 | 0.038 |
| 1994 | 0.025 | 0.055 |
| 1995 | -0.044 | 0.041 |
| 1996 | -0.065 | 0.027 |
| 1997 | 0.019 | 0.012 |
| 1998 | 0.054 | -0.128 |
| 1999 | -0.020 | -0.078 |
| 2000 | 0.005 | -0.034 |
| 2001 | 0.002 | 0.047 |
| 2002 | -0.012 | 0.032 |
| 2003 | 0.059 | 0.096 |
| 2004 | -0.022 | 0.083 |
| 2005 | -0.040 | -0.022 |
| 2006 | -0.030 | -0.028 |
| 2007 | 0.078 | -0.055 |
| 2008 | -0.005 | 0.084 |
| 2009 | -0.007 | -0.132 |
| 2010 | 0.048 | -0.021 |
| 2011 | 0.000 | -0.001 |
| 2012 | -0.031 | 0.025 |
| 2013 | 0.039 | -0.120 |
| 2014 | -0.022 | -0.132 |
| 2015 | -0.052 | -0.105 |
| 2016 | 0.007 | 0.024 |
| 2017 | -0.022 | -0.050 |
| 2018 | -0.035 | -0.010 |
| 2019 | -0.020 | -0.017 |
| 2020 | -0.014 | -0.012 |
| 2021 | 0.013 | -0.010 |
| 2022 | 0.012 | -0.095 |
| 2023 | 0.004 | -0.110 |
| 2024 | -0.026 | -0.037 |
| 2025 | 0.046 | -0.026 |
| 2026 | 0.001 | -0.041 |
| 2027 | -0.001 | -0.047 |
| 2028 | 0.013 | -0.017 |
| 2029 | -0.017 | -0.052 |
| 2030 | -0.025 | 0.031 |
| 2031 | -0.071 | 0.042 |
| 2032 | -0.023 | 0.089 |
| 2033 | 0.047 | -0.104 |

| | | |
|---|---|---|
| 2034 | 0.054 | 0.226 |
| 2035 | 0.001 | 0.040 |
| 2036 | -0.028 | -0.070 |
| 2037 | 0.027 | 0.006 |
| 2038 | -0.054 | -0.040 |
| 2039 | -0.015 | -0.147 |
| 2040 | 0.077 | 0.038 |
| 2041 | 0.011 | 0.075 |
| 2042 | 0.010 | -0.036 |
| 2043 | 0.003 | -0.077 |
| 2044 | -0.053 | -0.046 |
| 2045 | -0.066 | 0.096 |
| 2046 | -0.071 | 0.233 |
| 2047 | -0.011 | -0.053 |
| 2048 | -0.011 | 0.022 |
| 2049 | 0.046 | -0.037 |
| 2050 | 0.003 | 0.018 |
| 2051 | 0.008 | 0.081 |
| 2052 | 0.013 | -0.028 |
| 2053 | -0.032 | 0.062 |
| 2054 | -0.027 | -0.031 |
| 2055 | -0.023 | 0.051 |
| 2056 | 0.032 | -0.006 |
| 2057 | -0.023 | -0.073 |
| 2058 | -0.060 | 0.080 |
| 2059 | 0.052 | 0.028 |
| 2060 | -0.012 | -0.074 |
| 2061 | -0.021 | 0.036 |
| 2062 | -0.014 | -0.107 |
| 2063 | 0.068 | -0.094 |
| 2064 | -0.024 | -0.052 |
| 2065 | 0.031 | -0.059 |
| 2066 | -0.010 | 0.021 |
| 2067 | 0.024 | 0.135 |
| 2068 | 0.003 | -0.125 |
| 2069 | -0.035 | 0.076 |
| 2070 | -0.055 | 0.038 |
| 2071 | -0.041 | 0.054 |
| 2072 | 0.028 | 0.043 |
| 2073 | 0.060 | -0.042 |
| 2074 | -0.005 | -0.065 |
| 2075 | -0.051 | 0.084 |
| 2076 | 0.027 | 0.023 |
| 2077 | 0.021 | 0.000 |
| 2078 | -0.052 | -0.045 |
| 2079 | 0.038 | -0.134 |
| 2080 | -0.003 | 0.106 |
| 2081 | 0.023 | -0.132 |
| 2082 | 0.011 | -0.040 |
| 2083 | -0.022 | -0.008 |
| 2084 | -0.014 | 0.089 |

| | | |
|---|---|---|
| 2085 | 0.008 | 0.051 |
| 2086 | 0.048 | 0.035 |
| 2087 | 0.052 | 0.030 |
| 2088 | -0.026 | -0.068 |
| 2089 | 0.039 | 0.030 |
| 2090 | -0.001 | 0.004 |
| 2091 | -0.049 | -0.002 |
| 2092 | 0.004 | -0.003 |
| 2093 | 0.042 | -0.040 |
| 2094 | -0.007 | -0.020 |
| 2095 | -0.017 | -0.019 |
| 2096 | -0.004 | -0.065 |
| 2097 | -0.008 | -0.028 |
| 2098 | 0.002 | -0.028 |
| 2099 | -0.014 | 0.002 |
| 2100 | 0.028 | -0.033 |
| 2101 | -0.012 | 0.055 |
| 2102 | 0.011 | 0.079 |
| 2103 | 0.026 | 0.026 |
| 2104 | 0.020 | 0.000 |
| 2105 | -0.050 | 0.049 |
| 2106 | -0.013 | 0.017 |
| 2107 | -0.023 | 0.013 |
| 2108 | -0.025 | 0.017 |
| 2109 | -0.010 | -0.102 |
| 2110 | -0.016 | 0.026 |
| 2111 | 0.060 | 0.016 |
| 2112 | 0.001 | 0.094 |
| 2113 | 0.024 | -0.027 |
| 2114 | 0.023 | 0.015 |
| 2115 | 0.010 | -0.033 |
| 2116 | -0.014 | -0.094 |
| 2117 | 0.006 | 0.004 |
| 2118 | 0.036 | -0.064 |
| 2119 | 0.044 | -0.116 |
| 2120 | -0.067 | -0.089 |
| 2121 | -0.025 | 0.117 |
| 2122 | -0.022 | -0.020 |
| 2123 | -0.032 | 0.012 |
| 2124 | 0.078 | 0.016 |
| 2125 | 0.038 | 0.003 |
| 2126 | -0.064 | 0.121 |
| 2127 | -0.017 | 0.056 |
| 2128 | -0.012 | -0.027 |
| 2129 | -0.024 | -0.033 |
| 2130 | -0.047 | 0.052 |
| 2131 | -0.032 | 0.011 |
| 2132 | 0.012 | 0.006 |
| 2133 | 0.004 | -0.093 |
| 2134 | -0.013 | -0.040 |
| 2135 | 0.041 | 0.206 |

| | | |
|---|---|---|
| 2136 | 0.033 | 0.057 |
| 2137 | 0.009 | -0.001 |
| 2138 | 0.021 | -0.094 |
| 2139 | -0.023 | -0.066 |
| 2140 | -0.015 | -0.066 |
| 2141 | -0.010 | -0.106 |
| 2142 | -0.046 | 0.082 |
| 2143 | 0.006 | -0.072 |
| 2144 | -0.024 | 0.146 |
| 2145 | -0.004 | -0.004 |
| 2146 | 0.029 | -0.106 |
| 2147 | -0.006 | 0.095 |
| 2148 | -0.078 | -0.054 |
| 2149 | 0.072 | -0.042 |
| 2150 | -0.028 | -0.074 |
| 2151 | -0.020 | -0.178 |
| 2152 | -0.042 | -0.071 |
| 2153 | -0.006 | 0.013 |
| 2154 | -0.017 | -0.053 |
| 2155 | -0.028 | -0.023 |
| 2156 | -0.006 | 0.057 |
| 2157 | -0.033 | 0.000 |
| 2158 | 0.040 | 0.057 |
| 2159 | -0.020 | 0.063 |
| 2160 | 0.019 | -0.054 |
| 2161 | -0.025 | -0.063 |
| 2162 | 0.077 | -0.030 |
| 2163 | -0.024 | -0.033 |
| 2164 | -0.034 | 0.056 |
| 2165 | 0.066 | -0.010 |
| 2166 | 0.009 | -0.025 |
| 2167 | -0.086 | 0.004 |
| 2168 | -0.076 | 0.037 |
| 2169 | 0.032 | 0.092 |
| 2170 | -0.011 | -0.098 |
| 2171 | -0.046 | 0.046 |
| 2172 | 0.069 | -0.046 |
| 2173 | 0.029 | -0.069 |
| 2174 | -0.048 | -0.028 |
| 2175 | -0.020 | 0.058 |
| 2176 | -0.015 | -0.060 |
| 2177 | 0.004 | 0.085 |
| 2178 | 0.079 | 0.088 |
| 2179 | -0.025 | 0.056 |
| 2180 | 0.008 | 0.054 |
| 2181 | -0.017 | -0.016 |
| 2182 | -0.035 | -0.007 |
| 2183 | -0.022 | 0.092 |
| 2184 | 0.000 | 0.005 |
| 2185 | -0.020 | 0.148 |
| 2186 | -0.019 | 0.003 |

| | | |
|---|---|---|
| 2187 | -0.009 | 0.074 |
| 2188 | 0.031 | 0.064 |
| 2189 | 0.058 | -0.028 |
| 2190 | -0.016 | 0.021 |
| 2191 | 0.024 | -0.001 |
| 2192 | 0.039 | -0.003 |
| 2193 | -0.028 | -0.042 |
| 2194 | 0.028 | -0.055 |
| 2195 | 0.018 | 0.006 |
| 2196 | -0.062 | -0.110 |
| 2197 | 0.019 | -0.031 |
| 2198 | 0.036 | -0.008 |
| 2199 | 0.005 | -0.110 |
| 2200 | -0.029 | -0.074 |
| 2201 | 0.065 | -0.102 |
| 2202 | 0.031 | -0.088 |
| 2203 | -0.034 | -0.055 |
| 2204 | -0.001 | 0.146 |
| 2205 | 0.048 | -0.100 |
| 2206 | 0.013 | -0.122 |
| 2207 | -0.005 | 0.133 |
| 2208 | -0.008 | 0.045 |
| 2209 | 0.005 | 0.039 |
| 2210 | 0.032 | -0.011 |
| 2211 | 0.034 | 0.072 |
| 2212 | 0.005 | -0.040 |
| 2213 | 0.042 | -0.146 |
| 2214 | -0.026 | -0.012 |
| 2215 | -0.025 | 0.012 |
| 2216 | 0.026 | 0.079 |
| 2217 | 0.019 | -0.092 |
| 2218 | 0.007 | -0.038 |
| 2219 | 0.019 | 0.057 |
| 2220 | 0.007 | 0.030 |
| 2221 | -0.001 | -0.004 |
| 2222 | 0.019 | 0.000 |
| 2223 | 0.003 | -0.012 |
| 2224 | 0.047 | 0.013 |
| 2225 | 0.081 | -0.008 |
| 2226 | -0.003 | -0.137 |
| 2227 | 0.027 | 0.045 |
| 2228 | 0.010 | -0.015 |
| 2229 | 0.034 | 0.007 |
| 2230 | 0.047 | -0.141 |
| 2231 | 0.019 | -0.033 |
| 2232 | 0.002 | -0.036 |
| 2233 | 0.033 | 0.084 |
| 2234 | 0.058 | -0.004 |
| 2235 | -0.008 | 0.057 |
| 2236 | -0.055 | -0.040 |
| 2237 | 0.017 | -0.010 |

| | | |
|---|---|---|
| 2238 | -0.032 | -0.118 |
| 2239 | -0.018 | -0.034 |
| 2240 | -0.033 | 0.021 |
| 2241 | 0.001 | 0.165 |
| 2242 | -0.005 | -0.071 |
| 2243 | 0.044 | 0.083 |
| 2244 | 0.004 | 0.082 |
| 2245 | 0.017 | 0.038 |
| 2246 | -0.034 | 0.068 |
| 2247 | -0.003 | 0.020 |
| 2248 | 0.034 | 0.053 |
| 2249 | -0.004 | 0.077 |
| 2250 | -0.070 | 0.087 |
| 2251 | -0.028 | 0.046 |
| 2252 | 0.026 | -0.094 |
| 2253 | -0.009 | 0.091 |
| 2254 | 0.029 | -0.068 |
| 2255 | 0.038 | 0.041 |
| 2256 | 0.016 | 0.054 |
| 2257 | 0.058 | 0.005 |
| 2258 | -0.005 | -0.031 |
| 2259 | 0.026 | -0.040 |
| 2260 | -0.032 | 0.071 |
| 2261 | -0.024 | 0.027 |
| 2262 | 0.009 | 0.050 |
| 2263 | 0.022 | 0.071 |
| 2264 | -0.011 | 0.036 |
| 2265 | 0.065 | -0.017 |
| 2266 | 0.024 | -0.136 |
| 2267 | 0.045 | -0.102 |
| 2268 | 0.018 | 0.014 |
| 2269 | -0.001 | -0.107 |
| 2270 | -0.002 | -0.124 |
| 2271 | 0.024 | 0.041 |
| 2272 | -0.015 | -0.041 |
| 2273 | 0.004 | -0.005 |
| 2274 | 0.023 | -0.030 |
| 2275 | 0.000 | 0.096 |
| 2276 | 0.024 | 0.104 |
| 2277 | -0.042 | -0.029 |
| 2278 | 0.073 | -0.031 |
| 2279 | 0.046 | -0.017 |
| 2280 | -0.006 | 0.023 |
| 2281 | -0.021 | -0.028 |
| 2282 | 0.027 | -0.028 |
| 2283 | 0.028 | -0.009 |
| 2284 | 0.048 | -0.075 |
| 2285 | 0.018 | -0.042 |
| 2286 | -0.003 | -0.060 |
| 2287 | 0.048 | 0.013 |
| 2288 | -0.007 | -0.148 |

| | | |
|---|---|---|
| 2289 | 0.010 | -0.053 |
| 2290 | -0.005 | -0.052 |
| 2291 | -0.018 | -0.016 |
| 2292 | -0.035 | 0.043 |
| 2293 | -0.014 | -0.022 |
| 2294 | -0.022 | -0.126 |
| 2295 | -0.018 | -0.031 |
| 2296 | 0.027 | -0.125 |
| 2297 | -0.019 | -0.016 |
| 2298 | -0.033 | 0.251 |
| 2299 | -0.014 | -0.088 |
| 2300 | -0.036 | 0.040 |
| 2301 | -0.025 | -0.046 |
| 2302 | -0.021 | 0.044 |
| 2303 | -0.076 | 0.080 |
| 2304 | -0.037 | 0.158 |
| 2305 | -0.065 | 0.104 |
| 2306 | -0.020 | 0.042 |
| 2307 | 0.063 | 0.066 |
| 2308 | -0.009 | 0.082 |
| 2309 | -0.077 | 0.014 |
| 2310 | 0.037 | -0.007 |
| 2311 | 0.014 | 0.087 |
| 2312 | 0.030 | 0.001 |
| 2313 | 0.028 | -0.046 |
| 2314 | -0.025 | -0.013 |
| 2315 | 0.117 | -0.145 |
| 2316 | -0.071 | 0.133 |
| 2317 | 0.004 | 0.083 |
| 2318 | 0.022 | 0.096 |
| 2319 | -0.037 | -0.019 |
| 2320 | -0.008 | 0.051 |
| 2321 | 0.030 | 0.067 |
| 2322 | 0.034 | 0.025 |
| 2323 | -0.018 | 0.051 |
| 2324 | 0.067 | 0.069 |
| 2325 | -0.003 | -0.129 |
| 2326 | 0.046 | -0.003 |
| 2327 | -0.006 | 0.050 |
| 2328 | 0.031 | -0.022 |
| 2329 | -0.016 | 0.047 |
| 2330 | -0.041 | -0.032 |
| 2331 | -0.089 | -0.089 |
| 2332 | -0.050 | -0.064 |
| 2333 | -0.034 | -0.074 |
| 2334 | 0.020 | 0.105 |
| 2335 | 0.019 | -0.046 |
| 2336 | -0.021 | 0.063 |
| 2337 | -0.020 | 0.019 |
| 2338 | 0.038 | 0.049 |
| 2339 | -0.025 | 0.020 |

| | | |
|---|---|---|
| 2340 | 0.013 | -0.001 |
| 2341 | 0.032 | -0.059 |
| 2342 | 0.015 | 0.057 |
| 2343 | -0.039 | 0.059 |
| 2344 | 0.019 | 0.014 |
| 2345 | -0.060 | 0.120 |
| 2346 | -0.069 | 0.004 |
| 2347 | -0.040 | 0.031 |
| 2348 | -0.024 | 0.027 |
| 2349 | -0.024 | 0.058 |
| 2350 | -0.034 | 0.038 |
| 2351 | -0.042 | 0.011 |
| 2352 | 0.003 | 0.013 |
| 2353 | 0.039 | 0.023 |
| 2354 | 0.026 | 0.006 |
| 2355 | -0.013 | 0.015 |
| 2356 | -0.033 | 0.165 |
| 2357 | 0.024 | 0.036 |
| 2358 | 0.030 | 0.128 |
| 2359 | 0.037 | 0.004 |
| 2360 | -0.034 | -0.022 |
| 2361 | 0.032 | -0.074 |
| 2362 | -0.026 | 0.011 |
| 2363 | 0.036 | 0.014 |
| 2364 | -0.028 | -0.034 |
| 2365 | -0.027 | 0.018 |
| 2366 | -0.025 | 0.007 |
| 2367 | 0.054 | -0.162 |
| 2368 | -0.052 | 0.086 |
| 2369 | 0.014 | -0.024 |
| 2370 | 0.008 | 0.091 |
| 2371 | -0.022 | 0.029 |
| 2372 | 0.015 | 0.087 |
| 2373 | 0.048 | -0.022 |
| 2374 | 0.022 | -0.170 |
| 2375 | 0.001 | -0.042 |
| 2376 | -0.041 | -0.076 |
| 2377 | 0.021 | -0.001 |
| 2378 | -0.034 | -0.103 |
| 2379 | -0.065 | 0.020 |
| 2380 | 0.012 | 0.089 |
| 2381 | -0.020 | -0.087 |
| 2382 | -0.022 | -0.118 |
| 2383 | -0.029 | 0.005 |
| 2384 | -0.001 | -0.029 |
| 2385 | -0.027 | 0.100 |
| 2386 | 0.019 | -0.177 |
| 2387 | -0.030 | -0.015 |
| 2388 | -0.008 | 0.096 |
| 2389 | 0.029 | 0.028 |
| 2390 | -0.018 | 0.067 |

| | | |
|---|---|---|
| 2391 | -0.016 | -0.011 |
| 2392 | 0.045 | 0.066 |
| 2393 | -0.039 | -0.092 |
| 2394 | 0.014 | 0.104 |
| 2395 | -0.076 | -0.008 |
| 2396 | 0.045 | -0.134 |
| 2397 | 0.015 | -0.057 |
| 2398 | -0.036 | -0.033 |
| 2399 | -0.023 | 0.031 |
| 2400 | 0.040 | 0.062 |
| 2401 | 0.012 | 0.024 |
| 2402 | -0.058 | 0.016 |
| 2403 | 0.031 | 0.081 |
| 2404 | 0.062 | -0.040 |
| 2405 | 0.028 | -0.022 |
| 2406 | 0.010 | 0.052 |
| 2407 | 0.001 | 0.052 |
| 2408 | -0.006 | -0.073 |
| 2409 | -0.051 | -0.042 |
| 2410 | -0.027 | 0.032 |
| 2411 | 0.055 | -0.038 |
| 2412 | -0.023 | 0.070 |
| 2413 | 0.030 | -0.033 |
| 2414 | -0.025 | 0.014 |
| 2415 | 0.040 | -0.072 |
| 2416 | 0.029 | -0.043 |
| 2417 | 0.015 | -0.013 |
| 2418 | 0.043 | 0.031 |
| 2419 | 0.004 | -0.023 |
| 2420 | 0.021 | 0.038 |
| 2421 | -0.056 | 0.088 |
| 2422 | 0.069 | 0.151 |
| 2423 | 0.028 | 0.072 |
| 2424 | -0.010 | -0.022 |
| 2425 | -0.107 | -0.023 |
| 2426 | 0.007 | -0.047 |
| 2427 | -0.044 | 0.062 |
| 2428 | 0.061 | -0.034 |
| 2429 | 0.007 | -0.005 |
| 2430 | 0.007 | -0.001 |
| 2431 | 0.012 | -0.076 |
| 2432 | -0.004 | 0.087 |
| 2433 | 0.004 | 0.057 |
| 2434 | 0.029 | -0.136 |
| 2435 | -0.005 | -0.041 |
| 2436 | -0.014 | 0.012 |
| 2437 | 0.026 | 0.102 |
| 2438 | -0.009 | -0.045 |
| 2439 | -0.013 | 0.056 |
| 2440 | 0.015 | -0.096 |
| 2441 | 0.006 | 0.061 |

| | | |
|---|---|---|
| 2442 | -0.006 | 0.052 |
| 2443 | -0.022 | -0.006 |
| 2444 | -0.020 | -0.091 |
| 2445 | 0.008 | -0.039 |
| 2446 | 0.023 | -0.026 |
| 2447 | 0.071 | 0.042 |
| 2448 | -0.046 | -0.029 |
| 2449 | -0.024 | -0.024 |
| 2450 | 0.000 | 0.024 |
| 2451 | -0.034 | -0.045 |
| 2452 | -0.080 | -0.022 |
| 2453 | 0.020 | -0.019 |
| 2454 | 0.001 | -0.004 |
| 2455 | -0.046 | 0.156 |
| 2456 | 0.005 | 0.113 |
| 2457 | -0.037 | 0.025 |
| 2458 | -0.006 | 0.000 |
| 2459 | -0.012 | 0.002 |
| 2460 | 0.018 | -0.035 |
| 2461 | 0.025 | 0.155 |
| 2462 | 0.040 | 0.071 |
| 2463 | 0.024 | 0.076 |
| 2464 | 0.054 | -0.053 |
| 2465 | 0.032 | -0.047 |
| 2466 | 0.010 | 0.058 |
| 2467 | 0.009 | -0.005 |
| 2468 | 0.007 | -0.005 |
| 2469 | -0.017 | -0.001 |
| 2470 | -0.006 | -0.010 |
| 2471 | -0.028 | 0.073 |
| 2472 | 0.002 | 0.039 |
| 2473 | -0.020 | -0.002 |
| 2474 | -0.004 | -0.093 |
| 2475 | 0.022 | 0.052 |
| 2476 | 0.012 | 0.036 |
| 2477 | -0.014 | 0.075 |
| 2478 | 0.002 | -0.106 |
| 2479 | 0.058 | 0.072 |
| 2480 | -0.038 | 0.038 |
| 2481 | 0.025 | 0.033 |
| 2482 | -0.015 | -0.010 |
| 2483 | -0.006 | 0.097 |
| 2484 | 0.040 | -0.006 |
| 2485 | -0.013 | 0.058 |
| 2486 | 0.013 | 0.082 |
| 2487 | 0.031 | 0.112 |
| 2488 | -0.019 | 0.078 |
| 2489 | 0.049 | 0.115 |
| 2490 | -0.048 | -0.110 |
| 2491 | -0.012 | 0.007 |
| 2492 | 0.000 | -0.003 |

| | | |
|---|---|---|
| 2493 | 0.017 | -0.079 |
| 2494 | -0.041 | -0.050 |
| 2495 | 0.000 | 0.024 |
| 2496 | -0.083 | 0.045 |
| 2497 | -0.006 | 0.100 |
| 2498 | 0.013 | 0.049 |
| 2499 | -0.019 | -0.089 |
| 2500 | 0.028 | -0.058 |
| 2501 | 0.028 | 0.035 |
| 2502 | -0.003 | 0.034 |
| 2503 | 0.014 | -0.078 |
| 2504 | -0.026 | 0.066 |
| 2505 | -0.029 | 0.031 |
| 2506 | 0.040 | 0.120 |
| 2507 | 0.030 | 0.075 |
| 2508 | 0.021 | -0.073 |
| 2509 | 0.030 | 0.033 |
| 2510 | -0.073 | 0.129 |
| 2511 | 0.014 | 0.166 |
| 2512 | 0.081 | -0.053 |
| 2513 | -0.004 | 0.045 |
| 2514 | 0.037 | -0.040 |
| 2515 | -0.027 | 0.035 |
| 2516 | 0.003 | -0.168 |
| 2517 | -0.027 | 0.032 |
| 2518 | 0.011 | -0.107 |
| 2519 | -0.004 | -0.112 |
| 2520 | 0.005 | -0.002 |
| 2521 | -0.041 | 0.052 |
| 2522 | -0.007 | -0.045 |
| 2523 | 0.014 | 0.090 |
| 2524 | 0.016 | 0.025 |
| 2525 | -0.002 | -0.059 |
| 2526 | 0.041 | 0.096 |
| 2527 | 0.069 | 0.033 |
| 2528 | -0.001 | 0.044 |
| 2529 | -0.003 | 0.053 |
| 2530 | 0.002 | 0.026 |
| 2531 | -0.027 | -0.079 |
| 2532 | 0.003 | -0.110 |
| 2533 | -0.004 | 0.011 |
| 2534 | -0.020 | -0.015 |
| 2535 | -0.031 | -0.053 |
| 2536 | -0.007 | 0.009 |
| 2537 | 0.002 | -0.106 |
| 2538 | -0.068 | -0.028 |
| 2539 | -0.018 | 0.081 |
| 2540 | 0.044 | 0.097 |
| 2541 | 0.041 | -0.014 |
| 2542 | 0.043 | 0.030 |
| 2543 | -0.062 | -0.149 |

| | | |
|---|---|---|
| 2544 | -0.045 | 0.198 |
| 2545 | -0.015 | 0.086 |
| 2546 | -0.002 | 0.019 |
| 2547 | 0.008 | 0.060 |
| 2548 | -0.027 | -0.092 |
| 2549 | 0.030 | 0.092 |
| 2550 | 0.015 | 0.013 |
| 2551 | 0.017 | -0.081 |
| 2552 | 0.045 | -0.157 |
| 2553 | -0.020 | 0.009 |
| 2554 | -0.074 | -0.003 |
| 2555 | 0.004 | -0.029 |
| 2556 | 0.050 | 0.110 |
| 2557 | 0.046 | 0.019 |
| 2558 | 0.051 | 0.061 |
| 2559 | -0.019 | -0.021 |
| 2560 | -0.010 | -0.012 |
| 2561 | 0.008 | -0.024 |
| 2562 | 0.007 | 0.039 |
| 2563 | 0.011 | 0.081 |
| 2564 | 0.032 | 0.038 |
| 2565 | -0.005 | -0.067 |
| 2566 | -0.002 | 0.007 |
| 2567 | -0.010 | 0.003 |
| 2568 | -0.060 | -0.122 |
| 2569 | 0.016 | -0.085 |
| 2570 | -0.005 | -0.010 |
| 2571 | -0.040 | -0.004 |
| 2572 | -0.002 | -0.053 |
| 2573 | 0.016 | -0.007 |
| 2574 | 0.004 | 0.025 |
| 2575 | 0.024 | -0.001 |
| 2576 | -0.012 | -0.046 |
| 2577 | 0.013 | 0.014 |
| 2578 | -0.027 | -0.015 |
| 2579 | 0.043 | -0.021 |
| 2580 | 0.048 | -0.021 |
| 2581 | -0.021 | 0.083 |
| 2582 | 0.020 | -0.050 |
| 2583 | 0.031 | 0.043 |
| 2584 | -0.009 | 0.090 |
| 2585 | -0.047 | 0.065 |
| 2586 | -0.015 | -0.092 |
| 2587 | -0.001 | -0.002 |
| 2588 | 0.008 | -0.010 |
| 2589 | -0.048 | 0.040 |
| 2590 | 0.024 | -0.019 |
| 2591 | 0.001 | 0.015 |
| 2592 | -0.025 | 0.033 |
| 2593 | -0.015 | -0.077 |
| 2594 | 0.038 | 0.055 |

| | | |
|---|---|---|
| 2595 | 0.028 | -0.060 |
| 2596 | 0.018 | -0.027 |
| 2597 | 0.034 | 0.076 |
| 2598 | 0.016 | 0.090 |
| 2599 | -0.008 | -0.050 |
| 2600 | 0.039 | 0.116 |
| 2601 | 0.010 | -0.043 |
| 2602 | 0.032 | 0.042 |
| 2603 | 0.051 | 0.204 |
| 2604 | 0.028 | -0.051 |
| 2605 | 0.020 | -0.022 |
| 2606 | -0.020 | 0.132 |
| 2607 | -0.014 | 0.015 |
| 2608 | 0.064 | 0.058 |
| 2609 | 0.004 | 0.047 |
| 2610 | 0.026 | 0.146 |
| 2611 | -0.038 | 0.065 |
| 2612 | -0.001 | -0.071 |
| 2613 | -0.070 | -0.004 |
| 2614 | 0.003 | -0.074 |
| 2615 | 0.018 | -0.008 |
| 2616 | -0.022 | 0.053 |
| 2617 | -0.014 | -0.013 |
| 2618 | -0.015 | 0.046 |
| 2619 | -0.024 | -0.090 |
| 2620 | -0.021 | 0.066 |
| 2621 | 0.034 | 0.005 |
| 2622 | 0.038 | -0.020 |
| 2623 | 0.013 | 0.039 |
| 2624 | -0.091 | 0.093 |
| 2625 | -0.007 | -0.085 |
| 2626 | -0.005 | -0.030 |
| 2627 | 0.008 | -0.009 |
| 2628 | -0.009 | 0.003 |
| 2629 | 0.000 | 0.032 |
| 2630 | 0.038 | -0.165 |
| 2631 | -0.006 | 0.082 |
| 2632 | -0.023 | -0.065 |
| 2633 | -0.008 | 0.170 |
| 2634 | -0.071 | -0.068 |
| 2635 | -0.046 | -0.032 |
| 2636 | 0.030 | -0.062 |
| 2637 | -0.034 | -0.005 |
| 2638 | 0.006 | -0.057 |
| 2639 | -0.055 | 0.047 |
| 2640 | -0.032 | -0.076 |
| 2641 | -0.048 | 0.031 |
| 2642 | -0.036 | -0.144 |
| 2643 | -0.009 | -0.153 |
| 2644 | 0.023 | 0.171 |
| 2645 | -0.061 | 0.071 |

| | | |
|---|---|---|
| 2646 | -0.050 | 0.060 |
| 2647 | -0.003 | -0.019 |
| 2648 | 0.033 | -0.092 |
| 2649 | -0.021 | -0.082 |
| 2650 | 0.022 | 0.076 |
| 2651 | 0.021 | 0.052 |
| 2652 | -0.011 | 0.021 |
| 2653 | 0.052 | -0.064 |
| 2654 | 0.024 | -0.033 |
| 2655 | -0.010 | 0.008 |
| 2656 | 0.025 | -0.040 |
| 2657 | 0.047 | -0.044 |
| 2658 | 0.016 | -0.018 |
| 2659 | -0.058 | 0.006 |
| 2660 | -0.012 | -0.036 |
| 2661 | 0.010 | -0.046 |
| 2662 | -0.021 | -0.100 |
| 2663 | 0.004 | -0.065 |
| 2664 | -0.017 | 0.080 |
| 2665 | 0.005 | 0.058 |
| 2666 | -0.011 | -0.076 |
| 2667 | -0.002 | -0.113 |
| 2668 | -0.024 | 0.028 |
| 2669 | -0.009 | -0.042 |
| 2670 | 0.010 | 0.068 |
| 2671 | -0.028 | 0.134 |
| 2672 | 0.029 | -0.103 |
| 2673 | 0.053 | -0.038 |
| 2674 | 0.023 | 0.031 |
| 2675 | 0.039 | -0.075 |
| 2676 | -0.006 | -0.007 |
| 2677 | -0.001 | 0.035 |
| 2678 | 0.000 | 0.024 |
| 2679 | 0.054 | -0.115 |
| 2680 | 0.029 | -0.020 |
| 2681 | 0.010 | -0.063 |
| 2682 | -0.037 | 0.076 |
| 2683 | 0.072 | 0.003 |
| 2684 | -0.001 | -0.058 |
| 2685 | -0.002 | 0.039 |
| 2686 | -0.031 | 0.030 |
| 2687 | 0.013 | -0.052 |
| 2688 | -0.003 | 0.076 |
| 2689 | 0.051 | 0.023 |
| 2690 | 0.003 | -0.015 |
| 2691 | -0.027 | 0.066 |
| 2692 | 0.011 | -0.011 |
| 2693 | 0.000 | 0.018 |
| 2694 | 0.065 | 0.022 |
| 2695 | 0.005 | 0.132 |
| 2696 | 0.003 | 0.062 |

| | | |
|---|---|---|
| 2697 | 0.021 | 0.029 |
| 2698 | 0.018 | 0.034 |
| 2699 | -0.010 | 0.030 |
| 2700 | -0.035 | 0.118 |
| 2701 | -0.057 | 0.052 |
| 2702 | 0.024 | -0.043 |
| 2703 | -0.005 | 0.045 |
| 2704 | 0.007 | 0.059 |
| 2705 | 0.056 | 0.049 |
| 2706 | -0.010 | -0.071 |
| 2707 | 0.003 | 0.018 |
| 2708 | 0.041 | 0.059 |
| 2709 | 0.019 | -0.127 |
| 2710 | -0.011 | 0.109 |
| 2711 | -0.025 | 0.052 |
| 2712 | 0.010 | 0.065 |
| 2713 | 0.017 | 0.097 |
| 2714 | 0.023 | 0.062 |
| 2715 | 0.058 | 0.040 |
| 2716 | -0.025 | -0.078 |
| 2717 | -0.008 | 0.004 |
| 2718 | -0.053 | 0.091 |
| 2719 | 0.017 | -0.110 |
| 2720 | -0.005 | 0.023 |
| 2721 | 0.010 | -0.042 |
| 2722 | -0.020 | -0.004 |
| 2723 | -0.072 | -0.072 |
| 2724 | -0.012 | -0.022 |
| 2725 | -0.018 | 0.004 |
| 2726 | -0.054 | 0.080 |
| 2727 | 0.069 | -0.065 |
| 2728 | 0.011 | 0.193 |
| 2729 | -0.027 | 0.050 |
| 2730 | 0.103 | -0.114 |
| 2731 | 0.074 | -0.068 |
| 2732 | -0.041 | -0.118 |
| 2733 | 0.002 | -0.012 |
| 2734 | -0.089 | 0.030 |
| 2735 | -0.012 | 0.053 |
| 2736 | 0.057 | -0.120 |
| 2737 | 0.012 | 0.006 |
| 2738 | -0.028 | -0.035 |
| 2739 | -0.033 | -0.041 |
| 2740 | 0.005 | -0.022 |
| 2741 | -0.033 | 0.024 |
| 2742 | -0.018 | 0.096 |
| 2743 | 0.036 | 0.073 |
| 2744 | -0.030 | -0.042 |
| 2745 | 0.036 | -0.019 |
| 2746 | -0.045 | -0.043 |
| 2747 | 0.042 | -0.010 |

| | | |
|---|---|---|
| 2748 | -0.041 | -0.023 |
| 2749 | 0.010 | -0.074 |
| 2750 | 0.019 | 0.065 |
| 2751 | 0.000 | -0.208 |
| 2752 | 0.028 | 0.013 |
| 2753 | -0.027 | 0.019 |
| 2754 | 0.001 | -0.012 |
| 2755 | 0.001 | 0.047 |
| 2756 | 0.044 | -0.038 |
| 2757 | -0.005 | -0.080 |
| 2758 | -0.012 | -0.038 |
| 2759 | 0.050 | -0.080 |
| 2760 | 0.078 | -0.172 |
| 2761 | 0.040 | -0.019 |
| 2762 | -0.006 | 0.034 |
| 2763 | 0.009 | 0.014 |
| 2764 | 0.010 | 0.131 |
| 2765 | -0.021 | 0.052 |
| 2766 | -0.050 | -0.036 |
| 2767 | 0.019 | -0.076 |
| 2768 | -0.004 | 0.166 |
| 2769 | 0.005 | 0.005 |
| 2770 | 0.028 | -0.002 |
| 2771 | 0.047 | 0.083 |
| 2772 | -0.047 | -0.023 |
| 2773 | 0.042 | -0.059 |
| 2774 | 0.017 | 0.014 |
| 2775 | 0.053 | 0.032 |
| 2776 | -0.035 | 0.032 |
| 2777 | 0.028 | 0.006 |
| 2778 | -0.034 | 0.098 |
| 2779 | 0.007 | -0.009 |
| 2780 | 0.042 | -0.011 |
| 2781 | 0.006 | -0.007 |
| 2782 | -0.024 | 0.244 |
| 2783 | -0.045 | -0.107 |
| 2784 | 0.002 | 0.109 |
| 2785 | -0.071 | 0.086 |
| 2786 | 0.029 | 0.037 |
| 2787 | -0.007 | 0.121 |
| 2788 | -0.043 | 0.019 |
| 2789 | -0.030 | -0.001 |
| 2790 | 0.002 | -0.088 |
| 2791 | -0.011 | 0.150 |
| 2792 | -0.020 | -0.090 |
| 2793 | -0.059 | -0.005 |
| 2794 | -0.044 | 0.000 |
| 2795 | -0.003 | 0.082 |
| 2796 | 0.031 | -0.073 |
| 2797 | 0.006 | -0.054 |
| 2798 | -0.014 | -0.122 |

| | | |
|---|---|---|
| 2799 | 0.027 | -0.025 |
| 2800 | -0.015 | 0.066 |
| 2801 | -0.072 | -0.086 |
| 2802 | 0.017 | -0.012 |
| 2803 | -0.018 | 0.028 |
| 2804 | 0.040 | -0.037 |
| 2805 | -0.028 | 0.039 |
| 2806 | 0.010 | -0.073 |
| 2807 | -0.021 | -0.036 |
| 2808 | 0.004 | -0.057 |
| 2809 | 0.016 | -0.098 |
| 2810 | -0.055 | 0.033 |
| 2811 | -0.001 | 0.096 |
| 2812 | -0.029 | -0.108 |
| 2813 | -0.008 | 0.051 |
| 2814 | 0.007 | 0.006 |
| 2815 | 0.017 | -0.123 |
| 2816 | -0.012 | -0.073 |
| 2817 | 0.014 | 0.035 |
| 2818 | 0.029 | 0.040 |
| 2819 | 0.000 | -0.014 |
| 2820 | -0.051 | 0.007 |
| 2821 | 0.007 | 0.077 |
| 2822 | 0.033 | 0.107 |
| 2823 | -0.025 | -0.034 |
| 2824 | 0.035 | 0.031 |
| 2825 | -0.009 | -0.107 |
| 2826 | 0.046 | 0.052 |
| 2827 | -0.052 | 0.008 |
| 2828 | -0.013 | -0.006 |
| 2829 | -0.048 | -0.208 |
| 2830 | 0.017 | 0.075 |
| 2831 | 0.009 | 0.031 |
| 2832 | -0.005 | -0.043 |
| 2833 | 0.007 | -0.089 |
| 2834 | -0.036 | 0.006 |
| 2835 | 0.029 | -0.018 |
| 2836 | 0.016 | -0.003 |
| 2837 | -0.014 | -0.042 |
| 2838 | 0.013 | -0.116 |
| 2839 | 0.105 | -0.097 |
| 2840 | 0.000 | -0.020 |
| 2841 | -0.061 | 0.078 |
| 2842 | 0.003 | -0.028 |
| 2843 | -0.082 | 0.088 |
| 2844 | 0.076 | 0.017 |
| 2845 | -0.005 | 0.078 |
| 2846 | 0.000 | -0.092 |
| 2847 | 0.002 | -0.075 |
| 2848 | -0.017 | 0.002 |
| 2849 | 0.031 | 0.124 |

| | | |
|---|---|---|
| 2850 | 0.023 | -0.058 |
| 2851 | 0.026 | 0.054 |
| 2852 | 0.003 | -0.010 |
| 2853 | -0.025 | -0.092 |
| 2854 | -0.010 | 0.064 |
| 2855 | -0.022 | -0.012 |
| 2856 | 0.019 | -0.043 |
| 2857 | -0.046 | -0.043 |
| 2858 | 0.020 | -0.141 |
| 2859 | 0.016 | -0.048 |
| 2860 | 0.043 | 0.002 |
| 2861 | 0.006 | 0.058 |
| 2862 | 0.015 | 0.034 |
| 2863 | -0.010 | -0.067 |
| 2864 | 0.008 | 0.011 |
| 2865 | 0.010 | 0.068 |
| 2866 | 0.059 | 0.010 |
| 2867 | 0.066 | 0.120 |
| 2868 | 0.025 | 0.053 |
| 2869 | 0.000 | -0.059 |
| 2870 | 0.017 | 0.128 |
| 2871 | 0.016 | -0.011 |
| 2872 | -0.009 | 0.020 |
| 2873 | -0.013 | -0.029 |
| 2874 | -0.031 | -0.082 |
| 2875 | -0.006 | -0.042 |
| 2876 | -0.008 | 0.046 |
| 2877 | 0.008 | -0.016 |
| 2878 | -0.002 | 0.022 |
| 2879 | -0.032 | -0.032 |
| 2880 | -0.034 | -0.053 |
| 2881 | 0.034 | 0.012 |
| 2882 | -0.008 | 0.051 |
| 2883 | -0.038 | -0.179 |
| 2884 | 0.058 | 0.002 |
| 2885 | 0.018 | -0.101 |
| 2886 | -0.076 | -0.095 |
| 2887 | -0.017 | 0.001 |
| 2888 | 0.075 | 0.027 |
| 2889 | -0.029 | -0.178 |
| 2890 | 0.024 | 0.011 |
| 2891 | -0.055 | -0.008 |
| 2892 | 0.005 | -0.001 |
| 2893 | 0.070 | 0.038 |
| 2894 | -0.048 | -0.055 |
| 2895 | 0.011 | 0.122 |
| 2896 | 0.027 | -0.098 |
| 2897 | 0.044 | -0.001 |
| 2898 | -0.044 | 0.005 |
| 2899 | 0.002 | -0.039 |
| 2900 | -0.030 | -0.063 |

| | | |
|---|---|---|
| 2901 | -0.005 | 0.057 |
| 2902 | 0.045 | 0.040 |
| 2903 | 0.071 | -0.094 |
| 2904 | -0.039 | 0.030 |
| 2905 | -0.020 | 0.082 |
| 2906 | 0.043 | 0.049 |
| 2907 | -0.009 | -0.040 |
| 2908 | -0.024 | 0.148 |
| 2909 | 0.058 | -0.089 |
| 2910 | -0.032 | -0.058 |
| 2911 | -0.019 | -0.046 |
| 2912 | -0.026 | -0.083 |
| 2913 | 0.017 | 0.019 |
| 2914 | 0.010 | -0.038 |
| 2915 | -0.028 | 0.094 |
| 2916 | -0.029 | -0.049 |
| 2917 | -0.065 | 0.083 |
| 2918 | -0.023 | 0.050 |
| 2919 | -0.033 | -0.110 |
| 2920 | 0.024 | 0.014 |
| 2921 | -0.028 | 0.029 |
| 2922 | 0.045 | 0.180 |
| 2923 | 0.075 | 0.085 |
| 2924 | -0.026 | 0.068 |
| 2925 | 0.008 | 0.068 |
| 2926 | -0.024 | 0.054 |
| 2927 | 0.024 | -0.018 |
| 2928 | 0.005 | -0.057 |
| 2929 | -0.012 | -0.012 |
| 2930 | 0.021 | 0.158 |
| 2931 | -0.029 | 0.065 |
| 2932 | -0.018 | -0.022 |
| 2933 | 0.026 | -0.084 |
| 2934 | 0.010 | -0.028 |
| 2935 | 0.045 | -0.039 |
| 2936 | -0.025 | -0.033 |
| 2937 | -0.034 | -0.011 |
| 2938 | 0.017 | 0.049 |
| 2939 | 0.000 | -0.164 |
| 2940 | -0.023 | 0.031 |
| 2941 | -0.021 | -0.076 |
| 2942 | -0.066 | 0.015 |
| 2943 | 0.056 | 0.094 |
| 2944 | 0.008 | -0.011 |
| 2945 | 0.041 | 0.066 |
| 2946 | 0.055 | -0.040 |
| 2947 | 0.028 | 0.028 |
| 2948 | -0.018 | 0.042 |
| 2949 | -0.040 | 0.076 |
| 2950 | 0.006 | -0.042 |
| 2951 | 0.008 | 0.001 |

| | | |
|---|---|---|
| 2952 | 0.006 | -0.075 |
| 2953 | 0.007 | 0.008 |
| 2954 | -0.011 | -0.035 |
| 2955 | 0.055 | 0.052 |
| 2956 | -0.042 | -0.048 |
| 2957 | -0.050 | 0.078 |
| 2958 | 0.025 | -0.002 |
| 2959 | -0.088 | 0.129 |
| 2960 | 0.002 | -0.015 |
| 2961 | 0.015 | -0.078 |
| 2962 | -0.002 | -0.026 |
| 2963 | 0.003 | -0.040 |
| 2964 | 0.005 | -0.046 |
| 2965 | 0.011 | -0.042 |
| 2966 | 0.007 | 0.093 |
| 2967 | 0.039 | -0.055 |
| 2968 | 0.014 | -0.114 |
| 2969 | 0.003 | -0.068 |
| 2970 | 0.014 | -0.058 |
| 2971 | 0.056 | 0.059 |
| 2972 | -0.021 | 0.038 |
| 2973 | 0.004 | 0.095 |
| 2974 | -0.046 | -0.017 |
| 2975 | 0.041 | 0.081 |
| 2976 | -0.001 | 0.025 |
| 2977 | 0.004 | 0.094 |
| 2978 | 0.037 | -0.054 |
| 2979 | -0.062 | -0.085 |
| 2980 | -0.009 | 0.087 |
| 2981 | -0.014 | -0.017 |
| 2982 | 0.043 | 0.001 |
| 2983 | 0.077 | 0.082 |
| 2984 | -0.028 | -0.124 |
| 2985 | -0.046 | 0.112 |
| 2986 | 0.032 | 0.027 |
| 2987 | -0.024 | 0.033 |
| 2988 | -0.011 | 0.051 |
| 2989 | -0.004 | 0.006 |
| 2990 | -0.026 | -0.023 |
| 2991 | -0.034 | 0.022 |
| 2992 | -0.036 | 0.027 |
| 2993 | -0.042 | 0.062 |
| 2994 | 0.013 | 0.038 |
| 2995 | 0.005 | 0.004 |
| 2996 | 0.005 | 0.155 |
| 2997 | -0.003 | -0.043 |
| 2998 | -0.012 | 0.063 |
| 2999 | -0.067 | -0.185 |
| 3000 | -0.074 | 0.054 |
| 3001 | -0.026 | 0.010 |
| 3002 | 0.009 | -0.011 |

| | | |
|---|---|---|
| 3003 | 0.011 | -0.012 |
| 3004 | -0.030 | 0.009 |
| 3005 | -0.014 | -0.072 |
| 3006 | 0.059 | -0.044 |
| 3007 | -0.015 | -0.029 |
| 3008 | -0.056 | 0.096 |
| 3009 | -0.022 | -0.046 |
| 3010 | 0.012 | -0.133 |
| 3011 | -0.001 | 0.145 |
| 3012 | -0.004 | -0.088 |
| 3013 | -0.052 | 0.046 |
| 3014 | 0.040 | -0.099 |
| 3015 | -0.025 | 0.019 |
| 3016 | -0.028 | -0.038 |
| 3017 | 0.008 | -0.035 |
| 3018 | -0.062 | 0.037 |
| 3019 | -0.025 | -0.055 |
| 3020 | 0.000 | -0.087 |
| 3021 | 0.005 | 0.043 |
| 3022 | 0.008 | -0.129 |
| 3023 | -0.042 | -0.083 |
| 3024 | 0.038 | -0.013 |
| 3025 | 0.035 | 0.038 |
| 3026 | -0.017 | 0.002 |
| 3027 | 0.039 | -0.058 |
| 3028 | -0.029 | -0.135 |
| 3029 | 0.053 | -0.043 |
| 3030 | -0.026 | 0.022 |
| 3031 | 0.016 | 0.010 |
| 3032 | 0.006 | -0.004 |
| 3033 | -0.029 | -0.007 |
| 3034 | 0.029 | 0.020 |
| 3035 | -0.009 | -0.038 |
| 3036 | 0.059 | -0.062 |
| 3037 | 0.040 | 0.059 |
| 3038 | -0.033 | -0.047 |
| 3039 | 0.010 | -0.054 |
| 3040 | 0.002 | -0.077 |
| 3041 | 0.007 | 0.002 |
| 3042 | -0.023 | -0.022 |
| 3043 | 0.022 | -0.019 |
| 3044 | 0.028 | -0.045 |
| 3045 | -0.012 | -0.073 |
| 3046 | -0.011 | -0.031 |
| 3047 | 0.017 | -0.047 |
| 3048 | 0.006 | -0.045 |
| 3049 | 0.066 | -0.090 |
| 3050 | -0.018 | 0.013 |
| 3051 | 0.022 | 0.004 |
| 3052 | 0.022 | 0.022 |
| 3053 | -0.056 | -0.048 |

| | | |
|---|---|---|
| 3054 | -0.073 | -0.010 |
| 3055 | 0.001 | -0.006 |
| 3056 | 0.001 | 0.051 |
| 3057 | 0.026 | 0.042 |
| 3058 | -0.047 | -0.018 |
| 3059 | 0.003 | 0.026 |
| 3060 | 0.053 | 0.081 |
| 3061 | -0.041 | 0.034 |
| 3062 | 0.026 | 0.039 |
| 3063 | -0.010 | 0.028 |
| 3064 | 0.080 | 0.238 |
| 3065 | -0.011 | 0.028 |
| 3066 | 0.017 | -0.006 |
| 3067 | 0.007 | 0.064 |
| 3068 | -0.009 | -0.073 |
| 3069 | 0.008 | 0.087 |
| 3070 | 0.017 | 0.022 |
| 3071 | -0.004 | 0.097 |
| 3072 | -0.007 | 0.106 |
| 3073 | -0.018 | 0.103 |
| 3074 | -0.062 | 0.003 |
| 3075 | -0.034 | 0.035 |
| 3076 | 0.012 | -0.026 |
| 3077 | -0.046 | 0.115 |
| 3078 | 0.031 | 0.016 |
| 3079 | 0.002 | -0.028 |
| 3080 | 0.009 | 0.124 |
| 3081 | -0.083 | 0.026 |
| 3082 | -0.031 | -0.016 |
| 3083 | -0.047 | -0.108 |
| 3084 | 0.011 | 0.005 |
| 3085 | 0.022 | 0.029 |
| 3086 | -0.019 | 0.009 |
| 3087 | 0.007 | 0.090 |
| 3088 | 0.003 | 0.055 |
| 3089 | -0.063 | 0.069 |
| 3090 | 0.026 | 0.091 |
| 3091 | -0.027 | -0.022 |
| 3092 | -0.051 | 0.050 |
| 3093 | 0.038 | -0.041 |
| 3094 | 0.023 | 0.115 |
| 3095 | -0.024 | -0.005 |
| 3096 | 0.048 | 0.102 |
| 3097 | -0.008 | 0.083 |
| 3098 | 0.003 | 0.043 |
| 3099 | 0.060 | 0.022 |
| 3100 | 0.013 | 0.028 |
| 3101 | -0.015 | 0.044 |
| 3102 | 0.007 | -0.022 |
| 3103 | -0.079 | 0.099 |
| 3104 | -0.028 | 0.006 |

| | | |
|---|---|---|
| 3105 | -0.027 | -0.054 |
| 3106 | 0.007 | -0.013 |
| 3107 | -0.034 | 0.009 |
| 3108 | -0.044 | -0.089 |
| 3109 | -0.030 | -0.089 |
| 3110 | 0.010 | 0.003 |
| 3111 | -0.061 | 0.047 |
| 3112 | 0.010 | -0.152 |
| 3113 | -0.067 | 0.011 |
| 3114 | 0.010 | -0.159 |
| 3115 | -0.005 | 0.050 |
| 3116 | 0.004 | 0.112 |
| 3117 | -0.026 | -0.020 |
| 3118 | 0.054 | -0.032 |
| 3119 | 0.027 | -0.015 |
| 3120 | -0.061 | 0.062 |
| 3121 | -0.011 | -0.091 |
| 3122 | 0.025 | -0.071 |
| 3123 | 0.009 | -0.051 |
| 3124 | -0.003 | 0.018 |
| 3125 | 0.004 | -0.233 |
| 3126 | 0.002 | -0.010 |
| 3127 | 0.029 | 0.003 |
| 3128 | 0.006 | -0.093 |
| 3129 | 0.048 | -0.046 |
| 3130 | 0.015 | -0.012 |
| 3131 | 0.001 | -0.013 |
| 3132 | -0.066 | 0.010 |
| 3133 | -0.011 | -0.004 |
| 3134 | 0.016 | 0.090 |
| 3135 | -0.049 | 0.077 |
| 3136 | 0.019 | -0.029 |
| 3137 | -0.032 | -0.063 |
| 3138 | 0.012 | 0.150 |
| 3139 | 0.006 | 0.099 |
| 3140 | -0.036 | 0.024 |
| 3141 | -0.004 | 0.018 |
| 3142 | -0.071 | -0.018 |
| 3143 | -0.059 | -0.012 |
| 3144 | 0.036 | 0.084 |
| 3145 | -0.082 | -0.016 |
| 3146 | -0.058 | -0.049 |
| 3147 | 0.052 | 0.065 |
| 3148 | -0.027 | 0.014 |
| 3149 | -0.008 | 0.072 |
| 3150 | -0.019 | -0.005 |
| 3151 | 0.017 | -0.028 |
| 3152 | -0.054 | 0.051 |
| 3153 | -0.009 | 0.136 |
| 3154 | -0.005 | -0.060 |
| 3155 | -0.013 | -0.077 |

| | | |
|---|---|---|
| 3156 | 0.040 | 0.039 |
| 3157 | 0.026 | 0.114 |
| 3158 | -0.020 | 0.007 |
| 3159 | 0.061 | -0.039 |
| 3160 | 0.014 | 0.068 |
| 3161 | -0.009 | -0.034 |
| 3162 | -0.016 | 0.009 |
| 3163 | 0.019 | 0.079 |
| 3164 | -0.022 | -0.009 |
| 3165 | 0.030 | 0.055 |
| 3166 | 0.025 | 0.089 |
| 3167 | 0.081 | -0.044 |
| 3168 | -0.012 | 0.016 |
| 3169 | -0.062 | 0.049 |
| 3170 | -0.036 | 0.038 |
| 3171 | -0.052 | -0.004 |
| 3172 | 0.006 | -0.002 |
| 3173 | -0.013 | -0.015 |
| 3174 | -0.010 | 0.057 |
| 3175 | 0.044 | -0.110 |
| 3176 | 0.046 | -0.111 |
| 3177 | 0.023 | -0.113 |
| 3178 | -0.024 | 0.047 |
| 3179 | 0.037 | -0.100 |
| 3180 | 0.038 | 0.006 |
| 3181 | 0.021 | 0.018 |
| 3182 | -0.030 | 0.102 |
| 3183 | 0.000 | 0.070 |
| 3184 | 0.003 | 0.124 |
| 3185 | -0.047 | -0.083 |
| 3186 | -0.018 | 0.034 |
| 3187 | 0.006 | -0.055 |
| 3188 | -0.015 | -0.088 |
| 3189 | -0.030 | 0.071 |
| 3190 | -0.025 | -0.017 |
| 3191 | 0.037 | -0.028 |
| 3192 | -0.068 | -0.047 |
| 3193 | 0.029 | -0.138 |
| 3194 | -0.007 | -0.105 |
| 3195 | -0.005 | -0.003 |
| 3196 | -0.038 | -0.076 |
| 3197 | -0.044 | 0.112 |
| 3198 | 0.009 | -0.038 |
| 3199 | -0.014 | 0.032 |
| 3200 | 0.006 | 0.012 |
| 3201 | 0.010 | 0.028 |
| 3202 | -0.026 | 0.109 |
| 3203 | 0.010 | -0.055 |
| 3204 | 0.015 | 0.027 |
| 3205 | 0.032 | -0.022 |
| 3206 | -0.016 | -0.040 |

| | | |
|---|---|---|
| 3207 | -0.053 | -0.083 |
| 3208 | 0.074 | -0.127 |
| 3209 | 0.034 | -0.083 |
| 3210 | -0.034 | 0.011 |
| 3211 | -0.011 | -0.082 |
| 3212 | 0.047 | -0.079 |
| 3213 | 0.058 | 0.114 |
| 3214 | 0.040 | -0.007 |
| 3215 | 0.043 | -0.074 |
| 3216 | 0.012 | 0.066 |
| 3217 | 0.008 | 0.126 |
| 3218 | 0.016 | -0.091 |
| 3219 | 0.002 | -0.035 |
| 3220 | -0.005 | 0.073 |
| 3221 | -0.045 | -0.057 |
| 3222 | 0.016 | 0.032 |
| 3223 | -0.003 | -0.130 |
| 3224 | -0.001 | -0.026 |
| 3225 | 0.022 | 0.112 |
| 3226 | -0.047 | 0.005 |
| 3227 | -0.026 | 0.056 |
| 3228 | -0.010 | -0.019 |
| 3229 | -0.031 | -0.105 |
| 3230 | -0.062 | 0.045 |
| 3231 | 0.058 | 0.154 |
| 3232 | -0.026 | -0.108 |
| 3233 | 0.037 | 0.074 |
| 3234 | -0.024 | -0.003 |
| 3235 | -0.051 | -0.044 |
| 3236 | -0.010 | -0.037 |
| 3237 | -0.054 | 0.150 |
| 3238 | -0.013 | -0.085 |
| 3239 | 0.048 | -0.026 |
| 3240 | 0.017 | -0.049 |
| 3241 | -0.025 | 0.012 |
| 3242 | -0.096 | 0.117 |
| 3243 | -0.020 | -0.090 |
| 3244 | 0.000 | 0.094 |
| 3245 | 0.011 | -0.174 |
| 3246 | 0.036 | -0.106 |
| 3247 | -0.040 | 0.056 |
| 3248 | 0.045 | 0.069 |
| 3249 | 0.045 | -0.044 |
| 3250 | -0.018 | 0.004 |
| 3251 | 0.017 | 0.079 |
| 3252 | -0.014 | -0.004 |
| 3253 | 0.010 | 0.039 |
| 3254 | 0.002 | 0.070 |
| 3255 | 0.028 | -0.027 |
| 3256 | 0.008 | 0.021 |
| 3257 | 0.009 | 0.017 |

| | | |
|---|---|---|
| 3258 | -0.037 | -0.108 |
| 3259 | 0.047 | 0.053 |
| 3260 | 0.031 | -0.059 |
| 3261 | 0.002 | 0.026 |
| 3262 | -0.023 | 0.106 |
| 3263 | 0.000 | 0.033 |
| 3264 | 0.042 | 0.091 |
| 3265 | -0.014 | -0.076 |
| 3266 | 0.050 | 0.064 |
| 3267 | 0.019 | -0.010 |
| 3268 | -0.003 | -0.108 |
| 3269 | -0.025 | 0.049 |
| 3270 | -0.051 | 0.117 |
| 3271 | -0.011 | -0.044 |
| 3272 | 0.039 | 0.009 |
| 3273 | 0.039 | 0.048 |
| 3274 | -0.047 | -0.003 |
| 3275 | 0.018 | -0.042 |
| 3276 | 0.019 | 0.024 |
| 3277 | -0.018 | -0.028 |
| 3278 | 0.059 | 0.003 |
| 3279 | 0.056 | -0.006 |
| 3280 | 0.021 | -0.027 |
| 3281 | -0.015 | -0.048 |
| 3282 | -0.030 | -0.057 |
| 3283 | 0.014 | -0.050 |
| 3284 | 0.020 | 0.030 |
| 3285 | 0.006 | -0.015 |
| 3286 | -0.013 | 0.019 |
| 3287 | -0.028 | -0.122 |
| 3288 | -0.020 | 0.005 |
| 3289 | -0.044 | 0.193 |
| 3290 | -0.065 | 0.139 |
| 3291 | -0.015 | 0.039 |
| 3292 | -0.024 | -0.068 |
| 3293 | 0.005 | -0.055 |
| 3294 | 0.008 | -0.045 |
| 3295 | -0.017 | -0.002 |
| 3296 | 0.042 | 0.071 |
| 3297 | -0.033 | 0.015 |
| 3298 | -0.031 | -0.017 |
| 3299 | 0.022 | -0.012 |
| 3300 | -0.005 | 0.002 |
| 3301 | 0.038 | -0.209 |
| 3302 | -0.024 | -0.083 |
| 3303 | 0.007 | 0.010 |
| 3304 | 0.020 | 0.051 |
| 3305 | 0.038 | -0.039 |
| 3306 | -0.057 | -0.043 |
| 3307 | 0.009 | 0.098 |
| 3308 | 0.016 | 0.000 |

| | | |
|---|---|---|
| 3309 | -0.019 | -0.111 |
| 3310 | 0.022 | 0.037 |
| 3311 | -0.029 | -0.032 |
| 3312 | 0.012 | 0.010 |
| 3313 | -0.045 | 0.081 |
| 3314 | 0.045 | 0.080 |
| 3315 | -0.026 | -0.053 |
| 3316 | -0.008 | 0.106 |
| 3317 | -0.038 | 0.000 |
| 3318 | 0.029 | -0.090 |
| 3319 | 0.008 | -0.139 |
| 3320 | -0.006 | -0.034 |
| 3321 | 0.017 | -0.064 |
| 3322 | -0.035 | 0.142 |
| 3323 | -0.043 | -0.004 |
| 3324 | -0.012 | 0.158 |
| 3325 | 0.021 | -0.038 |
| 3326 | -0.013 | 0.016 |
| 3327 | 0.019 | -0.105 |
| 3328 | 0.069 | 0.041 |
| 3329 | -0.012 | -0.091 |
| 3330 | 0.035 | 0.034 |
| 3331 | 0.009 | -0.059 |
| 3332 | 0.009 | 0.026 |
| 3333 | 0.065 | -0.028 |
| 3334 | -0.040 | 0.026 |
| 3335 | -0.042 | -0.069 |
| 3336 | -0.024 | 0.063 |
| 3337 | -0.008 | -0.053 |
| 3338 | -0.001 | -0.027 |
| 3339 | 0.016 | 0.023 |
| 3340 | -0.016 | -0.050 |
| 3341 | -0.022 | -0.080 |
| 3342 | 0.022 | -0.023 |
| 3343 | 0.031 | -0.133 |
| 3344 | 0.031 | 0.136 |
| 3345 | -0.080 | -0.027 |
| 3346 | 0.001 | -0.042 |
| 3347 | -0.009 | -0.013 |
| 3348 | 0.022 | 0.000 |
| 3349 | 0.034 | -0.048 |
| 3350 | -0.064 | -0.007 |
| 3351 | -0.034 | -0.048 |
| 3352 | -0.085 | -0.094 |
| 3353 | 0.027 | -0.003 |
| 3354 | -0.071 | 0.037 |
| 3355 | -0.015 | -0.068 |
| 3356 | -0.012 | 0.031 |
| 3357 | 0.054 | 0.004 |
| 3358 | 0.023 | -0.022 |
| 3359 | -0.006 | 0.052 |

| | | |
|---|---|---|
| 3360 | 0.022 | 0.045 |
| 3361 | 0.014 | -0.080 |
| 3362 | -0.018 | 0.080 |
| 3363 | 0.036 | -0.191 |
| 3364 | -0.010 | -0.002 |
| 3365 | 0.018 | -0.047 |
| 3366 | -0.042 | -0.076 |
| 3367 | 0.002 | -0.004 |
| 3368 | -0.055 | -0.061 |
| 3369 | 0.088 | -0.053 |
| 3370 | 0.055 | 0.060 |
| 3371 | -0.004 | -0.041 |
| 3372 | 0.067 | -0.131 |
| 3373 | 0.001 | -0.069 |
| 3374 | -0.047 | 0.030 |
| 3375 | 0.102 | 0.083 |
| 3376 | 0.020 | 0.043 |
| 3377 | -0.030 | 0.032 |
| 3378 | -0.028 | -0.012 |
| 3379 | -0.039 | 0.019 |
| 3380 | 0.014 | 0.017 |
| 3381 | 0.056 | 0.059 |
| 3382 | -0.015 | 0.041 |
| 3383 | 0.003 | -0.064 |
| 3384 | 0.007 | 0.089 |
| 3385 | -0.055 | 0.100 |
| 3386 | -0.008 | 0.127 |
| 3387 | -0.029 | -0.052 |
| 3388 | 0.059 | 0.044 |
| 3389 | -0.043 | -0.010 |
| 3390 | 0.004 | -0.005 |
| 3391 | -0.023 | 0.057 |
| 3392 | 0.044 | -0.015 |
| 3393 | -0.035 | -0.032 |
| 3394 | -0.046 | -0.060 |
| 3395 | -0.017 | 0.003 |
| 3396 | -0.032 | 0.013 |
| 3397 | 0.002 | -0.020 |
| 3398 | -0.018 | -0.013 |
| 3399 | -0.051 | -0.017 |
| 3400 | -0.024 | 0.014 |
| 3401 | -0.037 | 0.099 |
| 3402 | 0.018 | 0.011 |
| 3403 | 0.024 | 0.082 |
| 3404 | -0.001 | 0.049 |
| 3405 | -0.062 | -0.020 |
| 3406 | 0.057 | 0.076 |
| 3407 | 0.009 | -0.088 |
| 3408 | 0.004 | 0.043 |
| 3409 | -0.050 | -0.126 |
| 3410 | -0.041 | 0.018 |

| | | |
|---|---|---|
| 3411 | -0.043 | -0.070 |
| 3412 | 0.031 | 0.077 |
| 3413 | 0.029 | -0.121 |
| 3414 | -0.048 | -0.025 |
| 3415 | 0.002 | 0.020 |
| 3416 | -0.038 | 0.081 |
| 3417 | -0.003 | 0.014 |
| 3418 | 0.042 | 0.062 |
| 3419 | -0.031 | -0.028 |
| 3420 | 0.010 | -0.045 |
| 3421 | 0.015 | 0.030 |
| 3422 | 0.040 | 0.055 |
| 3423 | -0.014 | 0.039 |
| 3424 | 0.024 | -0.067 |
| 3425 | 0.030 | -0.080 |
| 3426 | -0.004 | 0.044 |
| 3427 | -0.030 | -0.004 |
| 3428 | 0.006 | -0.103 |
| 3429 | 0.018 | 0.001 |
| 3430 | -0.009 | 0.098 |
| 3431 | 0.005 | -0.016 |
| 3432 | 0.004 | 0.013 |
| 3433 | -0.012 | 0.038 |
| 3434 | -0.025 | -0.046 |
| 3435 | -0.041 | 0.034 |
| 3436 | -0.053 | 0.072 |
| 3437 | -0.020 | 0.107 |
| 3438 | 0.024 | 0.027 |
| 3439 | -0.075 | 0.118 |
| 3440 | 0.015 | 0.032 |
| 3441 | 0.040 | 0.018 |
| 3442 | 0.070 | 0.107 |
| 3443 | -0.006 | 0.029 |
| 3444 | 0.042 | -0.180 |
| 3445 | 0.038 | 0.038 |
| 3446 | -0.031 | 0.038 |
| 3447 | 0.016 | 0.037 |
| 3448 | 0.038 | 0.104 |
| 3449 | -0.025 | 0.021 |
| 3450 | -0.029 | 0.055 |
| 3451 | 0.026 | 0.082 |
| 3452 | -0.050 | 0.060 |
| 3453 | 0.024 | -0.049 |
| 3454 | -0.061 | 0.111 |
| 3455 | 0.021 | 0.055 |
| 3456 | 0.050 | 0.100 |
| 3457 | 0.024 | 0.048 |
| 3458 | 0.013 | -0.091 |
| 3459 | 0.013 | -0.015 |
| 3460 | 0.007 | -0.072 |
| 3461 | -0.015 | -0.003 |

| | | |
|---|---|---|
| 3462 | -0.050 | 0.064 |
| 3463 | 0.104 | -0.059 |
| 3464 | -0.018 | -0.047 |
| 3465 | 0.000 | 0.125 |
| 3466 | -0.054 | 0.064 |
| 3467 | -0.028 | -0.004 |
| 3468 | 0.036 | 0.086 |
| 3469 | 0.003 | -0.150 |
| 3470 | -0.011 | -0.037 |
| 3471 | -0.043 | -0.109 |
| 3472 | -0.065 | 0.016 |
| 3473 | 0.082 | -0.180 |
| 3474 | 0.013 | 0.064 |
| 3475 | -0.004 | 0.076 |
| 3476 | -0.016 | 0.067 |
| 3477 | -0.015 | -0.007 |
| 3478 | 0.013 | -0.014 |
| 3479 | -0.035 | 0.026 |
| 3480 | -0.009 | -0.047 |
| 3481 | -0.016 | 0.027 |
| 3482 | -0.017 | 0.051 |
| 3483 | -0.015 | 0.100 |
| 3484 | 0.052 | 0.014 |
| 3485 | 0.013 | -0.009 |
| 3486 | -0.028 | 0.014 |
| 3487 | -0.010 | -0.081 |
| 3488 | 0.064 | 0.084 |
| 3489 | 0.014 | -0.035 |
| 3490 | 0.006 | -0.003 |
| 3491 | -0.002 | -0.062 |
| 3492 | -0.109 | -0.028 |
| 3493 | 0.000 | 0.024 |
| 3494 | 0.021 | 0.016 |
| 3495 | -0.015 | -0.075 |
| 3496 | -0.019 | 0.065 |
| 3497 | 0.010 | -0.053 |
| 3498 | -0.050 | -0.020 |
| 3499 | 0.045 | -0.016 |
| 3500 | -0.046 | -0.117 |
| 3501 | 0.025 | -0.024 |
| 3502 | 0.000 | 0.044 |
| 3503 | -0.073 | -0.105 |
| 3504 | 0.061 | 0.073 |
| 3505 | 0.006 | -0.018 |
| 3506 | 0.007 | 0.013 |
| 3507 | -0.018 | -0.106 |
| 3508 | -0.033 | -0.094 |
| 3509 | -0.032 | 0.079 |
| 3510 | 0.034 | -0.076 |
| 3511 | -0.003 | -0.146 |
| 3512 | 0.058 | 0.055 |

| | | |
|---|---|---|
| 3513 | 0.005 | 0.115 |
| 3514 | -0.006 | -0.003 |
| 3515 | -0.014 | 0.200 |
| 3516 | 0.011 | 0.115 |
| 3517 | -0.076 | 0.121 |
| 3518 | -0.010 | -0.054 |
| 3519 | -0.003 | -0.113 |
| 3520 | 0.030 | -0.035 |
| 3521 | -0.028 | -0.118 |
| 3522 | -0.003 | -0.144 |
| 3523 | -0.037 | 0.102 |
| 3524 | 0.022 | -0.023 |
| 3525 | 0.001 | -0.030 |
| 3526 | -0.016 | 0.108 |
| 3527 | 0.022 | 0.017 |
| 3528 | 0.011 | 0.044 |
| 3529 | 0.035 | -0.130 |
| 3530 | 0.008 | 0.040 |
| 3531 | -0.004 | 0.014 |
| 3532 | 0.005 | -0.045 |
| 3533 | 0.016 | 0.050 |
| 3534 | 0.047 | 0.031 |
| 3535 | 0.021 | 0.000 |
| 3536 | -0.014 | 0.047 |
| 3537 | 0.029 | 0.018 |
| 3538 | 0.025 | 0.061 |
| 3539 | -0.075 | -0.069 |
| 3540 | -0.035 | -0.025 |
| 3541 | 0.008 | -0.045 |
| 3542 | -0.073 | -0.154 |
| 3543 | 0.044 | 0.021 |
| 3544 | 0.010 | -0.079 |
| 3545 | 0.033 | 0.078 |
| 3546 | 0.041 | -0.014 |
| 3547 | -0.030 | 0.032 |
| 3548 | 0.009 | 0.189 |
| 3549 | -0.109 | -0.004 |
| 3550 | 0.055 | 0.094 |
| 3551 | -0.012 | -0.001 |
| 3552 | 0.036 | -0.047 |
| 3553 | 0.046 | 0.048 |
| 3554 | -0.006 | -0.030 |
| 3555 | 0.007 | 0.080 |
| 3556 | 0.013 | -0.047 |
| 3557 | 0.023 | -0.027 |
| 3558 | 0.021 | 0.023 |
| 3559 | -0.024 | 0.033 |
| 3560 | 0.034 | -0.090 |
| 3561 | 0.017 | 0.127 |
| 3562 | 0.032 | 0.031 |
| 3563 | 0.063 | -0.074 |

| | | |
|---|---|---|
| 3564 | 0.029 | 0.044 |
| 3565 | -0.057 | 0.016 |
| 3566 | 0.035 | -0.066 |
| 3567 | 0.016 | -0.110 |
| 3568 | 0.026 | 0.030 |
| 3569 | -0.051 | -0.102 |
| 3570 | -0.052 | 0.143 |
| 3571 | -0.012 | -0.043 |
| 3572 | -0.002 | -0.017 |
| 3573 | 0.007 | 0.045 |
| 3574 | 0.020 | -0.056 |
| 3575 | -0.054 | 0.049 |
| 3576 | -0.004 | 0.011 |
| 3577 | -0.021 | -0.011 |
| 3578 | 0.003 | 0.031 |
| 3579 | 0.029 | 0.065 |
| 3580 | 0.057 | 0.047 |
| 3581 | 0.023 | -0.024 |
| 3582 | 0.007 | -0.007 |
| 3583 | -0.025 | 0.096 |
| 3584 | 0.019 | 0.016 |
| 3585 | 0.023 | -0.083 |
| 3586 | -0.057 | 0.026 |
| 3587 | -0.037 | 0.018 |
| 3588 | -0.049 | 0.041 |
| 3589 | 0.060 | -0.024 |
| 3590 | 0.015 | 0.050 |
| 3591 | -0.006 | -0.024 |
| 3592 | -0.025 | -0.155 |
| 3593 | 0.081 | 0.026 |
| 3594 | -0.023 | -0.037 |
| 3595 | 0.016 | 0.044 |
| 3596 | 0.002 | -0.071 |
| 3597 | 0.026 | -0.080 |
| 3598 | 0.017 | -0.068 |
| 3599 | -0.087 | 0.017 |
| 3600 | 0.062 | 0.028 |
| 3601 | 0.011 | 0.003 |
| 3602 | -0.030 | -0.070 |
| 3603 | -0.032 | 0.089 |
| 3604 | -0.095 | -0.036 |
| 3605 | -0.024 | -0.018 |
| 3606 | -0.046 | -0.146 |
| 3607 | -0.015 | -0.099 |
| 3608 | -0.031 | -0.075 |
| 3609 | 0.046 | 0.077 |
| 3610 | -0.026 | 0.040 |
| 3611 | 0.009 | -0.038 |
| 3612 | 0.014 | 0.043 |
| 3613 | 0.006 | -0.063 |
| 3614 | -0.009 | 0.071 |

| | | |
|---|---|---|
| 3615 | -0.045 | -0.023 |
| 3616 | 0.056 | 0.154 |
| 3617 | 0.040 | 0.022 |
| 3618 | -0.068 | -0.034 |
| 3619 | -0.098 | 0.053 |
| 3620 | -0.012 | -0.097 |
| 3621 | 0.012 | 0.006 |
| 3622 | -0.033 | -0.028 |
| 3623 | 0.062 | 0.069 |
| 3624 | 0.001 | -0.023 |
| 3625 | 0.005 | 0.121 |
| 3626 | -0.004 | 0.026 |
| 3627 | 0.025 | 0.043 |
| 3628 | 0.072 | 0.011 |
| 3629 | -0.025 | -0.048 |
| 3630 | 0.018 | 0.039 |
| 3631 | 0.003 | 0.008 |
| 3632 | 0.023 | -0.076 |
| 3633 | 0.003 | -0.014 |
| 3634 | 0.011 | -0.075 |
| 3635 | 0.020 | -0.160 |
| 3636 | 0.050 | -0.178 |
| 3637 | 0.008 | -0.136 |
| 3638 | -0.003 | 0.049 |
| 3639 | -0.005 | -0.126 |
| 3640 | 0.006 | -0.092 |
| 3641 | 0.019 | 0.005 |
| 3642 | 0.003 | 0.006 |
| 3643 | 0.003 | 0.004 |
| 3644 | 0.007 | 0.057 |
| 3645 | 0.008 | 0.021 |
| 3646 | 0.025 | 0.087 |
| 3647 | 0.056 | 0.029 |
| 3648 | -0.002 | 0.085 |
| 3649 | -0.049 | 0.051 |
| 3650 | 0.058 | 0.017 |
| 3651 | -0.028 | -0.227 |
| 3652 | -0.097 | 0.118 |
| 3653 | 0.041 | 0.120 |
| 3654 | -0.034 | -0.032 |
| 3655 | 0.049 | 0.082 |
| 3656 | 0.015 | 0.062 |
| 3657 | -0.046 | -0.063 |
| 3658 | 0.036 | 0.011 |
| 3659 | 0.025 | 0.068 |
| 3660 | 0.021 | -0.041 |
| 3661 | -0.023 | 0.028 |
| 3662 | -0.039 | -0.025 |
| 3663 | -0.013 | -0.153 |
| 3664 | -0.003 | 0.084 |
| 3665 | 0.032 | -0.041 |

| | | |
|---|---|---|
| 3666 | -0.001 | -0.021 |
| 3667 | -0.032 | 0.123 |
| 3668 | 0.015 | 0.042 |
| 3669 | 0.009 | 0.106 |
| 3670 | -0.044 | -0.040 |
| 3671 | -0.058 | -0.077 |
| 3672 | 0.035 | -0.048 |
| 3673 | -0.019 | 0.041 |
| 3674 | -0.026 | -0.080 |
| 3675 | 0.019 | 0.086 |
| 3676 | -0.014 | -0.023 |
| 3677 | 0.084 | 0.186 |
| 3678 | -0.032 | -0.002 |
| 3679 | -0.008 | 0.008 |
| 3680 | 0.053 | -0.094 |
| 3681 | 0.007 | -0.066 |
| 3682 | -0.036 | 0.032 |
| 3683 | 0.011 | -0.147 |
| 3684 | 0.076 | -0.062 |
| 3685 | -0.013 | 0.017 |
| 3686 | 0.008 | 0.118 |
| 3687 | -0.004 | -0.016 |
| 3688 | -0.002 | 0.051 |
| 3689 | -0.039 | -0.036 |
| 3690 | -0.023 | 0.017 |
| 3691 | -0.026 | 0.071 |
| 3692 | 0.011 | 0.081 |
| 3693 | -0.023 | -0.102 |
| 3694 | -0.002 | 0.057 |
| 3695 | 0.033 | -0.074 |
| 3696 | -0.055 | -0.070 |
| 3697 | 0.083 | 0.014 |
| 3698 | 0.013 | 0.002 |
| 3699 | 0.044 | 0.119 |
| 3700 | 0.052 | 0.031 |
| 3701 | 0.032 | -0.029 |
| 3702 | -0.055 | -0.023 |
| 3703 | -0.037 | -0.220 |
| 3704 | -0.012 | 0.017 |
| 3705 | -0.017 | 0.048 |
| 3706 | -0.040 | -0.013 |
| 3707 | 0.024 | -0.075 |
| 3708 | -0.021 | -0.034 |
| 3709 | -0.069 | -0.036 |
| 3710 | 0.038 | 0.045 |
| 3711 | 0.063 | 0.033 |
| 3712 | -0.013 | -0.007 |
| 3713 | 0.067 | -0.027 |
| 3714 | 0.056 | -0.046 |
| 3715 | -0.028 | -0.029 |
| 3716 | -0.013 | 0.004 |

| | | |
|---|---|---|
| 3717 | 0.017 | -0.138 |
| 3718 | 0.052 | -0.040 |
| 3719 | -0.074 | 0.004 |
| 3720 | 0.032 | -0.108 |
| 3721 | -0.005 | -0.004 |
| 3722 | -0.033 | -0.082 |
| 3723 | 0.102 | 0.002 |
| 3724 | 0.013 | 0.105 |
| 3725 | 0.025 | 0.008 |
| 3726 | -0.049 | -0.069 |
| 3727 | -0.014 | 0.001 |
| 3728 | 0.020 | -0.057 |
| 3729 | 0.022 | 0.040 |
| 3730 | -0.032 | -0.098 |
| 3731 | -0.026 | -0.055 |
| 3732 | -0.040 | -0.009 |
| 3733 | -0.024 | 0.007 |
| 3734 | 0.036 | 0.123 |
| 3735 | -0.051 | -0.025 |
| 3736 | 0.030 | 0.029 |
| 3737 | -0.018 | 0.020 |
| 3738 | 0.052 | -0.069 |
| 3739 | 0.037 | 0.065 |
| 3740 | -0.006 | -0.053 |
| 3741 | 0.030 | -0.158 |
| 3742 | -0.046 | 0.026 |
| 3743 | -0.029 | 0.131 |
| 3744 | -0.029 | -0.038 |
| 3745 | -0.011 | 0.137 |
| 3746 | 0.020 | 0.066 |
| 3747 | -0.011 | -0.092 |
| 3748 | -0.026 | 0.125 |
| 3749 | -0.058 | 0.042 |
| 3750 | 0.008 | -0.045 |
| 3751 | 0.005 | 0.005 |
| 3752 | 0.039 | -0.037 |
| 3753 | -0.049 | -0.023 |
| 3754 | -0.025 | 0.020 |
| 3755 | 0.013 | -0.024 |
| 3756 | 0.002 | -0.114 |
| 3757 | -0.011 | 0.018 |
| 3758 | -0.037 | 0.167 |
| 3759 | -0.033 | 0.034 |
| 3760 | 0.009 | -0.024 |
| 3761 | 0.021 | -0.030 |
| 3762 | 0.030 | 0.018 |
| 3763 | -0.034 | -0.028 |
| 3764 | 0.016 | -0.090 |
| 3765 | 0.040 | -0.113 |
| 3766 | -0.006 | 0.156 |
| 3767 | -0.037 | 0.068 |

| | | |
|---|---|---|
| 3768 | 0.026 | 0.027 |
| 3769 | 0.011 | -0.010 |
| 3770 | 0.039 | 0.030 |
| 3771 | -0.050 | 0.050 |
| 3772 | -0.008 | -0.019 |
| 3773 | 0.064 | -0.038 |
| 3774 | 0.016 | 0.020 |
| 3775 | 0.001 | -0.030 |
| 3776 | 0.017 | 0.072 |
| 3777 | -0.006 | 0.048 |
| 3778 | -0.007 | 0.015 |
| 3779 | -0.006 | -0.028 |
| 3780 | 0.064 | -0.076 |
| 3781 | 0.004 | -0.027 |
| 3782 | 0.044 | 0.036 |
| 3783 | 0.002 | 0.062 |
| 3784 | 0.013 | -0.114 |
| 3785 | 0.030 | -0.005 |
| 3786 | 0.039 | 0.072 |
| 3787 | 0.014 | -0.119 |
| 3788 | 0.037 | 0.041 |
| 3789 | 0.045 | 0.041 |
| 3790 | 0.024 | 0.007 |
| 3791 | -0.017 | -0.026 |
| 3792 | -0.076 | 0.013 |
| 3793 | 0.003 | -0.016 |
| 3794 | -0.021 | -0.008 |
| 3795 | 0.012 | 0.139 |
| 3796 | -0.066 | -0.035 |
| 3797 | 0.008 | -0.036 |
| 3798 | -0.004 | -0.037 |
| 3799 | 0.040 | 0.001 |
| 3800 | -0.029 | -0.034 |
| 3801 | 0.044 | -0.031 |
| 3802 | 0.007 | -0.070 |
| 3803 | 0.057 | -0.173 |
| 3804 | 0.002 | 0.023 |
| 3805 | 0.015 | -0.041 |
| 3806 | -0.049 | -0.044 |
| 3807 | -0.024 | -0.036 |
| 3808 | 0.032 | 0.048 |
| 3809 | -0.059 | -0.037 |
| 3810 | -0.001 | 0.052 |
| 3811 | 0.018 | -0.139 |
| 3812 | 0.031 | 0.122 |
| 3813 | -0.063 | 0.002 |
| 3814 | 0.036 | 0.029 |
| 3815 | -0.031 | 0.030 |
| 3816 | -0.007 | 0.041 |
| 3817 | 0.077 | -0.003 |
| 3818 | 0.016 | -0.027 |

| | | |
|---|---|---|
| 3819 | 0.006 | 0.015 |
| 3820 | -0.035 | -0.035 |
| 3821 | 0.054 | 0.071 |
| 3822 | -0.038 | 0.046 |
| 3823 | 0.021 | -0.140 |
| 3824 | 0.053 | 0.053 |
| 3825 | -0.001 | -0.012 |
| 3826 | 0.068 | -0.006 |
| 3827 | -0.023 | 0.011 |
| 3828 | -0.018 | -0.090 |
| 3829 | 0.023 | 0.117 |
| 3830 | 0.040 | 0.034 |
| 3831 | -0.018 | 0.039 |
| 3832 | 0.023 | 0.107 |
| 3833 | -0.021 | 0.053 |
| 3834 | -0.016 | -0.078 |
| 3835 | -0.099 | 0.132 |
| 3836 | -0.007 | -0.122 |
| 3837 | -0.001 | 0.045 |
| 3838 | -0.097 | -0.071 |
| 3839 | -0.032 | 0.054 |
| 3840 | -0.013 | 0.020 |
| 3841 | -0.010 | -0.102 |
| 3842 | 0.015 | 0.025 |
| 3843 | -0.051 | -0.002 |
| 3844 | 0.021 | -0.075 |
| 3845 | -0.027 | -0.102 |
| 3846 | 0.032 | 0.006 |
| 3847 | 0.005 | 0.000 |
| 3848 | 0.001 | -0.134 |
| 3849 | 0.019 | -0.098 |
| 3850 | -0.010 | -0.023 |
| 3851 | -0.057 | 0.011 |
| 3852 | 0.016 | 0.001 |
| 3853 | -0.036 | -0.031 |
| 3854 | -0.030 | 0.053 |
| 3855 | -0.003 | 0.010 |
| 3856 | 0.037 | 0.068 |
| 3857 | 0.024 | 0.029 |
| 3858 | 0.003 | -0.018 |
| 3859 | 0.004 | -0.020 |
| 3860 | 0.068 | 0.039 |
| 3861 | -0.001 | -0.111 |
| 3862 | -0.011 | -0.032 |
| 3863 | 0.016 | 0.058 |
| 3864 | 0.027 | 0.010 |
| 3865 | -0.005 | -0.082 |
| 3866 | 0.008 | -0.079 |
| 3867 | 0.008 | 0.055 |
| 3868 | 0.003 | 0.043 |
| 3869 | 0.010 | 0.097 |

| | | |
|---|---|---|
| 3870 | -0.053 | 0.023 |
| 3871 | 0.062 | 0.003 |
| 3872 | -0.042 | 0.066 |
| 3873 | 0.013 | 0.028 |
| 3874 | -0.008 | 0.091 |
| 3875 | -0.027 | -0.001 |
| 3876 | -0.023 | -0.241 |
| 3877 | -0.008 | 0.053 |
| 3878 | -0.064 | -0.027 |
| 3879 | -0.028 | -0.046 |
| 3880 | -0.042 | 0.047 |
| 3881 | 0.034 | -0.043 |
| 3882 | 0.002 | 0.020 |
| 3883 | 0.045 | 0.014 |
| 3884 | 0.008 | 0.023 |
| 3885 | 0.085 | 0.062 |
| 3886 | 0.001 | -0.086 |
| 3887 | -0.098 | 0.082 |
| 3888 | 0.000 | 0.037 |
| 3889 | -0.049 | 0.009 |
| 3890 | -0.028 | -0.099 |
| 3891 | -0.081 | -0.021 |
| 3892 | -0.039 | -0.066 |
| 3893 | -0.013 | 0.021 |
| 3894 | -0.054 | 0.108 |
| 3895 | 0.061 | 0.019 |
| 3896 | -0.014 | -0.006 |
| 3897 | 0.043 | 0.045 |
| 3898 | -0.042 | 0.003 |
| 3899 | 0.035 | -0.156 |
| 3900 | 0.025 | 0.130 |
| 3901 | -0.031 | 0.195 |
| 3902 | -0.017 | 0.098 |
| 3903 | 0.125 | -0.049 |
| 3904 | 0.063 | 0.043 |
| 3905 | -0.034 | 0.048 |
| 3906 | -0.007 | 0.036 |
| 3907 | -0.032 | -0.020 |
| 3908 | -0.010 | 0.046 |
| 3909 | 0.042 | -0.031 |
| 3910 | 0.020 | -0.038 |
| 3911 | -0.038 | -0.029 |
| 3912 | -0.084 | 0.044 |
| 3913 | -0.142 | 0.129 |
| 3914 | 0.010 | -0.083 |
| 3915 | 0.017 | 0.094 |
| 3916 | 0.041 | -0.015 |
| 3917 | 0.029 | 0.090 |
| 3918 | 0.058 | 0.051 |
| 3919 | 0.007 | 0.059 |
| 3920 | -0.055 | -0.031 |

| | | |
|---|---|---|
| 3921 | 0.070 | 0.016 |
| 3922 | 0.010 | 0.091 |
| 3923 | -0.073 | -0.016 |
| 3924 | 0.022 | -0.047 |
| 3925 | -0.013 | 0.102 |
| 3926 | 0.011 | -0.086 |
| 3927 | 0.019 | 0.016 |
| 3928 | 0.037 | -0.009 |
| 3929 | 0.000 | -0.081 |
| 3930 | 0.031 | 0.076 |
| 3931 | 0.026 | 0.135 |
| 3932 | -0.051 | -0.082 |
| 3933 | 0.034 | -0.011 |
| 3934 | 0.047 | -0.114 |
| 3935 | -0.015 | 0.024 |
| 3936 | 0.013 | -0.002 |
| 3937 | -0.002 | 0.092 |
| 3938 | 0.004 | 0.085 |
| 3939 | 0.050 | -0.038 |
| 3940 | -0.030 | -0.082 |
| 3941 | 0.005 | -0.033 |
| 3942 | -0.048 | -0.014 |
| 3943 | 0.014 | -0.047 |
| 3944 | 0.022 | -0.099 |
| 3945 | 0.019 | 0.052 |
| 3946 | -0.020 | 0.063 |
| 3947 | 0.003 | 0.053 |
| 3948 | 0.098 | -0.027 |
| 3949 | 0.042 | 0.116 |
| 3950 | 0.029 | 0.056 |
| 3951 | -0.029 | 0.048 |
| 3952 | -0.023 | -0.102 |
| 3953 | 0.000 | 0.114 |
| 3954 | -0.058 | -0.063 |
| 3955 | -0.027 | -0.065 |
| 3956 | 0.044 | -0.149 |
| 3957 | 0.007 | -0.150 |
| 3958 | -0.050 | 0.089 |
| 3959 | -0.017 | -0.095 |
| 3960 | -0.009 | -0.022 |
| 3961 | 0.039 | 0.089 |
| 3962 | 0.018 | -0.073 |
| 3963 | -0.066 | 0.067 |
| 3964 | 0.010 | 0.049 |
| 3965 | 0.041 | 0.030 |
| 3966 | -0.113 | 0.021 |
| 3967 | 0.022 | -0.003 |
| 3968 | 0.099 | 0.017 |
| 3969 | -0.001 | -0.085 |
| 3970 | 0.007 | 0.003 |
| 3971 | -0.019 | -0.040 |

| | | |
|---|---|---|
| 3972 | 0.021 | 0.078 |
| 3973 | 0.029 | -0.017 |
| 3974 | -0.021 | -0.074 |
| 3975 | 0.037 | -0.098 |
| 3976 | 0.011 | 0.113 |
| 3977 | 0.018 | -0.102 |
| 3978 | 0.026 | -0.007 |
| 3979 | 0.008 | 0.010 |
| 3980 | 0.008 | 0.043 |
| 3981 | 0.007 | -0.030 |
| 3982 | 0.006 | -0.049 |
| 3983 | 0.006 | -0.008 |
| 3984 | 0.064 | 0.046 |
| 3985 | 0.066 | -0.113 |
| 3986 | -0.019 | -0.102 |
| 3987 | -0.068 | -0.005 |
| 3988 | -0.007 | -0.019 |
| 3989 | 0.013 | 0.001 |
| 3990 | -0.015 | 0.000 |
| 3991 | 0.049 | 0.056 |
| 3992 | 0.017 | -0.048 |
| 3993 | -0.032 | -0.112 |
| 3994 | -0.060 | -0.062 |
| 3995 | 0.073 | 0.109 |
| 3996 | 0.080 | 0.060 |
| 3997 | 0.074 | 0.070 |
| 3998 | -0.015 | 0.031 |
| 3999 | -0.017 | 0.027 |
| 4000 | -0.003 | -0.011 |
| 4001 | -0.009 | -0.104 |
| 4002 | -0.052 | 0.065 |
| 4003 | 0.018 | 0.116 |
| 4004 | 0.031 | 0.088 |
| 4005 | -0.001 | 0.035 |
| 4006 | -0.045 | 0.086 |
| 4007 | 0.004 | -0.031 |
| 4008 | 0.049 | 0.002 |
| 4009 | -0.056 | -0.067 |
| 4010 | -0.021 | -0.039 |
| 4011 | 0.002 | 0.020 |
| 4012 | 0.102 | -0.066 |
| 4013 | -0.030 | 0.010 |
| 4014 | 0.019 | -0.031 |
| 4015 | 0.000 | -0.088 |
| 4016 | 0.015 | 0.039 |
| 4017 | -0.007 | 0.008 |
| 4018 | -0.009 | 0.011 |
| 4019 | -0.002 | -0.045 |
| 4020 | -0.079 | -0.006 |
| 4021 | -0.060 | -0.097 |
| 4022 | 0.013 | -0.016 |

| | | |
|---|---|---|
| 4023 | -0.045 | -0.123 |
| 4024 | -0.008 | -0.049 |
| 4025 | -0.023 | -0.021 |
| 4026 | 0.037 | 0.023 |
| 4027 | 0.001 | -0.058 |
| 4028 | -0.030 | -0.028 |
| 4029 | -0.067 | -0.081 |
| 4030 | 0.025 | 0.012 |
| 4031 | 0.032 | 0.072 |
| 4032 | 0.045 | -0.047 |
| 4033 | 0.009 | -0.035 |
| 4034 | -0.024 | 0.126 |
| 4035 | -0.032 | 0.041 |
| 4036 | -0.020 | -0.081 |
| 4037 | 0.044 | -0.126 |
| 4038 | 0.007 | -0.195 |
| 4039 | 0.003 | 0.059 |
| 4040 | 0.045 | 0.026 |
| 4041 | -0.019 | -0.096 |
| 4042 | 0.016 | -0.036 |
| 4043 | -0.019 | 0.028 |
| 4044 | 0.001 | -0.017 |
| 4045 | -0.011 | 0.053 |
| 4046 | 0.005 | 0.063 |
| 4047 | 0.014 | -0.195 |
| 4048 | 0.008 | 0.040 |
| 4049 | -0.006 | 0.214 |
| 4050 | -0.009 | 0.038 |
| 4051 | -0.060 | 0.034 |
| 4052 | -0.002 | 0.093 |
| 4053 | -0.005 | -0.080 |
| 4054 | 0.065 | 0.033 |
| 4055 | 0.011 | 0.086 |
| 4056 | 0.083 | -0.028 |
| 4057 | 0.080 | -0.041 |
| 4058 | 0.007 | -0.195 |
| 4059 | 0.029 | -0.067 |
| 4060 | -0.062 | 0.012 |
| 4061 | 0.013 | 0.060 |
| 4062 | -0.028 | 0.094 |
| 4063 | 0.038 | -0.108 |
| 4064 | 0.052 | 0.031 |
| 4065 | -0.030 | -0.014 |
| 4066 | 0.021 | 0.048 |
| 4067 | 0.087 | -0.007 |
| 4068 | 0.083 | 0.061 |
| 4069 | 0.056 | -0.053 |
| 4070 | -0.051 | -0.031 |
| 4071 | -0.062 | -0.018 |
| 4072 | 0.017 | 0.004 |
| 4073 | 0.023 | -0.019 |

| | | |
|---|---|---|
| 4074 | 0.099 | 0.035 |
| 4075 | 0.008 | 0.064 |
| 4076 | -0.031 | 0.061 |
| 4077 | -0.002 | -0.032 |
| 4078 | 0.004 | -0.051 |
| 4079 | 0.044 | -0.022 |
| 4080 | 0.049 | -0.033 |
| 4081 | -0.005 | -0.106 |
| 4082 | 0.010 | 0.044 |
| 4083 | -0.047 | -0.058 |
| 4084 | -0.031 | -0.133 |
| 4085 | 0.036 | -0.056 |
| 4086 | 0.001 | -0.020 |
| 4087 | 0.009 | 0.007 |
| 4088 | 0.022 | 0.057 |
| 4089 | -0.001 | 0.103 |
| 4090 | 0.003 | -0.053 |
| 4091 | 0.037 | -0.006 |
| 4092 | -0.044 | 0.021 |
| 4093 | -0.015 | -0.009 |
| 4094 | 0.007 | -0.029 |
| 4095 | 0.011 | -0.110 |
| 4096 | 0.030 | 0.033 |
| 4097 | 0.017 | 0.087 |
| 4098 | -0.044 | -0.094 |
| 4099 | -0.023 | 0.056 |
| 4100 | -0.048 | -0.050 |
| 4101 | -0.080 | 0.002 |
| 4102 | 0.021 | -0.059 |
| 4103 | -0.010 | 0.028 |
| 4104 | -0.029 | -0.066 |
| 4105 | 0.015 | 0.179 |
| 4106 | -0.060 | -0.044 |
| 4107 | 0.028 | -0.024 |
| 4108 | -0.010 | -0.023 |
| 4109 | -0.030 | -0.098 |
| 4110 | -0.017 | -0.059 |
| 4111 | 0.067 | 0.012 |
| 4112 | 0.006 | -0.095 |
| 4113 | -0.034 | 0.069 |
| 4114 | -0.003 | -0.015 |
| 4115 | -0.009 | -0.008 |
| 4116 | -0.044 | 0.121 |
| 4117 | -0.047 | -0.021 |
| 4118 | 0.039 | 0.000 |
| 4119 | 0.015 | -0.120 |
| 4120 | -0.034 | -0.021 |
| 4121 | -0.040 | -0.071 |
| 4122 | -0.023 | 0.030 |
| 4123 | 0.008 | 0.068 |
| 4124 | -0.053 | -0.010 |

| | | |
|---|---|---|
| 4125 | 0.030 | -0.044 |
| 4126 | -0.017 | -0.011 |
| 4127 | 0.029 | -0.065 |
| 4128 | -0.063 | 0.103 |
| 4129 | 0.006 | 0.023 |
| 4130 | -0.017 | -0.028 |
| 4131 | 0.049 | -0.017 |
| 4132 | 0.007 | -0.027 |
| 4133 | 0.003 | 0.013 |
| 4134 | 0.047 | 0.022 |
| 4135 | 0.026 | 0.006 |
| 4136 | -0.070 | 0.037 |
| 4137 | 0.002 | 0.062 |
| 4138 | 0.012 | 0.050 |
| 4139 | 0.000 | 0.013 |
| 4140 | 0.013 | -0.028 |
| 4141 | 0.060 | -0.017 |
| 4142 | 0.027 | -0.123 |
| 4143 | -0.008 | 0.031 |
| 4144 | 0.009 | -0.092 |
| 4145 | 0.017 | 0.105 |
| 4146 | -0.070 | -0.041 |
| 4147 | 0.008 | 0.025 |
| 4148 | 0.021 | 0.060 |
| 4149 | -0.009 | -0.015 |
| 4150 | 0.031 | 0.001 |
| 4151 | -0.021 | 0.106 |
| 4152 | 0.000 | 0.061 |
| 4153 | 0.066 | -0.007 |
| 4154 | 0.032 | -0.006 |
| 4155 | -0.040 | 0.098 |
| 4156 | 0.022 | -0.099 |
| 4157 | 0.023 | -0.097 |
| 4158 | -0.029 | -0.048 |
| 4159 | -0.003 | 0.012 |
| 4160 | -0.004 | -0.108 |
| 4161 | 0.050 | -0.109 |
| 4162 | 0.015 | -0.054 |
| 4163 | 0.022 | 0.015 |
| 4164 | 0.026 | 0.028 |
| 4165 | -0.014 | 0.105 |
| 4166 | -0.071 | 0.014 |
| 4167 | 0.012 | 0.021 |
| 4168 | 0.028 | 0.107 |
| 4169 | 0.039 | -0.106 |
| 4170 | -0.038 | 0.025 |
| 4171 | 0.005 | 0.134 |
| 4172 | 0.008 | -0.006 |
| 4173 | 0.029 | 0.027 |
| 4174 | 0.025 | 0.138 |
| 4175 | -0.049 | 0.006 |

| | | |
|---|---|---|
| 4176 | -0.004 | 0.096 |
| 4177 | -0.019 | 0.000 |
| 4178 | -0.018 | -0.041 |
| 4179 | 0.010 | -0.067 |
| 4180 | -0.039 | 0.078 |
| 4181 | -0.013 | -0.058 |
| 4182 | -0.012 | 0.142 |
| 4183 | -0.004 | -0.142 |
| 4184 | 0.024 | -0.010 |
| 4185 | -0.009 | 0.008 |
| 4186 | 0.034 | 0.046 |
| 4187 | 0.080 | -0.076 |
| 4188 | -0.027 | 0.009 |
| 4189 | -0.023 | -0.051 |
| 4190 | 0.022 | -0.018 |
| 4191 | 0.095 | -0.051 |
| 4192 | 0.027 | -0.088 |
| 4193 | 0.026 | 0.079 |
| 4194 | -0.012 | -0.055 |
| 4195 | 0.005 | 0.133 |
| 4196 | -0.014 | -0.060 |
| 4197 | -0.029 | 0.064 |
| 4198 | 0.009 | -0.004 |
| 4199 | 0.018 | 0.073 |
| 4200 | 0.005 | 0.057 |
| 4201 | -0.041 | 0.031 |
| 4202 | 0.012 | 0.041 |
| 4203 | 0.029 | 0.015 |
| 4204 | 0.027 | -0.049 |
| 4205 | -0.023 | -0.017 |
| 4206 | 0.005 | -0.081 |
| 4207 | -0.064 | 0.064 |
| 4208 | 0.050 | 0.058 |
| 4209 | -0.014 | -0.074 |
| 4210 | 0.103 | 0.120 |
| 4211 | -0.022 | 0.019 |
| 4212 | 0.008 | 0.020 |
| 4213 | 0.026 | -0.090 |
| 4214 | 0.009 | 0.045 |
| 4215 | -0.052 | -0.163 |
| 4216 | 0.000 | -0.214 |
| 4217 | 0.021 | -0.030 |
| 4218 | 0.009 | 0.035 |
| 4219 | -0.013 | 0.002 |
| 4220 | -0.059 | -0.057 |
| 4221 | -0.005 | 0.013 |
| 4222 | -0.065 | -0.014 |
| 4223 | 0.025 | 0.015 |
| 4224 | 0.006 | -0.011 |
| 4225 | -0.002 | -0.029 |
| 4226 | -0.022 | -0.005 |

| | | |
|---|---|---|
| 4227 | 0.031 | -0.062 |
| 4228 | -0.001 | 0.052 |
| 4229 | 0.016 | 0.091 |
| 4230 | -0.012 | -0.080 |
| 4231 | 0.008 | -0.068 |
| 4232 | -0.019 | 0.084 |
| 4233 | 0.027 | -0.104 |
| 4234 | 0.013 | 0.038 |
| 4235 | 0.036 | 0.005 |
| 4236 | 0.011 | -0.027 |
| 4237 | -0.024 | 0.090 |
| 4238 | 0.018 | -0.045 |
| 4239 | -0.029 | -0.029 |
| 4240 | -0.007 | 0.004 |
| 4241 | 0.003 | 0.033 |
| 4242 | -0.011 | 0.082 |
| 4243 | -0.044 | 0.057 |
| 4244 | 0.016 | 0.009 |
| 4245 | 0.052 | 0.041 |
| 4246 | 0.031 | 0.028 |
| 4247 | 0.013 | 0.033 |
| 4248 | 0.006 | -0.146 |
| 4249 | -0.038 | 0.009 |
| 4250 | 0.028 | 0.023 |
| 4251 | -0.034 | -0.133 |
| 4252 | -0.010 | 0.008 |
| 4253 | 0.024 | -0.063 |
| 4254 | 0.012 | 0.008 |
| 4255 | -0.021 | -0.042 |
| 4256 | -0.051 | -0.010 |
| 4257 | 0.026 | 0.022 |
| 4258 | 0.002 | 0.009 |
| 4259 | 0.037 | -0.049 |
| 4260 | -0.016 | -0.014 |
| 4261 | 0.004 | -0.074 |
| 4262 | 0.004 | 0.046 |
| 4263 | -0.047 | -0.044 |
| 4264 | -0.013 | 0.032 |
| 4265 | 0.047 | -0.102 |
| 4266 | -0.014 | 0.009 |
| 4267 | -0.009 | -0.084 |
| 4268 | -0.001 | -0.026 |
| 4269 | -0.016 | -0.141 |
| 4270 | -0.084 | -0.066 |
| 4271 | 0.033 | -0.075 |
| 4272 | -0.028 | 0.071 |
| 4273 | -0.066 | 0.041 |
| 4274 | 0.076 | -0.013 |
| 4275 | -0.013 | 0.009 |
| 4276 | 0.001 | -0.059 |
| 4277 | -0.019 | 0.144 |

| | | |
|---|---|---|
| 4278 | -0.014 | 0.025 |
| 4279 | 0.036 | 0.050 |
| 4280 | 0.006 | 0.038 |
| 4281 | -0.006 | -0.031 |
| 4282 | -0.026 | 0.042 |
| 4283 | 0.029 | 0.064 |
| 4284 | 0.033 | -0.045 |
| 4285 | -0.006 | 0.203 |
| 4286 | 0.013 | -0.051 |
| 4287 | -0.006 | 0.022 |
| 4288 | 0.009 | -0.036 |
| 4289 | -0.027 | 0.019 |
| 4290 | -0.072 | -0.011 |
| 4291 | 0.005 | 0.076 |
| 4292 | 0.036 | 0.071 |
| 4293 | -0.033 | -0.085 |
| 4294 | 0.034 | -0.049 |
| 4295 | -0.004 | 0.022 |
| 4296 | 0.063 | -0.110 |
| 4297 | 0.023 | 0.080 |
| 4298 | -0.052 | 0.000 |
| 4299 | 0.087 | 0.127 |
| 4300 | -0.016 | 0.123 |
| 4301 | 0.010 | 0.016 |
| 4302 | 0.011 | 0.068 |
| 4303 | -0.043 | 0.059 |
| 4304 | -0.007 | 0.089 |
| 4305 | -0.035 | 0.003 |
| 4306 | -0.012 | 0.016 |
| 4307 | -0.005 | -0.008 |
| 4308 | 0.006 | 0.009 |
| 4309 | -0.029 | 0.009 |
| 4310 | 0.024 | 0.041 |
| 4311 | 0.000 | -0.080 |
| 4312 | -0.005 | 0.085 |
| 4313 | -0.020 | 0.141 |
| 4314 | 0.011 | -0.014 |
| 4315 | 0.065 | -0.079 |
| 4316 | -0.013 | -0.015 |
| 4317 | -0.036 | 0.054 |
| 4318 | -0.022 | 0.029 |
| 4319 | -0.002 | -0.059 |
| 4320 | 0.006 | -0.029 |
| 4321 | 0.053 | -0.026 |
| 4322 | 0.021 | -0.077 |
| 4323 | -0.007 | -0.020 |
| 4324 | 0.042 | -0.189 |
| 4325 | 0.018 | 0.001 |
| 4326 | -0.010 | -0.010 |
| 4327 | -0.018 | 0.002 |
| 4328 | -0.014 | 0.156 |

| | | |
|---|---|---|
| 4329 | -0.010 | -0.012 |
| 4330 | 0.004 | 0.111 |
| 4331 | 0.036 | 0.038 |
| 4332 | -0.060 | -0.065 |
| 4333 | -0.001 | -0.182 |
| 4334 | 0.013 | -0.084 |
| 4335 | -0.002 | 0.000 |
| 4336 | 0.052 | -0.084 |
| 4337 | -0.002 | -0.129 |
| 4338 | -0.033 | 0.031 |
| 4339 | 0.019 | -0.194 |
| 4340 | 0.023 | -0.124 |
| 4341 | -0.014 | -0.014 |
| 4342 | 0.017 | -0.050 |
| 4343 | -0.038 | -0.041 |
| 4344 | -0.012 | -0.039 |
| 4345 | -0.043 | 0.030 |
| 4346 | -0.034 | 0.027 |
| 4347 | -0.033 | -0.092 |
| 4348 | -0.061 | 0.158 |
| 4349 | 0.017 | 0.058 |
| 4350 | 0.025 | -0.005 |
| 4351 | 0.065 | 0.083 |
| 4352 | 0.015 | -0.085 |
| 4353 | -0.009 | -0.085 |
| 4354 | 0.043 | 0.008 |
| 4355 | 0.019 | -0.065 |
| 4356 | 0.045 | 0.100 |
| 4357 | -0.020 | 0.063 |
| 4358 | 0.022 | -0.015 |
| 4359 | -0.026 | 0.022 |
| 4360 | -0.008 | 0.044 |
| 4361 | -0.011 | 0.151 |
| 4362 | -0.041 | -0.043 |
| 4363 | 0.061 | -0.053 |
| 4364 | 0.003 | -0.015 |
| 4365 | 0.007 | 0.040 |
| 4366 | -0.018 | 0.004 |
| 4367 | -0.023 | 0.039 |
| 4368 | 0.003 | -0.040 |
| 4369 | 0.064 | 0.026 |
| 4370 | 0.069 | 0.024 |
| 4371 | 0.016 | 0.187 |
| 4372 | 0.037 | -0.057 |
| 4373 | -0.046 | 0.079 |
| 4374 | -0.049 | 0.029 |
| 4375 | 0.012 | 0.002 |
| 4376 | -0.021 | 0.015 |
| 4377 | -0.113 | 0.051 |
| 4378 | 0.001 | 0.000 |
| 4379 | 0.044 | 0.065 |

| | | |
|---|---|---|
| 4380 | 0.050 | 0.077 |
| 4381 | 0.059 | 0.020 |
| 4382 | -0.025 | 0.001 |
| 4383 | -0.004 | 0.011 |
| 4384 | 0.033 | 0.040 |
| 4385 | -0.025 | -0.022 |
| 4386 | -0.033 | 0.047 |
| 4387 | 0.016 | 0.099 |
| 4388 | 0.027 | -0.009 |
| 4389 | 0.008 | 0.096 |
| 4390 | 0.059 | -0.075 |
| 4391 | 0.046 | -0.102 |
| 4392 | -0.016 | 0.090 |
| 4393 | -0.015 | -0.008 |
| 4394 | 0.017 | 0.048 |
| 4395 | 0.064 | -0.034 |
| 4396 | 0.020 | -0.023 |
| 4397 | -0.007 | 0.121 |
| 4398 | -0.013 | -0.038 |
| 4399 | -0.033 | 0.173 |
| 4400 | -0.003 | 0.089 |
| 4401 | 0.002 | 0.002 |
| 4402 | -0.009 | -0.017 |
| 4403 | 0.033 | 0.061 |
| 4404 | -0.040 | 0.030 |
| 4405 | 0.011 | -0.071 |
| 4406 | 0.027 | 0.106 |
| 4407 | 0.009 | -0.095 |
| 4408 | 0.037 | 0.020 |
| 4409 | -0.048 | -0.030 |
| 4410 | -0.034 | 0.015 |
| 4411 | -0.011 | 0.145 |
| 4412 | 0.005 | 0.081 |
| 4413 | 0.040 | -0.111 |
| 4414 | -0.006 | 0.033 |
| 4415 | -0.021 | 0.013 |
| 4416 | -0.040 | -0.126 |
| 4417 | 0.033 | -0.060 |
| 4418 | -0.050 | -0.073 |
| 4419 | -0.015 | -0.099 |
| 4420 | 0.038 | -0.021 |
| 4421 | 0.042 | 0.045 |
| 4422 | -0.001 | -0.055 |
| 4423 | 0.013 | 0.002 |
| 4424 | 0.074 | 0.057 |
| 4425 | 0.036 | -0.020 |
| 4426 | 0.010 | 0.093 |
| 4427 | 0.026 | -0.035 |
| 4428 | 0.015 | 0.028 |
| 4429 | 0.057 | 0.163 |
| 4430 | 0.040 | 0.025 |

| | | |
|---|---|---|
| 4431 | -0.011 | -0.050 |
| 4432 | -0.021 | 0.008 |
| 4433 | 0.015 | -0.110 |
| 4434 | 0.035 | 0.038 |
| 4435 | -0.032 | -0.064 |
| 4436 | 0.000 | -0.044 |
| 4437 | 0.000 | -0.162 |
| 4438 | 0.011 | -0.001 |
| 4439 | -0.055 | -0.028 |
| 4440 | -0.009 | 0.020 |
| 4441 | -0.016 | -0.059 |
| 4442 | 0.048 | 0.027 |
| 4443 | 0.027 | 0.020 |
| 4444 | 0.037 | 0.068 |
| 4445 | -0.039 | -0.042 |
| 4446 | -0.016 | -0.025 |
| 4447 | 0.022 | -0.034 |
| 4448 | -0.025 | 0.001 |
| 4449 | 0.022 | -0.047 |
| 4450 | 0.025 | -0.004 |
| 4451 | -0.013 | -0.083 |
| 4452 | 0.010 | -0.069 |
| 4453 | 0.022 | 0.061 |
| 4454 | 0.040 | -0.071 |
| 4455 | 0.040 | 0.044 |
| 4456 | 0.025 | 0.027 |
| 4457 | 0.004 | 0.028 |
| 4458 | 0.004 | 0.059 |
| 4459 | -0.024 | -0.065 |
| 4460 | -0.029 | -0.026 |
| 4461 | -0.019 | -0.106 |
| 4462 | -0.003 | 0.002 |
| 4463 | -0.016 | 0.080 |
| 4464 | 0.065 | 0.105 |
| 4465 | -0.028 | 0.025 |
| 4466 | -0.019 | 0.001 |
| 4467 | -0.051 | 0.127 |
| 4468 | 0.024 | 0.130 |
| 4469 | -0.016 | 0.048 |
| 4470 | 0.055 | -0.002 |
| 4471 | 0.020 | 0.026 |
| 4472 | 0.067 | 0.055 |
| 4473 | 0.013 | 0.016 |
| 4474 | 0.003 | -0.047 |
| 4475 | -0.048 | -0.021 |
| 4476 | -0.006 | -0.002 |
| 4477 | 0.028 | 0.046 |
| 4478 | 0.015 | 0.034 |
| 4479 | -0.016 | -0.064 |
| 4480 | 0.017 | -0.045 |
| 4481 | 0.022 | -0.106 |

| | | |
|---|---|---|
| 4482 | -0.030 | -0.039 |
| 4483 | -0.043 | 0.053 |
| 4484 | -0.052 | -0.091 |
| 4485 | -0.002 | -0.015 |
| 4486 | 0.033 | 0.068 |
| 4487 | -0.001 | 0.015 |
| 4488 | -0.020 | -0.105 |
| 4489 | 0.019 | 0.027 |
| 4490 | 0.027 | -0.129 |
| 4491 | -0.002 | -0.086 |
| 4492 | 0.001 | -0.105 |
| 4493 | -0.019 | -0.048 |
| 4494 | -0.008 | -0.015 |
| 4495 | -0.070 | 0.030 |
| 4496 | 0.015 | -0.059 |
| 4497 | -0.057 | -0.024 |
| 4498 | -0.013 | 0.008 |
| 4499 | -0.002 | 0.073 |
| 4500 | -0.044 | 0.017 |
| 4501 | -0.044 | -0.063 |
| 4502 | 0.012 | 0.020 |
| 4503 | -0.026 | 0.023 |
| 4504 | -0.035 | -0.075 |
| 4505 | 0.016 | -0.055 |
| 4506 | 0.023 | 0.003 |
| 4507 | 0.010 | 0.122 |
| 4508 | 0.074 | -0.004 |
| 4509 | -0.001 | 0.022 |
| 4510 | 0.012 | -0.168 |
| 4511 | -0.009 | 0.107 |
| 4512 | 0.058 | -0.113 |
| 4513 | -0.008 | 0.065 |
| 4514 | 0.039 | -0.161 |
| 4515 | 0.034 | -0.126 |
| 4516 | -0.058 | 0.144 |
| 4517 | -0.032 | -0.033 |
| 4518 | 0.030 | -0.137 |
| 4519 | 0.047 | -0.009 |
| 4520 | -0.069 | -0.026 |
| 4521 | -0.003 | 0.115 |
| 4522 | 0.027 | 0.049 |
| 4523 | 0.026 | -0.084 |
| 4524 | -0.006 | 0.013 |
| 4525 | 0.014 | -0.032 |
| 4526 | 0.037 | 0.018 |
| 4527 | -0.010 | 0.062 |
| 4528 | 0.022 | 0.055 |
| 4529 | -0.049 | 0.029 |
| 4530 | -0.052 | -0.015 |
| 4531 | 0.011 | -0.011 |
| 4532 | 0.040 | 0.008 |

| | | |
|---|---|---|
| 4533 | -0.016 | -0.053 |
| 4534 | 0.000 | 0.106 |
| 4535 | 0.093 | 0.028 |
| 4536 | 0.027 | -0.055 |
| 4537 | -0.013 | 0.062 |
| 4538 | 0.046 | -0.131 |
| 4539 | 0.016 | 0.090 |
| 4540 | -0.066 | -0.089 |
| 4541 | 0.056 | -0.040 |
| 4542 | -0.062 | -0.016 |
| 4543 | 0.038 | -0.021 |
| 4544 | -0.051 | -0.047 |
| 4545 | 0.041 | -0.057 |
| 4546 | 0.013 | 0.076 |
| 4547 | 0.034 | 0.046 |
| 4548 | 0.039 | -0.052 |
| 4549 | -0.053 | 0.067 |
| 4550 | -0.037 | -0.039 |
| 4551 | 0.022 | 0.077 |
| 4552 | 0.026 | 0.053 |
| 4553 | 0.047 | -0.109 |
| 4554 | 0.008 | -0.076 |
| 4555 | 0.009 | -0.116 |
| 4556 | 0.005 | -0.051 |
| 4557 | -0.026 | 0.043 |
| 4558 | 0.011 | -0.010 |
| 4559 | -0.003 | -0.042 |
| 4560 | 0.012 | -0.018 |
| 4561 | 0.037 | 0.004 |
| 4562 | 0.033 | -0.048 |
| 4563 | 0.001 | 0.051 |
| 4564 | -0.004 | -0.030 |
| 4565 | 0.037 | 0.015 |
| 4566 | -0.012 | -0.104 |
| 4567 | -0.052 | -0.042 |
| 4568 | -0.023 | 0.102 |
| 4569 | -0.005 | -0.107 |
| 4570 | 0.018 | -0.115 |
| 4571 | 0.010 | 0.134 |
| 4572 | -0.053 | -0.176 |
| 4573 | -0.024 | -0.030 |
| 4574 | -0.045 | 0.062 |
| 4575 | 0.051 | -0.051 |
| 4576 | -0.014 | -0.102 |
| 4577 | 0.038 | -0.061 |
| 4578 | 0.095 | -0.085 |
| 4579 | -0.078 | -0.026 |
| 4580 | -0.025 | 0.131 |
| 4581 | 0.073 | -0.116 |
| 4582 | -0.019 | -0.005 |
| 4583 | -0.059 | -0.088 |

| | | |
|---|---|---|
| 4584 | -0.063 | 0.049 |
| 4585 | -0.028 | -0.050 |
| 4586 | -0.027 | 0.038 |
| 4587 | 0.029 | 0.020 |
| 4588 | 0.010 | 0.104 |
| 4589 | -0.080 | 0.103 |
| 4590 | -0.006 | 0.243 |
| 4591 | -0.003 | 0.089 |
| 4592 | -0.019 | -0.077 |
| 4593 | 0.011 | 0.090 |
| 4594 | 0.016 | -0.012 |
| 4595 | 0.054 | 0.012 |
| 4596 | -0.027 | -0.040 |
| 4597 | -0.019 | -0.061 |
| 4598 | -0.010 | -0.019 |
| 4599 | 0.015 | 0.087 |
| 4600 | -0.068 | 0.026 |
| 4601 | 0.035 | -0.052 |
| 4602 | 0.036 | 0.013 |
| 4603 | 0.020 | 0.024 |
| 4604 | 0.022 | -0.017 |
| 4605 | -0.032 | -0.097 |
| 4606 | -0.001 | -0.052 |
| 4607 | 0.000 | 0.025 |
| 4608 | -0.043 | 0.071 |
| 4609 | 0.019 | 0.097 |
| 4610 | 0.038 | -0.040 |
| 4611 | -0.085 | 0.014 |
| 4612 | -0.011 | -0.007 |
| 4613 | 0.018 | -0.036 |
| 4614 | 0.005 | -0.065 |
| 4615 | 0.051 | 0.051 |
| 4616 | -0.050 | 0.060 |
| 4617 | 0.024 | -0.006 |
| 4618 | 0.008 | -0.078 |
| 4619 | 0.004 | 0.032 |
| 4620 | -0.070 | -0.065 |
| 4621 | -0.015 | -0.034 |
| 4622 | 0.088 | 0.005 |
| 4623 | 0.037 | -0.063 |
| 4624 | 0.002 | 0.262 |
| 4625 | 0.030 | 0.062 |
| 4626 | -0.006 | 0.036 |
| 4627 | -0.002 | 0.103 |
| 4628 | 0.003 | 0.041 |
| 4629 | -0.017 | 0.044 |
| 4630 | -0.024 | 0.037 |
| 4631 | -0.038 | -0.036 |
| 4632 | -0.016 | 0.060 |
| 4633 | 0.004 | 0.068 |
| 4634 | -0.010 | -0.087 |

| | | |
|---|---|---|
| 4635 | -0.004 | 0.033 |
| 4636 | -0.007 | -0.032 |
| 4637 | -0.029 | 0.004 |
| 4638 | -0.007 | 0.021 |
| 4639 | 0.041 | 0.088 |
| 4640 | -0.021 | 0.019 |
| 4641 | 0.016 | -0.011 |
| 4642 | 0.007 | -0.078 |
| 4643 | -0.054 | -0.020 |
| 4644 | 0.006 | -0.027 |
| 4645 | 0.018 | 0.051 |
| 4646 | -0.031 | -0.056 |
| 4647 | -0.021 | -0.003 |
| 4648 | -0.019 | 0.007 |
| 4649 | 0.016 | -0.034 |
| 4650 | -0.023 | 0.150 |
| 4651 | 0.080 | 0.111 |
| 4652 | -0.013 | 0.017 |
| 4653 | -0.053 | 0.070 |
| 4654 | 0.033 | -0.035 |
| 4655 | 0.037 | 0.024 |
| 4656 | 0.014 | -0.039 |
| 4657 | 0.040 | -0.021 |
| 4658 | -0.028 | 0.097 |
| 4659 | -0.052 | -0.006 |
| 4660 | 0.043 | -0.038 |
| 4661 | -0.017 | 0.038 |
| 4662 | -0.068 | -0.002 |
| 4663 | -0.006 | -0.060 |
| 4664 | 0.005 | -0.111 |
| 4665 | 0.003 | -0.016 |
| 4666 | 0.009 | 0.000 |
| 4667 | 0.019 | 0.017 |
| 4668 | -0.001 | 0.051 |
| 4669 | -0.044 | -0.022 |
| 4670 | -0.035 | 0.054 |
| 4671 | 0.008 | 0.038 |
| 4672 | -0.054 | 0.119 |
| 4673 | -0.028 | 0.019 |
| 4674 | -0.043 | -0.007 |
| 4675 | 0.007 | 0.058 |
| 4676 | 0.042 | -0.075 |
| 4677 | -0.003 | -0.084 |
| 4678 | 0.045 | 0.129 |
| 4679 | -0.033 | 0.014 |
| 4680 | 0.015 | -0.075 |
| 4681 | 0.011 | 0.091 |
| 4682 | -0.058 | -0.076 |
| 4683 | 0.018 | -0.132 |
| 4684 | 0.005 | 0.009 |
| 4685 | -0.039 | 0.069 |

| | | |
|---|---|---|
| 4686 | -0.039 | -0.054 |
| 4687 | -0.019 | -0.008 |
| 4688 | -0.025 | 0.068 |
| 4689 | -0.075 | -0.019 |
| 4690 | -0.038 | -0.027 |
| 4691 | 0.011 | -0.057 |
| 4692 | 0.006 | -0.033 |
| 4693 | -0.064 | -0.038 |
| 4694 | -0.021 | 0.033 |
| 4695 | 0.001 | -0.040 |
| 4696 | 0.037 | 0.074 |
| 4697 | -0.020 | 0.074 |
| 4698 | 0.044 | 0.076 |
| 4699 | -0.002 | 0.091 |
| 4700 | 0.026 | -0.036 |
| 4701 | -0.058 | -0.022 |
| 4702 | 0.046 | -0.034 |
| 4703 | -0.016 | -0.085 |
| 4704 | 0.026 | -0.027 |
| 4705 | -0.079 | 0.012 |
| 4706 | 0.059 | 0.043 |
| 4707 | -0.011 | -0.012 |
| 4708 | -0.036 | 0.022 |
| 4709 | 0.001 | 0.079 |
| 4710 | 0.000 | -0.020 |
| 4711 | 0.037 | -0.068 |
| 4712 | -0.002 | -0.040 |
| 4713 | -0.047 | -0.082 |
| 4714 | -0.013 | -0.010 |
| 4715 | -0.029 | -0.012 |
| 4716 | 0.009 | -0.163 |
| 4717 | -0.047 | -0.061 |
| 4718 | -0.007 | -0.043 |
| 4719 | -0.016 | 0.069 |
| 4720 | 0.054 | -0.045 |
| 4721 | -0.026 | -0.067 |
| 4722 | -0.011 | 0.076 |
| 4723 | -0.054 | 0.065 |
| 4724 | 0.018 | -0.077 |
| 4725 | -0.031 | 0.007 |
| 4726 | 0.019 | -0.057 |
| 4727 | -0.034 | 0.040 |
| 4728 | 0.004 | -0.083 |
| 4729 | 0.024 | 0.034 |
| 4730 | -0.036 | 0.000 |
| 4731 | 0.010 | -0.050 |
| 4732 | 0.018 | 0.108 |
| 4733 | -0.018 | 0.058 |
| 4734 | -0.010 | 0.067 |
| 4735 | 0.001 | -0.074 |
| 4736 | 0.019 | 0.042 |

| | | |
|---|---|---|
| 4737 | -0.052 | 0.058 |
| 4738 | 0.008 | 0.030 |
| 4739 | -0.009 | -0.001 |
| 4740 | 0.053 | 0.002 |
| 4741 | 0.021 | 0.039 |
| 4742 | 0.024 | 0.028 |
| 4743 | -0.051 | -0.021 |
| 4744 | 0.035 | -0.046 |
| 4745 | -0.029 | -0.085 |
| 4746 | -0.018 | 0.025 |
| 4747 | -0.020 | 0.063 |
| 4748 | 0.035 | -0.091 |
| 4749 | -0.036 | 0.006 |
| 4750 | 0.088 | 0.050 |
| 4751 | 0.044 | -0.039 |
| 4752 | 0.017 | -0.065 |
| 4753 | 0.020 | 0.210 |
| 4754 | -0.024 | -0.050 |
| 4755 | -0.021 | 0.000 |
| 4756 | 0.063 | 0.108 |
| 4757 | 0.062 | -0.072 |
| 4758 | -0.005 | -0.023 |
| 4759 | 0.069 | -0.012 |
| 4760 | -0.045 | 0.016 |
| 4761 | -0.017 | -0.064 |
| 4762 | 0.023 | 0.068 |
| 4763 | 0.000 | 0.013 |
| 4764 | -0.009 | -0.013 |
| 4765 | -0.053 | 0.077 |
| 4766 | 0.001 | 0.053 |
| 4767 | -0.033 | -0.026 |
| 4768 | 0.004 | 0.050 |
| 4769 | 0.011 | 0.045 |
| 4770 | -0.009 | -0.036 |
| 4771 | 0.040 | 0.047 |
| 4772 | 0.027 | -0.037 |
| 4773 | 0.018 | -0.058 |
| 4774 | -0.015 | 0.058 |
| 4775 | 0.003 | 0.155 |
| 4776 | -0.010 | 0.064 |
| 4777 | 0.000 | 0.010 |
| 4778 | -0.023 | 0.030 |
| 4779 | -0.018 | -0.048 |
| 4780 | -0.006 | 0.004 |
| 4781 | 0.060 | -0.032 |
| 4782 | 0.016 | -0.074 |
| 4783 | -0.001 | 0.024 |
| 4784 | 0.110 | -0.033 |
| 4785 | 0.028 | 0.015 |
| 4786 | -0.007 | 0.007 |
| 4787 | -0.027 | -0.028 |

| | | |
|---|---|---|
| 4788 | -0.002 | 0.114 |
| 4789 | -0.001 | 0.055 |
| 4790 | 0.067 | 0.053 |
| 4791 | -0.036 | 0.036 |
| 4792 | -0.043 | -0.075 |
| 4793 | -0.033 | -0.012 |
| 4794 | 0.087 | -0.024 |
| 4795 | 0.016 | 0.001 |
| 4796 | 0.010 | -0.055 |
| 4797 | 0.043 | 0.191 |
| 4798 | 0.039 | 0.019 |
| 4799 | 0.021 | -0.135 |
| 4800 | -0.028 | -0.069 |
| 4801 | -0.015 | 0.149 |
| 4802 | 0.045 | 0.031 |
| 4803 | 0.045 | 0.095 |
| 4804 | -0.009 | -0.146 |
| 4805 | -0.063 | 0.007 |
| 4806 | 0.028 | 0.060 |
| 4807 | 0.021 | 0.063 |
| 4808 | -0.039 | 0.104 |
| 4809 | 0.018 | -0.077 |
| 4810 | -0.033 | -0.047 |
| 4811 | 0.030 | -0.003 |
| 4812 | 0.017 | -0.003 |
| 4813 | -0.027 | 0.087 |
| 4814 | 0.037 | 0.003 |
| 4815 | 0.006 | 0.095 |
| 4816 | -0.013 | 0.099 |
| 4817 | -0.004 | 0.059 |
| 4818 | -0.007 | 0.008 |
| 4819 | 0.041 | -0.016 |
| 4820 | -0.123 | 0.133 |
| 4821 | -0.039 | 0.081 |
| 4822 | -0.030 | 0.109 |
| 4823 | 0.008 | -0.133 |
| 4824 | 0.029 | 0.118 |
| 4825 | 0.002 | 0.004 |
| 4826 | 0.032 | -0.065 |
| 4827 | -0.009 | 0.141 |
| 4828 | 0.012 | -0.037 |
| 4829 | 0.009 | 0.029 |
| 4830 | 0.007 | -0.001 |
| 4831 | -0.002 | -0.035 |
| 4832 | -0.011 | 0.069 |
| 4833 | 0.015 | 0.067 |
| 4834 | 0.015 | 0.090 |
| 4835 | 0.001 | 0.029 |
| 4836 | -0.019 | -0.010 |
| 4837 | 0.031 | 0.101 |
| 4838 | 0.029 | 0.107 |

| | | |
|---|---|---|
| 4839 | 0.035 | -0.047 |
| 4840 | -0.008 | -0.050 |
| 4841 | -0.031 | -0.087 |
| 4842 | -0.044 | -0.133 |
| 4843 | -0.025 | -0.015 |
| 4844 | 0.057 | 0.079 |
| 4845 | -0.009 | 0.035 |
| 4846 | -0.026 | 0.053 |
| 4847 | 0.000 | 0.153 |
| 4848 | -0.004 | 0.085 |
| 4849 | -0.012 | -0.080 |
| 4850 | 0.005 | 0.132 |
| 4851 | 0.025 | 0.133 |
| 4852 | 0.002 | -0.010 |
| 4853 | -0.007 | -0.055 |
| 4854 | -0.001 | 0.108 |
| 4855 | 0.008 | 0.030 |
| 4856 | 0.028 | -0.042 |
| 4857 | 0.017 | -0.032 |
| 4858 | 0.069 | -0.010 |
| 4859 | -0.029 | -0.007 |
| 4860 | -0.003 | -0.166 |
| 4861 | -0.060 | -0.016 |
| 4862 | 0.007 | 0.078 |
| 4863 | 0.024 | -0.015 |
| 4864 | 0.006 | -0.118 |
| 4865 | -0.033 | -0.069 |
| 4866 | 0.001 | 0.111 |
| 4867 | -0.003 | 0.004 |
| 4868 | 0.011 | 0.052 |
| 4869 | -0.015 | 0.028 |
| 4870 | -0.041 | -0.028 |
| 4871 | 0.023 | -0.015 |
| 4872 | -0.082 | -0.020 |
| 4873 | -0.031 | -0.027 |
| 4874 | -0.036 | 0.000 |
| 4875 | -0.036 | 0.029 |
| 4876 | 0.011 | -0.108 |
| 4877 | -0.071 | -0.035 |
| 4878 | -0.037 | -0.040 |
| 4879 | -0.045 | 0.070 |
| 4880 | -0.011 | 0.003 |
| 4881 | 0.038 | -0.052 |
| 4882 | -0.049 | 0.036 |
| 4883 | -0.002 | 0.038 |
| 4884 | -0.001 | -0.138 |
| 4885 | -0.016 | 0.015 |
| 4886 | 0.016 | -0.014 |
| 4887 | 0.006 | 0.034 |
| 4888 | 0.023 | 0.007 |
| 4889 | 0.035 | 0.000 |

| | | |
|---|---|---|
| 4890 | 0.011 | -0.158 |
| 4891 | 0.053 | 0.051 |
| 4892 | -0.002 | -0.015 |
| 4893 | 0.008 | -0.104 |
| 4894 | 0.018 | -0.047 |
| 4895 | -0.023 | -0.047 |
| 4896 | 0.015 | 0.014 |
| 4897 | 0.067 | 0.244 |
| 4898 | -0.041 | -0.071 |
| 4899 | 0.037 | -0.082 |
| 4900 | 0.005 | -0.062 |
| 4901 | 0.012 | -0.144 |
| 4902 | -0.007 | 0.155 |
| 4903 | 0.060 | 0.078 |
| 4904 | -0.025 | 0.024 |
| 4905 | 0.016 | 0.115 |
| 4906 | 0.001 | -0.020 |
| 4907 | -0.065 | -0.023 |
| 4908 | -0.021 | 0.000 |
| 4909 | -0.044 | -0.003 |
| 4910 | 0.030 | 0.101 |
| 4911 | -0.026 | 0.052 |
| 4912 | 0.022 | 0.009 |
| 4913 | -0.033 | -0.054 |
| 4914 | -0.002 | -0.012 |
| 4915 | 0.033 | 0.038 |
| 4916 | -0.055 | 0.045 |
| 4917 | 0.044 | -0.003 |
| 4918 | 0.033 | -0.020 |
| 4919 | -0.032 | -0.089 |
| 4920 | 0.010 | -0.046 |
| 4921 | 0.015 | -0.063 |
| 4922 | 0.053 | 0.012 |
| 4923 | -0.031 | 0.042 |
| 4924 | -0.018 | -0.020 |
| 4925 | -0.039 | 0.066 |
| 4926 | -0.021 | 0.111 |
| 4927 | -0.010 | -0.118 |
| 4928 | 0.032 | -0.013 |
| 4929 | 0.078 | -0.112 |
| 4930 | -0.021 | -0.035 |
| 4931 | -0.056 | 0.090 |
| 4932 | -0.037 | -0.039 |
| 4933 | 0.031 | -0.024 |
| 4934 | -0.017 | -0.033 |
| 4935 | -0.027 | -0.055 |
| 4936 | 0.050 | -0.039 |
| 4937 | 0.057 | -0.036 |
| 4938 | 0.038 | -0.005 |
| 4939 | 0.001 | -0.135 |
| 4940 | -0.026 | -0.145 |

| | | |
|---|---|---|
| 4941 | 0.027 | 0.087 |
| 4942 | -0.001 | -0.032 |
| 4943 | -0.029 | -0.019 |
| 4944 | -0.010 | -0.009 |
| 4945 | -0.038 | 0.036 |
| 4946 | 0.023 | -0.170 |
| 4947 | -0.003 | 0.001 |
| 4948 | 0.034 | -0.010 |
| 4949 | 0.064 | 0.158 |
| 4950 | 0.021 | 0.185 |
| 4951 | -0.052 | 0.077 |
| 4952 | -0.011 | -0.106 |
| 4953 | 0.018 | -0.011 |
| 4954 | -0.043 | -0.117 |
| 4955 | 0.050 | 0.102 |
| 4956 | -0.035 | 0.013 |
| 4957 | -0.012 | 0.072 |
| 4958 | -0.038 | -0.011 |
| 4959 | 0.007 | 0.073 |
| 4960 | 0.003 | -0.037 |
| 4961 | -0.044 | -0.074 |
| 4962 | -0.030 | 0.030 |
| 4963 | -0.007 | -0.038 |
| 4964 | -0.023 | -0.122 |
| 4965 | -0.044 | -0.058 |
| 4966 | -0.046 | 0.060 |
| 4967 | 0.008 | -0.080 |
| 4968 | 0.032 | -0.099 |
| 4969 | 0.030 | 0.200 |
| 4970 | -0.042 | 0.085 |
| 4971 | 0.011 | 0.059 |
| 4972 | -0.042 | 0.049 |
| 4973 | -0.035 | -0.119 |
| 4974 | 0.003 | 0.122 |
| 4975 | 0.025 | 0.019 |
| 4976 | -0.010 | -0.023 |
| 4977 | 0.039 | 0.064 |
| 4978 | -0.007 | 0.026 |
| 4979 | 0.008 | -0.027 |
| 4980 | -0.018 | -0.059 |
| 4981 | 0.043 | -0.018 |
| 4982 | -0.018 | 0.064 |
| 4983 | 0.043 | -0.060 |
| 4984 | 0.006 | 0.049 |
| 4985 | -0.008 | -0.104 |
| 4986 | 0.042 | 0.082 |
| 4987 | 0.038 | -0.035 |
| 4988 | -0.011 | 0.089 |
| 4989 | 0.007 | -0.043 |
| 4990 | -0.066 | -0.076 |
| 4991 | -0.029 | -0.162 |

| | | |
|---|---|---|
| 4992 | -0.035 | -0.012 |
| 4993 | 0.027 | -0.190 |
| 4994 | 0.057 | 0.012 |
| 4995 | -0.050 | -0.013 |
| 4996 | 0.049 | -0.056 |
| 4997 | 0.049 | 0.016 |
| 4998 | -0.023 | -0.048 |
| 4999 | -0.011 | -0.100 |
| 5000 | 0.001 | -0.037 |
| 5001 | -0.020 | 0.007 |
| 5002 | 0.026 | 0.036 |
| 5003 | 0.011 | 0.002 |
| 5004 | -0.019 | -0.032 |
| 5005 | -0.002 | -0.014 |
| 5006 | 0.007 | -0.102 |
| 5007 | -0.070 | -0.053 |
| 5008 | -0.056 | 0.058 |
| 5009 | -0.049 | 0.022 |
| 5010 | -0.016 | -0.007 |
| 5011 | 0.014 | -0.057 |
| 5012 | -0.033 | 0.014 |
| 5013 | 0.001 | 0.113 |
| 5014 | 0.068 | -0.026 |
| 5015 | -0.001 | -0.012 |
| 5016 | -0.026 | 0.007 |
| 5017 | -0.010 | -0.048 |
| 5018 | -0.077 | -0.034 |
| 5019 | 0.027 | -0.026 |
| 5020 | -0.026 | 0.089 |
| 5021 | -0.006 | -0.018 |
| 5022 | -0.031 | -0.040 |
| 5023 | 0.037 | 0.043 |
| 5024 | 0.005 | 0.100 |
| 5025 | 0.008 | 0.007 |
| 5026 | -0.017 | 0.000 |
| 5027 | 0.000 | 0.009 |
| 5028 | 0.071 | -0.060 |
| 5029 | -0.056 | -0.036 |
| 5030 | -0.003 | 0.066 |
| 5031 | 0.003 | -0.021 |
| 5032 | 0.065 | -0.035 |
| 5033 | 0.028 | -0.066 |
| 5034 | 0.009 | -0.068 |
| 5035 | 0.010 | -0.046 |
| 5036 | -0.036 | -0.107 |
| 5037 | -0.005 | 0.069 |
| 5038 | -0.006 | 0.018 |
| 5039 | 0.031 | -0.006 |
| 5040 | 0.018 | -0.061 |
| 5041 | 0.032 | 0.071 |
| 5042 | 0.019 | -0.108 |

| | | |
|---|---|---|
| 5043 | 0.041 | -0.099 |
| 5044 | -0.018 | 0.073 |
| 5045 | -0.030 | -0.026 |
| 5046 | 0.020 | -0.097 |
| 5047 | -0.024 | -0.034 |
| 5048 | -0.049 | -0.096 |
| 5049 | 0.017 | -0.118 |
| 5050 | -0.008 | 0.026 |
| 5051 | 0.012 | 0.005 |
| 5052 | -0.006 | -0.018 |
| 5053 | 0.063 | 0.021 |
| 5054 | -0.021 | -0.162 |
| 5055 | -0.013 | 0.034 |
| 5056 | -0.014 | -0.154 |
| 5057 | 0.019 | -0.129 |
| 5058 | 0.000 | -0.054 |
| 5059 | 0.001 | -0.028 |
| 5060 | -0.008 | -0.026 |
| 5061 | -0.020 | -0.007 |
| 5062 | -0.034 | 0.046 |
| 5063 | -0.006 | 0.010 |
| 5064 | 0.005 | 0.041 |
| 5065 | 0.045 | 0.002 |
| 5066 | -0.031 | -0.018 |
| 5067 | 0.008 | 0.109 |
| 5068 | 0.114 | 0.082 |
| 5069 | -0.011 | -0.042 |
| 5070 | -0.038 | 0.003 |
| 5071 | 0.029 | -0.059 |
| 5072 | -0.024 | -0.026 |
| 5073 | 0.007 | 0.031 |
| 5074 | 0.005 | 0.003 |
| 5075 | -0.018 | -0.044 |
| 5076 | -0.038 | 0.171 |
| 5077 | 0.003 | -0.117 |
| 5078 | -0.006 | -0.026 |
| 5079 | -0.002 | -0.009 |
| 5080 | -0.012 | 0.044 |
| 5081 | -0.028 | -0.044 |
| 5082 | -0.015 | -0.049 |
| 5083 | -0.035 | 0.024 |
| 5084 | -0.001 | -0.115 |
| 5085 | -0.055 | 0.032 |
| 5086 | -0.025 | -0.039 |
| 5087 | 0.066 | -0.037 |
| 5088 | -0.029 | -0.008 |
| 5089 | 0.022 | -0.101 |
| 5090 | -0.095 | 0.107 |
| 5091 | -0.020 | -0.042 |
| 5092 | -0.029 | -0.037 |
| 5093 | 0.063 | -0.121 |

| | | |
|---|---|---|
| 5094 | 0.033 | -0.001 |
| 5095 | 0.029 | 0.002 |
| 5096 | 0.077 | -0.051 |
| 5097 | 0.025 | -0.114 |
| 5098 | -0.011 | -0.091 |
| 5099 | 0.055 | 0.015 |
| 5100 | -0.013 | 0.060 |
| 5101 | 0.014 | 0.048 |
| 5102 | 0.056 | -0.063 |
| 5103 | 0.011 | -0.038 |
| 5104 | 0.053 | -0.007 |
| 5105 | -0.042 | 0.022 |
| 5106 | 0.033 | -0.138 |
| 5107 | 0.015 | -0.072 |
| 5108 | -0.010 | -0.176 |
| 5109 | -0.021 | 0.073 |
| 5110 | 0.017 | 0.002 |
| 5111 | 0.003 | -0.056 |
| 5112 | 0.000 | 0.038 |
| 5113 | 0.027 | -0.031 |
| 5114 | 0.040 | -0.059 |
| 5115 | 0.039 | 0.024 |
| 5116 | 0.011 | 0.058 |
| 5117 | -0.021 | -0.095 |
| 5118 | 0.036 | -0.064 |
| 5119 | 0.058 | -0.115 |
| 5120 | -0.047 | -0.085 |
| 5121 | 0.027 | -0.003 |
| 5122 | 0.003 | -0.080 |
| 5123 | 0.011 | -0.107 |
| 5124 | 0.016 | 0.124 |
| 5125 | 0.035 | 0.020 |
| 5126 | 0.036 | -0.100 |
| 5127 | 0.007 | 0.019 |
| 5128 | -0.021 | 0.011 |
| 5129 | 0.010 | -0.103 |
| 5130 | 0.002 | -0.085 |
| 5131 | -0.028 | -0.086 |
| 5132 | -0.002 | 0.061 |
| 5133 | -0.059 | -0.085 |
| 5134 | 0.001 | -0.073 |
| 5135 | 0.025 | -0.076 |
| 5136 | -0.014 | -0.091 |
| 5137 | 0.022 | -0.198 |
| 5138 | 0.002 | 0.132 |
| 5139 | -0.028 | -0.142 |
| 5140 | 0.013 | -0.137 |
| 5141 | -0.048 | -0.036 |
| 5142 | -0.024 | -0.079 |
| 5143 | -0.028 | 0.028 |
| 5144 | 0.019 | 0.016 |

| | | |
|---|---|---|
| 5145 | 0.005 | 0.158 |
| 5146 | 0.045 | 0.035 |
| 5147 | 0.021 | 0.042 |
| 5148 | 0.003 | -0.014 |
| 5149 | -0.006 | -0.010 |
| 5150 | 0.018 | 0.117 |
| 5151 | 0.018 | 0.021 |
| 5152 | 0.036 | -0.020 |
| 5153 | 0.016 | 0.003 |
| 5154 | -0.003 | -0.115 |
| 5155 | 0.006 | -0.032 |
| 5156 | 0.006 | 0.024 |
| 5157 | 0.024 | 0.127 |
| 5158 | 0.023 | -0.021 |
| 5159 | 0.010 | 0.093 |
| 5160 | -0.001 | 0.059 |
| 5161 | 0.016 | 0.013 |
| 5162 | -0.006 | -0.042 |
| 5163 | -0.022 | 0.014 |
| 5164 | -0.004 | -0.065 |
| 5165 | -0.002 | 0.091 |
| 5166 | 0.010 | -0.055 |
| 5167 | 0.026 | 0.057 |
| 5168 | -0.037 | 0.051 |
| 5169 | 0.005 | -0.108 |
| 5170 | 0.030 | 0.037 |
| 5171 | -0.034 | -0.025 |
| 5172 | 0.008 | -0.196 |
| 5173 | -0.003 | -0.034 |
| 5174 | -0.031 | 0.067 |
| 5175 | 0.090 | -0.018 |
| 5176 | 0.061 | 0.020 |
| 5177 | -0.020 | 0.060 |
| 5178 | -0.039 | 0.010 |
| 5179 | 0.052 | -0.153 |
| 5180 | -0.019 | -0.081 |
| 5181 | 0.029 | -0.004 |
| 5182 | -0.049 | -0.052 |
| 5183 | -0.064 | -0.013 |
| 5184 | 0.009 | 0.115 |
| 5185 | -0.020 | 0.198 |
| 5186 | 0.051 | -0.006 |
| 5187 | -0.001 | -0.079 |
| 5188 | -0.028 | -0.014 |
| 5189 | 0.020 | -0.090 |
| 5190 | -0.009 | -0.010 |
| 5191 | -0.016 | 0.031 |
| 5192 | 0.020 | 0.003 |
| 5193 | 0.043 | -0.035 |
| 5194 | -0.014 | 0.144 |
| 5195 | -0.026 | -0.080 |

| | | |
|---|---|---|
| 5196 | -0.036 | -0.062 |
| 5197 | -0.019 | 0.035 |
| 5198 | -0.013 | -0.068 |
| 5199 | 0.036 | 0.009 |
| 5200 | -0.089 | 0.102 |
| 5201 | 0.057 | -0.004 |
| 5202 | -0.015 | -0.039 |
| 5203 | 0.038 | -0.160 |
| 5204 | 0.014 | 0.088 |
| 5205 | -0.024 | 0.066 |
| 5206 | -0.054 | -0.029 |
| 5207 | 0.016 | -0.011 |
| 5208 | 0.027 | -0.006 |
| 5209 | 0.007 | -0.012 |
| 5210 | 0.009 | -0.004 |
| 5211 | 0.002 | 0.001 |
| 5212 | 0.009 | -0.061 |
| 5213 | -0.018 | 0.038 |
| 5214 | 0.010 | 0.050 |
| 5215 | -0.007 | -0.170 |
| 5216 | -0.009 | -0.027 |
| 5217 | -0.009 | 0.049 |
| 5218 | -0.004 | -0.012 |
| 5219 | -0.005 | 0.041 |
| 5220 | -0.044 | 0.083 |
| 5221 | 0.051 | 0.017 |
| 5222 | 0.009 | -0.060 |
| 5223 | -0.098 | -0.029 |
| 5224 | 0.031 | -0.003 |
| 5225 | 0.044 | -0.001 |
| 5226 | 0.067 | -0.037 |
| 5227 | -0.051 | 0.112 |
| 5228 | -0.063 | 0.053 |
| 5229 | -0.062 | -0.106 |
| 5230 | 0.000 | 0.140 |
| 5231 | 0.024 | -0.013 |
| 5232 | 0.003 | -0.051 |
| 5233 | 0.011 | 0.018 |
| 5234 | -0.011 | 0.043 |
| 5235 | 0.008 | -0.117 |
| 5236 | 0.009 | -0.059 |
| 5237 | 0.001 | 0.067 |
| 5238 | -0.041 | -0.136 |
| 5239 | -0.001 | -0.053 |
| 5240 | -0.028 | 0.126 |
| 5241 | -0.009 | 0.060 |
| 5242 | -0.003 | 0.015 |
| 5243 | 0.040 | 0.121 |
| 5244 | -0.014 | 0.043 |
| 5245 | -0.031 | 0.007 |
| 5246 | 0.013 | 0.046 |

| | | |
|---|---|---|
| 5247 | 0.055 | -0.063 |
| 5248 | 0.041 | -0.148 |
| 5249 | -0.022 | 0.048 |
| 5250 | 0.017 | 0.065 |
| 5251 | -0.060 | 0.026 |
| 5252 | -0.045 | -0.126 |
| 5253 | -0.028 | -0.091 |
| 5254 | -0.064 | -0.046 |
| 5255 | 0.011 | -0.023 |
| 5256 | -0.003 | 0.148 |
| 5257 | 0.064 | 0.028 |
| 5258 | 0.027 | -0.065 |
| 5259 | -0.026 | 0.203 |
| 5260 | 0.023 | -0.053 |
| 5261 | 0.033 | -0.087 |
| 5262 | 0.007 | -0.073 |
| 5263 | 0.002 | 0.109 |
| 5264 | -0.012 | -0.059 |
| 5265 | -0.030 | -0.125 |
| 5266 | -0.018 | -0.013 |
| 5267 | 0.004 | -0.033 |
| 5268 | 0.061 | -0.019 |
| 5269 | -0.073 | 0.071 |
| 5270 | 0.021 | -0.147 |
| 5271 | -0.051 | 0.005 |
| 5272 | -0.001 | 0.022 |
| 5273 | -0.040 | 0.137 |
| 5274 | -0.042 | 0.043 |
| 5275 | -0.046 | 0.000 |
| 5276 | 0.020 | 0.008 |
| 5277 | 0.043 | -0.023 |
| 5278 | 0.025 | -0.031 |
| 5279 | 0.000 | 0.050 |
| 5280 | -0.026 | -0.204 |
| 5281 | 0.022 | 0.103 |
| 5282 | -0.019 | -0.052 |
| 5283 | -0.012 | -0.005 |
| 5284 | -0.043 | -0.082 |
| 5285 | -0.034 | 0.036 |
| 5286 | 0.010 | 0.078 |
| 5287 | 0.029 | -0.002 |
| 5288 | -0.060 | 0.029 |
| 5289 | -0.007 | -0.117 |
| 5290 | 0.001 | 0.076 |
| 5291 | -0.023 | 0.069 |
| 5292 | -0.008 | -0.028 |
| 5293 | -0.022 | -0.072 |
| 5294 | -0.045 | -0.072 |
| 5295 | 0.022 | 0.141 |
| 5296 | -0.035 | -0.172 |
| 5297 | 0.002 | 0.043 |

| | | |
|---|---|---|
| 5298 | -0.027 | -0.053 |
| 5299 | -0.005 | -0.054 |
| 5300 | 0.026 | 0.034 |
| 5301 | -0.031 | 0.013 |
| 5302 | -0.033 | -0.002 |
| 5303 | 0.015 | -0.012 |
| 5304 | 0.039 | 0.062 |
| 5305 | -0.040 | -0.069 |
| 5306 | 0.035 | 0.044 |
| 5307 | 0.010 | -0.026 |
| 5308 | 0.027 | 0.005 |
| 5309 | 0.002 | 0.009 |
| 5310 | -0.053 | -0.137 |
| 5311 | 0.079 | 0.026 |
| 5312 | -0.022 | -0.143 |
| 5313 | -0.029 | 0.014 |
| 5314 | 0.025 | -0.134 |
| 5315 | 0.043 | 0.105 |
| 5316 | -0.011 | 0.040 |
| 5317 | 0.003 | 0.033 |
| 5318 | -0.006 | -0.136 |
| 5319 | 0.031 | -0.065 |
| 5320 | -0.026 | -0.097 |
| 5321 | 0.004 | -0.011 |
| 5322 | -0.023 | 0.035 |
| 5323 | 0.000 | -0.050 |
| 5324 | -0.025 | 0.075 |
| 5325 | 0.044 | 0.070 |
| 5326 | -0.057 | -0.116 |
| 5327 | -0.020 | 0.053 |
| 5328 | -0.010 | 0.042 |
| 5329 | -0.003 | 0.019 |
| 5330 | -0.025 | -0.043 |
| 5331 | -0.005 | 0.028 |
| 5332 | 0.049 | -0.074 |
| 5333 | -0.051 | 0.000 |
| 5334 | -0.070 | 0.049 |
| 5335 | 0.056 | 0.012 |
| 5336 | -0.013 | -0.027 |
| 5337 | 0.039 | -0.011 |
| 5338 | -0.020 | -0.019 |
| 5339 | 0.003 | -0.038 |
| 5340 | -0.009 | 0.079 |
| 5341 | 0.004 | -0.065 |
| 5342 | -0.021 | -0.147 |
| 5343 | 0.022 | 0.084 |
| 5344 | 0.023 | 0.133 |
| 5345 | -0.021 | 0.013 |
| 5346 | 0.035 | -0.037 |
| 5347 | 0.022 | 0.050 |
| 5348 | 0.011 | 0.041 |

| | | |
|---|---|---|
| 5349 | -0.006 | -0.103 |
| 5350 | 0.003 | -0.008 |
| 5351 | -0.044 | -0.038 |
| 5352 | 0.025 | -0.045 |
| 5353 | -0.027 | 0.131 |
| 5354 | -0.033 | -0.040 |
| 5355 | 0.001 | -0.043 |
| 5356 | 0.038 | 0.007 |
| 5357 | 0.016 | 0.095 |
| 5358 | -0.046 | -0.015 |
| 5359 | -0.015 | -0.113 |
| 5360 | -0.031 | -0.041 |
| 5361 | -0.002 | 0.082 |
| 5362 | 0.008 | -0.073 |
| 5363 | -0.035 | -0.009 |
| 5364 | 0.031 | -0.133 |
| 5365 | 0.030 | -0.065 |
| 5366 | 0.048 | 0.077 |
| 5367 | -0.031 | -0.051 |
| 5368 | 0.008 | 0.090 |
| 5369 | -0.012 | -0.003 |
| 5370 | 0.062 | -0.172 |
| 5371 | -0.018 | -0.082 |
| 5372 | 0.038 | 0.093 |
| 5373 | 0.043 | 0.000 |
| 5374 | 0.052 | 0.045 |
| 5375 | 0.057 | -0.070 |
| 5376 | -0.008 | -0.025 |
| 5377 | -0.013 | -0.070 |
| 5378 | 0.037 | 0.074 |
| 5379 | -0.004 | 0.035 |
| 5380 | 0.024 | 0.113 |
| 5381 | -0.035 | -0.045 |
| 5382 | -0.055 | 0.092 |
| 5383 | 0.038 | -0.060 |
| 5384 | 0.039 | 0.034 |
| 5385 | 0.062 | 0.057 |
| 5386 | -0.091 | 0.048 |
| 5387 | 0.008 | -0.124 |
| 5388 | -0.026 | -0.022 |
| 5389 | 0.059 | 0.082 |
| 5390 | 0.046 | -0.163 |
| 5391 | -0.025 | -0.042 |
| 5392 | 0.046 | 0.122 |
| 5393 | -0.072 | 0.023 |
| 5394 | 0.000 | 0.023 |
| 5395 | 0.035 | -0.019 |
| 5396 | 0.018 | -0.057 |
| 5397 | -0.032 | 0.031 |
| 5398 | 0.028 | 0.049 |
| 5399 | 0.016 | -0.055 |

| | | |
|---|---|---|
| 5400 | -0.006 | -0.032 |
| 5401 | 0.060 | 0.040 |
| 5402 | 0.058 | -0.073 |
| 5403 | -0.019 | 0.048 |
| 5404 | -0.018 | -0.105 |
| 5405 | -0.014 | 0.070 |
| 5406 | -0.016 | 0.085 |
| 5407 | -0.002 | -0.030 |
| 5408 | -0.053 | 0.021 |
| 5409 | 0.052 | -0.235 |
| 5410 | 0.018 | -0.052 |
| 5411 | 0.051 | -0.038 |
| 5412 | 0.031 | -0.003 |
| 5413 | 0.031 | 0.061 |
| 5414 | 0.059 | -0.054 |
| 5415 | 0.026 | -0.001 |
| 5416 | -0.052 | 0.014 |
| 5417 | -0.044 | -0.052 |
| 5418 | -0.044 | -0.119 |
| 5419 | 0.025 | 0.037 |
| 5420 | -0.001 | 0.024 |
| 5421 | -0.019 | -0.036 |
| 5422 | -0.043 | -0.078 |
| 5423 | -0.078 | 0.048 |
| 5424 | 0.050 | -0.019 |
| 5425 | -0.008 | 0.066 |
| 5426 | 0.019 | -0.057 |
| 5427 | 0.049 | -0.011 |
| 5428 | 0.009 | -0.131 |
| 5429 | -0.004 | 0.095 |
| 5430 | -0.041 | 0.024 |
| 5431 | -0.026 | 0.101 |
| 5432 | -0.014 | 0.021 |
| 5433 | 0.008 | -0.004 |
| 5434 | -0.070 | 0.099 |
| 5435 | -0.004 | -0.017 |
| 5436 | -0.052 | 0.102 |
| 5437 | -0.052 | 0.063 |
| 5438 | -0.008 | 0.153 |
| 5439 | -0.014 | -0.017 |
| 5440 | 0.075 | -0.005 |
| 5441 | 0.066 | -0.018 |
| 5442 | -0.016 | -0.046 |
| 5443 | -0.013 | 0.057 |
| 5444 | -0.023 | -0.097 |
| 5445 | -0.008 | 0.021 |
| 5446 | -0.025 | -0.011 |
| 5447 | -0.054 | 0.068 |
| 5448 | 0.018 | -0.138 |
| 5449 | -0.007 | -0.006 |
| 5450 | -0.039 | 0.001 |

| | | |
|---|---|---|
| 5451 | 0.011 | -0.035 |
| 5452 | 0.012 | 0.000 |
| 5453 | 0.036 | 0.015 |
| 5454 | -0.025 | 0.039 |
| 5455 | -0.003 | 0.112 |
| 5456 | 0.017 | 0.015 |
| 5457 | -0.008 | -0.036 |
| 5458 | 0.000 | 0.066 |
| 5459 | -0.010 | 0.017 |
| 5460 | 0.024 | 0.021 |
| 5461 | -0.014 | 0.067 |
| 5462 | 0.058 | 0.043 |
| 5463 | 0.032 | 0.013 |
| 5464 | 0.047 | -0.052 |
| 5465 | 0.059 | -0.017 |
| 5466 | -0.009 | -0.138 |
| 5467 | -0.045 | -0.065 |
| 5468 | -0.022 | -0.109 |
| 5469 | 0.036 | 0.025 |
| 5470 | -0.080 | -0.068 |
| 5471 | -0.039 | -0.032 |
| 5472 | 0.035 | -0.189 |
| 5473 | -0.001 | 0.090 |
| 5474 | 0.058 | 0.000 |
| 5475 | 0.031 | -0.078 |
| 5476 | -0.003 | -0.098 |
| 5477 | -0.032 | 0.094 |
| 5478 | -0.028 | -0.069 |
| 5479 | -0.005 | -0.015 |
| 5480 | 0.065 | 0.051 |
| 5481 | -0.039 | -0.013 |
| 5482 | -0.006 | -0.050 |
| 5483 | 0.005 | -0.031 |
| 5484 | 0.032 | 0.039 |
| 5485 | -0.013 | -0.052 |
| 5486 | -0.035 | -0.006 |
| 5487 | -0.025 | 0.053 |
| 5488 | 0.018 | 0.000 |
| 5489 | -0.002 | -0.128 |
| 5490 | 0.014 | -0.086 |
| 5491 | 0.028 | -0.044 |
| 5492 | -0.029 | -0.006 |
| 5493 | 0.011 | 0.011 |
| 5494 | 0.059 | 0.099 |
| 5495 | -0.056 | -0.059 |
| 5496 | 0.006 | -0.015 |
| 5497 | 0.048 | 0.015 |
| 5498 | -0.097 | 0.024 |
| 5499 | -0.001 | -0.015 |
| 5500 | 0.043 | 0.050 |
| 5501 | 0.003 | -0.020 |

| | | |
|---|---|---|
| 5502 | 0.050 | -0.055 |
| 5503 | -0.012 | 0.129 |
| 5504 | 0.011 | -0.175 |
| 5505 | -0.053 | 0.084 |
| 5506 | 0.010 | 0.083 |
| 5507 | 0.001 | -0.091 |
| 5508 | 0.077 | 0.071 |
| 5509 | -0.031 | 0.023 |
| 5510 | -0.024 | -0.028 |
| 5511 | 0.017 | -0.063 |
| 5512 | 0.010 | 0.096 |
| 5513 | 0.004 | -0.087 |
| 5514 | -0.039 | 0.093 |
| 5515 | 0.011 | -0.082 |
| 5516 | 0.022 | 0.037 |
| 5517 | -0.060 | 0.026 |
| 5518 | -0.041 | 0.070 |
| 5519 | 0.025 | -0.031 |
| 5520 | 0.036 | 0.032 |
| 5521 | 0.086 | -0.161 |
| 5522 | -0.046 | 0.000 |
| 5523 | 0.082 | 0.001 |
| 5524 | -0.001 | -0.003 |
| 5525 | -0.007 | 0.017 |
| 5526 | 0.003 | 0.016 |
| 5527 | 0.081 | -0.086 |
| 5528 | -0.043 | -0.099 |
| 5529 | 0.027 | -0.097 |
| 5530 | -0.026 | -0.040 |
| 5531 | -0.035 | 0.009 |
| 5532 | -0.012 | 0.113 |
| 5533 | 0.008 | 0.003 |
| 5534 | -0.008 | -0.097 |
| 5535 | 0.020 | -0.012 |
| 5536 | 0.015 | -0.139 |
| 5537 | 0.036 | -0.077 |
| 5538 | 0.019 | -0.063 |
| 5539 | -0.001 | 0.058 |
| 5540 | -0.044 | -0.017 |
| 5541 | 0.015 | -0.088 |
| 5542 | 0.020 | -0.051 |
| 5543 | -0.038 | -0.094 |
| 5544 | 0.048 | -0.025 |
| 5545 | -0.016 | 0.129 |
| 5546 | 0.010 | -0.021 |
| 5547 | -0.002 | -0.058 |
| 5548 | -0.055 | -0.032 |
| 5549 | -0.006 | -0.097 |
| 5550 | -0.003 | -0.113 |
| 5551 | 0.002 | -0.012 |
| 5552 | -0.020 | -0.101 |

| | | |
|---|---|---|
| 5553 | 0.034 | 0.151 |
| 5554 | 0.037 | -0.008 |
| 5555 | 0.018 | 0.058 |
| 5556 | 0.009 | -0.022 |
| 5557 | 0.002 | 0.049 |
| 5558 | -0.057 | -0.027 |
| 5559 | 0.029 | -0.029 |
| 5560 | -0.027 | -0.094 |
| 5561 | -0.014 | 0.021 |
| 5562 | 0.024 | 0.076 |
| 5563 | -0.011 | 0.038 |
| 5564 | 0.002 | -0.003 |
| 5565 | 0.100 | -0.023 |
| 5566 | 0.030 | -0.038 |
| 5567 | 0.025 | -0.090 |
| 5568 | -0.014 | 0.066 |
| 5569 | 0.025 | 0.016 |
| 5570 | -0.012 | -0.072 |
| 5571 | 0.006 | -0.014 |
| 5572 | -0.056 | -0.051 |
| 5573 | -0.076 | 0.041 |
| 5574 | -0.085 | -0.027 |
| 5575 | 0.012 | 0.106 |
| 5576 | -0.038 | -0.097 |
| 5577 | -0.004 | 0.138 |
| 5578 | -0.054 | 0.033 |
| 5579 | 0.010 | 0.091 |
| 5580 | 0.056 | 0.044 |
| 5581 | 0.027 | 0.028 |
| 5582 | -0.020 | -0.066 |
| 5583 | 0.044 | 0.023 |
| 5584 | 0.047 | 0.021 |
| 5585 | 0.013 | 0.057 |
| 5586 | 0.019 | 0.026 |
| 5587 | -0.021 | 0.116 |
| 5588 | 0.013 | 0.028 |
| 5589 | 0.022 | 0.079 |
| 5590 | 0.033 | 0.056 |
| 5591 | -0.020 | 0.057 |
| 5592 | 0.037 | 0.018 |
| 5593 | 0.054 | -0.035 |
| 5594 | -0.023 | 0.016 |
| 5595 | 0.013 | -0.049 |
| 5596 | -0.008 | 0.063 |
| 5597 | 0.088 | 0.006 |
| 5598 | -0.022 | 0.065 |
| 5599 | -0.047 | 0.139 |
| 5600 | 0.053 | 0.079 |
| 5601 | -0.051 | -0.033 |
| 5602 | 0.024 | -0.026 |
| 5603 | 0.023 | -0.037 |

| | | |
|---|---|---|
| 5604 | 0.014 | -0.039 |
| 5605 | 0.022 | 0.016 |
| 5606 | 0.039 | -0.055 |
| 5607 | -0.025 | -0.089 |
| 5608 | -0.014 | 0.155 |
| 5609 | 0.021 | -0.024 |
| 5610 | 0.000 | 0.154 |
| 5611 | -0.037 | 0.022 |
| 5612 | 0.005 | 0.033 |
| 5613 | -0.084 | 0.045 |
| 5614 | 0.036 | 0.086 |
| 5615 | 0.021 | -0.027 |
| 5616 | 0.025 | -0.001 |
| 5617 | -0.001 | -0.177 |
| 5618 | 0.104 | -0.069 |
| 5619 | 0.014 | 0.097 |
| 5620 | 0.003 | 0.119 |
| 5621 | 0.034 | 0.053 |
| 5622 | 0.056 | 0.027 |
| 5623 | -0.033 | -0.044 |
| 5624 | 0.012 | -0.066 |
| 5625 | 0.016 | -0.021 |
| 5626 | 0.059 | 0.083 |
| 5627 | 0.003 | -0.026 |
| 5628 | 0.026 | 0.018 |
| 5629 | 0.063 | 0.181 |
| 5630 | -0.019 | 0.072 |
| 5631 | -0.005 | 0.085 |
| 5632 | 0.012 | 0.030 |
| 5633 | 0.026 | -0.014 |
| 5634 | -0.027 | 0.063 |
| 5635 | 0.048 | 0.087 |
| 5636 | -0.001 | -0.024 |
| 5637 | 0.002 | -0.023 |
| 5638 | 0.021 | -0.142 |
| 5639 | -0.033 | 0.083 |
| 5640 | 0.015 | -0.016 |
| 5641 | 0.044 | 0.027 |
| 5642 | -0.003 | 0.019 |
| 5643 | 0.001 | 0.070 |
| 5644 | -0.069 | -0.011 |
| 5645 | 0.007 | -0.065 |
| 5646 | 0.029 | 0.013 |
| 5647 | -0.003 | -0.099 |
| 5648 | 0.037 | -0.109 |
| 5649 | -0.067 | -0.090 |
| 5650 | -0.018 | -0.002 |
| 5651 | -0.022 | -0.015 |
| 5652 | -0.021 | -0.101 |
| 5653 | -0.021 | -0.060 |
| 5654 | 0.064 | 0.059 |

| | | |
|---|---|---|
| 5655 | -0.044 | 0.025 |
| 5656 | -0.017 | -0.079 |
| 5657 | -0.020 | 0.098 |
| 5658 | 0.022 | -0.008 |
| 5659 | 0.015 | -0.005 |
| 5660 | -0.008 | -0.012 |
| 5661 | 0.061 | 0.118 |
| 5662 | 0.025 | -0.056 |
| 5663 | -0.011 | -0.111 |
| 5664 | 0.030 | -0.121 |
| 5665 | 0.009 | -0.031 |
| 5666 | -0.003 | -0.163 |
| 5667 | -0.008 | -0.004 |
| 5668 | -0.014 | 0.155 |
| 5669 | 0.076 | -0.114 |
| 5670 | 0.068 | -0.041 |
| 5671 | 0.091 | -0.090 |
| 5672 | 0.002 | 0.157 |
| 5673 | 0.013 | -0.172 |
| 5674 | 0.034 | 0.009 |
| 5675 | 0.025 | -0.001 |
| 5676 | -0.018 | 0.041 |
| 5677 | 0.036 | 0.066 |
| 5678 | 0.030 | 0.052 |
| 5679 | 0.045 | 0.055 |
| 5680 | -0.063 | -0.081 |
| 5681 | 0.011 | -0.072 |
| 5682 | -0.037 | 0.025 |
| 5683 | -0.002 | -0.032 |
| 5684 | -0.018 | 0.052 |
| 5685 | -0.050 | -0.232 |
| 5686 | 0.008 | -0.032 |
| 5687 | 0.033 | -0.057 |
| 5688 | -0.015 | 0.120 |
| 5689 | -0.033 | -0.072 |
| 5690 | -0.002 | -0.050 |
| 5691 | 0.010 | -0.006 |
| 5692 | -0.010 | 0.029 |
| 5693 | 0.060 | 0.061 |
| 5694 | 0.019 | -0.031 |
| 5695 | -0.031 | 0.143 |
| 5696 | 0.026 | -0.060 |
| 5697 | -0.027 | 0.038 |
| 5698 | -0.018 | 0.071 |
| 5699 | 0.026 | -0.054 |
| 5700 | -0.025 | -0.028 |
| 5701 | -0.017 | -0.063 |
| 5702 | 0.001 | 0.065 |
| 5703 | -0.004 | -0.118 |
| 5704 | -0.089 | 0.030 |
| 5705 | -0.002 | -0.103 |

| | | |
|---|---|---|
| 5706 | -0.020 | -0.143 |
| 5707 | 0.004 | -0.038 |
| 5708 | 0.048 | 0.024 |
| 5709 | 0.001 | 0.255 |
| 5710 | 0.016 | -0.073 |
| 5711 | -0.002 | 0.069 |
| 5712 | 0.025 | -0.042 |
| 5713 | 0.026 | -0.124 |
| 5714 | -0.035 | -0.054 |
| 5715 | 0.010 | 0.070 |
| 5716 | -0.049 | 0.087 |
| 5717 | -0.010 | -0.069 |
| 5718 | 0.036 | -0.035 |
| 5719 | -0.076 | 0.029 |
| 5720 | 0.018 | -0.063 |
| 5721 | 0.029 | -0.108 |
| 5722 | 0.032 | 0.136 |
| 5723 | 0.055 | -0.021 |
| 5724 | -0.011 | 0.021 |
| 5725 | -0.034 | 0.112 |
| 5726 | 0.004 | -0.037 |
| 5727 | -0.054 | -0.003 |
| 5728 | 0.019 | 0.003 |
| 5729 | -0.049 | -0.048 |
| 5730 | 0.018 | 0.038 |
| 5731 | -0.038 | -0.016 |
| 5732 | 0.027 | 0.018 |
| 5733 | 0.007 | 0.028 |
| 5734 | -0.096 | -0.134 |
| 5735 | -0.021 | -0.087 |
| 5736 | -0.070 | -0.025 |
| 5737 | -0.023 | -0.066 |
| 5738 | -0.031 | 0.015 |
| 5739 | -0.008 | -0.058 |
| 5740 | -0.034 | 0.126 |
| 5741 | 0.021 | 0.073 |
| 5742 | 0.001 | -0.029 |
| 5743 | 0.041 | 0.057 |
| 5744 | 0.006 | -0.048 |
| 5745 | -0.032 | -0.114 |
| 5746 | 0.039 | -0.013 |
| 5747 | 0.006 | -0.083 |
| 5748 | 0.017 | -0.105 |
| 5749 | 0.040 | 0.063 |
| 5750 | -0.026 | -0.145 |
| 5751 | -0.049 | 0.025 |
| 5752 | -0.018 | 0.075 |
| 5753 | 0.016 | -0.008 |
| 5754 | 0.024 | 0.039 |
| 5755 | -0.017 | 0.003 |
| 5756 | 0.051 | -0.078 |

| | | |
|---|---|---|
| 5757 | 0.037 | -0.054 |
| 5758 | -0.072 | -0.034 |
| 5759 | 0.025 | -0.031 |
| 5760 | -0.072 | -0.063 |
| 5761 | -0.001 | -0.109 |
| 5762 | 0.103 | 0.018 |
| 5763 | -0.012 | 0.180 |
| 5764 | -0.016 | -0.026 |
| 5765 | -0.013 | 0.018 |
| 5766 | -0.029 | -0.033 |
| 5767 | 0.025 | -0.101 |
| 5768 | 0.012 | -0.066 |
| 5769 | 0.037 | 0.070 |
| 5770 | 0.068 | -0.027 |
| 5771 | 0.039 | 0.118 |
| 5772 | 0.005 | -0.024 |
| 5773 | -0.012 | -0.037 |
| 5774 | 0.032 | 0.031 |
| 5775 | 0.014 | 0.036 |
| 5776 | 0.033 | 0.023 |
| 5777 | -0.043 | -0.073 |
| 5778 | -0.083 | -0.038 |
| 5779 | 0.000 | -0.101 |
| 5780 | 0.029 | -0.048 |
| 5781 | 0.057 | 0.048 |
| 5782 | -0.010 | 0.027 |
| 5783 | -0.043 | -0.078 |
| 5784 | 0.035 | -0.002 |
| 5785 | -0.039 | 0.056 |
| 5786 | 0.020 | -0.109 |
| 5787 | -0.020 | -0.074 |
| 5788 | -0.001 | 0.018 |
| 5789 | -0.032 | 0.065 |
| 5790 | 0.019 | -0.052 |
| 5791 | 0.035 | 0.175 |
| 5792 | 0.010 | -0.154 |
| 5793 | 0.005 | -0.060 |
| 5794 | -0.015 | -0.055 |
| 5795 | 0.044 | 0.029 |
| 5796 | -0.029 | 0.090 |
| 5797 | -0.061 | -0.016 |
| 5798 | -0.012 | 0.056 |
| 5799 | 0.037 | 0.088 |
| 5800 | 0.040 | -0.043 |
| 5801 | -0.005 | 0.024 |
| 5802 | -0.031 | -0.008 |
| 5803 | 0.052 | -0.044 |
| 5804 | -0.024 | -0.031 |
| 5805 | 0.028 | 0.025 |
| 5806 | -0.022 | 0.047 |
| 5807 | 0.024 | -0.067 |

| | | |
|---|---|---|
| 5808 | 0.036 | 0.024 |
| 5809 | -0.032 | 0.102 |
| 5810 | -0.012 | 0.121 |
| 5811 | 0.024 | -0.058 |
| 5812 | -0.028 | -0.067 |
| 5813 | 0.001 | 0.071 |
| 5814 | -0.017 | -0.097 |
| 5815 | -0.016 | -0.016 |
| 5816 | -0.031 | 0.003 |
| 5817 | 0.017 | 0.093 |
| 5818 | 0.034 | 0.069 |
| 5819 | 0.014 | -0.038 |
| 5820 | 0.046 | -0.046 |
| 5821 | -0.035 | -0.086 |
| 5822 | -0.029 | 0.006 |
| 5823 | -0.032 | -0.047 |
| 5824 | 0.032 | 0.087 |
| 5825 | 0.050 | 0.071 |
| 5826 | -0.024 | 0.067 |
| 5827 | 0.017 | -0.018 |
| 5828 | -0.010 | -0.018 |
| 5829 | -0.006 | 0.005 |
| 5830 | -0.034 | -0.011 |
| 5831 | -0.030 | -0.003 |
| 5832 | -0.036 | -0.078 |
| 5833 | -0.013 | 0.072 |
| 5834 | -0.004 | -0.013 |
| 5835 | -0.051 | -0.095 |
| 5836 | 0.079 | 0.053 |
| 5837 | -0.015 | 0.043 |
| 5838 | 0.064 | 0.003 |
| 5839 | 0.058 | 0.069 |
| 5840 | -0.054 | 0.098 |
| 5841 | -0.011 | -0.116 |
| 5842 | 0.024 | 0.043 |
| 5843 | 0.053 | -0.041 |
| 5844 | 0.053 | -0.033 |
| 5845 | 0.013 | 0.053 |
| 5846 | 0.034 | 0.000 |
| 5847 | -0.024 | 0.025 |
| 5848 | -0.027 | 0.027 |
| 5849 | -0.019 | 0.023 |
| 5850 | 0.040 | 0.019 |
| 5851 | -0.029 | 0.022 |
| 5852 | -0.069 | -0.019 |
| 5853 | 0.085 | -0.021 |
| 5854 | 0.030 | -0.110 |
| 5855 | -0.037 | 0.030 |
| 5856 | 0.039 | 0.009 |
| 5857 | 0.055 | 0.032 |
| 5858 | -0.030 | 0.015 |

| | | |
|---|---|---|
| 5859 | 0.004 | -0.081 |
| 5860 | 0.021 | -0.049 |
| 5861 | -0.027 | 0.123 |
| 5862 | 0.022 | 0.010 |
| 5863 | 0.001 | -0.075 |
| 5864 | -0.011 | 0.049 |
| 5865 | -0.023 | 0.050 |
| 5866 | -0.039 | -0.048 |
| 5867 | 0.054 | 0.085 |
| 5868 | 0.001 | -0.043 |
| 5869 | -0.003 | -0.052 |
| 5870 | -0.019 | 0.071 |
| 5871 | -0.037 | 0.010 |
| 5872 | -0.010 | 0.141 |
| 5873 | -0.042 | 0.072 |
| 5874 | -0.001 | 0.105 |
| 5875 | 0.039 | -0.039 |
| 5876 | 0.059 | 0.011 |
| 5877 | 0.030 | -0.063 |
| 5878 | -0.002 | -0.088 |
| 5879 | -0.001 | -0.063 |
| 5880 | -0.001 | 0.030 |
| 5881 | -0.011 | 0.001 |
| 5882 | -0.016 | 0.040 |
| 5883 | 0.068 | 0.098 |
| 5884 | 0.042 | 0.010 |
| 5885 | 0.025 | -0.002 |
| 5886 | 0.040 | -0.030 |
| 5887 | -0.025 | -0.080 |
| 5888 | 0.035 | -0.016 |
| 5889 | 0.013 | -0.001 |
| 5890 | -0.094 | -0.003 |
| 5891 | 0.017 | -0.066 |
| 5892 | 0.055 | 0.053 |
| 5893 | -0.046 | -0.036 |
| 5894 | -0.022 | 0.031 |
| 5895 | 0.042 | 0.042 |
| 5896 | -0.039 | 0.004 |
| 5897 | -0.033 | -0.139 |
| 5898 | -0.009 | -0.088 |
| 5899 | -0.031 | 0.008 |
| 5900 | 0.065 | -0.087 |
| 5901 | -0.010 | -0.158 |
| 5902 | 0.042 | -0.076 |
| 5903 | 0.027 | -0.098 |
| 5904 | 0.011 | -0.096 |
| 5905 | 0.041 | -0.050 |
| 5906 | -0.036 | 0.040 |
| 5907 | -0.025 | -0.056 |
| 5908 | 0.010 | 0.046 |
| 5909 | 0.009 | -0.036 |

| | | |
|---|---|---|
| 5910 | -0.009 | -0.013 |
| 5911 | 0.014 | -0.101 |
| 5912 | -0.003 | 0.017 |
| 5913 | 0.042 | 0.067 |
| 5914 | -0.024 | -0.062 |
| 5915 | 0.006 | -0.079 |
| 5916 | 0.023 | -0.028 |
| 5917 | 0.028 | -0.068 |
| 5918 | -0.028 | 0.077 |
| 5919 | 0.034 | -0.105 |
| 5920 | -0.063 | -0.162 |
| 5921 | 0.013 | -0.043 |
| 5922 | -0.009 | -0.024 |
| 5923 | 0.075 | -0.027 |
| 5924 | -0.026 | 0.035 |
| 5925 | -0.038 | -0.026 |
| 5926 | 0.013 | -0.026 |
| 5927 | -0.033 | -0.024 |
| 5928 | -0.039 | -0.019 |
| 5929 | 0.037 | 0.080 |
| 5930 | 0.095 | -0.128 |
| 5931 | -0.004 | 0.003 |
| 5932 | 0.085 | 0.034 |
| 5933 | -0.051 | 0.079 |
| 5934 | 0.003 | -0.155 |
| 5935 | -0.012 | -0.036 |
| 5936 | 0.025 | 0.096 |
| 5937 | -0.010 | -0.046 |
| 5938 | 0.084 | 0.127 |
| 5939 | -0.017 | 0.127 |
| 5940 | -0.023 | -0.020 |
| 5941 | 0.041 | -0.041 |
| 5942 | -0.027 | 0.142 |
| 5943 | 0.056 | -0.089 |
| 5944 | -0.053 | 0.066 |
| 5945 | 0.040 | -0.045 |
| 5946 | -0.003 | -0.001 |
| 5947 | 0.039 | 0.028 |
| 5948 | 0.079 | -0.048 |
| 5949 | -0.028 | 0.034 |
| 5950 | -0.023 | 0.053 |
| 5951 | -0.004 | -0.058 |
| 5952 | -0.046 | 0.057 |
| 5953 | -0.055 | 0.026 |
| 5954 | -0.027 | 0.001 |
| 5955 | 0.000 | 0.008 |
| 5956 | 0.004 | 0.073 |
| 5957 | -0.027 | -0.151 |
| 5958 | 0.029 | 0.067 |
| 5959 | 0.036 | -0.102 |
| 5960 | -0.039 | 0.010 |

| | | |
|---|---|---|
| 5961 | -0.010 | -0.096 |
| 5962 | 0.007 | -0.129 |
| 5963 | -0.043 | 0.023 |
| 5964 | -0.061 | -0.001 |
| 5965 | 0.030 | -0.097 |
| 5966 | -0.004 | 0.085 |
| 5967 | 0.056 | -0.069 |
| 5968 | 0.027 | 0.040 |
| 5969 | -0.043 | -0.015 |
| 5970 | 0.010 | 0.071 |
| 5971 | 0.025 | -0.110 |
| 5972 | 0.007 | -0.087 |
| 5973 | -0.002 | -0.070 |
| 5974 | -0.043 | 0.001 |
| 5975 | 0.063 | -0.048 |
| 5976 | 0.054 | 0.135 |
| 5977 | 0.023 | -0.044 |
| 5978 | 0.048 | 0.040 |
| 5979 | 0.059 | 0.015 |
| 5980 | 0.008 | 0.087 |
| 5981 | -0.023 | -0.083 |
| 5982 | 0.042 | 0.084 |
| 5983 | -0.025 | 0.097 |
| 5984 | -0.001 | 0.047 |
| 5985 | -0.030 | -0.027 |
| 5986 | -0.023 | 0.099 |
| 5987 | 0.007 | 0.054 |
| 5988 | 0.012 | -0.023 |
| 5989 | -0.019 | 0.080 |
| 5990 | 0.001 | 0.014 |
| 5991 | -0.011 | 0.004 |
| 5992 | 0.061 | -0.053 |
| 5993 | -0.018 | 0.029 |
| 5994 | 0.034 | -0.063 |
| 5995 | -0.055 | 0.002 |
| 5996 | -0.023 | -0.037 |
| 5997 | 0.013 | 0.080 |
| 5998 | -0.003 | -0.085 |
| 5999 | -0.004 | -0.014 |
| 6000 | -0.044 | 0.020 |
| 6001 | -0.047 | 0.004 |
| 6002 | -0.015 | -0.037 |
| 6003 | -0.004 | -0.180 |
| 6004 | -0.044 | 0.056 |
| 6005 | -0.031 | -0.033 |
| 6006 | -0.007 | -0.039 |
| 6007 | -0.017 | 0.080 |
| 6008 | -0.043 | -0.117 |
| 6009 | -0.056 | 0.145 |
| 6010 | 0.004 | -0.022 |
| 6011 | -0.046 | 0.044 |

| | | |
|---|---|---|
| 6012 | -0.003 | -0.016 |
| 6013 | 0.002 | -0.064 |
| 6014 | 0.032 | -0.039 |
| 6015 | -0.018 | 0.000 |
| 6016 | 0.039 | 0.202 |
| 6017 | -0.028 | 0.035 |
| 6018 | 0.028 | -0.052 |
| 6019 | 0.045 | 0.110 |
| 6020 | -0.020 | -0.037 |
| 6021 | -0.051 | 0.011 |
| 6022 | -0.017 | 0.037 |
| 6023 | -0.002 | -0.084 |
| 6024 | -0.011 | -0.027 |
| 6025 | 0.014 | 0.048 |
| 6026 | 0.091 | -0.082 |
| 6027 | -0.007 | 0.055 |
| 6028 | -0.038 | 0.046 |
| 6029 | 0.020 | 0.062 |
| 6030 | -0.085 | 0.005 |
| 6031 | -0.017 | 0.019 |
| 6032 | 0.085 | 0.017 |
| 6033 | -0.054 | 0.050 |
| 6034 | -0.027 | 0.040 |
| 6035 | -0.063 | -0.029 |
| 6036 | 0.073 | -0.007 |
| 6037 | -0.039 | -0.136 |
| 6038 | 0.030 | -0.084 |
| 6039 | -0.009 | -0.002 |
| 6040 | 0.010 | -0.044 |
| 6041 | 0.036 | 0.052 |
| 6042 | -0.023 | 0.019 |
| 6043 | -0.060 | 0.109 |
| 6044 | 0.060 | 0.069 |
| 6045 | -0.024 | -0.001 |
| 6046 | 0.031 | 0.070 |
| 6047 | -0.015 | 0.027 |
| 6048 | 0.009 | -0.037 |
| 6049 | 0.030 | 0.031 |
| 6050 | -0.031 | -0.036 |
| 6051 | 0.011 | -0.082 |
| 6052 | -0.055 | -0.003 |
| 6053 | 0.035 | 0.164 |
| 6054 | 0.019 | -0.070 |
| 6055 | -0.047 | 0.034 |
| 6056 | -0.007 | -0.053 |
| 6057 | 0.008 | 0.045 |
| 6058 | -0.038 | -0.072 |
| 6059 | -0.001 | 0.078 |
| 6060 | 0.099 | -0.012 |
| 6061 | -0.028 | -0.136 |
| 6062 | 0.070 | 0.070 |

| | | |
|---|---|---|
| 6063 | 0.011 | 0.018 |
| 6064 | 0.027 | -0.035 |
| 6065 | -0.043 | -0.074 |
| 6066 | -0.064 | -0.037 |
| 6067 | -0.042 | -0.050 |
| 6068 | 0.021 | 0.074 |
| 6069 | 0.020 | -0.013 |
| 6070 | -0.003 | 0.106 |
| 6071 | 0.009 | 0.035 |
| 6072 | -0.038 | 0.157 |
| 6073 | -0.022 | 0.044 |
| 6074 | -0.044 | -0.164 |
| 6075 | -0.036 | -0.015 |
| 6076 | -0.043 | -0.035 |
| 6077 | 0.044 | 0.110 |
| 6078 | -0.030 | 0.095 |
| 6079 | -0.033 | 0.106 |
| 6080 | 0.022 | 0.021 |
| 6081 | 0.028 | 0.055 |
| 6082 | -0.008 | -0.023 |
| 6083 | -0.029 | -0.074 |
| 6084 | 0.001 | 0.061 |
| 6085 | -0.041 | 0.109 |
| 6086 | -0.047 | 0.060 |
| 6087 | 0.001 | 0.008 |
| 6088 | 0.011 | 0.004 |
| 6089 | -0.012 | -0.047 |
| 6090 | -0.045 | 0.084 |
| 6091 | -0.023 | 0.061 |
| 6092 | 0.028 | -0.102 |
| 6093 | 0.000 | -0.063 |
| 6094 | 0.041 | -0.077 |
| 6095 | -0.006 | -0.050 |
| 6096 | 0.024 | 0.013 |
| 6097 | -0.018 | -0.008 |
| 6098 | -0.044 | 0.029 |
| 6099 | 0.064 | -0.125 |
| 6100 | 0.020 | 0.021 |
| 6101 | 0.033 | 0.063 |
| 6102 | 0.020 | -0.031 |
| 6103 | -0.044 | -0.015 |
| 6104 | -0.085 | -0.108 |
| 6105 | 0.067 | 0.101 |
| 6106 | -0.010 | 0.120 |
| 6107 | 0.013 | -0.092 |
| 6108 | -0.007 | -0.121 |
| 6109 | 0.000 | 0.077 |
| 6110 | 0.025 | -0.033 |
| 6111 | -0.015 | 0.005 |
| 6112 | -0.010 | 0.042 |
| 6113 | -0.014 | -0.036 |

| | | |
|---|---|---|
| 6114 | 0.014 | 0.166 |
| 6115 | -0.006 | -0.024 |
| 6116 | 0.009 | 0.062 |
| 6117 | 0.035 | 0.194 |
| 6118 | 0.040 | 0.008 |
| 6119 | -0.015 | 0.071 |
| 6120 | -0.008 | -0.132 |
| 6121 | -0.049 | 0.113 |
| 6122 | -0.036 | -0.077 |
| 6123 | -0.008 | -0.041 |
| 6124 | -0.005 | -0.035 |
| 6125 | -0.054 | -0.030 |
| 6126 | 0.026 | 0.029 |
| 6127 | -0.046 | -0.090 |
| 6128 | -0.013 | -0.055 |
| 6129 | 0.003 | 0.184 |
| 6130 | 0.053 | 0.084 |
| 6131 | -0.024 | 0.069 |
| 6132 | -0.012 | 0.078 |
| 6133 | -0.053 | 0.094 |
| 6134 | -0.003 | -0.044 |
| 6135 | 0.048 | -0.062 |
| 6136 | -0.052 | 0.077 |
| 6137 | -0.004 | -0.116 |
| 6138 | -0.005 | 0.087 |
| 6139 | 0.014 | 0.158 |
| 6140 | 0.009 | -0.005 |
| 6141 | -0.006 | -0.184 |
| 6142 | 0.028 | -0.010 |
| 6143 | -0.010 | 0.020 |
| 6144 | -0.034 | -0.066 |
| 6145 | -0.002 | 0.039 |
| 6146 | -0.038 | 0.022 |
| 6147 | -0.024 | -0.108 |
| 6148 | 0.045 | -0.149 |
| 6149 | -0.007 | 0.064 |
| 6150 | -0.047 | -0.070 |
| 6151 | 0.012 | -0.053 |
| 6152 | 0.010 | -0.069 |
| 6153 | 0.032 | -0.099 |
| 6154 | 0.034 | -0.087 |
| 6155 | -0.014 | 0.034 |
| 6156 | -0.003 | -0.051 |
| 6157 | -0.027 | 0.083 |
| 6158 | 0.044 | 0.069 |
| 6159 | 0.031 | 0.026 |
| 6160 | -0.046 | 0.009 |
| 6161 | -0.053 | 0.022 |
| 6162 | -0.003 | -0.016 |
| 6163 | -0.035 | 0.007 |
| 6164 | -0.040 | 0.016 |

| | | |
|---|---|---|
| 6165 | -0.021 | -0.093 |
| 6166 | 0.046 | 0.048 |
| 6167 | 0.011 | 0.025 |
| 6168 | -0.027 | -0.022 |
| 6169 | -0.009 | 0.020 |
| 6170 | -0.051 | 0.102 |
| 6171 | 0.009 | -0.028 |
| 6172 | -0.014 | -0.142 |
| 6173 | -0.004 | -0.014 |
| 6174 | -0.030 | 0.007 |
| 6175 | -0.005 | 0.094 |
| 6176 | -0.023 | 0.024 |
| 6177 | -0.009 | -0.132 |
| 6178 | -0.007 | -0.097 |
| 6179 | -0.002 | 0.046 |
| 6180 | 0.020 | -0.039 |
| 6181 | 0.020 | 0.046 |
| 6182 | -0.002 | 0.022 |
| 6183 | -0.008 | -0.065 |
| 6184 | 0.085 | 0.099 |
| 6185 | 0.032 | -0.097 |
| 6186 | 0.015 | 0.005 |
| 6187 | 0.006 | 0.003 |
| 6188 | 0.006 | -0.049 |
| 6189 | 0.012 | 0.064 |
| 6190 | 0.039 | -0.062 |
| 6191 | -0.028 | 0.008 |
| 6192 | 0.014 | -0.141 |
| 6193 | 0.032 | 0.075 |
| 6194 | -0.030 | 0.070 |
| 6195 | 0.020 | -0.090 |
| 6196 | -0.007 | 0.061 |
| 6197 | 0.037 | -0.070 |
| 6198 | -0.052 | -0.109 |
| 6199 | -0.027 | 0.013 |
| 6200 | 0.039 | 0.070 |
| 6201 | 0.046 | 0.008 |
| 6202 | -0.005 | 0.089 |
| 6203 | -0.012 | 0.065 |
| 6204 | -0.046 | -0.031 |
| 6205 | 0.049 | -0.129 |
| 6206 | -0.016 | 0.055 |
| 6207 | 0.026 | -0.104 |
| 6208 | 0.021 | 0.130 |
| 6209 | 0.025 | -0.033 |
| 6210 | -0.039 | 0.051 |
| 6211 | 0.039 | 0.193 |
| 6212 | -0.009 | -0.013 |
| 6213 | 0.039 | 0.146 |
| 6214 | -0.012 | 0.037 |
| 6215 | -0.004 | 0.043 |

| | | |
|---|---|---|
| 6216 | 0.035 | -0.006 |
| 6217 | -0.022 | 0.025 |
| 6218 | 0.018 | -0.044 |
| 6219 | -0.050 | -0.017 |
| 6220 | 0.060 | -0.033 |
| 6221 | -0.009 | -0.053 |
| 6222 | 0.000 | 0.001 |
| 6223 | -0.027 | 0.011 |
| 6224 | -0.070 | 0.026 |
| 6225 | 0.002 | -0.054 |
| 6226 | 0.052 | -0.041 |
| 6227 | -0.015 | 0.001 |
| 6228 | 0.068 | 0.046 |
| 6229 | 0.053 | 0.049 |
| 6230 | 0.017 | -0.051 |
| 6231 | -0.066 | 0.059 |
| 6232 | -0.006 | 0.033 |
| 6233 | -0.037 | -0.181 |
| 6234 | 0.007 | -0.091 |
| 6235 | 0.009 | 0.040 |
| 6236 | 0.016 | 0.003 |
| 6237 | 0.011 | -0.074 |
| 6238 | -0.021 | 0.005 |
| 6239 | -0.013 | -0.054 |
| 6240 | -0.019 | -0.026 |
| 6241 | 0.005 | 0.074 |
| 6242 | 0.014 | -0.126 |
| 6243 | 0.029 | 0.065 |
| 6244 | -0.016 | 0.144 |
| 6245 | -0.057 | -0.014 |
| 6246 | -0.026 | 0.077 |
| 6247 | -0.003 | -0.101 |
| 6248 | 0.045 | 0.200 |
| 6249 | 0.007 | 0.143 |
| 6250 | 0.010 | 0.107 |
| 6251 | 0.067 | -0.054 |
| 6252 | 0.022 | -0.041 |
| 6253 | -0.078 | 0.005 |
| 6254 | 0.026 | 0.235 |
| 6255 | 0.057 | -0.006 |
| 6256 | -0.032 | 0.120 |
| 6257 | -0.017 | 0.039 |
| 6258 | 0.041 | -0.062 |
| 6259 | -0.004 | -0.070 |
| 6260 | -0.032 | -0.027 |
| 6261 | 0.009 | -0.010 |
| 6262 | -0.018 | -0.079 |
| 6263 | -0.012 | -0.017 |
| 6264 | -0.027 | 0.052 |
| 6265 | 0.028 | 0.038 |
| 6266 | 0.032 | -0.105 |

| | | |
|---|---|---|
| 6267 | -0.003 | 0.058 |
| 6268 | 0.034 | -0.142 |
| 6269 | -0.021 | 0.102 |
| 6270 | 0.034 | -0.084 |
| 6271 | -0.005 | 0.007 |
| 6272 | -0.005 | -0.040 |
| 6273 | -0.009 | 0.019 |
| 6274 | 0.015 | -0.086 |
| 6275 | -0.059 | -0.004 |
| 6276 | -0.014 | -0.109 |
| 6277 | 0.008 | 0.011 |
| 6278 | 0.015 | -0.059 |
| 6279 | 0.043 | 0.151 |
| 6280 | -0.020 | 0.102 |
| 6281 | 0.016 | -0.076 |
| 6282 | -0.015 | -0.036 |
| 6283 | -0.008 | 0.026 |
| 6284 | -0.004 | 0.097 |
| 6285 | -0.002 | -0.158 |
| 6286 | 0.033 | -0.037 |
| 6287 | -0.055 | 0.036 |
| 6288 | 0.066 | 0.168 |
| 6289 | 0.029 | 0.022 |
| 6290 | -0.012 | -0.030 |
| 6291 | -0.004 | -0.095 |
| 6292 | -0.055 | -0.115 |
| 6293 | -0.051 | 0.112 |
| 6294 | -0.008 | 0.163 |
| 6295 | -0.047 | 0.093 |
| 6296 | 0.009 | -0.042 |
| 6297 | 0.018 | -0.072 |
| 6298 | -0.028 | 0.008 |
| 6299 | -0.008 | -0.024 |
| 6300 | 0.010 | 0.034 |
| 6301 | 0.015 | -0.014 |
| 6302 | 0.065 | 0.077 |
| 6303 | 0.015 | -0.049 |
| 6304 | -0.026 | -0.087 |
| 6305 | 0.011 | 0.135 |
| 6306 | -0.026 | 0.028 |
| 6307 | -0.008 | -0.038 |
| 6308 | 0.015 | -0.142 |
| 6309 | -0.001 | -0.105 |
| 6310 | -0.027 | 0.130 |
| 6311 | -0.062 | 0.032 |
| 6312 | -0.034 | -0.135 |
| 6313 | 0.034 | 0.080 |
| 6314 | -0.005 | 0.048 |
| 6315 | -0.009 | 0.154 |
| 6316 | -0.025 | 0.024 |
| 6317 | 0.034 | -0.045 |

| | | |
|---|---|---|
| 6318 | 0.072 | -0.043 |
| 6319 | 0.010 | -0.087 |
| 6320 | 0.009 | -0.017 |
| 6321 | 0.002 | -0.038 |
| 6322 | -0.018 | 0.084 |
| 6323 | 0.050 | 0.024 |
| 6324 | -0.022 | 0.070 |
| 6325 | -0.013 | 0.056 |
| 6326 | 0.056 | 0.064 |
| 6327 | 0.126 | -0.046 |
| 6328 | 0.025 | 0.055 |
| 6329 | -0.055 | -0.020 |
| 6330 | 0.007 | 0.041 |
| 6331 | -0.005 | -0.078 |
| 6332 | 0.000 | -0.079 |
| 6333 | -0.016 | -0.058 |
| 6334 | -0.046 | 0.032 |
| 6335 | 0.069 | 0.087 |
| 6336 | -0.019 | -0.012 |
| 6337 | -0.002 | 0.037 |
| 6338 | -0.019 | 0.055 |
| 6339 | 0.003 | -0.012 |
| 6340 | -0.033 | -0.149 |
| 6341 | -0.005 | -0.080 |
| 6342 | -0.010 | 0.119 |
| 6343 | -0.002 | -0.079 |
| 6344 | -0.008 | -0.095 |
| 6345 | 0.006 | -0.072 |
| 6346 | 0.011 | 0.204 |
| 6347 | 0.000 | -0.039 |
| 6348 | 0.008 | -0.038 |
| 6349 | -0.027 | 0.031 |
| 6350 | 0.008 | 0.140 |
| 6351 | -0.016 | -0.054 |
| 6352 | 0.079 | 0.109 |
| 6353 | -0.015 | 0.084 |
| 6354 | 0.027 | 0.085 |
| 6355 | -0.007 | -0.006 |
| 6356 | -0.024 | 0.026 |
| 6357 | 0.004 | 0.070 |
| 6358 | 0.020 | -0.027 |
| 6359 | -0.038 | -0.039 |
| 6360 | -0.006 | -0.078 |
| 6361 | -0.011 | 0.045 |
| 6362 | 0.008 | 0.056 |
| 6363 | 0.048 | 0.030 |
| 6364 | 0.060 | -0.016 |
| 6365 | -0.046 | 0.001 |
| 6366 | -0.029 | -0.014 |
| 6367 | -0.046 | -0.089 |
| 6368 | -0.040 | -0.105 |

| | | |
|---|---|---|
| 6369 | 0.007 | -0.056 |
| 6370 | -0.033 | -0.047 |
| 6371 | -0.047 | -0.009 |
| 6372 | 0.034 | 0.078 |
| 6373 | 0.061 | -0.042 |
| 6374 | -0.021 | 0.038 |
| 6375 | 0.000 | 0.098 |
| 6376 | -0.022 | 0.016 |
| 6377 | -0.081 | 0.058 |
| 6378 | -0.083 | 0.084 |
| 6379 | -0.011 | -0.040 |
| 6380 | -0.034 | -0.002 |
| 6381 | -0.027 | -0.005 |
| 6382 | 0.042 | 0.002 |
| 6383 | -0.029 | -0.110 |
| 6384 | -0.006 | 0.076 |
| 6385 | 0.005 | 0.086 |
| 6386 | -0.019 | -0.037 |
| 6387 | -0.010 | 0.217 |
| 6388 | -0.024 | -0.032 |
| 6389 | 0.003 | -0.065 |
| 6390 | -0.075 | 0.084 |
| 6391 | -0.059 | 0.001 |
| 6392 | -0.031 | -0.007 |
| 6393 | -0.017 | 0.019 |
| 6394 | -0.024 | 0.113 |
| 6395 | 0.025 | 0.109 |
| 6396 | -0.011 | 0.044 |
| 6397 | 0.012 | -0.048 |
| 6398 | -0.022 | 0.107 |
| 6399 | -0.023 | -0.084 |
| 6400 | 0.023 | 0.036 |
| 6401 | -0.042 | -0.060 |
| 6402 | -0.028 | 0.002 |
| 6403 | 0.018 | 0.152 |
| 6404 | 0.035 | 0.022 |
| 6405 | 0.004 | 0.040 |
| 6406 | -0.026 | 0.014 |
| 6407 | -0.048 | -0.038 |
| 6408 | 0.049 | -0.027 |
| 6409 | 0.009 | -0.014 |
| 6410 | -0.007 | 0.106 |
| 6411 | 0.010 | -0.122 |
| 6412 | 0.019 | 0.003 |
| 6413 | -0.016 | -0.074 |
| 6414 | 0.048 | 0.055 |
| 6415 | 0.017 | 0.200 |
| 6416 | 0.041 | -0.175 |
| 6417 | -0.011 | 0.167 |
| 6418 | -0.036 | -0.001 |
| 6419 | -0.028 | 0.019 |

| | | |
|---|---|---|
| 6420 | 0.055 | 0.045 |
| 6421 | 0.016 | -0.022 |
| 6422 | 0.014 | 0.015 |
| 6423 | -0.031 | 0.028 |
| 6424 | 0.035 | -0.091 |
| 6425 | -0.010 | 0.035 |
| 6426 | 0.034 | -0.068 |
| 6427 | 0.016 | -0.116 |
| 6428 | 0.019 | -0.053 |
| 6429 | -0.020 | 0.031 |
| 6430 | 0.006 | 0.066 |
| 6431 | -0.066 | -0.015 |
| 6432 | 0.039 | -0.038 |
| 6433 | 0.026 | -0.004 |
| 6434 | -0.035 | -0.050 |
| 6435 | 0.017 | 0.054 |
| 6436 | -0.005 | 0.016 |
| 6437 | 0.066 | -0.008 |
| 6438 | 0.005 | 0.015 |
| 6439 | 0.029 | -0.029 |
| 6440 | 0.012 | -0.016 |
| 6441 | 0.015 | -0.026 |
| 6442 | -0.030 | -0.082 |
| 6443 | 0.029 | 0.054 |
| 6444 | 0.044 | -0.041 |
| 6445 | 0.047 | 0.067 |
| 6446 | -0.042 | 0.146 |
| 6447 | 0.011 | 0.061 |
| 6448 | -0.040 | 0.020 |
| 6449 | -0.051 | 0.109 |
| 6450 | -0.036 | -0.046 |
| 6451 | 0.011 | -0.059 |
| 6452 | -0.044 | -0.057 |
| 6453 | -0.007 | 0.063 |
| 6454 | -0.022 | 0.008 |
| 6455 | -0.018 | -0.049 |
| 6456 | -0.020 | -0.056 |
| 6457 | 0.047 | 0.027 |
| 6458 | -0.043 | -0.055 |
| 6459 | 0.037 | 0.046 |
| 6460 | -0.036 | -0.064 |
| 6461 | -0.039 | 0.005 |
| 6462 | 0.021 | -0.027 |
| 6463 | 0.004 | -0.083 |
| 6464 | 0.034 | -0.063 |
| 6465 | -0.049 | 0.036 |
| 6466 | -0.075 | -0.046 |
| 6467 | -0.053 | -0.026 |
| 6468 | -0.014 | -0.074 |
| 6469 | 0.027 | 0.111 |
| 6470 | 0.054 | -0.024 |

| | | |
|---|---|---|
| 6471 | -0.013 | 0.037 |
| 6472 | 0.020 | 0.032 |
| 6473 | 0.018 | 0.050 |
| 6474 | -0.028 | -0.029 |
| 6475 | -0.054 | 0.062 |
| 6476 | -0.023 | -0.055 |
| 6477 | -0.005 | 0.062 |
| 6478 | 0.039 | 0.007 |
| 6479 | -0.052 | 0.139 |
| 6480 | 0.041 | -0.055 |
| 6481 | -0.011 | 0.048 |
| 6482 | -0.017 | -0.029 |
| 6483 | -0.012 | 0.164 |
| 6484 | -0.065 | -0.151 |
| 6485 | -0.041 | -0.085 |
| 6486 | 0.035 | 0.037 |
| 6487 | 0.021 | 0.067 |
| 6488 | -0.011 | -0.099 |
| 6489 | 0.069 | -0.097 |
| 6490 | 0.042 | -0.006 |
| 6491 | 0.001 | -0.022 |
| 6492 | 0.026 | -0.092 |
| 6493 | -0.022 | 0.130 |
| 6494 | 0.019 | -0.087 |
| 6495 | -0.041 | 0.096 |
| 6496 | 0.012 | -0.025 |
| 6497 | -0.023 | -0.010 |
| 6498 | -0.012 | 0.006 |
| 6499 | -0.015 | -0.117 |
| 6500 | -0.021 | 0.159 |
| 6501 | 0.052 | -0.009 |
| 6502 | 0.022 | 0.088 |
| 6503 | -0.063 | -0.029 |
| 6504 | -0.042 | 0.060 |
| 6505 | 0.025 | -0.026 |
| 6506 | -0.040 | 0.067 |
| 6507 | 0.013 | 0.013 |
| 6508 | -0.061 | -0.015 |
| 6509 | -0.014 | 0.014 |
| 6510 | 0.027 | -0.054 |
| 6511 | -0.033 | -0.014 |
| 6512 | 0.027 | -0.005 |
| 6513 | -0.009 | 0.036 |
| 6514 | -0.075 | 0.037 |
| 6515 | -0.041 | 0.084 |
| 6516 | 0.021 | -0.051 |
| 6517 | 0.025 | 0.021 |
| 6518 | -0.029 | 0.050 |
| 6519 | -0.014 | -0.138 |
| 6520 | -0.006 | 0.049 |
| 6521 | 0.042 | -0.091 |

| | | |
|---|---|---|
| 6522 | 0.046 | -0.123 |
| 6523 | -0.018 | -0.052 |
| 6524 | 0.018 | -0.022 |
| 6525 | -0.018 | -0.001 |
| 6526 | 0.021 | 0.162 |
| 6527 | 0.025 | -0.018 |
| 6528 | -0.048 | -0.038 |
| 6529 | -0.014 | -0.036 |
| 6530 | 0.010 | -0.007 |
| 6531 | -0.003 | 0.049 |
| 6532 | 0.051 | 0.127 |
| 6533 | 0.007 | -0.005 |
| 6534 | -0.020 | 0.035 |
| 6535 | -0.034 | 0.116 |
| 6536 | 0.003 | -0.007 |
| 6537 | 0.062 | -0.066 |
| 6538 | -0.075 | 0.083 |
| 6539 | -0.033 | -0.020 |
| 6540 | 0.063 | 0.044 |
| 6541 | -0.003 | 0.054 |
| 6542 | -0.008 | -0.027 |
| 6543 | 0.021 | -0.153 |
| 6544 | -0.001 | -0.050 |
| 6545 | -0.033 | 0.048 |
| 6546 | 0.005 | -0.022 |
| 6547 | 0.047 | -0.063 |
| 6548 | 0.026 | 0.135 |
| 6549 | -0.024 | -0.127 |
| 6550 | -0.022 | -0.056 |
| 6551 | -0.021 | -0.212 |
| 6552 | 0.027 | -0.047 |
| 6553 | 0.009 | -0.085 |
| 6554 | -0.009 | -0.179 |
| 6555 | 0.024 | 0.149 |
| 6556 | -0.020 | -0.130 |
| 6557 | 0.007 | -0.067 |
| 6558 | 0.016 | -0.004 |
| 6559 | -0.008 | 0.083 |
| 6560 | -0.025 | 0.026 |
| 6561 | 0.054 | 0.039 |
| 6562 | 0.041 | 0.039 |
| 6563 | 0.005 | -0.071 |
| 6564 | 0.003 | -0.061 |
| 6565 | 0.028 | -0.013 |
| 6566 | -0.003 | 0.043 |
| 6567 | 0.019 | -0.048 |
| 6568 | 0.019 | 0.074 |
| 6569 | -0.013 | 0.057 |
| 6570 | -0.019 | -0.043 |
| 6571 | -0.006 | 0.049 |
| 6572 | 0.024 | -0.093 |

| | | |
|---|---|---|
| 6573 | 0.056 | 0.002 |
| 6574 | 0.073 | 0.022 |
| 6575 | -0.030 | 0.005 |
| 6576 | -0.028 | -0.054 |
| 6577 | 0.023 | 0.058 |
| 6578 | 0.049 | -0.242 |
| 6579 | -0.015 | 0.010 |
| 6580 | -0.022 | -0.166 |
| 6581 | -0.060 | -0.108 |
| 6582 | -0.039 | 0.012 |
| 6583 | -0.010 | 0.024 |
| 6584 | 0.044 | 0.029 |
| 6585 | 0.017 | 0.076 |
| 6586 | -0.004 | -0.058 |
| 6587 | -0.026 | -0.060 |
| 6588 | -0.015 | 0.107 |
| 6589 | 0.009 | -0.001 |
| 6590 | -0.029 | 0.019 |
| 6591 | 0.013 | -0.030 |
| 6592 | -0.009 | 0.070 |
| 6593 | 0.063 | -0.081 |
| 6594 | -0.071 | 0.023 |
| 6595 | -0.025 | 0.039 |
| 6596 | 0.016 | -0.119 |
| 6597 | 0.016 | 0.123 |
| 6598 | 0.031 | 0.036 |
| 6599 | 0.006 | -0.065 |
| 6600 | 0.015 | -0.038 |
| 6601 | 0.043 | 0.115 |
| 6602 | -0.100 | -0.057 |
| 6603 | 0.013 | -0.057 |
| 6604 | 0.003 | 0.022 |
| 6605 | 0.025 | -0.102 |
| 6606 | 0.022 | -0.114 |
| 6607 | -0.025 | -0.062 |
| 6608 | 0.048 | -0.073 |
| 6609 | -0.018 | 0.000 |
| 6610 | 0.020 | 0.000 |
| 6611 | 0.015 | 0.160 |
| 6612 | 0.026 | -0.004 |
| 6613 | -0.021 | -0.013 |
| 6614 | -0.071 | 0.057 |
| 6615 | 0.091 | -0.059 |
| 6616 | 0.012 | 0.077 |
| 6617 | 0.042 | 0.092 |
| 6618 | -0.060 | 0.018 |
| 6619 | 0.051 | -0.029 |
| 6620 | -0.066 | 0.071 |
| 6621 | -0.036 | 0.140 |
| 6622 | -0.027 | -0.097 |
| 6623 | -0.042 | 0.073 |

| | | |
|---|---|---|
| 6624 | 0.030 | 0.098 |
| 6625 | -0.011 | 0.027 |
| 6626 | 0.035 | 0.017 |
| 6627 | 0.083 | 0.014 |
| 6628 | 0.034 | 0.020 |
| 6629 | 0.071 | 0.006 |
| 6630 | 0.000 | 0.016 |
| 6631 | 0.005 | -0.091 |
| 6632 | 0.042 | -0.028 |
| 6633 | 0.027 | -0.140 |
| 6634 | 0.010 | 0.008 |
| 6635 | -0.055 | 0.110 |
| 6636 | 0.033 | 0.074 |
| 6637 | -0.087 | -0.031 |
| 6638 | -0.004 | -0.086 |
| 6639 | 0.024 | -0.130 |
| 6640 | -0.057 | -0.040 |
| 6641 | 0.027 | 0.094 |
| 6642 | 0.002 | 0.072 |
| 6643 | -0.020 | -0.133 |
| 6644 | 0.003 | -0.108 |
| 6645 | 0.024 | 0.052 |
| 6646 | 0.085 | -0.065 |
| 6647 | -0.020 | 0.181 |
| 6648 | 0.096 | 0.027 |
| 6649 | 0.003 | 0.033 |
| 6650 | -0.003 | -0.019 |
| 6651 | -0.024 | -0.106 |
| 6652 | -0.002 | -0.106 |
| 6653 | 0.034 | 0.118 |
| 6654 | -0.022 | -0.051 |
| 6655 | 0.026 | 0.056 |
| 6656 | -0.050 | 0.011 |
| 6657 | 0.007 | -0.019 |
| 6658 | 0.009 | -0.052 |
| 6659 | 0.061 | 0.032 |
| 6660 | -0.029 | 0.072 |
| 6661 | -0.044 | 0.068 |
| 6662 | -0.040 | -0.098 |
| 6663 | -0.033 | -0.045 |
| 6664 | 0.041 | -0.094 |
| 6665 | 0.022 | -0.018 |
| 6666 | -0.021 | 0.005 |
| 6667 | -0.003 | 0.123 |
| 6668 | 0.025 | -0.066 |
| 6669 | 0.005 | -0.049 |
| 6670 | -0.014 | 0.137 |
| 6671 | 0.064 | -0.128 |
| 6672 | -0.022 | 0.003 |
| 6673 | 0.003 | -0.109 |
| 6674 | 0.008 | -0.081 |

| | | |
|---|---|---|
| 6675 | -0.014 | 0.016 |
| 6676 | 0.009 | 0.026 |
| 6677 | 0.026 | 0.035 |
| 6678 | -0.025 | 0.015 |
| 6679 | 0.048 | -0.025 |
| 6680 | 0.009 | 0.057 |
| 6681 | -0.019 | -0.013 |
| 6682 | 0.042 | 0.086 |
| 6683 | 0.003 | 0.044 |
| 6684 | -0.008 | -0.085 |
| 6685 | -0.024 | 0.099 |
| 6686 | 0.003 | 0.074 |
| 6687 | 0.024 | 0.040 |
| 6688 | 0.021 | 0.030 |
| 6689 | 0.021 | 0.076 |
| 6690 | 0.018 | 0.179 |
| 6691 | 0.068 | -0.035 |
| 6692 | 0.088 | -0.084 |
| 6693 | 0.005 | -0.100 |
| 6694 | -0.040 | 0.114 |
| 6695 | 0.015 | -0.124 |
| 6696 | -0.010 | 0.048 |
| 6697 | -0.073 | -0.109 |
| 6698 | 0.008 | -0.112 |
| 6699 | -0.030 | 0.040 |
| 6700 | 0.042 | -0.060 |
| 6701 | 0.052 | -0.037 |
| 6702 | -0.027 | 0.119 |
| 6703 | -0.001 | -0.075 |
| 6704 | 0.019 | 0.009 |
| 6705 | -0.015 | 0.046 |
| 6706 | 0.003 | 0.062 |
| 6707 | -0.022 | -0.071 |
| 6708 | 0.029 | -0.047 |
| 6709 | -0.009 | 0.002 |
| 6710 | -0.077 | -0.085 |
| 6711 | 0.011 | -0.036 |
| 6712 | -0.029 | -0.008 |
| 6713 | -0.018 | -0.042 |
| 6714 | 0.001 | 0.073 |
| 6715 | -0.001 | -0.120 |
| 6716 | 0.050 | -0.062 |
| 6717 | -0.011 | 0.021 |
| 6718 | 0.032 | -0.022 |
| 6719 | 0.060 | 0.109 |
| 6720 | 0.023 | -0.117 |
| 6721 | 0.060 | 0.071 |
| 6722 | -0.006 | -0.106 |
| 6723 | -0.036 | -0.002 |
| 6724 | -0.023 | -0.027 |
| 6725 | -0.039 | -0.015 |

| | | |
|---|---|---|
| 6726 | 0.011 | -0.025 |
| 6727 | 0.000 | 0.024 |
| 6728 | 0.011 | -0.036 |
| 6729 | 0.040 | -0.042 |
| 6730 | 0.001 | 0.036 |
| 6731 | -0.076 | -0.048 |
| 6732 | 0.003 | -0.032 |
| 6733 | 0.044 | -0.079 |
| 6734 | 0.021 | 0.068 |
| 6735 | -0.039 | -0.042 |
| 6736 | -0.015 | -0.050 |
| 6737 | -0.006 | -0.022 |
| 6738 | -0.035 | -0.074 |
| 6739 | 0.002 | 0.079 |
| 6740 | -0.021 | 0.013 |
| 6741 | 0.005 | -0.046 |
| 6742 | 0.016 | 0.061 |
| 6743 | -0.005 | 0.053 |
| 6744 | -0.009 | -0.045 |
| 6745 | -0.012 | -0.065 |
| 6746 | 0.012 | -0.099 |
| 6747 | -0.023 | -0.037 |
| 6748 | 0.022 | -0.035 |
| 6749 | -0.036 | 0.133 |
| 6750 | 0.032 | 0.000 |
| 6751 | -0.027 | -0.155 |
| 6752 | 0.049 | 0.064 |
| 6753 | 0.023 | -0.042 |
| 6754 | 0.070 | 0.002 |
| 6755 | -0.025 | -0.114 |
| 6756 | 0.019 | -0.031 |
| 6757 | -0.016 | 0.079 |
| 6758 | -0.013 | -0.059 |
| 6759 | -0.036 | -0.104 |
| 6760 | -0.012 | 0.062 |
| 6761 | -0.010 | 0.063 |
| 6762 | 0.027 | -0.070 |
| 6763 | -0.057 | 0.015 |
| 6764 | 0.057 | -0.006 |
| 6765 | -0.011 | 0.066 |
| 6766 | 0.025 | -0.091 |
| 6767 | -0.022 | 0.034 |
| 6768 | 0.005 | 0.182 |
| 6769 | 0.009 | 0.104 |
| 6770 | 0.011 | 0.059 |
| 6771 | 0.000 | 0.030 |
| 6772 | 0.005 | 0.123 |
| 6773 | -0.046 | 0.094 |
| 6774 | -0.004 | 0.028 |
| 6775 | -0.003 | -0.025 |
| 6776 | 0.010 | 0.010 |

| | | |
|---|---|---|
| 6777 | 0.025 | -0.015 |
| 6778 | -0.037 | -0.047 |
| 6779 | -0.010 | -0.024 |
| 6780 | 0.029 | 0.130 |
| 6781 | -0.030 | -0.014 |
| 6782 | 0.009 | 0.014 |
| 6783 | -0.001 | 0.086 |
| 6784 | -0.004 | 0.018 |
| 6785 | -0.019 | -0.065 |
| 6786 | 0.012 | -0.163 |
| 6787 | -0.013 | 0.016 |
| 6788 | -0.051 | -0.019 |
| 6789 | 0.006 | 0.013 |
| 6790 | 0.003 | -0.002 |
| 6791 | -0.013 | -0.041 |
| 6792 | -0.010 | -0.019 |
| 6793 | 0.017 | 0.017 |
| 6794 | -0.021 | -0.070 |
| 6795 | -0.012 | -0.027 |
| 6796 | -0.019 | 0.024 |
| 6797 | 0.019 | -0.018 |
| 6798 | 0.002 | -0.039 |
| 6799 | 0.021 | 0.019 |
| 6800 | 0.014 | 0.117 |
| 6801 | -0.017 | 0.048 |
| 6802 | 0.040 | -0.121 |
| 6803 | 0.015 | 0.043 |
| 6804 | 0.027 | 0.084 |
| 6805 | -0.011 | 0.006 |
| 6806 | 0.005 | 0.009 |
| 6807 | -0.005 | 0.004 |
| 6808 | -0.009 | 0.049 |
| 6809 | 0.030 | -0.048 |
| 6810 | 0.008 | 0.067 |
| 6811 | -0.003 | -0.014 |
| 6812 | 0.048 | 0.045 |
| 6813 | -0.021 | -0.076 |
| 6814 | 0.051 | 0.048 |
| 6815 | 0.005 | -0.055 |
| 6816 | 0.028 | 0.077 |
| 6817 | -0.030 | 0.040 |
| 6818 | -0.013 | 0.066 |
| 6819 | -0.011 | 0.066 |
| 6820 | 0.071 | -0.027 |
| 6821 | 0.021 | -0.034 |
| 6822 | -0.013 | -0.081 |
| 6823 | -0.024 | -0.077 |
| 6824 | -0.051 | 0.087 |
| 6825 | -0.003 | 0.218 |
| 6826 | 0.000 | -0.021 |
| 6827 | 0.007 | -0.028 |

| | | |
|---|---|---|
| 6828 | 0.002 | 0.072 |
| 6829 | 0.031 | 0.005 |
| 6830 | 0.005 | -0.041 |
| 6831 | -0.032 | -0.130 |
| 6832 | -0.037 | -0.061 |
| 6833 | 0.013 | 0.002 |
| 6834 | 0.045 | 0.020 |
| 6835 | 0.025 | 0.008 |
| 6836 | 0.005 | 0.033 |
| 6837 | 0.020 | -0.001 |
| 6838 | -0.020 | 0.103 |
| 6839 | 0.030 | 0.027 |
| 6840 | 0.018 | -0.030 |
| 6841 | -0.002 | -0.015 |
| 6842 | -0.034 | -0.090 |
| 6843 | 0.045 | -0.011 |
| 6844 | -0.024 | 0.043 |
| 6845 | -0.005 | -0.084 |
| 6846 | -0.105 | 0.077 |
| 6847 | -0.040 | -0.012 |
| 6848 | 0.028 | 0.089 |
| 6849 | 0.015 | -0.091 |
| 6850 | 0.029 | 0.042 |
| 6851 | 0.008 | 0.009 |
| 6852 | -0.031 | 0.018 |
| 6853 | -0.055 | -0.140 |
| 6854 | 0.049 | 0.029 |
| 6855 | 0.045 | -0.089 |
| 6856 | -0.049 | 0.001 |
| 6857 | -0.014 | -0.044 |
| 6858 | 0.025 | -0.085 |
| 6859 | 0.032 | -0.018 |
| 6860 | 0.020 | 0.111 |
| 6861 | -0.016 | -0.106 |
| 6862 | 0.058 | -0.017 |
| 6863 | 0.006 | 0.166 |
| 6864 | 0.026 | -0.045 |
| 6865 | -0.002 | -0.070 |
| 6866 | 0.051 | 0.038 |
| 6867 | -0.054 | 0.007 |
| 6868 | 0.057 | 0.024 |
| 6869 | 0.015 | 0.076 |
| 6870 | 0.066 | -0.087 |
| 6871 | 0.012 | -0.065 |
| 6872 | -0.010 | -0.086 |
| 6873 | -0.021 | 0.009 |
| 6874 | -0.034 | -0.029 |
| 6875 | -0.048 | -0.043 |
| 6876 | 0.008 | 0.090 |
| 6877 | 0.043 | 0.022 |
| 6878 | 0.033 | -0.098 |

| | | |
|---|---|---|
| 6879 | -0.016 | -0.028 |
| 6880 | 0.008 | 0.061 |
| 6881 | -0.045 | 0.080 |
| 6882 | -0.027 | -0.090 |
| 6883 | -0.006 | -0.030 |
| 6884 | -0.046 | -0.087 |
| 6885 | 0.009 | 0.124 |
| 6886 | 0.027 | -0.028 |
| 6887 | -0.024 | 0.004 |
| 6888 | 0.042 | 0.033 |
| 6889 | 0.009 | -0.003 |
| 6890 | 0.008 | -0.051 |
| 6891 | 0.002 | 0.096 |
| 6892 | -0.040 | -0.026 |
| 6893 | 0.044 | -0.034 |
| 6894 | 0.022 | -0.132 |
| 6895 | -0.021 | 0.034 |
| 6896 | 0.009 | -0.169 |
| 6897 | 0.009 | 0.147 |
| 6898 | -0.046 | 0.029 |
| 6899 | -0.040 | 0.025 |
| 6900 | 0.037 | 0.002 |
| 6901 | 0.016 | 0.145 |
| 6902 | 0.082 | -0.107 |
| 6903 | -0.011 | 0.038 |
| 6904 | 0.005 | 0.052 |
| 6905 | -0.008 | -0.040 |
| 6906 | 0.064 | 0.065 |
| 6907 | -0.011 | -0.057 |
| 6908 | 0.039 | 0.195 |
| 6909 | 0.028 | 0.036 |
| 6910 | -0.012 | 0.009 |
| 6911 | -0.039 | 0.012 |
| 6912 | 0.005 | 0.042 |
| 6913 | -0.028 | 0.031 |
| 6914 | -0.054 | 0.141 |
| 6915 | -0.050 | -0.020 |
| 6916 | 0.023 | 0.078 |
| 6917 | 0.055 | -0.107 |
| 6918 | 0.002 | 0.034 |
| 6919 | 0.005 | 0.092 |
| 6920 | 0.007 | 0.020 |
| 6921 | 0.041 | 0.166 |
| 6922 | 0.015 | -0.030 |
| 6923 | 0.019 | -0.156 |
| 6924 | -0.015 | -0.061 |
| 6925 | -0.017 | 0.078 |
| 6926 | 0.011 | -0.014 |
| 6927 | 0.020 | 0.044 |
| 6928 | -0.031 | 0.120 |
| 6929 | 0.019 | -0.008 |

| | | |
|---|---|---|
| 6930 | 0.029 | 0.076 |
| 6931 | 0.007 | -0.009 |
| 6932 | -0.041 | -0.001 |
| 6933 | 0.001 | 0.001 |
| 6934 | -0.077 | -0.086 |
| 6935 | 0.028 | 0.141 |
| 6936 | 0.057 | -0.106 |
| 6937 | 0.039 | -0.049 |
| 6938 | -0.005 | -0.054 |
| 6939 | -0.029 | 0.027 |
| 6940 | -0.012 | 0.080 |
| 6941 | -0.008 | -0.096 |
| 6942 | -0.025 | -0.009 |
| 6943 | -0.036 | 0.102 |
| 6944 | -0.021 | 0.013 |
| 6945 | -0.011 | 0.130 |
| 6946 | -0.017 | 0.021 |
| 6947 | -0.007 | -0.006 |
| 6948 | -0.002 | 0.047 |
| 6949 | -0.052 | -0.027 |
| 6950 | -0.011 | 0.101 |
| 6951 | -0.001 | -0.033 |
| 6952 | 0.031 | -0.056 |
| 6953 | -0.004 | 0.011 |
| 6954 | 0.022 | 0.020 |
| 6955 | 0.008 | 0.016 |
| 6956 | -0.039 | -0.063 |
| 6957 | -0.043 | 0.038 |
| 6958 | 0.005 | -0.033 |
| 6959 | 0.030 | 0.024 |
| 6960 | -0.014 | 0.079 |
| 6961 | -0.084 | 0.145 |
| 6962 | 0.037 | -0.016 |
| 6963 | 0.048 | -0.008 |
| 6964 | -0.010 | -0.028 |
| 6965 | 0.007 | 0.032 |
| 6966 | 0.007 | -0.009 |
| 6967 | 0.042 | -0.047 |
| 6968 | -0.035 | -0.038 |
| 6969 | 0.015 | 0.018 |
| 6970 | -0.043 | -0.033 |
| 6971 | 0.067 | 0.081 |
| 6972 | 0.036 | 0.015 |
| 6973 | -0.087 | -0.081 |
| 6974 | -0.014 | 0.000 |
| 6975 | -0.013 | 0.071 |
| 6976 | -0.012 | 0.069 |
| 6977 | 0.042 | 0.156 |
| 6978 | -0.003 | -0.023 |
| 6979 | -0.039 | 0.027 |
| 6980 | 0.028 | 0.010 |

| | | |
|---|---|---|
| 6981 | 0.028 | -0.039 |
| 6982 | 0.031 | 0.031 |
| 6983 | -0.081 | -0.031 |
| 6984 | 0.000 | 0.047 |
| 6985 | 0.038 | -0.011 |
| 6986 | 0.011 | 0.047 |
| 6987 | -0.072 | 0.090 |
| 6988 | -0.034 | -0.036 |
| 6989 | -0.048 | 0.077 |
| 6990 | 0.016 | 0.113 |
| 6991 | 0.022 | 0.083 |
| 6992 | -0.030 | 0.061 |
| 6993 | -0.022 | -0.052 |
| 6994 | 0.036 | -0.151 |
| 6995 | -0.002 | -0.060 |
| 6996 | 0.024 | 0.113 |
| 6997 | -0.029 | -0.030 |
| 6998 | -0.020 | 0.041 |
| 6999 | 0.005 | -0.060 |
| 7000 | 0.039 | -0.025 |
| 7001 | -0.002 | 0.062 |
| 7002 | 0.001 | -0.051 |
| 7003 | 0.030 | 0.095 |
| 7004 | -0.032 | 0.038 |
| 7005 | -0.024 | -0.039 |
| 7006 | 0.029 | 0.049 |
| 7007 | -0.049 | 0.022 |
| 7008 | 0.027 | -0.022 |
| 7009 | 0.005 | -0.040 |
| 7010 | 0.022 | -0.148 |
| 7011 | 0.004 | -0.030 |
| 7012 | -0.059 | 0.069 |
| 7013 | 0.026 | 0.062 |
| 7014 | 0.015 | -0.004 |
| 7015 | -0.103 | 0.014 |
| 7016 | 0.021 | 0.076 |
| 7017 | -0.042 | 0.033 |
| 7018 | -0.004 | -0.037 |
| 7019 | 0.019 | 0.045 |
| 7020 | -0.017 | -0.022 |
| 7021 | 0.060 | -0.056 |
| 7022 | 0.033 | -0.073 |
| 7023 | 0.029 | 0.129 |
| 7024 | -0.011 | 0.027 |
| 7025 | -0.087 | -0.051 |
| 7026 | 0.022 | -0.038 |
| 7027 | 0.000 | -0.020 |
| 7028 | 0.040 | 0.053 |
| 7029 | 0.013 | 0.061 |
| 7030 | -0.023 | -0.024 |
| 7031 | 0.085 | -0.116 |

| | | |
|---|---|---|
| 7032 | -0.031 | -0.020 |
| 7033 | -0.057 | 0.047 |
| 7034 | -0.011 | 0.074 |
| 7035 | 0.025 | 0.166 |
| 7036 | 0.039 | -0.036 |
| 7037 | -0.029 | 0.005 |
| 7038 | -0.001 | 0.119 |
| 7039 | -0.039 | -0.008 |
| 7040 | 0.022 | -0.022 |
| 7041 | -0.089 | 0.099 |
| 7042 | 0.019 | -0.078 |
| 7043 | -0.039 | -0.030 |
| 7044 | 0.007 | -0.078 |
| 7045 | -0.008 | -0.039 |
| 7046 | -0.026 | 0.011 |
| 7047 | 0.019 | 0.156 |
| 7048 | -0.025 | -0.068 |
| 7049 | 0.017 | -0.172 |
| 7050 | -0.062 | -0.026 |
| 7051 | -0.054 | -0.041 |
| 7052 | -0.020 | 0.080 |
| 7053 | -0.033 | -0.012 |
| 7054 | -0.043 | -0.017 |
| 7055 | 0.017 | 0.039 |
| 7056 | 0.065 | 0.131 |
| 7057 | 0.024 | -0.067 |
| 7058 | -0.048 | -0.064 |
| 7059 | 0.017 | -0.084 |
| 7060 | -0.050 | -0.054 |
| 7061 | 0.061 | 0.118 |
| 7062 | -0.010 | 0.115 |
| 7063 | 0.007 | -0.009 |
| 7064 | -0.034 | -0.009 |
| 7065 | 0.052 | -0.010 |
| 7066 | -0.033 | 0.133 |
| 7067 | 0.019 | -0.023 |
| 7068 | -0.090 | 0.028 |
| 7069 | -0.023 | -0.043 |
| 7070 | 0.032 | 0.025 |
| 7071 | -0.057 | 0.017 |
| 7072 | -0.003 | 0.000 |
| 7073 | -0.031 | -0.027 |
| 7074 | 0.017 | -0.034 |
| 7075 | -0.084 | 0.104 |
| 7076 | -0.006 | 0.027 |
| 7077 | 0.020 | 0.164 |
| 7078 | -0.045 | -0.108 |
| 7079 | -0.054 | -0.062 |
| 7080 | 0.050 | -0.096 |
| 7081 | 0.040 | -0.058 |
| 7082 | -0.025 | -0.024 |

| | | |
|---|---|---|
| 7083 | -0.050 | -0.051 |
| 7084 | -0.019 | -0.037 |
| 7085 | -0.028 | 0.075 |
| 7086 | -0.017 | 0.001 |
| 7087 | 0.036 | 0.020 |
| 7088 | 0.034 | -0.003 |
| 7089 | 0.019 | -0.153 |
| 7090 | -0.002 | -0.074 |
| 7091 | -0.047 | -0.016 |
| 7092 | -0.004 | -0.103 |
| 7093 | 0.034 | 0.009 |
| 7094 | 0.025 | -0.004 |
| 7095 | -0.053 | -0.088 |
| 7096 | -0.012 | -0.135 |
| 7097 | -0.049 | -0.039 |
| 7098 | -0.013 | 0.008 |
| 7099 | 0.037 | 0.018 |
| 7100 | -0.009 | 0.015 |
| 7101 | 0.015 | -0.001 |
| 7102 | 0.025 | 0.025 |
| 7103 | -0.057 | 0.007 |
| 7104 | -0.026 | 0.064 |
| 7105 | -0.047 | -0.002 |
| 7106 | 0.000 | -0.070 |
| 7107 | -0.060 | -0.064 |
| 7108 | 0.000 | 0.018 |
| 7109 | 0.003 | -0.046 |
| 7110 | -0.003 | -0.013 |
| 7111 | -0.056 | -0.072 |
| 7112 | 0.009 | 0.054 |
| 7113 | 0.034 | 0.107 |
| 7114 | 0.035 | 0.021 |
| 7115 | 0.074 | -0.021 |
| 7116 | -0.040 | -0.042 |
| 7117 | -0.007 | 0.071 |
| 7118 | -0.013 | 0.016 |
| 7119 | -0.003 | 0.088 |
| 7120 | 0.002 | -0.032 |
| 7121 | 0.019 | -0.004 |
| 7122 | 0.012 | 0.002 |
| 7123 | 0.077 | -0.133 |
| 7124 | -0.028 | 0.084 |
| 7125 | -0.040 | 0.040 |
| 7126 | -0.086 | 0.044 |
| 7127 | -0.007 | 0.074 |
| 7128 | 0.053 | 0.030 |
| 7129 | -0.007 | -0.103 |
| 7130 | -0.035 | -0.182 |
| 7131 | 0.010 | -0.111 |
| 7132 | 0.049 | -0.030 |
| 7133 | 0.013 | -0.042 |

| | | |
|---|---|---|
| 7134 | -0.013 | -0.026 |
| 7135 | -0.044 | -0.134 |
| 7136 | 0.014 | 0.035 |
| 7137 | 0.036 | -0.005 |
| 7138 | -0.016 | 0.004 |
| 7139 | 0.035 | 0.188 |
| 7140 | 0.044 | -0.092 |
| 7141 | 0.017 | -0.003 |
| 7142 | -0.008 | 0.018 |
| 7143 | 0.049 | 0.018 |
| 7144 | -0.133 | 0.065 |
| 7145 | -0.019 | 0.003 |
| 7146 | 0.012 | -0.068 |
| 7147 | -0.022 | -0.033 |
| 7148 | 0.029 | -0.017 |
| 7149 | -0.067 | -0.062 |
| 7150 | -0.016 | -0.125 |
| 7151 | 0.013 | -0.060 |
| 7152 | -0.022 | -0.045 |
| 7153 | -0.080 | 0.049 |
| 7154 | -0.043 | -0.063 |
| 7155 | 0.085 | -0.039 |
| 7156 | 0.004 | 0.018 |
| 7157 | -0.041 | 0.005 |
| 7158 | 0.005 | 0.129 |
| 7159 | 0.011 | -0.021 |
| 7160 | -0.043 | 0.102 |
| 7161 | -0.017 | -0.060 |
| 7162 | 0.006 | -0.002 |
| 7163 | 0.012 | 0.026 |
| 7164 | 0.034 | -0.049 |
| 7165 | -0.044 | -0.106 |
| 7166 | 0.062 | -0.044 |
| 7167 | 0.038 | 0.115 |
| 7168 | -0.032 | 0.078 |
| 7169 | 0.030 | 0.044 |
| 7170 | 0.054 | 0.013 |
| 7171 | 0.018 | -0.130 |
| 7172 | 0.033 | 0.012 |
| 7173 | -0.050 | 0.157 |
| 7174 | 0.071 | -0.118 |
| 7175 | 0.037 | 0.016 |
| 7176 | 0.022 | -0.014 |
| 7177 | 0.013 | -0.011 |
| 7178 | 0.062 | 0.082 |
| 7179 | -0.088 | 0.081 |
| 7180 | 0.044 | -0.095 |
| 7181 | 0.001 | 0.007 |
| 7182 | 0.016 | 0.105 |
| 7183 | 0.040 | -0.158 |
| 7184 | 0.002 | -0.009 |

| | | |
|---|---|---|
| 7185 | -0.075 | -0.051 |
| 7186 | 0.011 | -0.081 |
| 7187 | -0.045 | -0.091 |
| 7188 | -0.033 | 0.033 |
| 7189 | -0.010 | 0.030 |
| 7190 | -0.019 | 0.019 |
| 7191 | -0.025 | 0.028 |
| 7192 | 0.047 | 0.017 |
| 7193 | 0.006 | 0.059 |
| 7194 | -0.025 | -0.009 |
| 7195 | -0.011 | 0.063 |
| 7196 | 0.006 | -0.010 |
| 7197 | 0.006 | 0.050 |
| 7198 | -0.035 | -0.032 |
| 7199 | -0.068 | -0.032 |
| 7200 | 0.052 | 0.078 |
| 7201 | -0.028 | -0.053 |
| 7202 | 0.046 | 0.001 |
| 7203 | 0.005 | 0.099 |
| 7204 | 0.117 | -0.014 |
| 7205 | 0.016 | -0.042 |
| 7206 | -0.024 | 0.058 |
| 7207 | 0.013 | 0.071 |
| 7208 | 0.058 | -0.037 |
| 7209 | -0.036 | -0.022 |
| 7210 | -0.069 | -0.005 |
| 7211 | -0.110 | -0.019 |
| 7212 | 0.052 | 0.036 |
| 7213 | 0.016 | -0.152 |
| 7214 | 0.028 | -0.001 |
| 7215 | -0.034 | -0.053 |
| 7216 | -0.011 | -0.084 |
| 7217 | 0.033 | -0.021 |
| 7218 | 0.083 | 0.071 |
| 7219 | 0.100 | -0.111 |
| 7220 | -0.026 | -0.086 |
| 7221 | 0.017 | 0.132 |
| 7222 | 0.026 | -0.040 |
| 7223 | 0.013 | -0.047 |
| 7224 | 0.025 | -0.086 |
| 7225 | -0.037 | 0.116 |
| 7226 | 0.024 | 0.001 |
| 7227 | 0.016 | 0.051 |
| 7228 | -0.035 | -0.043 |
| 7229 | 0.028 | 0.115 |
| 7230 | -0.065 | 0.136 |
| 7231 | 0.037 | 0.054 |
| 7232 | -0.052 | -0.002 |
| 7233 | 0.017 | -0.027 |
| 7234 | -0.033 | 0.020 |
| 7235 | 0.030 | -0.007 |

| | | |
|---|---|---|
| 7236 | -0.015 | -0.021 |
| 7237 | -0.047 | -0.077 |
| 7238 | -0.031 | 0.006 |
| 7239 | 0.025 | -0.029 |
| 7240 | -0.041 | 0.001 |
| 7241 | -0.073 | 0.035 |
| 7242 | -0.052 | 0.094 |
| 7243 | -0.046 | 0.010 |
| 7244 | -0.011 | -0.017 |
| 7245 | -0.008 | 0.084 |
| 7246 | 0.004 | -0.030 |
| 7247 | -0.043 | 0.036 |
| 7248 | 0.014 | 0.047 |
| 7249 | -0.057 | -0.034 |
| 7250 | 0.038 | -0.053 |
| 7251 | -0.091 | -0.018 |
| 7252 | 0.022 | 0.008 |
| 7253 | 0.007 | -0.103 |
| 7254 | 0.013 | 0.080 |
| 7255 | -0.034 | 0.019 |
| 7256 | 0.019 | 0.023 |
| 7257 | -0.018 | 0.063 |
| 7258 | 0.023 | 0.041 |
| 7259 | 0.016 | -0.018 |
| 7260 | -0.035 | -0.015 |
| 7261 | 0.010 | 0.101 |
| 7262 | -0.039 | -0.057 |
| 7263 | 0.055 | -0.059 |
| 7264 | -0.014 | -0.022 |
| 7265 | 0.003 | 0.008 |
| 7266 | -0.012 | -0.106 |
| 7267 | -0.006 | -0.047 |
| 7268 | 0.057 | -0.004 |
| 7269 | 0.044 | 0.080 |
| 7270 | -0.003 | -0.006 |
| 7271 | -0.011 | -0.049 |
| 7272 | 0.008 | -0.062 |
| 7273 | -0.002 | -0.051 |
| 7274 | 0.055 | 0.056 |
| 7275 | -0.043 | 0.041 |
| 7276 | 0.010 | 0.174 |
| 7277 | -0.009 | -0.068 |
| 7278 | 0.041 | 0.070 |
| 7279 | -0.014 | 0.047 |
| 7280 | 0.016 | 0.036 |
| 7281 | 0.040 | -0.010 |
| 7282 | -0.003 | 0.065 |
| 7283 | 0.022 | -0.027 |
| 7284 | 0.034 | 0.041 |
| 7285 | -0.021 | 0.033 |
| 7286 | -0.013 | -0.096 |

| | | |
|---|---|---|
| 7287 | -0.011 | 0.179 |
| 7288 | -0.053 | 0.002 |
| 7289 | -0.026 | 0.087 |
| 7290 | 0.009 | -0.132 |
| 7291 | -0.018 | -0.002 |
| 7292 | 0.019 | 0.050 |
| 7293 | -0.020 | -0.072 |
| 7294 | -0.032 | -0.005 |
| 7295 | 0.003 | -0.038 |
| 7296 | -0.026 | 0.125 |
| 7297 | -0.032 | -0.041 |
| 7298 | 0.011 | -0.040 |
| 7299 | 0.028 | 0.080 |
| 7300 | 0.014 | -0.064 |
| 7301 | -0.032 | 0.022 |
| 7302 | 0.040 | 0.022 |
| 7303 | 0.013 | -0.093 |
| 7304 | 0.012 | -0.026 |
| 7305 | -0.071 | -0.037 |
| 7306 | -0.014 | 0.043 |
| 7307 | -0.009 | -0.012 |
| 7308 | -0.023 | 0.013 |
| 7309 | 0.006 | -0.018 |
| 7310 | -0.012 | 0.099 |
| 7311 | 0.029 | 0.021 |
| 7312 | 0.021 | 0.059 |
| 7313 | 0.060 | -0.047 |
| 7314 | 0.020 | -0.134 |
| 7315 | -0.008 | -0.054 |
| 7316 | 0.058 | -0.026 |
| 7317 | 0.004 | -0.012 |
| 7318 | 0.019 | -0.037 |
| 7319 | -0.028 | 0.156 |
| 7320 | 0.028 | 0.057 |
| 7321 | -0.012 | 0.055 |
| 7322 | -0.007 | -0.053 |
| 7323 | 0.050 | 0.022 |
| 7324 | -0.046 | -0.064 |
| 7325 | -0.058 | -0.130 |
| 7326 | -0.048 | 0.002 |
| 7327 | 0.064 | 0.053 |
| 7328 | 0.064 | 0.077 |
| 7329 | 0.056 | -0.042 |
| 7330 | 0.006 | -0.055 |
| 7331 | -0.003 | 0.072 |
| 7332 | 0.019 | 0.017 |
| 7333 | -0.019 | -0.011 |
| 7334 | -0.005 | -0.031 |
| 7335 | 0.060 | 0.031 |
| 7336 | -0.020 | 0.094 |
| 7337 | -0.014 | 0.059 |

| | | |
|---|---|---|
| 7338 | 0.019 | -0.062 |
| 7339 | -0.043 | -0.029 |
| 7340 | 0.006 | 0.062 |
| 7341 | -0.032 | -0.049 |
| 7342 | 0.017 | -0.063 |
| 7343 | -0.058 | -0.077 |
| 7344 | 0.023 | 0.059 |
| 7345 | 0.008 | 0.037 |
| 7346 | 0.003 | 0.008 |
| 7347 | 0.014 | 0.089 |
| 7348 | 0.006 | 0.106 |
| 7349 | 0.024 | -0.027 |
| 7350 | -0.064 | 0.012 |
| 7351 | -0.017 | 0.096 |
| 7352 | 0.083 | -0.076 |
| 7353 | 0.004 | -0.094 |
| 7354 | -0.070 | 0.001 |
| 7355 | -0.023 | -0.088 |
| 7356 | -0.017 | 0.081 |
| 7357 | 0.023 | 0.030 |
| 7358 | -0.018 | 0.057 |
| 7359 | 0.010 | 0.029 |
| 7360 | 0.013 | -0.026 |
| 7361 | -0.046 | 0.077 |
| 7362 | -0.011 | 0.066 |
| 7363 | 0.001 | 0.121 |
| 7364 | 0.024 | 0.052 |
| 7365 | -0.045 | -0.032 |
| 7366 | 0.010 | 0.154 |
| 7367 | 0.094 | 0.072 |
| 7368 | -0.042 | 0.119 |
| 7369 | 0.021 | -0.020 |
| 7370 | -0.018 | -0.059 |
| 7371 | 0.006 | -0.003 |
| 7372 | 0.026 | 0.259 |
| 7373 | 0.017 | -0.026 |
| 7374 | -0.004 | 0.013 |
| 7375 | 0.022 | -0.036 |
| 7376 | 0.003 | -0.010 |
| 7377 | -0.003 | -0.112 |
| 7378 | -0.054 | -0.130 |
| 7379 | -0.018 | 0.098 |
| 7380 | -0.023 | 0.133 |
| 7381 | -0.050 | 0.131 |
| 7382 | -0.024 | 0.002 |
| 7383 | 0.016 | 0.008 |
| 7384 | 0.017 | 0.078 |
| 7385 | 0.037 | -0.103 |
| 7386 | -0.002 | 0.003 |
| 7387 | -0.007 | -0.035 |
| 7388 | 0.056 | -0.083 |

| | | |
|---|---|---|
| 7389 | 0.063 | 0.058 |
| 7390 | -0.001 | -0.063 |
| 7391 | 0.047 | -0.053 |
| 7392 | -0.014 | 0.093 |
| 7393 | -0.017 | -0.088 |
| 7394 | -0.036 | 0.119 |
| 7395 | -0.004 | -0.005 |
| 7396 | 0.016 | -0.013 |
| 7397 | -0.066 | -0.078 |
| 7398 | -0.003 | 0.046 |
| 7399 | 0.044 | -0.010 |
| 7400 | -0.050 | 0.061 |
| 7401 | -0.015 | -0.046 |
| 7402 | -0.020 | 0.072 |
| 7403 | -0.006 | -0.040 |
| 7404 | 0.030 | -0.054 |
| 7405 | -0.022 | 0.020 |
| 7406 | -0.026 | -0.007 |
| 7407 | -0.003 | 0.011 |
| 7408 | 0.022 | 0.086 |
| 7409 | -0.024 | -0.028 |
| 7410 | 0.017 | 0.114 |
| 7411 | 0.088 | 0.061 |
| 7412 | 0.036 | -0.127 |
| 7413 | -0.029 | 0.096 |
| 7414 | 0.001 | 0.047 |
| 7415 | 0.003 | -0.018 |
| 7416 | -0.033 | -0.009 |
| 7417 | 0.068 | 0.072 |
| 7418 | -0.078 | 0.080 |
| 7419 | -0.005 | -0.041 |
| 7420 | -0.024 | 0.007 |
| 7421 | 0.014 | 0.012 |
| 7422 | -0.035 | 0.029 |
| 7423 | 0.004 | -0.131 |
| 7424 | 0.050 | 0.176 |
| 7425 | -0.052 | 0.065 |
| 7426 | 0.002 | 0.036 |
| 7427 | 0.017 | -0.063 |
| 7428 | -0.005 | 0.012 |
| 7429 | 0.003 | 0.046 |
| 7430 | 0.009 | 0.071 |
| 7431 | -0.039 | -0.103 |
| 7432 | 0.004 | -0.021 |
| 7433 | -0.022 | -0.041 |
| 7434 | -0.057 | -0.072 |
| 7435 | 0.001 | 0.107 |
| 7436 | -0.013 | 0.069 |
| 7437 | -0.003 | -0.042 |
| 7438 | 0.013 | -0.049 |
| 7439 | 0.003 | -0.106 |

| | | |
|---|---|---|
| 7440 | -0.050 | -0.037 |
| 7441 | -0.026 | -0.009 |
| 7442 | -0.011 | -0.118 |
| 7443 | 0.071 | 0.047 |
| 7444 | -0.021 | 0.065 |
| 7445 | -0.018 | -0.025 |
| 7446 | -0.006 | -0.009 |
| 7447 | -0.018 | -0.032 |
| 7448 | -0.034 | 0.074 |
| 7449 | -0.053 | -0.007 |
| 7450 | -0.056 | 0.103 |
| 7451 | 0.000 | -0.013 |
| 7452 | 0.002 | -0.082 |
| 7453 | 0.013 | 0.182 |
| 7454 | 0.057 | 0.028 |
| 7455 | -0.040 | -0.054 |
| 7456 | -0.021 | 0.145 |
| 7457 | 0.051 | -0.003 |
| 7458 | -0.062 | 0.081 |
| 7459 | -0.030 | 0.030 |
| 7460 | -0.043 | -0.115 |
| 7461 | 0.061 | 0.080 |
| 7462 | -0.035 | 0.085 |
| 7463 | -0.021 | 0.005 |
| 7464 | 0.000 | 0.032 |
| 7465 | 0.017 | -0.226 |
| 7466 | -0.046 | -0.077 |
| 7467 | 0.004 | -0.055 |
| 7468 | -0.074 | 0.113 |
| 7469 | -0.020 | -0.036 |
| 7470 | 0.044 | -0.132 |
| 7471 | -0.033 | 0.066 |
| 7472 | 0.020 | -0.022 |
| 7473 | 0.003 | 0.128 |
| 7474 | -0.037 | -0.018 |
| 7475 | 0.009 | -0.019 |
| 7476 | 0.010 | 0.107 |
| 7477 | 0.024 | 0.040 |
| 7478 | 0.008 | 0.029 |
| 7479 | -0.004 | 0.083 |
| 7480 | -0.076 | -0.003 |
| 7481 | 0.012 | 0.117 |
| 7482 | 0.010 | -0.039 |
| 7483 | 0.017 | 0.071 |
| 7484 | 0.039 | -0.118 |
| 7485 | -0.083 | 0.032 |
| 7486 | -0.018 | -0.043 |
| 7487 | -0.027 | -0.054 |
| 7488 | 0.035 | -0.082 |
| 7489 | 0.128 | -0.015 |
| 7490 | -0.078 | 0.126 |

| | | |
|---|---|---|
| 7491 | -0.038 | -0.079 |
| 7492 | 0.024 | -0.026 |
| 7493 | -0.027 | 0.059 |
| 7494 | 0.035 | 0.085 |
| 7495 | -0.013 | -0.181 |
| 7496 | 0.001 | 0.174 |
| 7497 | -0.021 | 0.006 |
| 7498 | -0.028 | -0.140 |
| 7499 | -0.023 | 0.053 |
| 7500 | 0.046 | 0.100 |
| 7501 | 0.011 | 0.056 |
| 7502 | -0.006 | -0.044 |
| 7503 | 0.022 | -0.034 |
| 7504 | -0.013 | -0.017 |
| 7505 | -0.008 | -0.079 |
| 7506 | 0.056 | -0.022 |
| 7507 | -0.005 | 0.065 |
| 7508 | -0.035 | 0.046 |
| 7509 | -0.082 | -0.011 |
| 7510 | -0.068 | -0.079 |
| 7511 | 0.070 | -0.126 |
| 7512 | 0.018 | -0.006 |
| 7513 | 0.013 | 0.050 |
| 7514 | 0.031 | 0.050 |
| 7515 | 0.020 | -0.075 |
| 7516 | -0.043 | -0.003 |
| 7517 | -0.034 | 0.054 |
| 7518 | -0.073 | 0.052 |
| 7519 | -0.045 | -0.042 |
| 7520 | 0.002 | -0.029 |
| 7521 | 0.014 | 0.039 |
| 7522 | -0.021 | 0.047 |
| 7523 | 0.014 | -0.005 |
| 7524 | -0.010 | -0.055 |
| 7525 | -0.025 | -0.004 |
| 7526 | 0.020 | 0.045 |
| 7527 | -0.017 | -0.049 |
| 7528 | -0.020 | -0.076 |
| 7529 | 0.053 | 0.133 |
| 7530 | 0.005 | -0.050 |
| 7531 | -0.017 | 0.001 |
| 7532 | -0.029 | -0.074 |
| 7533 | -0.013 | -0.025 |
| 7534 | 0.021 | 0.102 |
| 7535 | -0.059 | 0.015 |
| 7536 | 0.002 | 0.167 |
| 7537 | 0.003 | 0.125 |
| 7538 | 0.057 | -0.037 |
| 7539 | -0.043 | 0.040 |
| 7540 | 0.027 | -0.056 |
| 7541 | 0.014 | 0.006 |

| | | |
|---|---|---|
| 7542 | -0.060 | -0.097 |
| 7543 | 0.027 | -0.153 |
| 7544 | 0.064 | -0.119 |
| 7545 | 0.021 | -0.060 |
| 7546 | -0.026 | -0.024 |
| 7547 | 0.030 | -0.109 |
| 7548 | 0.023 | -0.072 |
| 7549 | -0.020 | -0.123 |
| 7550 | -0.001 | -0.035 |
| 7551 | -0.029 | 0.029 |
| 7552 | -0.072 | 0.018 |
| 7553 | 0.007 | -0.085 |
| 7554 | -0.049 | 0.008 |
| 7555 | 0.001 | -0.123 |
| 7556 | 0.068 | 0.146 |
| 7557 | -0.004 | -0.073 |
| 7558 | 0.012 | -0.087 |
| 7559 | 0.041 | -0.070 |
| 7560 | 0.028 | -0.034 |
| 7561 | 0.001 | -0.108 |
| 7562 | -0.003 | 0.139 |
| 7563 | 0.042 | -0.109 |
| 7564 | -0.043 | -0.125 |
| 7565 | 0.039 | -0.007 |
| 7566 | 0.021 | 0.028 |
| 7567 | -0.046 | 0.000 |
| 7568 | -0.069 | -0.060 |
| 7569 | -0.001 | 0.002 |
| 7570 | 0.019 | 0.011 |
| 7571 | -0.006 | 0.068 |
| 7572 | 0.031 | -0.042 |
| 7573 | 0.031 | -0.041 |
| 7574 | -0.032 | 0.012 |
| 7575 | -0.013 | -0.064 |
| 7576 | -0.015 | 0.094 |
| 7577 | 0.032 | 0.054 |
| 7578 | -0.006 | -0.164 |
| 7579 | -0.006 | 0.103 |
| 7580 | -0.044 | -0.137 |
| 7581 | -0.085 | 0.001 |
| 7582 | -0.002 | 0.038 |
| 7583 | 0.026 | -0.044 |
| 7584 | 0.042 | -0.082 |
| 7585 | 0.052 | -0.031 |
| 7586 | 0.018 | -0.138 |
| 7587 | 0.024 | -0.028 |
| 7588 | -0.014 | 0.035 |
| 7589 | -0.011 | -0.033 |
| 7590 | 0.017 | 0.017 |
| 7591 | -0.026 | -0.046 |
| 7592 | 0.049 | 0.063 |

| | | |
|---|---|---|
| 7593 | 0.027 | 0.061 |
| 7594 | 0.023 | -0.026 |
| 7595 | 0.044 | 0.108 |
| 7596 | 0.005 | -0.002 |
| 7597 | 0.032 | -0.120 |
| 7598 | 0.024 | 0.014 |
| 7599 | 0.035 | 0.138 |
| 7600 | -0.072 | 0.006 |
| 7601 | 0.010 | 0.057 |
| 7602 | 0.003 | 0.004 |
| 7603 | 0.011 | -0.020 |
| 7604 | 0.009 | 0.000 |
| 7605 | 0.023 | -0.023 |
| 7606 | -0.034 | 0.032 |
| 7607 | 0.046 | 0.050 |
| 7608 | -0.065 | -0.017 |
| 7609 | -0.008 | 0.060 |
| 7610 | 0.017 | -0.037 |
| 7611 | -0.023 | -0.043 |
| 7612 | -0.061 | 0.036 |
| 7613 | 0.013 | 0.028 |
| 7614 | -0.032 | 0.069 |
| 7615 | -0.005 | -0.027 |
| 7616 | 0.013 | -0.017 |
| 7617 | -0.015 | -0.079 |
| 7618 | 0.000 | 0.016 |
| 7619 | -0.012 | 0.044 |
| 7620 | 0.021 | 0.032 |
| 7621 | 0.013 | 0.054 |
| 7622 | 0.074 | 0.042 |
| 7623 | 0.000 | -0.049 |
| 7624 | -0.021 | 0.048 |
| 7625 | 0.024 | 0.095 |
| 7626 | 0.037 | 0.077 |
| 7627 | 0.007 | 0.096 |
| 7628 | -0.069 | 0.203 |
| 7629 | 0.040 | 0.107 |
| 7630 | 0.031 | -0.033 |
| 7631 | 0.048 | 0.105 |
| 7632 | -0.009 | -0.036 |
| 7633 | -0.032 | 0.041 |
| 7634 | -0.044 | -0.048 |
| 7635 | 0.004 | -0.015 |
| 7636 | -0.010 | 0.028 |
| 7637 | -0.022 | -0.048 |
| 7638 | -0.002 | 0.167 |
| 7639 | 0.008 | -0.057 |
| 7640 | -0.004 | -0.033 |
| 7641 | 0.023 | -0.011 |
| 7642 | -0.025 | 0.021 |
| 7643 | -0.013 | -0.072 |

| | | |
|---|---|---|
| 7644 | -0.057 | -0.046 |
| 7645 | -0.001 | 0.027 |
| 7646 | -0.003 | 0.129 |
| 7647 | 0.048 | -0.005 |
| 7648 | 0.000 | -0.154 |
| 7649 | 0.003 | 0.036 |
| 7650 | -0.027 | -0.015 |
| 7651 | -0.059 | -0.080 |
| 7652 | -0.020 | -0.097 |
| 7653 | -0.008 | 0.040 |
| 7654 | 0.008 | 0.065 |
| 7655 | -0.018 | 0.069 |
| 7656 | 0.000 | 0.132 |
| 7657 | -0.054 | 0.053 |
| 7658 | 0.046 | -0.012 |
| 7659 | -0.037 | -0.018 |
| 7660 | 0.018 | 0.022 |
| 7661 | -0.036 | 0.014 |
| 7662 | -0.046 | 0.170 |
| 7663 | 0.009 | 0.019 |
| 7664 | -0.033 | -0.048 |
| 7665 | 0.031 | -0.131 |
| 7666 | 0.030 | -0.051 |
| 7667 | 0.055 | 0.117 |
| 7668 | -0.006 | -0.045 |
| 7669 | 0.040 | -0.133 |
| 7670 | -0.001 | -0.022 |
| 7671 | -0.021 | -0.032 |
| 7672 | -0.001 | -0.107 |
| 7673 | 0.004 | -0.133 |
| 7674 | -0.015 | 0.041 |
| 7675 | -0.002 | -0.106 |
| 7676 | 0.039 | -0.007 |
| 7677 | -0.011 | 0.023 |
| 7678 | 0.006 | 0.030 |
| 7679 | -0.026 | -0.026 |
| 7680 | 0.081 | 0.047 |
| 7681 | -0.028 | -0.032 |
| 7682 | -0.050 | 0.072 |
| 7683 | 0.031 | 0.013 |
| 7684 | -0.074 | -0.026 |
| 7685 | -0.020 | 0.126 |
| 7686 | 0.010 | -0.061 |
| 7687 | -0.031 | 0.060 |
| 7688 | -0.016 | -0.065 |
| 7689 | -0.043 | 0.069 |
| 7690 | -0.026 | -0.045 |
| 7691 | -0.045 | -0.104 |
| 7692 | -0.022 | 0.027 |
| 7693 | 0.005 | -0.055 |
| 7694 | -0.025 | 0.001 |

| | | |
|---|---|---|
| 7695 | 0.035 | -0.019 |
| 7696 | 0.046 | -0.067 |
| 7697 | 0.028 | 0.068 |
| 7698 | 0.022 | -0.114 |
| 7699 | 0.023 | 0.100 |
| 7700 | 0.035 | 0.067 |
| 7701 | -0.056 | -0.021 |
| 7702 | 0.039 | -0.030 |
| 7703 | 0.002 | 0.104 |
| 7704 | 0.015 | 0.010 |
| 7705 | -0.067 | 0.075 |
| 7706 | -0.016 | -0.150 |
| 7707 | -0.010 | 0.052 |
| 7708 | 0.007 | -0.020 |
| 7709 | 0.032 | 0.081 |
| 7710 | -0.061 | -0.078 |
| 7711 | 0.040 | -0.125 |
| 7712 | 0.042 | 0.059 |
| 7713 | 0.004 | 0.000 |
| 7714 | -0.034 | -0.004 |
| 7715 | -0.003 | -0.039 |
| 7716 | 0.022 | -0.003 |
| 7717 | 0.022 | -0.022 |
| 7718 | -0.014 | 0.093 |
| 7719 | 0.049 | 0.109 |
| 7720 | -0.017 | 0.112 |
| 7721 | 0.061 | 0.009 |
| 7722 | 0.040 | -0.008 |
| 7723 | -0.037 | -0.034 |
| 7724 | -0.081 | 0.056 |
| 7725 | -0.048 | -0.035 |
| 7726 | 0.013 | -0.091 |
| 7727 | 0.012 | 0.044 |
| 7728 | 0.043 | 0.014 |
| 7729 | -0.019 | -0.051 |
| 7730 | -0.034 | 0.020 |
| 7731 | -0.021 | -0.035 |
| 7732 | -0.002 | 0.056 |
| 7733 | 0.024 | 0.086 |
| 7734 | 0.013 | -0.057 |
| 7735 | 0.041 | 0.171 |
| 7736 | 0.052 | 0.123 |
| 7737 | -0.027 | -0.059 |
| 7738 | -0.089 | 0.001 |
| 7739 | -0.042 | 0.079 |
| 7740 | -0.013 | -0.083 |
| 7741 | 0.038 | 0.031 |
| 7742 | 0.018 | 0.091 |
| 7743 | -0.020 | -0.092 |
| 7744 | 0.024 | -0.008 |
| 7745 | 0.081 | 0.078 |

| | | |
|---|---|---|
| 7746 | 0.036 | -0.029 |
| 7747 | -0.096 | -0.088 |
| 7748 | 0.004 | -0.225 |
| 7749 | 0.031 | 0.166 |
| 7750 | -0.037 | 0.040 |
| 7751 | -0.017 | 0.027 |
| 7752 | -0.015 | -0.056 |
| 7753 | 0.053 | 0.006 |
| 7754 | 0.037 | 0.003 |
| 7755 | 0.011 | -0.019 |
| 7756 | 0.075 | -0.034 |
| 7757 | 0.006 | -0.123 |
| 7758 | 0.029 | -0.002 |
| 7759 | -0.053 | -0.065 |
| 7760 | 0.030 | 0.047 |
| 7761 | -0.011 | -0.099 |
| 7762 | -0.008 | -0.018 |
| 7763 | 0.064 | -0.049 |
| 7764 | 0.038 | -0.039 |
| 7765 | 0.030 | 0.027 |
| 7766 | -0.036 | -0.030 |
| 7767 | 0.026 | 0.098 |
| 7768 | -0.026 | -0.087 |
| 7769 | 0.015 | 0.029 |
| 7770 | 0.049 | 0.023 |
| 7771 | -0.010 | 0.083 |
| 7772 | -0.003 | 0.008 |
| 7773 | -0.033 | 0.136 |
| 7774 | 0.057 | 0.081 |
| 7775 | 0.034 | 0.014 |
| 7776 | 0.114 | 0.111 |
| 7777 | 0.054 | 0.054 |
| 7778 | -0.062 | 0.138 |
| 7779 | -0.029 | 0.043 |
| 7780 | 0.039 | 0.015 |
| 7781 | 0.052 | 0.147 |
| 7782 | -0.004 | -0.170 |
| 7783 | 0.059 | 0.042 |
| 7784 | 0.005 | -0.035 |
| 7785 | -0.016 | -0.021 |
| 7786 | -0.042 | -0.061 |
| 7787 | 0.021 | 0.032 |
| 7788 | 0.013 | 0.008 |
| 7789 | 0.048 | 0.068 |
| 7790 | 0.051 | 0.125 |
| 7791 | -0.009 | 0.023 |
| 7792 | 0.015 | 0.051 |
| 7793 | -0.062 | -0.070 |
| 7794 | -0.052 | 0.061 |
| 7795 | 0.016 | -0.107 |
| 7796 | 0.039 | 0.011 |

| | | |
|---|---|---|
| 7797 | 0.069 | 0.124 |
| 7798 | 0.041 | 0.105 |
| 7799 | -0.044 | 0.084 |
| 7800 | -0.029 | -0.011 |
| 7801 | 0.011 | 0.033 |
| 7802 | 0.020 | -0.025 |
| 7803 | 0.040 | -0.012 |
| 7804 | 0.001 | -0.020 |
| 7805 | -0.006 | -0.019 |
| 7806 | -0.020 | -0.021 |
| 7807 | -0.020 | 0.016 |
| 7808 | -0.025 | -0.066 |
| 7809 | -0.031 | -0.003 |
| 7810 | -0.042 | -0.080 |
| 7811 | -0.027 | 0.097 |
| 7812 | -0.020 | -0.032 |
| 7813 | 0.016 | 0.101 |
| 7814 | 0.004 | 0.030 |
| 7815 | 0.020 | 0.063 |
| 7816 | -0.005 | -0.004 |
| 7817 | 0.017 | 0.069 |
| 7818 | 0.012 | 0.024 |
| 7819 | 0.025 | -0.013 |
| 7820 | -0.031 | -0.053 |
| 7821 | 0.028 | 0.047 |
| 7822 | 0.044 | -0.053 |
| 7823 | 0.027 | 0.099 |
| 7824 | -0.035 | -0.046 |
| 7825 | 0.004 | -0.018 |
| 7826 | 0.056 | -0.027 |
| 7827 | -0.028 | -0.018 |
| 7828 | -0.077 | 0.008 |
| 7829 | 0.006 | 0.001 |
| 7830 | -0.045 | 0.081 |
| 7831 | -0.050 | -0.033 |
| 7832 | -0.018 | -0.007 |
| 7833 | -0.011 | -0.037 |
| 7834 | -0.023 | -0.140 |
| 7835 | 0.005 | -0.020 |
| 7836 | -0.010 | -0.085 |
| 7837 | -0.046 | 0.080 |
| 7838 | 0.001 | 0.030 |
| 7839 | 0.019 | -0.046 |
| 7840 | 0.023 | -0.053 |
| 7841 | 0.050 | 0.031 |
| 7842 | 0.089 | 0.023 |
| 7843 | 0.012 | 0.084 |
| 7844 | -0.009 | 0.026 |
| 7845 | -0.023 | 0.024 |
| 7846 | -0.001 | 0.174 |
| 7847 | -0.064 | 0.050 |

| | | |
|---|---|---|
| 7848 | -0.059 | -0.005 |
| 7849 | 0.025 | 0.101 |
| 7850 | 0.065 | 0.034 |
| 7851 | 0.012 | 0.040 |
| 7852 | -0.047 | -0.064 |
| 7853 | 0.003 | 0.047 |
| 7854 | -0.035 | -0.033 |
| 7855 | -0.065 | -0.034 |
| 7856 | 0.034 | 0.093 |
| 7857 | -0.012 | -0.024 |
| 7858 | -0.023 | -0.006 |
| 7859 | -0.092 | 0.092 |
| 7860 | -0.033 | -0.117 |
| 7861 | 0.001 | -0.097 |
| 7862 | -0.013 | 0.055 |
| 7863 | -0.030 | -0.120 |
| 7864 | -0.031 | 0.078 |
| 7865 | -0.040 | -0.128 |
| 7866 | -0.048 | -0.005 |
| 7867 | 0.004 | 0.088 |
| 7868 | -0.032 | -0.059 |
| 7869 | -0.037 | 0.057 |
| 7870 | 0.002 | 0.016 |
| 7871 | 0.014 | -0.038 |
| 7872 | -0.020 | 0.035 |
| 7873 | 0.026 | 0.041 |
| 7874 | 0.020 | 0.087 |
| 7875 | -0.044 | 0.058 |
| 7876 | 0.016 | 0.016 |
| 7877 | 0.016 | 0.040 |
| 7878 | 0.006 | -0.013 |
| 7879 | -0.021 | 0.156 |
| 7880 | -0.052 | 0.032 |
| 7881 | 0.014 | 0.063 |
| 7882 | 0.037 | 0.044 |
| 7883 | -0.007 | 0.010 |
| 7884 | 0.064 | -0.001 |
| 7885 | 0.084 | 0.050 |
| 7886 | -0.012 | -0.031 |
| 7887 | -0.029 | 0.008 |
| 7888 | -0.042 | 0.002 |
| 7889 | 0.064 | 0.072 |
| 7890 | -0.022 | -0.079 |
| 7891 | 0.009 | 0.012 |
| 7892 | 0.030 | 0.083 |
| 7893 | 0.045 | -0.056 |
| 7894 | 0.040 | -0.041 |
| 7895 | 0.036 | -0.111 |
| 7896 | 0.023 | -0.108 |
| 7897 | 0.001 | -0.051 |
| 7898 | -0.079 | -0.021 |

| | | |
|---|---|---|
| 7899 | -0.030 | 0.063 |
| 7900 | -0.012 | 0.027 |
| 7901 | 0.008 | 0.158 |
| 7902 | 0.042 | -0.070 |
| 7903 | 0.009 | 0.017 |
| 7904 | 0.003 | -0.148 |
| 7905 | 0.008 | -0.038 |
| 7906 | -0.010 | 0.028 |
| 7907 | -0.001 | 0.006 |
| 7908 | -0.010 | 0.006 |
| 7909 | 0.006 | -0.049 |
| 7910 | -0.057 | -0.009 |
| 7911 | 0.025 | -0.007 |
| 7912 | 0.001 | -0.070 |
| 7913 | 0.039 | -0.010 |
| 7914 | -0.046 | -0.032 |
| 7915 | 0.050 | 0.077 |
| 7916 | -0.061 | -0.006 |
| 7917 | -0.006 | -0.054 |
| 7918 | -0.045 | 0.011 |
| 7919 | -0.050 | -0.033 |
| 7920 | 0.006 | 0.076 |
| 7921 | -0.054 | -0.047 |
| 7922 | 0.017 | 0.067 |
| 7923 | -0.022 | -0.082 |
| 7924 | 0.021 | -0.104 |
| 7925 | -0.034 | -0.120 |
| 7926 | -0.013 | -0.068 |
| 7927 | 0.026 | -0.029 |
| 7928 | -0.017 | 0.028 |
| 7929 | -0.050 | 0.056 |
| 7930 | -0.027 | -0.123 |
| 7931 | 0.041 | 0.024 |
| 7932 | -0.006 | 0.084 |
| 7933 | -0.009 | -0.115 |
| 7934 | -0.023 | 0.083 |
| 7935 | -0.033 | 0.020 |
| 7936 | -0.049 | -0.033 |
| 7937 | -0.086 | -0.027 |
| 7938 | -0.026 | -0.164 |
| 7939 | 0.001 | -0.048 |
| 7940 | 0.031 | 0.106 |
| 7941 | -0.048 | 0.023 |
| 7942 | 0.015 | -0.108 |
| 7943 | -0.037 | 0.003 |
| 7944 | -0.010 | -0.052 |
| 7945 | -0.032 | -0.001 |
| 7946 | -0.003 | -0.076 |
| 7947 | -0.030 | 0.007 |
| 7948 | -0.051 | 0.073 |
| 7949 | 0.018 | 0.020 |

| | | |
|---|---|---|
| 7950 | 0.035 | 0.065 |
| 7951 | 0.063 | 0.046 |
| 7952 | -0.002 | 0.050 |
| 7953 | 0.042 | -0.056 |
| 7954 | 0.043 | -0.122 |
| 7955 | -0.050 | -0.001 |
| 7956 | 0.001 | -0.074 |
| 7957 | 0.069 | 0.052 |
| 7958 | -0.014 | -0.027 |
| 7959 | -0.054 | 0.109 |
| 7960 | -0.069 | -0.063 |
| 7961 | -0.018 | 0.091 |
| 7962 | -0.066 | 0.009 |
| 7963 | -0.039 | -0.023 |
| 7964 | 0.002 | -0.026 |
| 7965 | 0.075 | 0.019 |
| 7966 | 0.035 | 0.032 |
| 7967 | -0.052 | -0.003 |
| 7968 | -0.011 | 0.013 |
| 7969 | -0.002 | -0.080 |
| 7970 | -0.024 | -0.044 |
| 7971 | -0.010 | -0.065 |
| 7972 | 0.103 | 0.032 |
| 7973 | -0.014 | 0.044 |
| 7974 | 0.018 | -0.013 |
| 7975 | 0.022 | -0.015 |
| 7976 | 0.026 | -0.058 |
| 7977 | 0.068 | -0.072 |
| 7978 | 0.048 | 0.107 |
| 7979 | 0.059 | -0.065 |
| 7980 | 0.027 | -0.079 |
| 7981 | 0.008 | 0.013 |
| 7982 | 0.037 | -0.004 |
| 7983 | -0.065 | 0.134 |
| 7984 | -0.043 | -0.009 |
| 7985 | -0.016 | -0.040 |
| 7986 | -0.024 | -0.161 |
| 7987 | 0.034 | -0.044 |
| 7988 | 0.016 | 0.038 |
| 7989 | 0.005 | 0.116 |
| 7990 | -0.003 | -0.019 |
| 7991 | -0.045 | -0.026 |
| 7992 | -0.007 | 0.037 |
| 7993 | 0.017 | -0.004 |
| 7994 | -0.010 | -0.109 |
| 7995 | -0.028 | 0.079 |
| 7996 | -0.002 | 0.072 |
| 7997 | 0.008 | -0.085 |
| 7998 | -0.016 | -0.094 |
| 7999 | -0.020 | 0.009 |
| 8000 | 0.044 | -0.068 |

| | | |
|---|---|---|
| 8001 | -0.023 | -0.052 |
| 8002 | 0.014 | -0.086 |
| 8003 | -0.046 | 0.088 |
| 8004 | 0.007 | -0.006 |
| 8005 | -0.033 | 0.016 |
| 8006 | 0.034 | -0.014 |
| 8007 | -0.053 | -0.128 |
| 8008 | -0.029 | -0.013 |
| 8009 | 0.043 | 0.021 |
| 8010 | 0.006 | 0.021 |
| 8011 | -0.048 | 0.084 |
| 8012 | -0.017 | -0.005 |
| 8013 | 0.000 | 0.039 |
| 8014 | 0.011 | -0.086 |
| 8015 | 0.030 | 0.024 |
| 8016 | -0.013 | 0.007 |
| 8017 | -0.022 | -0.089 |
| 8018 | -0.033 | 0.023 |
| 8019 | -0.028 | -0.103 |
| 8020 | -0.025 | -0.215 |
| 8021 | -0.009 | 0.048 |
| 8022 | 0.022 | -0.042 |
| 8023 | -0.011 | -0.037 |
| 8024 | -0.007 | -0.064 |
| 8025 | -0.016 | -0.070 |
| 8026 | -0.001 | -0.127 |
| 8027 | 0.024 | -0.018 |
| 8028 | 0.009 | 0.012 |
| 8029 | 0.030 | -0.018 |
| 8030 | -0.007 | -0.086 |
| 8031 | 0.012 | -0.021 |
| 8032 | -0.008 | -0.096 |
| 8033 | 0.005 | -0.013 |
| 8034 | -0.022 | -0.060 |
| 8035 | -0.019 | -0.088 |
| 8036 | -0.018 | 0.061 |
| 8037 | 0.030 | -0.054 |
| 8038 | 0.013 | -0.110 |
| 8039 | 0.083 | -0.001 |
| 8040 | -0.020 | 0.095 |
| 8041 | -0.018 | 0.073 |
| 8042 | -0.032 | 0.042 |
| 8043 | 0.005 | -0.079 |
| 8044 | -0.037 | -0.089 |
| 8045 | 0.043 | 0.003 |
| 8046 | 0.010 | 0.127 |
| 8047 | 0.043 | -0.081 |
| 8048 | -0.015 | 0.059 |
| 8049 | 0.048 | 0.047 |
| 8050 | -0.022 | 0.060 |
| 8051 | -0.039 | 0.034 |

| | | |
|---|---|---|
| 8052 | 0.019 | -0.060 |
| 8053 | 0.059 | 0.047 |
| 8054 | 0.002 | 0.065 |
| 8055 | 0.024 | -0.033 |
| 8056 | 0.037 | 0.036 |
| 8057 | 0.067 | 0.062 |
| 8058 | 0.030 | 0.051 |
| 8059 | 0.018 | 0.033 |
| 8060 | 0.021 | -0.141 |
| 8061 | 0.016 | -0.013 |
| 8062 | 0.031 | -0.007 |
| 8063 | 0.034 | 0.018 |
| 8064 | -0.021 | 0.037 |
| 8065 | 0.011 | -0.104 |
| 8066 | 0.032 | -0.071 |
| 8067 | -0.017 | -0.009 |
| 8068 | 0.023 | -0.019 |
| 8069 | -0.019 | 0.084 |
| 8070 | -0.035 | 0.039 |
| 8071 | -0.026 | -0.140 |
| 8072 | -0.007 | -0.001 |
| 8073 | -0.011 | -0.042 |
| 8074 | -0.007 | 0.031 |
| 8075 | -0.009 | 0.089 |
| 8076 | -0.003 | -0.032 |
| 8077 | 0.027 | -0.120 |
| 8078 | 0.047 | -0.050 |
| 8079 | -0.013 | -0.043 |
| 8080 | 0.016 | -0.036 |
| 8081 | -0.008 | -0.003 |
| 8082 | -0.009 | 0.007 |
| 8083 | -0.060 | 0.051 |
| 8084 | 0.002 | 0.015 |
| 8085 | -0.001 | -0.048 |
| 8086 | -0.058 | 0.153 |
| 8087 | -0.054 | -0.081 |
| 8088 | -0.011 | -0.039 |
| 8089 | 0.075 | -0.066 |
| 8090 | -0.051 | -0.026 |
| 8091 | -0.023 | -0.020 |
| 8092 | -0.015 | -0.095 |
| 8093 | 0.020 | 0.007 |
| 8094 | 0.069 | -0.111 |
| 8095 | -0.069 | -0.101 |
| 8096 | 0.050 | -0.052 |
| 8097 | -0.019 | 0.139 |
| 8098 | -0.007 | -0.091 |
| 8099 | -0.089 | -0.002 |
| 8100 | 0.002 | 0.023 |
| 8101 | 0.067 | -0.092 |
| 8102 | -0.023 | 0.045 |

| | | |
|---|---|---|
| 8103 | 0.025 | -0.047 |
| 8104 | -0.014 | 0.102 |
| 8105 | 0.021 | -0.005 |
| 8106 | 0.015 | 0.059 |
| 8107 | 0.013 | 0.014 |
| 8108 | 0.015 | 0.025 |
| 8109 | 0.019 | -0.008 |
| 8110 | 0.001 | 0.004 |
| 8111 | 0.035 | 0.084 |
| 8112 | 0.056 | 0.074 |
| 8113 | -0.020 | 0.058 |
| 8114 | 0.005 | -0.077 |
| 8115 | 0.019 | 0.019 |
| 8116 | -0.021 | -0.062 |
| 8117 | -0.002 | -0.009 |
| 8118 | 0.007 | -0.050 |
| 8119 | -0.005 | 0.091 |
| 8120 | 0.048 | 0.115 |
| 8121 | 0.002 | 0.187 |
| 8122 | 0.061 | 0.032 |
| 8123 | -0.012 | 0.026 |
| 8124 | -0.018 | 0.067 |
| 8125 | -0.027 | -0.124 |
| 8126 | 0.020 | 0.112 |
| 8127 | 0.005 | -0.056 |
| 8128 | -0.024 | 0.047 |
| 8129 | 0.033 | -0.034 |
| 8130 | 0.003 | 0.075 |
| 8131 | -0.048 | -0.092 |
| 8132 | 0.008 | -0.003 |
| 8133 | 0.050 | 0.040 |
| 8134 | -0.053 | -0.041 |
| 8135 | 0.018 | -0.031 |
| 8136 | -0.039 | -0.129 |
| 8137 | -0.010 | 0.071 |
| 8138 | -0.034 | 0.002 |
| 8139 | 0.010 | 0.017 |
| 8140 | -0.019 | 0.104 |
| 8141 | 0.024 | 0.042 |
| 8142 | -0.021 | 0.038 |
| 8143 | 0.031 | 0.026 |
| 8144 | -0.038 | -0.026 |
| 8145 | -0.009 | -0.056 |
| 8146 | 0.014 | -0.060 |
| 8147 | -0.026 | -0.081 |
| 8148 | -0.060 | 0.159 |
| 8149 | -0.003 | -0.020 |
| 8150 | -0.034 | -0.016 |
| 8151 | -0.043 | 0.099 |
| 8152 | -0.051 | 0.117 |
| 8153 | -0.025 | -0.047 |

| | | |
|---|---|---|
| 8154 | 0.059 | -0.110 |
| 8155 | 0.001 | -0.112 |
| 8156 | -0.006 | 0.013 |
| 8157 | 0.038 | 0.036 |
| 8158 | -0.003 | -0.074 |
| 8159 | 0.021 | -0.012 |
| 8160 | 0.026 | -0.012 |
| 8161 | -0.036 | -0.033 |
| 8162 | 0.060 | -0.223 |
| 8163 | -0.029 | -0.033 |
| 8164 | 0.023 | -0.095 |
| 8165 | -0.006 | 0.024 |
| 8166 | 0.020 | -0.030 |
| 8167 | -0.072 | -0.004 |
| 8168 | -0.027 | -0.011 |
| 8169 | -0.011 | 0.042 |
| 8170 | -0.003 | 0.028 |
| 8171 | -0.043 | -0.113 |
| 8172 | 0.055 | 0.121 |
| 8173 | -0.035 | -0.105 |
| 8174 | 0.002 | 0.057 |
| 8175 | -0.016 | -0.040 |
| 8176 | 0.009 | -0.089 |
| 8177 | -0.046 | 0.086 |
| 8178 | -0.005 | 0.101 |
| 8179 | 0.056 | 0.009 |
| 8180 | 0.030 | -0.064 |
| 8181 | 0.015 | 0.025 |
| 8182 | 0.007 | -0.056 |
| 8183 | -0.019 | 0.061 |
| 8184 | -0.012 | -0.066 |
| 8185 | -0.001 | -0.059 |
| 8186 | -0.008 | -0.119 |
| 8187 | 0.040 | 0.038 |
| 8188 | -0.045 | 0.038 |
| 8189 | 0.033 | 0.081 |
| 8190 | 0.036 | -0.035 |
| 8191 | -0.008 | -0.083 |
| 8192 | 0.033 | -0.063 |
| 8193 | -0.028 | -0.039 |
| 8194 | -0.004 | 0.091 |
| 8195 | -0.032 | -0.023 |
| 8196 | -0.047 | 0.165 |
| 8197 | -0.067 | 0.053 |
| 8198 | -0.060 | 0.018 |
| 8199 | 0.018 | 0.058 |
| 8200 | 0.036 | -0.006 |
| 8201 | 0.022 | -0.020 |
| 8202 | 0.029 | -0.090 |
| 8203 | 0.016 | -0.055 |
| 8204 | 0.046 | 0.080 |

| | | |
|---|---|---|
| 8205 | 0.032 | 0.094 |
| 8206 | -0.015 | 0.089 |
| 8207 | 0.048 | 0.038 |
| 8208 | 0.036 | -0.021 |
| 8209 | -0.017 | 0.076 |
| 8210 | -0.040 | 0.128 |
| 8211 | -0.012 | 0.067 |
| 8212 | -0.003 | -0.088 |
| 8213 | 0.013 | -0.089 |
| 8214 | 0.000 | 0.106 |
| 8215 | 0.020 | -0.016 |
| 8216 | -0.022 | -0.035 |
| 8217 | -0.031 | 0.040 |
| 8218 | -0.026 | -0.065 |
| 8219 | -0.051 | -0.041 |
| 8220 | 0.029 | 0.009 |
| 8221 | -0.002 | 0.022 |
| 8222 | -0.012 | 0.093 |
| 8223 | 0.005 | 0.001 |
| 8224 | -0.009 | 0.062 |
| 8225 | -0.016 | 0.047 |
| 8226 | -0.033 | 0.032 |
| 8227 | 0.008 | -0.073 |
| 8228 | 0.038 | 0.083 |
| 8229 | -0.059 | -0.034 |
| 8230 | 0.002 | -0.070 |
| 8231 | 0.042 | 0.048 |
| 8232 | 0.036 | 0.005 |
| 8233 | 0.002 | 0.072 |
| 8234 | 0.026 | 0.075 |
| 8235 | 0.063 | -0.024 |
| 8236 | 0.023 | -0.012 |
| 8237 | -0.038 | -0.110 |
| 8238 | 0.008 | 0.054 |
| 8239 | 0.035 | 0.026 |
| 8240 | -0.054 | -0.082 |
| 8241 | 0.031 | -0.007 |
| 8242 | 0.008 | -0.050 |
| 8243 | 0.013 | -0.089 |
| 8244 | 0.051 | -0.084 |
| 8245 | 0.033 | 0.107 |
| 8246 | 0.028 | 0.005 |
| 8247 | -0.093 | -0.040 |
| 8248 | 0.014 | -0.018 |
| 8249 | -0.012 | -0.041 |
| 8250 | 0.031 | 0.008 |
| 8251 | -0.066 | -0.019 |
| 8252 | -0.018 | 0.105 |
| 8253 | 0.045 | -0.049 |
| 8254 | 0.026 | 0.016 |
| 8255 | 0.005 | 0.087 |

| | | |
|---|---|---|
| 8256 | 0.018 | 0.040 |
| 8257 | -0.018 | -0.114 |
| 8258 | -0.016 | 0.014 |
| 8259 | 0.021 | 0.046 |
| 8260 | -0.029 | -0.041 |
| 8261 | -0.005 | 0.090 |
| 8262 | 0.004 | -0.006 |
| 8263 | -0.013 | -0.084 |
| 8264 | -0.009 | -0.016 |
| 8265 | -0.003 | 0.072 |
| 8266 | -0.040 | -0.047 |
| 8267 | -0.019 | -0.039 |
| 8268 | 0.045 | -0.070 |
| 8269 | 0.005 | 0.046 |
| 8270 | -0.020 | 0.042 |
| 8271 | 0.023 | -0.017 |
| 8272 | 0.040 | 0.051 |
| 8273 | 0.003 | -0.054 |
| 8274 | -0.020 | -0.031 |
| 8275 | 0.033 | 0.045 |
| 8276 | 0.076 | -0.019 |
| 8277 | -0.031 | -0.028 |
| 8278 | 0.032 | 0.010 |
| 8279 | 0.004 | -0.119 |
| 8280 | -0.046 | 0.114 |
| 8281 | -0.067 | 0.132 |
| 8282 | 0.048 | 0.019 |
| 8283 | -0.010 | -0.127 |
| 8284 | -0.035 | 0.066 |
| 8285 | 0.020 | 0.037 |
| 8286 | 0.056 | 0.003 |
| 8287 | -0.067 | 0.071 |
| 8288 | 0.075 | -0.021 |
| 8289 | 0.002 | 0.109 |
| 8290 | 0.039 | 0.059 |
| 8291 | 0.020 | 0.046 |
| 8292 | -0.084 | -0.051 |
| 8293 | 0.079 | -0.025 |
| 8294 | -0.017 | 0.018 |
| 8295 | -0.061 | 0.006 |
| 8296 | 0.006 | -0.042 |
| 8297 | -0.066 | 0.050 |
| 8298 | 0.037 | 0.063 |
| 8299 | 0.017 | 0.019 |
| 8300 | -0.017 | -0.132 |
| 8301 | -0.031 | 0.146 |
| 8302 | -0.056 | 0.057 |
| 8303 | -0.021 | -0.002 |
| 8304 | -0.033 | -0.084 |
| 8305 | 0.006 | -0.098 |
| 8306 | 0.022 | 0.056 |

| | | |
|---|---|---|
| 8307 | -0.005 | -0.018 |
| 8308 | 0.015 | 0.012 |
| 8309 | -0.010 | 0.067 |
| 8310 | 0.004 | 0.045 |
| 8311 | 0.039 | -0.086 |
| 8312 | 0.033 | 0.049 |
| 8313 | 0.038 | 0.105 |
| 8314 | -0.007 | 0.009 |
| 8315 | -0.030 | 0.070 |
| 8316 | 0.051 | 0.132 |
| 8317 | -0.017 | 0.069 |
| 8318 | 0.043 | -0.041 |
| 8319 | -0.052 | -0.006 |
| 8320 | -0.071 | -0.095 |
| 8321 | -0.059 | 0.080 |
| 8322 | 0.061 | 0.045 |
| 8323 | 0.069 | 0.145 |
| 8324 | 0.014 | -0.099 |
| 8325 | -0.006 | -0.163 |
| 8326 | -0.029 | -0.054 |
| 8327 | -0.007 | -0.082 |
| 8328 | -0.051 | -0.018 |
| 8329 | -0.027 | -0.055 |
| 8330 | -0.005 | -0.021 |
| 8331 | -0.044 | 0.054 |
| 8332 | -0.026 | 0.108 |
| 8333 | 0.021 | 0.020 |
| 8334 | 0.040 | 0.105 |
| 8335 | 0.004 | -0.026 |
| 8336 | 0.057 | 0.044 |
| 8337 | -0.009 | 0.022 |
| 8338 | -0.009 | -0.042 |
| 8339 | -0.033 | 0.017 |
| 8340 | -0.023 | 0.029 |
| 8341 | -0.028 | 0.066 |
| 8342 | 0.022 | -0.073 |
| 8343 | -0.013 | -0.112 |
| 8344 | -0.040 | 0.010 |
| 8345 | 0.086 | 0.027 |
| 8346 | -0.024 | 0.069 |
| 8347 | 0.011 | -0.029 |
| 8348 | -0.002 | 0.140 |
| 8349 | 0.028 | 0.014 |
| 8350 | -0.005 | -0.074 |
| 8351 | 0.040 | -0.039 |
| 8352 | 0.022 | 0.127 |
| 8353 | -0.035 | -0.025 |
| 8354 | 0.015 | -0.056 |
| 8355 | -0.015 | 0.013 |
| 8356 | -0.076 | -0.080 |
| 8357 | -0.055 | 0.036 |

| | | |
|---|---|---|
| 8358 | -0.007 | 0.069 |
| 8359 | 0.057 | -0.083 |
| 8360 | 0.004 | -0.039 |
| 8361 | -0.048 | -0.015 |
| 8362 | 0.002 | 0.022 |
| 8363 | 0.102 | -0.059 |
| 8364 | -0.016 | 0.082 |
| 8365 | 0.051 | -0.020 |
| 8366 | -0.017 | 0.059 |
| 8367 | -0.011 | -0.071 |
| 8368 | -0.043 | 0.061 |
| 8369 | 0.050 | 0.003 |
| 8370 | 0.030 | 0.068 |
| 8371 | 0.013 | 0.105 |
| 8372 | 0.015 | -0.045 |
| 8373 | -0.001 | -0.064 |
| 8374 | -0.034 | -0.094 |
| 8375 | -0.032 | 0.107 |
| 8376 | 0.016 | 0.097 |
| 8377 | 0.011 | -0.079 |
| 8378 | -0.010 | 0.071 |
| 8379 | 0.000 | 0.169 |
| 8380 | -0.009 | 0.108 |
| 8381 | 0.000 | -0.044 |
| 8382 | -0.017 | -0.080 |
| 8383 | -0.014 | 0.163 |
| 8384 | 0.008 | -0.038 |
| 8385 | 0.007 | -0.113 |
| 8386 | -0.041 | 0.121 |
| 8387 | 0.020 | 0.068 |
| 8388 | 0.024 | 0.015 |
| 8389 | -0.024 | 0.047 |
| 8390 | 0.012 | 0.015 |
| 8391 | -0.011 | -0.020 |
| 8392 | -0.012 | 0.141 |
| 8393 | 0.001 | 0.029 |
| 8394 | 0.008 | 0.078 |
| 8395 | 0.003 | 0.071 |
| 8396 | -0.057 | 0.118 |
| 8397 | 0.021 | 0.054 |
| 8398 | -0.016 | -0.001 |
| 8399 | 0.015 | -0.059 |
| 8400 | -0.047 | -0.036 |
| 8401 | 0.044 | -0.010 |
| 8402 | 0.031 | 0.006 |
| 8403 | 0.027 | -0.019 |
| 8404 | -0.018 | -0.062 |
| 8405 | -0.070 | 0.052 |
| 8406 | -0.044 | 0.120 |
| 8407 | -0.068 | -0.007 |
| 8408 | -0.010 | -0.079 |

| | | |
|---|---|---|
| 8409 | -0.038 | 0.046 |
| 8410 | -0.057 | 0.065 |
| 8411 | -0.007 | 0.003 |
| 8412 | 0.025 | 0.026 |
| 8413 | -0.013 | 0.004 |
| 8414 | 0.029 | -0.023 |
| 8415 | -0.016 | -0.009 |
| 8416 | 0.002 | -0.031 |
| 8417 | -0.048 | 0.015 |
| 8418 | 0.006 | -0.046 |
| 8419 | -0.042 | 0.013 |
| 8420 | 0.016 | 0.066 |
| 8421 | 0.029 | 0.040 |
| 8422 | 0.003 | 0.012 |
| 8423 | -0.032 | 0.061 |
| 8424 | 0.068 | 0.002 |
| 8425 | -0.008 | 0.006 |
| 8426 | -0.032 | 0.010 |
| 8427 | 0.020 | -0.057 |
| 8428 | -0.013 | 0.024 |
| 8429 | -0.035 | 0.004 |
| 8430 | 0.042 | 0.025 |
| 8431 | 0.037 | -0.020 |
| 8432 | -0.021 | 0.025 |
| 8433 | -0.026 | 0.121 |
| 8434 | -0.017 | -0.027 |
| 8435 | -0.021 | 0.025 |
| 8436 | -0.034 | 0.065 |
| 8437 | 0.048 | 0.095 |
| 8438 | 0.000 | 0.098 |
| 8439 | -0.033 | -0.031 |
| 8440 | 0.033 | 0.012 |
| 8441 | 0.105 | -0.088 |
| 8442 | -0.053 | -0.025 |
| 8443 | -0.043 | -0.043 |
| 8444 | 0.017 | -0.029 |
| 8445 | 0.061 | 0.034 |
| 8446 | 0.005 | 0.062 |
| 8447 | -0.006 | 0.087 |
| 8448 | -0.032 | -0.100 |
| 8449 | -0.015 | 0.023 |
| 8450 | -0.013 | 0.001 |
| 8451 | 0.071 | 0.101 |
| 8452 | 0.014 | 0.074 |
| 8453 | 0.045 | 0.065 |
| 8454 | 0.027 | -0.162 |
| 8455 | -0.003 | -0.027 |
| 8456 | -0.026 | -0.022 |
| 8457 | -0.011 | -0.103 |
| 8458 | 0.003 | -0.028 |
| 8459 | -0.063 | -0.041 |

| | | |
|---|---|---|
| 8460 | 0.033 | 0.051 |
| 8461 | 0.052 | 0.075 |
| 8462 | 0.049 | 0.108 |
| 8463 | -0.010 | 0.090 |
| 8464 | 0.000 | -0.025 |
| 8465 | 0.029 | -0.121 |
| 8466 | -0.017 | -0.031 |
| 8467 | -0.019 | 0.008 |
| 8468 | -0.010 | -0.055 |
| 8469 | 0.010 | -0.086 |
| 8470 | 0.056 | -0.039 |
| 8471 | -0.012 | -0.067 |
| 8472 | 0.026 | -0.018 |
| 8473 | -0.020 | -0.117 |
| 8474 | 0.022 | -0.075 |
| 8475 | -0.030 | 0.160 |
| 8476 | -0.073 | 0.034 |
| 8477 | -0.007 | -0.095 |
| 8478 | -0.027 | -0.197 |
| 8479 | 0.009 | 0.256 |
| 8480 | 0.003 | -0.203 |
| 8481 | -0.018 | -0.077 |
| 8482 | -0.040 | -0.019 |
| 8483 | -0.025 | -0.017 |
| 8484 | -0.029 | 0.089 |
| 8485 | -0.003 | -0.025 |
| 8486 | -0.021 | 0.016 |
| 8487 | 0.003 | 0.176 |
| 8488 | 0.019 | 0.038 |
| 8489 | -0.041 | -0.050 |
| 8490 | 0.009 | 0.000 |
| 8491 | 0.008 | 0.036 |
| 8492 | -0.013 | 0.025 |
| 8493 | -0.018 | 0.026 |
| 8494 | 0.008 | 0.013 |
| 8495 | -0.020 | -0.008 |
| 8496 | 0.012 | 0.035 |
| 8497 | 0.017 | 0.101 |
| 8498 | 0.065 | 0.015 |
| 8499 | 0.008 | 0.027 |
| 8500 | -0.009 | -0.090 |
| 8501 | 0.031 | 0.020 |
| 8502 | -0.011 | 0.041 |
| 8503 | -0.021 | 0.028 |
| 8504 | -0.004 | -0.050 |
| 8505 | -0.008 | -0.059 |
| 8506 | 0.029 | 0.058 |
| 8507 | 0.002 | -0.037 |
| 8508 | 0.006 | 0.043 |
| 8509 | 0.007 | 0.009 |
| 8510 | 0.035 | -0.047 |

| | | |
|---|---|---|
| 8511 | -0.004 | -0.034 |
| 8512 | -0.032 | 0.021 |
| 8513 | 0.031 | 0.024 |
| 8514 | 0.013 | 0.028 |
| 8515 | 0.076 | -0.071 |
| 8516 | -0.075 | 0.061 |
| 8517 | -0.011 | -0.110 |
| 8518 | -0.052 | 0.070 |
| 8519 | -0.017 | -0.045 |
| 8520 | -0.008 | 0.084 |
| 8521 | -0.021 | 0.004 |
| 8522 | 0.000 | 0.054 |
| 8523 | 0.023 | -0.049 |
| 8524 | -0.027 | 0.082 |
| 8525 | 0.018 | 0.024 |
| 8526 | -0.006 | -0.034 |
| 8527 | 0.012 | 0.121 |
| 8528 | -0.031 | 0.032 |
| 8529 | -0.032 | 0.101 |
| 8530 | -0.005 | 0.078 |
| 8531 | 0.008 | -0.032 |
| 8532 | 0.025 | 0.002 |
| 8533 | 0.010 | -0.180 |
| 8534 | 0.019 | -0.029 |
| 8535 | -0.061 | -0.018 |
| 8536 | -0.027 | -0.032 |
| 8537 | -0.067 | 0.062 |
| 8538 | -0.063 | 0.021 |
| 8539 | -0.041 | 0.117 |
| 8540 | 0.029 | 0.058 |
| 8541 | 0.018 | 0.071 |
| 8542 | -0.058 | -0.097 |
| 8543 | 0.004 | -0.002 |
| 8544 | 0.074 | 0.078 |
| 8545 | 0.020 | -0.049 |
| 8546 | 0.035 | -0.017 |
| 8547 | 0.046 | 0.083 |
| 8548 | -0.020 | 0.074 |
| 8549 | -0.017 | -0.024 |
| 8550 | -0.041 | 0.034 |
| 8551 | -0.013 | -0.042 |
| 8552 | 0.079 | 0.036 |
| 8553 | -0.043 | -0.110 |
| 8554 | -0.041 | 0.205 |
| 8555 | -0.014 | -0.004 |
| 8556 | -0.037 | 0.085 |
| 8557 | -0.048 | 0.132 |
| 8558 | 0.010 | 0.035 |
| 8559 | -0.041 | 0.012 |
| 8560 | 0.008 | -0.021 |
| 8561 | 0.008 | 0.011 |

| | | |
|---|---|---|
| 8562 | -0.012 | -0.102 |
| 8563 | -0.045 | 0.018 |
| 8564 | 0.008 | -0.005 |
| 8565 | -0.047 | -0.058 |
| 8566 | -0.014 | -0.024 |
| 8567 | -0.007 | 0.050 |
| 8568 | 0.021 | -0.035 |
| 8569 | 0.028 | 0.067 |
| 8570 | 0.021 | 0.100 |
| 8571 | 0.044 | -0.021 |
| 8572 | 0.028 | 0.112 |
| 8573 | -0.015 | 0.125 |
| 8574 | -0.027 | -0.111 |
| 8575 | -0.020 | 0.045 |
| 8576 | 0.015 | -0.091 |
| 8577 | 0.024 | 0.057 |
| 8578 | -0.037 | -0.002 |
| 8579 | -0.021 | 0.022 |
| 8580 | 0.047 | 0.031 |
| 8581 | -0.038 | 0.050 |
| 8582 | 0.022 | -0.015 |
| 8583 | 0.020 | 0.098 |
| 8584 | -0.004 | 0.149 |
| 8585 | -0.028 | -0.058 |
| 8586 | -0.013 | 0.004 |
| 8587 | -0.002 | 0.018 |
| 8588 | -0.022 | -0.110 |
| 8589 | 0.026 | 0.069 |
| 8590 | 0.015 | 0.141 |
| 8591 | 0.077 | -0.012 |
| 8592 | -0.023 | -0.029 |
| 8593 | 0.032 | 0.227 |
| 8594 | 0.019 | 0.095 |
| 8595 | -0.074 | 0.006 |
| 8596 | 0.038 | -0.027 |
| 8597 | -0.029 | -0.027 |
| 8598 | -0.029 | -0.034 |
| 8599 | 0.002 | -0.152 |
| 8600 | -0.028 | -0.066 |
| 8601 | 0.051 | -0.107 |
| 8602 | 0.017 | 0.060 |
| 8603 | 0.000 | 0.003 |
| 8604 | 0.009 | 0.160 |
| 8605 | -0.088 | -0.032 |
| 8606 | 0.010 | 0.072 |
| 8607 | -0.027 | -0.088 |
| 8608 | 0.019 | -0.031 |
| 8609 | -0.068 | 0.016 |
| 8610 | 0.038 | 0.020 |
| 8611 | -0.012 | 0.094 |
| 8612 | 0.019 | -0.019 |

| | | |
|---|---|---|
| 8613 | -0.026 | -0.049 |
| 8614 | 0.120 | -0.041 |
| 8615 | 0.004 | 0.128 |
| 8616 | 0.021 | -0.004 |
| 8617 | -0.029 | 0.073 |
| 8618 | 0.048 | 0.087 |
| 8619 | -0.001 | 0.031 |
| 8620 | -0.016 | 0.124 |
| 8621 | -0.034 | 0.005 |
| 8622 | 0.056 | 0.100 |
| 8623 | 0.048 | -0.123 |
| 8624 | -0.043 | -0.073 |
| 8625 | -0.027 | -0.065 |
| 8626 | 0.045 | -0.089 |
| 8627 | 0.018 | 0.090 |
| 8628 | 0.036 | 0.117 |
| 8629 | -0.020 | -0.087 |
| 8630 | 0.006 | 0.025 |
| 8631 | -0.008 | 0.064 |
| 8632 | -0.087 | -0.090 |
| 8633 | -0.031 | 0.024 |
| 8634 | -0.010 | 0.014 |
| 8635 | -0.009 | -0.059 |
| 8636 | 0.056 | -0.040 |
| 8637 | 0.040 | 0.001 |
| 8638 | 0.012 | 0.006 |
| 8639 | -0.008 | -0.041 |
| 8640 | 0.028 | 0.057 |
| 8641 | 0.096 | -0.013 |
| 8642 | -0.021 | -0.075 |
| 8643 | 0.004 | -0.082 |
| 8644 | 0.016 | -0.105 |
| 8645 | 0.047 | -0.129 |
| 8646 | 0.026 | -0.073 |
| 8647 | -0.001 | 0.056 |
| 8648 | -0.003 | -0.155 |
| 8649 | 0.031 | 0.121 |
| 8650 | 0.014 | 0.008 |
| 8651 | 0.013 | -0.024 |
| 8652 | -0.079 | 0.061 |
| 8653 | 0.042 | 0.015 |
| 8654 | 0.097 | -0.067 |
| 8655 | -0.021 | -0.076 |
| 8656 | -0.005 | -0.102 |
| 8657 | 0.064 | -0.065 |
| 8658 | -0.049 | 0.054 |
| 8659 | -0.010 | -0.090 |
| 8660 | 0.010 | 0.100 |
| 8661 | -0.029 | 0.005 |
| 8662 | 0.025 | -0.065 |
| 8663 | -0.082 | -0.014 |

| | | |
|---|---|---|
| 8664 | -0.005 | -0.014 |
| 8665 | 0.002 | -0.081 |
| 8666 | -0.004 | 0.040 |
| 8667 | 0.020 | -0.007 |
| 8668 | 0.042 | -0.158 |
| 8669 | 0.004 | -0.026 |
| 8670 | -0.055 | -0.088 |
| 8671 | -0.006 | -0.038 |
| 8672 | -0.046 | 0.023 |
| 8673 | 0.059 | -0.035 |
| 8674 | 0.000 | -0.106 |
| 8675 | -0.004 | -0.082 |
| 8676 | 0.076 | 0.037 |
| 8677 | -0.047 | -0.071 |
| 8678 | 0.039 | -0.156 |
| 8679 | -0.077 | -0.023 |
| 8680 | -0.033 | 0.098 |
| 8681 | -0.027 | 0.094 |
| 8682 | 0.021 | 0.071 |
| 8683 | -0.045 | -0.113 |
| 8684 | 0.035 | -0.140 |
| 8685 | 0.036 | 0.177 |
| 8686 | -0.008 | -0.113 |
| 8687 | 0.032 | 0.045 |
| 8688 | 0.050 | -0.075 |
| 8689 | -0.022 | -0.030 |
| 8690 | 0.012 | -0.044 |
| 8691 | -0.001 | -0.012 |
| 8692 | 0.039 | 0.016 |
| 8693 | 0.060 | -0.013 |
| 8694 | 0.023 | 0.075 |
| 8695 | 0.021 | -0.075 |
| 8696 | 0.034 | -0.159 |
| 8697 | -0.010 | 0.008 |
| 8698 | -0.035 | 0.030 |
| 8699 | 0.012 | 0.088 |
| 8700 | 0.031 | -0.064 |
| 8701 | 0.009 | -0.051 |
| 8702 | -0.005 | -0.004 |
| 8703 | 0.032 | -0.141 |
| 8704 | 0.004 | -0.076 |
| 8705 | -0.004 | 0.048 |
| 8706 | 0.014 | -0.023 |
| 8707 | -0.034 | 0.072 |
| 8708 | 0.028 | -0.064 |
| 8709 | 0.036 | -0.036 |
| 8710 | -0.009 | -0.062 |
| 8711 | 0.021 | -0.044 |
| 8712 | 0.008 | -0.068 |
| 8713 | -0.078 | -0.111 |
| 8714 | 0.001 | 0.014 |

| | | |
|---|---|---|
| 8715 | 0.010 | 0.014 |
| 8716 | -0.007 | 0.042 |
| 8717 | 0.057 | 0.211 |
| 8718 | 0.008 | -0.011 |
| 8719 | 0.075 | -0.062 |
| 8720 | -0.003 | -0.015 |
| 8721 | -0.066 | 0.066 |
| 8722 | -0.015 | 0.026 |
| 8723 | 0.036 | 0.007 |
| 8724 | -0.009 | -0.027 |
| 8725 | 0.027 | 0.062 |
| 8726 | -0.023 | -0.053 |
| 8727 | -0.029 | 0.021 |
| 8728 | 0.022 | 0.040 |
| 8729 | 0.045 | -0.006 |
| 8730 | 0.013 | 0.080 |
| 8731 | 0.039 | -0.139 |
| 8732 | -0.001 | -0.018 |
| 8733 | 0.011 | -0.081 |
| 8734 | -0.023 | -0.011 |
| 8735 | -0.017 | -0.115 |
| 8736 | 0.046 | 0.100 |
| 8737 | -0.057 | 0.063 |
| 8738 | 0.001 | 0.037 |
| 8739 | -0.011 | -0.033 |
| 8740 | -0.033 | 0.138 |
| 8741 | -0.006 | -0.053 |
| 8742 | 0.010 | 0.061 |
| 8743 | 0.020 | 0.006 |
| 8744 | 0.018 | 0.012 |
| 8745 | -0.059 | -0.134 |
| 8746 | -0.033 | -0.054 |
| 8747 | 0.035 | -0.117 |
| 8748 | -0.049 | -0.193 |
| 8749 | 0.003 | 0.017 |
| 8750 | -0.039 | -0.028 |
| 8751 | -0.046 | -0.025 |
| 8752 | 0.067 | -0.056 |
| 8753 | 0.022 | 0.084 |
| 8754 | 0.045 | 0.020 |
| 8755 | 0.009 | 0.001 |
| 8756 | -0.009 | 0.107 |
| 8757 | 0.051 | -0.095 |
| 8758 | -0.009 | -0.120 |
| 8759 | -0.053 | -0.108 |
| 8760 | -0.061 | -0.118 |
| 8761 | -0.082 | 0.020 |
| 8762 | -0.015 | 0.138 |
| 8763 | -0.042 | -0.029 |
| 8764 | 0.014 | -0.061 |
| 8765 | -0.014 | 0.021 |

| | | |
|---|---|---|
| 8766 | 0.039 | -0.003 |
| 8767 | 0.002 | 0.044 |
| 8768 | 0.029 | -0.010 |
| 8769 | -0.021 | 0.042 |
| 8770 | -0.019 | -0.003 |
| 8771 | -0.022 | -0.123 |
| 8772 | -0.021 | 0.073 |
| 8773 | 0.007 | -0.104 |
| 8774 | 0.001 | 0.006 |
| 8775 | -0.053 | 0.036 |
| 8776 | 0.036 | -0.050 |
| 8777 | -0.035 | 0.052 |
| 8778 | -0.006 | -0.055 |
| 8779 | 0.002 | 0.070 |
| 8780 | 0.034 | 0.024 |
| 8781 | 0.016 | 0.008 |
| 8782 | 0.085 | 0.059 |
| 8783 | 0.005 | 0.024 |
| 8784 | -0.031 | -0.041 |
| 8785 | 0.027 | -0.084 |
| 8786 | -0.020 | 0.017 |
| 8787 | 0.009 | -0.040 |
| 8788 | 0.005 | 0.051 |
| 8789 | -0.045 | 0.048 |
| 8790 | -0.052 | -0.042 |
| 8791 | -0.009 | 0.067 |
| 8792 | -0.029 | -0.065 |
| 8793 | 0.027 | -0.046 |
| 8794 | 0.024 | -0.019 |
| 8795 | 0.045 | -0.081 |
| 8796 | 0.026 | -0.024 |
| 8797 | 0.044 | 0.037 |
| 8798 | 0.025 | -0.066 |
| 8799 | 0.033 | 0.095 |
| 8800 | 0.037 | -0.017 |
| 8801 | -0.076 | -0.040 |
| 8802 | -0.027 | 0.088 |
| 8803 | 0.013 | -0.001 |
| 8804 | -0.010 | -0.004 |
| 8805 | 0.071 | -0.041 |
| 8806 | -0.027 | -0.034 |
| 8807 | -0.012 | 0.006 |
| 8808 | -0.004 | -0.086 |
| 8809 | -0.002 | 0.011 |
| 8810 | 0.017 | 0.004 |
| 8811 | -0.002 | -0.049 |
| 8812 | -0.025 | 0.062 |
| 8813 | 0.071 | -0.027 |
| 8814 | 0.025 | -0.010 |
| 8815 | 0.043 | -0.046 |
| 8816 | 0.002 | -0.041 |

| | | |
|---|---|---|
| 8817 | 0.021 | 0.132 |
| 8818 | -0.020 | 0.052 |
| 8819 | -0.013 | 0.052 |
| 8820 | 0.014 | -0.030 |
| 8821 | 0.031 | -0.152 |
| 8822 | 0.006 | -0.151 |
| 8823 | 0.030 | 0.089 |
| 8824 | -0.036 | 0.051 |
| 8825 | -0.033 | -0.027 |
| 8826 | 0.016 | 0.037 |
| 8827 | 0.004 | 0.075 |
| 8828 | 0.015 | -0.046 |
| 8829 | -0.006 | 0.040 |
| 8830 | 0.072 | 0.021 |
| 8831 | 0.060 | -0.004 |
| 8832 | 0.060 | -0.006 |
| 8833 | -0.056 | -0.022 |
| 8834 | 0.016 | -0.072 |
| 8835 | -0.007 | -0.103 |
| 8836 | 0.053 | 0.074 |
| 8837 | 0.016 | 0.050 |
| 8838 | -0.102 | -0.153 |
| 8839 | -0.017 | 0.043 |
| 8840 | -0.033 | -0.023 |
| 8841 | 0.016 | 0.020 |
| 8842 | 0.019 | 0.107 |
| 8843 | 0.001 | -0.118 |
| 8844 | -0.045 | -0.096 |
| 8845 | 0.000 | 0.121 |
| 8846 | -0.044 | -0.077 |
| 8847 | 0.030 | -0.033 |
| 8848 | -0.020 | -0.076 |
| 8849 | 0.008 | 0.071 |
| 8850 | -0.008 | -0.017 |
| 8851 | -0.003 | -0.049 |
| 8852 | -0.014 | -0.105 |
| 8853 | 0.041 | 0.035 |
| 8854 | 0.100 | -0.116 |
| 8855 | -0.027 | 0.038 |
| 8856 | 0.025 | 0.011 |
| 8857 | 0.014 | -0.020 |
| 8858 | -0.009 | 0.018 |
| 8859 | -0.029 | -0.012 |
| 8860 | 0.022 | 0.073 |
| 8861 | -0.030 | -0.084 |
| 8862 | -0.019 | -0.141 |
| 8863 | -0.039 | 0.034 |
| 8864 | 0.022 | 0.037 |
| 8865 | -0.006 | -0.066 |
| 8866 | -0.030 | -0.009 |
| 8867 | 0.023 | -0.102 |

| | | |
|---|---|---|
| 8868 | -0.058 | 0.073 |
| 8869 | -0.056 | 0.113 |
| 8870 | 0.008 | -0.061 |
| 8871 | -0.019 | -0.128 |
| 8872 | -0.038 | -0.006 |
| 8873 | 0.006 | 0.093 |
| 8874 | 0.021 | -0.089 |
| 8875 | -0.024 | -0.014 |
| 8876 | 0.006 | 0.090 |
| 8877 | -0.033 | 0.056 |
| 8878 | 0.011 | -0.106 |
| 8879 | -0.030 | -0.080 |
| 8880 | 0.002 | 0.000 |
| 8881 | -0.018 | 0.016 |
| 8882 | -0.031 | 0.063 |
| 8883 | -0.006 | 0.029 |
| 8884 | 0.055 | 0.028 |
| 8885 | 0.032 | 0.047 |
| 8886 | -0.030 | -0.075 |
| 8887 | 0.049 | 0.068 |
| 8888 | -0.027 | -0.070 |
| 8889 | 0.052 | 0.003 |
| 8890 | 0.003 | 0.053 |
| 8891 | 0.033 | 0.148 |
| 8892 | 0.039 | -0.024 |
| 8893 | -0.044 | 0.073 |
| 8894 | 0.033 | -0.089 |
| 8895 | 0.030 | 0.096 |
| 8896 | 0.012 | 0.101 |
| 8897 | 0.016 | 0.046 |
| 8898 | -0.058 | -0.029 |
| 8899 | 0.028 | 0.135 |
| 8900 | 0.024 | -0.119 |
| 8901 | -0.008 | -0.030 |
| 8902 | -0.029 | -0.013 |
| 8903 | -0.002 | -0.103 |
| 8904 | -0.069 | -0.195 |
| 8905 | 0.030 | 0.097 |
| 8906 | 0.001 | 0.105 |
| 8907 | 0.026 | -0.113 |
| 8908 | 0.000 | -0.032 |
| 8909 | 0.025 | 0.064 |
| 8910 | -0.014 | -0.027 |
| 8911 | 0.017 | 0.019 |
| 8912 | 0.014 | -0.004 |
| 8913 | -0.021 | 0.104 |
| 8914 | 0.002 | -0.020 |
| 8915 | -0.028 | 0.096 |
| 8916 | -0.054 | 0.050 |
| 8917 | -0.014 | 0.068 |
| 8918 | 0.017 | -0.107 |

| | | |
|---|---|---|
| 8919 | -0.053 | -0.001 |
| 8920 | -0.029 | -0.014 |
| 8921 | -0.029 | 0.078 |
| 8922 | 0.024 | -0.069 |
| 8923 | -0.006 | 0.052 |
| 8924 | 0.019 | -0.050 |
| 8925 | -0.072 | 0.017 |
| 8926 | -0.006 | -0.012 |
| 8927 | 0.002 | 0.146 |
| 8928 | -0.026 | 0.030 |
| 8929 | 0.041 | 0.036 |
| 8930 | 0.005 | -0.034 |
| 8931 | -0.001 | -0.077 |
| 8932 | 0.020 | 0.012 |
| 8933 | -0.032 | -0.026 |
| 8934 | 0.040 | 0.115 |
| 8935 | -0.004 | 0.049 |
| 8936 | -0.034 | 0.022 |
| 8937 | -0.003 | 0.043 |
| 8938 | 0.026 | 0.046 |
| 8939 | -0.007 | 0.044 |
| 8940 | -0.054 | -0.061 |
| 8941 | -0.051 | -0.139 |
| 8942 | 0.032 | 0.080 |
| 8943 | 0.019 | -0.087 |
| 8944 | 0.020 | 0.034 |
| 8945 | 0.028 | 0.045 |
| 8946 | -0.062 | 0.040 |
| 8947 | -0.039 | -0.087 |
| 8948 | 0.062 | 0.137 |
| 8949 | -0.033 | 0.114 |
| 8950 | -0.068 | -0.022 |
| 8951 | -0.026 | -0.014 |
| 8952 | -0.001 | 0.086 |
| 8953 | -0.034 | -0.074 |
| 8954 | 0.023 | 0.114 |
| 8955 | -0.021 | 0.008 |
| 8956 | 0.022 | -0.047 |
| 8957 | 0.037 | 0.102 |
| 8958 | -0.030 | -0.037 |
| 8959 | 0.005 | -0.062 |
| 8960 | 0.006 | 0.043 |
| 8961 | -0.004 | 0.223 |
| 8962 | -0.058 | -0.028 |
| 8963 | 0.009 | 0.014 |
| 8964 | -0.032 | 0.122 |
| 8965 | -0.011 | -0.070 |
| 8966 | 0.020 | -0.079 |
| 8967 | 0.000 | -0.067 |
| 8968 | -0.026 | 0.000 |
| 8969 | 0.017 | -0.013 |

| | | |
|---|---|---|
| 8970 | 0.019 | -0.155 |
| 8971 | -0.011 | -0.060 |
| 8972 | 0.014 | -0.028 |
| 8973 | 0.016 | 0.001 |
| 8974 | 0.010 | 0.051 |
| 8975 | -0.047 | 0.016 |
| 8976 | -0.055 | -0.015 |
| 8977 | 0.052 | 0.069 |
| 8978 | 0.035 | -0.029 |
| 8979 | 0.033 | -0.012 |
| 8980 | 0.037 | -0.098 |
| 8981 | 0.005 | 0.057 |
| 8982 | -0.034 | 0.035 |
| 8983 | -0.067 | 0.089 |
| 8984 | 0.004 | 0.030 |
| 8985 | -0.023 | 0.061 |
| 8986 | -0.086 | 0.165 |
| 8987 | -0.001 | 0.038 |
| 8988 | 0.039 | -0.038 |
| 8989 | -0.006 | 0.001 |
| 8990 | 0.038 | -0.058 |
| 8991 | -0.008 | -0.067 |
| 8992 | -0.020 | 0.037 |
| 8993 | 0.017 | 0.045 |
| 8994 | 0.041 | 0.026 |
| 8995 | 0.010 | -0.048 |
| 8996 | -0.003 | -0.046 |
| 8997 | -0.017 | -0.097 |
| 8998 | -0.006 | -0.054 |
| 8999 | 0.064 | -0.118 |
| 9000 | 0.001 | 0.018 |
| 9001 | -0.032 | -0.072 |
| 9002 | 0.034 | -0.031 |
| 9003 | -0.007 | 0.012 |
| 9004 | -0.025 | -0.029 |
| 9005 | 0.027 | 0.030 |
| 9006 | -0.014 | -0.028 |
| 9007 | 0.005 | -0.215 |
| 9008 | 0.033 | 0.148 |
| 9009 | 0.028 | -0.010 |
| 9010 | -0.001 | 0.013 |
| 9011 | -0.039 | -0.002 |
| 9012 | 0.016 | -0.081 |
| 9013 | 0.032 | 0.055 |
| 9014 | -0.006 | 0.037 |
| 9015 | -0.021 | 0.039 |
| 9016 | -0.005 | 0.021 |
| 9017 | -0.031 | -0.021 |
| 9018 | 0.024 | -0.095 |
| 9019 | -0.041 | 0.044 |
| 9020 | -0.019 | 0.098 |

| | | |
|---|---|---|
| 9021 | 0.060 | 0.061 |
| 9022 | -0.004 | 0.030 |
| 9023 | -0.055 | -0.060 |
| 9024 | 0.078 | 0.166 |
| 9025 | -0.039 | 0.010 |
| 9026 | 0.043 | -0.006 |
| 9027 | 0.019 | -0.030 |
| 9028 | -0.067 | -0.097 |
| 9029 | -0.055 | 0.002 |
| 9030 | 0.019 | -0.059 |
| 9031 | 0.026 | -0.044 |
| 9032 | 0.029 | 0.030 |
| 9033 | -0.040 | -0.119 |
| 9034 | -0.072 | -0.034 |
| 9035 | -0.008 | -0.025 |
| 9036 | 0.019 | -0.059 |
| 9037 | 0.028 | 0.069 |
| 9038 | -0.036 | 0.025 |
| 9039 | -0.011 | -0.048 |
| 9040 | 0.014 | 0.126 |
| 9041 | 0.032 | -0.134 |
| 9042 | 0.088 | 0.031 |
| 9043 | 0.069 | -0.066 |
| 9044 | 0.013 | -0.049 |
| 9045 | 0.014 | -0.029 |
| 9046 | -0.024 | -0.018 |
| 9047 | 0.013 | 0.013 |
| 9048 | 0.022 | -0.014 |
| 9049 | -0.007 | 0.017 |
| 9050 | -0.021 | -0.043 |
| 9051 | 0.053 | -0.035 |
| 9052 | -0.062 | 0.171 |
| 9053 | 0.032 | -0.025 |
| 9054 | -0.001 | 0.022 |
| 9055 | -0.011 | -0.009 |
| 9056 | -0.011 | 0.037 |
| 9057 | 0.041 | 0.111 |
| 9058 | -0.005 | 0.101 |
| 9059 | 0.030 | -0.006 |
| 9060 | -0.026 | -0.015 |
| 9061 | 0.092 | 0.045 |
| 9062 | -0.029 | 0.016 |
| 9063 | -0.006 | 0.096 |
| 9064 | -0.005 | -0.011 |
| 9065 | 0.028 | -0.012 |
| 9066 | 0.043 | 0.028 |
| 9067 | -0.045 | 0.077 |
| 9068 | 0.067 | 0.034 |
| 9069 | 0.058 | 0.035 |
| 9070 | -0.090 | -0.053 |
| 9071 | 0.015 | -0.051 |

| | | |
|---|---|---|
| 9072 | -0.035 | 0.089 |
| 9073 | 0.010 | -0.041 |
| 9074 | -0.016 | -0.068 |
| 9075 | -0.030 | -0.041 |
| 9076 | -0.034 | 0.027 |
| 9077 | 0.019 | 0.025 |
| 9078 | -0.057 | -0.024 |
| 9079 | 0.026 | 0.001 |
| 9080 | 0.029 | -0.082 |
| 9081 | 0.000 | -0.066 |
| 9082 | -0.087 | -0.042 |
| 9083 | 0.049 | -0.038 |
| 9084 | 0.041 | -0.103 |
| 9085 | -0.012 | -0.019 |
| 9086 | -0.028 | 0.130 |
| 9087 | 0.032 | -0.044 |
| 9088 | 0.049 | -0.055 |
| 9089 | -0.017 | -0.116 |
| 9090 | 0.008 | -0.063 |
| 9091 | -0.012 | -0.054 |
| 9092 | 0.007 | 0.007 |
| 9093 | 0.010 | -0.029 |
| 9094 | -0.019 | -0.061 |
| 9095 | 0.033 | -0.029 |
| 9096 | 0.012 | -0.076 |
| 9097 | 0.034 | 0.008 |
| 9098 | 0.010 | 0.111 |
| 9099 | 0.068 | -0.029 |
| 9100 | -0.016 | 0.077 |
| 9101 | 0.045 | 0.060 |
| 9102 | -0.062 | 0.009 |
| 9103 | 0.032 | -0.007 |
| 9104 | 0.001 | -0.018 |
| 9105 | 0.018 | 0.000 |
| 9106 | -0.116 | 0.006 |
| 9107 | 0.006 | -0.099 |
| 9108 | 0.018 | -0.006 |
| 9109 | -0.004 | -0.159 |
| 9110 | 0.001 | 0.187 |
| 9111 | 0.030 | -0.007 |
| 9112 | 0.028 | -0.062 |
| 9113 | -0.009 | 0.068 |
| 9114 | 0.000 | 0.035 |
| 9115 | -0.041 | -0.033 |
| 9116 | 0.013 | -0.019 |
| 9117 | 0.009 | -0.026 |
| 9118 | -0.075 | 0.126 |
| 9119 | 0.031 | 0.007 |
| 9120 | 0.038 | 0.095 |
| 9121 | -0.007 | 0.056 |
| 9122 | 0.008 | -0.086 |

| | | |
|---|---|---|
| 9123 | 0.023 | 0.034 |
| 9124 | 0.058 | -0.091 |
| 9125 | -0.018 | 0.022 |
| 9126 | -0.017 | -0.111 |
| 9127 | 0.048 | -0.108 |
| 9128 | 0.004 | -0.021 |
| 9129 | -0.038 | -0.050 |
| 9130 | 0.005 | -0.069 |
| 9131 | 0.005 | 0.034 |
| 9132 | -0.041 | 0.014 |
| 9133 | -0.003 | -0.048 |
| 9134 | 0.065 | -0.011 |
| 9135 | -0.011 | 0.016 |
| 9136 | 0.023 | -0.060 |
| 9137 | -0.015 | 0.071 |
| 9138 | -0.058 | 0.042 |
| 9139 | -0.035 | -0.027 |
| 9140 | 0.025 | 0.006 |
| 9141 | -0.043 | 0.111 |
| 9142 | -0.018 | -0.021 |
| 9143 | 0.000 | 0.029 |
| 9144 | 0.079 | 0.004 |
| 9145 | -0.035 | -0.033 |
| 9146 | -0.054 | -0.036 |
| 9147 | -0.003 | -0.080 |
| 9148 | 0.020 | -0.017 |
| 9149 | -0.004 | 0.100 |
| 9150 | -0.038 | 0.080 |
| 9151 | 0.051 | -0.057 |
| 9152 | -0.045 | -0.033 |
| 9153 | -0.064 | 0.118 |
| 9154 | -0.017 | 0.055 |
| 9155 | -0.013 | -0.098 |
| 9156 | -0.027 | -0.033 |
| 9157 | -0.029 | -0.055 |
| 9158 | -0.009 | -0.047 |
| 9159 | 0.063 | -0.046 |
| 9160 | 0.041 | -0.085 |
| 9161 | -0.028 | -0.069 |
| 9162 | 0.030 | -0.016 |
| 9163 | -0.042 | -0.060 |
| 9164 | -0.022 | -0.123 |
| 9165 | 0.037 | 0.153 |
| 9166 | -0.061 | -0.038 |
| 9167 | 0.011 | 0.054 |
| 9168 | 0.028 | 0.029 |
| 9169 | 0.025 | -0.186 |
| 9170 | 0.014 | 0.044 |
| 9171 | 0.033 | 0.070 |
| 9172 | -0.009 | -0.149 |
| 9173 | 0.019 | 0.017 |

| | | |
|---|---|---|
| 9174 | 0.047 | -0.059 |
| 9175 | -0.032 | 0.017 |
| 9176 | 0.022 | -0.020 |
| 9177 | -0.044 | -0.051 |
| 9178 | 0.033 | 0.057 |
| 9179 | 0.053 | 0.048 |
| 9180 | -0.011 | -0.151 |
| 9181 | 0.076 | -0.036 |
| 9182 | -0.018 | 0.007 |
| 9183 | 0.050 | -0.096 |
| 9184 | -0.014 | 0.074 |
| 9185 | -0.018 | -0.123 |
| 9186 | 0.030 | 0.108 |
| 9187 | -0.051 | -0.096 |
| 9188 | 0.004 | 0.091 |
| 9189 | -0.005 | 0.071 |
| 9190 | 0.006 | -0.009 |
| 9191 | -0.004 | 0.020 |
| 9192 | -0.003 | 0.107 |
| 9193 | -0.009 | -0.087 |
| 9194 | -0.020 | -0.024 |
| 9195 | 0.008 | -0.044 |
| 9196 | 0.017 | -0.030 |
| 9197 | -0.014 | -0.068 |
| 9198 | -0.050 | -0.066 |
| 9199 | -0.075 | 0.059 |
| 9200 | 0.034 | 0.061 |
| 9201 | -0.069 | -0.040 |
| 9202 | 0.052 | 0.021 |
| 9203 | 0.028 | -0.049 |
| 9204 | 0.009 | 0.046 |
| 9205 | -0.024 | -0.021 |
| 9206 | -0.034 | -0.074 |
| 9207 | 0.036 | 0.023 |
| 9208 | 0.027 | -0.008 |
| 9209 | -0.008 | -0.093 |
| 9210 | 0.031 | -0.033 |
| 9211 | -0.028 | 0.029 |
| 9212 | 0.012 | -0.034 |
| 9213 | -0.056 | 0.004 |
| 9214 | 0.029 | -0.035 |
| 9215 | 0.030 | 0.049 |
| 9216 | 0.031 | 0.028 |
| 9217 | -0.084 | 0.021 |
| 9218 | -0.008 | -0.051 |
| 9219 | 0.023 | 0.049 |
| 9220 | 0.013 | 0.057 |
| 9221 | -0.034 | -0.021 |
| 9222 | 0.049 | -0.022 |
| 9223 | 0.039 | 0.121 |
| 9224 | 0.006 | -0.016 |

| | | |
|---|---|---|
| 9225 | -0.063 | 0.170 |
| 9226 | -0.003 | -0.090 |
| 9227 | 0.012 | 0.030 |
| 9228 | -0.029 | -0.020 |
| 9229 | -0.047 | 0.039 |
| 9230 | 0.015 | -0.020 |
| 9231 | -0.001 | -0.073 |
| 9232 | 0.012 | 0.068 |
| 9233 | -0.003 | 0.091 |
| 9234 | -0.030 | -0.104 |
| 9235 | -0.005 | -0.027 |
| 9236 | -0.023 | -0.019 |
| 9237 | -0.041 | -0.039 |
| 9238 | 0.062 | 0.028 |
| 9239 | -0.034 | 0.103 |
| 9240 | -0.057 | -0.027 |
| 9241 | -0.053 | 0.119 |
| 9242 | -0.041 | 0.081 |
| 9243 | 0.045 | 0.028 |
| 9244 | -0.020 | -0.105 |
| 9245 | -0.003 | -0.081 |
| 9246 | -0.056 | 0.104 |
| 9247 | 0.052 | 0.094 |
| 9248 | 0.005 | 0.132 |
| 9249 | 0.027 | -0.022 |
| 9250 | 0.019 | 0.027 |
| 9251 | 0.043 | 0.094 |
| 9252 | -0.056 | 0.038 |
| 9253 | -0.002 | 0.006 |
| 9254 | -0.026 | -0.016 |
| 9255 | -0.038 | 0.006 |
| 9256 | 0.071 | 0.011 |
| 9257 | 0.009 | -0.104 |
| 9258 | 0.003 | -0.001 |
| 9259 | 0.080 | -0.001 |
| 9260 | 0.008 | -0.028 |
| 9261 | -0.011 | -0.115 |
| 9262 | 0.008 | -0.090 |
| 9263 | -0.053 | -0.023 |
| 9264 | 0.001 | 0.038 |
| 9265 | 0.003 | 0.040 |
| 9266 | -0.011 | 0.074 |
| 9267 | -0.016 | -0.001 |
| 9268 | -0.010 | 0.117 |
| 9269 | -0.067 | 0.111 |
| 9270 | -0.061 | -0.086 |
| 9271 | 0.007 | 0.072 |
| 9272 | -0.018 | 0.000 |
| 9273 | -0.029 | 0.069 |
| 9274 | -0.074 | -0.125 |
| 9275 | 0.025 | 0.190 |

| | | |
|---|---|---|
| 9276 | -0.006 | -0.030 |
| 9277 | -0.013 | 0.016 |
| 9278 | -0.034 | 0.027 |
| 9279 | -0.027 | -0.044 |
| 9280 | -0.008 | 0.060 |
| 9281 | -0.039 | -0.089 |
| 9282 | 0.007 | 0.018 |
| 9283 | 0.044 | 0.066 |
| 9284 | -0.010 | -0.003 |
| 9285 | 0.041 | -0.057 |
| 9286 | 0.015 | -0.074 |
| 9287 | 0.002 | -0.008 |
| 9288 | -0.039 | -0.023 |
| 9289 | -0.023 | 0.021 |
| 9290 | -0.006 | -0.038 |
| 9291 | -0.009 | -0.057 |
| 9292 | -0.015 | 0.032 |
| 9293 | 0.009 | -0.035 |
| 9294 | -0.017 | 0.073 |
| 9295 | 0.024 | 0.077 |
| 9296 | 0.045 | 0.062 |
| 9297 | -0.022 | -0.014 |
| 9298 | 0.044 | 0.044 |
| 9299 | 0.046 | 0.164 |
| 9300 | 0.049 | 0.020 |
| 9301 | 0.000 | 0.085 |
| 9302 | -0.010 | 0.042 |
| 9303 | -0.059 | -0.086 |
| 9304 | 0.013 | -0.024 |
| 9305 | 0.009 | 0.058 |
| 9306 | -0.065 | -0.041 |
| 9307 | 0.062 | -0.054 |
| 9308 | 0.016 | 0.017 |
| 9309 | 0.016 | -0.104 |
| 9310 | -0.001 | -0.068 |
| 9311 | 0.031 | -0.065 |
| 9312 | -0.008 | -0.055 |
| 9313 | -0.058 | -0.048 |
| 9314 | -0.039 | 0.122 |
| 9315 | -0.042 | -0.078 |
| 9316 | -0.024 | -0.018 |
| 9317 | -0.014 | 0.063 |
| 9318 | 0.029 | 0.027 |
| 9319 | -0.008 | 0.006 |
| 9320 | -0.009 | 0.053 |
| 9321 | -0.019 | 0.052 |
| 9322 | -0.033 | -0.047 |
| 9323 | -0.030 | 0.036 |
| 9324 | 0.011 | 0.026 |
| 9325 | 0.039 | 0.010 |
| 9326 | 0.059 | -0.054 |

| | | |
|---|---|---|
| 9327 | 0.022 | 0.026 |
| 9328 | -0.011 | 0.060 |
| 9329 | 0.051 | -0.041 |
| 9330 | 0.009 | -0.094 |
| 9331 | 0.038 | 0.068 |
| 9332 | 0.014 | 0.096 |
| 9333 | -0.041 | -0.027 |
| 9334 | -0.014 | 0.027 |
| 9335 | -0.023 | 0.023 |
| 9336 | 0.010 | 0.015 |
| 9337 | -0.026 | 0.030 |
| 9338 | 0.077 | -0.089 |
| 9339 | -0.071 | 0.001 |
| 9340 | -0.041 | -0.061 |
| 9341 | 0.059 | -0.023 |
| 9342 | -0.019 | -0.010 |
| 9343 | 0.033 | 0.031 |
| 9344 | -0.027 | 0.091 |
| 9345 | -0.006 | 0.005 |
| 9346 | -0.026 | -0.021 |
| 9347 | 0.027 | 0.015 |
| 9348 | 0.003 | 0.056 |
| 9349 | -0.042 | -0.023 |
| 9350 | 0.032 | 0.108 |
| 9351 | 0.037 | 0.029 |
| 9352 | -0.010 | 0.078 |
| 9353 | 0.024 | 0.066 |
| 9354 | 0.015 | 0.006 |
| 9355 | 0.004 | -0.023 |
| 9356 | 0.032 | -0.187 |
| 9357 | -0.005 | 0.071 |
| 9358 | -0.008 | -0.016 |
| 9359 | 0.035 | -0.114 |
| 9360 | 0.007 | 0.067 |
| 9361 | -0.066 | 0.078 |
| 9362 | -0.013 | 0.115 |
| 9363 | -0.030 | -0.011 |
| 9364 | -0.001 | 0.118 |
| 9365 | -0.032 | 0.028 |
| 9366 | 0.026 | 0.030 |
| 9367 | 0.012 | 0.031 |
| 9368 | -0.038 | 0.073 |
| 9369 | 0.015 | 0.016 |
| 9370 | 0.044 | -0.014 |
| 9371 | -0.015 | -0.043 |
| 9372 | -0.009 | 0.085 |
| 9373 | -0.062 | -0.034 |
| 9374 | 0.059 | 0.061 |
| 9375 | -0.026 | -0.089 |
| 9376 | -0.006 | 0.016 |
| 9377 | -0.081 | -0.003 |

| | | |
|---|---|---|
| 9378 | -0.002 | 0.001 |
| 9379 | 0.092 | 0.019 |
| 9380 | 0.014 | 0.030 |
| 9381 | 0.025 | 0.010 |
| 9382 | 0.002 | 0.084 |
| 9383 | -0.009 | 0.097 |
| 9384 | -0.030 | 0.085 |
| 9385 | -0.017 | -0.042 |
| 9386 | 0.033 | -0.002 |
| 9387 | 0.027 | -0.073 |
| 9388 | 0.015 | 0.102 |
| 9389 | -0.021 | 0.015 |
| 9390 | 0.010 | 0.035 |
| 9391 | 0.027 | 0.026 |
| 9392 | 0.030 | 0.090 |
| 9393 | -0.010 | 0.088 |
| 9394 | 0.065 | -0.090 |
| 9395 | -0.017 | -0.061 |
| 9396 | 0.014 | 0.097 |
| 9397 | 0.046 | 0.024 |
| 9398 | -0.029 | 0.066 |
| 9399 | -0.017 | -0.076 |
| 9400 | 0.038 | -0.084 |
| 9401 | -0.022 | -0.012 |
| 9402 | 0.013 | -0.069 |
| 9403 | -0.058 | -0.113 |
| 9404 | -0.020 | 0.100 |
| 9405 | 0.034 | -0.020 |
| 9406 | 0.066 | -0.004 |
| 9407 | 0.005 | -0.016 |
| 9408 | -0.065 | 0.039 |
| 9409 | -0.056 | 0.053 |
| 9410 | 0.001 | -0.098 |
| 9411 | -0.009 | 0.126 |
| 9412 | 0.057 | -0.059 |
| 9413 | -0.043 | 0.018 |
| 9414 | -0.012 | -0.028 |
| 9415 | 0.000 | -0.033 |
| 9416 | 0.007 | -0.075 |
| 9417 | -0.038 | -0.084 |
| 9418 | -0.055 | 0.075 |
| 9419 | 0.048 | -0.128 |
| 9420 | -0.020 | 0.002 |
| 9421 | -0.019 | 0.094 |
| 9422 | 0.031 | 0.157 |
| 9423 | 0.041 | 0.218 |
| 9424 | 0.012 | -0.129 |
| 9425 | -0.012 | -0.058 |
| 9426 | 0.004 | -0.044 |
| 9427 | 0.013 | 0.016 |
| 9428 | -0.031 | 0.057 |

| | | |
|---|---|---|
| 9429 | 0.035 | -0.020 |
| 9430 | -0.046 | -0.006 |
| 9431 | -0.005 | -0.006 |
| 9432 | 0.026 | -0.033 |
| 9433 | 0.020 | -0.102 |
| 9434 | -0.021 | -0.047 |
| 9435 | 0.018 | -0.005 |
| 9436 | 0.007 | -0.018 |
| 9437 | -0.008 | 0.066 |
| 9438 | 0.048 | -0.019 |
| 9439 | -0.005 | 0.057 |
| 9440 | -0.006 | -0.080 |
| 9441 | 0.005 | 0.047 |
| 9442 | 0.004 | 0.049 |
| 9443 | 0.055 | 0.012 |
| 9444 | -0.004 | 0.006 |
| 9445 | 0.015 | 0.014 |
| 9446 | -0.031 | 0.039 |
| 9447 | -0.016 | 0.066 |
| 9448 | 0.083 | -0.123 |
| 9449 | 0.067 | 0.062 |
| 9450 | -0.010 | -0.101 |
| 9451 | 0.021 | -0.033 |
| 9452 | -0.011 | 0.006 |
| 9453 | 0.034 | 0.070 |
| 9454 | 0.033 | -0.022 |
| 9455 | 0.043 | 0.047 |
| 9456 | 0.056 | -0.046 |
| 9457 | -0.033 | -0.037 |
| 9458 | 0.098 | 0.035 |
| 9459 | 0.036 | -0.002 |
| 9460 | -0.011 | -0.041 |
| 9461 | -0.072 | -0.048 |
| 9462 | -0.026 | 0.018 |
| 9463 | -0.006 | -0.044 |
| 9464 | 0.012 | 0.039 |
| 9465 | -0.022 | 0.080 |
| 9466 | 0.027 | 0.078 |
| 9467 | -0.016 | -0.001 |
| 9468 | -0.010 | 0.038 |
| 9469 | 0.039 | -0.007 |
| 9470 | 0.029 | 0.018 |
| 9471 | -0.003 | -0.017 |
| 9472 | 0.019 | 0.033 |
| 9473 | 0.027 | 0.033 |
| 9474 | -0.037 | -0.065 |
| 9475 | 0.052 | -0.047 |
| 9476 | 0.002 | -0.093 |
| 9477 | -0.036 | 0.067 |
| 9478 | 0.046 | 0.110 |
| 9479 | -0.021 | -0.009 |

| | | |
|---|---|---|
| 9480 | -0.006 | -0.054 |
| 9481 | -0.054 | -0.206 |
| 9482 | -0.046 | -0.032 |
| 9483 | -0.028 | -0.131 |
| 9484 | -0.013 | 0.061 |
| 9485 | 0.034 | 0.004 |
| 9486 | -0.017 | 0.049 |
| 9487 | -0.021 | 0.053 |
| 9488 | -0.064 | 0.039 |
| 9489 | -0.020 | 0.135 |
| 9490 | 0.046 | 0.022 |
| 9491 | 0.066 | -0.071 |
| 9492 | 0.008 | 0.054 |
| 9493 | -0.027 | 0.014 |
| 9494 | -0.064 | -0.022 |
| 9495 | -0.027 | 0.111 |
| 9496 | 0.031 | 0.006 |
| 9497 | 0.047 | -0.123 |
| 9498 | -0.001 | 0.028 |
| 9499 | 0.010 | 0.101 |
| 9500 | -0.002 | -0.070 |
| 9501 | -0.041 | 0.090 |
| 9502 | -0.024 | -0.002 |
| 9503 | 0.019 | -0.036 |
| 9504 | -0.015 | -0.085 |
| 9505 | -0.021 | 0.010 |
| 9506 | 0.048 | -0.136 |
| 9507 | -0.033 | -0.145 |
| 9508 | 0.051 | -0.079 |
| 9509 | 0.009 | 0.044 |
| 9510 | 0.006 | 0.063 |
| 9511 | 0.050 | 0.035 |
| 9512 | -0.069 | -0.036 |
| 9513 | 0.027 | -0.022 |
| 9514 | -0.006 | -0.063 |
| 9515 | 0.043 | 0.187 |
| 9516 | -0.061 | 0.113 |
| 9517 | 0.020 | 0.097 |
| 9518 | 0.003 | -0.049 |
| 9519 | -0.035 | 0.057 |
| 9520 | 0.042 | -0.051 |
| 9521 | -0.036 | -0.127 |
| 9522 | 0.059 | -0.066 |
| 9523 | -0.004 | 0.077 |
| 9524 | -0.007 | -0.150 |
| 9525 | -0.017 | 0.025 |
| 9526 | -0.044 | -0.034 |
| 9527 | -0.068 | -0.080 |
| 9528 | 0.020 | -0.028 |
| 9529 | -0.031 | -0.008 |
| 9530 | -0.077 | -0.089 |

| | | |
|---|---|---|
| 9531 | 0.021 | -0.076 |
| 9532 | 0.007 | 0.045 |
| 9533 | 0.018 | 0.055 |
| 9534 | -0.029 | 0.024 |
| 9535 | 0.003 | 0.008 |
| 9536 | -0.033 | 0.094 |
| 9537 | -0.021 | 0.023 |
| 9538 | -0.058 | -0.044 |
| 9539 | -0.025 | 0.050 |
| 9540 | 0.005 | -0.151 |
| 9541 | 0.028 | 0.010 |
| 9542 | -0.003 | -0.028 |
| 9543 | 0.073 | 0.091 |
| 9544 | -0.015 | 0.008 |
| 9545 | -0.025 | 0.020 |
| 9546 | 0.056 | -0.082 |
| 9547 | -0.050 | -0.051 |
| 9548 | 0.047 | 0.044 |
| 9549 | 0.011 | 0.027 |
| 9550 | -0.016 | -0.052 |
| 9551 | 0.009 | 0.017 |
| 9552 | -0.056 | 0.045 |
| 9553 | 0.008 | 0.033 |
| 9554 | 0.003 | -0.038 |
| 9555 | 0.050 | -0.043 |
| 9556 | -0.039 | 0.008 |
| 9557 | 0.041 | -0.039 |
| 9558 | 0.051 | 0.004 |
| 9559 | -0.080 | -0.063 |
| 9560 | 0.074 | -0.001 |
| 9561 | 0.012 | -0.076 |
| 9562 | 0.036 | -0.018 |
| 9563 | 0.045 | -0.062 |
| 9564 | 0.012 | -0.056 |
| 9565 | 0.005 | 0.008 |
| 9566 | -0.026 | -0.084 |
| 9567 | -0.014 | 0.027 |
| 9568 | -0.063 | -0.089 |
| 9569 | 0.005 | -0.074 |
| 9570 | 0.011 | 0.020 |
| 9571 | -0.006 | 0.052 |
| 9572 | 0.030 | -0.021 |
| 9573 | 0.069 | 0.077 |
| 9574 | 0.022 | 0.011 |
| 9575 | 0.040 | -0.100 |
| 9576 | -0.012 | 0.104 |
| 9577 | 0.017 | 0.086 |
| 9578 | -0.003 | -0.080 |
| 9579 | 0.054 | -0.027 |
| 9580 | -0.022 | 0.113 |
| 9581 | 0.007 | -0.094 |

| | | |
|---|---|---|
| 9582 | 0.001 | 0.011 |
| 9583 | 0.034 | 0.173 |
| 9584 | 0.011 | -0.111 |
| 9585 | 0.010 | -0.028 |
| 9586 | -0.028 | -0.156 |
| 9587 | 0.007 | 0.007 |
| 9588 | 0.020 | 0.019 |
| 9589 | -0.016 | -0.038 |
| 9590 | 0.016 | 0.040 |
| 9591 | -0.006 | -0.052 |
| 9592 | 0.030 | -0.002 |
| 9593 | 0.048 | 0.035 |
| 9594 | 0.011 | -0.088 |
| 9595 | 0.046 | -0.121 |
| 9596 | -0.064 | -0.082 |
| 9597 | -0.025 | 0.046 |
| 9598 | -0.023 | 0.026 |
| 9599 | -0.023 | -0.010 |
| 9600 | 0.025 | -0.057 |
| 9601 | -0.002 | 0.044 |
| 9602 | 0.073 | -0.014 |
| 9603 | 0.033 | -0.022 |
| 9604 | -0.045 | 0.035 |
| 9605 | 0.013 | 0.058 |
| 9606 | 0.017 | 0.187 |
| 9607 | -0.017 | -0.042 |
| 9608 | 0.050 | -0.068 |
| 9609 | -0.049 | -0.108 |
| 9610 | -0.026 | 0.024 |
| 9611 | -0.020 | -0.025 |
| 9612 | 0.013 | 0.088 |
| 9613 | -0.005 | -0.006 |
| 9614 | 0.016 | -0.074 |
| 9615 | -0.028 | 0.096 |
| 9616 | 0.028 | -0.049 |
| 9617 | -0.046 | -0.066 |
| 9618 | 0.039 | -0.072 |
| 9619 | -0.014 | 0.032 |
| 9620 | -0.027 | 0.002 |
| 9621 | -0.026 | 0.057 |
| 9622 | 0.000 | -0.011 |
| 9623 | -0.076 | -0.041 |
| 9624 | -0.028 | 0.047 |
| 9625 | -0.024 | 0.004 |
| 9626 | -0.002 | 0.016 |
| 9627 | 0.037 | -0.028 |
| 9628 | -0.008 | -0.141 |
| 9629 | 0.012 | 0.137 |
| 9630 | -0.008 | 0.019 |
| 9631 | -0.053 | 0.007 |
| 9632 | -0.010 | 0.000 |

| | | |
|---|---|---|
| 9633 | -0.021 | -0.097 |
| 9634 | -0.017 | 0.114 |
| 9635 | -0.017 | -0.135 |
| 9636 | 0.026 | 0.065 |
| 9637 | 0.001 | -0.095 |
| 9638 | 0.021 | -0.007 |
| 9639 | 0.048 | -0.116 |
| 9640 | -0.056 | -0.038 |
| 9641 | -0.008 | -0.159 |
| 9642 | 0.031 | 0.010 |
| 9643 | 0.014 | -0.036 |
| 9644 | -0.043 | -0.021 |
| 9645 | 0.056 | 0.087 |
| 9646 | -0.066 | -0.092 |
| 9647 | -0.019 | -0.040 |
| 9648 | -0.011 | 0.008 |
| 9649 | 0.030 | 0.057 |
| 9650 | -0.016 | 0.075 |
| 9651 | -0.022 | -0.066 |
| 9652 | -0.031 | 0.050 |
| 9653 | -0.006 | -0.003 |
| 9654 | -0.011 | 0.023 |
| 9655 | -0.046 | 0.030 |
| 9656 | 0.019 | 0.012 |
| 9657 | 0.004 | -0.011 |
| 9658 | 0.021 | -0.152 |
| 9659 | -0.005 | -0.054 |
| 9660 | -0.023 | 0.040 |
| 9661 | -0.029 | 0.058 |
| 9662 | 0.043 | 0.120 |
| 9663 | -0.022 | 0.032 |
| 9664 | -0.024 | 0.002 |
| 9665 | 0.016 | -0.062 |
| 9666 | 0.000 | 0.026 |
| 9667 | -0.006 | -0.052 |
| 9668 | -0.011 | -0.146 |
| 9669 | 0.028 | 0.161 |
| 9670 | 0.013 | 0.025 |
| 9671 | -0.022 | 0.068 |
| 9672 | 0.010 | 0.072 |
| 9673 | -0.011 | 0.055 |
| 9674 | -0.050 | 0.031 |
| 9675 | -0.011 | -0.046 |
| 9676 | -0.008 | 0.068 |
| 9677 | -0.036 | -0.004 |
| 9678 | -0.025 | -0.038 |
| 9679 | -0.037 | -0.113 |
| 9680 | 0.006 | 0.016 |
| 9681 | -0.045 | 0.064 |
| 9682 | 0.031 | 0.019 |
| 9683 | 0.001 | -0.057 |

| | | |
|---|---|---|
| 9684 | 0.011 | -0.031 |
| 9685 | 0.022 | -0.103 |
| 9686 | 0.008 | -0.208 |
| 9687 | -0.009 | -0.035 |
| 9688 | -0.008 | 0.000 |
| 9689 | 0.003 | 0.057 |
| 9690 | -0.040 | 0.173 |
| 9691 | 0.026 | -0.047 |
| 9692 | 0.051 | -0.004 |
| 9693 | 0.029 | -0.102 |
| 9694 | 0.028 | -0.002 |
| 9695 | -0.020 | -0.064 |
| 9696 | 0.025 | 0.002 |
| 9697 | -0.029 | 0.117 |
| 9698 | 0.000 | 0.036 |
| 9699 | 0.016 | 0.039 |
| 9700 | 0.001 | 0.000 |
| 9701 | 0.045 | 0.033 |
| 9702 | -0.011 | -0.022 |
| 9703 | -0.011 | 0.126 |
| 9704 | 0.064 | -0.168 |
| 9705 | 0.045 | -0.112 |
| 9706 | 0.032 | -0.028 |
| 9707 | 0.027 | -0.014 |
| 9708 | -0.023 | 0.069 |
| 9709 | 0.079 | -0.025 |
| 9710 | -0.024 | -0.007 |
| 9711 | -0.078 | 0.159 |
| 9712 | -0.056 | 0.107 |
| 9713 | -0.044 | 0.010 |
| 9714 | -0.013 | 0.043 |
| 9715 | -0.084 | -0.048 |
| 9716 | -0.025 | -0.027 |
| 9717 | 0.024 | -0.067 |
| 9718 | -0.016 | -0.036 |
| 9719 | 0.018 | -0.015 |
| 9720 | -0.083 | -0.031 |
| 9721 | 0.017 | 0.034 |
| 9722 | 0.020 | 0.034 |
| 9723 | -0.022 | -0.065 |
| 9724 | 0.065 | 0.097 |
| 9725 | 0.023 | 0.029 |
| 9726 | -0.025 | -0.076 |
| 9727 | 0.020 | 0.109 |
| 9728 | 0.030 | 0.100 |
| 9729 | -0.001 | -0.046 |
| 9730 | 0.055 | -0.046 |
| 9731 | -0.029 | -0.058 |
| 9732 | 0.033 | 0.066 |
| 9733 | -0.001 | -0.007 |
| 9734 | 0.011 | -0.018 |

| | | |
|---|---|---|
| 9735 | -0.063 | 0.083 |
| 9736 | -0.009 | 0.029 |
| 9737 | -0.022 | 0.010 |
| 9738 | -0.013 | -0.007 |
| 9739 | 0.001 | 0.031 |
| 9740 | -0.003 | -0.068 |
| 9741 | -0.003 | 0.001 |
| 9742 | 0.022 | 0.031 |
| 9743 | 0.000 | -0.056 |
| 9744 | 0.013 | -0.009 |
| 9745 | 0.013 | -0.015 |
| 9746 | 0.060 | -0.038 |
| 9747 | -0.058 | 0.005 |
| 9748 | -0.067 | 0.005 |
| 9749 | 0.024 | 0.024 |
| 9750 | 0.009 | 0.059 |
| 9751 | -0.041 | 0.088 |
| 9752 | 0.030 | -0.067 |
| 9753 | 0.001 | 0.032 |
| 9754 | -0.020 | 0.127 |
| 9755 | -0.032 | -0.049 |
| 9756 | -0.024 | 0.007 |
| 9757 | 0.014 | -0.118 |
| 9758 | 0.045 | 0.067 |
| 9759 | 0.026 | -0.025 |
| 9760 | 0.000 | 0.060 |
| 9761 | 0.037 | 0.038 |
| 9762 | 0.001 | -0.062 |
| 9763 | -0.064 | 0.045 |
| 9764 | -0.024 | -0.020 |
| 9765 | 0.019 | -0.061 |
| 9766 | -0.035 | 0.029 |
| 9767 | -0.011 | 0.097 |
| 9768 | 0.036 | 0.013 |
| 9769 | -0.040 | 0.038 |
| 9770 | -0.006 | 0.030 |
| 9771 | 0.035 | -0.128 |
| 9772 | 0.076 | -0.019 |
| 9773 | 0.033 | 0.059 |
| 9774 | -0.049 | -0.049 |
| 9775 | -0.041 | 0.043 |
| 9776 | -0.061 | -0.022 |
| 9777 | -0.058 | 0.098 |
| 9778 | 0.043 | 0.003 |
| 9779 | 0.044 | 0.162 |
| 9780 | 0.015 | 0.051 |
| 9781 | -0.009 | -0.049 |
| 9782 | -0.050 | 0.039 |
| 9783 | -0.055 | -0.060 |
| 9784 | 0.001 | -0.137 |
| 9785 | -0.007 | 0.047 |

| | | |
|---|---|---|
| 9786 | 0.024 | -0.010 |
| 9787 | 0.063 | -0.008 |
| 9788 | -0.044 | -0.036 |
| 9789 | -0.019 | 0.020 |
| 9790 | 0.034 | -0.046 |
| 9791 | -0.038 | 0.009 |
| 9792 | -0.008 | -0.031 |
| 9793 | 0.000 | -0.006 |
| 9794 | -0.012 | 0.146 |
| 9795 | 0.021 | 0.029 |
| 9796 | 0.070 | 0.139 |
| 9797 | 0.023 | -0.115 |
| 9798 | 0.033 | 0.092 |
| 9799 | -0.025 | -0.080 |
| 9800 | -0.007 | -0.045 |
| 9801 | -0.035 | -0.031 |
| 9802 | -0.014 | -0.081 |
| 9803 | 0.055 | -0.074 |
| 9804 | -0.002 | -0.058 |
| 9805 | -0.054 | 0.058 |
| 9806 | -0.022 | -0.061 |
| 9807 | 0.001 | 0.102 |
| 9808 | 0.031 | -0.071 |
| 9809 | -0.023 | 0.036 |
| 9810 | -0.004 | 0.054 |
| 9811 | -0.016 | 0.021 |
| 9812 | -0.004 | -0.091 |
| 9813 | -0.014 | 0.046 |
| 9814 | -0.025 | 0.142 |
| 9815 | -0.045 | -0.042 |
| 9816 | -0.013 | -0.113 |
| 9817 | 0.050 | -0.063 |
| 9818 | 0.011 | -0.034 |
| 9819 | 0.006 | 0.129 |
| 9820 | -0.059 | -0.008 |
| 9821 | -0.017 | -0.022 |
| 9822 | 0.048 | -0.133 |
| 9823 | 0.030 | 0.063 |
| 9824 | 0.009 | 0.024 |
| 9825 | -0.034 | 0.155 |
| 9826 | 0.014 | -0.070 |
| 9827 | -0.011 | -0.105 |
| 9828 | -0.036 | -0.080 |
| 9829 | -0.038 | 0.002 |
| 9830 | -0.001 | 0.035 |
| 9831 | 0.021 | -0.009 |
| 9832 | -0.004 | 0.012 |
| 9833 | 0.029 | 0.040 |
| 9834 | -0.031 | -0.108 |
| 9835 | -0.037 | 0.048 |
| 9836 | 0.012 | -0.010 |

| | | |
|---|---|---|
| 9837 | 0.029 | -0.007 |
| 9838 | -0.014 | -0.073 |
| 9839 | 0.024 | 0.037 |
| 9840 | -0.054 | 0.015 |
| 9841 | -0.002 | 0.077 |
| 9842 | 0.003 | -0.077 |
| 9843 | -0.036 | 0.019 |
| 9844 | -0.064 | -0.010 |
| 9845 | 0.000 | 0.032 |
| 9846 | -0.027 | -0.084 |
| 9847 | -0.004 | -0.062 |
| 9848 | 0.051 | -0.018 |
| 9849 | 0.040 | 0.033 |
| 9850 | 0.012 | -0.066 |
| 9851 | 0.050 | -0.167 |
| 9852 | -0.022 | 0.077 |
| 9853 | 0.029 | -0.011 |
| 9854 | 0.043 | -0.002 |
| 9855 | -0.019 | -0.033 |
| 9856 | -0.037 | 0.058 |
| 9857 | 0.022 | 0.052 |
| 9858 | 0.062 | 0.032 |
| 9859 | -0.018 | -0.086 |
| 9860 | 0.036 | 0.085 |
| 9861 | -0.034 | 0.007 |
| 9862 | 0.019 | -0.061 |
| 9863 | 0.031 | -0.021 |
| 9864 | 0.008 | -0.007 |
| 9865 | -0.076 | 0.100 |
| 9866 | -0.029 | 0.029 |
| 9867 | -0.038 | 0.045 |
| 9868 | 0.006 | -0.063 |
| 9869 | 0.001 | -0.079 |
| 9870 | -0.057 | -0.048 |
| 9871 | 0.020 | -0.022 |
| 9872 | -0.062 | 0.050 |
| 9873 | -0.020 | 0.077 |
| 9874 | 0.043 | -0.105 |
| 9875 | 0.031 | 0.133 |
| 9876 | -0.038 | 0.028 |
| 9877 | 0.027 | -0.141 |
| 9878 | 0.005 | -0.098 |
| 9879 | -0.002 | 0.096 |
| 9880 | -0.043 | -0.040 |
| 9881 | -0.026 | 0.013 |
| 9882 | 0.016 | 0.021 |
| 9883 | -0.021 | 0.111 |
| 9884 | 0.036 | 0.006 |
| 9885 | 0.017 | 0.033 |
| 9886 | -0.020 | 0.127 |
| 9887 | -0.020 | 0.026 |

| | | |
|---|---|---|
| 9888 | 0.009 | 0.024 |
| 9889 | -0.031 | -0.105 |
| 9890 | -0.031 | 0.027 |
| 9891 | 0.033 | -0.034 |
| 9892 | -0.052 | -0.011 |
| 9893 | 0.016 | 0.051 |
| 9894 | 0.003 | -0.005 |
| 9895 | -0.010 | 0.001 |
| 9896 | -0.057 | 0.088 |
| 9897 | -0.004 | -0.044 |
| 9898 | -0.012 | -0.061 |
| 9899 | 0.028 | 0.004 |
| 9900 | -0.038 | -0.066 |
| 9901 | 0.011 | -0.069 |
| 9902 | -0.049 | 0.023 |
| 9903 | 0.002 | -0.110 |
| 9904 | -0.008 | -0.015 |
| 9905 | 0.041 | -0.046 |
| 9906 | -0.046 | -0.017 |
| 9907 | 0.049 | 0.138 |
| 9908 | 0.020 | 0.025 |
| 9909 | 0.043 | 0.034 |
| 9910 | -0.025 | -0.054 |
| 9911 | -0.031 | -0.102 |
| 9912 | 0.021 | 0.027 |
| 9913 | 0.042 | 0.025 |
| 9914 | -0.017 | 0.047 |
| 9915 | -0.006 | -0.015 |
| 9916 | 0.020 | -0.114 |
| 9917 | -0.013 | -0.022 |
| 9918 | -0.024 | -0.119 |
| 9919 | 0.058 | -0.010 |
| 9920 | 0.015 | -0.053 |
| 9921 | 0.095 | -0.042 |
| 9922 | 0.037 | -0.046 |
| 9923 | 0.029 | -0.063 |
| 9924 | -0.038 | -0.029 |
| 9925 | 0.022 | -0.030 |
| 9926 | -0.074 | -0.047 |
| 9927 | 0.023 | -0.010 |
| 9928 | -0.049 | -0.061 |
| 9929 | 0.015 | -0.056 |
| 9930 | -0.013 | -0.024 |
| 9931 | 0.037 | 0.107 |
| 9932 | -0.039 | 0.096 |
| 9933 | -0.041 | -0.007 |
| 9934 | -0.072 | -0.103 |
| 9935 | 0.041 | 0.192 |
| 9936 | -0.010 | 0.057 |
| 9937 | -0.031 | 0.120 |
| 9938 | 0.013 | -0.073 |

| | | |
|---|---|---|
| 9939 | -0.009 | -0.088 |
| 9940 | 0.026 | 0.013 |
| 9941 | 0.063 | 0.090 |
| 9942 | 0.021 | -0.007 |
| 9943 | -0.019 | 0.005 |
| 9944 | 0.039 | 0.079 |
| 9945 | 0.004 | 0.007 |
| 9946 | -0.008 | 0.100 |
| 9947 | -0.092 | 0.042 |
| 9948 | 0.011 | 0.023 |
| 9949 | 0.045 | -0.098 |
| 9950 | 0.014 | 0.040 |
| 9951 | -0.010 | 0.037 |
| 9952 | 0.004 | 0.095 |
| 9953 | 0.017 | 0.008 |
| 9954 | -0.012 | -0.011 |
| 9955 | -0.018 | 0.030 |
| 9956 | 0.017 | -0.129 |
| 9957 | 0.018 | -0.025 |
| 9958 | 0.004 | 0.008 |
| 9959 | -0.074 | -0.046 |
| 9960 | -0.016 | -0.130 |
| 9961 | -0.005 | -0.065 |
| 9962 | 0.017 | -0.118 |
| 9963 | -0.055 | 0.021 |
| 9964 | -0.044 | -0.084 |
| 9965 | 0.009 | 0.079 |
| 9966 | 0.014 | -0.062 |
| 9967 | -0.015 | 0.084 |
| 9968 | 0.010 | -0.075 |
| 9969 | 0.039 | 0.050 |
| 9970 | -0.034 | 0.034 |
| 9971 | -0.015 | -0.090 |
| 9972 | -0.045 | 0.052 |
| 9973 | 0.008 | 0.086 |
| 9974 | 0.024 | -0.086 |
| 9975 | -0.005 | 0.076 |
| 9976 | -0.097 | 0.026 |
| 9977 | -0.027 | -0.068 |
| 9978 | 0.015 | 0.089 |
| 9979 | -0.012 | -0.041 |
| 9980 | 0.037 | 0.116 |
| 9981 | -0.040 | 0.060 |
| 9982 | -0.025 | -0.102 |
| 9983 | 0.039 | 0.003 |
| 9984 | -0.014 | -0.015 |
| 9985 | -0.062 | -0.007 |
| 9986 | 0.018 | -0.049 |
| 9987 | -0.019 | 0.016 |
| 9988 | 0.044 | -0.017 |
| 9989 | -0.010 | 0.018 |

| | | |
|---|---|---|
| 9990 | 0.021 | 0.029 |
| 9991 | -0.012 | 0.053 |
| 9992 | 0.065 | 0.077 |
| 9993 | -0.020 | 0.080 |
| 9994 | -0.031 | -0.081 |
| 9995 | 0.051 | 0.101 |
| 9996 | 0.006 | 0.062 |
| 9997 | 0.025 | -0.147 |
| 9998 | -0.027 | -0.123 |
| 9999 | -0.001 | -0.058 |
| 10000 | -0.059 | 0.046 |

**Table S10. Correlation values for the abundance of TRiC complex subunits with age in GV oocytes. Age-Correlated Trends: |0.3| ≤ R< |0.5|; p≤0.05. Age-Correlated: R≥ |0.5|, p.adj≤0.2.**

| | Uniprot ID | Protein name | R | P-value | P.adjusted |
|---|---|---|---|---|---|
| | Q99832 | TCPH | -0.578 | 0.002 | 0.159 |
| | P17987 | TCPA | -0.502 | 0.009 | 0.343 |
| | P50990 | TCPQ | -0.419 | 0.033 | 0.442 |
| | P48643 | TCPE | -0.401 | 0.042 | 0.450 |
| | P50991 | TCPD | -0.356 | 0.074 | 0.509 |
| | P78371 | TCPB | -0.365 | 0.067 | 0.509 |
| | P40227 | TCPZ | -0.297 | 0.140 | 0.632 |
| | P49368 | TCPG | 0.003 | 0.989 | 0.999 |
| $R_{mean}$ | | | -0.364 | | |

**Table S11. Identified changes in GO (Biological Processes, BP; Cellular Compartments, CC) and KEGG pathways. GV oocytes: Young *vs.* AMA**

| | Terms | Overlap | Ratio | P-value | P.adj | Log2 (FC) | Proteins |
|---|---|---|---|---|---|---|---|
| BP | Wnt signaling pathway, planar cell polarity pathway (GO:0060071) | 42/85 | 0.5 | 0.0002 | 0.037 | 0.131 | PSD10; PSD12; PSD11; PSDE; PSD13; CLH1; PSA7; PSMD8; CDC42; PSB6; PSB7; PSMD6; PSB4; PSMD7; PSB5; PSMD4; PSB2; PSMD5; PSB3; PSMD2; PSMD3; PSB1; PSMD1; RAC1; PSMF1; AP2M1; AP2B1; RHOA; PSA5; PSA6; PSA3; PRS8; PSA4; PRS10; PSA1; PRS6A; PSA2; PRS6B; PRS4; PRS7; PSME1; PROF1 |
| | regulation of establishment of planar polarity (GO:0090175) | 42/88 | 0.5 | 0.0002 | 0.037 | 0.131 | PSD10; PSD12; PSD11; PSDE; PSD13; CLH1; PSA7; PSMD8; CDC42; PSB6; PSB7; PSMD6; PSB4; PSMD7; PSB5; PSMD4; PSB2; PSMD5; PSB3; PSMD2; PSMD3; PSB1; PSMD1; RAC1; PSMF1; AP2M1; AP2B1; RHOA; PSA5; PSA6; PSA3; PRS8; PSA4; PRS10; PSA1; PRS6A; PSA2; PRS6B; PRS4; PRS7; PSME1; PROF1 |
| | positive regulation of telomerase RNA localization to Cajal body (GO:1904874) | 10/15 | 0.7 | 0.0001 | 0.031 | 0.249 | TCPG; TCPZ; TCPB; RUVB2; RUVB1; TCPA; TCPQ; TCPH; TCPE; TCPD |
| | Arp2/3 complex-mediated actin nucleation (GO:0034314) | 08/15 | 0.5 | 0.0003 | 0.039 | 0.281 | ARP3; ARP2; ARPC2; ARPC3; ARC1B; ARP5L; ARPC4; ARPC5 |
| CC | ficolin-1-rich granule lumen (GO:1904813) | 65/123 | 0.528455285 | 0.0009 | 0.021 | 0.125 | G6PI; DDX3X; HS90B; A1AT; ACPH; SRP14; PSMD6; PSMD7; ACTY; FRIH; PSMD2; PSMD3; COTL1; OSTF1; GLYG; CATD; PGM1; CATB; ARP2; HS90A; DYLT1; PGAM1; K2C1; NDKB; MIF; PSA5; ACLY; KPYM; PSA2; ALDOC; ALDOA; PPIA; IMB1; VINC; PA1B2; CYTB; PSD12; TERA; ASAH1; PSD11; PSDE; PSD13; GSTP1; MVP; GNS; PSB7; PRDX4; CAND1; PSB1; MK01; LKHA4; TCPQ; HSP7C; XRCC6; GELS; IDHC; HUWE1; HAP28; ARPC5; EF2; PFKAL; PRS6A; BGAL; IMDH2; PRS7 |

**Table S12. Proteins showing correlation trends with age in MII oocytes. (|0.3| ≤ R< |0.5|; p≤0.05)**

| Uniprot ID | Protein name | R | P-value | P.adjusted |
|---|---|---|---|---|
| P09972 | ALDOC | -0.567 | 0.002 | 0.262 |
| O60825 | F262 | -0.555 | 0.002 | 0.288 |
| Q9BV73 | CP250 | -0.554 | 0.002 | 0.288 |
| P45974 | UBP5 | -0.544 | 0.003 | 0.331 |
| Q92905 | CSN5 | -0.532 | 0.004 | 0.364 |
| Q9HCN8 | SDF2L | -0.509 | 0.006 | 0.455 |
| Q15814 | TBCC | -0.505 | 0.006 | 0.462 |
| Q9P2B4 | CT2NL | -0.500 | 0.009 | 0.492 |
| Q99805 | TM9S2 | -0.494 | 0.008 | 0.485 |
| P15121 | ALDR | -0.490 | 0.008 | 0.485 |
| Q9H3K6 | BOLA2 | -0.488 | 0.008 | 0.485 |
| Q9UJ70-2 | NAGK | -0.482 | 0.011 | 0.502 |
| O60825-2 | F262 | -0.477 | 0.018 | 0.570 |
| P35080-2 | PROF2 | -0.474 | 0.011 | 0.502 |
| P06748 | NPM | -0.471 | 0.011 | 0.502 |
| P54652 | HSP72 | -0.467 | 0.012 | 0.502 |
| Q92506 | DHB8 | -0.467 | 0.016 | 0.549 |
| O95433 | AHSA1 | -0.465 | 0.013 | 0.502 |
| P60520 | GBRL2 | -0.464 | 0.013 | 0.502 |
| Q9UHD1 | CHRD1 | -0.463 | 0.013 | 0.502 |
| P61077 | UB2D3 | -0.459 | 0.014 | 0.511 |
| P50991 | TCPD | -0.451 | 0.016 | 0.549 |
| Q13310 | PABP4 | -0.446 | 0.017 | 0.549 |
| P46379 | BAG6 | -0.428 | 0.023 | 0.643 |
| Q8NA58 | PNDC1 | -0.421 | 0.032 | 0.676 |
| Q86TP1 | PRUN1 | -0.420 | 0.041 | 0.676 |
| Q8TD30 | ALAT2 | -0.419 | 0.026 | 0.651 |
| P62805 | H4 | -0.418 | 0.027 | 0.656 |
| Q9H832 | UBE2Z | -0.416 | 0.038 | 0.676 |
| P22061-2 | PIMT | -0.407 | 0.032 | 0.676 |
| Q99653 | CHP1 | -0.405 | 0.045 | 0.676 |
| Q969H8 | MYDGF | -0.403 | 0.034 | 0.676 |
| Q13368 | MPP3 | -0.402 | 0.038 | 0.676 |
| Q6ZMR3 | LDH6A | -0.398 | 0.036 | 0.676 |
| P60842 | IF4A1 | -0.396 | 0.037 | 0.676 |
| Q10471 | GALT2 | -0.393 | 0.042 | 0.676 |
| P42126 | ECI1 | -0.392 | 0.039 | 0.676 |
| P61088 | UBE2N | -0.389 | 0.045 | 0.676 |
| P62191 | PRS4 | -0.388 | 0.041 | 0.676 |
| P26358-2 | DNMT1 | -0.387 | 0.042 | 0.676 |
| P22234 | PUR6 | -0.386 | 0.043 | 0.676 |
| P04075 | ALDOA | -0.384 | 0.043 | 0.676 |
| O60701 | UGDH | -0.384 | 0.044 | 0.676 |
| Q96BN8 | OTUL | -0.379 | 0.047 | 0.676 |
| Q13561 | DCTN2 | -0.378 | 0.047 | 0.676 |
| P02792 | FRIL | -0.377 | 0.048 | 0.676 |
| Q96TC7 | RMD3 | -0.375 | 0.049 | 0.676 |
| P20618 | PSB1 | -0.373 | 0.050 | 0.676 |

| | | | | |
|---|---|---|---|---|
| P62328 | TYB4 | 0.375 | 0.049 | 0.676 |
| Q9Y294 | ASF1A | 0.376 | 0.049 | 0.676 |
| O95292 | VAPB | 0.377 | 0.048 | 0.676 |
| P04004 | VTNC | 0.381 | 0.046 | 0.676 |
| P63172 | DYLT1 | 0.381 | 0.050 | 0.676 |
| P19823 | ITIH2 | 0.388 | 0.042 | 0.676 |
| P60852 | ZP1 | 0.388 | 0.041 | 0.676 |
| P13473 | LAMP2 | 0.391 | 0.048 | 0.676 |
| P60033 | CD81 | 0.391 | 0.044 | 0.676 |
| O00232 | PSD12 | 0.395 | 0.046 | 0.676 |
| P10909 | CLUS | 0.395 | 0.038 | 0.676 |
| P53618 | COPB | 0.395 | 0.038 | 0.676 |
| A6NNZ2 | TBB8B | 0.395 | 0.037 | 0.676 |
| Q8IWA4 | MFN1 | 0.398 | 0.040 | 0.676 |
| Q92542 | NICA | 0.399 | 0.048 | 0.676 |
| Q9HD67 | MYO10 | 0.399 | 0.039 | 0.676 |
| P42025 | ACTY | 0.400 | 0.035 | 0.676 |
| Q9P1Z2 | CACO1 | 0.400 | 0.039 | 0.676 |
| P05556 | ITB1 | 0.400 | 0.035 | 0.676 |
| P13611 | CSPG2 | 0.401 | 0.034 | 0.676 |
| Q14677 | EPN4 | 0.401 | 0.034 | 0.676 |
| Q8N5K1 | CISD2 | 0.402 | 0.042 | 0.676 |
| P43121 | MUC18 | 0.403 | 0.041 | 0.676 |
| P41222 | PTGDS | 0.407 | 0.032 | 0.676 |
| P07942 | LAMB1 | 0.419 | 0.026 | 0.651 |
| Q14203 | DCTN1 | 0.422 | 0.028 | 0.671 |
| Q9BRQ6 | MIC25 | 0.423 | 0.025 | 0.647 |
| P82909 | KGD4 | 0.425 | 0.024 | 0.645 |
| P06733 | ENOA | 0.427 | 0.024 | 0.645 |
| O43504 | LTOR5 | 0.427 | 0.026 | 0.651 |
| P04114 | APOB | 0.432 | 0.022 | 0.617 |
| P40926 | MDHM | 0.433 | 0.021 | 0.617 |
| Q99816 | TS101 | 0.441 | 0.019 | 0.570 |
| Q9NX76 | CKLF6 | 0.449 | 0.024 | 0.645 |
| P21754 | ZP3 | 0.458 | 0.014 | 0.511 |
| P47914 | RL29 | 0.460 | 0.014 | 0.511 |
| Q9UK76 | JUPI1 | 0.464 | 0.017 | 0.549 |
| Q14451 | GRB7 | 0.465 | 0.013 | 0.502 |
| Q6P1M0 | S27A4 | 0.465 | 0.017 | 0.549 |
| Q16658 | FSCN1 | 0.469 | 0.012 | 0.502 |
| Q92530 | PSMF1 | 0.471 | 0.020 | 0.602 |
| P07437 | TBB5 | 0.473 | 0.013 | 0.502 |
| Q8IZA3 | H18 | 0.486 | 0.009 | 0.485 |
| P04899 | GNAI2 | 0.491 | 0.008 | 0.485 |
| P02748 | CO9 | 0.494 | 0.009 | 0.485 |
| Q12836 | ZP4 | 0.494 | 0.008 | 0.485 |
| Q13505 | MTX1 | 0.515 | 0.005 | 0.455 |
| P46776 | RL27A | 0.519 | 0.006 | 0.455 |
| P09497-2 | CLCB | 0.532 | 0.004 | 0.364 |
| P18084 | ITB5 | 0.542 | 0.005 | 0.455 |
| O60493 | SNX3 | 0.570 | 0.002 | 0.262 |

**Table S13. Correlation values for the abundance of TRiC complex subunits with age in MII oocytes. Age-Correlated Trends: |0.3| ≤ R< |0.5|; p≤0.05. Age-Correlated: R≥ |0.5|, p.adj≤0.2.**

| Uniprot ID | Protein name | R | P-value | P.adjusted |
|---|---|---|---|---|
| P50991 | TCPD | -0.451 | 0.016 | 0.549 |
| P48643 | TCPE | -0.358 | 0.061 | 0.696 |
| P49368 | TCPG | -0.311 | 0.107 | 0.723 |
| P78371 | TCPB | -0.341 | 0.076 | 0.723 |
| P50990 | TCPQ | -0.294 | 0.128 | 0.744 |
| P17987 | TCPA | -0.222 | 0.256 | 0.801 |
| P40227 | TCPZ | -0.065 | 0.744 | 0.967 |
| Q99832 | TCPH | -0.055 | 0.781 | 0.968 |
| $R_{mean}$ | | -0.262 | | |

**Table S14. Permutation Results for randomized correlations of Rmean**

| | MII |
|---|---|
| Randomization # | Rmean for TRiC subunits |
| 1 | -0.068 |
| 2 | 0.030 |
| 3 | -0.027 |
| 4 | 0.046 |
| 5 | 0.105 |
| 6 | -0.009 |
| 7 | 0.026 |
| 8 | 0.018 |
| 9 | -0.073 |
| 10 | -0.003 |
| 11 | -0.009 |
| 12 | 0.040 |
| 13 | 0.001 |
| 14 | -0.029 |
| 15 | -0.024 |
| 16 | 0.060 |
| 17 | -0.004 |
| 18 | 0.135 |
| 19 | 0.091 |
| 20 | 0.007 |
| 21 | 0.089 |
| 22 | -0.141 |
| 23 | -0.042 |
| 24 | -0.013 |
| 25 | -0.065 |
| 26 | -0.027 |
| 27 | 0.011 |
| 28 | -0.008 |
| 29 | -0.069 |
| 30 | -0.086 |
| 31 | 0.061 |
| 32 | 0.089 |
| 33 | 0.091 |
| 34 | -0.066 |
| 35 | 0.049 |
| 36 | 0.027 |
| 37 | -0.087 |
| 38 | -0.121 |
| 39 | -0.081 |
| 40 | -0.016 |
| 41 | -0.180 |
| 42 | -0.107 |
| 43 | -0.031 |
| 44 | -0.007 |
| 45 | -0.020 |
| 46 | 0.149 |

| | |
|---|---|
| 47 | -0.063 |
| 48 | 0.046 |
| 49 | 0.015 |
| 50 | -0.144 |
| 51 | 0.035 |
| 52 | 0.158 |
| 53 | 0.089 |
| 54 | 0.084 |
| 55 | -0.079 |
| 56 | -0.083 |
| 57 | 0.044 |
| 58 | 0.019 |
| 59 | -0.001 |
| 60 | -0.012 |
| 61 | 0.078 |
| 62 | 0.049 |
| 63 | 0.018 |
| 64 | 0.074 |
| 65 | -0.083 |
| 66 | -0.008 |
| 67 | 0.123 |
| 68 | 0.016 |
| 69 | 0.027 |
| 70 | 0.046 |
| 71 | 0.114 |
| 72 | -0.124 |
| 73 | 0.000 |
| 74 | 0.033 |
| 75 | -0.121 |
| 76 | 0.034 |
| 77 | 0.043 |
| 78 | -0.015 |
| 79 | -0.002 |
| 80 | -0.001 |
| 81 | -0.026 |
| 82 | -0.076 |
| 83 | -0.095 |
| 84 | 0.043 |
| 85 | -0.011 |
| 86 | 0.080 |
| 87 | -0.135 |
| 88 | 0.157 |
| 89 | -0.059 |
| 90 | -0.011 |
| 91 | -0.038 |
| 92 | -0.009 |
| 93 | -0.043 |
| 94 | 0.006 |
| 95 | 0.070 |
| 96 | -0.103 |

| | |
|---|---|
| 97 | -0.017 |
| 98 | 0.024 |
| 99 | 0.061 |
| 100 | 0.050 |
| 101 | -0.043 |
| 102 | 0.090 |
| 103 | -0.026 |
| 104 | 0.000 |
| 105 | 0.100 |
| 106 | -0.094 |
| 107 | 0.015 |
| 108 | -0.074 |
| 109 | -0.050 |
| 110 | -0.076 |
| 111 | -0.019 |
| 112 | -0.065 |
| 113 | 0.047 |
| 114 | -0.187 |
| 115 | -0.069 |
| 116 | 0.098 |
| 117 | 0.061 |
| 118 | -0.027 |
| 119 | 0.134 |
| 120 | 0.073 |
| 121 | 0.074 |
| 122 | 0.013 |
| 123 | -0.056 |
| 124 | -0.012 |
| 125 | 0.028 |
| 126 | 0.006 |
| 127 | 0.024 |
| 128 | -0.011 |
| 129 | 0.039 |
| 130 | -0.036 |
| 131 | 0.002 |
| 132 | -0.089 |
| 133 | 0.044 |
| 134 | -0.073 |
| 135 | -0.066 |
| 136 | 0.005 |
| 137 | 0.026 |
| 138 | -0.010 |
| 139 | 0.105 |
| 140 | 0.109 |
| 141 | -0.044 |
| 142 | 0.001 |
| 143 | -0.008 |
| 144 | -0.004 |
| 145 | 0.012 |
| 146 | 0.029 |

| | |
|---|---|
| 147 | -0.064 |
| 148 | 0.004 |
| 149 | 0.008 |
| 150 | -0.141 |
| 151 | -0.014 |
| 152 | 0.120 |
| 153 | -0.045 |
| 154 | 0.012 |
| 155 | -0.081 |
| 156 | 0.061 |
| 157 | -0.086 |
| 158 | -0.022 |
| 159 | -0.090 |
| 160 | -0.024 |
| 161 | 0.019 |
| 162 | 0.031 |
| 163 | -0.004 |
| 164 | 0.085 |
| 165 | -0.003 |
| 166 | 0.036 |
| 167 | 0.037 |
| 168 | -0.047 |
| 169 | -0.019 |
| 170 | -0.020 |
| 171 | -0.045 |
| 172 | 0.066 |
| 173 | -0.047 |
| 174 | -0.013 |
| 175 | -0.015 |
| 176 | 0.008 |
| 177 | -0.017 |
| 178 | -0.135 |
| 179 | -0.103 |
| 180 | -0.022 |
| 181 | -0.013 |
| 182 | -0.070 |
| 183 | -0.054 |
| 184 | 0.051 |
| 185 | 0.069 |
| 186 | -0.013 |
| 187 | -0.005 |
| 188 | -0.129 |
| 189 | 0.072 |
| 190 | 0.027 |
| 191 | 0.067 |
| 192 | 0.035 |
| 193 | -0.043 |
| 194 | 0.038 |
| 195 | -0.092 |
| 196 | 0.003 |

| | |
|---|---|
| 197 | -0.010 |
| 198 | -0.059 |
| 199 | -0.054 |
| 200 | 0.093 |
| 201 | 0.100 |
| 202 | -0.008 |
| 203 | 0.004 |
| 204 | 0.003 |
| 205 | -0.064 |
| 206 | 0.013 |
| 207 | 0.044 |
| 208 | -0.015 |
| 209 | 0.076 |
| 210 | 0.094 |
| 211 | 0.024 |
| 212 | -0.083 |
| 213 | 0.130 |
| 214 | 0.079 |
| 215 | -0.044 |
| 216 | 0.054 |
| 217 | 0.075 |
| 218 | -0.052 |
| 219 | 0.048 |
| 220 | -0.019 |
| 221 | 0.020 |
| 222 | 0.055 |
| 223 | 0.035 |
| 224 | 0.066 |
| 225 | -0.016 |
| 226 | 0.002 |
| 227 | 0.027 |
| 228 | 0.073 |
| 229 | 0.039 |
| 230 | 0.020 |
| 231 | 0.029 |
| 232 | -0.067 |
| 233 | -0.013 |
| 234 | -0.048 |
| 235 | -0.014 |
| 236 | 0.124 |
| 237 | 0.020 |
| 238 | 0.048 |
| 239 | -0.004 |
| 240 | -0.067 |
| 241 | 0.011 |
| 242 | -0.029 |
| 243 | 0.143 |
| 244 | -0.095 |
| 245 | 0.091 |
| 246 | 0.100 |

| | |
|---|---|
| 247 | 0.062 |
| 248 | 0.057 |
| 249 | 0.038 |
| 250 | -0.113 |
| 251 | -0.129 |
| 252 | -0.002 |
| 253 | 0.104 |
| 254 | 0.061 |
| 255 | -0.051 |
| 256 | 0.001 |
| 257 | 0.004 |
| 258 | -0.031 |
| 259 | -0.016 |
| 260 | -0.033 |
| 261 | -0.025 |
| 262 | 0.069 |
| 263 | -0.016 |
| 264 | 0.103 |
| 265 | 0.037 |
| 266 | 0.109 |
| 267 | -0.012 |
| 268 | 0.104 |
| 269 | -0.010 |
| 270 | -0.017 |
| 271 | -0.050 |
| 272 | -0.028 |
| 273 | -0.078 |
| 274 | 0.070 |
| 275 | 0.029 |
| 276 | -0.066 |
| 277 | -0.062 |
| 278 | -0.041 |
| 279 | -0.009 |
| 280 | 0.023 |
| 281 | -0.095 |
| 282 | 0.004 |
| 283 | -0.112 |
| 284 | 0.021 |
| 285 | 0.012 |
| 286 | 0.025 |
| 287 | -0.037 |
| 288 | -0.071 |
| 289 | 0.005 |
| 290 | -0.022 |
| 291 | -0.192 |
| 292 | 0.176 |
| 293 | -0.086 |
| 294 | -0.077 |
| 295 | 0.074 |
| 296 | -0.061 |

| | |
|---|---|
| 297 | 0.064 |
| 298 | 0.002 |
| 299 | -0.035 |
| 300 | 0.034 |
| 301 | -0.006 |
| 302 | -0.017 |
| 303 | -0.008 |
| 304 | 0.198 |
| 305 | 0.018 |
| 306 | -0.003 |
| 307 | -0.029 |
| 308 | -0.106 |
| 309 | 0.057 |
| 310 | 0.035 |
| 311 | -0.081 |
| 312 | -0.037 |
| 313 | -0.121 |
| 314 | 0.071 |
| 315 | -0.035 |
| 316 | -0.107 |
| 317 | -0.016 |
| 318 | 0.018 |
| 319 | 0.097 |
| 320 | 0.054 |
| 321 | -0.063 |
| 322 | -0.006 |
| 323 | 0.004 |
| 324 | 0.138 |
| 325 | 0.060 |
| 326 | -0.059 |
| 327 | 0.000 |
| 328 | -0.050 |
| 329 | -0.046 |
| 330 | 0.002 |
| 331 | 0.074 |
| 332 | 0.089 |
| 333 | 0.032 |
| 334 | -0.006 |
| 335 | -0.061 |
| 336 | -0.028 |
| 337 | 0.026 |
| 338 | -0.037 |
| 339 | 0.053 |
| 340 | -0.069 |
| 341 | 0.115 |
| 342 | -0.078 |
| 343 | 0.043 |
| 344 | 0.110 |
| 345 | -0.034 |
| 346 | -0.118 |

| | |
|---|---|
| 347 | -0.076 |
| 348 | -0.007 |
| 349 | -0.054 |
| 350 | 0.020 |
| 351 | -0.142 |
| 352 | -0.054 |
| 353 | 0.103 |
| 354 | -0.037 |
| 355 | -0.025 |
| 356 | -0.027 |
| 357 | 0.039 |
| 358 | 0.010 |
| 359 | 0.051 |
| 360 | -0.018 |
| 361 | -0.034 |
| 362 | 0.058 |
| 363 | 0.062 |
| 364 | 0.121 |
| 365 | 0.065 |
| 366 | -0.016 |
| 367 | 0.000 |
| 368 | -0.074 |
| 369 | -0.017 |
| 370 | -0.086 |
| 371 | -0.096 |
| 372 | 0.117 |
| 373 | -0.089 |
| 374 | 0.059 |
| 375 | 0.041 |
| 376 | 0.102 |
| 377 | 0.120 |
| 378 | 0.081 |
| 379 | 0.023 |
| 380 | -0.043 |
| 381 | 0.162 |
| 382 | 0.003 |
| 383 | 0.026 |
| 384 | -0.032 |
| 385 | 0.003 |
| 386 | -0.062 |
| 387 | 0.020 |
| 388 | 0.049 |
| 389 | -0.067 |
| 390 | 0.025 |
| 391 | 0.005 |
| 392 | 0.007 |
| 393 | -0.068 |
| 394 | 0.077 |
| 395 | 0.038 |
| 396 | 0.000 |

| | |
|---|---|
| 397 | 0.045 |
| 398 | 0.054 |
| 399 | -0.083 |
| 400 | 0.048 |
| 401 | 0.170 |
| 402 | 0.123 |
| 403 | -0.004 |
| 404 | 0.066 |
| 405 | 0.030 |
| 406 | 0.041 |
| 407 | 0.031 |
| 408 | -0.058 |
| 409 | 0.064 |
| 410 | 0.027 |
| 411 | 0.060 |
| 412 | -0.012 |
| 413 | 0.043 |
| 414 | -0.017 |
| 415 | 0.069 |
| 416 | -0.048 |
| 417 | 0.069 |
| 418 | -0.008 |
| 419 | 0.104 |
| 420 | 0.053 |
| 421 | -0.060 |
| 422 | -0.017 |
| 423 | -0.014 |
| 424 | -0.116 |
| 425 | -0.098 |
| 426 | -0.005 |
| 427 | 0.063 |
| 428 | 0.061 |
| 429 | 0.070 |
| 430 | -0.024 |
| 431 | 0.074 |
| 432 | -0.030 |
| 433 | -0.040 |
| 434 | -0.029 |
| 435 | 0.072 |
| 436 | -0.112 |
| 437 | -0.055 |
| 438 | -0.075 |
| 439 | -0.054 |
| 440 | -0.015 |
| 441 | -0.040 |
| 442 | -0.080 |
| 443 | 0.057 |
| 444 | -0.049 |
| 445 | -0.142 |
| 446 | -0.002 |

| | |
|---|---|
| 447 | 0.077 |
| 448 | -0.027 |
| 449 | -0.113 |
| 450 | -0.127 |
| 451 | -0.045 |
| 452 | 0.021 |
| 453 | 0.005 |
| 454 | -0.051 |
| 455 | -0.059 |
| 456 | -0.120 |
| 457 | 0.103 |
| 458 | -0.060 |
| 459 | 0.068 |
| 460 | 0.076 |
| 461 | -0.152 |
| 462 | 0.064 |
| 463 | -0.030 |
| 464 | -0.055 |
| 465 | -0.028 |
| 466 | -0.024 |
| 467 | -0.052 |
| 468 | 0.102 |
| 469 | -0.024 |
| 470 | -0.043 |
| 471 | 0.036 |
| 472 | 0.139 |
| 473 | -0.091 |
| 474 | -0.031 |
| 475 | 0.007 |
| 476 | -0.036 |
| 477 | 0.180 |
| 478 | -0.088 |
| 479 | 0.118 |
| 480 | 0.033 |
| 481 | 0.116 |
| 482 | 0.002 |
| 483 | 0.048 |
| 484 | 0.017 |
| 485 | -0.040 |
| 486 | 0.036 |
| 487 | 0.128 |
| 488 | -0.084 |
| 489 | 0.004 |
| 490 | -0.002 |
| 491 | 0.050 |
| 492 | -0.066 |
| 493 | -0.081 |
| 494 | -0.051 |
| 495 | -0.072 |
| 496 | -0.112 |

| | |
|---|---|
| 497 | 0.049 |
| 498 | -0.101 |
| 499 | -0.118 |
| 500 | -0.038 |
| 501 | 0.041 |
| 502 | -0.015 |
| 503 | 0.054 |
| 504 | 0.027 |
| 505 | -0.038 |
| 506 | -0.091 |
| 507 | -0.049 |
| 508 | -0.007 |
| 509 | -0.055 |
| 510 | -0.131 |
| 511 | 0.090 |
| 512 | 0.054 |
| 513 | -0.154 |
| 514 | -0.081 |
| 515 | 0.013 |
| 516 | -0.044 |
| 517 | 0.156 |
| 518 | 0.093 |
| 519 | -0.217 |
| 520 | 0.054 |
| 521 | -0.171 |
| 522 | 0.006 |
| 523 | 0.009 |
| 524 | -0.036 |
| 525 | -0.055 |
| 526 | -0.033 |
| 527 | 0.030 |
| 528 | -0.087 |
| 529 | 0.035 |
| 530 | -0.031 |
| 531 | -0.035 |
| 532 | 0.160 |
| 533 | 0.019 |
| 534 | 0.046 |
| 535 | 0.021 |
| 536 | -0.173 |
| 537 | -0.097 |
| 538 | -0.001 |
| 539 | -0.106 |
| 540 | 0.024 |
| 541 | 0.177 |
| 542 | 0.058 |
| 543 | -0.012 |
| 544 | -0.008 |
| 545 | -0.005 |
| 546 | 0.071 |

| | |
|---|---|
| 547 | 0.003 |
| 548 | -0.031 |
| 549 | 0.018 |
| 550 | -0.044 |
| 551 | 0.020 |
| 552 | 0.153 |
| 553 | -0.114 |
| 554 | 0.027 |
| 555 | 0.088 |
| 556 | 0.107 |
| 557 | -0.063 |
| 558 | -0.047 |
| 559 | -0.079 |
| 560 | 0.003 |
| 561 | 0.135 |
| 562 | 0.117 |
| 563 | 0.057 |
| 564 | 0.003 |
| 565 | -0.044 |
| 566 | 0.048 |
| 567 | 0.036 |
| 568 | -0.187 |
| 569 | 0.072 |
| 570 | -0.012 |
| 571 | -0.076 |
| 572 | 0.008 |
| 573 | -0.167 |
| 574 | -0.068 |
| 575 | -0.033 |
| 576 | 0.009 |
| 577 | 0.049 |
| 578 | 0.017 |
| 579 | 0.036 |
| 580 | 0.020 |
| 581 | -0.029 |
| 582 | -0.037 |
| 583 | 0.045 |
| 584 | -0.037 |
| 585 | 0.040 |
| 586 | 0.022 |
| 587 | 0.079 |
| 588 | -0.186 |
| 589 | -0.100 |
| 590 | 0.066 |
| 591 | 0.078 |
| 592 | 0.022 |
| 593 | 0.037 |
| 594 | 0.054 |
| 595 | -0.049 |
| 596 | -0.042 |

| | |
|---|---|
| 597 | 0.027 |
| 598 | -0.065 |
| 599 | 0.015 |
| 600 | -0.119 |
| 601 | -0.015 |
| 602 | -0.184 |
| 603 | -0.080 |
| 604 | -0.015 |
| 605 | -0.070 |
| 606 | 0.028 |
| 607 | -0.046 |
| 608 | 0.091 |
| 609 | 0.042 |
| 610 | -0.021 |
| 611 | 0.124 |
| 612 | 0.069 |
| 613 | 0.006 |
| 614 | -0.081 |
| 615 | 0.049 |
| 616 | 0.043 |
| 617 | -0.030 |
| 618 | 0.021 |
| 619 | -0.061 |
| 620 | -0.055 |
| 621 | -0.057 |
| 622 | -0.038 |
| 623 | 0.008 |
| 624 | -0.113 |
| 625 | -0.079 |
| 626 | -0.060 |
| 627 | 0.041 |
| 628 | 0.029 |
| 629 | -0.011 |
| 630 | -0.005 |
| 631 | 0.071 |
| 632 | 0.040 |
| 633 | 0.012 |
| 634 | 0.125 |
| 635 | -0.027 |
| 636 | -0.036 |
| 637 | -0.015 |
| 638 | -0.030 |
| 639 | 0.077 |
| 640 | -0.042 |
| 641 | -0.051 |
| 642 | -0.162 |
| 643 | 0.074 |
| 644 | -0.083 |
| 645 | 0.073 |
| 646 | 0.052 |

| | |
|---|---|
| 647 | -0.012 |
| 648 | 0.003 |
| 649 | -0.113 |
| 650 | 0.086 |
| 651 | -0.027 |
| 652 | 0.005 |
| 653 | -0.114 |
| 654 | -0.002 |
| 655 | -0.107 |
| 656 | 0.054 |
| 657 | 0.058 |
| 658 | 0.030 |
| 659 | -0.043 |
| 660 | 0.170 |
| 661 | 0.001 |
| 662 | 0.116 |
| 663 | -0.101 |
| 664 | 0.112 |
| 665 | 0.078 |
| 666 | -0.021 |
| 667 | 0.031 |
| 668 | -0.039 |
| 669 | -0.025 |
| 670 | -0.082 |
| 671 | -0.005 |
| 672 | 0.000 |
| 673 | 0.006 |
| 674 | 0.003 |
| 675 | -0.052 |
| 676 | 0.034 |
| 677 | -0.069 |
| 678 | 0.035 |
| 679 | -0.082 |
| 680 | 0.024 |
| 681 | 0.044 |
| 682 | -0.098 |
| 683 | 0.016 |
| 684 | -0.038 |
| 685 | 0.135 |
| 686 | -0.021 |
| 687 | -0.105 |
| 688 | -0.027 |
| 689 | 0.166 |
| 690 | 0.001 |
| 691 | -0.074 |
| 692 | -0.021 |
| 693 | 0.098 |
| 694 | -0.015 |
| 695 | 0.065 |
| 696 | 0.090 |

| | |
|---|---|
| 697 | 0.261 |
| 698 | -0.003 |
| 699 | 0.008 |
| 700 | -0.034 |
| 701 | -0.022 |
| 702 | -0.058 |
| 703 | 0.031 |
| 704 | 0.012 |
| 705 | -0.019 |
| 706 | -0.047 |
| 707 | -0.050 |
| 708 | -0.023 |
| 709 | 0.085 |
| 710 | -0.023 |
| 711 | 0.101 |
| 712 | -0.073 |
| 713 | -0.024 |
| 714 | 0.023 |
| 715 | 0.165 |
| 716 | 0.001 |
| 717 | -0.030 |
| 718 | 0.017 |
| 719 | -0.058 |
| 720 | 0.003 |
| 721 | 0.011 |
| 722 | -0.024 |
| 723 | 0.033 |
| 724 | -0.098 |
| 725 | -0.020 |
| 726 | 0.051 |
| 727 | 0.108 |
| 728 | 0.157 |
| 729 | 0.006 |
| 730 | -0.004 |
| 731 | 0.078 |
| 732 | -0.073 |
| 733 | 0.040 |
| 734 | -0.148 |
| 735 | 0.029 |
| 736 | 0.052 |
| 737 | 0.006 |
| 738 | 0.088 |
| 739 | -0.034 |
| 740 | -0.050 |
| 741 | 0.070 |
| 742 | 0.034 |
| 743 | 0.100 |
| 744 | 0.101 |
| 745 | -0.106 |
| 746 | 0.000 |

| | |
|---|---|
| 747 | 0.032 |
| 748 | 0.055 |
| 749 | -0.034 |
| 750 | 0.085 |
| 751 | -0.043 |
| 752 | -0.013 |
| 753 | 0.039 |
| 754 | 0.063 |
| 755 | 0.060 |
| 756 | 0.043 |
| 757 | 0.006 |
| 758 | 0.043 |
| 759 | 0.004 |
| 760 | 0.039 |
| 761 | 0.008 |
| 762 | 0.037 |
| 763 | 0.002 |
| 764 | 0.046 |
| 765 | -0.040 |
| 766 | -0.024 |
| 767 | -0.030 |
| 768 | -0.026 |
| 769 | 0.060 |
| 770 | -0.037 |
| 771 | 0.068 |
| 772 | 0.085 |
| 773 | 0.004 |
| 774 | 0.000 |
| 775 | 0.021 |
| 776 | -0.085 |
| 777 | -0.164 |
| 778 | -0.015 |
| 779 | -0.053 |
| 780 | 0.068 |
| 781 | 0.001 |
| 782 | 0.226 |
| 783 | -0.036 |
| 784 | 0.152 |
| 785 | -0.020 |
| 786 | -0.060 |
| 787 | 0.050 |
| 788 | -0.023 |
| 789 | -0.004 |
| 790 | 0.000 |
| 791 | 0.019 |
| 792 | 0.070 |
| 793 | 0.028 |
| 794 | 0.022 |
| 795 | -0.042 |
| 796 | -0.020 |

| | |
|---|---|
| 797 | 0.109 |
| 798 | 0.071 |
| 799 | 0.088 |
| 800 | 0.112 |
| 801 | -0.085 |
| 802 | -0.091 |
| 803 | 0.050 |
| 804 | -0.149 |
| 805 | 0.008 |
| 806 | 0.098 |
| 807 | 0.052 |
| 808 | -0.006 |
| 809 | -0.069 |
| 810 | 0.070 |
| 811 | -0.009 |
| 812 | 0.080 |
| 813 | 0.082 |
| 814 | -0.041 |
| 815 | -0.009 |
| 816 | 0.043 |
| 817 | 0.022 |
| 818 | -0.042 |
| 819 | 0.017 |
| 820 | -0.012 |
| 821 | -0.088 |
| 822 | 0.014 |
| 823 | -0.002 |
| 824 | 0.040 |
| 825 | 0.030 |
| 826 | -0.019 |
| 827 | -0.012 |
| 828 | 0.076 |
| 829 | 0.113 |
| 830 | -0.109 |
| 831 | 0.042 |
| 832 | -0.040 |
| 833 | 0.058 |
| 834 | 0.069 |
| 835 | -0.009 |
| 836 | -0.198 |
| 837 | -0.021 |
| 838 | -0.087 |
| 839 | 0.036 |
| 840 | 0.040 |
| 841 | -0.123 |
| 842 | 0.006 |
| 843 | -0.050 |
| 844 | 0.004 |
| 845 | 0.013 |
| 846 | 0.128 |

| | |
|---|---|
| 847 | -0.054 |
| 848 | 0.101 |
| 849 | -0.014 |
| 850 | -0.035 |
| 851 | -0.081 |
| 852 | 0.098 |
| 853 | 0.079 |
| 854 | -0.042 |
| 855 | 0.003 |
| 856 | -0.112 |
| 857 | -0.048 |
| 858 | -0.112 |
| 859 | 0.007 |
| 860 | 0.098 |
| 861 | -0.057 |
| 862 | -0.045 |
| 863 | 0.048 |
| 864 | 0.063 |
| 865 | 0.019 |
| 866 | -0.064 |
| 867 | -0.016 |
| 868 | 0.118 |
| 869 | -0.046 |
| 870 | -0.072 |
| 871 | -0.041 |
| 872 | -0.041 |
| 873 | -0.089 |
| 874 | 0.074 |
| 875 | 0.032 |
| 876 | 0.044 |
| 877 | -0.115 |
| 878 | 0.001 |
| 879 | -0.061 |
| 880 | -0.118 |
| 881 | 0.081 |
| 882 | 0.179 |
| 883 | 0.097 |
| 884 | -0.083 |
| 885 | -0.011 |
| 886 | 0.068 |
| 887 | 0.052 |
| 888 | -0.097 |
| 889 | -0.044 |
| 890 | -0.013 |
| 891 | -0.104 |
| 892 | 0.042 |
| 893 | -0.003 |
| 894 | 0.108 |
| 895 | -0.096 |
| 896 | 0.010 |

| | |
|---|---|
| 897 | -0.067 |
| 898 | 0.061 |
| 899 | 0.066 |
| 900 | 0.050 |
| 901 | 0.064 |
| 902 | 0.017 |
| 903 | 0.039 |
| 904 | 0.014 |
| 905 | -0.042 |
| 906 | 0.068 |
| 907 | -0.008 |
| 908 | 0.007 |
| 909 | -0.032 |
| 910 | 0.021 |
| 911 | -0.064 |
| 912 | 0.031 |
| 913 | 0.111 |
| 914 | 0.046 |
| 915 | -0.001 |
| 916 | 0.063 |
| 917 | -0.039 |
| 918 | 0.115 |
| 919 | -0.120 |
| 920 | 0.052 |
| 921 | 0.057 |
| 922 | 0.056 |
| 923 | -0.035 |
| 924 | 0.105 |
| 925 | 0.112 |
| 926 | -0.075 |
| 927 | 0.070 |
| 928 | -0.022 |
| 929 | -0.007 |
| 930 | -0.082 |
| 931 | -0.051 |
| 932 | -0.060 |
| 933 | -0.048 |
| 934 | -0.134 |
| 935 | 0.003 |
| 936 | -0.072 |
| 937 | -0.031 |
| 938 | 0.071 |
| 939 | -0.015 |
| 940 | 0.086 |
| 941 | -0.066 |
| 942 | 0.035 |
| 943 | 0.034 |
| 944 | 0.038 |
| 945 | -0.111 |
| 946 | 0.080 |

| | |
|---|---|
| 947 | 0.005 |
| 948 | -0.098 |
| 949 | 0.011 |
| 950 | 0.160 |
| 951 | 0.023 |
| 952 | -0.034 |
| 953 | -0.095 |
| 954 | -0.062 |
| 955 | 0.088 |
| 956 | 0.119 |
| 957 | 0.022 |
| 958 | -0.059 |
| 959 | -0.035 |
| 960 | -0.058 |
| 961 | -0.074 |
| 962 | -0.004 |
| 963 | 0.092 |
| 964 | -0.026 |
| 965 | 0.068 |
| 966 | -0.011 |
| 967 | 0.062 |
| 968 | 0.093 |
| 969 | 0.103 |
| 970 | 0.015 |
| 971 | 0.152 |
| 972 | 0.029 |
| 973 | -0.027 |
| 974 | -0.024 |
| 975 | 0.035 |
| 976 | 0.069 |
| 977 | -0.030 |
| 978 | -0.124 |
| 979 | 0.022 |
| 980 | -0.014 |
| 981 | 0.096 |
| 982 | 0.021 |
| 983 | 0.015 |
| 984 | 0.029 |
| 985 | -0.122 |
| 986 | 0.068 |
| 987 | -0.020 |
| 988 | -0.054 |
| 989 | 0.051 |
| 990 | -0.055 |
| 991 | 0.012 |
| 992 | 0.030 |
| 993 | -0.061 |
| 994 | -0.124 |
| 995 | -0.088 |
| 996 | 0.058 |

| | |
|---|---|
| 997 | -0.019 |
| 998 | 0.070 |
| 999 | 0.107 |
| 1000 | -0.017 |
| 1001 | 0.005 |
| 1002 | 0.015 |
| 1003 | -0.021 |
| 1004 | -0.005 |
| 1005 | -0.070 |
| 1006 | -0.039 |
| 1007 | 0.056 |
| 1008 | 0.003 |
| 1009 | -0.052 |
| 1010 | 0.023 |
| 1011 | 0.007 |
| 1012 | 0.122 |
| 1013 | -0.036 |
| 1014 | -0.008 |
| 1015 | 0.152 |
| 1016 | -0.097 |
| 1017 | 0.131 |
| 1018 | 0.156 |
| 1019 | -0.003 |
| 1020 | -0.021 |
| 1021 | 0.009 |
| 1022 | -0.119 |
| 1023 | 0.077 |
| 1024 | 0.061 |
| 1025 | 0.058 |
| 1026 | -0.045 |
| 1027 | 0.055 |
| 1028 | 0.000 |
| 1029 | 0.100 |
| 1030 | -0.047 |
| 1031 | -0.042 |
| 1032 | -0.006 |
| 1033 | 0.082 |
| 1034 | 0.064 |
| 1035 | -0.012 |
| 1036 | 0.023 |
| 1037 | -0.088 |
| 1038 | 0.002 |
| 1039 | -0.052 |
| 1040 | -0.051 |
| 1041 | -0.122 |
| 1042 | -0.133 |
| 1043 | 0.107 |
| 1044 | 0.097 |
| 1045 | -0.117 |
| 1046 | 0.123 |

| | |
|---|---|
| 1047 | -0.031 |
| 1048 | -0.009 |
| 1049 | 0.059 |
| 1050 | -0.046 |
| 1051 | 0.067 |
| 1052 | -0.066 |
| 1053 | -0.071 |
| 1054 | 0.045 |
| 1055 | -0.104 |
| 1056 | -0.022 |
| 1057 | 0.034 |
| 1058 | 0.029 |
| 1059 | -0.031 |
| 1060 | 0.007 |
| 1061 | 0.008 |
| 1062 | 0.085 |
| 1063 | -0.048 |
| 1064 | 0.018 |
| 1065 | 0.004 |
| 1066 | -0.009 |
| 1067 | -0.018 |
| 1068 | -0.192 |
| 1069 | 0.052 |
| 1070 | -0.019 |
| 1071 | -0.107 |
| 1072 | 0.065 |
| 1073 | -0.122 |
| 1074 | 0.023 |
| 1075 | 0.063 |
| 1076 | -0.036 |
| 1077 | 0.215 |
| 1078 | 0.006 |
| 1079 | -0.004 |
| 1080 | 0.073 |
| 1081 | -0.073 |
| 1082 | -0.019 |
| 1083 | -0.006 |
| 1084 | -0.050 |
| 1085 | 0.017 |
| 1086 | 0.005 |
| 1087 | 0.021 |
| 1088 | 0.067 |
| 1089 | 0.055 |
| 1090 | -0.009 |
| 1091 | 0.059 |
| 1092 | -0.020 |
| 1093 | -0.033 |
| 1094 | 0.113 |
| 1095 | 0.078 |
| 1096 | -0.028 |

| | |
|---|---|
| 1097 | -0.020 |
| 1098 | -0.008 |
| 1099 | -0.009 |
| 1100 | -0.054 |
| 1101 | -0.110 |
| 1102 | -0.023 |
| 1103 | 0.210 |
| 1104 | -0.007 |
| 1105 | 0.130 |
| 1106 | -0.122 |
| 1107 | 0.018 |
| 1108 | -0.172 |
| 1109 | -0.013 |
| 1110 | 0.041 |
| 1111 | 0.026 |
| 1112 | 0.043 |
| 1113 | 0.028 |
| 1114 | 0.054 |
| 1115 | 0.112 |
| 1116 | 0.021 |
| 1117 | -0.023 |
| 1118 | -0.234 |
| 1119 | -0.010 |
| 1120 | 0.052 |
| 1121 | -0.129 |
| 1122 | -0.104 |
| 1123 | 0.024 |
| 1124 | -0.113 |
| 1125 | 0.039 |
| 1126 | -0.059 |
| 1127 | 0.089 |
| 1128 | 0.106 |
| 1129 | 0.019 |
| 1130 | -0.050 |
| 1131 | 0.150 |
| 1132 | 0.035 |
| 1133 | 0.129 |
| 1134 | 0.150 |
| 1135 | -0.053 |
| 1136 | 0.032 |
| 1137 | 0.043 |
| 1138 | 0.136 |
| 1139 | 0.108 |
| 1140 | 0.000 |
| 1141 | -0.011 |
| 1142 | -0.006 |
| 1143 | 0.053 |
| 1144 | 0.054 |
| 1145 | -0.111 |
| 1146 | 0.060 |

| | |
|---|---|
| 1147 | -0.049 |
| 1148 | 0.089 |
| 1149 | 0.043 |
| 1150 | -0.036 |
| 1151 | 0.056 |
| 1152 | -0.013 |
| 1153 | 0.100 |
| 1154 | -0.004 |
| 1155 | -0.015 |
| 1156 | -0.118 |
| 1157 | 0.076 |
| 1158 | -0.021 |
| 1159 | -0.030 |
| 1160 | 0.038 |
| 1161 | -0.055 |
| 1162 | -0.040 |
| 1163 | 0.130 |
| 1164 | -0.055 |
| 1165 | -0.001 |
| 1166 | -0.191 |
| 1167 | 0.035 |
| 1168 | -0.106 |
| 1169 | -0.076 |
| 1170 | 0.120 |
| 1171 | 0.061 |
| 1172 | 0.049 |
| 1173 | 0.019 |
| 1174 | -0.051 |
| 1175 | -0.004 |
| 1176 | -0.051 |
| 1177 | -0.006 |
| 1178 | -0.035 |
| 1179 | -0.083 |
| 1180 | 0.061 |
| 1181 | -0.089 |
| 1182 | 0.005 |
| 1183 | -0.086 |
| 1184 | -0.067 |
| 1185 | 0.027 |
| 1186 | 0.019 |
| 1187 | 0.026 |
| 1188 | -0.062 |
| 1189 | -0.165 |
| 1190 | 0.078 |
| 1191 | 0.068 |
| 1192 | 0.072 |
| 1193 | -0.044 |
| 1194 | -0.025 |
| 1195 | 0.077 |
| 1196 | 0.050 |

| | |
|---|---|
| 1197 | -0.005 |
| 1198 | 0.149 |
| 1199 | -0.107 |
| 1200 | -0.009 |
| 1201 | 0.078 |
| 1202 | 0.081 |
| 1203 | -0.005 |
| 1204 | 0.080 |
| 1205 | -0.106 |
| 1206 | 0.025 |
| 1207 | 0.047 |
| 1208 | -0.062 |
| 1209 | 0.013 |
| 1210 | 0.050 |
| 1211 | -0.075 |
| 1212 | 0.011 |
| 1213 | 0.069 |
| 1214 | 0.082 |
| 1215 | 0.020 |
| 1216 | 0.020 |
| 1217 | 0.071 |
| 1218 | 0.000 |
| 1219 | -0.082 |
| 1220 | -0.015 |
| 1221 | 0.090 |
| 1222 | -0.045 |
| 1223 | -0.011 |
| 1224 | -0.048 |
| 1225 | 0.046 |
| 1226 | 0.034 |
| 1227 | 0.047 |
| 1228 | 0.040 |
| 1229 | -0.032 |
| 1230 | -0.020 |
| 1231 | -0.093 |
| 1232 | 0.084 |
| 1233 | -0.055 |
| 1234 | -0.001 |
| 1235 | 0.117 |
| 1236 | -0.008 |
| 1237 | -0.013 |
| 1238 | -0.025 |
| 1239 | -0.076 |
| 1240 | 0.008 |
| 1241 | 0.096 |
| 1242 | -0.044 |
| 1243 | 0.076 |
| 1244 | 0.028 |
| 1245 | -0.026 |
| 1246 | -0.001 |

| | |
|---|---|
| 1247 | 0.112 |
| 1248 | -0.043 |
| 1249 | 0.052 |
| 1250 | 0.112 |
| 1251 | -0.054 |
| 1252 | 0.113 |
| 1253 | -0.072 |
| 1254 | -0.103 |
| 1255 | -0.040 |
| 1256 | -0.094 |
| 1257 | 0.071 |
| 1258 | -0.028 |
| 1259 | 0.041 |
| 1260 | -0.073 |
| 1261 | -0.010 |
| 1262 | -0.055 |
| 1263 | -0.120 |
| 1264 | 0.029 |
| 1265 | 0.073 |
| 1266 | 0.081 |
| 1267 | 0.048 |
| 1268 | -0.079 |
| 1269 | 0.037 |
| 1270 | 0.111 |
| 1271 | -0.039 |
| 1272 | -0.035 |
| 1273 | -0.028 |
| 1274 | -0.073 |
| 1275 | -0.006 |
| 1276 | -0.123 |
| 1277 | 0.110 |
| 1278 | 0.051 |
| 1279 | 0.048 |
| 1280 | -0.018 |
| 1281 | -0.011 |
| 1282 | 0.041 |
| 1283 | -0.005 |
| 1284 | 0.036 |
| 1285 | 0.035 |
| 1286 | 0.120 |
| 1287 | 0.135 |
| 1288 | 0.039 |
| 1289 | -0.016 |
| 1290 | -0.073 |
| 1291 | 0.084 |
| 1292 | -0.094 |
| 1293 | 0.016 |
| 1294 | -0.100 |
| 1295 | 0.079 |
| 1296 | 0.000 |

| | |
|---|---|
| 1297 | -0.036 |
| 1298 | 0.069 |
| 1299 | -0.027 |
| 1300 | -0.067 |
| 1301 | 0.074 |
| 1302 | 0.099 |
| 1303 | 0.029 |
| 1304 | 0.084 |
| 1305 | 0.022 |
| 1306 | -0.050 |
| 1307 | 0.034 |
| 1308 | -0.097 |
| 1309 | 0.113 |
| 1310 | -0.098 |
| 1311 | 0.028 |
| 1312 | -0.045 |
| 1313 | -0.067 |
| 1314 | 0.147 |
| 1315 | 0.003 |
| 1316 | -0.164 |
| 1317 | -0.064 |
| 1318 | 0.080 |
| 1319 | -0.065 |
| 1320 | -0.046 |
| 1321 | 0.094 |
| 1322 | 0.002 |
| 1323 | -0.066 |
| 1324 | 0.041 |
| 1325 | 0.107 |
| 1326 | 0.050 |
| 1327 | -0.103 |
| 1328 | 0.015 |
| 1329 | 0.064 |
| 1330 | -0.087 |
| 1331 | 0.016 |
| 1332 | 0.022 |
| 1333 | -0.030 |
| 1334 | -0.051 |
| 1335 | -0.004 |
| 1336 | 0.116 |
| 1337 | 0.032 |
| 1338 | 0.056 |
| 1339 | -0.022 |
| 1340 | 0.110 |
| 1341 | 0.154 |
| 1342 | 0.052 |
| 1343 | 0.019 |
| 1344 | 0.059 |
| 1345 | 0.068 |
| 1346 | 0.081 |

| | |
|---|---|
| 1347 | 0.038 |
| 1348 | 0.005 |
| 1349 | -0.221 |
| 1350 | -0.004 |
| 1351 | -0.110 |
| 1352 | -0.045 |
| 1353 | -0.013 |
| 1354 | -0.033 |
| 1355 | 0.007 |
| 1356 | 0.033 |
| 1357 | 0.029 |
| 1358 | 0.044 |
| 1359 | -0.012 |
| 1360 | 0.024 |
| 1361 | -0.004 |
| 1362 | 0.104 |
| 1363 | 0.016 |
| 1364 | 0.026 |
| 1365 | -0.123 |
| 1366 | -0.057 |
| 1367 | 0.088 |
| 1368 | -0.025 |
| 1369 | 0.093 |
| 1370 | 0.050 |
| 1371 | -0.004 |
| 1372 | 0.071 |
| 1373 | 0.003 |
| 1374 | 0.123 |
| 1375 | -0.061 |
| 1376 | -0.003 |
| 1377 | -0.015 |
| 1378 | 0.051 |
| 1379 | -0.075 |
| 1380 | 0.030 |
| 1381 | -0.044 |
| 1382 | -0.124 |
| 1383 | 0.063 |
| 1384 | -0.043 |
| 1385 | -0.063 |
| 1386 | -0.005 |
| 1387 | -0.060 |
| 1388 | 0.109 |
| 1389 | 0.039 |
| 1390 | 0.043 |
| 1391 | -0.179 |
| 1392 | 0.013 |
| 1393 | -0.032 |
| 1394 | 0.099 |
| 1395 | -0.066 |
| 1396 | -0.096 |

| | |
|---|---|
| 1397 | -0.026 |
| 1398 | 0.002 |
| 1399 | 0.021 |
| 1400 | -0.034 |
| 1401 | -0.073 |
| 1402 | 0.070 |
| 1403 | -0.055 |
| 1404 | 0.062 |
| 1405 | 0.058 |
| 1406 | -0.016 |
| 1407 | 0.069 |
| 1408 | 0.212 |
| 1409 | -0.046 |
| 1410 | 0.152 |
| 1411 | -0.009 |
| 1412 | -0.057 |
| 1413 | 0.015 |
| 1414 | -0.039 |
| 1415 | 0.037 |
| 1416 | -0.069 |
| 1417 | 0.001 |
| 1418 | -0.092 |
| 1419 | -0.056 |
| 1420 | 0.038 |
| 1421 | -0.018 |
| 1422 | 0.002 |
| 1423 | -0.004 |
| 1424 | 0.058 |
| 1425 | -0.032 |
| 1426 | -0.110 |
| 1427 | 0.182 |
| 1428 | -0.004 |
| 1429 | -0.063 |
| 1430 | 0.063 |
| 1431 | 0.064 |
| 1432 | -0.166 |
| 1433 | 0.044 |
| 1434 | 0.128 |
| 1435 | 0.002 |
| 1436 | -0.086 |
| 1437 | -0.033 |
| 1438 | 0.041 |
| 1439 | -0.052 |
| 1440 | 0.009 |
| 1441 | 0.007 |
| 1442 | -0.027 |
| 1443 | -0.002 |
| 1444 | -0.018 |
| 1445 | 0.060 |
| 1446 | -0.025 |

| | |
|---|---|
| 1447 | 0.023 |
| 1448 | -0.039 |
| 1449 | 0.070 |
| 1450 | -0.034 |
| 1451 | 0.043 |
| 1452 | 0.140 |
| 1453 | 0.030 |
| 1454 | -0.084 |
| 1455 | 0.018 |
| 1456 | 0.021 |
| 1457 | -0.005 |
| 1458 | -0.035 |
| 1459 | -0.003 |
| 1460 | 0.025 |
| 1461 | 0.133 |
| 1462 | -0.052 |
| 1463 | 0.095 |
| 1464 | -0.023 |
| 1465 | 0.045 |
| 1466 | -0.016 |
| 1467 | -0.105 |
| 1468 | 0.007 |
| 1469 | -0.035 |
| 1470 | -0.015 |
| 1471 | 0.060 |
| 1472 | 0.048 |
| 1473 | 0.063 |
| 1474 | 0.088 |
| 1475 | 0.048 |
| 1476 | 0.078 |
| 1477 | -0.083 |
| 1478 | -0.167 |
| 1479 | -0.004 |
| 1480 | 0.039 |
| 1481 | 0.051 |
| 1482 | -0.019 |
| 1483 | 0.074 |
| 1484 | -0.060 |
| 1485 | 0.060 |
| 1486 | 0.023 |
| 1487 | 0.013 |
| 1488 | 0.112 |
| 1489 | -0.026 |
| 1490 | -0.081 |
| 1491 | -0.066 |
| 1492 | 0.021 |
| 1493 | 0.043 |
| 1494 | 0.014 |
| 1495 | -0.015 |
| 1496 | -0.069 |

| | |
|---|---|
| 1497 | 0.057 |
| 1498 | -0.035 |
| 1499 | -0.089 |
| 1500 | 0.078 |
| 1501 | 0.045 |
| 1502 | 0.074 |
| 1503 | 0.036 |
| 1504 | 0.082 |
| 1505 | -0.050 |
| 1506 | -0.059 |
| 1507 | -0.040 |
| 1508 | 0.026 |
| 1509 | -0.042 |
| 1510 | -0.016 |
| 1511 | 0.100 |
| 1512 | 0.025 |
| 1513 | 0.062 |
| 1514 | 0.090 |
| 1515 | -0.145 |
| 1516 | 0.061 |
| 1517 | -0.175 |
| 1518 | 0.034 |
| 1519 | 0.049 |
| 1520 | 0.049 |
| 1521 | -0.021 |
| 1522 | 0.077 |
| 1523 | 0.045 |
| 1524 | -0.009 |
| 1525 | 0.072 |
| 1526 | 0.070 |
| 1527 | -0.054 |
| 1528 | -0.227 |
| 1529 | 0.067 |
| 1530 | 0.001 |
| 1531 | -0.033 |
| 1532 | -0.054 |
| 1533 | -0.022 |
| 1534 | 0.112 |
| 1535 | 0.035 |
| 1536 | -0.051 |
| 1537 | -0.068 |
| 1538 | -0.067 |
| 1539 | 0.122 |
| 1540 | -0.064 |
| 1541 | 0.041 |
| 1542 | -0.029 |
| 1543 | 0.047 |
| 1544 | 0.109 |
| 1545 | 0.142 |
| 1546 | -0.044 |

| | |
|---|---|
| 1547 | 0.078 |
| 1548 | 0.013 |
| 1549 | 0.117 |
| 1550 | -0.011 |
| 1551 | -0.064 |
| 1552 | -0.016 |
| 1553 | -0.086 |
| 1554 | -0.013 |
| 1555 | 0.088 |
| 1556 | 0.137 |
| 1557 | 0.027 |
| 1558 | -0.067 |
| 1559 | -0.054 |
| 1560 | -0.022 |
| 1561 | 0.048 |
| 1562 | -0.011 |
| 1563 | 0.073 |
| 1564 | -0.012 |
| 1565 | 0.192 |
| 1566 | -0.044 |
| 1567 | 0.050 |
| 1568 | -0.055 |
| 1569 | 0.095 |
| 1570 | -0.022 |
| 1571 | -0.091 |
| 1572 | -0.061 |
| 1573 | -0.033 |
| 1574 | 0.027 |
| 1575 | 0.211 |
| 1576 | -0.087 |
| 1577 | 0.058 |
| 1578 | 0.028 |
| 1579 | 0.067 |
| 1580 | -0.024 |
| 1581 | 0.007 |
| 1582 | 0.039 |
| 1583 | -0.121 |
| 1584 | 0.117 |
| 1585 | 0.068 |
| 1586 | 0.064 |
| 1587 | 0.055 |
| 1588 | -0.088 |
| 1589 | -0.035 |
| 1590 | 0.111 |
| 1591 | -0.070 |
| 1592 | -0.081 |
| 1593 | 0.095 |
| 1594 | -0.118 |
| 1595 | 0.028 |
| 1596 | 0.015 |

| | |
|---|---|
| 1597 | 0.042 |
| 1598 | 0.022 |
| 1599 | 0.029 |
| 1600 | 0.124 |
| 1601 | 0.092 |
| 1602 | -0.053 |
| 1603 | 0.011 |
| 1604 | -0.078 |
| 1605 | 0.065 |
| 1606 | 0.021 |
| 1607 | 0.007 |
| 1608 | 0.013 |
| 1609 | -0.010 |
| 1610 | 0.008 |
| 1611 | 0.014 |
| 1612 | -0.192 |
| 1613 | -0.032 |
| 1614 | -0.031 |
| 1615 | 0.032 |
| 1616 | 0.050 |
| 1617 | -0.016 |
| 1618 | -0.117 |
| 1619 | 0.125 |
| 1620 | -0.050 |
| 1621 | -0.068 |
| 1622 | -0.061 |
| 1623 | -0.002 |
| 1624 | -0.013 |
| 1625 | 0.070 |
| 1626 | 0.016 |
| 1627 | -0.077 |
| 1628 | -0.038 |
| 1629 | -0.037 |
| 1630 | -0.040 |
| 1631 | 0.047 |
| 1632 | 0.028 |
| 1633 | 0.073 |
| 1634 | -0.091 |
| 1635 | -0.007 |
| 1636 | -0.011 |
| 1637 | -0.024 |
| 1638 | -0.014 |
| 1639 | 0.084 |
| 1640 | -0.006 |
| 1641 | -0.100 |
| 1642 | -0.068 |
| 1643 | -0.037 |
| 1644 | 0.040 |
| 1645 | -0.006 |
| 1646 | 0.114 |

| | |
|---|---|
| 1647 | 0.074 |
| 1648 | -0.047 |
| 1649 | -0.004 |
| 1650 | 0.112 |
| 1651 | -0.069 |
| 1652 | -0.054 |
| 1653 | 0.049 |
| 1654 | 0.081 |
| 1655 | -0.023 |
| 1656 | -0.120 |
| 1657 | 0.109 |
| 1658 | -0.139 |
| 1659 | 0.131 |
| 1660 | 0.120 |
| 1661 | -0.121 |
| 1662 | -0.072 |
| 1663 | 0.073 |
| 1664 | -0.053 |
| 1665 | 0.064 |
| 1666 | -0.028 |
| 1667 | -0.054 |
| 1668 | 0.096 |
| 1669 | -0.020 |
| 1670 | 0.007 |
| 1671 | 0.012 |
| 1672 | -0.016 |
| 1673 | 0.037 |
| 1674 | 0.103 |
| 1675 | -0.039 |
| 1676 | 0.019 |
| 1677 | 0.006 |
| 1678 | 0.052 |
| 1679 | 0.007 |
| 1680 | 0.007 |
| 1681 | 0.029 |
| 1682 | -0.045 |
| 1683 | 0.044 |
| 1684 | 0.023 |
| 1685 | 0.007 |
| 1686 | -0.111 |
| 1687 | 0.041 |
| 1688 | -0.032 |
| 1689 | -0.023 |
| 1690 | 0.028 |
| 1691 | -0.075 |
| 1692 | -0.066 |
| 1693 | -0.079 |
| 1694 | -0.101 |
| 1695 | -0.095 |
| 1696 | 0.055 |

| | |
|---|---|
| 1697 | -0.079 |
| 1698 | -0.050 |
| 1699 | 0.112 |
| 1700 | 0.041 |
| 1701 | -0.007 |
| 1702 | -0.107 |
| 1703 | 0.119 |
| 1704 | -0.054 |
| 1705 | 0.067 |
| 1706 | -0.030 |
| 1707 | 0.003 |
| 1708 | -0.086 |
| 1709 | -0.041 |
| 1710 | -0.038 |
| 1711 | -0.011 |
| 1712 | -0.060 |
| 1713 | 0.116 |
| 1714 | 0.046 |
| 1715 | 0.018 |
| 1716 | 0.017 |
| 1717 | 0.016 |
| 1718 | -0.072 |
| 1719 | 0.040 |
| 1720 | 0.050 |
| 1721 | -0.077 |
| 1722 | 0.054 |
| 1723 | -0.060 |
| 1724 | -0.025 |
| 1725 | -0.022 |
| 1726 | 0.081 |
| 1727 | 0.013 |
| 1728 | 0.072 |
| 1729 | -0.061 |
| 1730 | -0.016 |
| 1731 | 0.087 |
| 1732 | -0.050 |
| 1733 | 0.022 |
| 1734 | 0.005 |
| 1735 | 0.088 |
| 1736 | -0.013 |
| 1737 | -0.009 |
| 1738 | 0.013 |
| 1739 | -0.180 |
| 1740 | 0.028 |
| 1741 | 0.066 |
| 1742 | -0.044 |
| 1743 | 0.034 |
| 1744 | 0.016 |
| 1745 | 0.093 |
| 1746 | -0.060 |

| | |
|---|---|
| 1747 | -0.037 |
| 1748 | 0.003 |
| 1749 | 0.089 |
| 1750 | 0.076 |
| 1751 | -0.072 |
| 1752 | 0.038 |
| 1753 | -0.066 |
| 1754 | 0.062 |
| 1755 | -0.007 |
| 1756 | -0.022 |
| 1757 | 0.053 |
| 1758 | -0.044 |
| 1759 | -0.039 |
| 1760 | 0.001 |
| 1761 | 0.000 |
| 1762 | 0.015 |
| 1763 | -0.036 |
| 1764 | 0.043 |
| 1765 | -0.029 |
| 1766 | -0.075 |
| 1767 | 0.009 |
| 1768 | 0.140 |
| 1769 | -0.105 |
| 1770 | 0.059 |
| 1771 | -0.112 |
| 1772 | -0.096 |
| 1773 | 0.072 |
| 1774 | 0.022 |
| 1775 | -0.008 |
| 1776 | 0.047 |
| 1777 | -0.037 |
| 1778 | 0.091 |
| 1779 | -0.119 |
| 1780 | 0.075 |
| 1781 | 0.147 |
| 1782 | 0.027 |
| 1783 | -0.022 |
| 1784 | -0.023 |
| 1785 | 0.039 |
| 1786 | 0.038 |
| 1787 | 0.018 |
| 1788 | 0.024 |
| 1789 | 0.000 |
| 1790 | 0.025 |
| 1791 | 0.064 |
| 1792 | -0.119 |
| 1793 | -0.071 |
| 1794 | -0.032 |
| 1795 | 0.035 |
| 1796 | -0.087 |

| | |
|---|---|
| 1797 | -0.044 |
| 1798 | -0.028 |
| 1799 | -0.022 |
| 1800 | 0.030 |
| 1801 | 0.176 |
| 1802 | -0.002 |
| 1803 | 0.051 |
| 1804 | -0.031 |
| 1805 | 0.040 |
| 1806 | -0.089 |
| 1807 | 0.028 |
| 1808 | -0.043 |
| 1809 | 0.083 |
| 1810 | 0.056 |
| 1811 | 0.075 |
| 1812 | -0.012 |
| 1813 | 0.045 |
| 1814 | 0.029 |
| 1815 | 0.046 |
| 1816 | -0.011 |
| 1817 | -0.078 |
| 1818 | -0.026 |
| 1819 | 0.020 |
| 1820 | -0.079 |
| 1821 | -0.021 |
| 1822 | -0.082 |
| 1823 | -0.101 |
| 1824 | 0.101 |
| 1825 | 0.103 |
| 1826 | 0.040 |
| 1827 | -0.101 |
| 1828 | 0.004 |
| 1829 | 0.047 |
| 1830 | -0.035 |
| 1831 | 0.030 |
| 1832 | -0.044 |
| 1833 | -0.053 |
| 1834 | 0.081 |
| 1835 | 0.045 |
| 1836 | -0.101 |
| 1837 | -0.002 |
| 1838 | -0.043 |
| 1839 | 0.072 |
| 1840 | 0.153 |
| 1841 | -0.023 |
| 1842 | -0.063 |
| 1843 | 0.009 |
| 1844 | 0.040 |
| 1845 | 0.015 |
| 1846 | 0.109 |

| | |
|---|---|
| 1847 | -0.067 |
| 1848 | -0.164 |
| 1849 | -0.090 |
| 1850 | -0.089 |
| 1851 | 0.032 |
| 1852 | -0.005 |
| 1853 | -0.127 |
| 1854 | -0.104 |
| 1855 | 0.037 |
| 1856 | -0.025 |
| 1857 | 0.035 |
| 1858 | -0.128 |
| 1859 | -0.096 |
| 1860 | 0.089 |
| 1861 | 0.102 |
| 1862 | -0.047 |
| 1863 | 0.003 |
| 1864 | 0.081 |
| 1865 | 0.046 |
| 1866 | -0.044 |
| 1867 | 0.031 |
| 1868 | 0.019 |
| 1869 | 0.023 |
| 1870 | -0.027 |
| 1871 | 0.183 |
| 1872 | -0.031 |
| 1873 | 0.007 |
| 1874 | 0.140 |
| 1875 | 0.104 |
| 1876 | -0.029 |
| 1877 | 0.009 |
| 1878 | -0.118 |
| 1879 | 0.051 |
| 1880 | -0.023 |
| 1881 | 0.073 |
| 1882 | -0.050 |
| 1883 | -0.059 |
| 1884 | 0.027 |
| 1885 | -0.219 |
| 1886 | -0.106 |
| 1887 | 0.053 |
| 1888 | 0.006 |
| 1889 | 0.026 |
| 1890 | 0.017 |
| 1891 | 0.005 |
| 1892 | 0.145 |
| 1893 | -0.056 |
| 1894 | 0.018 |
| 1895 | -0.026 |
| 1896 | 0.051 |

| | |
|---|---|
| 1897 | 0.007 |
| 1898 | -0.004 |
| 1899 | -0.045 |
| 1900 | 0.048 |
| 1901 | -0.091 |
| 1902 | -0.108 |
| 1903 | 0.008 |
| 1904 | -0.073 |
| 1905 | -0.085 |
| 1906 | 0.014 |
| 1907 | -0.061 |
| 1908 | 0.131 |
| 1909 | 0.022 |
| 1910 | -0.123 |
| 1911 | 0.077 |
| 1912 | -0.072 |
| 1913 | 0.006 |
| 1914 | 0.060 |
| 1915 | -0.007 |
| 1916 | -0.022 |
| 1917 | -0.027 |
| 1918 | 0.050 |
| 1919 | -0.076 |
| 1920 | -0.069 |
| 1921 | -0.033 |
| 1922 | -0.131 |
| 1923 | -0.089 |
| 1924 | 0.043 |
| 1925 | 0.021 |
| 1926 | 0.099 |
| 1927 | 0.065 |
| 1928 | -0.021 |
| 1929 | -0.023 |
| 1930 | -0.105 |
| 1931 | -0.093 |
| 1932 | 0.083 |
| 1933 | 0.102 |
| 1934 | 0.012 |
| 1935 | 0.043 |
| 1936 | 0.021 |
| 1937 | -0.044 |
| 1938 | -0.107 |
| 1939 | -0.099 |
| 1940 | -0.133 |
| 1941 | -0.064 |
| 1942 | 0.166 |
| 1943 | 0.000 |
| 1944 | 0.018 |
| 1945 | 0.037 |
| 1946 | 0.003 |

| | |
|---|---|
| 1947 | -0.049 |
| 1948 | -0.012 |
| 1949 | -0.036 |
| 1950 | -0.134 |
| 1951 | -0.030 |
| 1952 | 0.108 |
| 1953 | -0.062 |
| 1954 | -0.092 |
| 1955 | 0.009 |
| 1956 | -0.040 |
| 1957 | 0.019 |
| 1958 | 0.052 |
| 1959 | -0.004 |
| 1960 | 0.152 |
| 1961 | 0.062 |
| 1962 | -0.155 |
| 1963 | -0.006 |
| 1964 | -0.015 |
| 1965 | 0.064 |
| 1966 | 0.098 |
| 1967 | 0.101 |
| 1968 | -0.059 |
| 1969 | -0.005 |
| 1970 | -0.071 |
| 1971 | 0.017 |
| 1972 | -0.034 |
| 1973 | 0.005 |
| 1974 | -0.094 |
| 1975 | 0.012 |
| 1976 | -0.034 |
| 1977 | -0.029 |
| 1978 | 0.046 |
| 1979 | 0.040 |
| 1980 | -0.099 |
| 1981 | 0.051 |
| 1982 | 0.015 |
| 1983 | -0.041 |
| 1984 | -0.108 |
| 1985 | 0.055 |
| 1986 | -0.030 |
| 1987 | -0.084 |
| 1988 | -0.051 |
| 1989 | 0.001 |
| 1990 | 0.035 |
| 1991 | 0.095 |
| 1992 | 0.034 |
| 1993 | -0.115 |
| 1994 | 0.078 |
| 1995 | -0.125 |
| 1996 | 0.015 |

| | |
|---|---|
| 1997 | -0.028 |
| 1998 | -0.012 |
| 1999 | 0.057 |
| 2000 | -0.048 |
| 2001 | -0.071 |
| 2002 | -0.004 |
| 2003 | 0.111 |
| 2004 | 0.072 |
| 2005 | 0.005 |
| 2006 | 0.018 |
| 2007 | -0.067 |
| 2008 | 0.055 |
| 2009 | -0.019 |
| 2010 | -0.112 |
| 2011 | 0.098 |
| 2012 | 0.128 |
| 2013 | 0.049 |
| 2014 | -0.121 |
| 2015 | 0.020 |
| 2016 | 0.067 |
| 2017 | 0.113 |
| 2018 | 0.013 |
| 2019 | -0.067 |
| 2020 | -0.005 |
| 2021 | -0.055 |
| 2022 | -0.007 |
| 2023 | 0.114 |
| 2024 | -0.053 |
| 2025 | 0.032 |
| 2026 | -0.130 |
| 2027 | 0.006 |
| 2028 | 0.076 |
| 2029 | 0.014 |
| 2030 | 0.021 |
| 2031 | 0.149 |
| 2032 | 0.046 |
| 2033 | 0.002 |
| 2034 | 0.044 |
| 2035 | -0.060 |
| 2036 | 0.121 |
| 2037 | 0.062 |
| 2038 | 0.023 |
| 2039 | 0.084 |
| 2040 | -0.107 |
| 2041 | -0.083 |
| 2042 | -0.042 |
| 2043 | -0.041 |
| 2044 | 0.004 |
| 2045 | 0.028 |
| 2046 | 0.003 |

| | |
|---|---|
| 2047 | 0.063 |
| 2048 | 0.130 |
| 2049 | -0.011 |
| 2050 | -0.007 |
| 2051 | 0.016 |
| 2052 | 0.014 |
| 2053 | 0.064 |
| 2054 | 0.012 |
| 2055 | -0.013 |
| 2056 | 0.007 |
| 2057 | 0.101 |
| 2058 | -0.016 |
| 2059 | 0.010 |
| 2060 | 0.151 |
| 2061 | -0.095 |
| 2062 | 0.092 |
| 2063 | -0.017 |
| 2064 | -0.016 |
| 2065 | 0.037 |
| 2066 | -0.029 |
| 2067 | 0.005 |
| 2068 | -0.018 |
| 2069 | -0.052 |
| 2070 | -0.062 |
| 2071 | 0.037 |
| 2072 | 0.012 |
| 2073 | -0.078 |
| 2074 | 0.212 |
| 2075 | 0.012 |
| 2076 | 0.083 |
| 2077 | 0.056 |
| 2078 | 0.121 |
| 2079 | -0.098 |
| 2080 | -0.054 |
| 2081 | -0.096 |
| 2082 | -0.053 |
| 2083 | 0.046 |
| 2084 | 0.055 |
| 2085 | -0.069 |
| 2086 | 0.058 |
| 2087 | -0.003 |
| 2088 | 0.082 |
| 2089 | 0.031 |
| 2090 | -0.021 |
| 2091 | 0.008 |
| 2092 | 0.033 |
| 2093 | 0.083 |
| 2094 | 0.134 |
| 2095 | -0.009 |
| 2096 | -0.074 |

| | |
|---|---|
| 2097 | -0.044 |
| 2098 | -0.003 |
| 2099 | -0.014 |
| 2100 | 0.000 |
| 2101 | 0.040 |
| 2102 | -0.055 |
| 2103 | 0.106 |
| 2104 | 0.051 |
| 2105 | 0.048 |
| 2106 | 0.079 |
| 2107 | 0.001 |
| 2108 | -0.008 |
| 2109 | -0.076 |
| 2110 | -0.077 |
| 2111 | 0.017 |
| 2112 | 0.044 |
| 2113 | 0.051 |
| 2114 | -0.092 |
| 2115 | -0.097 |
| 2116 | -0.142 |
| 2117 | -0.002 |
| 2118 | 0.049 |
| 2119 | 0.038 |
| 2120 | -0.014 |
| 2121 | -0.052 |
| 2122 | 0.042 |
| 2123 | 0.042 |
| 2124 | -0.075 |
| 2125 | -0.049 |
| 2126 | -0.069 |
| 2127 | 0.032 |
| 2128 | -0.140 |
| 2129 | 0.079 |
| 2130 | 0.042 |
| 2131 | 0.002 |
| 2132 | 0.110 |
| 2133 | -0.032 |
| 2134 | -0.112 |
| 2135 | 0.140 |
| 2136 | 0.072 |
| 2137 | -0.095 |
| 2138 | 0.017 |
| 2139 | 0.011 |
| 2140 | -0.019 |
| 2141 | -0.057 |
| 2142 | -0.128 |
| 2143 | -0.017 |
| 2144 | 0.020 |
| 2145 | -0.045 |
| 2146 | -0.013 |

| | |
|---|---|
| 2147 | -0.083 |
| 2148 | 0.116 |
| 2149 | 0.065 |
| 2150 | 0.034 |
| 2151 | 0.089 |
| 2152 | 0.021 |
| 2153 | 0.102 |
| 2154 | 0.007 |
| 2155 | 0.039 |
| 2156 | -0.103 |
| 2157 | 0.039 |
| 2158 | 0.015 |
| 2159 | 0.080 |
| 2160 | 0.060 |
| 2161 | 0.073 |
| 2162 | -0.021 |
| 2163 | 0.024 |
| 2164 | -0.038 |
| 2165 | -0.016 |
| 2166 | -0.122 |
| 2167 | 0.041 |
| 2168 | -0.048 |
| 2169 | -0.003 |
| 2170 | -0.114 |
| 2171 | 0.062 |
| 2172 | -0.026 |
| 2173 | -0.086 |
| 2174 | 0.026 |
| 2175 | 0.070 |
| 2176 | 0.063 |
| 2177 | -0.053 |
| 2178 | 0.012 |
| 2179 | -0.103 |
| 2180 | 0.000 |
| 2181 | -0.063 |
| 2182 | -0.001 |
| 2183 | 0.020 |
| 2184 | 0.025 |
| 2185 | 0.045 |
| 2186 | 0.079 |
| 2187 | 0.042 |
| 2188 | 0.072 |
| 2189 | 0.011 |
| 2190 | -0.085 |
| 2191 | 0.060 |
| 2192 | -0.069 |
| 2193 | -0.033 |
| 2194 | -0.102 |
| 2195 | -0.075 |
| 2196 | -0.093 |

| | |
|---|---|
| 2197 | 0.034 |
| 2198 | 0.069 |
| 2199 | -0.077 |
| 2200 | 0.075 |
| 2201 | 0.003 |
| 2202 | -0.063 |
| 2203 | 0.060 |
| 2204 | -0.024 |
| 2205 | -0.028 |
| 2206 | 0.019 |
| 2207 | -0.113 |
| 2208 | -0.071 |
| 2209 | -0.031 |
| 2210 | -0.127 |
| 2211 | -0.044 |
| 2212 | 0.061 |
| 2213 | -0.084 |
| 2214 | 0.063 |
| 2215 | -0.098 |
| 2216 | -0.025 |
| 2217 | -0.096 |
| 2218 | 0.012 |
| 2219 | -0.141 |
| 2220 | -0.053 |
| 2221 | 0.017 |
| 2222 | -0.040 |
| 2223 | 0.065 |
| 2224 | 0.020 |
| 2225 | 0.063 |
| 2226 | 0.052 |
| 2227 | 0.048 |
| 2228 | -0.199 |
| 2229 | -0.023 |
| 2230 | -0.089 |
| 2231 | 0.084 |
| 2232 | 0.034 |
| 2233 | -0.047 |
| 2234 | 0.037 |
| 2235 | 0.151 |
| 2236 | 0.068 |
| 2237 | -0.075 |
| 2238 | -0.007 |
| 2239 | -0.009 |
| 2240 | 0.062 |
| 2241 | -0.010 |
| 2242 | -0.013 |
| 2243 | -0.077 |
| 2244 | -0.060 |
| 2245 | -0.093 |
| 2246 | -0.119 |

| | |
|---|---|
| 2247 | 0.024 |
| 2248 | 0.086 |
| 2249 | 0.042 |
| 2250 | -0.013 |
| 2251 | 0.017 |
| 2252 | 0.082 |
| 2253 | 0.041 |
| 2254 | 0.045 |
| 2255 | -0.022 |
| 2256 | 0.005 |
| 2257 | 0.002 |
| 2258 | 0.029 |
| 2259 | 0.001 |
| 2260 | 0.074 |
| 2261 | 0.132 |
| 2262 | -0.044 |
| 2263 | -0.056 |
| 2264 | -0.063 |
| 2265 | 0.021 |
| 2266 | -0.030 |
| 2267 | -0.028 |
| 2268 | -0.033 |
| 2269 | -0.039 |
| 2270 | -0.043 |
| 2271 | 0.024 |
| 2272 | 0.055 |
| 2273 | -0.051 |
| 2274 | -0.037 |
| 2275 | -0.102 |
| 2276 | 0.071 |
| 2277 | 0.030 |
| 2278 | 0.028 |
| 2279 | -0.073 |
| 2280 | -0.103 |
| 2281 | -0.043 |
| 2282 | -0.019 |
| 2283 | -0.087 |
| 2284 | -0.054 |
| 2285 | 0.074 |
| 2286 | -0.072 |
| 2287 | -0.116 |
| 2288 | 0.035 |
| 2289 | -0.022 |
| 2290 | -0.001 |
| 2291 | -0.043 |
| 2292 | -0.049 |
| 2293 | 0.092 |
| 2294 | 0.002 |
| 2295 | -0.069 |
| 2296 | -0.007 |

| | |
|---|---|
| 2297 | -0.099 |
| 2298 | -0.116 |
| 2299 | -0.055 |
| 2300 | 0.063 |
| 2301 | 0.071 |
| 2302 | 0.062 |
| 2303 | -0.010 |
| 2304 | -0.035 |
| 2305 | 0.073 |
| 2306 | 0.065 |
| 2307 | -0.133 |
| 2308 | 0.021 |
| 2309 | 0.007 |
| 2310 | -0.074 |
| 2311 | 0.016 |
| 2312 | -0.022 |
| 2313 | 0.009 |
| 2314 | -0.003 |
| 2315 | 0.034 |
| 2316 | 0.028 |
| 2317 | -0.064 |
| 2318 | 0.106 |
| 2319 | -0.042 |
| 2320 | -0.007 |
| 2321 | 0.090 |
| 2322 | 0.047 |
| 2323 | -0.101 |
| 2324 | -0.014 |
| 2325 | -0.050 |
| 2326 | 0.075 |
| 2327 | -0.012 |
| 2328 | 0.028 |
| 2329 | 0.034 |
| 2330 | -0.150 |
| 2331 | 0.027 |
| 2332 | -0.053 |
| 2333 | -0.032 |
| 2334 | -0.151 |
| 2335 | 0.013 |
| 2336 | 0.169 |
| 2337 | 0.078 |
| 2338 | -0.199 |
| 2339 | -0.001 |
| 2340 | 0.049 |
| 2341 | -0.141 |
| 2342 | -0.062 |
| 2343 | 0.031 |
| 2344 | 0.025 |
| 2345 | 0.091 |
| 2346 | -0.039 |

| | |
|---|---|
| 2347 | -0.083 |
| 2348 | -0.099 |
| 2349 | 0.093 |
| 2350 | 0.047 |
| 2351 | 0.018 |
| 2352 | 0.140 |
| 2353 | -0.089 |
| 2354 | 0.088 |
| 2355 | -0.075 |
| 2356 | -0.005 |
| 2357 | 0.050 |
| 2358 | -0.066 |
| 2359 | 0.105 |
| 2360 | -0.004 |
| 2361 | 0.074 |
| 2362 | 0.072 |
| 2363 | -0.076 |
| 2364 | -0.168 |
| 2365 | -0.025 |
| 2366 | 0.140 |
| 2367 | 0.071 |
| 2368 | -0.016 |
| 2369 | -0.108 |
| 2370 | 0.068 |
| 2371 | -0.013 |
| 2372 | 0.039 |
| 2373 | 0.100 |
| 2374 | 0.048 |
| 2375 | 0.060 |
| 2376 | -0.018 |
| 2377 | 0.036 |
| 2378 | 0.072 |
| 2379 | -0.047 |
| 2380 | -0.056 |
| 2381 | -0.089 |
| 2382 | 0.007 |
| 2383 | -0.015 |
| 2384 | 0.027 |
| 2385 | 0.084 |
| 2386 | -0.136 |
| 2387 | 0.017 |
| 2388 | -0.171 |
| 2389 | 0.039 |
| 2390 | -0.011 |
| 2391 | 0.099 |
| 2392 | 0.040 |
| 2393 | -0.043 |
| 2394 | 0.048 |
| 2395 | 0.022 |
| 2396 | -0.023 |

| | |
|---|---|
| 2397 | 0.046 |
| 2398 | -0.007 |
| 2399 | -0.038 |
| 2400 | -0.029 |
| 2401 | 0.125 |
| 2402 | -0.009 |
| 2403 | -0.144 |
| 2404 | 0.130 |
| 2405 | -0.059 |
| 2406 | -0.104 |
| 2407 | -0.049 |
| 2408 | 0.075 |
| 2409 | -0.070 |
| 2410 | 0.064 |
| 2411 | 0.060 |
| 2412 | -0.041 |
| 2413 | 0.043 |
| 2414 | -0.004 |
| 2415 | -0.063 |
| 2416 | -0.037 |
| 2417 | -0.148 |
| 2418 | 0.048 |
| 2419 | -0.041 |
| 2420 | -0.027 |
| 2421 | -0.021 |
| 2422 | -0.009 |
| 2423 | -0.051 |
| 2424 | -0.174 |
| 2425 | -0.093 |
| 2426 | 0.088 |
| 2427 | 0.072 |
| 2428 | 0.003 |
| 2429 | 0.117 |
| 2430 | -0.072 |
| 2431 | -0.023 |
| 2432 | 0.117 |
| 2433 | 0.039 |
| 2434 | -0.052 |
| 2435 | 0.023 |
| 2436 | -0.010 |
| 2437 | 0.053 |
| 2438 | -0.113 |
| 2439 | 0.033 |
| 2440 | 0.040 |
| 2441 | 0.086 |
| 2442 | -0.051 |
| 2443 | -0.037 |
| 2444 | 0.023 |
| 2445 | -0.025 |
| 2446 | -0.116 |

| | |
|---|---|
| 2447 | 0.007 |
| 2448 | 0.106 |
| 2449 | -0.147 |
| 2450 | 0.089 |
| 2451 | 0.070 |
| 2452 | 0.011 |
| 2453 | 0.016 |
| 2454 | -0.099 |
| 2455 | 0.057 |
| 2456 | 0.120 |
| 2457 | -0.030 |
| 2458 | -0.103 |
| 2459 | -0.083 |
| 2460 | -0.116 |
| 2461 | -0.073 |
| 2462 | -0.001 |
| 2463 | -0.077 |
| 2464 | -0.007 |
| 2465 | 0.122 |
| 2466 | -0.052 |
| 2467 | -0.110 |
| 2468 | -0.008 |
| 2469 | -0.006 |
| 2470 | -0.057 |
| 2471 | 0.051 |
| 2472 | -0.072 |
| 2473 | 0.006 |
| 2474 | 0.016 |
| 2475 | -0.005 |
| 2476 | 0.020 |
| 2477 | -0.018 |
| 2478 | 0.063 |
| 2479 | 0.039 |
| 2480 | 0.015 |
| 2481 | -0.057 |
| 2482 | -0.061 |
| 2483 | -0.016 |
| 2484 | 0.088 |
| 2485 | 0.000 |
| 2486 | 0.025 |
| 2487 | 0.037 |
| 2488 | -0.081 |
| 2489 | -0.087 |
| 2490 | 0.010 |
| 2491 | 0.020 |
| 2492 | -0.061 |
| 2493 | 0.046 |
| 2494 | 0.023 |
| 2495 | 0.056 |
| 2496 | -0.002 |

| | |
|---|---|
| 2497 | 0.057 |
| 2498 | 0.039 |
| 2499 | -0.030 |
| 2500 | -0.004 |
| 2501 | -0.023 |
| 2502 | -0.006 |
| 2503 | 0.030 |
| 2504 | 0.095 |
| 2505 | 0.081 |
| 2506 | 0.005 |
| 2507 | 0.011 |
| 2508 | -0.029 |
| 2509 | 0.027 |
| 2510 | 0.055 |
| 2511 | -0.029 |
| 2512 | 0.099 |
| 2513 | -0.073 |
| 2514 | 0.008 |
| 2515 | 0.052 |
| 2516 | -0.060 |
| 2517 | 0.026 |
| 2518 | -0.088 |
| 2519 | 0.072 |
| 2520 | 0.112 |
| 2521 | -0.130 |
| 2522 | -0.128 |
| 2523 | -0.056 |
| 2524 | -0.116 |
| 2525 | 0.145 |
| 2526 | 0.031 |
| 2527 | 0.063 |
| 2528 | -0.056 |
| 2529 | 0.037 |
| 2530 | -0.061 |
| 2531 | -0.142 |
| 2532 | 0.058 |
| 2533 | 0.029 |
| 2534 | 0.066 |
| 2535 | 0.010 |
| 2536 | 0.048 |
| 2537 | -0.002 |
| 2538 | 0.104 |
| 2539 | -0.022 |
| 2540 | 0.083 |
| 2541 | 0.046 |
| 2542 | -0.013 |
| 2543 | 0.087 |
| 2544 | -0.059 |
| 2545 | 0.095 |
| 2546 | 0.040 |

| | |
|---|---|
| 2547 | 0.037 |
| 2548 | -0.037 |
| 2549 | -0.083 |
| 2550 | 0.032 |
| 2551 | -0.119 |
| 2552 | 0.046 |
| 2553 | 0.048 |
| 2554 | 0.017 |
| 2555 | -0.112 |
| 2556 | 0.017 |
| 2557 | -0.143 |
| 2558 | 0.039 |
| 2559 | -0.040 |
| 2560 | -0.057 |
| 2561 | -0.027 |
| 2562 | 0.085 |
| 2563 | 0.038 |
| 2564 | 0.022 |
| 2565 | 0.004 |
| 2566 | -0.068 |
| 2567 | 0.019 |
| 2568 | -0.023 |
| 2569 | -0.073 |
| 2570 | -0.073 |
| 2571 | 0.052 |
| 2572 | -0.018 |
| 2573 | -0.068 |
| 2574 | -0.038 |
| 2575 | -0.037 |
| 2576 | 0.030 |
| 2577 | -0.056 |
| 2578 | -0.092 |
| 2579 | -0.072 |
| 2580 | 0.058 |
| 2581 | 0.143 |
| 2582 | -0.123 |
| 2583 | 0.102 |
| 2584 | 0.026 |
| 2585 | 0.069 |
| 2586 | -0.021 |
| 2587 | -0.033 |
| 2588 | 0.053 |
| 2589 | 0.060 |
| 2590 | -0.005 |
| 2591 | -0.005 |
| 2592 | -0.144 |
| 2593 | 0.009 |
| 2594 | -0.087 |
| 2595 | 0.054 |
| 2596 | 0.001 |

| | |
|---|---|
| 2597 | -0.090 |
| 2598 | 0.032 |
| 2599 | -0.042 |
| 2600 | -0.021 |
| 2601 | -0.123 |
| 2602 | 0.100 |
| 2603 | 0.107 |
| 2604 | -0.053 |
| 2605 | 0.000 |
| 2606 | 0.030 |
| 2607 | -0.099 |
| 2608 | -0.109 |
| 2609 | -0.069 |
| 2610 | -0.114 |
| 2611 | -0.072 |
| 2612 | 0.036 |
| 2613 | -0.040 |
| 2614 | 0.037 |
| 2615 | 0.039 |
| 2616 | 0.047 |
| 2617 | 0.004 |
| 2618 | 0.019 |
| 2619 | 0.052 |
| 2620 | 0.028 |
| 2621 | 0.005 |
| 2622 | 0.032 |
| 2623 | -0.009 |
| 2624 | 0.062 |
| 2625 | -0.015 |
| 2626 | 0.013 |
| 2627 | -0.119 |
| 2628 | 0.064 |
| 2629 | 0.020 |
| 2630 | 0.055 |
| 2631 | -0.154 |
| 2632 | -0.003 |
| 2633 | 0.005 |
| 2634 | -0.012 |
| 2635 | 0.007 |
| 2636 | 0.074 |
| 2637 | 0.046 |
| 2638 | 0.029 |
| 2639 | 0.042 |
| 2640 | 0.025 |
| 2641 | -0.165 |
| 2642 | -0.002 |
| 2643 | 0.001 |
| 2644 | -0.067 |
| 2645 | -0.058 |
| 2646 | 0.046 |

| | |
|---|---|
| 2647 | 0.119 |
| 2648 | 0.096 |
| 2649 | -0.003 |
| 2650 | -0.047 |
| 2651 | -0.037 |
| 2652 | 0.072 |
| 2653 | -0.164 |
| 2654 | 0.098 |
| 2655 | -0.033 |
| 2656 | 0.041 |
| 2657 | -0.041 |
| 2658 | -0.046 |
| 2659 | 0.031 |
| 2660 | 0.128 |
| 2661 | 0.031 |
| 2662 | 0.065 |
| 2663 | 0.102 |
| 2664 | 0.095 |
| 2665 | 0.068 |
| 2666 | 0.023 |
| 2667 | 0.006 |
| 2668 | -0.011 |
| 2669 | 0.065 |
| 2670 | 0.001 |
| 2671 | -0.081 |
| 2672 | 0.070 |
| 2673 | -0.089 |
| 2674 | -0.112 |
| 2675 | -0.006 |
| 2676 | 0.056 |
| 2677 | -0.013 |
| 2678 | 0.024 |
| 2679 | 0.045 |
| 2680 | 0.016 |
| 2681 | 0.098 |
| 2682 | 0.054 |
| 2683 | 0.102 |
| 2684 | -0.072 |
| 2685 | -0.009 |
| 2686 | 0.019 |
| 2687 | -0.039 |
| 2688 | 0.063 |
| 2689 | -0.056 |
| 2690 | 0.094 |
| 2691 | -0.056 |
| 2692 | 0.075 |
| 2693 | -0.122 |
| 2694 | 0.058 |
| 2695 | 0.020 |
| 2696 | 0.085 |

| | |
|---|---|
| 2697 | 0.044 |
| 2698 | -0.061 |
| 2699 | -0.118 |
| 2700 | -0.091 |
| 2701 | 0.130 |
| 2702 | -0.001 |
| 2703 | 0.013 |
| 2704 | 0.076 |
| 2705 | -0.065 |
| 2706 | 0.009 |
| 2707 | -0.111 |
| 2708 | -0.015 |
| 2709 | 0.059 |
| 2710 | 0.071 |
| 2711 | 0.020 |
| 2712 | 0.029 |
| 2713 | 0.080 |
| 2714 | -0.121 |
| 2715 | -0.015 |
| 2716 | 0.051 |
| 2717 | 0.101 |
| 2718 | 0.027 |
| 2719 | 0.008 |
| 2720 | 0.033 |
| 2721 | -0.006 |
| 2722 | 0.135 |
| 2723 | 0.041 |
| 2724 | -0.006 |
| 2725 | -0.104 |
| 2726 | 0.053 |
| 2727 | 0.022 |
| 2728 | 0.071 |
| 2729 | -0.005 |
| 2730 | 0.059 |
| 2731 | -0.059 |
| 2732 | 0.024 |
| 2733 | -0.034 |
| 2734 | -0.014 |
| 2735 | -0.060 |
| 2736 | -0.045 |
| 2737 | 0.109 |
| 2738 | 0.000 |
| 2739 | 0.086 |
| 2740 | -0.125 |
| 2741 | -0.003 |
| 2742 | -0.039 |
| 2743 | -0.080 |
| 2744 | -0.072 |
| 2745 | 0.064 |
| 2746 | -0.104 |

| | |
|---|---|
| 2747 | 0.051 |
| 2748 | 0.036 |
| 2749 | -0.007 |
| 2750 | 0.124 |
| 2751 | -0.077 |
| 2752 | 0.158 |
| 2753 | 0.067 |
| 2754 | -0.040 |
| 2755 | -0.054 |
| 2756 | 0.047 |
| 2757 | 0.029 |
| 2758 | -0.056 |
| 2759 | 0.116 |
| 2760 | -0.057 |
| 2761 | 0.064 |
| 2762 | -0.022 |
| 2763 | 0.004 |
| 2764 | -0.147 |
| 2765 | 0.076 |
| 2766 | -0.030 |
| 2767 | -0.022 |
| 2768 | -0.053 |
| 2769 | 0.091 |
| 2770 | -0.011 |
| 2771 | 0.022 |
| 2772 | -0.001 |
| 2773 | 0.005 |
| 2774 | -0.023 |
| 2775 | -0.089 |
| 2776 | 0.015 |
| 2777 | 0.003 |
| 2778 | -0.005 |
| 2779 | 0.109 |
| 2780 | -0.130 |
| 2781 | -0.029 |
| 2782 | -0.112 |
| 2783 | 0.096 |
| 2784 | 0.018 |
| 2785 | 0.091 |
| 2786 | 0.065 |
| 2787 | -0.054 |
| 2788 | -0.103 |
| 2789 | 0.063 |
| 2790 | -0.014 |
| 2791 | 0.073 |
| 2792 | -0.064 |
| 2793 | -0.101 |
| 2794 | -0.047 |
| 2795 | -0.011 |
| 2796 | 0.144 |

| | |
|---|---|
| 2797 | 0.089 |
| 2798 | 0.001 |
| 2799 | 0.092 |
| 2800 | 0.056 |
| 2801 | 0.006 |
| 2802 | 0.021 |
| 2803 | -0.026 |
| 2804 | -0.006 |
| 2805 | -0.094 |
| 2806 | 0.007 |
| 2807 | -0.003 |
| 2808 | -0.005 |
| 2809 | -0.008 |
| 2810 | 0.080 |
| 2811 | -0.122 |
| 2812 | -0.013 |
| 2813 | 0.085 |
| 2814 | -0.006 |
| 2815 | -0.085 |
| 2816 | 0.028 |
| 2817 | -0.042 |
| 2818 | 0.148 |
| 2819 | -0.042 |
| 2820 | 0.053 |
| 2821 | 0.067 |
| 2822 | 0.152 |
| 2823 | -0.010 |
| 2824 | 0.030 |
| 2825 | 0.052 |
| 2826 | -0.025 |
| 2827 | -0.026 |
| 2828 | 0.024 |
| 2829 | -0.151 |
| 2830 | -0.002 |
| 2831 | -0.019 |
| 2832 | 0.018 |
| 2833 | 0.163 |
| 2834 | 0.050 |
| 2835 | -0.062 |
| 2836 | -0.041 |
| 2837 | 0.060 |
| 2838 | 0.128 |
| 2839 | -0.055 |
| 2840 | 0.080 |
| 2841 | -0.029 |
| 2842 | 0.025 |
| 2843 | -0.003 |
| 2844 | 0.044 |
| 2845 | -0.016 |
| 2846 | 0.052 |

| | |
|---|---|
| 2847 | -0.011 |
| 2848 | -0.031 |
| 2849 | -0.001 |
| 2850 | -0.078 |
| 2851 | 0.062 |
| 2852 | -0.021 |
| 2853 | 0.001 |
| 2854 | 0.035 |
| 2855 | -0.116 |
| 2856 | -0.057 |
| 2857 | 0.047 |
| 2858 | -0.030 |
| 2859 | -0.010 |
| 2860 | 0.000 |
| 2861 | -0.077 |
| 2862 | -0.036 |
| 2863 | -0.038 |
| 2864 | -0.096 |
| 2865 | 0.069 |
| 2866 | 0.124 |
| 2867 | 0.035 |
| 2868 | -0.108 |
| 2869 | 0.016 |
| 2870 | 0.024 |
| 2871 | 0.068 |
| 2872 | -0.024 |
| 2873 | -0.023 |
| 2874 | -0.001 |
| 2875 | -0.039 |
| 2876 | 0.107 |
| 2877 | 0.090 |
| 2878 | -0.072 |
| 2879 | -0.133 |
| 2880 | -0.019 |
| 2881 | 0.024 |
| 2882 | 0.060 |
| 2883 | 0.001 |
| 2884 | 0.015 |
| 2885 | -0.089 |
| 2886 | 0.067 |
| 2887 | -0.160 |
| 2888 | -0.079 |
| 2889 | -0.002 |
| 2890 | 0.009 |
| 2891 | -0.032 |
| 2892 | -0.017 |
| 2893 | 0.098 |
| 2894 | -0.109 |
| 2895 | 0.001 |
| 2896 | -0.102 |

| | |
|---|---|
| 2897 | 0.031 |
| 2898 | 0.002 |
| 2899 | -0.134 |
| 2900 | -0.014 |
| 2901 | 0.074 |
| 2902 | 0.032 |
| 2903 | -0.027 |
| 2904 | 0.011 |
| 2905 | 0.037 |
| 2906 | 0.060 |
| 2907 | 0.110 |
| 2908 | 0.145 |
| 2909 | -0.088 |
| 2910 | 0.052 |
| 2911 | 0.031 |
| 2912 | 0.074 |
| 2913 | 0.042 |
| 2914 | 0.075 |
| 2915 | 0.028 |
| 2916 | -0.035 |
| 2917 | 0.054 |
| 2918 | 0.009 |
| 2919 | -0.047 |
| 2920 | 0.019 |
| 2921 | -0.113 |
| 2922 | 0.033 |
| 2923 | -0.051 |
| 2924 | -0.062 |
| 2925 | 0.038 |
| 2926 | 0.020 |
| 2927 | -0.015 |
| 2928 | -0.054 |
| 2929 | -0.182 |
| 2930 | -0.037 |
| 2931 | -0.005 |
| 2932 | 0.047 |
| 2933 | -0.117 |
| 2934 | 0.096 |
| 2935 | -0.022 |
| 2936 | 0.125 |
| 2937 | 0.103 |
| 2938 | 0.008 |
| 2939 | -0.032 |
| 2940 | -0.068 |
| 2941 | 0.035 |
| 2942 | 0.075 |
| 2943 | 0.000 |
| 2944 | 0.110 |
| 2945 | 0.035 |
| 2946 | 0.009 |

| | |
|---|---|
| 2947 | -0.060 |
| 2948 | -0.017 |
| 2949 | 0.058 |
| 2950 | 0.076 |
| 2951 | 0.049 |
| 2952 | 0.107 |
| 2953 | -0.195 |
| 2954 | -0.002 |
| 2955 | -0.095 |
| 2956 | -0.008 |
| 2957 | -0.005 |
| 2958 | -0.097 |
| 2959 | 0.262 |
| 2960 | 0.052 |
| 2961 | -0.093 |
| 2962 | 0.128 |
| 2963 | -0.011 |
| 2964 | 0.057 |
| 2965 | -0.050 |
| 2966 | 0.068 |
| 2967 | 0.025 |
| 2968 | -0.021 |
| 2969 | 0.030 |
| 2970 | 0.057 |
| 2971 | 0.029 |
| 2972 | 0.014 |
| 2973 | -0.045 |
| 2974 | 0.025 |
| 2975 | 0.044 |
| 2976 | -0.093 |
| 2977 | -0.016 |
| 2978 | 0.099 |
| 2979 | 0.010 |
| 2980 | 0.075 |
| 2981 | -0.003 |
| 2982 | -0.074 |
| 2983 | 0.000 |
| 2984 | 0.033 |
| 2985 | 0.015 |
| 2986 | 0.040 |
| 2987 | -0.052 |
| 2988 | -0.048 |
| 2989 | -0.071 |
| 2990 | 0.043 |
| 2991 | -0.002 |
| 2992 | 0.021 |
| 2993 | 0.032 |
| 2994 | 0.049 |
| 2995 | -0.033 |
| 2996 | 0.065 |

| | |
|---|---|
| 2997 | -0.022 |
| 2998 | -0.038 |
| 2999 | 0.060 |
| 3000 | -0.074 |
| 3001 | 0.014 |
| 3002 | 0.003 |
| 3003 | -0.038 |
| 3004 | -0.126 |
| 3005 | 0.098 |
| 3006 | -0.046 |
| 3007 | 0.005 |
| 3008 | -0.008 |
| 3009 | -0.066 |
| 3010 | 0.006 |
| 3011 | -0.001 |
| 3012 | 0.029 |
| 3013 | -0.112 |
| 3014 | 0.001 |
| 3015 | -0.036 |
| 3016 | -0.038 |
| 3017 | -0.051 |
| 3018 | -0.010 |
| 3019 | 0.069 |
| 3020 | 0.073 |
| 3021 | 0.133 |
| 3022 | 0.025 |
| 3023 | -0.004 |
| 3024 | -0.039 |
| 3025 | -0.025 |
| 3026 | -0.013 |
| 3027 | -0.035 |
| 3028 | -0.087 |
| 3029 | -0.047 |
| 3030 | 0.035 |
| 3031 | -0.006 |
| 3032 | -0.026 |
| 3033 | 0.048 |
| 3034 | -0.006 |
| 3035 | 0.040 |
| 3036 | -0.097 |
| 3037 | 0.041 |
| 3038 | 0.007 |
| 3039 | 0.088 |
| 3040 | -0.042 |
| 3041 | 0.012 |
| 3042 | -0.075 |
| 3043 | -0.066 |
| 3044 | -0.035 |
| 3045 | -0.019 |
| 3046 | -0.096 |

| | |
|---|---|
| 3047 | 0.102 |
| 3048 | -0.083 |
| 3049 | 0.022 |
| 3050 | 0.009 |
| 3051 | -0.009 |
| 3052 | -0.113 |
| 3053 | 0.110 |
| 3054 | -0.052 |
| 3055 | 0.170 |
| 3056 | -0.037 |
| 3057 | 0.050 |
| 3058 | 0.056 |
| 3059 | 0.077 |
| 3060 | -0.054 |
| 3061 | -0.041 |
| 3062 | -0.018 |
| 3063 | 0.057 |
| 3064 | -0.074 |
| 3065 | 0.063 |
| 3066 | 0.038 |
| 3067 | -0.031 |
| 3068 | -0.017 |
| 3069 | -0.098 |
| 3070 | -0.022 |
| 3071 | -0.042 |
| 3072 | -0.118 |
| 3073 | -0.085 |
| 3074 | -0.089 |
| 3075 | -0.038 |
| 3076 | -0.067 |
| 3077 | 0.140 |
| 3078 | -0.020 |
| 3079 | -0.057 |
| 3080 | 0.003 |
| 3081 | -0.023 |
| 3082 | 0.082 |
| 3083 | -0.060 |
| 3084 | 0.057 |
| 3085 | -0.017 |
| 3086 | 0.074 |
| 3087 | 0.100 |
| 3088 | -0.065 |
| 3089 | -0.080 |
| 3090 | 0.001 |
| 3091 | 0.024 |
| 3092 | -0.031 |
| 3093 | 0.000 |
| 3094 | 0.045 |
| 3095 | 0.160 |
| 3096 | -0.075 |

| | |
|---|---|
| 3097 | 0.025 |
| 3098 | -0.026 |
| 3099 | 0.018 |
| 3100 | 0.115 |
| 3101 | 0.075 |
| 3102 | 0.022 |
| 3103 | 0.050 |
| 3104 | -0.079 |
| 3105 | 0.033 |
| 3106 | 0.080 |
| 3107 | -0.089 |
| 3108 | 0.001 |
| 3109 | 0.011 |
| 3110 | 0.073 |
| 3111 | -0.046 |
| 3112 | -0.097 |
| 3113 | -0.100 |
| 3114 | -0.124 |
| 3115 | 0.041 |
| 3116 | -0.006 |
| 3117 | -0.009 |
| 3118 | -0.068 |
| 3119 | -0.011 |
| 3120 | 0.098 |
| 3121 | 0.052 |
| 3122 | -0.016 |
| 3123 | 0.097 |
| 3124 | -0.046 |
| 3125 | -0.063 |
| 3126 | 0.015 |
| 3127 | 0.027 |
| 3128 | 0.120 |
| 3129 | -0.070 |
| 3130 | -0.063 |
| 3131 | 0.037 |
| 3132 | 0.032 |
| 3133 | 0.062 |
| 3134 | 0.043 |
| 3135 | 0.075 |
| 3136 | 0.003 |
| 3137 | 0.043 |
| 3138 | -0.041 |
| 3139 | 0.173 |
| 3140 | -0.002 |
| 3141 | 0.043 |
| 3142 | -0.013 |
| 3143 | 0.052 |
| 3144 | -0.067 |
| 3145 | 0.030 |
| 3146 | -0.016 |

| | |
|---|---|
| 3147 | 0.096 |
| 3148 | -0.006 |
| 3149 | -0.039 |
| 3150 | 0.146 |
| 3151 | 0.029 |
| 3152 | -0.190 |
| 3153 | -0.004 |
| 3154 | -0.008 |
| 3155 | 0.040 |
| 3156 | 0.060 |
| 3157 | 0.044 |
| 3158 | -0.008 |
| 3159 | 0.117 |
| 3160 | -0.058 |
| 3161 | -0.099 |
| 3162 | 0.066 |
| 3163 | -0.036 |
| 3164 | -0.044 |
| 3165 | 0.033 |
| 3166 | -0.033 |
| 3167 | 0.003 |
| 3168 | -0.025 |
| 3169 | 0.014 |
| 3170 | 0.152 |
| 3171 | 0.029 |
| 3172 | 0.040 |
| 3173 | 0.016 |
| 3174 | 0.059 |
| 3175 | 0.037 |
| 3176 | 0.068 |
| 3177 | -0.057 |
| 3178 | 0.009 |
| 3179 | 0.038 |
| 3180 | -0.028 |
| 3181 | -0.067 |
| 3182 | 0.099 |
| 3183 | 0.076 |
| 3184 | -0.016 |
| 3185 | 0.067 |
| 3186 | 0.053 |
| 3187 | -0.048 |
| 3188 | 0.022 |
| 3189 | 0.037 |
| 3190 | -0.166 |
| 3191 | -0.022 |
| 3192 | -0.002 |
| 3193 | -0.098 |
| 3194 | -0.052 |
| 3195 | 0.107 |
| 3196 | -0.037 |

| | |
|---|---|
| 3197 | -0.046 |
| 3198 | -0.119 |
| 3199 | 0.021 |
| 3200 | -0.049 |
| 3201 | 0.008 |
| 3202 | 0.040 |
| 3203 | 0.033 |
| 3204 | -0.032 |
| 3205 | -0.025 |
| 3206 | -0.065 |
| 3207 | 0.054 |
| 3208 | 0.015 |
| 3209 | -0.074 |
| 3210 | -0.026 |
| 3211 | -0.021 |
| 3212 | 0.054 |
| 3213 | -0.045 |
| 3214 | -0.022 |
| 3215 | 0.034 |
| 3216 | -0.008 |
| 3217 | -0.007 |
| 3218 | -0.086 |
| 3219 | -0.133 |
| 3220 | 0.076 |
| 3221 | 0.142 |
| 3222 | -0.116 |
| 3223 | 0.038 |
| 3224 | 0.086 |
| 3225 | -0.003 |
| 3226 | -0.137 |
| 3227 | 0.082 |
| 3228 | -0.001 |
| 3229 | -0.007 |
| 3230 | -0.004 |
| 3231 | 0.018 |
| 3232 | 0.049 |
| 3233 | 0.084 |
| 3234 | -0.086 |
| 3235 | 0.085 |
| 3236 | -0.041 |
| 3237 | -0.071 |
| 3238 | 0.105 |
| 3239 | 0.034 |
| 3240 | 0.091 |
| 3241 | 0.021 |
| 3242 | 0.038 |
| 3243 | -0.010 |
| 3244 | 0.021 |
| 3245 | -0.018 |
| 3246 | 0.065 |

| | |
|---|---|
| 3247 | -0.090 |
| 3248 | -0.036 |
| 3249 | -0.042 |
| 3250 | -0.006 |
| 3251 | 0.028 |
| 3252 | 0.010 |
| 3253 | -0.039 |
| 3254 | 0.085 |
| 3255 | -0.026 |
| 3256 | -0.194 |
| 3257 | -0.007 |
| 3258 | 0.076 |
| 3259 | -0.050 |
| 3260 | 0.048 |
| 3261 | -0.042 |
| 3262 | -0.066 |
| 3263 | 0.024 |
| 3264 | 0.080 |
| 3265 | -0.044 |
| 3266 | -0.067 |
| 3267 | 0.085 |
| 3268 | -0.049 |
| 3269 | 0.033 |
| 3270 | 0.088 |
| 3271 | 0.093 |
| 3272 | -0.053 |
| 3273 | 0.076 |
| 3274 | -0.080 |
| 3275 | -0.030 |
| 3276 | 0.037 |
| 3277 | 0.065 |
| 3278 | 0.030 |
| 3279 | 0.069 |
| 3280 | -0.055 |
| 3281 | -0.087 |
| 3282 | 0.104 |
| 3283 | 0.014 |
| 3284 | 0.047 |
| 3285 | -0.007 |
| 3286 | 0.085 |
| 3287 | 0.018 |
| 3288 | 0.070 |
| 3289 | -0.045 |
| 3290 | -0.016 |
| 3291 | -0.008 |
| 3292 | 0.037 |
| 3293 | -0.092 |
| 3294 | -0.019 |
| 3295 | -0.174 |
| 3296 | 0.005 |

| | |
|---|---|
| 3297 | 0.016 |
| 3298 | 0.044 |
| 3299 | 0.110 |
| 3300 | -0.002 |
| 3301 | -0.042 |
| 3302 | -0.016 |
| 3303 | 0.073 |
| 3304 | 0.037 |
| 3305 | -0.025 |
| 3306 | 0.005 |
| 3307 | -0.017 |
| 3308 | -0.055 |
| 3309 | 0.054 |
| 3310 | -0.026 |
| 3311 | 0.043 |
| 3312 | 0.003 |
| 3313 | -0.017 |
| 3314 | -0.014 |
| 3315 | 0.059 |
| 3316 | 0.010 |
| 3317 | -0.102 |
| 3318 | 0.050 |
| 3319 | -0.138 |
| 3320 | 0.178 |
| 3321 | -0.015 |
| 3322 | 0.134 |
| 3323 | 0.060 |
| 3324 | -0.068 |
| 3325 | -0.018 |
| 3326 | -0.003 |
| 3327 | -0.016 |
| 3328 | 0.038 |
| 3329 | 0.027 |
| 3330 | 0.069 |
| 3331 | 0.063 |
| 3332 | -0.041 |
| 3333 | -0.027 |
| 3334 | 0.114 |
| 3335 | -0.083 |
| 3336 | -0.022 |
| 3337 | 0.074 |
| 3338 | -0.139 |
| 3339 | 0.081 |
| 3340 | 0.098 |
| 3341 | -0.048 |
| 3342 | -0.073 |
| 3343 | -0.012 |
| 3344 | 0.124 |
| 3345 | -0.056 |
| 3346 | 0.075 |

| | |
|---|---|
| 3347 | -0.126 |
| 3348 | -0.043 |
| 3349 | 0.032 |
| 3350 | 0.009 |
| 3351 | 0.032 |
| 3352 | 0.101 |
| 3353 | -0.023 |
| 3354 | -0.129 |
| 3355 | -0.008 |
| 3356 | -0.003 |
| 3357 | -0.019 |
| 3358 | -0.029 |
| 3359 | -0.018 |
| 3360 | -0.036 |
| 3361 | 0.032 |
| 3362 | -0.020 |
| 3363 | -0.116 |
| 3364 | 0.013 |
| 3365 | -0.004 |
| 3366 | 0.085 |
| 3367 | 0.019 |
| 3368 | -0.057 |
| 3369 | 0.056 |
| 3370 | -0.012 |
| 3371 | -0.059 |
| 3372 | 0.091 |
| 3373 | 0.096 |
| 3374 | 0.002 |
| 3375 | 0.026 |
| 3376 | 0.084 |
| 3377 | 0.053 |
| 3378 | 0.012 |
| 3379 | 0.034 |
| 3380 | -0.011 |
| 3381 | 0.007 |
| 3382 | -0.131 |
| 3383 | 0.035 |
| 3384 | -0.108 |
| 3385 | 0.027 |
| 3386 | 0.014 |
| 3387 | -0.139 |
| 3388 | -0.019 |
| 3389 | 0.129 |
| 3390 | 0.121 |
| 3391 | 0.003 |
| 3392 | -0.117 |
| 3393 | 0.016 |
| 3394 | -0.094 |
| 3395 | -0.004 |
| 3396 | -0.068 |

| | |
|---|---|
| 3397 | 0.000 |
| 3398 | 0.078 |
| 3399 | -0.021 |
| 3400 | 0.134 |
| 3401 | -0.105 |
| 3402 | 0.073 |
| 3403 | 0.041 |
| 3404 | -0.033 |
| 3405 | 0.011 |
| 3406 | 0.069 |
| 3407 | -0.050 |
| 3408 | -0.064 |
| 3409 | 0.037 |
| 3410 | 0.035 |
| 3411 | 0.129 |
| 3412 | 0.023 |
| 3413 | -0.018 |
| 3414 | 0.036 |
| 3415 | -0.112 |
| 3416 | 0.097 |
| 3417 | 0.105 |
| 3418 | -0.009 |
| 3419 | -0.041 |
| 3420 | 0.045 |
| 3421 | 0.015 |
| 3422 | -0.092 |
| 3423 | -0.066 |
| 3424 | 0.056 |
| 3425 | -0.091 |
| 3426 | 0.028 |
| 3427 | 0.055 |
| 3428 | 0.045 |
| 3429 | 0.006 |
| 3430 | -0.007 |
| 3431 | 0.052 |
| 3432 | 0.016 |
| 3433 | -0.027 |
| 3434 | 0.077 |
| 3435 | -0.061 |
| 3436 | 0.011 |
| 3437 | 0.005 |
| 3438 | 0.008 |
| 3439 | 0.026 |
| 3440 | 0.109 |
| 3441 | -0.043 |
| 3442 | -0.037 |
| 3443 | 0.020 |
| 3444 | -0.035 |
| 3445 | -0.005 |
| 3446 | -0.037 |

| | |
|---|---|
| 3447 | -0.062 |
| 3448 | 0.052 |
| 3449 | -0.062 |
| 3450 | -0.144 |
| 3451 | 0.093 |
| 3452 | -0.065 |
| 3453 | 0.076 |
| 3454 | 0.091 |
| 3455 | 0.035 |
| 3456 | 0.090 |
| 3457 | 0.044 |
| 3458 | 0.111 |
| 3459 | -0.003 |
| 3460 | -0.002 |
| 3461 | -0.035 |
| 3462 | 0.065 |
| 3463 | 0.036 |
| 3464 | 0.037 |
| 3465 | 0.054 |
| 3466 | -0.012 |
| 3467 | 0.039 |
| 3468 | 0.057 |
| 3469 | 0.096 |
| 3470 | -0.047 |
| 3471 | -0.004 |
| 3472 | -0.061 |
| 3473 | 0.111 |
| 3474 | -0.001 |
| 3475 | 0.003 |
| 3476 | -0.030 |
| 3477 | -0.011 |
| 3478 | -0.065 |
| 3479 | -0.059 |
| 3480 | -0.087 |
| 3481 | 0.173 |
| 3482 | -0.010 |
| 3483 | -0.109 |
| 3484 | -0.025 |
| 3485 | 0.053 |
| 3486 | 0.065 |
| 3487 | -0.042 |
| 3488 | 0.019 |
| 3489 | 0.031 |
| 3490 | 0.046 |
| 3491 | -0.047 |
| 3492 | 0.047 |
| 3493 | -0.128 |
| 3494 | -0.109 |
| 3495 | 0.038 |
| 3496 | -0.061 |

| | |
|---|---|
| 3497 | 0.027 |
| 3498 | -0.128 |
| 3499 | -0.089 |
| 3500 | -0.088 |
| 3501 | -0.134 |
| 3502 | 0.038 |
| 3503 | -0.055 |
| 3504 | 0.014 |
| 3505 | 0.008 |
| 3506 | 0.109 |
| 3507 | 0.035 |
| 3508 | -0.024 |
| 3509 | 0.036 |
| 3510 | -0.004 |
| 3511 | 0.002 |
| 3512 | -0.125 |
| 3513 | 0.007 |
| 3514 | -0.098 |
| 3515 | 0.023 |
| 3516 | -0.008 |
| 3517 | -0.065 |
| 3518 | 0.061 |
| 3519 | -0.045 |
| 3520 | 0.097 |
| 3521 | 0.057 |
| 3522 | -0.061 |
| 3523 | 0.003 |
| 3524 | 0.204 |
| 3525 | 0.024 |
| 3526 | 0.179 |
| 3527 | 0.057 |
| 3528 | 0.006 |
| 3529 | -0.090 |
| 3530 | 0.043 |
| 3531 | 0.028 |
| 3532 | 0.052 |
| 3533 | 0.029 |
| 3534 | -0.100 |
| 3535 | 0.067 |
| 3536 | -0.051 |
| 3537 | -0.091 |
| 3538 | -0.123 |
| 3539 | 0.004 |
| 3540 | -0.029 |
| 3541 | -0.068 |
| 3542 | -0.022 |
| 3543 | 0.029 |
| 3544 | -0.143 |
| 3545 | -0.030 |
| 3546 | 0.059 |

| | |
|---|---|
| 3547 | 0.025 |
| 3548 | -0.016 |
| 3549 | 0.070 |
| 3550 | 0.051 |
| 3551 | -0.109 |
| 3552 | -0.059 |
| 3553 | 0.070 |
| 3554 | -0.041 |
| 3555 | -0.083 |
| 3556 | -0.092 |
| 3557 | -0.041 |
| 3558 | 0.038 |
| 3559 | -0.026 |
| 3560 | -0.132 |
| 3561 | 0.051 |
| 3562 | 0.055 |
| 3563 | 0.093 |
| 3564 | 0.036 |
| 3565 | 0.037 |
| 3566 | -0.003 |
| 3567 | -0.027 |
| 3568 | 0.116 |
| 3569 | -0.136 |
| 3570 | 0.112 |
| 3571 | 0.016 |
| 3572 | -0.016 |
| 3573 | -0.085 |
| 3574 | -0.086 |
| 3575 | 0.035 |
| 3576 | -0.009 |
| 3577 | -0.046 |
| 3578 | 0.021 |
| 3579 | 0.009 |
| 3580 | 0.015 |
| 3581 | 0.063 |
| 3582 | 0.014 |
| 3583 | 0.127 |
| 3584 | 0.048 |
| 3585 | -0.013 |
| 3586 | -0.002 |
| 3587 | 0.049 |
| 3588 | -0.024 |
| 3589 | 0.114 |
| 3590 | 0.081 |
| 3591 | 0.048 |
| 3592 | -0.096 |
| 3593 | -0.018 |
| 3594 | 0.001 |
| 3595 | 0.021 |
| 3596 | -0.064 |

| | |
|---|---|
| 3597 | 0.067 |
| 3598 | -0.028 |
| 3599 | 0.023 |
| 3600 | 0.029 |
| 3601 | 0.065 |
| 3602 | -0.125 |
| 3603 | -0.084 |
| 3604 | -0.012 |
| 3605 | 0.030 |
| 3606 | 0.091 |
| 3607 | -0.038 |
| 3608 | -0.005 |
| 3609 | 0.062 |
| 3610 | 0.170 |
| 3611 | -0.097 |
| 3612 | 0.026 |
| 3613 | 0.036 |
| 3614 | -0.060 |
| 3615 | 0.031 |
| 3616 | -0.064 |
| 3617 | 0.017 |
| 3618 | -0.004 |
| 3619 | 0.039 |
| 3620 | -0.005 |
| 3621 | 0.002 |
| 3622 | 0.009 |
| 3623 | -0.061 |
| 3624 | -0.047 |
| 3625 | 0.122 |
| 3626 | 0.098 |
| 3627 | 0.010 |
| 3628 | -0.009 |
| 3629 | 0.090 |
| 3630 | 0.044 |
| 3631 | 0.015 |
| 3632 | 0.015 |
| 3633 | -0.009 |
| 3634 | -0.077 |
| 3635 | -0.026 |
| 3636 | 0.082 |
| 3637 | 0.041 |
| 3638 | -0.009 |
| 3639 | -0.034 |
| 3640 | -0.026 |
| 3641 | -0.109 |
| 3642 | -0.137 |
| 3643 | 0.007 |
| 3644 | 0.036 |
| 3645 | -0.095 |
| 3646 | -0.137 |

| | |
|---|---|
| 3647 | 0.054 |
| 3648 | -0.043 |
| 3649 | -0.027 |
| 3650 | 0.046 |
| 3651 | 0.039 |
| 3652 | 0.183 |
| 3653 | 0.079 |
| 3654 | -0.086 |
| 3655 | 0.041 |
| 3656 | 0.047 |
| 3657 | 0.038 |
| 3658 | -0.151 |
| 3659 | 0.044 |
| 3660 | -0.097 |
| 3661 | -0.047 |
| 3662 | -0.159 |
| 3663 | 0.125 |
| 3664 | -0.029 |
| 3665 | 0.115 |
| 3666 | -0.160 |
| 3667 | -0.073 |
| 3668 | -0.136 |
| 3669 | -0.081 |
| 3670 | -0.027 |
| 3671 | 0.008 |
| 3672 | 0.095 |
| 3673 | -0.012 |
| 3674 | 0.032 |
| 3675 | -0.004 |
| 3676 | 0.037 |
| 3677 | 0.043 |
| 3678 | -0.011 |
| 3679 | 0.012 |
| 3680 | 0.013 |
| 3681 | 0.063 |
| 3682 | 0.022 |
| 3683 | -0.096 |
| 3684 | 0.039 |
| 3685 | 0.006 |
| 3686 | 0.088 |
| 3687 | 0.004 |
| 3688 | 0.007 |
| 3689 | -0.003 |
| 3690 | -0.028 |
| 3691 | -0.086 |
| 3692 | 0.090 |
| 3693 | 0.089 |
| 3694 | 0.186 |
| 3695 | 0.038 |
| 3696 | -0.007 |

| | |
|---|---|
| 3697 | -0.007 |
| 3698 | 0.014 |
| 3699 | 0.060 |
| 3700 | 0.004 |
| 3701 | -0.040 |
| 3702 | 0.162 |
| 3703 | -0.039 |
| 3704 | -0.029 |
| 3705 | 0.086 |
| 3706 | 0.075 |
| 3707 | 0.045 |
| 3708 | 0.039 |
| 3709 | -0.152 |
| 3710 | -0.009 |
| 3711 | -0.043 |
| 3712 | -0.097 |
| 3713 | 0.053 |
| 3714 | 0.008 |
| 3715 | 0.002 |
| 3716 | -0.114 |
| 3717 | -0.047 |
| 3718 | -0.044 |
| 3719 | 0.065 |
| 3720 | -0.098 |
| 3721 | -0.098 |
| 3722 | -0.111 |
| 3723 | 0.055 |
| 3724 | 0.003 |
| 3725 | 0.174 |
| 3726 | -0.026 |
| 3727 | -0.066 |
| 3728 | -0.178 |
| 3729 | 0.070 |
| 3730 | -0.068 |
| 3731 | 0.004 |
| 3732 | -0.106 |
| 3733 | -0.072 |
| 3734 | -0.003 |
| 3735 | -0.172 |
| 3736 | -0.164 |
| 3737 | -0.063 |
| 3738 | -0.040 |
| 3739 | -0.089 |
| 3740 | -0.060 |
| 3741 | -0.054 |
| 3742 | -0.053 |
| 3743 | -0.014 |
| 3744 | 0.043 |
| 3745 | -0.012 |
| 3746 | 0.074 |

| | |
|---|---|
| 3747 | -0.106 |
| 3748 | -0.120 |
| 3749 | 0.085 |
| 3750 | -0.191 |
| 3751 | -0.046 |
| 3752 | 0.010 |
| 3753 | -0.139 |
| 3754 | 0.017 |
| 3755 | 0.073 |
| 3756 | 0.067 |
| 3757 | 0.023 |
| 3758 | -0.027 |
| 3759 | -0.032 |
| 3760 | -0.018 |
| 3761 | -0.014 |
| 3762 | 0.006 |
| 3763 | -0.067 |
| 3764 | 0.102 |
| 3765 | 0.019 |
| 3766 | -0.030 |
| 3767 | -0.095 |
| 3768 | -0.001 |
| 3769 | 0.018 |
| 3770 | 0.128 |
| 3771 | -0.048 |
| 3772 | 0.032 |
| 3773 | -0.011 |
| 3774 | 0.094 |
| 3775 | 0.048 |
| 3776 | 0.077 |
| 3777 | -0.010 |
| 3778 | -0.071 |
| 3779 | 0.137 |
| 3780 | 0.153 |
| 3781 | -0.186 |
| 3782 | 0.053 |
| 3783 | -0.004 |
| 3784 | 0.112 |
| 3785 | 0.011 |
| 3786 | -0.122 |
| 3787 | -0.058 |
| 3788 | 0.020 |
| 3789 | 0.078 |
| 3790 | 0.032 |
| 3791 | 0.093 |
| 3792 | 0.045 |
| 3793 | 0.013 |
| 3794 | 0.117 |
| 3795 | 0.012 |
| 3796 | 0.078 |

| | |
|---|---|
| 3797 | 0.100 |
| 3798 | -0.081 |
| 3799 | 0.111 |
| 3800 | -0.053 |
| 3801 | -0.015 |
| 3802 | -0.054 |
| 3803 | -0.024 |
| 3804 | 0.006 |
| 3805 | 0.018 |
| 3806 | -0.108 |
| 3807 | -0.086 |
| 3808 | 0.086 |
| 3809 | 0.011 |
| 3810 | -0.129 |
| 3811 | -0.021 |
| 3812 | 0.095 |
| 3813 | 0.010 |
| 3814 | 0.092 |
| 3815 | -0.037 |
| 3816 | -0.083 |
| 3817 | 0.103 |
| 3818 | 0.038 |
| 3819 | -0.096 |
| 3820 | 0.082 |
| 3821 | -0.074 |
| 3822 | 0.048 |
| 3823 | 0.046 |
| 3824 | 0.092 |
| 3825 | -0.011 |
| 3826 | 0.043 |
| 3827 | 0.011 |
| 3828 | 0.047 |
| 3829 | -0.105 |
| 3830 | 0.016 |
| 3831 | -0.135 |
| 3832 | 0.029 |
| 3833 | 0.018 |
| 3834 | 0.027 |
| 3835 | -0.066 |
| 3836 | -0.051 |
| 3837 | -0.116 |
| 3838 | -0.071 |
| 3839 | 0.048 |
| 3840 | 0.041 |
| 3841 | -0.001 |
| 3842 | -0.065 |
| 3843 | -0.010 |
| 3844 | -0.045 |
| 3845 | -0.013 |
| 3846 | 0.122 |

| | |
|---|---|
| 3847 | -0.006 |
| 3848 | -0.067 |
| 3849 | 0.117 |
| 3850 | -0.072 |
| 3851 | -0.052 |
| 3852 | -0.087 |
| 3853 | -0.105 |
| 3854 | -0.160 |
| 3855 | -0.022 |
| 3856 | -0.006 |
| 3857 | 0.019 |
| 3858 | 0.050 |
| 3859 | 0.070 |
| 3860 | 0.053 |
| 3861 | -0.072 |
| 3862 | -0.052 |
| 3863 | 0.084 |
| 3864 | 0.029 |
| 3865 | 0.110 |
| 3866 | -0.040 |
| 3867 | -0.025 |
| 3868 | -0.011 |
| 3869 | -0.080 |
| 3870 | 0.034 |
| 3871 | 0.050 |
| 3872 | 0.003 |
| 3873 | 0.016 |
| 3874 | -0.044 |
| 3875 | -0.027 |
| 3876 | 0.022 |
| 3877 | -0.053 |
| 3878 | -0.054 |
| 3879 | -0.014 |
| 3880 | -0.025 |
| 3881 | -0.045 |
| 3882 | -0.081 |
| 3883 | 0.016 |
| 3884 | 0.061 |
| 3885 | 0.107 |
| 3886 | -0.017 |
| 3887 | -0.069 |
| 3888 | 0.107 |
| 3889 | 0.116 |
| 3890 | 0.022 |
| 3891 | 0.080 |
| 3892 | 0.161 |
| 3893 | -0.101 |
| 3894 | -0.057 |
| 3895 | 0.076 |
| 3896 | -0.033 |

| | |
|---|---|
| 3897 | 0.035 |
| 3898 | -0.067 |
| 3899 | -0.102 |
| 3900 | -0.006 |
| 3901 | 0.122 |
| 3902 | 0.108 |
| 3903 | 0.080 |
| 3904 | -0.111 |
| 3905 | -0.009 |
| 3906 | 0.095 |
| 3907 | -0.008 |
| 3908 | 0.026 |
| 3909 | 0.013 |
| 3910 | -0.066 |
| 3911 | 0.103 |
| 3912 | -0.063 |
| 3913 | -0.057 |
| 3914 | 0.089 |
| 3915 | -0.020 |
| 3916 | -0.026 |
| 3917 | 0.005 |
| 3918 | -0.009 |
| 3919 | -0.027 |
| 3920 | 0.017 |
| 3921 | -0.039 |
| 3922 | 0.035 |
| 3923 | 0.043 |
| 3924 | 0.003 |
| 3925 | -0.032 |
| 3926 | -0.075 |
| 3927 | 0.006 |
| 3928 | 0.110 |
| 3929 | 0.035 |
| 3930 | -0.055 |
| 3931 | -0.079 |
| 3932 | 0.005 |
| 3933 | 0.057 |
| 3934 | -0.021 |
| 3935 | 0.029 |
| 3936 | 0.009 |
| 3937 | 0.048 |
| 3938 | 0.132 |
| 3939 | -0.164 |
| 3940 | -0.040 |
| 3941 | -0.038 |
| 3942 | 0.000 |
| 3943 | 0.022 |
| 3944 | -0.053 |
| 3945 | 0.008 |
| 3946 | -0.006 |

| | |
|---|---|
| 3947 | 0.017 |
| 3948 | -0.041 |
| 3949 | -0.008 |
| 3950 | 0.022 |
| 3951 | 0.037 |
| 3952 | -0.020 |
| 3953 | 0.040 |
| 3954 | -0.028 |
| 3955 | 0.005 |
| 3956 | 0.081 |
| 3957 | -0.083 |
| 3958 | 0.017 |
| 3959 | 0.014 |
| 3960 | -0.084 |
| 3961 | 0.015 |
| 3962 | 0.076 |
| 3963 | -0.033 |
| 3964 | -0.024 |
| 3965 | -0.022 |
| 3966 | -0.089 |
| 3967 | -0.004 |
| 3968 | -0.054 |
| 3969 | 0.066 |
| 3970 | -0.018 |
| 3971 | 0.085 |
| 3972 | 0.013 |
| 3973 | 0.075 |
| 3974 | -0.043 |
| 3975 | 0.008 |
| 3976 | 0.016 |
| 3977 | 0.018 |
| 3978 | 0.004 |
| 3979 | 0.143 |
| 3980 | 0.072 |
| 3981 | 0.053 |
| 3982 | -0.096 |
| 3983 | -0.079 |
| 3984 | 0.050 |
| 3985 | 0.099 |
| 3986 | -0.053 |
| 3987 | -0.018 |
| 3988 | 0.033 |
| 3989 | -0.084 |
| 3990 | 0.081 |
| 3991 | -0.113 |
| 3992 | 0.047 |
| 3993 | -0.053 |
| 3994 | 0.008 |
| 3995 | -0.017 |
| 3996 | -0.083 |

| | |
|---|---|
| 3997 | -0.005 |
| 3998 | -0.111 |
| 3999 | 0.058 |
| 4000 | -0.007 |
| 4001 | -0.051 |
| 4002 | -0.047 |
| 4003 | 0.052 |
| 4004 | -0.117 |
| 4005 | -0.021 |
| 4006 | 0.018 |
| 4007 | -0.081 |
| 4008 | 0.060 |
| 4009 | 0.041 |
| 4010 | -0.011 |
| 4011 | 0.043 |
| 4012 | 0.095 |
| 4013 | 0.022 |
| 4014 | 0.009 |
| 4015 | 0.151 |
| 4016 | 0.048 |
| 4017 | 0.016 |
| 4018 | -0.011 |
| 4019 | -0.065 |
| 4020 | -0.033 |
| 4021 | -0.020 |
| 4022 | -0.070 |
| 4023 | 0.134 |
| 4024 | 0.068 |
| 4025 | 0.088 |
| 4026 | -0.006 |
| 4027 | 0.047 |
| 4028 | 0.054 |
| 4029 | -0.145 |
| 4030 | -0.016 |
| 4031 | -0.036 |
| 4032 | -0.017 |
| 4033 | -0.154 |
| 4034 | 0.063 |
| 4035 | 0.011 |
| 4036 | 0.038 |
| 4037 | 0.023 |
| 4038 | 0.057 |
| 4039 | -0.040 |
| 4040 | 0.016 |
| 4041 | 0.014 |
| 4042 | -0.019 |
| 4043 | -0.163 |
| 4044 | -0.015 |
| 4045 | 0.066 |
| 4046 | -0.114 |

| | |
|---|---|
| 4047 | -0.160 |
| 4048 | -0.044 |
| 4049 | 0.056 |
| 4050 | -0.022 |
| 4051 | 0.023 |
| 4052 | -0.112 |
| 4053 | -0.077 |
| 4054 | -0.050 |
| 4055 | 0.070 |
| 4056 | -0.158 |
| 4057 | 0.072 |
| 4058 | 0.028 |
| 4059 | 0.003 |
| 4060 | 0.037 |
| 4061 | -0.030 |
| 4062 | -0.007 |
| 4063 | -0.004 |
| 4064 | 0.019 |
| 4065 | -0.041 |
| 4066 | 0.001 |
| 4067 | -0.110 |
| 4068 | -0.110 |
| 4069 | 0.006 |
| 4070 | -0.066 |
| 4071 | 0.031 |
| 4072 | 0.112 |
| 4073 | -0.100 |
| 4074 | 0.026 |
| 4075 | 0.052 |
| 4076 | 0.102 |
| 4077 | 0.089 |
| 4078 | -0.045 |
| 4079 | 0.095 |
| 4080 | 0.046 |
| 4081 | -0.026 |
| 4082 | -0.057 |
| 4083 | -0.002 |
| 4084 | 0.056 |
| 4085 | -0.029 |
| 4086 | 0.115 |
| 4087 | -0.028 |
| 4088 | -0.135 |
| 4089 | 0.015 |
| 4090 | 0.072 |
| 4091 | 0.007 |
| 4092 | -0.045 |
| 4093 | 0.007 |
| 4094 | -0.055 |
| 4095 | -0.043 |
| 4096 | 0.048 |

| | |
|---|---|
| 4097 | -0.111 |
| 4098 | -0.026 |
| 4099 | 0.062 |
| 4100 | -0.152 |
| 4101 | -0.070 |
| 4102 | -0.001 |
| 4103 | -0.043 |
| 4104 | -0.074 |
| 4105 | -0.096 |
| 4106 | -0.064 |
| 4107 | -0.035 |
| 4108 | 0.001 |
| 4109 | 0.044 |
| 4110 | 0.026 |
| 4111 | 0.021 |
| 4112 | 0.045 |
| 4113 | -0.023 |
| 4114 | -0.009 |
| 4115 | 0.091 |
| 4116 | 0.090 |
| 4117 | -0.064 |
| 4118 | 0.056 |
| 4119 | 0.036 |
| 4120 | -0.041 |
| 4121 | 0.059 |
| 4122 | -0.080 |
| 4123 | 0.060 |
| 4124 | 0.137 |
| 4125 | -0.051 |
| 4126 | -0.093 |
| 4127 | 0.030 |
| 4128 | -0.021 |
| 4129 | -0.007 |
| 4130 | -0.086 |
| 4131 | 0.001 |
| 4132 | -0.032 |
| 4133 | 0.075 |
| 4134 | 0.130 |
| 4135 | 0.056 |
| 4136 | -0.126 |
| 4137 | 0.202 |
| 4138 | -0.034 |
| 4139 | -0.040 |
| 4140 | -0.014 |
| 4141 | -0.087 |
| 4142 | 0.016 |
| 4143 | -0.054 |
| 4144 | 0.003 |
| 4145 | 0.068 |
| 4146 | 0.024 |

| | |
|---|---|
| 4147 | 0.015 |
| 4148 | -0.060 |
| 4149 | -0.037 |
| 4150 | 0.010 |
| 4151 | 0.080 |
| 4152 | 0.173 |
| 4153 | -0.043 |
| 4154 | 0.029 |
| 4155 | 0.043 |
| 4156 | -0.048 |
| 4157 | 0.111 |
| 4158 | -0.112 |
| 4159 | -0.055 |
| 4160 | 0.136 |
| 4161 | -0.002 |
| 4162 | -0.072 |
| 4163 | -0.041 |
| 4164 | 0.153 |
| 4165 | 0.074 |
| 4166 | 0.093 |
| 4167 | 0.003 |
| 4168 | 0.099 |
| 4169 | 0.041 |
| 4170 | -0.102 |
| 4171 | 0.078 |
| 4172 | -0.011 |
| 4173 | 0.120 |
| 4174 | -0.016 |
| 4175 | -0.066 |
| 4176 | 0.054 |
| 4177 | 0.018 |
| 4178 | 0.094 |
| 4179 | 0.101 |
| 4180 | 0.028 |
| 4181 | -0.028 |
| 4182 | 0.122 |
| 4183 | 0.013 |
| 4184 | 0.049 |
| 4185 | 0.010 |
| 4186 | -0.029 |
| 4187 | -0.083 |
| 4188 | -0.014 |
| 4189 | -0.020 |
| 4190 | 0.070 |
| 4191 | 0.085 |
| 4192 | -0.043 |
| 4193 | -0.043 |
| 4194 | -0.066 |
| 4195 | -0.037 |
| 4196 | 0.036 |

| | |
|---|---|
| 4197 | -0.015 |
| 4198 | 0.064 |
| 4199 | 0.133 |
| 4200 | 0.179 |
| 4201 | 0.036 |
| 4202 | -0.036 |
| 4203 | 0.049 |
| 4204 | 0.048 |
| 4205 | 0.082 |
| 4206 | -0.062 |
| 4207 | -0.133 |
| 4208 | -0.006 |
| 4209 | 0.054 |
| 4210 | 0.062 |
| 4211 | 0.011 |
| 4212 | 0.035 |
| 4213 | -0.120 |
| 4214 | 0.002 |
| 4215 | -0.052 |
| 4216 | -0.051 |
| 4217 | -0.064 |
| 4218 | 0.006 |
| 4219 | -0.104 |
| 4220 | 0.075 |
| 4221 | -0.048 |
| 4222 | -0.062 |
| 4223 | -0.063 |
| 4224 | -0.013 |
| 4225 | 0.011 |
| 4226 | 0.028 |
| 4227 | -0.023 |
| 4228 | -0.049 |
| 4229 | 0.008 |
| 4230 | 0.090 |
| 4231 | 0.066 |
| 4232 | -0.032 |
| 4233 | 0.049 |
| 4234 | 0.030 |
| 4235 | -0.062 |
| 4236 | 0.032 |
| 4237 | -0.142 |
| 4238 | -0.026 |
| 4239 | -0.020 |
| 4240 | -0.005 |
| 4241 | -0.057 |
| 4242 | 0.033 |
| 4243 | -0.061 |
| 4244 | 0.023 |
| 4245 | -0.001 |
| 4246 | -0.018 |

| | |
|---|---|
| 4247 | -0.020 |
| 4248 | 0.086 |
| 4249 | -0.009 |
| 4250 | 0.033 |
| 4251 | 0.076 |
| 4252 | -0.091 |
| 4253 | -0.043 |
| 4254 | -0.045 |
| 4255 | 0.024 |
| 4256 | 0.114 |
| 4257 | 0.085 |
| 4258 | 0.047 |
| 4259 | 0.042 |
| 4260 | 0.011 |
| 4261 | 0.038 |
| 4262 | -0.138 |
| 4263 | -0.070 |
| 4264 | -0.067 |
| 4265 | -0.074 |
| 4266 | 0.081 |
| 4267 | -0.097 |
| 4268 | -0.067 |
| 4269 | -0.047 |
| 4270 | -0.088 |
| 4271 | 0.010 |
| 4272 | -0.024 |
| 4273 | -0.039 |
| 4274 | -0.021 |
| 4275 | 0.087 |
| 4276 | 0.022 |
| 4277 | 0.036 |
| 4278 | 0.016 |
| 4279 | 0.075 |
| 4280 | -0.090 |
| 4281 | 0.058 |
| 4282 | -0.032 |
| 4283 | 0.055 |
| 4284 | 0.071 |
| 4285 | -0.048 |
| 4286 | -0.042 |
| 4287 | 0.019 |
| 4288 | -0.008 |
| 4289 | 0.032 |
| 4290 | 0.008 |
| 4291 | -0.061 |
| 4292 | -0.030 |
| 4293 | 0.007 |
| 4294 | -0.033 |
| 4295 | 0.056 |
| 4296 | -0.110 |

| | |
|---|---|
| 4297 | 0.003 |
| 4298 | -0.056 |
| 4299 | 0.011 |
| 4300 | -0.031 |
| 4301 | -0.024 |
| 4302 | -0.011 |
| 4303 | -0.021 |
| 4304 | 0.008 |
| 4305 | -0.047 |
| 4306 | 0.087 |
| 4307 | 0.091 |
| 4308 | 0.027 |
| 4309 | -0.023 |
| 4310 | -0.075 |
| 4311 | 0.051 |
| 4312 | 0.107 |
| 4313 | 0.049 |
| 4314 | 0.062 |
| 4315 | 0.098 |
| 4316 | 0.022 |
| 4317 | 0.000 |
| 4318 | 0.036 |
| 4319 | -0.039 |
| 4320 | -0.112 |
| 4321 | 0.033 |
| 4322 | -0.069 |
| 4323 | 0.068 |
| 4324 | -0.094 |
| 4325 | -0.059 |
| 4326 | 0.032 |
| 4327 | -0.123 |
| 4328 | -0.004 |
| 4329 | -0.043 |
| 4330 | -0.057 |
| 4331 | 0.071 |
| 4332 | 0.037 |
| 4333 | -0.030 |
| 4334 | -0.226 |
| 4335 | -0.090 |
| 4336 | -0.105 |
| 4337 | 0.014 |
| 4338 | 0.007 |
| 4339 | -0.064 |
| 4340 | -0.003 |
| 4341 | -0.045 |
| 4342 | 0.027 |
| 4343 | -0.035 |
| 4344 | 0.073 |
| 4345 | 0.035 |
| 4346 | 0.125 |

| | |
|---|---|
| 4347 | 0.050 |
| 4348 | 0.029 |
| 4349 | 0.005 |
| 4350 | 0.103 |
| 4351 | -0.053 |
| 4352 | 0.054 |
| 4353 | -0.049 |
| 4354 | 0.114 |
| 4355 | 0.003 |
| 4356 | -0.008 |
| 4357 | -0.085 |
| 4358 | 0.002 |
| 4359 | -0.034 |
| 4360 | -0.039 |
| 4361 | -0.056 |
| 4362 | -0.072 |
| 4363 | -0.037 |
| 4364 | -0.028 |
| 4365 | 0.025 |
| 4366 | 0.010 |
| 4367 | 0.060 |
| 4368 | 0.121 |
| 4369 | -0.009 |
| 4370 | 0.060 |
| 4371 | -0.041 |
| 4372 | -0.084 |
| 4373 | 0.069 |
| 4374 | 0.060 |
| 4375 | -0.002 |
| 4376 | -0.069 |
| 4377 | 0.013 |
| 4378 | -0.026 |
| 4379 | 0.093 |
| 4380 | -0.030 |
| 4381 | 0.017 |
| 4382 | 0.050 |
| 4383 | -0.102 |
| 4384 | 0.028 |
| 4385 | -0.040 |
| 4386 | -0.040 |
| 4387 | -0.008 |
| 4388 | -0.076 |
| 4389 | -0.023 |
| 4390 | 0.037 |
| 4391 | 0.052 |
| 4392 | 0.033 |
| 4393 | 0.043 |
| 4394 | -0.025 |
| 4395 | -0.029 |
| 4396 | 0.047 |

| | |
|---|---|
| 4397 | 0.052 |
| 4398 | -0.010 |
| 4399 | -0.028 |
| 4400 | -0.060 |
| 4401 | 0.028 |
| 4402 | -0.066 |
| 4403 | -0.009 |
| 4404 | -0.063 |
| 4405 | -0.039 |
| 4406 | 0.117 |
| 4407 | -0.079 |
| 4408 | 0.041 |
| 4409 | -0.013 |
| 4410 | -0.047 |
| 4411 | -0.129 |
| 4412 | 0.032 |
| 4413 | 0.005 |
| 4414 | 0.016 |
| 4415 | -0.005 |
| 4416 | -0.017 |
| 4417 | -0.020 |
| 4418 | 0.004 |
| 4419 | 0.017 |
| 4420 | 0.015 |
| 4421 | 0.077 |
| 4422 | 0.032 |
| 4423 | -0.026 |
| 4424 | 0.096 |
| 4425 | -0.041 |
| 4426 | -0.049 |
| 4427 | -0.055 |
| 4428 | 0.012 |
| 4429 | 0.016 |
| 4430 | 0.050 |
| 4431 | 0.098 |
| 4432 | -0.119 |
| 4433 | -0.030 |
| 4434 | 0.040 |
| 4435 | 0.018 |
| 4436 | -0.139 |
| 4437 | 0.058 |
| 4438 | 0.090 |
| 4439 | -0.015 |
| 4440 | -0.113 |
| 4441 | -0.031 |
| 4442 | -0.037 |
| 4443 | -0.169 |
| 4444 | 0.045 |
| 4445 | -0.010 |
| 4446 | -0.034 |

| | |
|---|---|
| 4447 | -0.001 |
| 4448 | 0.026 |
| 4449 | 0.139 |
| 4450 | 0.001 |
| 4451 | -0.005 |
| 4452 | 0.152 |
| 4453 | 0.141 |
| 4454 | 0.004 |
| 4455 | -0.018 |
| 4456 | -0.115 |
| 4457 | -0.013 |
| 4458 | 0.095 |
| 4459 | 0.020 |
| 4460 | 0.079 |
| 4461 | 0.032 |
| 4462 | 0.038 |
| 4463 | -0.001 |
| 4464 | -0.031 |
| 4465 | 0.056 |
| 4466 | 0.027 |
| 4467 | -0.092 |
| 4468 | -0.009 |
| 4469 | 0.042 |
| 4470 | -0.140 |
| 4471 | 0.047 |
| 4472 | -0.068 |
| 4473 | 0.016 |
| 4474 | -0.036 |
| 4475 | 0.024 |
| 4476 | 0.086 |
| 4477 | 0.039 |
| 4478 | -0.008 |
| 4479 | -0.048 |
| 4480 | 0.108 |
| 4481 | -0.019 |
| 4482 | -0.058 |
| 4483 | 0.031 |
| 4484 | -0.008 |
| 4485 | 0.046 |
| 4486 | -0.070 |
| 4487 | 0.078 |
| 4488 | -0.041 |
| 4489 | 0.031 |
| 4490 | 0.060 |
| 4491 | 0.117 |
| 4492 | -0.069 |
| 4493 | -0.012 |
| 4494 | 0.040 |
| 4495 | 0.018 |
| 4496 | -0.135 |

| | |
|---|---|
| 4497 | 0.053 |
| 4498 | -0.118 |
| 4499 | -0.079 |
| 4500 | 0.076 |
| 4501 | 0.015 |
| 4502 | 0.061 |
| 4503 | 0.050 |
| 4504 | 0.119 |
| 4505 | -0.060 |
| 4506 | 0.021 |
| 4507 | 0.028 |
| 4508 | -0.001 |
| 4509 | -0.023 |
| 4510 | -0.059 |
| 4511 | 0.105 |
| 4512 | -0.045 |
| 4513 | -0.048 |
| 4514 | -0.014 |
| 4515 | -0.067 |
| 4516 | 0.118 |
| 4517 | -0.013 |
| 4518 | -0.167 |
| 4519 | -0.005 |
| 4520 | 0.045 |
| 4521 | -0.049 |
| 4522 | 0.142 |
| 4523 | 0.040 |
| 4524 | 0.005 |
| 4525 | 0.072 |
| 4526 | -0.044 |
| 4527 | -0.151 |
| 4528 | 0.065 |
| 4529 | -0.118 |
| 4530 | -0.141 |
| 4531 | 0.023 |
| 4532 | -0.038 |
| 4533 | 0.034 |
| 4534 | -0.006 |
| 4535 | 0.019 |
| 4536 | -0.069 |
| 4537 | -0.243 |
| 4538 | -0.004 |
| 4539 | -0.033 |
| 4540 | -0.055 |
| 4541 | 0.102 |
| 4542 | 0.027 |
| 4543 | -0.128 |
| 4544 | -0.064 |
| 4545 | 0.093 |
| 4546 | -0.054 |

| | |
|---|---|
| 4547 | -0.054 |
| 4548 | -0.008 |
| 4549 | 0.151 |
| 4550 | -0.017 |
| 4551 | -0.031 |
| 4552 | 0.127 |
| 4553 | -0.022 |
| 4554 | 0.034 |
| 4555 | 0.011 |
| 4556 | -0.023 |
| 4557 | 0.026 |
| 4558 | 0.100 |
| 4559 | -0.001 |
| 4560 | 0.008 |
| 4561 | -0.146 |
| 4562 | 0.137 |
| 4563 | 0.077 |
| 4564 | 0.022 |
| 4565 | -0.012 |
| 4566 | -0.109 |
| 4567 | 0.025 |
| 4568 | -0.072 |
| 4569 | 0.093 |
| 4570 | -0.003 |
| 4571 | 0.006 |
| 4572 | 0.048 |
| 4573 | 0.070 |
| 4574 | 0.053 |
| 4575 | -0.001 |
| 4576 | 0.037 |
| 4577 | -0.095 |
| 4578 | 0.063 |
| 4579 | 0.120 |
| 4580 | 0.028 |
| 4581 | -0.027 |
| 4582 | -0.045 |
| 4583 | -0.055 |
| 4584 | 0.022 |
| 4585 | 0.002 |
| 4586 | -0.026 |
| 4587 | -0.035 |
| 4588 | -0.039 |
| 4589 | 0.038 |
| 4590 | 0.128 |
| 4591 | 0.052 |
| 4592 | -0.081 |
| 4593 | 0.014 |
| 4594 | 0.028 |
| 4595 | -0.076 |
| 4596 | 0.036 |

| | |
|---|---|
| 4597 | 0.075 |
| 4598 | 0.060 |
| 4599 | -0.035 |
| 4600 | -0.047 |
| 4601 | -0.075 |
| 4602 | 0.107 |
| 4603 | 0.023 |
| 4604 | 0.013 |
| 4605 | 0.035 |
| 4606 | 0.087 |
| 4607 | 0.025 |
| 4608 | 0.000 |
| 4609 | -0.015 |
| 4610 | -0.026 |
| 4611 | -0.011 |
| 4612 | 0.080 |
| 4613 | -0.053 |
| 4614 | -0.049 |
| 4615 | -0.012 |
| 4616 | 0.053 |
| 4617 | -0.086 |
| 4618 | -0.032 |
| 4619 | -0.041 |
| 4620 | -0.099 |
| 4621 | 0.006 |
| 4622 | -0.001 |
| 4623 | -0.005 |
| 4624 | 0.062 |
| 4625 | 0.028 |
| 4626 | -0.005 |
| 4627 | 0.021 |
| 4628 | -0.063 |
| 4629 | -0.059 |
| 4630 | -0.034 |
| 4631 | 0.064 |
| 4632 | 0.109 |
| 4633 | 0.020 |
| 4634 | -0.066 |
| 4635 | 0.098 |
| 4636 | 0.028 |
| 4637 | -0.150 |
| 4638 | 0.118 |
| 4639 | -0.005 |
| 4640 | 0.074 |
| 4641 | -0.088 |
| 4642 | -0.091 |
| 4643 | 0.032 |
| 4644 | 0.133 |
| 4645 | -0.034 |
| 4646 | -0.050 |

| | |
|---|---|
| 4647 | -0.023 |
| 4648 | 0.009 |
| 4649 | 0.056 |
| 4650 | -0.014 |
| 4651 | 0.065 |
| 4652 | -0.125 |
| 4653 | -0.052 |
| 4654 | 0.013 |
| 4655 | 0.009 |
| 4656 | 0.044 |
| 4657 | -0.006 |
| 4658 | 0.003 |
| 4659 | -0.002 |
| 4660 | 0.035 |
| 4661 | -0.066 |
| 4662 | 0.106 |
| 4663 | -0.002 |
| 4664 | -0.046 |
| 4665 | -0.017 |
| 4666 | 0.189 |
| 4667 | 0.018 |
| 4668 | 0.061 |
| 4669 | 0.020 |
| 4670 | 0.074 |
| 4671 | -0.053 |
| 4672 | 0.031 |
| 4673 | -0.007 |
| 4674 | 0.005 |
| 4675 | -0.081 |
| 4676 | 0.022 |
| 4677 | -0.037 |
| 4678 | 0.001 |
| 4679 | 0.090 |
| 4680 | 0.006 |
| 4681 | 0.085 |
| 4682 | -0.025 |
| 4683 | 0.053 |
| 4684 | 0.014 |
| 4685 | -0.091 |
| 4686 | -0.042 |
| 4687 | 0.158 |
| 4688 | 0.076 |
| 4689 | -0.066 |
| 4690 | 0.031 |
| 4691 | -0.023 |
| 4692 | 0.043 |
| 4693 | -0.012 |
| 4694 | -0.086 |
| 4695 | -0.045 |
| 4696 | -0.050 |

| | |
|---|---|
| 4697 | 0.086 |
| 4698 | -0.003 |
| 4699 | -0.065 |
| 4700 | -0.026 |
| 4701 | 0.038 |
| 4702 | -0.126 |
| 4703 | 0.141 |
| 4704 | 0.024 |
| 4705 | 0.062 |
| 4706 | -0.017 |
| 4707 | 0.039 |
| 4708 | 0.017 |
| 4709 | -0.046 |
| 4710 | -0.053 |
| 4711 | -0.003 |
| 4712 | -0.079 |
| 4713 | 0.038 |
| 4714 | -0.003 |
| 4715 | -0.073 |
| 4716 | -0.023 |
| 4717 | -0.074 |
| 4718 | -0.102 |
| 4719 | 0.058 |
| 4720 | -0.022 |
| 4721 | -0.013 |
| 4722 | 0.030 |
| 4723 | 0.044 |
| 4724 | -0.063 |
| 4725 | 0.126 |
| 4726 | 0.007 |
| 4727 | 0.023 |
| 4728 | -0.063 |
| 4729 | 0.103 |
| 4730 | -0.034 |
| 4731 | -0.070 |
| 4732 | 0.150 |
| 4733 | -0.122 |
| 4734 | 0.044 |
| 4735 | 0.019 |
| 4736 | -0.017 |
| 4737 | -0.174 |
| 4738 | -0.021 |
| 4739 | 0.018 |
| 4740 | 0.080 |
| 4741 | -0.007 |
| 4742 | -0.121 |
| 4743 | 0.103 |
| 4744 | -0.071 |
| 4745 | 0.054 |
| 4746 | 0.043 |

| | |
|---|---|
| 4747 | -0.031 |
| 4748 | -0.134 |
| 4749 | 0.004 |
| 4750 | 0.054 |
| 4751 | 0.015 |
| 4752 | 0.011 |
| 4753 | 0.070 |
| 4754 | 0.030 |
| 4755 | 0.091 |
| 4756 | 0.051 |
| 4757 | -0.023 |
| 4758 | 0.035 |
| 4759 | 0.031 |
| 4760 | 0.011 |
| 4761 | 0.051 |
| 4762 | 0.002 |
| 4763 | -0.025 |
| 4764 | -0.180 |
| 4765 | 0.008 |
| 4766 | 0.070 |
| 4767 | 0.003 |
| 4768 | -0.004 |
| 4769 | -0.007 |
| 4770 | 0.030 |
| 4771 | 0.075 |
| 4772 | -0.019 |
| 4773 | 0.157 |
| 4774 | -0.119 |
| 4775 | -0.040 |
| 4776 | 0.052 |
| 4777 | -0.007 |
| 4778 | -0.123 |
| 4779 | -0.040 |
| 4780 | -0.058 |
| 4781 | -0.110 |
| 4782 | 0.046 |
| 4783 | -0.131 |
| 4784 | 0.084 |
| 4785 | 0.056 |
| 4786 | -0.109 |
| 4787 | 0.090 |
| 4788 | -0.067 |
| 4789 | 0.089 |
| 4790 | 0.080 |
| 4791 | -0.058 |
| 4792 | 0.003 |
| 4793 | 0.181 |
| 4794 | -0.009 |
| 4795 | -0.011 |
| 4796 | 0.002 |

| | |
|---|---|
| 4797 | 0.005 |
| 4798 | -0.044 |
| 4799 | 0.072 |
| 4800 | -0.019 |
| 4801 | 0.038 |
| 4802 | 0.064 |
| 4803 | -0.101 |
| 4804 | 0.057 |
| 4805 | 0.025 |
| 4806 | 0.011 |
| 4807 | 0.054 |
| 4808 | -0.001 |
| 4809 | -0.080 |
| 4810 | -0.032 |
| 4811 | 0.029 |
| 4812 | 0.137 |
| 4813 | 0.002 |
| 4814 | -0.106 |
| 4815 | -0.042 |
| 4816 | 0.107 |
| 4817 | 0.047 |
| 4818 | -0.019 |
| 4819 | -0.050 |
| 4820 | 0.094 |
| 4821 | 0.058 |
| 4822 | -0.037 |
| 4823 | 0.039 |
| 4824 | -0.055 |
| 4825 | -0.062 |
| 4826 | 0.044 |
| 4827 | 0.002 |
| 4828 | 0.069 |
| 4829 | -0.015 |
| 4830 | -0.008 |
| 4831 | 0.014 |
| 4832 | -0.105 |
| 4833 | 0.092 |
| 4834 | -0.024 |
| 4835 | 0.076 |
| 4836 | -0.032 |
| 4837 | -0.113 |
| 4838 | 0.053 |
| 4839 | -0.025 |
| 4840 | -0.034 |
| 4841 | 0.073 |
| 4842 | 0.038 |
| 4843 | -0.021 |
| 4844 | -0.027 |
| 4845 | 0.072 |
| 4846 | 0.028 |

| | |
|---|---|
| 4847 | 0.011 |
| 4848 | 0.038 |
| 4849 | 0.015 |
| 4850 | -0.003 |
| 4851 | 0.055 |
| 4852 | 0.021 |
| 4853 | -0.013 |
| 4854 | -0.125 |
| 4855 | 0.030 |
| 4856 | -0.125 |
| 4857 | -0.076 |
| 4858 | -0.114 |
| 4859 | 0.038 |
| 4860 | -0.162 |
| 4861 | -0.021 |
| 4862 | -0.019 |
| 4863 | 0.036 |
| 4864 | -0.031 |
| 4865 | -0.013 |
| 4866 | -0.073 |
| 4867 | 0.094 |
| 4868 | 0.020 |
| 4869 | 0.135 |
| 4870 | 0.042 |
| 4871 | 0.080 |
| 4872 | -0.015 |
| 4873 | 0.040 |
| 4874 | 0.195 |
| 4875 | 0.095 |
| 4876 | 0.116 |
| 4877 | 0.037 |
| 4878 | 0.128 |
| 4879 | 0.117 |
| 4880 | 0.023 |
| 4881 | 0.022 |
| 4882 | 0.061 |
| 4883 | 0.117 |
| 4884 | 0.072 |
| 4885 | 0.016 |
| 4886 | -0.109 |
| 4887 | 0.185 |
| 4888 | 0.135 |
| 4889 | -0.009 |
| 4890 | -0.011 |
| 4891 | 0.023 |
| 4892 | 0.042 |
| 4893 | -0.032 |
| 4894 | 0.045 |
| 4895 | 0.028 |
| 4896 | 0.134 |

| | |
|---|---|
| 4897 | -0.073 |
| 4898 | -0.014 |
| 4899 | -0.086 |
| 4900 | -0.014 |
| 4901 | -0.016 |
| 4902 | -0.047 |
| 4903 | -0.035 |
| 4904 | -0.013 |
| 4905 | -0.047 |
| 4906 | -0.042 |
| 4907 | -0.017 |
| 4908 | 0.041 |
| 4909 | -0.029 |
| 4910 | -0.021 |
| 4911 | 0.112 |
| 4912 | -0.003 |
| 4913 | -0.024 |
| 4914 | 0.010 |
| 4915 | -0.049 |
| 4916 | 0.049 |
| 4917 | 0.071 |
| 4918 | -0.016 |
| 4919 | 0.048 |
| 4920 | 0.009 |
| 4921 | 0.030 |
| 4922 | 0.031 |
| 4923 | 0.097 |
| 4924 | -0.083 |
| 4925 | -0.034 |
| 4926 | -0.047 |
| 4927 | 0.066 |
| 4928 | 0.065 |
| 4929 | -0.051 |
| 4930 | -0.059 |
| 4931 | 0.048 |
| 4932 | 0.125 |
| 4933 | 0.073 |
| 4934 | 0.016 |
| 4935 | -0.043 |
| 4936 | -0.046 |
| 4937 | 0.049 |
| 4938 | 0.094 |
| 4939 | 0.067 |
| 4940 | -0.113 |
| 4941 | -0.133 |
| 4942 | 0.005 |
| 4943 | -0.002 |
| 4944 | -0.059 |
| 4945 | -0.046 |
| 4946 | -0.074 |

| | |
|---|---|
| 4947 | 0.011 |
| 4948 | 0.002 |
| 4949 | -0.096 |
| 4950 | 0.013 |
| 4951 | -0.017 |
| 4952 | -0.079 |
| 4953 | -0.011 |
| 4954 | 0.003 |
| 4955 | 0.074 |
| 4956 | 0.015 |
| 4957 | 0.140 |
| 4958 | 0.025 |
| 4959 | 0.040 |
| 4960 | 0.007 |
| 4961 | -0.049 |
| 4962 | -0.001 |
| 4963 | -0.124 |
| 4964 | -0.070 |
| 4965 | -0.011 |
| 4966 | -0.050 |
| 4967 | -0.143 |
| 4968 | 0.050 |
| 4969 | -0.012 |
| 4970 | -0.113 |
| 4971 | 0.083 |
| 4972 | 0.083 |
| 4973 | -0.088 |
| 4974 | 0.016 |
| 4975 | 0.068 |
| 4976 | -0.047 |
| 4977 | 0.029 |
| 4978 | -0.026 |
| 4979 | 0.010 |
| 4980 | 0.004 |
| 4981 | -0.043 |
| 4982 | -0.058 |
| 4983 | 0.037 |
| 4984 | 0.021 |
| 4985 | 0.064 |
| 4986 | 0.001 |
| 4987 | -0.017 |
| 4988 | 0.023 |
| 4989 | 0.159 |
| 4990 | -0.095 |
| 4991 | -0.067 |
| 4992 | -0.040 |
| 4993 | -0.104 |
| 4994 | 0.048 |
| 4995 | 0.046 |
| 4996 | -0.076 |

| | |
|---|---|
| 4997 | 0.121 |
| 4998 | -0.145 |
| 4999 | -0.015 |
| 5000 | -0.041 |
| 5001 | -0.012 |
| 5002 | 0.076 |
| 5003 | 0.015 |
| 5004 | -0.087 |
| 5005 | 0.010 |
| 5006 | -0.012 |
| 5007 | 0.128 |
| 5008 | -0.057 |
| 5009 | 0.087 |
| 5010 | -0.040 |
| 5011 | 0.068 |
| 5012 | 0.030 |
| 5013 | 0.036 |
| 5014 | 0.095 |
| 5015 | 0.000 |
| 5016 | 0.011 |
| 5017 | -0.086 |
| 5018 | 0.065 |
| 5019 | -0.073 |
| 5020 | 0.011 |
| 5021 | 0.034 |
| 5022 | -0.078 |
| 5023 | -0.033 |
| 5024 | 0.019 |
| 5025 | 0.033 |
| 5026 | 0.026 |
| 5027 | -0.102 |
| 5028 | 0.019 |
| 5029 | 0.091 |
| 5030 | -0.037 |
| 5031 | -0.025 |
| 5032 | 0.074 |
| 5033 | 0.029 |
| 5034 | 0.043 |
| 5035 | 0.146 |
| 5036 | 0.066 |
| 5037 | -0.119 |
| 5038 | 0.094 |
| 5039 | -0.039 |
| 5040 | -0.027 |
| 5041 | -0.011 |
| 5042 | -0.078 |
| 5043 | -0.119 |
| 5044 | 0.095 |
| 5045 | 0.058 |
| 5046 | -0.008 |

| | |
|---|---|
| 5047 | 0.039 |
| 5048 | 0.020 |
| 5049 | 0.040 |
| 5050 | 0.014 |
| 5051 | 0.103 |
| 5052 | 0.083 |
| 5053 | 0.014 |
| 5054 | -0.042 |
| 5055 | -0.101 |
| 5056 | -0.047 |
| 5057 | 0.003 |
| 5058 | 0.043 |
| 5059 | -0.025 |
| 5060 | -0.026 |
| 5061 | -0.100 |
| 5062 | -0.005 |
| 5063 | -0.024 |
| 5064 | 0.030 |
| 5065 | -0.037 |
| 5066 | 0.099 |
| 5067 | 0.077 |
| 5068 | 0.031 |
| 5069 | 0.096 |
| 5070 | 0.033 |
| 5071 | 0.003 |
| 5072 | -0.023 |
| 5073 | -0.019 |
| 5074 | 0.013 |
| 5075 | 0.033 |
| 5076 | -0.055 |
| 5077 | 0.024 |
| 5078 | 0.067 |
| 5079 | 0.069 |
| 5080 | -0.002 |
| 5081 | -0.065 |
| 5082 | -0.001 |
| 5083 | -0.008 |
| 5084 | -0.097 |
| 5085 | -0.040 |
| 5086 | -0.140 |
| 5087 | 0.090 |
| 5088 | -0.051 |
| 5089 | 0.050 |
| 5090 | -0.078 |
| 5091 | -0.008 |
| 5092 | 0.080 |
| 5093 | -0.035 |
| 5094 | -0.057 |
| 5095 | -0.004 |
| 5096 | 0.053 |

| | |
|---|---|
| 5097 | -0.063 |
| 5098 | -0.045 |
| 5099 | 0.019 |
| 5100 | -0.060 |
| 5101 | 0.069 |
| 5102 | -0.056 |
| 5103 | 0.043 |
| 5104 | -0.074 |
| 5105 | 0.045 |
| 5106 | 0.015 |
| 5107 | -0.028 |
| 5108 | -0.001 |
| 5109 | -0.045 |
| 5110 | -0.074 |
| 5111 | 0.040 |
| 5112 | 0.011 |
| 5113 | 0.022 |
| 5114 | 0.055 |
| 5115 | 0.040 |
| 5116 | -0.119 |
| 5117 | 0.081 |
| 5118 | 0.040 |
| 5119 | -0.021 |
| 5120 | -0.059 |
| 5121 | 0.046 |
| 5122 | 0.085 |
| 5123 | -0.043 |
| 5124 | -0.101 |
| 5125 | -0.042 |
| 5126 | 0.026 |
| 5127 | 0.050 |
| 5128 | 0.055 |
| 5129 | -0.006 |
| 5130 | -0.077 |
| 5131 | 0.007 |
| 5132 | 0.125 |
| 5133 | 0.033 |
| 5134 | 0.038 |
| 5135 | 0.154 |
| 5136 | -0.045 |
| 5137 | 0.034 |
| 5138 | 0.055 |
| 5139 | -0.032 |
| 5140 | -0.035 |
| 5141 | 0.038 |
| 5142 | -0.016 |
| 5143 | 0.036 |
| 5144 | -0.039 |
| 5145 | -0.001 |
| 5146 | 0.023 |

| | |
|---|---|
| 5147 | 0.088 |
| 5148 | 0.063 |
| 5149 | 0.046 |
| 5150 | -0.055 |
| 5151 | 0.090 |
| 5152 | 0.039 |
| 5153 | -0.079 |
| 5154 | -0.014 |
| 5155 | 0.194 |
| 5156 | 0.150 |
| 5157 | -0.018 |
| 5158 | -0.068 |
| 5159 | -0.081 |
| 5160 | -0.054 |
| 5161 | 0.161 |
| 5162 | -0.076 |
| 5163 | -0.009 |
| 5164 | -0.025 |
| 5165 | -0.032 |
| 5166 | 0.000 |
| 5167 | -0.010 |
| 5168 | -0.010 |
| 5169 | -0.022 |
| 5170 | -0.052 |
| 5171 | -0.072 |
| 5172 | -0.003 |
| 5173 | 0.009 |
| 5174 | 0.108 |
| 5175 | -0.021 |
| 5176 | 0.101 |
| 5177 | -0.092 |
| 5178 | 0.039 |
| 5179 | -0.011 |
| 5180 | -0.045 |
| 5181 | 0.034 |
| 5182 | -0.011 |
| 5183 | -0.069 |
| 5184 | -0.009 |
| 5185 | 0.047 |
| 5186 | 0.016 |
| 5187 | 0.077 |
| 5188 | 0.053 |
| 5189 | 0.062 |
| 5190 | 0.127 |
| 5191 | 0.004 |
| 5192 | -0.111 |
| 5193 | -0.015 |
| 5194 | 0.122 |
| 5195 | -0.105 |
| 5196 | 0.013 |

| | |
|---|---|
| 5197 | -0.099 |
| 5198 | 0.088 |
| 5199 | -0.012 |
| 5200 | 0.124 |
| 5201 | 0.059 |
| 5202 | -0.006 |
| 5203 | 0.094 |
| 5204 | -0.027 |
| 5205 | 0.013 |
| 5206 | -0.125 |
| 5207 | 0.071 |
| 5208 | -0.056 |
| 5209 | -0.011 |
| 5210 | -0.049 |
| 5211 | 0.016 |
| 5212 | 0.023 |
| 5213 | -0.108 |
| 5214 | 0.099 |
| 5215 | -0.035 |
| 5216 | -0.034 |
| 5217 | -0.051 |
| 5218 | -0.092 |
| 5219 | 0.039 |
| 5220 | 0.022 |
| 5221 | 0.077 |
| 5222 | 0.102 |
| 5223 | -0.118 |
| 5224 | 0.036 |
| 5225 | -0.099 |
| 5226 | 0.000 |
| 5227 | 0.174 |
| 5228 | -0.005 |
| 5229 | -0.086 |
| 5230 | 0.051 |
| 5231 | 0.021 |
| 5232 | 0.079 |
| 5233 | 0.061 |
| 5234 | 0.060 |
| 5235 | -0.070 |
| 5236 | 0.097 |
| 5237 | 0.088 |
| 5238 | -0.022 |
| 5239 | -0.053 |
| 5240 | -0.090 |
| 5241 | -0.060 |
| 5242 | -0.017 |
| 5243 | -0.022 |
| 5244 | 0.043 |
| 5245 | 0.089 |
| 5246 | -0.020 |

| | |
|---|---|
| 5247 | 0.019 |
| 5248 | -0.136 |
| 5249 | -0.044 |
| 5250 | -0.044 |
| 5251 | 0.052 |
| 5252 | -0.107 |
| 5253 | -0.082 |
| 5254 | -0.073 |
| 5255 | -0.078 |
| 5256 | -0.060 |
| 5257 | -0.017 |
| 5258 | -0.008 |
| 5259 | -0.006 |
| 5260 | 0.042 |
| 5261 | 0.079 |
| 5262 | -0.102 |
| 5263 | -0.002 |
| 5264 | 0.055 |
| 5265 | 0.028 |
| 5266 | -0.063 |
| 5267 | -0.047 |
| 5268 | 0.137 |
| 5269 | 0.020 |
| 5270 | 0.172 |
| 5271 | -0.049 |
| 5272 | -0.052 |
| 5273 | -0.092 |
| 5274 | -0.011 |
| 5275 | -0.022 |
| 5276 | 0.067 |
| 5277 | -0.034 |
| 5278 | 0.001 |
| 5279 | 0.050 |
| 5280 | 0.031 |
| 5281 | -0.007 |
| 5282 | -0.018 |
| 5283 | -0.029 |
| 5284 | -0.076 |
| 5285 | 0.105 |
| 5286 | -0.079 |
| 5287 | -0.049 |
| 5288 | -0.052 |
| 5289 | 0.077 |
| 5290 | -0.152 |
| 5291 | 0.047 |
| 5292 | 0.026 |
| 5293 | -0.112 |
| 5294 | 0.075 |
| 5295 | 0.043 |
| 5296 | -0.045 |

| | |
|---|---|
| 5297 | 0.128 |
| 5298 | -0.021 |
| 5299 | 0.156 |
| 5300 | 0.021 |
| 5301 | 0.141 |
| 5302 | -0.059 |
| 5303 | -0.019 |
| 5304 | 0.046 |
| 5305 | 0.129 |
| 5306 | -0.024 |
| 5307 | -0.005 |
| 5308 | -0.130 |
| 5309 | 0.117 |
| 5310 | 0.015 |
| 5311 | 0.105 |
| 5312 | 0.043 |
| 5313 | 0.078 |
| 5314 | 0.026 |
| 5315 | -0.068 |
| 5316 | 0.008 |
| 5317 | -0.020 |
| 5318 | 0.035 |
| 5319 | 0.034 |
| 5320 | 0.070 |
| 5321 | -0.073 |
| 5322 | -0.142 |
| 5323 | -0.006 |
| 5324 | -0.038 |
| 5325 | -0.060 |
| 5326 | 0.070 |
| 5327 | 0.037 |
| 5328 | -0.027 |
| 5329 | 0.059 |
| 5330 | 0.037 |
| 5331 | 0.025 |
| 5332 | 0.098 |
| 5333 | -0.065 |
| 5334 | -0.025 |
| 5335 | -0.033 |
| 5336 | 0.012 |
| 5337 | -0.013 |
| 5338 | 0.033 |
| 5339 | 0.077 |
| 5340 | -0.059 |
| 5341 | 0.095 |
| 5342 | -0.003 |
| 5343 | 0.015 |
| 5344 | -0.021 |
| 5345 | 0.095 |
| 5346 | 0.022 |

| | |
|---|---|
| 5347 | 0.020 |
| 5348 | -0.018 |
| 5349 | 0.099 |
| 5350 | -0.031 |
| 5351 | 0.029 |
| 5352 | -0.007 |
| 5353 | -0.065 |
| 5354 | -0.029 |
| 5355 | 0.021 |
| 5356 | -0.026 |
| 5357 | 0.091 |
| 5358 | 0.025 |
| 5359 | -0.012 |
| 5360 | 0.000 |
| 5361 | -0.085 |
| 5362 | 0.014 |
| 5363 | -0.038 |
| 5364 | 0.131 |
| 5365 | -0.094 |
| 5366 | -0.003 |
| 5367 | -0.182 |
| 5368 | -0.039 |
| 5369 | 0.066 |
| 5370 | -0.011 |
| 5371 | 0.129 |
| 5372 | 0.208 |
| 5373 | -0.122 |
| 5374 | 0.038 |
| 5375 | -0.006 |
| 5376 | 0.033 |
| 5377 | 0.029 |
| 5378 | -0.047 |
| 5379 | -0.023 |
| 5380 | 0.155 |
| 5381 | 0.006 |
| 5382 | 0.053 |
| 5383 | -0.086 |
| 5384 | 0.041 |
| 5385 | 0.039 |
| 5386 | 0.094 |
| 5387 | 0.140 |
| 5388 | 0.040 |
| 5389 | -0.159 |
| 5390 | -0.050 |
| 5391 | 0.087 |
| 5392 | 0.002 |
| 5393 | 0.057 |
| 5394 | -0.040 |
| 5395 | 0.150 |
| 5396 | 0.054 |

| | |
|---|---|
| 5397 | 0.053 |
| 5398 | -0.104 |
| 5399 | 0.066 |
| 5400 | -0.112 |
| 5401 | 0.147 |
| 5402 | -0.035 |
| 5403 | -0.026 |
| 5404 | -0.023 |
| 5405 | -0.048 |
| 5406 | -0.023 |
| 5407 | -0.009 |
| 5408 | -0.084 |
| 5409 | -0.069 |
| 5410 | -0.088 |
| 5411 | -0.071 |
| 5412 | 0.054 |
| 5413 | 0.102 |
| 5414 | 0.057 |
| 5415 | 0.129 |
| 5416 | -0.094 |
| 5417 | 0.096 |
| 5418 | 0.021 |
| 5419 | 0.034 |
| 5420 | -0.018 |
| 5421 | 0.091 |
| 5422 | 0.093 |
| 5423 | 0.006 |
| 5424 | -0.070 |
| 5425 | 0.061 |
| 5426 | -0.126 |
| 5427 | 0.048 |
| 5428 | -0.069 |
| 5429 | 0.166 |
| 5430 | -0.041 |
| 5431 | 0.052 |
| 5432 | 0.195 |
| 5433 | -0.011 |
| 5434 | 0.022 |
| 5435 | 0.069 |
| 5436 | -0.085 |
| 5437 | -0.005 |
| 5438 | 0.003 |
| 5439 | -0.112 |
| 5440 | -0.005 |
| 5441 | -0.043 |
| 5442 | 0.054 |
| 5443 | -0.038 |
| 5444 | 0.049 |
| 5445 | -0.086 |
| 5446 | -0.241 |

| | |
|---|---|
| 5447 | -0.100 |
| 5448 | 0.046 |
| 5449 | 0.059 |
| 5450 | 0.055 |
| 5451 | -0.008 |
| 5452 | -0.008 |
| 5453 | -0.016 |
| 5454 | -0.047 |
| 5455 | -0.026 |
| 5456 | 0.026 |
| 5457 | -0.137 |
| 5458 | -0.157 |
| 5459 | -0.020 |
| 5460 | -0.051 |
| 5461 | -0.117 |
| 5462 | -0.032 |
| 5463 | -0.012 |
| 5464 | 0.032 |
| 5465 | 0.040 |
| 5466 | -0.032 |
| 5467 | -0.032 |
| 5468 | 0.019 |
| 5469 | -0.118 |
| 5470 | 0.038 |
| 5471 | -0.078 |
| 5472 | -0.029 |
| 5473 | -0.044 |
| 5474 | -0.002 |
| 5475 | 0.052 |
| 5476 | -0.070 |
| 5477 | 0.090 |
| 5478 | -0.092 |
| 5479 | 0.069 |
| 5480 | 0.036 |
| 5481 | 0.048 |
| 5482 | -0.022 |
| 5483 | -0.092 |
| 5484 | -0.116 |
| 5485 | -0.023 |
| 5486 | -0.099 |
| 5487 | 0.048 |
| 5488 | -0.057 |
| 5489 | -0.074 |
| 5490 | -0.040 |
| 5491 | -0.091 |
| 5492 | -0.081 |
| 5493 | -0.011 |
| 5494 | -0.014 |
| 5495 | -0.060 |
| 5496 | -0.075 |

| | |
|---|---|
| 5497 | 0.079 |
| 5498 | 0.042 |
| 5499 | -0.005 |
| 5500 | -0.090 |
| 5501 | 0.016 |
| 5502 | 0.062 |
| 5503 | -0.092 |
| 5504 | -0.077 |
| 5505 | -0.089 |
| 5506 | -0.065 |
| 5507 | 0.073 |
| 5508 | 0.068 |
| 5509 | -0.105 |
| 5510 | 0.056 |
| 5511 | 0.002 |
| 5512 | 0.082 |
| 5513 | -0.091 |
| 5514 | -0.061 |
| 5515 | -0.052 |
| 5516 | 0.041 |
| 5517 | -0.040 |
| 5518 | 0.043 |
| 5519 | -0.082 |
| 5520 | 0.144 |
| 5521 | 0.001 |
| 5522 | -0.054 |
| 5523 | 0.049 |
| 5524 | 0.074 |
| 5525 | -0.028 |
| 5526 | 0.017 |
| 5527 | -0.084 |
| 5528 | -0.093 |
| 5529 | 0.005 |
| 5530 | -0.043 |
| 5531 | 0.011 |
| 5532 | -0.010 |
| 5533 | 0.046 |
| 5534 | -0.005 |
| 5535 | -0.036 |
| 5536 | -0.088 |
| 5537 | -0.039 |
| 5538 | 0.071 |
| 5539 | -0.085 |
| 5540 | 0.111 |
| 5541 | -0.055 |
| 5542 | -0.033 |
| 5543 | 0.131 |
| 5544 | 0.036 |
| 5545 | 0.013 |
| 5546 | 0.013 |

| | |
|---|---|
| 5547 | -0.074 |
| 5548 | -0.003 |
| 5549 | 0.193 |
| 5550 | 0.051 |
| 5551 | 0.020 |
| 5552 | 0.007 |
| 5553 | -0.017 |
| 5554 | 0.010 |
| 5555 | 0.037 |
| 5556 | -0.042 |
| 5557 | 0.040 |
| 5558 | 0.050 |
| 5559 | -0.020 |
| 5560 | -0.019 |
| 5561 | -0.084 |
| 5562 | 0.017 |
| 5563 | -0.026 |
| 5564 | 0.120 |
| 5565 | -0.017 |
| 5566 | 0.012 |
| 5567 | -0.048 |
| 5568 | 0.014 |
| 5569 | -0.052 |
| 5570 | 0.008 |
| 5571 | 0.011 |
| 5572 | -0.061 |
| 5573 | 0.079 |
| 5574 | 0.021 |
| 5575 | 0.079 |
| 5576 | 0.106 |
| 5577 | -0.106 |
| 5578 | -0.119 |
| 5579 | 0.032 |
| 5580 | -0.001 |
| 5581 | -0.081 |
| 5582 | -0.052 |
| 5583 | -0.072 |
| 5584 | 0.055 |
| 5585 | 0.021 |
| 5586 | -0.005 |
| 5587 | 0.048 |
| 5588 | 0.020 |
| 5589 | -0.020 |
| 5590 | -0.025 |
| 5591 | -0.057 |
| 5592 | 0.076 |
| 5593 | 0.010 |
| 5594 | -0.084 |
| 5595 | -0.009 |
| 5596 | 0.166 |

| | |
|---|---|
| 5597 | 0.015 |
| 5598 | -0.142 |
| 5599 | 0.022 |
| 5600 | -0.020 |
| 5601 | -0.051 |
| 5602 | -0.023 |
| 5603 | -0.202 |
| 5604 | 0.073 |
| 5605 | 0.003 |
| 5606 | -0.033 |
| 5607 | 0.027 |
| 5608 | -0.046 |
| 5609 | 0.035 |
| 5610 | -0.027 |
| 5611 | -0.048 |
| 5612 | 0.048 |
| 5613 | 0.041 |
| 5614 | 0.188 |
| 5615 | -0.053 |
| 5616 | 0.052 |
| 5617 | -0.013 |
| 5618 | 0.091 |
| 5619 | 0.006 |
| 5620 | 0.008 |
| 5621 | 0.049 |
| 5622 | -0.052 |
| 5623 | 0.081 |
| 5624 | -0.046 |
| 5625 | 0.062 |
| 5626 | 0.005 |
| 5627 | -0.120 |
| 5628 | -0.078 |
| 5629 | 0.025 |
| 5630 | -0.008 |
| 5631 | 0.081 |
| 5632 | 0.005 |
| 5633 | 0.009 |
| 5634 | 0.053 |
| 5635 | 0.028 |
| 5636 | 0.098 |
| 5637 | -0.003 |
| 5638 | 0.107 |
| 5639 | -0.111 |
| 5640 | -0.120 |
| 5641 | 0.065 |
| 5642 | -0.101 |
| 5643 | -0.024 |
| 5644 | -0.086 |
| 5645 | 0.016 |
| 5646 | -0.069 |

| | |
|---|---|
| 5647 | -0.007 |
| 5648 | 0.023 |
| 5649 | 0.003 |
| 5650 | -0.016 |
| 5651 | 0.017 |
| 5652 | 0.008 |
| 5653 | 0.017 |
| 5654 | -0.045 |
| 5655 | 0.089 |
| 5656 | 0.052 |
| 5657 | 0.016 |
| 5658 | 0.030 |
| 5659 | 0.065 |
| 5660 | 0.091 |
| 5661 | -0.008 |
| 5662 | -0.037 |
| 5663 | -0.031 |
| 5664 | 0.015 |
| 5665 | -0.005 |
| 5666 | -0.058 |
| 5667 | -0.026 |
| 5668 | 0.149 |
| 5669 | 0.082 |
| 5670 | 0.162 |
| 5671 | -0.037 |
| 5672 | 0.105 |
| 5673 | 0.246 |
| 5674 | 0.056 |
| 5675 | 0.003 |
| 5676 | 0.042 |
| 5677 | 0.010 |
| 5678 | -0.012 |
| 5679 | -0.118 |
| 5680 | 0.067 |
| 5681 | -0.110 |
| 5682 | 0.077 |
| 5683 | 0.064 |
| 5684 | -0.055 |
| 5685 | 0.061 |
| 5686 | 0.028 |
| 5687 | -0.010 |
| 5688 | 0.003 |
| 5689 | 0.022 |
| 5690 | 0.000 |
| 5691 | 0.094 |
| 5692 | 0.002 |
| 5693 | -0.010 |
| 5694 | 0.039 |
| 5695 | 0.137 |
| 5696 | 0.013 |

| | |
|---|---|
| 5697 | 0.054 |
| 5698 | -0.002 |
| 5699 | 0.002 |
| 5700 | -0.121 |
| 5701 | -0.020 |
| 5702 | 0.021 |
| 5703 | 0.002 |
| 5704 | -0.024 |
| 5705 | -0.138 |
| 5706 | 0.012 |
| 5707 | -0.043 |
| 5708 | 0.093 |
| 5709 | -0.155 |
| 5710 | 0.000 |
| 5711 | 0.030 |
| 5712 | -0.066 |
| 5713 | -0.057 |
| 5714 | 0.016 |
| 5715 | -0.069 |
| 5716 | -0.012 |
| 5717 | 0.009 |
| 5718 | -0.066 |
| 5719 | 0.028 |
| 5720 | 0.008 |
| 5721 | -0.055 |
| 5722 | -0.055 |
| 5723 | 0.034 |
| 5724 | -0.009 |
| 5725 | -0.008 |
| 5726 | -0.112 |
| 5727 | 0.016 |
| 5728 | 0.013 |
| 5729 | -0.025 |
| 5730 | 0.005 |
| 5731 | 0.070 |
| 5732 | -0.019 |
| 5733 | 0.000 |
| 5734 | 0.080 |
| 5735 | 0.185 |
| 5736 | -0.030 |
| 5737 | -0.045 |
| 5738 | -0.021 |
| 5739 | -0.013 |
| 5740 | -0.121 |
| 5741 | -0.061 |
| 5742 | -0.049 |
| 5743 | 0.084 |
| 5744 | 0.042 |
| 5745 | -0.088 |
| 5746 | -0.002 |

| | |
|---|---|
| 5747 | -0.032 |
| 5748 | -0.121 |
| 5749 | 0.063 |
| 5750 | 0.117 |
| 5751 | 0.092 |
| 5752 | -0.041 |
| 5753 | -0.035 |
| 5754 | 0.012 |
| 5755 | 0.017 |
| 5756 | 0.080 |
| 5757 | 0.073 |
| 5758 | 0.035 |
| 5759 | 0.025 |
| 5760 | 0.065 |
| 5761 | -0.038 |
| 5762 | -0.050 |
| 5763 | 0.056 |
| 5764 | -0.137 |
| 5765 | 0.074 |
| 5766 | -0.015 |
| 5767 | -0.035 |
| 5768 | -0.030 |
| 5769 | 0.011 |
| 5770 | 0.089 |
| 5771 | -0.151 |
| 5772 | 0.055 |
| 5773 | -0.060 |
| 5774 | 0.021 |
| 5775 | 0.023 |
| 5776 | -0.040 |
| 5777 | -0.102 |
| 5778 | -0.034 |
| 5779 | 0.049 |
| 5780 | -0.033 |
| 5781 | 0.108 |
| 5782 | 0.089 |
| 5783 | 0.112 |
| 5784 | -0.019 |
| 5785 | 0.034 |
| 5786 | -0.052 |
| 5787 | -0.073 |
| 5788 | 0.037 |
| 5789 | 0.004 |
| 5790 | 0.005 |
| 5791 | 0.031 |
| 5792 | -0.084 |
| 5793 | 0.004 |
| 5794 | -0.072 |
| 5795 | 0.096 |
| 5796 | 0.028 |

| | |
|---|---|
| 5797 | 0.022 |
| 5798 | -0.073 |
| 5799 | 0.084 |
| 5800 | -0.052 |
| 5801 | -0.028 |
| 5802 | 0.026 |
| 5803 | 0.064 |
| 5804 | -0.033 |
| 5805 | -0.017 |
| 5806 | 0.048 |
| 5807 | -0.033 |
| 5808 | 0.062 |
| 5809 | -0.094 |
| 5810 | -0.070 |
| 5811 | 0.015 |
| 5812 | -0.008 |
| 5813 | -0.218 |
| 5814 | 0.038 |
| 5815 | -0.100 |
| 5816 | 0.082 |
| 5817 | -0.041 |
| 5818 | -0.075 |
| 5819 | 0.042 |
| 5820 | 0.021 |
| 5821 | -0.041 |
| 5822 | 0.063 |
| 5823 | 0.006 |
| 5824 | 0.037 |
| 5825 | -0.047 |
| 5826 | 0.056 |
| 5827 | 0.092 |
| 5828 | 0.019 |
| 5829 | 0.048 |
| 5830 | 0.058 |
| 5831 | 0.011 |
| 5832 | -0.222 |
| 5833 | -0.071 |
| 5834 | -0.054 |
| 5835 | -0.019 |
| 5836 | 0.041 |
| 5837 | 0.055 |
| 5838 | 0.087 |
| 5839 | -0.003 |
| 5840 | -0.034 |
| 5841 | -0.078 |
| 5842 | 0.037 |
| 5843 | 0.127 |
| 5844 | -0.088 |
| 5845 | -0.057 |
| 5846 | 0.054 |

| | |
|---|---|
| 5847 | 0.051 |
| 5848 | 0.055 |
| 5849 | -0.033 |
| 5850 | -0.029 |
| 5851 | -0.150 |
| 5852 | -0.035 |
| 5853 | 0.094 |
| 5854 | -0.013 |
| 5855 | 0.042 |
| 5856 | 0.121 |
| 5857 | -0.059 |
| 5858 | 0.020 |
| 5859 | -0.108 |
| 5860 | -0.061 |
| 5861 | -0.070 |
| 5862 | -0.029 |
| 5863 | 0.097 |
| 5864 | -0.058 |
| 5865 | -0.016 |
| 5866 | 0.049 |
| 5867 | -0.118 |
| 5868 | 0.024 |
| 5869 | 0.100 |
| 5870 | -0.036 |
| 5871 | -0.035 |
| 5872 | 0.053 |
| 5873 | -0.070 |
| 5874 | 0.070 |
| 5875 | 0.086 |
| 5876 | -0.012 |
| 5877 | 0.009 |
| 5878 | -0.050 |
| 5879 | 0.113 |
| 5880 | 0.083 |
| 5881 | -0.048 |
| 5882 | 0.045 |
| 5883 | -0.051 |
| 5884 | -0.063 |
| 5885 | -0.015 |
| 5886 | -0.222 |
| 5887 | -0.128 |
| 5888 | -0.008 |
| 5889 | -0.074 |
| 5890 | -0.180 |
| 5891 | -0.061 |
| 5892 | -0.013 |
| 5893 | 0.012 |
| 5894 | 0.031 |
| 5895 | 0.049 |
| 5896 | -0.189 |

| | |
|---|---|
| 5897 | -0.040 |
| 5898 | -0.039 |
| 5899 | 0.044 |
| 5900 | -0.040 |
| 5901 | -0.050 |
| 5902 | -0.113 |
| 5903 | -0.039 |
| 5904 | 0.021 |
| 5905 | -0.010 |
| 5906 | 0.062 |
| 5907 | -0.018 |
| 5908 | -0.065 |
| 5909 | -0.136 |
| 5910 | 0.071 |
| 5911 | -0.012 |
| 5912 | -0.041 |
| 5913 | -0.163 |
| 5914 | -0.056 |
| 5915 | -0.014 |
| 5916 | -0.041 |
| 5917 | 0.124 |
| 5918 | 0.024 |
| 5919 | -0.017 |
| 5920 | 0.182 |
| 5921 | -0.025 |
| 5922 | -0.123 |
| 5923 | 0.029 |
| 5924 | 0.056 |
| 5925 | -0.007 |
| 5926 | -0.067 |
| 5927 | 0.014 |
| 5928 | 0.087 |
| 5929 | -0.042 |
| 5930 | 0.059 |
| 5931 | -0.100 |
| 5932 | -0.140 |
| 5933 | 0.008 |
| 5934 | -0.043 |
| 5935 | 0.010 |
| 5936 | 0.001 |
| 5937 | -0.014 |
| 5938 | -0.078 |
| 5939 | 0.120 |
| 5940 | -0.165 |
| 5941 | 0.038 |
| 5942 | -0.108 |
| 5943 | -0.040 |
| 5944 | 0.046 |
| 5945 | -0.052 |
| 5946 | -0.040 |

| | |
|---|---|
| 5947 | -0.022 |
| 5948 | -0.086 |
| 5949 | 0.093 |
| 5950 | 0.028 |
| 5951 | 0.014 |
| 5952 | 0.015 |
| 5953 | -0.027 |
| 5954 | 0.009 |
| 5955 | -0.033 |
| 5956 | 0.038 |
| 5957 | 0.073 |
| 5958 | 0.035 |
| 5959 | 0.064 |
| 5960 | 0.124 |
| 5961 | -0.084 |
| 5962 | -0.025 |
| 5963 | 0.003 |
| 5964 | -0.041 |
| 5965 | 0.058 |
| 5966 | -0.046 |
| 5967 | -0.091 |
| 5968 | -0.052 |
| 5969 | 0.006 |
| 5970 | -0.068 |
| 5971 | -0.123 |
| 5972 | -0.057 |
| 5973 | -0.184 |
| 5974 | -0.006 |
| 5975 | -0.042 |
| 5976 | 0.017 |
| 5977 | -0.104 |
| 5978 | 0.008 |
| 5979 | -0.013 |
| 5980 | 0.071 |
| 5981 | -0.038 |
| 5982 | 0.099 |
| 5983 | -0.082 |
| 5984 | 0.039 |
| 5985 | 0.015 |
| 5986 | -0.120 |
| 5987 | -0.056 |
| 5988 | 0.037 |
| 5989 | 0.094 |
| 5990 | 0.044 |
| 5991 | 0.075 |
| 5992 | 0.014 |
| 5993 | 0.029 |
| 5994 | -0.070 |
| 5995 | -0.059 |
| 5996 | -0.056 |

| | |
|---|---|
| 5997 | -0.046 |
| 5998 | 0.008 |
| 5999 | 0.039 |
| 6000 | -0.069 |
| 6001 | 0.067 |
| 6002 | 0.063 |
| 6003 | 0.035 |
| 6004 | -0.080 |
| 6005 | -0.036 |
| 6006 | 0.083 |
| 6007 | 0.016 |
| 6008 | 0.047 |
| 6009 | 0.106 |
| 6010 | 0.043 |
| 6011 | -0.169 |
| 6012 | 0.031 |
| 6013 | -0.051 |
| 6014 | 0.041 |
| 6015 | -0.152 |
| 6016 | -0.043 |
| 6017 | 0.117 |
| 6018 | 0.086 |
| 6019 | -0.030 |
| 6020 | -0.065 |
| 6021 | 0.071 |
| 6022 | 0.002 |
| 6023 | -0.011 |
| 6024 | 0.038 |
| 6025 | -0.037 |
| 6026 | 0.100 |
| 6027 | -0.100 |
| 6028 | -0.130 |
| 6029 | -0.052 |
| 6030 | 0.028 |
| 6031 | -0.092 |
| 6032 | -0.208 |
| 6033 | -0.018 |
| 6034 | -0.071 |
| 6035 | -0.072 |
| 6036 | -0.007 |
| 6037 | -0.009 |
| 6038 | -0.057 |
| 6039 | 0.077 |
| 6040 | 0.104 |
| 6041 | -0.140 |
| 6042 | 0.025 |
| 6043 | 0.018 |
| 6044 | -0.052 |
| 6045 | 0.079 |
| 6046 | 0.098 |

| | |
|---|---|
| 6047 | -0.025 |
| 6048 | 0.059 |
| 6049 | 0.004 |
| 6050 | -0.029 |
| 6051 | -0.031 |
| 6052 | -0.109 |
| 6053 | 0.069 |
| 6054 | 0.032 |
| 6055 | 0.132 |
| 6056 | -0.041 |
| 6057 | 0.057 |
| 6058 | 0.004 |
| 6059 | -0.011 |
| 6060 | 0.042 |
| 6061 | 0.088 |
| 6062 | -0.059 |
| 6063 | 0.108 |
| 6064 | 0.151 |
| 6065 | -0.024 |
| 6066 | 0.028 |
| 6067 | -0.027 |
| 6068 | 0.041 |
| 6069 | 0.060 |
| 6070 | 0.031 |
| 6071 | 0.039 |
| 6072 | 0.048 |
| 6073 | 0.039 |
| 6074 | 0.081 |
| 6075 | -0.032 |
| 6076 | 0.022 |
| 6077 | 0.076 |
| 6078 | 0.124 |
| 6079 | -0.078 |
| 6080 | 0.097 |
| 6081 | -0.101 |
| 6082 | 0.066 |
| 6083 | 0.058 |
| 6084 | -0.152 |
| 6085 | 0.000 |
| 6086 | 0.001 |
| 6087 | 0.046 |
| 6088 | -0.003 |
| 6089 | 0.033 |
| 6090 | -0.033 |
| 6091 | -0.074 |
| 6092 | -0.011 |
| 6093 | -0.007 |
| 6094 | 0.090 |
| 6095 | 0.059 |
| 6096 | -0.088 |

| | |
|---|---|
| 6097 | -0.008 |
| 6098 | 0.000 |
| 6099 | 0.019 |
| 6100 | 0.109 |
| 6101 | 0.111 |
| 6102 | -0.097 |
| 6103 | 0.072 |
| 6104 | -0.045 |
| 6105 | 0.047 |
| 6106 | 0.131 |
| 6107 | -0.033 |
| 6108 | -0.024 |
| 6109 | -0.001 |
| 6110 | -0.031 |
| 6111 | 0.045 |
| 6112 | -0.079 |
| 6113 | 0.067 |
| 6114 | 0.014 |
| 6115 | -0.027 |
| 6116 | 0.074 |
| 6117 | -0.032 |
| 6118 | -0.134 |
| 6119 | 0.062 |
| 6120 | 0.042 |
| 6121 | -0.028 |
| 6122 | -0.001 |
| 6123 | 0.036 |
| 6124 | 0.017 |
| 6125 | -0.066 |
| 6126 | 0.056 |
| 6127 | -0.023 |
| 6128 | 0.030 |
| 6129 | 0.020 |
| 6130 | 0.076 |
| 6131 | 0.026 |
| 6132 | 0.131 |
| 6133 | -0.073 |
| 6134 | 0.003 |
| 6135 | 0.129 |
| 6136 | -0.024 |
| 6137 | 0.067 |
| 6138 | -0.036 |
| 6139 | -0.021 |
| 6140 | 0.040 |
| 6141 | -0.017 |
| 6142 | -0.064 |
| 6143 | 0.034 |
| 6144 | 0.050 |
| 6145 | -0.025 |
| 6146 | 0.016 |

| | |
|---|---|
| 6147 | -0.003 |
| 6148 | -0.053 |
| 6149 | 0.024 |
| 6150 | 0.004 |
| 6151 | -0.041 |
| 6152 | 0.139 |
| 6153 | -0.032 |
| 6154 | -0.009 |
| 6155 | -0.085 |
| 6156 | -0.044 |
| 6157 | -0.050 |
| 6158 | -0.036 |
| 6159 | 0.051 |
| 6160 | 0.006 |
| 6161 | -0.171 |
| 6162 | 0.030 |
| 6163 | -0.002 |
| 6164 | -0.024 |
| 6165 | 0.038 |
| 6166 | -0.106 |
| 6167 | 0.065 |
| 6168 | 0.006 |
| 6169 | 0.143 |
| 6170 | 0.023 |
| 6171 | 0.099 |
| 6172 | -0.052 |
| 6173 | 0.014 |
| 6174 | -0.130 |
| 6175 | -0.003 |
| 6176 | -0.070 |
| 6177 | -0.018 |
| 6178 | 0.062 |
| 6179 | -0.079 |
| 6180 | 0.161 |
| 6181 | 0.038 |
| 6182 | 0.058 |
| 6183 | 0.177 |
| 6184 | -0.004 |
| 6185 | -0.121 |
| 6186 | 0.099 |
| 6187 | 0.111 |
| 6188 | 0.087 |
| 6189 | -0.097 |
| 6190 | 0.125 |
| 6191 | -0.026 |
| 6192 | 0.169 |
| 6193 | 0.062 |
| 6194 | 0.008 |
| 6195 | 0.071 |
| 6196 | -0.021 |

| | |
|---|---|
| 6197 | 0.051 |
| 6198 | -0.075 |
| 6199 | 0.028 |
| 6200 | 0.011 |
| 6201 | -0.005 |
| 6202 | 0.008 |
| 6203 | 0.029 |
| 6204 | 0.049 |
| 6205 | 0.041 |
| 6206 | -0.068 |
| 6207 | 0.067 |
| 6208 | -0.163 |
| 6209 | 0.039 |
| 6210 | 0.108 |
| 6211 | -0.046 |
| 6212 | 0.129 |
| 6213 | 0.040 |
| 6214 | -0.096 |
| 6215 | -0.040 |
| 6216 | 0.059 |
| 6217 | 0.000 |
| 6218 | -0.025 |
| 6219 | 0.006 |
| 6220 | -0.036 |
| 6221 | 0.036 |
| 6222 | -0.064 |
| 6223 | 0.054 |
| 6224 | -0.148 |
| 6225 | 0.044 |
| 6226 | 0.024 |
| 6227 | 0.040 |
| 6228 | 0.124 |
| 6229 | -0.081 |
| 6230 | 0.078 |
| 6231 | -0.129 |
| 6232 | 0.080 |
| 6233 | 0.007 |
| 6234 | 0.036 |
| 6235 | 0.052 |
| 6236 | -0.076 |
| 6237 | -0.056 |
| 6238 | -0.031 |
| 6239 | -0.073 |
| 6240 | 0.005 |
| 6241 | -0.153 |
| 6242 | 0.058 |
| 6243 | 0.057 |
| 6244 | 0.037 |
| 6245 | 0.039 |
| 6246 | 0.041 |

| | |
|---|---|
| 6247 | 0.094 |
| 6248 | 0.148 |
| 6249 | -0.017 |
| 6250 | 0.051 |
| 6251 | -0.057 |
| 6252 | -0.115 |
| 6253 | -0.158 |
| 6254 | -0.001 |
| 6255 | 0.071 |
| 6256 | 0.064 |
| 6257 | 0.099 |
| 6258 | 0.053 |
| 6259 | -0.024 |
| 6260 | 0.011 |
| 6261 | 0.071 |
| 6262 | -0.011 |
| 6263 | 0.009 |
| 6264 | 0.029 |
| 6265 | 0.014 |
| 6266 | 0.124 |
| 6267 | 0.100 |
| 6268 | 0.076 |
| 6269 | -0.082 |
| 6270 | 0.014 |
| 6271 | 0.106 |
| 6272 | 0.101 |
| 6273 | 0.027 |
| 6274 | 0.073 |
| 6275 | -0.005 |
| 6276 | 0.011 |
| 6277 | 0.001 |
| 6278 | -0.115 |
| 6279 | 0.065 |
| 6280 | -0.034 |
| 6281 | -0.035 |
| 6282 | -0.001 |
| 6283 | 0.029 |
| 6284 | -0.056 |
| 6285 | 0.025 |
| 6286 | -0.014 |
| 6287 | -0.012 |
| 6288 | -0.002 |
| 6289 | -0.032 |
| 6290 | 0.100 |
| 6291 | 0.119 |
| 6292 | 0.056 |
| 6293 | 0.043 |
| 6294 | -0.039 |
| 6295 | -0.047 |
| 6296 | 0.027 |

| | |
|---|---|
| 6297 | 0.069 |
| 6298 | 0.004 |
| 6299 | 0.074 |
| 6300 | 0.005 |
| 6301 | 0.061 |
| 6302 | -0.054 |
| 6303 | 0.011 |
| 6304 | 0.035 |
| 6305 | 0.131 |
| 6306 | -0.011 |
| 6307 | -0.058 |
| 6308 | -0.153 |
| 6309 | -0.033 |
| 6310 | 0.017 |
| 6311 | 0.013 |
| 6312 | -0.081 |
| 6313 | 0.126 |
| 6314 | 0.131 |
| 6315 | -0.050 |
| 6316 | 0.086 |
| 6317 | 0.038 |
| 6318 | -0.119 |
| 6319 | -0.004 |
| 6320 | -0.123 |
| 6321 | 0.026 |
| 6322 | 0.036 |
| 6323 | 0.067 |
| 6324 | 0.039 |
| 6325 | 0.097 |
| 6326 | -0.018 |
| 6327 | -0.006 |
| 6328 | -0.021 |
| 6329 | -0.010 |
| 6330 | 0.015 |
| 6331 | 0.007 |
| 6332 | 0.032 |
| 6333 | 0.012 |
| 6334 | 0.007 |
| 6335 | -0.081 |
| 6336 | -0.016 |
| 6337 | -0.021 |
| 6338 | 0.108 |
| 6339 | 0.152 |
| 6340 | 0.007 |
| 6341 | -0.023 |
| 6342 | -0.012 |
| 6343 | -0.016 |
| 6344 | -0.097 |
| 6345 | -0.013 |
| 6346 | 0.177 |

| | |
|---|---|
| 6347 | -0.057 |
| 6348 | 0.024 |
| 6349 | 0.042 |
| 6350 | 0.017 |
| 6351 | 0.143 |
| 6352 | 0.018 |
| 6353 | -0.058 |
| 6354 | 0.049 |
| 6355 | -0.071 |
| 6356 | -0.084 |
| 6357 | -0.011 |
| 6358 | 0.050 |
| 6359 | -0.065 |
| 6360 | -0.042 |
| 6361 | 0.027 |
| 6362 | -0.131 |
| 6363 | 0.108 |
| 6364 | -0.055 |
| 6365 | -0.075 |
| 6366 | -0.054 |
| 6367 | -0.063 |
| 6368 | -0.061 |
| 6369 | -0.033 |
| 6370 | -0.022 |
| 6371 | 0.095 |
| 6372 | -0.044 |
| 6373 | -0.001 |
| 6374 | 0.037 |
| 6375 | -0.019 |
| 6376 | -0.013 |
| 6377 | 0.079 |
| 6378 | -0.077 |
| 6379 | 0.010 |
| 6380 | -0.022 |
| 6381 | 0.001 |
| 6382 | 0.034 |
| 6383 | -0.086 |
| 6384 | 0.008 |
| 6385 | -0.116 |
| 6386 | 0.013 |
| 6387 | -0.024 |
| 6388 | -0.014 |
| 6389 | -0.045 |
| 6390 | -0.021 |
| 6391 | 0.041 |
| 6392 | -0.115 |
| 6393 | 0.185 |
| 6394 | 0.089 |
| 6395 | -0.075 |
| 6396 | 0.051 |

| | |
|---|---|
| 6397 | -0.050 |
| 6398 | 0.089 |
| 6399 | -0.045 |
| 6400 | 0.049 |
| 6401 | -0.088 |
| 6402 | 0.001 |
| 6403 | 0.028 |
| 6404 | -0.095 |
| 6405 | 0.073 |
| 6406 | -0.070 |
| 6407 | 0.032 |
| 6408 | 0.042 |
| 6409 | -0.014 |
| 6410 | 0.009 |
| 6411 | 0.004 |
| 6412 | 0.052 |
| 6413 | -0.046 |
| 6414 | -0.072 |
| 6415 | -0.019 |
| 6416 | 0.016 |
| 6417 | -0.005 |
| 6418 | -0.009 |
| 6419 | 0.064 |
| 6420 | -0.032 |
| 6421 | 0.083 |
| 6422 | -0.093 |
| 6423 | -0.027 |
| 6424 | 0.072 |
| 6425 | 0.012 |
| 6426 | 0.014 |
| 6427 | -0.084 |
| 6428 | 0.008 |
| 6429 | -0.059 |
| 6430 | 0.040 |
| 6431 | -0.007 |
| 6432 | 0.159 |
| 6433 | 0.037 |
| 6434 | -0.042 |
| 6435 | -0.014 |
| 6436 | -0.003 |
| 6437 | 0.003 |
| 6438 | 0.046 |
| 6439 | -0.007 |
| 6440 | 0.053 |
| 6441 | 0.060 |
| 6442 | -0.017 |
| 6443 | -0.063 |
| 6444 | -0.072 |
| 6445 | 0.030 |
| 6446 | -0.121 |

| | |
|---|---|
| 6447 | 0.025 |
| 6448 | -0.026 |
| 6449 | -0.121 |
| 6450 | 0.000 |
| 6451 | 0.111 |
| 6452 | -0.014 |
| 6453 | 0.035 |
| 6454 | -0.031 |
| 6455 | 0.067 |
| 6456 | -0.086 |
| 6457 | -0.078 |
| 6458 | -0.024 |
| 6459 | 0.133 |
| 6460 | -0.004 |
| 6461 | -0.004 |
| 6462 | -0.113 |
| 6463 | 0.031 |
| 6464 | -0.083 |
| 6465 | 0.042 |
| 6466 | 0.009 |
| 6467 | 0.094 |
| 6468 | -0.009 |
| 6469 | 0.136 |
| 6470 | 0.083 |
| 6471 | 0.134 |
| 6472 | -0.016 |
| 6473 | 0.083 |
| 6474 | 0.043 |
| 6475 | 0.064 |
| 6476 | -0.025 |
| 6477 | 0.113 |
| 6478 | 0.069 |
| 6479 | 0.005 |
| 6480 | 0.054 |
| 6481 | 0.026 |
| 6482 | -0.049 |
| 6483 | 0.023 |
| 6484 | -0.084 |
| 6485 | -0.016 |
| 6486 | -0.020 |
| 6487 | 0.045 |
| 6488 | -0.066 |
| 6489 | 0.097 |
| 6490 | -0.058 |
| 6491 | 0.045 |
| 6492 | -0.115 |
| 6493 | -0.034 |
| 6494 | 0.106 |
| 6495 | -0.006 |
| 6496 | 0.027 |

| | |
|---|---|
| 6497 | 0.081 |
| 6498 | 0.036 |
| 6499 | 0.039 |
| 6500 | -0.076 |
| 6501 | 0.026 |
| 6502 | -0.116 |
| 6503 | 0.031 |
| 6504 | 0.073 |
| 6505 | -0.027 |
| 6506 | 0.063 |
| 6507 | 0.030 |
| 6508 | -0.097 |
| 6509 | -0.092 |
| 6510 | -0.073 |
| 6511 | 0.024 |
| 6512 | 0.063 |
| 6513 | -0.072 |
| 6514 | 0.006 |
| 6515 | -0.008 |
| 6516 | 0.111 |
| 6517 | -0.061 |
| 6518 | 0.105 |
| 6519 | -0.051 |
| 6520 | -0.017 |
| 6521 | 0.059 |
| 6522 | -0.017 |
| 6523 | -0.088 |
| 6524 | 0.004 |
| 6525 | -0.008 |
| 6526 | 0.037 |
| 6527 | -0.050 |
| 6528 | 0.014 |
| 6529 | -0.017 |
| 6530 | -0.008 |
| 6531 | -0.102 |
| 6532 | 0.064 |
| 6533 | 0.045 |
| 6534 | -0.058 |
| 6535 | 0.053 |
| 6536 | -0.083 |
| 6537 | 0.007 |
| 6538 | -0.035 |
| 6539 | 0.128 |
| 6540 | -0.013 |
| 6541 | -0.015 |
| 6542 | 0.100 |
| 6543 | 0.129 |
| 6544 | -0.001 |
| 6545 | 0.053 |
| 6546 | 0.084 |

| | |
|---|---|
| 6547 | -0.007 |
| 6548 | 0.081 |
| 6549 | -0.087 |
| 6550 | 0.157 |
| 6551 | -0.069 |
| 6552 | -0.004 |
| 6553 | -0.114 |
| 6554 | 0.018 |
| 6555 | -0.078 |
| 6556 | -0.022 |
| 6557 | -0.052 |
| 6558 | -0.152 |
| 6559 | -0.038 |
| 6560 | -0.025 |
| 6561 | 0.017 |
| 6562 | -0.058 |
| 6563 | -0.117 |
| 6564 | -0.003 |
| 6565 | -0.027 |
| 6566 | -0.057 |
| 6567 | -0.004 |
| 6568 | 0.104 |
| 6569 | -0.029 |
| 6570 | -0.073 |
| 6571 | -0.019 |
| 6572 | 0.116 |
| 6573 | -0.035 |
| 6574 | 0.020 |
| 6575 | -0.025 |
| 6576 | -0.058 |
| 6577 | 0.102 |
| 6578 | -0.006 |
| 6579 | -0.091 |
| 6580 | 0.067 |
| 6581 | 0.004 |
| 6582 | -0.085 |
| 6583 | -0.022 |
| 6584 | 0.077 |
| 6585 | -0.063 |
| 6586 | 0.134 |
| 6587 | -0.045 |
| 6588 | -0.033 |
| 6589 | 0.004 |
| 6590 | -0.213 |
| 6591 | -0.064 |
| 6592 | -0.001 |
| 6593 | 0.085 |
| 6594 | 0.090 |
| 6595 | 0.003 |
| 6596 | 0.010 |

| | |
|---|---|
| 6597 | 0.032 |
| 6598 | -0.032 |
| 6599 | 0.127 |
| 6600 | -0.044 |
| 6601 | 0.020 |
| 6602 | -0.073 |
| 6603 | 0.013 |
| 6604 | 0.071 |
| 6605 | 0.064 |
| 6606 | -0.063 |
| 6607 | -0.006 |
| 6608 | 0.075 |
| 6609 | 0.016 |
| 6610 | -0.014 |
| 6611 | 0.004 |
| 6612 | 0.093 |
| 6613 | -0.095 |
| 6614 | -0.037 |
| 6615 | -0.006 |
| 6616 | -0.106 |
| 6617 | -0.053 |
| 6618 | -0.014 |
| 6619 | 0.031 |
| 6620 | -0.036 |
| 6621 | 0.001 |
| 6622 | 0.062 |
| 6623 | -0.034 |
| 6624 | -0.080 |
| 6625 | -0.036 |
| 6626 | 0.002 |
| 6627 | -0.033 |
| 6628 | 0.040 |
| 6629 | 0.080 |
| 6630 | 0.023 |
| 6631 | 0.045 |
| 6632 | -0.023 |
| 6633 | 0.099 |
| 6634 | 0.016 |
| 6635 | 0.059 |
| 6636 | -0.060 |
| 6637 | 0.094 |
| 6638 | -0.017 |
| 6639 | 0.027 |
| 6640 | 0.122 |
| 6641 | 0.086 |
| 6642 | 0.000 |
| 6643 | -0.063 |
| 6644 | -0.093 |
| 6645 | 0.069 |
| 6646 | -0.085 |

| | |
|---|---|
| 6647 | -0.027 |
| 6648 | -0.087 |
| 6649 | -0.055 |
| 6650 | 0.057 |
| 6651 | -0.067 |
| 6652 | 0.039 |
| 6653 | -0.014 |
| 6654 | 0.027 |
| 6655 | -0.047 |
| 6656 | -0.105 |
| 6657 | -0.107 |
| 6658 | -0.023 |
| 6659 | 0.124 |
| 6660 | 0.032 |
| 6661 | -0.029 |
| 6662 | -0.008 |
| 6663 | -0.034 |
| 6664 | -0.005 |
| 6665 | 0.032 |
| 6666 | 0.075 |
| 6667 | -0.095 |
| 6668 | 0.002 |
| 6669 | -0.056 |
| 6670 | -0.011 |
| 6671 | -0.075 |
| 6672 | 0.021 |
| 6673 | -0.001 |
| 6674 | -0.089 |
| 6675 | -0.108 |
| 6676 | -0.100 |
| 6677 | 0.001 |
| 6678 | -0.055 |
| 6679 | 0.059 |
| 6680 | 0.032 |
| 6681 | -0.075 |
| 6682 | 0.018 |
| 6683 | -0.066 |
| 6684 | 0.026 |
| 6685 | -0.032 |
| 6686 | 0.089 |
| 6687 | -0.027 |
| 6688 | 0.078 |
| 6689 | 0.026 |
| 6690 | -0.014 |
| 6691 | -0.043 |
| 6692 | 0.033 |
| 6693 | -0.014 |
| 6694 | -0.092 |
| 6695 | 0.066 |
| 6696 | -0.055 |

| | |
|---|---|
| 6697 | 0.081 |
| 6698 | -0.131 |
| 6699 | -0.061 |
| 6700 | -0.016 |
| 6701 | 0.025 |
| 6702 | -0.035 |
| 6703 | -0.077 |
| 6704 | 0.114 |
| 6705 | 0.159 |
| 6706 | 0.034 |
| 6707 | 0.001 |
| 6708 | 0.017 |
| 6709 | 0.104 |
| 6710 | -0.052 |
| 6711 | 0.051 |
| 6712 | -0.111 |
| 6713 | -0.078 |
| 6714 | -0.049 |
| 6715 | 0.003 |
| 6716 | 0.069 |
| 6717 | 0.011 |
| 6718 | -0.051 |
| 6719 | -0.007 |
| 6720 | 0.005 |
| 6721 | 0.105 |
| 6722 | -0.070 |
| 6723 | -0.037 |
| 6724 | 0.022 |
| 6725 | 0.031 |
| 6726 | 0.066 |
| 6727 | 0.093 |
| 6728 | 0.012 |
| 6729 | -0.018 |
| 6730 | 0.116 |
| 6731 | -0.094 |
| 6732 | 0.016 |
| 6733 | 0.057 |
| 6734 | 0.020 |
| 6735 | -0.074 |
| 6736 | 0.062 |
| 6737 | -0.068 |
| 6738 | -0.110 |
| 6739 | -0.143 |
| 6740 | -0.112 |
| 6741 | -0.019 |
| 6742 | -0.036 |
| 6743 | -0.036 |
| 6744 | -0.041 |
| 6745 | 0.057 |
| 6746 | 0.036 |

| | |
|---|---|
| 6747 | 0.035 |
| 6748 | -0.135 |
| 6749 | 0.059 |
| 6750 | 0.035 |
| 6751 | -0.021 |
| 6752 | 0.104 |
| 6753 | -0.031 |
| 6754 | -0.057 |
| 6755 | 0.072 |
| 6756 | -0.119 |
| 6757 | -0.130 |
| 6758 | 0.018 |
| 6759 | 0.138 |
| 6760 | 0.151 |
| 6761 | 0.024 |
| 6762 | 0.096 |
| 6763 | 0.046 |
| 6764 | 0.053 |
| 6765 | -0.006 |
| 6766 | 0.058 |
| 6767 | -0.054 |
| 6768 | -0.051 |
| 6769 | 0.037 |
| 6770 | -0.018 |
| 6771 | -0.098 |
| 6772 | 0.016 |
| 6773 | 0.036 |
| 6774 | 0.018 |
| 6775 | -0.096 |
| 6776 | 0.066 |
| 6777 | -0.076 |
| 6778 | 0.055 |
| 6779 | 0.042 |
| 6780 | -0.088 |
| 6781 | -0.044 |
| 6782 | 0.044 |
| 6783 | 0.051 |
| 6784 | -0.134 |
| 6785 | -0.006 |
| 6786 | 0.116 |
| 6787 | -0.049 |
| 6788 | -0.076 |
| 6789 | -0.100 |
| 6790 | 0.087 |
| 6791 | -0.062 |
| 6792 | 0.018 |
| 6793 | 0.014 |
| 6794 | -0.017 |
| 6795 | -0.109 |
| 6796 | -0.087 |

| | |
|---|---|
| 6797 | -0.016 |
| 6798 | -0.005 |
| 6799 | -0.016 |
| 6800 | -0.039 |
| 6801 | 0.015 |
| 6802 | 0.018 |
| 6803 | -0.267 |
| 6804 | 0.069 |
| 6805 | 0.058 |
| 6806 | -0.052 |
| 6807 | -0.004 |
| 6808 | 0.082 |
| 6809 | 0.004 |
| 6810 | 0.077 |
| 6811 | 0.085 |
| 6812 | 0.178 |
| 6813 | -0.061 |
| 6814 | 0.110 |
| 6815 | -0.108 |
| 6816 | -0.122 |
| 6817 | -0.073 |
| 6818 | 0.086 |
| 6819 | 0.136 |
| 6820 | -0.018 |
| 6821 | 0.002 |
| 6822 | 0.087 |
| 6823 | 0.050 |
| 6824 | 0.015 |
| 6825 | 0.072 |
| 6826 | -0.025 |
| 6827 | -0.045 |
| 6828 | -0.129 |
| 6829 | -0.113 |
| 6830 | -0.131 |
| 6831 | -0.106 |
| 6832 | 0.064 |
| 6833 | 0.065 |
| 6834 | 0.026 |
| 6835 | 0.013 |
| 6836 | -0.041 |
| 6837 | -0.144 |
| 6838 | 0.093 |
| 6839 | 0.030 |
| 6840 | 0.090 |
| 6841 | 0.088 |
| 6842 | 0.128 |
| 6843 | 0.026 |
| 6844 | 0.129 |
| 6845 | 0.120 |
| 6846 | 0.022 |

| | |
|---|---|
| 6847 | 0.029 |
| 6848 | -0.060 |
| 6849 | 0.117 |
| 6850 | 0.008 |
| 6851 | -0.068 |
| 6852 | -0.011 |
| 6853 | 0.084 |
| 6854 | -0.015 |
| 6855 | 0.028 |
| 6856 | 0.046 |
| 6857 | -0.052 |
| 6858 | -0.016 |
| 6859 | 0.043 |
| 6860 | -0.012 |
| 6861 | -0.029 |
| 6862 | -0.077 |
| 6863 | 0.032 |
| 6864 | 0.031 |
| 6865 | 0.027 |
| 6866 | 0.027 |
| 6867 | 0.123 |
| 6868 | -0.041 |
| 6869 | 0.046 |
| 6870 | -0.064 |
| 6871 | 0.106 |
| 6872 | -0.043 |
| 6873 | -0.073 |
| 6874 | 0.053 |
| 6875 | -0.084 |
| 6876 | -0.072 |
| 6877 | 0.031 |
| 6878 | 0.083 |
| 6879 | -0.015 |
| 6880 | 0.038 |
| 6881 | -0.015 |
| 6882 | 0.053 |
| 6883 | 0.015 |
| 6884 | -0.009 |
| 6885 | -0.120 |
| 6886 | 0.011 |
| 6887 | -0.084 |
| 6888 | 0.090 |
| 6889 | 0.083 |
| 6890 | 0.097 |
| 6891 | -0.032 |
| 6892 | -0.095 |
| 6893 | 0.022 |
| 6894 | -0.016 |
| 6895 | 0.030 |
| 6896 | -0.041 |

| | |
|---|---|
| 6897 | -0.054 |
| 6898 | 0.033 |
| 6899 | 0.070 |
| 6900 | -0.042 |
| 6901 | -0.046 |
| 6902 | 0.024 |
| 6903 | -0.039 |
| 6904 | 0.038 |
| 6905 | 0.025 |
| 6906 | -0.006 |
| 6907 | 0.150 |
| 6908 | -0.012 |
| 6909 | 0.014 |
| 6910 | -0.027 |
| 6911 | 0.100 |
| 6912 | -0.014 |
| 6913 | 0.004 |
| 6914 | -0.084 |
| 6915 | -0.102 |
| 6916 | 0.079 |
| 6917 | -0.025 |
| 6918 | -0.110 |
| 6919 | 0.013 |
| 6920 | -0.004 |
| 6921 | -0.039 |
| 6922 | 0.016 |
| 6923 | -0.054 |
| 6924 | -0.016 |
| 6925 | 0.038 |
| 6926 | -0.008 |
| 6927 | 0.045 |
| 6928 | 0.075 |
| 6929 | -0.053 |
| 6930 | 0.019 |
| 6931 | 0.039 |
| 6932 | 0.063 |
| 6933 | 0.034 |
| 6934 | -0.046 |
| 6935 | 0.038 |
| 6936 | -0.022 |
| 6937 | -0.135 |
| 6938 | -0.073 |
| 6939 | -0.094 |
| 6940 | 0.021 |
| 6941 | 0.027 |
| 6942 | 0.065 |
| 6943 | -0.025 |
| 6944 | -0.017 |
| 6945 | -0.031 |
| 6946 | 0.057 |

| | |
|---|---|
| 6947 | -0.034 |
| 6948 | -0.077 |
| 6949 | -0.087 |
| 6950 | -0.002 |
| 6951 | 0.027 |
| 6952 | 0.055 |
| 6953 | -0.025 |
| 6954 | 0.176 |
| 6955 | 0.057 |
| 6956 | -0.052 |
| 6957 | -0.079 |
| 6958 | 0.019 |
| 6959 | 0.010 |
| 6960 | -0.157 |
| 6961 | -0.069 |
| 6962 | -0.082 |
| 6963 | 0.115 |
| 6964 | -0.114 |
| 6965 | 0.090 |
| 6966 | 0.102 |
| 6967 | 0.025 |
| 6968 | 0.020 |
| 6969 | -0.076 |
| 6970 | -0.025 |
| 6971 | -0.017 |
| 6972 | -0.103 |
| 6973 | -0.130 |
| 6974 | -0.003 |
| 6975 | -0.178 |
| 6976 | 0.153 |
| 6977 | 0.007 |
| 6978 | -0.135 |
| 6979 | -0.023 |
| 6980 | 0.062 |
| 6981 | -0.006 |
| 6982 | -0.014 |
| 6983 | 0.086 |
| 6984 | -0.013 |
| 6985 | 0.061 |
| 6986 | -0.051 |
| 6987 | 0.111 |
| 6988 | 0.035 |
| 6989 | -0.078 |
| 6990 | 0.016 |
| 6991 | -0.001 |
| 6992 | 0.080 |
| 6993 | 0.023 |
| 6994 | -0.062 |
| 6995 | 0.022 |
| 6996 | -0.067 |

| | |
|---|---|
| 6997 | 0.077 |
| 6998 | 0.192 |
| 6999 | 0.020 |
| 7000 | -0.046 |
| 7001 | -0.190 |
| 7002 | 0.029 |
| 7003 | 0.102 |
| 7004 | 0.142 |
| 7005 | 0.097 |
| 7006 | -0.066 |
| 7007 | 0.033 |
| 7008 | 0.086 |
| 7009 | -0.053 |
| 7010 | -0.075 |
| 7011 | 0.017 |
| 7012 | -0.004 |
| 7013 | 0.047 |
| 7014 | -0.059 |
| 7015 | -0.124 |
| 7016 | -0.029 |
| 7017 | 0.033 |
| 7018 | -0.004 |
| 7019 | 0.107 |
| 7020 | -0.210 |
| 7021 | 0.133 |
| 7022 | -0.065 |
| 7023 | -0.074 |
| 7024 | 0.019 |
| 7025 | -0.101 |
| 7026 | 0.068 |
| 7027 | 0.009 |
| 7028 | 0.086 |
| 7029 | -0.025 |
| 7030 | 0.033 |
| 7031 | 0.053 |
| 7032 | -0.059 |
| 7033 | 0.097 |
| 7034 | 0.176 |
| 7035 | -0.068 |
| 7036 | 0.041 |
| 7037 | -0.141 |
| 7038 | 0.078 |
| 7039 | 0.012 |
| 7040 | 0.102 |
| 7041 | -0.014 |
| 7042 | -0.083 |
| 7043 | -0.045 |
| 7044 | 0.032 |
| 7045 | 0.014 |
| 7046 | 0.105 |

| | |
|---|---|
| 7047 | -0.003 |
| 7048 | 0.038 |
| 7049 | 0.055 |
| 7050 | 0.059 |
| 7051 | -0.121 |
| 7052 | 0.011 |
| 7053 | 0.012 |
| 7054 | 0.010 |
| 7055 | 0.084 |
| 7056 | -0.008 |
| 7057 | 0.032 |
| 7058 | 0.005 |
| 7059 | -0.120 |
| 7060 | 0.026 |
| 7061 | 0.103 |
| 7062 | -0.035 |
| 7063 | 0.016 |
| 7064 | 0.011 |
| 7065 | -0.064 |
| 7066 | -0.173 |
| 7067 | -0.116 |
| 7068 | -0.091 |
| 7069 | 0.123 |
| 7070 | 0.054 |
| 7071 | -0.166 |
| 7072 | 0.004 |
| 7073 | 0.012 |
| 7074 | -0.068 |
| 7075 | -0.106 |
| 7076 | -0.052 |
| 7077 | -0.003 |
| 7078 | -0.105 |
| 7079 | 0.027 |
| 7080 | 0.001 |
| 7081 | -0.105 |
| 7082 | -0.051 |
| 7083 | 0.095 |
| 7084 | 0.111 |
| 7085 | -0.074 |
| 7086 | -0.016 |
| 7087 | -0.028 |
| 7088 | 0.002 |
| 7089 | 0.041 |
| 7090 | 0.119 |
| 7091 | -0.015 |
| 7092 | 0.019 |
| 7093 | 0.061 |
| 7094 | 0.054 |
| 7095 | -0.002 |
| 7096 | 0.040 |

| | |
|---|---|
| 7097 | 0.049 |
| 7098 | 0.014 |
| 7099 | -0.037 |
| 7100 | 0.016 |
| 7101 | -0.009 |
| 7102 | -0.093 |
| 7103 | -0.037 |
| 7104 | -0.050 |
| 7105 | 0.080 |
| 7106 | 0.004 |
| 7107 | -0.036 |
| 7108 | 0.011 |
| 7109 | 0.004 |
| 7110 | 0.137 |
| 7111 | 0.122 |
| 7112 | -0.086 |
| 7113 | 0.042 |
| 7114 | 0.032 |
| 7115 | 0.014 |
| 7116 | -0.127 |
| 7117 | -0.050 |
| 7118 | -0.021 |
| 7119 | -0.093 |
| 7120 | -0.064 |
| 7121 | -0.005 |
| 7122 | 0.097 |
| 7123 | -0.051 |
| 7124 | -0.057 |
| 7125 | -0.056 |
| 7126 | 0.048 |
| 7127 | 0.247 |
| 7128 | 0.050 |
| 7129 | -0.186 |
| 7130 | -0.023 |
| 7131 | -0.082 |
| 7132 | 0.080 |
| 7133 | -0.008 |
| 7134 | -0.016 |
| 7135 | -0.121 |
| 7136 | 0.037 |
| 7137 | -0.090 |
| 7138 | -0.004 |
| 7139 | 0.029 |
| 7140 | 0.044 |
| 7141 | -0.005 |
| 7142 | 0.041 |
| 7143 | -0.046 |
| 7144 | -0.070 |
| 7145 | -0.047 |
| 7146 | -0.011 |

| | |
|---|---|
| 7147 | -0.007 |
| 7148 | -0.004 |
| 7149 | 0.027 |
| 7150 | -0.006 |
| 7151 | -0.089 |
| 7152 | -0.001 |
| 7153 | 0.029 |
| 7154 | 0.005 |
| 7155 | 0.022 |
| 7156 | 0.039 |
| 7157 | -0.012 |
| 7158 | -0.137 |
| 7159 | 0.023 |
| 7160 | -0.003 |
| 7161 | 0.012 |
| 7162 | -0.065 |
| 7163 | 0.015 |
| 7164 | 0.025 |
| 7165 | -0.127 |
| 7166 | 0.053 |
| 7167 | -0.027 |
| 7168 | -0.068 |
| 7169 | 0.070 |
| 7170 | 0.104 |
| 7171 | 0.026 |
| 7172 | -0.188 |
| 7173 | 0.016 |
| 7174 | 0.032 |
| 7175 | -0.047 |
| 7176 | 0.012 |
| 7177 | 0.047 |
| 7178 | -0.100 |
| 7179 | -0.059 |
| 7180 | -0.022 |
| 7181 | 0.018 |
| 7182 | 0.036 |
| 7183 | -0.032 |
| 7184 | -0.042 |
| 7185 | 0.041 |
| 7186 | -0.073 |
| 7187 | 0.043 |
| 7188 | 0.022 |
| 7189 | 0.034 |
| 7190 | 0.001 |
| 7191 | -0.030 |
| 7192 | -0.141 |
| 7193 | -0.039 |
| 7194 | -0.116 |
| 7195 | -0.028 |
| 7196 | 0.051 |

| | |
|---|---|
| 7197 | 0.088 |
| 7198 | -0.008 |
| 7199 | -0.006 |
| 7200 | -0.026 |
| 7201 | -0.136 |
| 7202 | 0.073 |
| 7203 | -0.057 |
| 7204 | 0.063 |
| 7205 | -0.010 |
| 7206 | -0.103 |
| 7207 | -0.043 |
| 7208 | -0.053 |
| 7209 | 0.059 |
| 7210 | -0.153 |
| 7211 | 0.072 |
| 7212 | -0.032 |
| 7213 | 0.081 |
| 7214 | 0.093 |
| 7215 | 0.051 |
| 7216 | 0.027 |
| 7217 | 0.034 |
| 7218 | 0.096 |
| 7219 | 0.140 |
| 7220 | 0.134 |
| 7221 | 0.017 |
| 7222 | -0.033 |
| 7223 | -0.075 |
| 7224 | 0.117 |
| 7225 | -0.093 |
| 7226 | -0.009 |
| 7227 | -0.069 |
| 7228 | 0.132 |
| 7229 | 0.101 |
| 7230 | -0.077 |
| 7231 | -0.092 |
| 7232 | -0.013 |
| 7233 | -0.068 |
| 7234 | 0.002 |
| 7235 | -0.043 |
| 7236 | 0.071 |
| 7237 | 0.142 |
| 7238 | -0.075 |
| 7239 | -0.076 |
| 7240 | 0.102 |
| 7241 | -0.070 |
| 7242 | 0.003 |
| 7243 | -0.083 |
| 7244 | 0.060 |
| 7245 | 0.035 |
| 7246 | 0.028 |

| | |
|---|---|
| 7247 | 0.011 |
| 7248 | 0.053 |
| 7249 | -0.037 |
| 7250 | 0.080 |
| 7251 | -0.039 |
| 7252 | -0.070 |
| 7253 | -0.049 |
| 7254 | 0.018 |
| 7255 | 0.004 |
| 7256 | 0.101 |
| 7257 | -0.032 |
| 7258 | 0.004 |
| 7259 | -0.050 |
| 7260 | 0.139 |
| 7261 | -0.047 |
| 7262 | 0.026 |
| 7263 | 0.074 |
| 7264 | 0.010 |
| 7265 | 0.109 |
| 7266 | -0.054 |
| 7267 | 0.068 |
| 7268 | -0.040 |
| 7269 | 0.090 |
| 7270 | -0.104 |
| 7271 | 0.102 |
| 7272 | -0.002 |
| 7273 | 0.044 |
| 7274 | -0.010 |
| 7275 | 0.014 |
| 7276 | 0.041 |
| 7277 | 0.022 |
| 7278 | 0.026 |
| 7279 | -0.080 |
| 7280 | 0.092 |
| 7281 | 0.096 |
| 7282 | 0.010 |
| 7283 | 0.027 |
| 7284 | -0.031 |
| 7285 | 0.191 |
| 7286 | -0.062 |
| 7287 | 0.146 |
| 7288 | 0.070 |
| 7289 | -0.042 |
| 7290 | -0.064 |
| 7291 | -0.044 |
| 7292 | 0.040 |
| 7293 | -0.030 |
| 7294 | 0.085 |
| 7295 | 0.073 |
| 7296 | 0.045 |

| | |
|---|---|
| 7297 | -0.113 |
| 7298 | -0.015 |
| 7299 | -0.012 |
| 7300 | -0.069 |
| 7301 | -0.020 |
| 7302 | 0.029 |
| 7303 | -0.037 |
| 7304 | -0.043 |
| 7305 | -0.043 |
| 7306 | 0.129 |
| 7307 | 0.045 |
| 7308 | -0.083 |
| 7309 | -0.106 |
| 7310 | 0.011 |
| 7311 | 0.044 |
| 7312 | 0.108 |
| 7313 | 0.016 |
| 7314 | -0.111 |
| 7315 | -0.058 |
| 7316 | 0.086 |
| 7317 | 0.084 |
| 7318 | 0.012 |
| 7319 | -0.116 |
| 7320 | -0.059 |
| 7321 | 0.090 |
| 7322 | 0.034 |
| 7323 | -0.045 |
| 7324 | 0.117 |
| 7325 | 0.044 |
| 7326 | 0.023 |
| 7327 | 0.028 |
| 7328 | 0.244 |
| 7329 | -0.135 |
| 7330 | 0.035 |
| 7331 | -0.003 |
| 7332 | -0.088 |
| 7333 | -0.073 |
| 7334 | -0.044 |
| 7335 | 0.024 |
| 7336 | -0.042 |
| 7337 | -0.047 |
| 7338 | -0.077 |
| 7339 | -0.038 |
| 7340 | -0.079 |
| 7341 | -0.102 |
| 7342 | 0.032 |
| 7343 | 0.112 |
| 7344 | 0.028 |
| 7345 | -0.101 |
| 7346 | 0.058 |

| | |
|---|---|
| 7347 | -0.121 |
| 7348 | 0.130 |
| 7349 | 0.035 |
| 7350 | 0.019 |
| 7351 | 0.061 |
| 7352 | 0.036 |
| 7353 | 0.080 |
| 7354 | 0.019 |
| 7355 | 0.001 |
| 7356 | -0.020 |
| 7357 | -0.128 |
| 7358 | 0.046 |
| 7359 | 0.022 |
| 7360 | 0.016 |
| 7361 | -0.067 |
| 7362 | -0.031 |
| 7363 | 0.045 |
| 7364 | 0.090 |
| 7365 | -0.025 |
| 7366 | -0.120 |
| 7367 | 0.005 |
| 7368 | 0.077 |
| 7369 | 0.063 |
| 7370 | -0.030 |
| 7371 | -0.021 |
| 7372 | 0.039 |
| 7373 | 0.073 |
| 7374 | -0.010 |
| 7375 | 0.006 |
| 7376 | 0.011 |
| 7377 | 0.082 |
| 7378 | -0.075 |
| 7379 | -0.015 |
| 7380 | 0.009 |
| 7381 | 0.073 |
| 7382 | 0.057 |
| 7383 | 0.051 |
| 7384 | -0.050 |
| 7385 | 0.070 |
| 7386 | -0.120 |
| 7387 | -0.129 |
| 7388 | -0.063 |
| 7389 | -0.048 |
| 7390 | -0.151 |
| 7391 | -0.092 |
| 7392 | 0.171 |
| 7393 | -0.053 |
| 7394 | -0.058 |
| 7395 | -0.034 |
| 7396 | 0.109 |

| | |
|---|---|
| 7397 | -0.046 |
| 7398 | -0.006 |
| 7399 | 0.134 |
| 7400 | 0.025 |
| 7401 | -0.062 |
| 7402 | 0.063 |
| 7403 | -0.067 |
| 7404 | -0.061 |
| 7405 | 0.019 |
| 7406 | -0.045 |
| 7407 | -0.073 |
| 7408 | -0.138 |
| 7409 | -0.031 |
| 7410 | -0.036 |
| 7411 | 0.119 |
| 7412 | 0.001 |
| 7413 | 0.004 |
| 7414 | -0.079 |
| 7415 | 0.063 |
| 7416 | 0.025 |
| 7417 | -0.001 |
| 7418 | -0.078 |
| 7419 | 0.019 |
| 7420 | 0.029 |
| 7421 | -0.144 |
| 7422 | -0.186 |
| 7423 | 0.064 |
| 7424 | 0.030 |
| 7425 | -0.069 |
| 7426 | 0.005 |
| 7427 | -0.030 |
| 7428 | 0.047 |
| 7429 | -0.009 |
| 7430 | -0.025 |
| 7431 | -0.095 |
| 7432 | -0.026 |
| 7433 | 0.041 |
| 7434 | -0.021 |
| 7435 | -0.100 |
| 7436 | 0.063 |
| 7437 | 0.001 |
| 7438 | -0.032 |
| 7439 | 0.038 |
| 7440 | -0.109 |
| 7441 | 0.067 |
| 7442 | -0.053 |
| 7443 | 0.025 |
| 7444 | -0.074 |
| 7445 | 0.151 |
| 7446 | -0.070 |

| | |
|---|---|
| 7447 | -0.014 |
| 7448 | -0.100 |
| 7449 | -0.029 |
| 7450 | -0.062 |
| 7451 | -0.060 |
| 7452 | 0.097 |
| 7453 | 0.025 |
| 7454 | -0.001 |
| 7455 | 0.022 |
| 7456 | 0.155 |
| 7457 | -0.099 |
| 7458 | 0.094 |
| 7459 | -0.055 |
| 7460 | 0.007 |
| 7461 | 0.014 |
| 7462 | -0.028 |
| 7463 | 0.039 |
| 7464 | 0.064 |
| 7465 | -0.079 |
| 7466 | 0.006 |
| 7467 | 0.112 |
| 7468 | -0.038 |
| 7469 | 0.032 |
| 7470 | -0.015 |
| 7471 | -0.017 |
| 7472 | -0.059 |
| 7473 | -0.048 |
| 7474 | -0.108 |
| 7475 | 0.029 |
| 7476 | -0.062 |
| 7477 | -0.021 |
| 7478 | 0.041 |
| 7479 | -0.034 |
| 7480 | 0.002 |
| 7481 | 0.045 |
| 7482 | 0.004 |
| 7483 | 0.022 |
| 7484 | 0.014 |
| 7485 | -0.015 |
| 7486 | -0.006 |
| 7487 | -0.070 |
| 7488 | -0.065 |
| 7489 | -0.065 |
| 7490 | 0.021 |
| 7491 | -0.007 |
| 7492 | 0.098 |
| 7493 | -0.005 |
| 7494 | 0.033 |
| 7495 | -0.075 |
| 7496 | 0.016 |

| | |
|---|---|
| 7497 | -0.070 |
| 7498 | -0.104 |
| 7499 | 0.007 |
| 7500 | 0.011 |
| 7501 | 0.027 |
| 7502 | 0.060 |
| 7503 | 0.042 |
| 7504 | 0.014 |
| 7505 | -0.067 |
| 7506 | -0.065 |
| 7507 | -0.055 |
| 7508 | 0.016 |
| 7509 | 0.071 |
| 7510 | 0.034 |
| 7511 | 0.071 |
| 7512 | 0.135 |
| 7513 | -0.013 |
| 7514 | 0.061 |
| 7515 | -0.099 |
| 7516 | -0.043 |
| 7517 | -0.001 |
| 7518 | -0.043 |
| 7519 | -0.192 |
| 7520 | 0.044 |
| 7521 | -0.062 |
| 7522 | -0.111 |
| 7523 | 0.051 |
| 7524 | 0.067 |
| 7525 | -0.119 |
| 7526 | -0.098 |
| 7527 | -0.041 |
| 7528 | -0.115 |
| 7529 | 0.006 |
| 7530 | 0.081 |
| 7531 | 0.109 |
| 7532 | 0.032 |
| 7533 | 0.090 |
| 7534 | 0.081 |
| 7535 | 0.092 |
| 7536 | -0.007 |
| 7537 | 0.022 |
| 7538 | -0.030 |
| 7539 | -0.118 |
| 7540 | 0.003 |
| 7541 | 0.004 |
| 7542 | -0.018 |
| 7543 | 0.059 |
| 7544 | 0.015 |
| 7545 | 0.022 |
| 7546 | 0.008 |

| | |
|---|---|
| 7547 | -0.064 |
| 7548 | -0.129 |
| 7549 | -0.025 |
| 7550 | -0.017 |
| 7551 | -0.135 |
| 7552 | 0.072 |
| 7553 | 0.111 |
| 7554 | -0.078 |
| 7555 | -0.074 |
| 7556 | -0.005 |
| 7557 | 0.078 |
| 7558 | -0.006 |
| 7559 | 0.031 |
| 7560 | 0.104 |
| 7561 | 0.038 |
| 7562 | 0.044 |
| 7563 | -0.017 |
| 7564 | -0.069 |
| 7565 | 0.076 |
| 7566 | -0.044 |
| 7567 | -0.004 |
| 7568 | 0.069 |
| 7569 | 0.059 |
| 7570 | 0.160 |
| 7571 | 0.114 |
| 7572 | -0.032 |
| 7573 | -0.072 |
| 7574 | 0.079 |
| 7575 | -0.028 |
| 7576 | -0.081 |
| 7577 | 0.003 |
| 7578 | 0.077 |
| 7579 | 0.000 |
| 7580 | 0.058 |
| 7581 | -0.037 |
| 7582 | 0.039 |
| 7583 | -0.030 |
| 7584 | -0.030 |
| 7585 | 0.018 |
| 7586 | 0.005 |
| 7587 | 0.098 |
| 7588 | 0.063 |
| 7589 | -0.013 |
| 7590 | 0.051 |
| 7591 | 0.076 |
| 7592 | -0.027 |
| 7593 | -0.038 |
| 7594 | 0.035 |
| 7595 | -0.033 |
| 7596 | 0.047 |

| | |
|---|---|
| 7597 | -0.005 |
| 7598 | 0.018 |
| 7599 | -0.064 |
| 7600 | 0.014 |
| 7601 | 0.033 |
| 7602 | -0.136 |
| 7603 | 0.046 |
| 7604 | -0.030 |
| 7605 | 0.021 |
| 7606 | 0.021 |
| 7607 | -0.027 |
| 7608 | -0.039 |
| 7609 | 0.132 |
| 7610 | -0.053 |
| 7611 | 0.018 |
| 7612 | -0.004 |
| 7613 | -0.007 |
| 7614 | 0.081 |
| 7615 | 0.015 |
| 7616 | 0.009 |
| 7617 | 0.121 |
| 7618 | -0.075 |
| 7619 | 0.075 |
| 7620 | 0.069 |
| 7621 | 0.025 |
| 7622 | 0.063 |
| 7623 | 0.039 |
| 7624 | 0.069 |
| 7625 | 0.006 |
| 7626 | 0.106 |
| 7627 | 0.019 |
| 7628 | 0.055 |
| 7629 | 0.016 |
| 7630 | 0.086 |
| 7631 | -0.046 |
| 7632 | 0.055 |
| 7633 | -0.063 |
| 7634 | 0.058 |
| 7635 | -0.007 |
| 7636 | 0.042 |
| 7637 | -0.039 |
| 7638 | 0.029 |
| 7639 | 0.052 |
| 7640 | 0.005 |
| 7641 | -0.002 |
| 7642 | -0.010 |
| 7643 | -0.022 |
| 7644 | -0.015 |
| 7645 | -0.152 |
| 7646 | -0.134 |

| | |
|---|---|
| 7647 | 0.066 |
| 7648 | 0.053 |
| 7649 | 0.079 |
| 7650 | 0.086 |
| 7651 | 0.031 |
| 7652 | 0.005 |
| 7653 | -0.009 |
| 7654 | -0.003 |
| 7655 | 0.176 |
| 7656 | 0.041 |
| 7657 | -0.115 |
| 7658 | 0.017 |
| 7659 | -0.044 |
| 7660 | 0.059 |
| 7661 | 0.031 |
| 7662 | 0.107 |
| 7663 | 0.108 |
| 7664 | 0.130 |
| 7665 | -0.023 |
| 7666 | 0.020 |
| 7667 | 0.021 |
| 7668 | 0.021 |
| 7669 | 0.140 |
| 7670 | -0.046 |
| 7671 | -0.163 |
| 7672 | -0.062 |
| 7673 | 0.065 |
| 7674 | -0.014 |
| 7675 | 0.142 |
| 7676 | 0.040 |
| 7677 | -0.025 |
| 7678 | -0.027 |
| 7679 | -0.033 |
| 7680 | -0.089 |
| 7681 | 0.013 |
| 7682 | -0.056 |
| 7683 | -0.125 |
| 7684 | 0.093 |
| 7685 | 0.019 |
| 7686 | 0.039 |
| 7687 | -0.025 |
| 7688 | 0.036 |
| 7689 | -0.004 |
| 7690 | 0.016 |
| 7691 | -0.096 |
| 7692 | 0.084 |
| 7693 | 0.051 |
| 7694 | 0.034 |
| 7695 | 0.039 |
| 7696 | 0.025 |

| | |
|---|---|
| 7697 | 0.037 |
| 7698 | 0.049 |
| 7699 | 0.005 |
| 7700 | 0.007 |
| 7701 | -0.082 |
| 7702 | 0.086 |
| 7703 | 0.087 |
| 7704 | -0.103 |
| 7705 | -0.063 |
| 7706 | 0.073 |
| 7707 | -0.120 |
| 7708 | -0.050 |
| 7709 | 0.015 |
| 7710 | 0.027 |
| 7711 | -0.027 |
| 7712 | 0.111 |
| 7713 | 0.039 |
| 7714 | 0.003 |
| 7715 | 0.021 |
| 7716 | 0.063 |
| 7717 | 0.021 |
| 7718 | -0.041 |
| 7719 | -0.118 |
| 7720 | -0.048 |
| 7721 | 0.061 |
| 7722 | 0.152 |
| 7723 | -0.014 |
| 7724 | 0.052 |
| 7725 | -0.020 |
| 7726 | 0.032 |
| 7727 | 0.082 |
| 7728 | 0.089 |
| 7729 | 0.124 |
| 7730 | 0.032 |
| 7731 | -0.057 |
| 7732 | -0.023 |
| 7733 | 0.072 |
| 7734 | -0.141 |
| 7735 | 0.127 |
| 7736 | -0.018 |
| 7737 | 0.047 |
| 7738 | -0.177 |
| 7739 | -0.044 |
| 7740 | -0.014 |
| 7741 | -0.057 |
| 7742 | 0.103 |
| 7743 | -0.115 |
| 7744 | 0.062 |
| 7745 | 0.136 |
| 7746 | -0.005 |

| | |
|---|---|
| 7747 | -0.060 |
| 7748 | 0.038 |
| 7749 | 0.088 |
| 7750 | -0.052 |
| 7751 | -0.090 |
| 7752 | 0.029 |
| 7753 | -0.056 |
| 7754 | 0.060 |
| 7755 | 0.093 |
| 7756 | -0.052 |
| 7757 | 0.035 |
| 7758 | -0.096 |
| 7759 | 0.042 |
| 7760 | -0.145 |
| 7761 | -0.007 |
| 7762 | 0.025 |
| 7763 | 0.072 |
| 7764 | -0.056 |
| 7765 | -0.022 |
| 7766 | 0.035 |
| 7767 | 0.039 |
| 7768 | -0.012 |
| 7769 | 0.001 |
| 7770 | 0.026 |
| 7771 | -0.063 |
| 7772 | -0.006 |
| 7773 | -0.019 |
| 7774 | 0.020 |
| 7775 | -0.057 |
| 7776 | -0.029 |
| 7777 | -0.064 |
| 7778 | 0.085 |
| 7779 | -0.011 |
| 7780 | -0.073 |
| 7781 | 0.010 |
| 7782 | -0.021 |
| 7783 | 0.067 |
| 7784 | -0.035 |
| 7785 | -0.057 |
| 7786 | 0.168 |
| 7787 | 0.014 |
| 7788 | 0.140 |
| 7789 | 0.038 |
| 7790 | 0.111 |
| 7791 | 0.055 |
| 7792 | -0.055 |
| 7793 | -0.080 |
| 7794 | 0.055 |
| 7795 | -0.038 |
| 7796 | -0.019 |

| | |
|---|---|
| 7797 | 0.103 |
| 7798 | -0.036 |
| 7799 | -0.032 |
| 7800 | -0.019 |
| 7801 | -0.007 |
| 7802 | -0.009 |
| 7803 | -0.005 |
| 7804 | 0.000 |
| 7805 | 0.143 |
| 7806 | -0.046 |
| 7807 | 0.078 |
| 7808 | -0.029 |
| 7809 | -0.103 |
| 7810 | -0.008 |
| 7811 | 0.050 |
| 7812 | -0.085 |
| 7813 | 0.037 |
| 7814 | 0.090 |
| 7815 | 0.005 |
| 7816 | 0.151 |
| 7817 | -0.087 |
| 7818 | 0.046 |
| 7819 | 0.049 |
| 7820 | 0.107 |
| 7821 | 0.066 |
| 7822 | 0.045 |
| 7823 | 0.042 |
| 7824 | 0.018 |
| 7825 | -0.051 |
| 7826 | 0.009 |
| 7827 | -0.151 |
| 7828 | -0.081 |
| 7829 | -0.074 |
| 7830 | -0.060 |
| 7831 | 0.126 |
| 7832 | -0.053 |
| 7833 | -0.115 |
| 7834 | -0.020 |
| 7835 | -0.038 |
| 7836 | 0.071 |
| 7837 | -0.087 |
| 7838 | -0.008 |
| 7839 | 0.040 |
| 7840 | 0.051 |
| 7841 | -0.066 |
| 7842 | 0.016 |
| 7843 | -0.022 |
| 7844 | -0.002 |
| 7845 | -0.102 |
| 7846 | -0.008 |

| | |
|---|---|
| 7847 | -0.076 |
| 7848 | -0.063 |
| 7849 | 0.087 |
| 7850 | -0.001 |
| 7851 | -0.029 |
| 7852 | -0.094 |
| 7853 | 0.040 |
| 7854 | 0.099 |
| 7855 | -0.079 |
| 7856 | 0.003 |
| 7857 | -0.113 |
| 7858 | -0.027 |
| 7859 | -0.079 |
| 7860 | -0.029 |
| 7861 | 0.041 |
| 7862 | 0.078 |
| 7863 | 0.079 |
| 7864 | 0.043 |
| 7865 | 0.042 |
| 7866 | 0.052 |
| 7867 | 0.083 |
| 7868 | 0.012 |
| 7869 | 0.091 |
| 7870 | 0.150 |
| 7871 | -0.053 |
| 7872 | 0.134 |
| 7873 | -0.077 |
| 7874 | -0.108 |
| 7875 | -0.008 |
| 7876 | -0.064 |
| 7877 | 0.023 |
| 7878 | 0.036 |
| 7879 | -0.054 |
| 7880 | -0.076 |
| 7881 | -0.004 |
| 7882 | -0.078 |
| 7883 | -0.113 |
| 7884 | -0.003 |
| 7885 | 0.008 |
| 7886 | 0.020 |
| 7887 | 0.022 |
| 7888 | -0.029 |
| 7889 | 0.071 |
| 7890 | 0.096 |
| 7891 | 0.083 |
| 7892 | -0.024 |
| 7893 | -0.002 |
| 7894 | 0.067 |
| 7895 | -0.026 |
| 7896 | 0.089 |

| | |
|---|---|
| 7897 | 0.035 |
| 7898 | 0.004 |
| 7899 | -0.050 |
| 7900 | -0.024 |
| 7901 | 0.038 |
| 7902 | -0.014 |
| 7903 | 0.069 |
| 7904 | 0.042 |
| 7905 | -0.025 |
| 7906 | -0.002 |
| 7907 | -0.095 |
| 7908 | -0.012 |
| 7909 | 0.005 |
| 7910 | 0.055 |
| 7911 | 0.011 |
| 7912 | -0.086 |
| 7913 | -0.024 |
| 7914 | -0.044 |
| 7915 | -0.073 |
| 7916 | 0.045 |
| 7917 | -0.024 |
| 7918 | 0.102 |
| 7919 | -0.043 |
| 7920 | -0.022 |
| 7921 | 0.031 |
| 7922 | 0.074 |
| 7923 | 0.071 |
| 7924 | -0.002 |
| 7925 | -0.035 |
| 7926 | -0.066 |
| 7927 | -0.071 |
| 7928 | -0.064 |
| 7929 | -0.015 |
| 7930 | -0.054 |
| 7931 | 0.129 |
| 7932 | 0.042 |
| 7933 | 0.049 |
| 7934 | -0.085 |
| 7935 | -0.107 |
| 7936 | 0.013 |
| 7937 | 0.109 |
| 7938 | -0.064 |
| 7939 | 0.029 |
| 7940 | 0.040 |
| 7941 | -0.010 |
| 7942 | 0.068 |
| 7943 | 0.021 |
| 7944 | -0.058 |
| 7945 | -0.028 |
| 7946 | -0.026 |

| | |
|---|---|
| 7947 | 0.009 |
| 7948 | -0.076 |
| 7949 | -0.059 |
| 7950 | 0.012 |
| 7951 | -0.084 |
| 7952 | 0.069 |
| 7953 | 0.017 |
| 7954 | 0.033 |
| 7955 | -0.036 |
| 7956 | 0.072 |
| 7957 | -0.053 |
| 7958 | -0.045 |
| 7959 | -0.050 |
| 7960 | -0.063 |
| 7961 | -0.044 |
| 7962 | 0.044 |
| 7963 | -0.069 |
| 7964 | -0.001 |
| 7965 | 0.069 |
| 7966 | 0.019 |
| 7967 | 0.014 |
| 7968 | 0.128 |
| 7969 | -0.034 |
| 7970 | -0.116 |
| 7971 | 0.013 |
| 7972 | 0.117 |
| 7973 | 0.030 |
| 7974 | 0.042 |
| 7975 | -0.081 |
| 7976 | 0.055 |
| 7977 | 0.086 |
| 7978 | -0.171 |
| 7979 | 0.047 |
| 7980 | 0.086 |
| 7981 | -0.020 |
| 7982 | 0.053 |
| 7983 | 0.077 |
| 7984 | 0.044 |
| 7985 | 0.136 |
| 7986 | 0.040 |
| 7987 | 0.064 |
| 7988 | 0.074 |
| 7989 | -0.035 |
| 7990 | 0.002 |
| 7991 | 0.072 |
| 7992 | -0.025 |
| 7993 | -0.010 |
| 7994 | 0.068 |
| 7995 | -0.147 |
| 7996 | 0.035 |

| | |
|---|---|
| 7997 | 0.042 |
| 7998 | 0.092 |
| 7999 | -0.032 |
| 8000 | -0.056 |
| 8001 | -0.026 |
| 8002 | -0.132 |
| 8003 | -0.023 |
| 8004 | -0.029 |
| 8005 | 0.015 |
| 8006 | -0.020 |
| 8007 | -0.009 |
| 8008 | 0.031 |
| 8009 | -0.072 |
| 8010 | -0.042 |
| 8011 | 0.030 |
| 8012 | 0.059 |
| 8013 | -0.012 |
| 8014 | 0.020 |
| 8015 | 0.073 |
| 8016 | 0.029 |
| 8017 | -0.037 |
| 8018 | -0.086 |
| 8019 | -0.035 |
| 8020 | -0.011 |
| 8021 | -0.124 |
| 8022 | 0.047 |
| 8023 | 0.114 |
| 8024 | 0.035 |
| 8025 | -0.038 |
| 8026 | -0.031 |
| 8027 | -0.038 |
| 8028 | -0.072 |
| 8029 | 0.037 |
| 8030 | 0.039 |
| 8031 | 0.039 |
| 8032 | 0.021 |
| 8033 | -0.030 |
| 8034 | 0.110 |
| 8035 | 0.019 |
| 8036 | -0.033 |
| 8037 | -0.110 |
| 8038 | 0.055 |
| 8039 | -0.010 |
| 8040 | -0.029 |
| 8041 | -0.010 |
| 8042 | 0.099 |
| 8043 | 0.054 |
| 8044 | -0.107 |
| 8045 | -0.038 |
| 8046 | 0.005 |

| | |
|---|---|
| 8047 | 0.004 |
| 8048 | -0.130 |
| 8049 | 0.019 |
| 8050 | 0.042 |
| 8051 | -0.029 |
| 8052 | -0.014 |
| 8053 | -0.041 |
| 8054 | 0.001 |
| 8055 | 0.043 |
| 8056 | -0.135 |
| 8057 | -0.029 |
| 8058 | 0.043 |
| 8059 | 0.065 |
| 8060 | 0.073 |
| 8061 | -0.147 |
| 8062 | 0.103 |
| 8063 | 0.047 |
| 8064 | -0.139 |
| 8065 | -0.103 |
| 8066 | -0.181 |
| 8067 | -0.055 |
| 8068 | 0.050 |
| 8069 | 0.078 |
| 8070 | -0.053 |
| 8071 | 0.009 |
| 8072 | 0.153 |
| 8073 | 0.062 |
| 8074 | -0.030 |
| 8075 | -0.125 |
| 8076 | -0.068 |
| 8077 | 0.034 |
| 8078 | 0.041 |
| 8079 | 0.032 |
| 8080 | 0.009 |
| 8081 | 0.037 |
| 8082 | -0.065 |
| 8083 | 0.023 |
| 8084 | -0.110 |
| 8085 | 0.108 |
| 8086 | -0.038 |
| 8087 | 0.058 |
| 8088 | -0.003 |
| 8089 | -0.002 |
| 8090 | -0.037 |
| 8091 | -0.065 |
| 8092 | -0.014 |
| 8093 | -0.075 |
| 8094 | -0.073 |
| 8095 | -0.135 |
| 8096 | 0.078 |

| | |
|---|---|
| 8097 | 0.023 |
| 8098 | 0.064 |
| 8099 | -0.041 |
| 8100 | -0.095 |
| 8101 | -0.110 |
| 8102 | -0.002 |
| 8103 | 0.110 |
| 8104 | 0.058 |
| 8105 | 0.081 |
| 8106 | 0.141 |
| 8107 | 0.103 |
| 8108 | 0.012 |
| 8109 | -0.202 |
| 8110 | -0.023 |
| 8111 | 0.080 |
| 8112 | -0.064 |
| 8113 | -0.187 |
| 8114 | 0.001 |
| 8115 | 0.050 |
| 8116 | 0.039 |
| 8117 | -0.043 |
| 8118 | 0.031 |
| 8119 | -0.028 |
| 8120 | 0.042 |
| 8121 | -0.049 |
| 8122 | -0.007 |
| 8123 | 0.009 |
| 8124 | -0.095 |
| 8125 | -0.117 |
| 8126 | -0.032 |
| 8127 | 0.085 |
| 8128 | 0.021 |
| 8129 | -0.057 |
| 8130 | 0.098 |
| 8131 | -0.074 |
| 8132 | -0.041 |
| 8133 | 0.107 |
| 8134 | 0.055 |
| 8135 | 0.092 |
| 8136 | 0.031 |
| 8137 | 0.065 |
| 8138 | 0.026 |
| 8139 | -0.008 |
| 8140 | -0.020 |
| 8141 | 0.062 |
| 8142 | 0.020 |
| 8143 | -0.145 |
| 8144 | -0.045 |
| 8145 | 0.052 |
| 8146 | -0.065 |

| | |
|---|---|
| 8147 | -0.005 |
| 8148 | 0.001 |
| 8149 | -0.041 |
| 8150 | -0.045 |
| 8151 | 0.072 |
| 8152 | 0.020 |
| 8153 | -0.157 |
| 8154 | -0.072 |
| 8155 | 0.003 |
| 8156 | -0.060 |
| 8157 | 0.043 |
| 8158 | -0.055 |
| 8159 | -0.013 |
| 8160 | 0.151 |
| 8161 | -0.032 |
| 8162 | 0.025 |
| 8163 | 0.001 |
| 8164 | 0.055 |
| 8165 | -0.036 |
| 8166 | 0.036 |
| 8167 | -0.035 |
| 8168 | -0.023 |
| 8169 | 0.044 |
| 8170 | 0.046 |
| 8171 | 0.029 |
| 8172 | 0.019 |
| 8173 | -0.072 |
| 8174 | -0.071 |
| 8175 | 0.015 |
| 8176 | -0.022 |
| 8177 | 0.045 |
| 8178 | -0.009 |
| 8179 | -0.191 |
| 8180 | -0.090 |
| 8181 | 0.026 |
| 8182 | 0.045 |
| 8183 | 0.049 |
| 8184 | -0.032 |
| 8185 | -0.048 |
| 8186 | 0.041 |
| 8187 | 0.073 |
| 8188 | 0.074 |
| 8189 | -0.107 |
| 8190 | 0.157 |
| 8191 | -0.059 |
| 8192 | -0.034 |
| 8193 | 0.084 |
| 8194 | -0.062 |
| 8195 | -0.034 |
| 8196 | -0.026 |

| | |
|---|---|
| 8197 | 0.033 |
| 8198 | -0.002 |
| 8199 | 0.069 |
| 8200 | 0.034 |
| 8201 | 0.097 |
| 8202 | 0.043 |
| 8203 | -0.004 |
| 8204 | 0.048 |
| 8205 | -0.038 |
| 8206 | -0.065 |
| 8207 | 0.073 |
| 8208 | 0.031 |
| 8209 | 0.064 |
| 8210 | 0.150 |
| 8211 | -0.022 |
| 8212 | -0.069 |
| 8213 | -0.029 |
| 8214 | 0.006 |
| 8215 | 0.036 |
| 8216 | -0.068 |
| 8217 | -0.007 |
| 8218 | 0.014 |
| 8219 | 0.093 |
| 8220 | -0.050 |
| 8221 | 0.060 |
| 8222 | 0.043 |
| 8223 | -0.046 |
| 8224 | 0.021 |
| 8225 | 0.066 |
| 8226 | -0.016 |
| 8227 | -0.089 |
| 8228 | -0.006 |
| 8229 | 0.001 |
| 8230 | -0.094 |
| 8231 | 0.065 |
| 8232 | -0.023 |
| 8233 | -0.037 |
| 8234 | -0.116 |
| 8235 | 0.106 |
| 8236 | -0.011 |
| 8237 | -0.081 |
| 8238 | -0.054 |
| 8239 | 0.108 |
| 8240 | -0.027 |
| 8241 | 0.085 |
| 8242 | 0.019 |
| 8243 | 0.014 |
| 8244 | 0.182 |
| 8245 | 0.178 |
| 8246 | 0.068 |

| | |
|---|---|
| 8247 | -0.009 |
| 8248 | 0.027 |
| 8249 | -0.001 |
| 8250 | 0.006 |
| 8251 | -0.038 |
| 8252 | 0.017 |
| 8253 | 0.096 |
| 8254 | 0.009 |
| 8255 | -0.126 |
| 8256 | 0.009 |
| 8257 | -0.082 |
| 8258 | -0.048 |
| 8259 | 0.030 |
| 8260 | -0.058 |
| 8261 | 0.013 |
| 8262 | 0.005 |
| 8263 | -0.052 |
| 8264 | 0.117 |
| 8265 | -0.004 |
| 8266 | -0.019 |
| 8267 | 0.027 |
| 8268 | -0.079 |
| 8269 | -0.011 |
| 8270 | 0.113 |
| 8271 | -0.056 |
| 8272 | 0.080 |
| 8273 | -0.019 |
| 8274 | 0.019 |
| 8275 | 0.030 |
| 8276 | 0.037 |
| 8277 | -0.050 |
| 8278 | -0.017 |
| 8279 | -0.013 |
| 8280 | -0.031 |
| 8281 | 0.042 |
| 8282 | -0.056 |
| 8283 | -0.108 |
| 8284 | 0.011 |
| 8285 | 0.043 |
| 8286 | 0.064 |
| 8287 | -0.080 |
| 8288 | 0.026 |
| 8289 | -0.015 |
| 8290 | 0.006 |
| 8291 | 0.012 |
| 8292 | 0.027 |
| 8293 | -0.001 |
| 8294 | -0.028 |
| 8295 | -0.041 |
| 8296 | -0.005 |

| | |
|---|---|
| 8297 | -0.086 |
| 8298 | 0.096 |
| 8299 | -0.081 |
| 8300 | -0.074 |
| 8301 | 0.050 |
| 8302 | -0.062 |
| 8303 | 0.023 |
| 8304 | 0.062 |
| 8305 | 0.112 |
| 8306 | -0.025 |
| 8307 | -0.065 |
| 8308 | -0.105 |
| 8309 | -0.087 |
| 8310 | 0.077 |
| 8311 | -0.053 |
| 8312 | 0.013 |
| 8313 | 0.038 |
| 8314 | -0.159 |
| 8315 | -0.036 |
| 8316 | 0.016 |
| 8317 | -0.083 |
| 8318 | -0.046 |
| 8319 | -0.050 |
| 8320 | 0.006 |
| 8321 | -0.051 |
| 8322 | 0.006 |
| 8323 | -0.074 |
| 8324 | -0.091 |
| 8325 | 0.110 |
| 8326 | -0.102 |
| 8327 | -0.016 |
| 8328 | -0.086 |
| 8329 | 0.008 |
| 8330 | 0.147 |
| 8331 | -0.048 |
| 8332 | -0.074 |
| 8333 | 0.046 |
| 8334 | -0.081 |
| 8335 | -0.072 |
| 8336 | -0.118 |
| 8337 | 0.148 |
| 8338 | 0.090 |
| 8339 | -0.014 |
| 8340 | 0.029 |
| 8341 | -0.127 |
| 8342 | 0.009 |
| 8343 | -0.017 |
| 8344 | 0.104 |
| 8345 | 0.126 |
| 8346 | 0.034 |

| | |
|---|---|
| 8347 | 0.157 |
| 8348 | 0.108 |
| 8349 | -0.106 |
| 8350 | -0.087 |
| 8351 | 0.065 |
| 8352 | -0.125 |
| 8353 | 0.041 |
| 8354 | -0.015 |
| 8355 | 0.071 |
| 8356 | 0.049 |
| 8357 | 0.063 |
| 8358 | -0.016 |
| 8359 | -0.051 |
| 8360 | -0.196 |
| 8361 | 0.153 |
| 8362 | 0.039 |
| 8363 | 0.053 |
| 8364 | 0.082 |
| 8365 | -0.013 |
| 8366 | 0.048 |
| 8367 | 0.005 |
| 8368 | 0.082 |
| 8369 | 0.001 |
| 8370 | 0.020 |
| 8371 | -0.069 |
| 8372 | 0.043 |
| 8373 | -0.041 |
| 8374 | -0.081 |
| 8375 | 0.001 |
| 8376 | -0.043 |
| 8377 | -0.004 |
| 8378 | 0.125 |
| 8379 | -0.002 |
| 8380 | 0.043 |
| 8381 | -0.153 |
| 8382 | 0.069 |
| 8383 | -0.019 |
| 8384 | 0.009 |
| 8385 | 0.180 |
| 8386 | -0.006 |
| 8387 | -0.058 |
| 8388 | 0.013 |
| 8389 | 0.131 |
| 8390 | 0.009 |
| 8391 | -0.023 |
| 8392 | 0.068 |
| 8393 | 0.060 |
| 8394 | 0.002 |
| 8395 | -0.119 |
| 8396 | 0.110 |

| | |
|---|---|
| 8397 | -0.034 |
| 8398 | 0.016 |
| 8399 | -0.027 |
| 8400 | 0.080 |
| 8401 | 0.047 |
| 8402 | -0.027 |
| 8403 | 0.044 |
| 8404 | 0.013 |
| 8405 | 0.086 |
| 8406 | -0.104 |
| 8407 | -0.104 |
| 8408 | -0.122 |
| 8409 | 0.155 |
| 8410 | -0.040 |
| 8411 | -0.098 |
| 8412 | 0.104 |
| 8413 | 0.008 |
| 8414 | -0.038 |
| 8415 | -0.011 |
| 8416 | -0.031 |
| 8417 | 0.013 |
| 8418 | -0.013 |
| 8419 | 0.028 |
| 8420 | -0.046 |
| 8421 | -0.058 |
| 8422 | -0.031 |
| 8423 | -0.009 |
| 8424 | -0.068 |
| 8425 | 0.015 |
| 8426 | 0.042 |
| 8427 | -0.073 |
| 8428 | 0.050 |
| 8429 | -0.082 |
| 8430 | 0.118 |
| 8431 | -0.019 |
| 8432 | 0.167 |
| 8433 | -0.055 |
| 8434 | -0.022 |
| 8435 | -0.032 |
| 8436 | 0.058 |
| 8437 | -0.048 |
| 8438 | 0.070 |
| 8439 | -0.002 |
| 8440 | -0.082 |
| 8441 | 0.115 |
| 8442 | 0.059 |
| 8443 | 0.012 |
| 8444 | -0.087 |
| 8445 | 0.107 |
| 8446 | -0.023 |

| | |
|---|---|
| 8447 | 0.003 |
| 8448 | -0.041 |
| 8449 | 0.100 |
| 8450 | 0.079 |
| 8451 | -0.073 |
| 8452 | -0.035 |
| 8453 | 0.049 |
| 8454 | 0.009 |
| 8455 | -0.079 |
| 8456 | -0.045 |
| 8457 | 0.021 |
| 8458 | -0.017 |
| 8459 | 0.073 |
| 8460 | -0.055 |
| 8461 | 0.000 |
| 8462 | 0.051 |
| 8463 | -0.060 |
| 8464 | 0.009 |
| 8465 | -0.022 |
| 8466 | 0.020 |
| 8467 | -0.047 |
| 8468 | -0.023 |
| 8469 | -0.148 |
| 8470 | 0.041 |
| 8471 | -0.010 |
| 8472 | 0.017 |
| 8473 | -0.080 |
| 8474 | -0.025 |
| 8475 | -0.053 |
| 8476 | -0.039 |
| 8477 | -0.060 |
| 8478 | 0.080 |
| 8479 | 0.010 |
| 8480 | -0.082 |
| 8481 | 0.105 |
| 8482 | -0.070 |
| 8483 | -0.002 |
| 8484 | -0.006 |
| 8485 | -0.061 |
| 8486 | -0.068 |
| 8487 | -0.039 |
| 8488 | 0.003 |
| 8489 | -0.068 |
| 8490 | 0.006 |
| 8491 | 0.073 |
| 8492 | -0.052 |
| 8493 | 0.088 |
| 8494 | 0.051 |
| 8495 | -0.013 |
| 8496 | 0.008 |

| | |
|---|---|
| 8497 | 0.064 |
| 8498 | 0.061 |
| 8499 | -0.011 |
| 8500 | -0.065 |
| 8501 | 0.026 |
| 8502 | -0.076 |
| 8503 | -0.067 |
| 8504 | 0.014 |
| 8505 | -0.026 |
| 8506 | 0.000 |
| 8507 | 0.009 |
| 8508 | 0.021 |
| 8509 | 0.080 |
| 8510 | -0.134 |
| 8511 | 0.064 |
| 8512 | 0.032 |
| 8513 | -0.004 |
| 8514 | -0.013 |
| 8515 | 0.143 |
| 8516 | 0.018 |
| 8517 | 0.107 |
| 8518 | 0.004 |
| 8519 | 0.015 |
| 8520 | 0.068 |
| 8521 | -0.006 |
| 8522 | -0.015 |
| 8523 | 0.031 |
| 8524 | -0.022 |
| 8525 | -0.035 |
| 8526 | 0.074 |
| 8527 | 0.046 |
| 8528 | -0.154 |
| 8529 | -0.009 |
| 8530 | 0.040 |
| 8531 | 0.116 |
| 8532 | 0.031 |
| 8533 | 0.055 |
| 8534 | 0.025 |
| 8535 | -0.028 |
| 8536 | -0.022 |
| 8537 | -0.097 |
| 8538 | -0.030 |
| 8539 | -0.073 |
| 8540 | -0.108 |
| 8541 | 0.019 |
| 8542 | 0.040 |
| 8543 | 0.135 |
| 8544 | -0.062 |
| 8545 | 0.030 |
| 8546 | 0.074 |

| | |
|---|---|
| 8547 | -0.007 |
| 8548 | -0.014 |
| 8549 | -0.066 |
| 8550 | -0.044 |
| 8551 | -0.037 |
| 8552 | -0.007 |
| 8553 | -0.043 |
| 8554 | -0.036 |
| 8555 | 0.021 |
| 8556 | 0.086 |
| 8557 | 0.020 |
| 8558 | -0.113 |
| 8559 | -0.033 |
| 8560 | 0.034 |
| 8561 | -0.080 |
| 8562 | 0.099 |
| 8563 | 0.191 |
| 8564 | -0.115 |
| 8565 | -0.020 |
| 8566 | 0.040 |
| 8567 | -0.006 |
| 8568 | 0.034 |
| 8569 | 0.030 |
| 8570 | 0.009 |
| 8571 | -0.014 |
| 8572 | 0.001 |
| 8573 | 0.044 |
| 8574 | -0.055 |
| 8575 | -0.027 |
| 8576 | -0.103 |
| 8577 | 0.026 |
| 8578 | -0.080 |
| 8579 | 0.071 |
| 8580 | -0.031 |
| 8581 | -0.091 |
| 8582 | 0.074 |
| 8583 | 0.024 |
| 8584 | -0.064 |
| 8585 | -0.023 |
| 8586 | 0.013 |
| 8587 | -0.019 |
| 8588 | 0.026 |
| 8589 | 0.019 |
| 8590 | 0.029 |
| 8591 | 0.019 |
| 8592 | -0.047 |
| 8593 | -0.010 |
| 8594 | -0.019 |
| 8595 | -0.012 |
| 8596 | -0.043 |

| | |
|---|---|
| 8597 | -0.017 |
| 8598 | 0.038 |
| 8599 | 0.040 |
| 8600 | 0.001 |
| 8601 | 0.041 |
| 8602 | 0.029 |
| 8603 | 0.014 |
| 8604 | -0.040 |
| 8605 | 0.058 |
| 8606 | 0.028 |
| 8607 | 0.004 |
| 8608 | 0.049 |
| 8609 | 0.111 |
| 8610 | 0.038 |
| 8611 | 0.008 |
| 8612 | -0.009 |
| 8613 | 0.042 |
| 8614 | 0.078 |
| 8615 | -0.133 |
| 8616 | 0.126 |
| 8617 | 0.127 |
| 8618 | -0.086 |
| 8619 | 0.069 |
| 8620 | -0.198 |
| 8621 | 0.155 |
| 8622 | -0.133 |
| 8623 | 0.074 |
| 8624 | -0.119 |
| 8625 | -0.100 |
| 8626 | -0.072 |
| 8627 | 0.105 |
| 8628 | -0.001 |
| 8629 | -0.042 |
| 8630 | 0.011 |
| 8631 | -0.044 |
| 8632 | 0.039 |
| 8633 | -0.025 |
| 8634 | 0.083 |
| 8635 | -0.005 |
| 8636 | 0.103 |
| 8637 | 0.007 |
| 8638 | -0.089 |
| 8639 | 0.028 |
| 8640 | 0.005 |
| 8641 | -0.022 |
| 8642 | -0.040 |
| 8643 | 0.028 |
| 8644 | 0.014 |
| 8645 | -0.021 |
| 8646 | -0.111 |

| | |
|---|---|
| 8647 | 0.022 |
| 8648 | -0.014 |
| 8649 | 0.041 |
| 8650 | -0.126 |
| 8651 | -0.068 |
| 8652 | 0.124 |
| 8653 | -0.091 |
| 8654 | -0.041 |
| 8655 | 0.021 |
| 8656 | 0.003 |
| 8657 | -0.017 |
| 8658 | -0.058 |
| 8659 | 0.045 |
| 8660 | -0.031 |
| 8661 | -0.041 |
| 8662 | -0.139 |
| 8663 | 0.080 |
| 8664 | -0.013 |
| 8665 | 0.074 |
| 8666 | 0.049 |
| 8667 | 0.053 |
| 8668 | 0.066 |
| 8669 | 0.121 |
| 8670 | 0.057 |
| 8671 | -0.041 |
| 8672 | 0.051 |
| 8673 | 0.010 |
| 8674 | 0.037 |
| 8675 | 0.151 |
| 8676 | 0.014 |
| 8677 | 0.021 |
| 8678 | 0.075 |
| 8679 | -0.082 |
| 8680 | -0.106 |
| 8681 | -0.062 |
| 8682 | 0.092 |
| 8683 | -0.016 |
| 8684 | -0.061 |
| 8685 | 0.014 |
| 8686 | 0.070 |
| 8687 | -0.022 |
| 8688 | -0.011 |
| 8689 | -0.061 |
| 8690 | 0.045 |
| 8691 | 0.017 |
| 8692 | -0.043 |
| 8693 | 0.005 |
| 8694 | 0.058 |
| 8695 | 0.022 |
| 8696 | 0.043 |

| | |
|---|---|
| 8697 | -0.105 |
| 8698 | 0.048 |
| 8699 | 0.079 |
| 8700 | -0.034 |
| 8701 | 0.035 |
| 8702 | 0.082 |
| 8703 | -0.080 |
| 8704 | -0.041 |
| 8705 | 0.085 |
| 8706 | -0.001 |
| 8707 | 0.124 |
| 8708 | -0.005 |
| 8709 | -0.027 |
| 8710 | -0.019 |
| 8711 | 0.154 |
| 8712 | -0.059 |
| 8713 | 0.079 |
| 8714 | -0.052 |
| 8715 | 0.021 |
| 8716 | -0.005 |
| 8717 | -0.033 |
| 8718 | -0.042 |
| 8719 | 0.130 |
| 8720 | 0.006 |
| 8721 | -0.059 |
| 8722 | -0.055 |
| 8723 | -0.050 |
| 8724 | 0.046 |
| 8725 | 0.045 |
| 8726 | 0.006 |
| 8727 | 0.022 |
| 8728 | 0.184 |
| 8729 | 0.058 |
| 8730 | -0.053 |
| 8731 | -0.009 |
| 8732 | 0.004 |
| 8733 | -0.051 |
| 8734 | 0.033 |
| 8735 | 0.034 |
| 8736 | -0.004 |
| 8737 | 0.045 |
| 8738 | -0.065 |
| 8739 | -0.072 |
| 8740 | 0.006 |
| 8741 | 0.090 |
| 8742 | -0.112 |
| 8743 | -0.077 |
| 8744 | -0.024 |
| 8745 | -0.070 |
| 8746 | -0.176 |

| | |
|---|---|
| 8747 | 0.014 |
| 8748 | 0.042 |
| 8749 | 0.056 |
| 8750 | -0.041 |
| 8751 | -0.087 |
| 8752 | 0.061 |
| 8753 | -0.092 |
| 8754 | 0.017 |
| 8755 | 0.040 |
| 8756 | -0.109 |
| 8757 | 0.025 |
| 8758 | -0.028 |
| 8759 | 0.071 |
| 8760 | -0.039 |
| 8761 | 0.066 |
| 8762 | -0.028 |
| 8763 | -0.053 |
| 8764 | 0.018 |
| 8765 | -0.061 |
| 8766 | 0.018 |
| 8767 | 0.029 |
| 8768 | 0.004 |
| 8769 | 0.072 |
| 8770 | 0.071 |
| 8771 | 0.001 |
| 8772 | -0.028 |
| 8773 | -0.024 |
| 8774 | 0.021 |
| 8775 | 0.058 |
| 8776 | -0.077 |
| 8777 | 0.087 |
| 8778 | 0.000 |
| 8779 | -0.062 |
| 8780 | -0.003 |
| 8781 | -0.117 |
| 8782 | -0.021 |
| 8783 | -0.023 |
| 8784 | 0.010 |
| 8785 | -0.026 |
| 8786 | -0.053 |
| 8787 | -0.078 |
| 8788 | -0.011 |
| 8789 | -0.030 |
| 8790 | 0.095 |
| 8791 | -0.128 |
| 8792 | -0.157 |
| 8793 | -0.016 |
| 8794 | -0.071 |
| 8795 | 0.009 |
| 8796 | -0.105 |

| | |
|---|---|
| 8797 | -0.068 |
| 8798 | -0.014 |
| 8799 | -0.013 |
| 8800 | 0.112 |
| 8801 | 0.013 |
| 8802 | 0.106 |
| 8803 | 0.106 |
| 8804 | -0.002 |
| 8805 | -0.089 |
| 8806 | 0.077 |
| 8807 | -0.017 |
| 8808 | -0.024 |
| 8809 | 0.152 |
| 8810 | -0.010 |
| 8811 | -0.060 |
| 8812 | 0.078 |
| 8813 | 0.054 |
| 8814 | 0.186 |
| 8815 | -0.049 |
| 8816 | -0.089 |
| 8817 | 0.107 |
| 8818 | 0.112 |
| 8819 | -0.040 |
| 8820 | 0.068 |
| 8821 | 0.032 |
| 8822 | -0.068 |
| 8823 | -0.032 |
| 8824 | -0.132 |
| 8825 | -0.027 |
| 8826 | 0.089 |
| 8827 | -0.005 |
| 8828 | 0.021 |
| 8829 | 0.012 |
| 8830 | -0.021 |
| 8831 | -0.024 |
| 8832 | -0.011 |
| 8833 | 0.043 |
| 8834 | -0.055 |
| 8835 | 0.132 |
| 8836 | 0.010 |
| 8837 | -0.019 |
| 8838 | -0.047 |
| 8839 | 0.003 |
| 8840 | -0.006 |
| 8841 | -0.033 |
| 8842 | 0.046 |
| 8843 | -0.146 |
| 8844 | 0.049 |
| 8845 | 0.089 |
| 8846 | 0.077 |

| | |
|---|---|
| 8847 | -0.073 |
| 8848 | -0.018 |
| 8849 | -0.014 |
| 8850 | 0.057 |
| 8851 | 0.019 |
| 8852 | -0.073 |
| 8853 | -0.077 |
| 8854 | -0.028 |
| 8855 | 0.037 |
| 8856 | -0.009 |
| 8857 | -0.003 |
| 8858 | 0.119 |
| 8859 | -0.021 |
| 8860 | 0.084 |
| 8861 | -0.054 |
| 8862 | 0.093 |
| 8863 | -0.090 |
| 8864 | -0.029 |
| 8865 | -0.001 |
| 8866 | 0.050 |
| 8867 | 0.065 |
| 8868 | 0.067 |
| 8869 | -0.073 |
| 8870 | -0.010 |
| 8871 | -0.093 |
| 8872 | -0.035 |
| 8873 | -0.017 |
| 8874 | 0.052 |
| 8875 | -0.035 |
| 8876 | -0.094 |
| 8877 | 0.102 |
| 8878 | -0.095 |
| 8879 | 0.013 |
| 8880 | 0.027 |
| 8881 | -0.028 |
| 8882 | 0.077 |
| 8883 | -0.096 |
| 8884 | -0.045 |
| 8885 | 0.062 |
| 8886 | 0.065 |
| 8887 | 0.061 |
| 8888 | -0.031 |
| 8889 | -0.034 |
| 8890 | -0.013 |
| 8891 | -0.025 |
| 8892 | 0.015 |
| 8893 | 0.095 |
| 8894 | -0.066 |
| 8895 | -0.056 |
| 8896 | -0.071 |

| | |
|---|---|
| 8897 | -0.066 |
| 8898 | -0.011 |
| 8899 | -0.057 |
| 8900 | 0.065 |
| 8901 | 0.058 |
| 8902 | 0.024 |
| 8903 | -0.121 |
| 8904 | -0.073 |
| 8905 | -0.005 |
| 8906 | 0.062 |
| 8907 | -0.059 |
| 8908 | -0.051 |
| 8909 | 0.043 |
| 8910 | 0.035 |
| 8911 | -0.032 |
| 8912 | -0.090 |
| 8913 | 0.027 |
| 8914 | -0.029 |
| 8915 | 0.063 |
| 8916 | 0.027 |
| 8917 | -0.074 |
| 8918 | 0.002 |
| 8919 | -0.051 |
| 8920 | -0.023 |
| 8921 | 0.013 |
| 8922 | -0.052 |
| 8923 | 0.069 |
| 8924 | -0.005 |
| 8925 | 0.172 |
| 8926 | -0.032 |
| 8927 | 0.060 |
| 8928 | -0.034 |
| 8929 | -0.026 |
| 8930 | -0.013 |
| 8931 | 0.114 |
| 8932 | -0.012 |
| 8933 | -0.010 |
| 8934 | 0.117 |
| 8935 | 0.136 |
| 8936 | -0.139 |
| 8937 | -0.053 |
| 8938 | 0.062 |
| 8939 | 0.016 |
| 8940 | 0.110 |
| 8941 | -0.027 |
| 8942 | 0.023 |
| 8943 | 0.106 |
| 8944 | -0.030 |
| 8945 | 0.031 |
| 8946 | 0.082 |

| | |
|---|---|
| 8947 | 0.009 |
| 8948 | -0.055 |
| 8949 | 0.099 |
| 8950 | 0.035 |
| 8951 | -0.041 |
| 8952 | -0.068 |
| 8953 | -0.059 |
| 8954 | 0.121 |
| 8955 | -0.017 |
| 8956 | 0.129 |
| 8957 | -0.067 |
| 8958 | 0.043 |
| 8959 | -0.010 |
| 8960 | 0.169 |
| 8961 | 0.053 |
| 8962 | 0.033 |
| 8963 | -0.072 |
| 8964 | 0.020 |
| 8965 | -0.077 |
| 8966 | 0.138 |
| 8967 | 0.024 |
| 8968 | -0.018 |
| 8969 | -0.034 |
| 8970 | -0.038 |
| 8971 | -0.005 |
| 8972 | 0.055 |
| 8973 | 0.028 |
| 8974 | 0.134 |
| 8975 | -0.009 |
| 8976 | 0.023 |
| 8977 | 0.091 |
| 8978 | 0.128 |
| 8979 | 0.010 |
| 8980 | 0.096 |
| 8981 | 0.072 |
| 8982 | 0.074 |
| 8983 | 0.149 |
| 8984 | 0.046 |
| 8985 | -0.063 |
| 8986 | 0.005 |
| 8987 | 0.019 |
| 8988 | 0.073 |
| 8989 | 0.056 |
| 8990 | -0.031 |
| 8991 | -0.031 |
| 8992 | 0.145 |
| 8993 | 0.147 |
| 8994 | -0.026 |
| 8995 | -0.070 |
| 8996 | 0.058 |

| | |
|---|---|
| 8997 | 0.056 |
| 8998 | -0.034 |
| 8999 | 0.059 |
| 9000 | 0.053 |
| 9001 | -0.085 |
| 9002 | 0.117 |
| 9003 | -0.059 |
| 9004 | -0.090 |
| 9005 | 0.085 |
| 9006 | -0.091 |
| 9007 | 0.106 |
| 9008 | -0.082 |
| 9009 | 0.023 |
| 9010 | 0.025 |
| 9011 | -0.097 |
| 9012 | -0.023 |
| 9013 | -0.194 |
| 9014 | 0.004 |
| 9015 | 0.118 |
| 9016 | 0.071 |
| 9017 | 0.064 |
| 9018 | 0.009 |
| 9019 | -0.034 |
| 9020 | 0.011 |
| 9021 | 0.045 |
| 9022 | 0.040 |
| 9023 | -0.091 |
| 9024 | 0.094 |
| 9025 | 0.027 |
| 9026 | -0.103 |
| 9027 | -0.001 |
| 9028 | 0.046 |
| 9029 | -0.014 |
| 9030 | 0.079 |
| 9031 | 0.089 |
| 9032 | -0.097 |
| 9033 | 0.027 |
| 9034 | -0.036 |
| 9035 | -0.106 |
| 9036 | -0.023 |
| 9037 | 0.012 |
| 9038 | -0.029 |
| 9039 | 0.013 |
| 9040 | 0.077 |
| 9041 | 0.030 |
| 9042 | 0.096 |
| 9043 | 0.114 |
| 9044 | -0.053 |
| 9045 | 0.073 |
| 9046 | -0.011 |

| | |
|---|---|
| 9047 | 0.039 |
| 9048 | -0.026 |
| 9049 | -0.061 |
| 9050 | 0.124 |
| 9051 | 0.091 |
| 9052 | -0.034 |
| 9053 | 0.006 |
| 9054 | -0.028 |
| 9055 | 0.000 |
| 9056 | -0.003 |
| 9057 | -0.024 |
| 9058 | 0.020 |
| 9059 | 0.004 |
| 9060 | 0.035 |
| 9061 | 0.038 |
| 9062 | -0.086 |
| 9063 | -0.053 |
| 9064 | -0.061 |
| 9065 | 0.084 |
| 9066 | -0.016 |
| 9067 | -0.038 |
| 9068 | 0.067 |
| 9069 | -0.003 |
| 9070 | -0.046 |
| 9071 | 0.061 |
| 9072 | -0.013 |
| 9073 | -0.002 |
| 9074 | 0.054 |
| 9075 | 0.005 |
| 9076 | 0.115 |
| 9077 | 0.143 |
| 9078 | -0.166 |
| 9079 | -0.089 |
| 9080 | 0.075 |
| 9081 | 0.089 |
| 9082 | -0.063 |
| 9083 | 0.116 |
| 9084 | -0.099 |
| 9085 | 0.005 |
| 9086 | -0.132 |
| 9087 | -0.061 |
| 9088 | -0.069 |
| 9089 | -0.059 |
| 9090 | -0.162 |
| 9091 | -0.157 |
| 9092 | 0.001 |
| 9093 | 0.028 |
| 9094 | 0.068 |
| 9095 | 0.098 |
| 9096 | 0.062 |

| | |
|---|---|
| 9097 | -0.020 |
| 9098 | 0.061 |
| 9099 | -0.105 |
| 9100 | 0.040 |
| 9101 | 0.094 |
| 9102 | -0.007 |
| 9103 | 0.008 |
| 9104 | -0.028 |
| 9105 | -0.062 |
| 9106 | -0.162 |
| 9107 | 0.010 |
| 9108 | -0.063 |
| 9109 | 0.016 |
| 9110 | -0.032 |
| 9111 | -0.003 |
| 9112 | -0.092 |
| 9113 | 0.086 |
| 9114 | 0.042 |
| 9115 | -0.005 |
| 9116 | -0.029 |
| 9117 | -0.054 |
| 9118 | -0.042 |
| 9119 | 0.070 |
| 9120 | -0.016 |
| 9121 | -0.069 |
| 9122 | 0.034 |
| 9123 | -0.063 |
| 9124 | 0.003 |
| 9125 | 0.114 |
| 9126 | 0.011 |
| 9127 | 0.131 |
| 9128 | 0.036 |
| 9129 | -0.007 |
| 9130 | 0.024 |
| 9131 | 0.039 |
| 9132 | -0.006 |
| 9133 | -0.008 |
| 9134 | -0.029 |
| 9135 | 0.006 |
| 9136 | 0.109 |
| 9137 | 0.009 |
| 9138 | -0.039 |
| 9139 | -0.078 |
| 9140 | 0.044 |
| 9141 | -0.076 |
| 9142 | 0.028 |
| 9143 | 0.025 |
| 9144 | -0.094 |
| 9145 | -0.083 |
| 9146 | 0.070 |

| | |
|---|---|
| 9147 | -0.013 |
| 9148 | -0.099 |
| 9149 | -0.025 |
| 9150 | -0.004 |
| 9151 | -0.015 |
| 9152 | 0.009 |
| 9153 | -0.028 |
| 9154 | -0.077 |
| 9155 | 0.155 |
| 9156 | -0.086 |
| 9157 | -0.071 |
| 9158 | 0.013 |
| 9159 | 0.025 |
| 9160 | -0.007 |
| 9161 | 0.139 |
| 9162 | -0.100 |
| 9163 | 0.043 |
| 9164 | -0.005 |
| 9165 | -0.061 |
| 9166 | -0.005 |
| 9167 | -0.074 |
| 9168 | 0.023 |
| 9169 | 0.051 |
| 9170 | -0.156 |
| 9171 | -0.032 |
| 9172 | 0.003 |
| 9173 | 0.060 |
| 9174 | 0.004 |
| 9175 | 0.014 |
| 9176 | -0.010 |
| 9177 | 0.037 |
| 9178 | -0.036 |
| 9179 | 0.036 |
| 9180 | -0.009 |
| 9181 | 0.032 |
| 9182 | -0.076 |
| 9183 | 0.053 |
| 9184 | 0.030 |
| 9185 | -0.039 |
| 9186 | 0.035 |
| 9187 | -0.089 |
| 9188 | 0.085 |
| 9189 | 0.015 |
| 9190 | 0.128 |
| 9191 | 0.051 |
| 9192 | 0.035 |
| 9193 | -0.032 |
| 9194 | -0.002 |
| 9195 | -0.081 |
| 9196 | 0.010 |

| | |
|---|---|
| 9197 | 0.019 |
| 9198 | -0.009 |
| 9199 | 0.135 |
| 9200 | -0.024 |
| 9201 | 0.027 |
| 9202 | -0.027 |
| 9203 | 0.003 |
| 9204 | -0.099 |
| 9205 | -0.014 |
| 9206 | -0.045 |
| 9207 | -0.075 |
| 9208 | 0.076 |
| 9209 | 0.097 |
| 9210 | -0.065 |
| 9211 | 0.157 |
| 9212 | -0.031 |
| 9213 | 0.097 |
| 9214 | 0.077 |
| 9215 | 0.049 |
| 9216 | -0.037 |
| 9217 | -0.119 |
| 9218 | -0.055 |
| 9219 | 0.076 |
| 9220 | -0.080 |
| 9221 | -0.042 |
| 9222 | 0.046 |
| 9223 | -0.025 |
| 9224 | 0.021 |
| 9225 | 0.074 |
| 9226 | -0.094 |
| 9227 | -0.078 |
| 9228 | -0.066 |
| 9229 | -0.119 |
| 9230 | -0.063 |
| 9231 | -0.037 |
| 9232 | 0.003 |
| 9233 | -0.064 |
| 9234 | -0.011 |
| 9235 | 0.038 |
| 9236 | -0.116 |
| 9237 | 0.048 |
| 9238 | 0.010 |
| 9239 | -0.043 |
| 9240 | -0.016 |
| 9241 | 0.117 |
| 9242 | -0.047 |
| 9243 | 0.070 |
| 9244 | 0.011 |
| 9245 | -0.076 |
| 9246 | -0.146 |

| | |
|---|---|
| 9247 | -0.059 |
| 9248 | -0.006 |
| 9249 | -0.059 |
| 9250 | 0.015 |
| 9251 | -0.069 |
| 9252 | -0.184 |
| 9253 | 0.069 |
| 9254 | 0.044 |
| 9255 | 0.024 |
| 9256 | -0.055 |
| 9257 | -0.003 |
| 9258 | 0.008 |
| 9259 | 0.044 |
| 9260 | 0.009 |
| 9261 | -0.024 |
| 9262 | 0.121 |
| 9263 | -0.049 |
| 9264 | -0.067 |
| 9265 | -0.105 |
| 9266 | -0.040 |
| 9267 | 0.122 |
| 9268 | 0.091 |
| 9269 | 0.065 |
| 9270 | 0.073 |
| 9271 | 0.156 |
| 9272 | -0.009 |
| 9273 | -0.039 |
| 9274 | 0.060 |
| 9275 | 0.062 |
| 9276 | -0.060 |
| 9277 | -0.066 |
| 9278 | 0.042 |
| 9279 | 0.041 |
| 9280 | 0.066 |
| 9281 | -0.143 |
| 9282 | 0.045 |
| 9283 | 0.105 |
| 9284 | 0.031 |
| 9285 | 0.062 |
| 9286 | -0.028 |
| 9287 | -0.046 |
| 9288 | 0.019 |
| 9289 | -0.010 |
| 9290 | -0.121 |
| 9291 | 0.111 |
| 9292 | 0.045 |
| 9293 | -0.077 |
| 9294 | 0.014 |
| 9295 | 0.009 |
| 9296 | -0.024 |

| | |
|---|---|
| 9297 | 0.052 |
| 9298 | 0.011 |
| 9299 | -0.001 |
| 9300 | 0.012 |
| 9301 | 0.031 |
| 9302 | 0.024 |
| 9303 | -0.018 |
| 9304 | -0.032 |
| 9305 | 0.047 |
| 9306 | 0.095 |
| 9307 | -0.003 |
| 9308 | 0.080 |
| 9309 | 0.078 |
| 9310 | 0.081 |
| 9311 | -0.035 |
| 9312 | -0.037 |
| 9313 | -0.085 |
| 9314 | 0.113 |
| 9315 | -0.071 |
| 9316 | 0.007 |
| 9317 | 0.053 |
| 9318 | -0.093 |
| 9319 | 0.124 |
| 9320 | -0.051 |
| 9321 | 0.048 |
| 9322 | -0.017 |
| 9323 | 0.053 |
| 9324 | -0.035 |
| 9325 | -0.119 |
| 9326 | 0.057 |
| 9327 | -0.025 |
| 9328 | 0.122 |
| 9329 | 0.088 |
| 9330 | -0.015 |
| 9331 | 0.017 |
| 9332 | -0.059 |
| 9333 | 0.071 |
| 9334 | 0.001 |
| 9335 | -0.077 |
| 9336 | -0.047 |
| 9337 | 0.047 |
| 9338 | -0.013 |
| 9339 | 0.058 |
| 9340 | -0.088 |
| 9341 | 0.071 |
| 9342 | -0.134 |
| 9343 | -0.047 |
| 9344 | -0.029 |
| 9345 | -0.011 |
| 9346 | -0.045 |

| | |
|---|---|
| 9347 | 0.008 |
| 9348 | -0.052 |
| 9349 | 0.036 |
| 9350 | 0.014 |
| 9351 | -0.049 |
| 9352 | -0.017 |
| 9353 | 0.043 |
| 9354 | 0.043 |
| 9355 | 0.005 |
| 9356 | -0.167 |
| 9357 | 0.013 |
| 9358 | 0.033 |
| 9359 | -0.093 |
| 9360 | 0.094 |
| 9361 | -0.045 |
| 9362 | 0.041 |
| 9363 | 0.136 |
| 9364 | 0.101 |
| 9365 | -0.027 |
| 9366 | 0.090 |
| 9367 | 0.037 |
| 9368 | -0.012 |
| 9369 | 0.108 |
| 9370 | 0.087 |
| 9371 | 0.038 |
| 9372 | 0.070 |
| 9373 | 0.020 |
| 9374 | -0.114 |
| 9375 | -0.005 |
| 9376 | 0.056 |
| 9377 | -0.017 |
| 9378 | 0.074 |
| 9379 | -0.001 |
| 9380 | 0.080 |
| 9381 | -0.008 |
| 9382 | -0.002 |
| 9383 | 0.052 |
| 9384 | 0.070 |
| 9385 | -0.069 |
| 9386 | -0.031 |
| 9387 | -0.045 |
| 9388 | -0.021 |
| 9389 | 0.086 |
| 9390 | -0.042 |
| 9391 | -0.021 |
| 9392 | 0.001 |
| 9393 | 0.025 |
| 9394 | -0.091 |
| 9395 | 0.047 |
| 9396 | 0.124 |

| | |
|---|---|
| 9397 | 0.009 |
| 9398 | 0.025 |
| 9399 | -0.059 |
| 9400 | -0.064 |
| 9401 | 0.041 |
| 9402 | -0.093 |
| 9403 | 0.035 |
| 9404 | 0.076 |
| 9405 | 0.069 |
| 9406 | 0.027 |
| 9407 | 0.087 |
| 9408 | -0.033 |
| 9409 | 0.071 |
| 9410 | -0.089 |
| 9411 | 0.028 |
| 9412 | -0.124 |
| 9413 | -0.018 |
| 9414 | 0.000 |
| 9415 | -0.019 |
| 9416 | 0.064 |
| 9417 | 0.034 |
| 9418 | -0.007 |
| 9419 | 0.062 |
| 9420 | -0.094 |
| 9421 | -0.051 |
| 9422 | 0.080 |
| 9423 | 0.067 |
| 9424 | -0.067 |
| 9425 | -0.167 |
| 9426 | -0.098 |
| 9427 | 0.034 |
| 9428 | 0.014 |
| 9429 | 0.000 |
| 9430 | -0.077 |
| 9431 | 0.165 |
| 9432 | 0.032 |
| 9433 | 0.052 |
| 9434 | -0.056 |
| 9435 | -0.032 |
| 9436 | -0.059 |
| 9437 | -0.005 |
| 9438 | 0.125 |
| 9439 | 0.082 |
| 9440 | 0.023 |
| 9441 | -0.010 |
| 9442 | -0.107 |
| 9443 | 0.101 |
| 9444 | -0.042 |
| 9445 | -0.122 |
| 9446 | -0.048 |

| | |
|---|---|
| 9447 | 0.051 |
| 9448 | -0.069 |
| 9449 | -0.030 |
| 9450 | 0.067 |
| 9451 | -0.013 |
| 9452 | 0.050 |
| 9453 | -0.056 |
| 9454 | -0.051 |
| 9455 | -0.037 |
| 9456 | 0.073 |
| 9457 | 0.036 |
| 9458 | 0.053 |
| 9459 | -0.151 |
| 9460 | 0.201 |
| 9461 | -0.161 |
| 9462 | 0.012 |
| 9463 | 0.054 |
| 9464 | 0.003 |
| 9465 | 0.032 |
| 9466 | 0.111 |
| 9467 | 0.060 |
| 9468 | 0.145 |
| 9469 | -0.043 |
| 9470 | 0.078 |
| 9471 | 0.039 |
| 9472 | 0.070 |
| 9473 | -0.060 |
| 9474 | 0.065 |
| 9475 | -0.075 |
| 9476 | 0.051 |
| 9477 | 0.019 |
| 9478 | 0.038 |
| 9479 | -0.056 |
| 9480 | 0.154 |
| 9481 | -0.011 |
| 9482 | 0.013 |
| 9483 | -0.069 |
| 9484 | 0.097 |
| 9485 | 0.026 |
| 9486 | 0.115 |
| 9487 | 0.049 |
| 9488 | -0.016 |
| 9489 | 0.023 |
| 9490 | 0.081 |
| 9491 | 0.050 |
| 9492 | -0.070 |
| 9493 | 0.028 |
| 9494 | 0.028 |
| 9495 | -0.145 |
| 9496 | -0.027 |

| | |
|---|---|
| 9497 | -0.003 |
| 9498 | -0.036 |
| 9499 | 0.018 |
| 9500 | -0.026 |
| 9501 | 0.016 |
| 9502 | -0.049 |
| 9503 | -0.071 |
| 9504 | 0.037 |
| 9505 | -0.116 |
| 9506 | -0.058 |
| 9507 | -0.070 |
| 9508 | 0.064 |
| 9509 | -0.129 |
| 9510 | -0.038 |
| 9511 | 0.188 |
| 9512 | -0.032 |
| 9513 | -0.056 |
| 9514 | -0.051 |
| 9515 | -0.034 |
| 9516 | -0.108 |
| 9517 | 0.014 |
| 9518 | 0.124 |
| 9519 | 0.031 |
| 9520 | 0.045 |
| 9521 | -0.092 |
| 9522 | -0.048 |
| 9523 | 0.036 |
| 9524 | 0.059 |
| 9525 | 0.055 |
| 9526 | -0.110 |
| 9527 | 0.039 |
| 9528 | -0.043 |
| 9529 | 0.000 |
| 9530 | -0.070 |
| 9531 | -0.020 |
| 9532 | -0.065 |
| 9533 | 0.005 |
| 9534 | -0.095 |
| 9535 | 0.014 |
| 9536 | 0.003 |
| 9537 | -0.086 |
| 9538 | -0.002 |
| 9539 | -0.005 |
| 9540 | -0.014 |
| 9541 | -0.048 |
| 9542 | 0.078 |
| 9543 | 0.065 |
| 9544 | 0.057 |
| 9545 | -0.019 |
| 9546 | -0.001 |

| | |
|---|---|
| 9547 | -0.115 |
| 9548 | -0.070 |
| 9549 | 0.005 |
| 9550 | 0.096 |
| 9551 | 0.097 |
| 9552 | -0.058 |
| 9553 | 0.022 |
| 9554 | 0.018 |
| 9555 | 0.117 |
| 9556 | -0.039 |
| 9557 | 0.003 |
| 9558 | 0.003 |
| 9559 | -0.098 |
| 9560 | 0.076 |
| 9561 | -0.077 |
| 9562 | -0.025 |
| 9563 | 0.068 |
| 9564 | -0.012 |
| 9565 | -0.024 |
| 9566 | -0.094 |
| 9567 | -0.179 |
| 9568 | 0.025 |
| 9569 | 0.094 |
| 9570 | 0.151 |
| 9571 | 0.078 |
| 9572 | -0.014 |
| 9573 | 0.019 |
| 9574 | -0.015 |
| 9575 | 0.047 |
| 9576 | -0.024 |
| 9577 | 0.044 |
| 9578 | 0.077 |
| 9579 | -0.007 |
| 9580 | -0.003 |
| 9581 | 0.001 |
| 9582 | 0.037 |
| 9583 | 0.052 |
| 9584 | 0.013 |
| 9585 | -0.053 |
| 9586 | 0.061 |
| 9587 | 0.025 |
| 9588 | 0.086 |
| 9589 | 0.102 |
| 9590 | 0.107 |
| 9591 | -0.080 |
| 9592 | 0.144 |
| 9593 | -0.186 |
| 9594 | -0.120 |
| 9595 | -0.139 |
| 9596 | -0.113 |

| | |
|---|---|
| 9597 | 0.007 |
| 9598 | -0.038 |
| 9599 | -0.055 |
| 9600 | 0.063 |
| 9601 | 0.055 |
| 9602 | -0.008 |
| 9603 | 0.021 |
| 9604 | 0.105 |
| 9605 | 0.033 |
| 9606 | 0.079 |
| 9607 | -0.003 |
| 9608 | 0.079 |
| 9609 | 0.082 |
| 9610 | 0.110 |
| 9611 | 0.063 |
| 9612 | -0.021 |
| 9613 | -0.067 |
| 9614 | -0.040 |
| 9615 | 0.079 |
| 9616 | -0.040 |
| 9617 | 0.035 |
| 9618 | 0.052 |
| 9619 | -0.081 |
| 9620 | 0.012 |
| 9621 | 0.012 |
| 9622 | 0.003 |
| 9623 | -0.025 |
| 9624 | -0.030 |
| 9625 | 0.019 |
| 9626 | -0.003 |
| 9627 | -0.034 |
| 9628 | 0.022 |
| 9629 | -0.075 |
| 9630 | 0.089 |
| 9631 | -0.082 |
| 9632 | -0.036 |
| 9633 | 0.020 |
| 9634 | -0.099 |
| 9635 | 0.039 |
| 9636 | -0.039 |
| 9637 | -0.032 |
| 9638 | -0.046 |
| 9639 | 0.010 |
| 9640 | 0.037 |
| 9641 | -0.061 |
| 9642 | 0.026 |
| 9643 | 0.000 |
| 9644 | -0.005 |
| 9645 | 0.032 |
| 9646 | 0.024 |

| | |
|---|---|
| 9647 | 0.065 |
| 9648 | -0.046 |
| 9649 | -0.086 |
| 9650 | -0.007 |
| 9651 | 0.080 |
| 9652 | -0.019 |
| 9653 | 0.068 |
| 9654 | 0.091 |
| 9655 | -0.018 |
| 9656 | 0.019 |
| 9657 | -0.151 |
| 9658 | -0.028 |
| 9659 | 0.040 |
| 9660 | 0.022 |
| 9661 | -0.047 |
| 9662 | 0.083 |
| 9663 | 0.005 |
| 9664 | 0.078 |
| 9665 | 0.045 |
| 9666 | 0.010 |
| 9667 | 0.076 |
| 9668 | -0.046 |
| 9669 | -0.045 |
| 9670 | -0.024 |
| 9671 | 0.107 |
| 9672 | 0.018 |
| 9673 | 0.022 |
| 9674 | -0.090 |
| 9675 | -0.043 |
| 9676 | 0.071 |
| 9677 | 0.146 |
| 9678 | -0.058 |
| 9679 | 0.048 |
| 9680 | 0.042 |
| 9681 | 0.049 |
| 9682 | 0.020 |
| 9683 | 0.011 |
| 9684 | 0.054 |
| 9685 | 0.104 |
| 9686 | -0.050 |
| 9687 | -0.008 |
| 9688 | -0.017 |
| 9689 | -0.030 |
| 9690 | -0.012 |
| 9691 | 0.007 |
| 9692 | -0.015 |
| 9693 | -0.092 |
| 9694 | -0.031 |
| 9695 | -0.010 |
| 9696 | 0.030 |

| | |
|---|---|
| 9697 | 0.080 |
| 9698 | 0.081 |
| 9699 | 0.093 |
| 9700 | 0.025 |
| 9701 | -0.019 |
| 9702 | -0.047 |
| 9703 | -0.062 |
| 9704 | -0.117 |
| 9705 | -0.032 |
| 9706 | 0.082 |
| 9707 | -0.216 |
| 9708 | 0.087 |
| 9709 | 0.026 |
| 9710 | -0.019 |
| 9711 | 0.059 |
| 9712 | -0.071 |
| 9713 | -0.042 |
| 9714 | -0.006 |
| 9715 | -0.031 |
| 9716 | -0.026 |
| 9717 | 0.029 |
| 9718 | 0.045 |
| 9719 | 0.115 |
| 9720 | -0.044 |
| 9721 | -0.082 |
| 9722 | -0.037 |
| 9723 | 0.005 |
| 9724 | 0.119 |
| 9725 | -0.031 |
| 9726 | 0.054 |
| 9727 | 0.037 |
| 9728 | -0.061 |
| 9729 | -0.001 |
| 9730 | 0.089 |
| 9731 | 0.089 |
| 9732 | 0.036 |
| 9733 | -0.119 |
| 9734 | 0.032 |
| 9735 | 0.058 |
| 9736 | -0.075 |
| 9737 | -0.177 |
| 9738 | -0.060 |
| 9739 | 0.099 |
| 9740 | -0.135 |
| 9741 | 0.088 |
| 9742 | 0.038 |
| 9743 | -0.057 |
| 9744 | 0.097 |
| 9745 | 0.115 |
| 9746 | 0.014 |

| | |
|---|---|
| 9747 | 0.077 |
| 9748 | -0.143 |
| 9749 | -0.059 |
| 9750 | -0.028 |
| 9751 | -0.061 |
| 9752 | -0.033 |
| 9753 | 0.041 |
| 9754 | 0.048 |
| 9755 | -0.071 |
| 9756 | 0.049 |
| 9757 | 0.027 |
| 9758 | 0.070 |
| 9759 | -0.037 |
| 9760 | 0.087 |
| 9761 | -0.036 |
| 9762 | -0.060 |
| 9763 | -0.020 |
| 9764 | -0.004 |
| 9765 | -0.002 |
| 9766 | 0.054 |
| 9767 | -0.007 |
| 9768 | 0.083 |
| 9769 | -0.018 |
| 9770 | -0.018 |
| 9771 | -0.119 |
| 9772 | 0.122 |
| 9773 | 0.007 |
| 9774 | -0.014 |
| 9775 | 0.121 |
| 9776 | 0.002 |
| 9777 | 0.083 |
| 9778 | -0.035 |
| 9779 | -0.031 |
| 9780 | 0.058 |
| 9781 | 0.168 |
| 9782 | 0.034 |
| 9783 | 0.144 |
| 9784 | -0.048 |
| 9785 | -0.016 |
| 9786 | 0.019 |
| 9787 | 0.050 |
| 9788 | 0.000 |
| 9789 | 0.014 |
| 9790 | -0.033 |
| 9791 | -0.106 |
| 9792 | -0.086 |
| 9793 | 0.030 |
| 9794 | 0.063 |
| 9795 | -0.092 |
| 9796 | -0.001 |

| | |
|---|---|
| 9797 | -0.029 |
| 9798 | -0.060 |
| 9799 | 0.052 |
| 9800 | -0.080 |
| 9801 | -0.061 |
| 9802 | 0.007 |
| 9803 | 0.135 |
| 9804 | 0.009 |
| 9805 | -0.051 |
| 9806 | 0.018 |
| 9807 | 0.181 |
| 9808 | -0.115 |
| 9809 | 0.025 |
| 9810 | 0.100 |
| 9811 | -0.006 |
| 9812 | -0.020 |
| 9813 | -0.036 |
| 9814 | 0.070 |
| 9815 | 0.015 |
| 9816 | -0.005 |
| 9817 | -0.032 |
| 9818 | -0.092 |
| 9819 | -0.062 |
| 9820 | 0.032 |
| 9821 | 0.133 |
| 9822 | -0.073 |
| 9823 | -0.120 |
| 9824 | -0.137 |
| 9825 | -0.013 |
| 9826 | 0.057 |
| 9827 | -0.083 |
| 9828 | -0.041 |
| 9829 | 0.061 |
| 9830 | -0.002 |
| 9831 | -0.032 |
| 9832 | 0.092 |
| 9833 | -0.011 |
| 9834 | -0.018 |
| 9835 | 0.031 |
| 9836 | 0.020 |
| 9837 | 0.067 |
| 9838 | -0.021 |
| 9839 | 0.040 |
| 9840 | 0.140 |
| 9841 | 0.001 |
| 9842 | 0.018 |
| 9843 | 0.055 |
| 9844 | 0.119 |
| 9845 | -0.083 |
| 9846 | -0.058 |

| | |
|---|---|
| 9847 | 0.027 |
| 9848 | 0.011 |
| 9849 | -0.014 |
| 9850 | 0.004 |
| 9851 | -0.017 |
| 9852 | -0.028 |
| 9853 | 0.130 |
| 9854 | -0.016 |
| 9855 | -0.011 |
| 9856 | -0.016 |
| 9857 | -0.011 |
| 9858 | 0.055 |
| 9859 | 0.059 |
| 9860 | -0.080 |
| 9861 | 0.084 |
| 9862 | 0.050 |
| 9863 | 0.038 |
| 9864 | 0.010 |
| 9865 | 0.020 |
| 9866 | 0.100 |
| 9867 | -0.014 |
| 9868 | 0.006 |
| 9869 | -0.121 |
| 9870 | -0.016 |
| 9871 | 0.001 |
| 9872 | 0.082 |
| 9873 | -0.049 |
| 9874 | 0.001 |
| 9875 | -0.110 |
| 9876 | -0.039 |
| 9877 | 0.086 |
| 9878 | 0.132 |
| 9879 | 0.065 |
| 9880 | 0.023 |
| 9881 | 0.014 |
| 9882 | -0.041 |
| 9883 | 0.061 |
| 9884 | -0.079 |
| 9885 | -0.014 |
| 9886 | -0.001 |
| 9887 | 0.032 |
| 9888 | -0.061 |
| 9889 | 0.071 |
| 9890 | 0.000 |
| 9891 | 0.096 |
| 9892 | 0.076 |
| 9893 | 0.096 |
| 9894 | 0.048 |
| 9895 | 0.063 |
| 9896 | -0.123 |

| | |
|---|---|
| 9897 | -0.022 |
| 9898 | -0.020 |
| 9899 | -0.067 |
| 9900 | 0.045 |
| 9901 | 0.024 |
| 9902 | -0.138 |
| 9903 | -0.054 |
| 9904 | 0.076 |
| 9905 | 0.074 |
| 9906 | -0.006 |
| 9907 | -0.129 |
| 9908 | -0.126 |
| 9909 | -0.016 |
| 9910 | 0.045 |
| 9911 | 0.054 |
| 9912 | -0.058 |
| 9913 | 0.008 |
| 9914 | 0.037 |
| 9915 | -0.030 |
| 9916 | -0.077 |
| 9917 | -0.092 |
| 9918 | 0.048 |
| 9919 | -0.017 |
| 9920 | -0.117 |
| 9921 | 0.049 |
| 9922 | 0.123 |
| 9923 | 0.049 |
| 9924 | 0.043 |
| 9925 | 0.095 |
| 9926 | 0.042 |
| 9927 | 0.037 |
| 9928 | 0.027 |
| 9929 | -0.008 |
| 9930 | -0.029 |
| 9931 | -0.073 |
| 9932 | 0.065 |
| 9933 | -0.019 |
| 9934 | -0.063 |
| 9935 | -0.054 |
| 9936 | -0.012 |
| 9937 | 0.054 |
| 9938 | 0.098 |
| 9939 | 0.052 |
| 9940 | -0.065 |
| 9941 | -0.057 |
| 9942 | 0.074 |
| 9943 | -0.002 |
| 9944 | 0.017 |
| 9945 | 0.084 |
| 9946 | 0.157 |

| | |
|---|---|
| 9947 | 0.036 |
| 9948 | -0.015 |
| 9949 | -0.065 |
| 9950 | -0.012 |
| 9951 | -0.006 |
| 9952 | 0.048 |
| 9953 | 0.024 |
| 9954 | -0.013 |
| 9955 | -0.043 |
| 9956 | -0.044 |
| 9957 | -0.161 |
| 9958 | -0.057 |
| 9959 | -0.025 |
| 9960 | -0.035 |
| 9961 | 0.040 |
| 9962 | 0.017 |
| 9963 | -0.079 |
| 9964 | 0.043 |
| 9965 | 0.053 |
| 9966 | 0.018 |
| 9967 | -0.058 |
| 9968 | -0.069 |
| 9969 | -0.050 |
| 9970 | 0.086 |
| 9971 | -0.050 |
| 9972 | 0.010 |
| 9973 | 0.004 |
| 9974 | -0.044 |
| 9975 | -0.043 |
| 9976 | -0.014 |
| 9977 | 0.075 |
| 9978 | 0.064 |
| 9979 | -0.051 |
| 9980 | 0.068 |
| 9981 | -0.071 |
| 9982 | -0.093 |
| 9983 | 0.054 |
| 9984 | 0.020 |
| 9985 | -0.015 |
| 9986 | -0.022 |
| 9987 | 0.020 |
| 9988 | -0.011 |
| 9989 | 0.079 |
| 9990 | -0.025 |
| 9991 | -0.018 |
| 9992 | 0.123 |
| 9993 | -0.047 |
| 9994 | 0.054 |
| 9995 | 0.069 |
| 9996 | -0.007 |

| | |
|---|---|
| 9997 | -0.116 |
| 9998 | 0.040 |
| 9999 | 0.036 |
| 10000 | -0.002 |

**Table S15. Identified changes in GO (Biological Processes, BP; Cellular Compartments, CC) and KEGG pathways. MII oocytes: Young *vs.* AMA**

| | Terms | Overlap | Ratio | P-value | P.adj | Log2 (FC) | Proteins |
|---|---|---|---|---|---|---|---|
| BP | positive regulation of telomerase RNA localization to Cajal body (GO:1904874) | 10/15 | 0.7 | 0.0002 | 0.030 | 0.115 | TCPG; TCPZ; TCPB; RUVB2; RUVB1; TCPA; TCPQ; TCPH; TCPE; TCPD |
| | positive regulation of establishment of protein localization to telomere (GO:190485 | 08/10 | 0.8 | 3.8E-05 | 0.024 | 0.135 | TCPG; TCPZ; TCPB; TCPA; TCPQ; TCPH; TCPE; TCPD |
| | regulation of telomerase RNA localization to Cajal body (GO:1904872) | 10/18 | 0.6 | 0.0002 | 0.030 | 0.115 | TCPG; TCPZ; TCPB; RUVB2; RUVB1; TCPA; TCPQ; TCPH; TCPE; TCPD |
| | positive regulation of protein localization to Cajal body (GO:1904871) | 08/11 | 0.7 | 3.8E-05 | 0.024 | 0.135 | TCPG; TCPZ; TCPB; TCPA; TCPQ; TCPH; TCPE; TCPD |
| | regulation of establishment of protein localization to telomere (GO:0070203) | 08/11 | 0.7 | 3.8E-05 | 0.024 | 0.135 | TCPG; TCPZ; TCPB; TCPA; TCPQ; TCPH; TCPE; TCPD |
| | regulation of protein localization to Cajal body (GO:1904869) | 08/11 | 0.7 | 3.8E-05 | 0.024 | 0.135 | TCPG; TCPZ; TCPB; TCPA; TCPQ; TCPH; TCPE; TCPD |
| | positive regulation of protein localization to chromosome, telomeric region (GO:19 | 08/12 | 0.7 | 3.8E-05 | 0.024 | 0.135 | TCPG; TCPZ; TCPB; TCPA; TCPQ; TCPH; TCPE; TCPD |
